# Evaluation of Expanded 2-Aminobenzothiazole Library for Inhibition of *Pseudomonas aeruginosa* Virulence Phenotypes

**DOI:** 10.1101/2023.05.02.539119

**Authors:** Conrad A. Fihn, Hannah K. Lembke, Jeffrey Gaulin, Patricia Bouchard, Alex R. Villarreal, Mitchell R. Penningroth, Kathryn K. Crone, Grace A. Vogt, Adam J. Gilbertsen, Yann Ayotte, Luciana Couthino de Oliveira, Michael H. Serrano-Wu, Nathalie Drouin, Deborah T. Hung, Ryan C. Hunter, Erin E. Carlson

## Abstract

Bacterial resistance to antibiotics is a rapidly increasing threat to human health. New strategies to combat resistant organisms are desperately needed. One potential avenue is targeting two-component systems, which are the main bacterial signal transduction pathways used to regulate development, metabolism, virulence, and antibiotic resistance. These systems consist of a homodimeric membrane-bound sensor histidine kinase, and a cognate effector, the response regulator. The high sequence conservation in the catalytic and adenosine triphosphate-binding (CA) domain of histidine kinases and their essential role in bacterial signal transduction could enable broad-spectrum antibacterial activity. Through this signal transduction, histidine kinases regulate multiple virulence mechanisms including toxin production, immune evasion, and antibiotic resistance. Targeting virulence, as opposed to development of bactericidal compounds, could reduce evolutionary pressure for acquired resistance. Additionally, compounds targeting the CA domain have the potential to impair multiple two-component systems that regulate virulence in one or more pathogens. We conducted structure-activity relationship studies of 2-aminobenzothiazole-based inhibitors designed to target the CA domain of histidine kinases. We found these compounds have anti-virulence activities in *Pseudomonas aeruginosa*, reducing motility phenotypes and toxin production associated with the pathogenic functions of this bacterium.

## Introduction

The World Health Organization (WHO) has declared antimicrobial resistance to be one of the “…top ten global public health threats facing humanity.”^1^ Various bacterial species have become resistant to all currently approved classes of antibiotics, most of which are derived from compounds developed over 40 years ago. Moreover, developing therapeutics that function through a new mechanism of action has been challenging^1^. Although the FDA approved 58 antibiotics during the past 30 years, only one was active against a novel target. Indeed, only 7 of the 43 antibiotics currently in the pipeline have been classified as novel.^2^ Even as new antibiotics are introduced, bacteria quickly develop resistance due to evolutionary pressures, accelerated by widespread use.^3-4^ In this manner, current strategies paradoxically promote resistance and new approaches are needed.

One promising alternative to traditional bactericidal or bacteriostatic antibiotics is targeting virulence factors, which renders bacteria less pathogenic. Virulence factors include a number of behaviors and processes such as motility, endotoxin production, excretion of proteolytic enzymes, secretion or injection systems, immune evasion, communication, signaling, adhesion, biofilm production, as well as antibiotic resistance mechanisms.^5^ Blocking one or more of these functions could reduce the severity of infection, enhancing the effectiveness of host immunity and efficacy of existing antibiotics.^6^ Such therapies can target virulence factors by directly inhibiting the regulatory systems that control them^6-10^ or virulence-associated proteolytic enzymes.^11-12^

One class of virulence regulators is the two-component systems (TCSs). TCSs are responsible for sensing changes in the bacterial environment and in turn regulating expression, and in some cases functionality, of the cellular machinery required to respond to these changes.^13-21^ TCSs are generally composed of a membrane-spanning histidine kinase (HK) and a cognate cytosolic response regulator (RR). The HK can have several auxiliary domains that affect function, but common to all is a variable periplasmic signaling domain that determines the stimulus/stimuli for each TCS (**Figure 1A**). In addition, HKs commonly possess a transmembrane domain, and a cytosolic kinase region composed of a dimerization and histidine phosphotransfer (DHp) domain, and a highly-conserved catalytic and adenosine triphosphate (ATP)-binding (CA) domain. Upon receiving a stimulus at the signaling domain, HKs undergo a conformational shift, dimerization, and ATP-binding event at the CA domain.^22-23^ The γ- phosphate of ATP is then transferred through a conserved histidine on the DHp domain to a conserved aspartate residue on the RR that can initiate a cellular response, generally acting as a transcription factor.^23-25^ HKs are categorized into eleven classes based on the sequence homology of their two major domains, with most HKs belonging to class-1.^20, 23^ Despite these sub-classifications, HKs all have conserved homology boxes, many of which are contained within the CA domain. The G1-, G2-, G3-, F-, and N-boxes are a series of conserved residues and are responsible for key interactions with ATP (**Figure 1B**).^22-23, 26^

**Figure 1.**
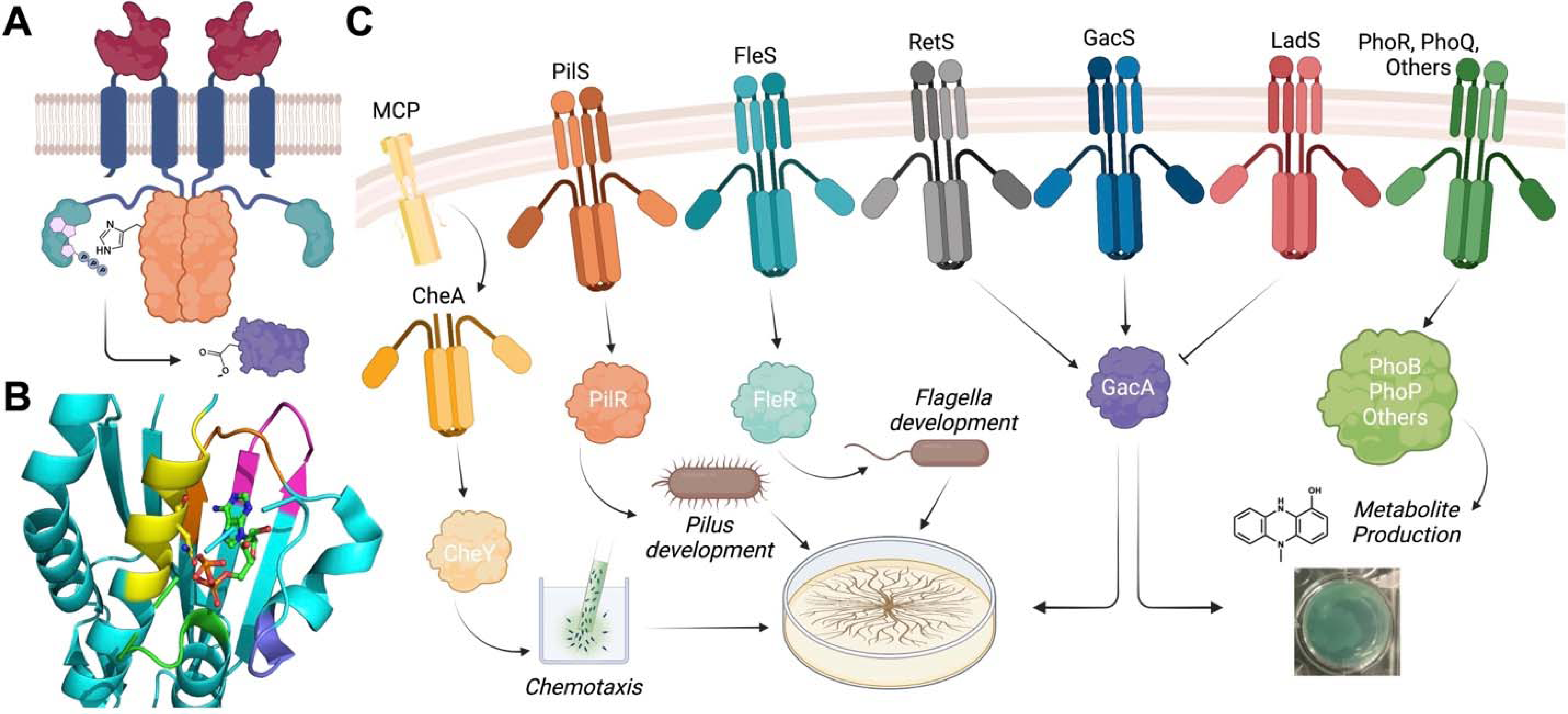
Histidine kinase structure and roles in *P. aeruginos*a virulence. A) Generic structure and domain organization of a TCS. HKs contain a periplasmic sensory domain (maroon) with a transmembrane domain (navy blue) to connect the cytosolic kinase domains comprised of the dimerization and histidine phosphorylation (DHp) domain with the catalytic histidine residue (orange) and the catalytic ATP-binding (CA) domain (teal). HKs generally have a cognate response regulator (purple) containing a conserved aspartate residue where phosphorylation occurs. B) CA domain, homology boxes that represent highly conserved sequence motifs. N-box (yellow), F-box (purple), G1-Box (orange), G2-box (green), G3-box (pink). C) Examples of HKs implicated in pyocyanin production and swarming motility in *P. aeruginosa*. CheA (yellow), a non-traditional cytosolic HK, receives signals through methyl-accepting chemotaxis proteins (MCPs).

HKs are an attractive target for anti-virulence therapy because of their roles in the regulation of multiple virulence mechanisms, their ubiquity among bacteria establishing the potential for broad-spectrum activity, and their absence in higher organisms. The latter suggests that targeting these proteins could produce fewer off-target effects in humans.^15, 27-28^ Depending on their native growth environment, bacteria can produce none to more than 100 different HKs with significant overlap, redundancy, and crosstalk between functionalities.^25, 29^ There are several potential approaches to blocking HK activity. Targeting the variable signaling domain could produce specific inhibition of single HKs while molecules directed to the highly-conserved CA domain could result in pan-inhibition of these proteins.^27, 30-43^

To maximize the potential for inhibiting critical virulence-associated HKs, we sought to develop pan-inhibitors. This strategy also has the theoretical advantage of reducing the emergence of resistance because it would require organisms to respond to a range of inhibitory factors.^44-45^ We previously developed a CA domain-targeted high-throughput screen and identified two compounds sharing a common 2-aminobenzothiazole core as inhibitors of the HKs (general structure in **Figure 2A**). One of these leads is riluzole (**Rilu-1**), an FDA-approved drug for the treatment of amyotrophic lateral sclerosis, and the other is a “dimer-like” version of this structure, **Rilu-2** (**Table 1**). Both molecules showed promising *in vitro* activity and relatively low toxicity in mammalian cells.^40, 46-47^ Consistent with this low toxicity, when **Rilu-2** and **Rilu-6** (0, 100, 200, 500 µM) were introduced intratracheally into wild-type BALB/c mice, none of the animals developed illness or died within 24 hours and there was only a slight increase in total white blood cell count (indicating a negligible inflammatory response; **SI Figure 1**). More recently, we examined these compounds, along with several others containing the same core, to determine whether they reduced multiple virulence-associated phenotypes in pathogenic bacteria without directly affecting viability or growth.^35^ Additionally, we tested the effects of these compounds on resensitization to polymyxin antibiotics in several Gram-negative pathogens and observed substantial reductions in resistance.^48-49^

**Figure 2.**
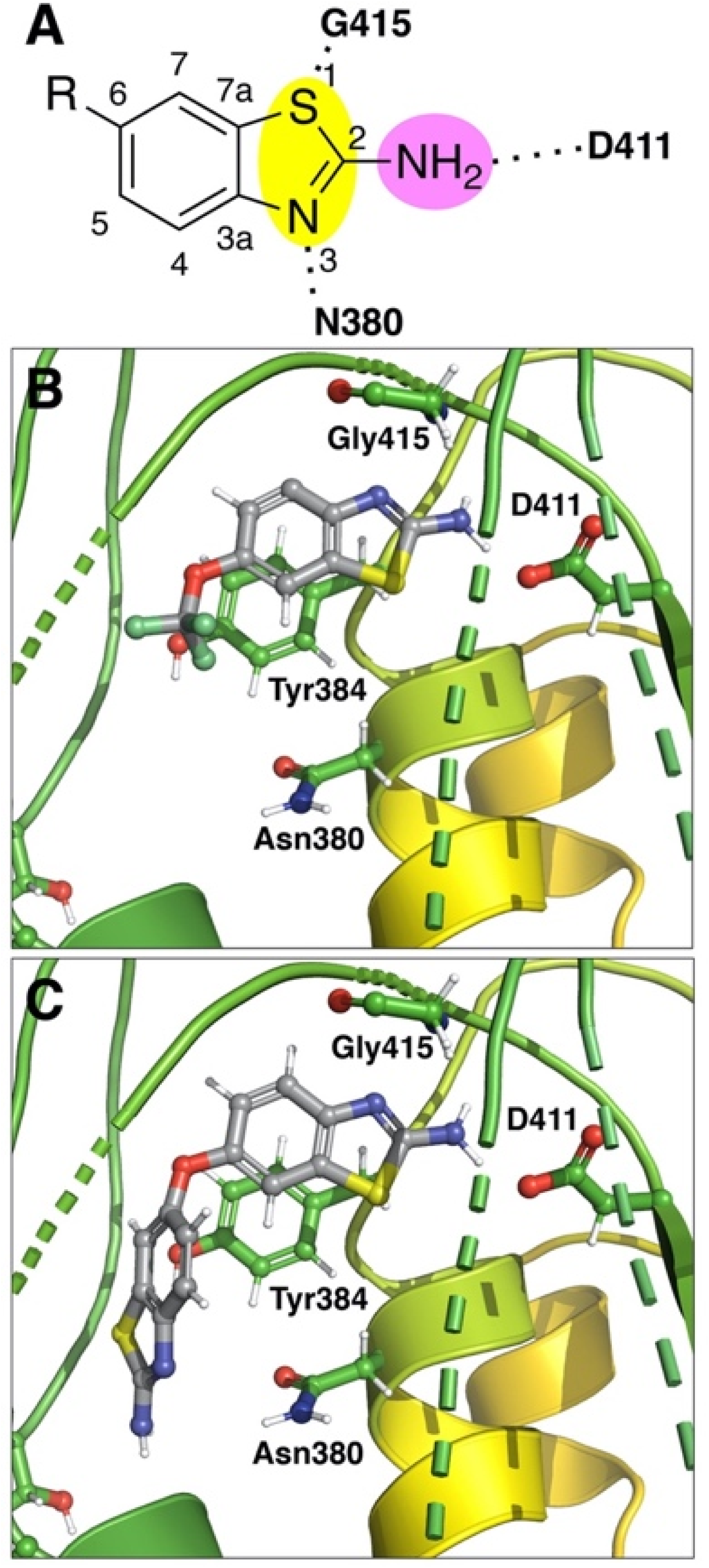
Predicted binding motifs of 2-aminobenzothiazole molecules. A) Numbering of 2- aminobenzothiazole inhibitors and hydrogen bond donor-acceptor-donor motif that is crucial for binding to key residues within the CA-domain binding pocket. Docked binding poses of Rilu-1 (B) and Rilu-2 (C) with HK853 (PDB: 3DGE). Both interact with highly-conserved Asp411 and have π-π stacking interactions with conserved Tyr384. The larger group off of the six-position in Rilu-2 occupies more space within the pocket. Red indicates oxygen, blue indicates nitrogen, yellow indicates sulfur, and gray indicates carbon for defined atoms.

**Table 1.**
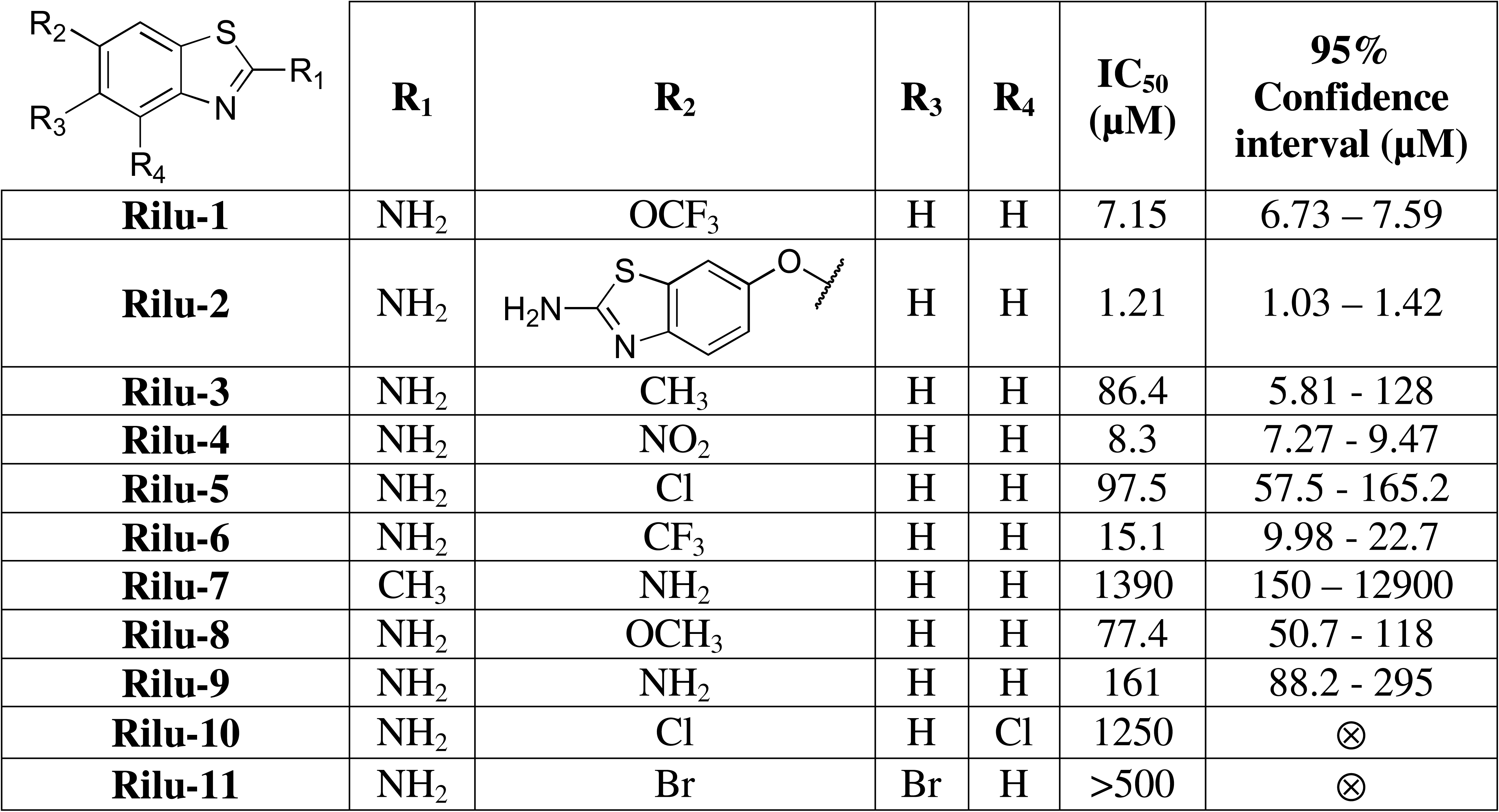
2-Aminobenzothiazole-based inhibitors *in vitro* IC_50_ values for original series. ⊗ indicates no confidence interval available for non-inhibitor.

|  | R<sub>1</sub> | R<sub>2</sub> | R<sub>3</sub> | R<sub>4</sub> | IC<sub>50</sub><br>(μM) | 95%<br>Confidence<br>interval (μM) |
| --- | --- | --- | --- | --- | --- | --- |
| **Rilu-1** | NH<sub>2</sub> | OCF<sub>3</sub> | H | H | 7.15 | 6.73 – 7.59 |
| **Rilu-2** | NH<sub>2</sub> |  | H | H | 1.21 | 1.03 – 1.42 |
| **Rilu-3** | NH<sub>2</sub> | CH<sub>3</sub> | H | H | 86.4 | 5.81 - 128 |
| **Rilu-4** | NH<sub>2</sub> | NO<sub>2</sub> | H | H | 8.3 | 7.27 - 9.47 |
| **Rilu-5** | NH<sub>2</sub> | Cl | H | H | 97.5 | 57.5 - 165.2 |
| **Rilu-6** | NH<sub>2</sub> | CF<sub>3</sub> | H | H | 15.1 | 9.98 - 22.7 |
| **Rilu-7** | CH<sub>3</sub> | NH<sub>2</sub> | H | H | 1390 | 150 – 12900 |
| **Rilu-8** | NH<sub>2</sub> | OCH<sub>3</sub> | H | H | 77.4 | 50.7 - 118 |
| **Rilu-9** | NH<sub>2</sub> | NH<sub>2</sub> | H | H | 161 | 88.2 - 295 |
| **Rilu-10** | NH<sub>2</sub> | Cl | H | Cl | 1250 | ⊗ |
| **Rilu-11** | NH<sub>2</sub> | Br | Br | H | >500 | ⊗ |

*Pseudomonas aeruginosa*, an opportunistic Gram-negative pathogen listed as a priority target for drug development by the WHO and the Center for Disease Control (CDC), showed distinct changes in a number of virulence-related phenotypes when exposed to our inhibitors.^50-51^ This organism is a serious pathogen, particularly among hospitalized patients and those with cystic fibrosis.^52^ *P. aeruginosa* can be particularly challenging to treat because of resistance to most antibiotics.^53-55^ TCSs are involved in the regulation of many virulence pathways in *P. aeruginosa* including swarming motility, which requires both pili and flagella, chemosensing, and the production of rhamnolipid, a surfactant.^56^ These processes are regulated through the following HKs: PilS, FleS, CheA, and BqsS (**Figure 1C**).^13, 16, 56^ Production of toxic secondary metabolites such as pyocyanin (PYO), a blue-green membrane permeable redox-active molecule, the synthesis of which correlates with numerous adverse patient outcomes and detrimental cellular effects,^57-59^ is also regulated by several TCSs, including PhoPQ and GacSA (**Figure 1C**).^13, 60-64^

Treatment of *P. aeruginosa* with our initial leads resulted in significant reductions in swarming behavior, which mimics the pathogen’s movement through mucosal environments such as the lungs, and is linked to greater antibiotic resistance.^63, 65-66^ Additionally, PYO production was markedly decreased. In this report, we further quantify the observed phenotypic changes and describe improvements of the *in vitro* and *in cellulo* activity of our leads based on structure-activity relationship (SAR) studies.^35^

## Results and Discussion

### Design and In Vitro Evaluation of New Compound Library

To improve molecule activity through SAR studies, analogs of our original riluzole-like compounds were synthesized to produce a pan-HK inhibitor that could be used as an anti-virulence agent. We set out to identify which portion(s) of the 2-aminobenzothiazole scaffold were essential for inhibitory activity and which moieties could be modified to improve inhibition and drug-like properties. Our prior work indicated that the hydrogen bond donor-acceptor-donor motif, which is hypothesized to interact with a highly conserved trio of residues (D411, G415, N380 in HK853 from *Thermatoga maritima*), is required for inhibition (**Figure 2**).^35-36^ Molecular docking experiments with **Rilu-1** and **Rilu-2** showed direct interaction of the 2-amino group with the aspartate residue in the G1-box (**Figures 1B** and **2B** and **C**). In addition, we found in our initial work that an analog in which this exocyclic amine is replaced with a methyl group (**Rilu-7**) possesses only mM activity *in vitro* and has little-to-no effect *in cellulo* (**Table 1**).^35^ Based on these observations, we expanded our molecule collection and performed preliminary screening using a gel-based assay to evaluate inhibition of HK activity as previously described (three concentrations up to 500 mM; **SI Figure 2**).^35-36, 40, 67^ We also investigated these compounds for their capacity to cause protein aggregation at relevant concentrations. Protein aggregators were excluded from further study (**SI Figure 3**).

Several analogs that have substituents branching from the exocyclic amine were generated by either direct *N*-alkylation of **Rilu-1** or 2-chloro-6-trifluoromethoxy benzothiazole, which had previously been synthesized.^68^ Neither secondary nor tertiary amines, including the addition of methyl, dimethyl, phenyl, benzyl, and acetyl (**C-1** to **C-6**), exhibited inhibition (**Table 2**). The benzothiazole heteroatom arrangement proved to be important for binding as the replacement of the sulfur with substituted or unsubstituted nitrogen atoms (**C-7** to **C-10**) substantially reduced *in vitro* activity (**Table 2**). Attempts to prepare the 2-aminobenzothiophene or 2-aminoindole analogs proved intractable as the final compounds were unstable. However, molecules that were still protected on the exocyclic nitrogen were tested and proved to be inactive *in vitro* (**SI Figure 2**).

**Table 2.**
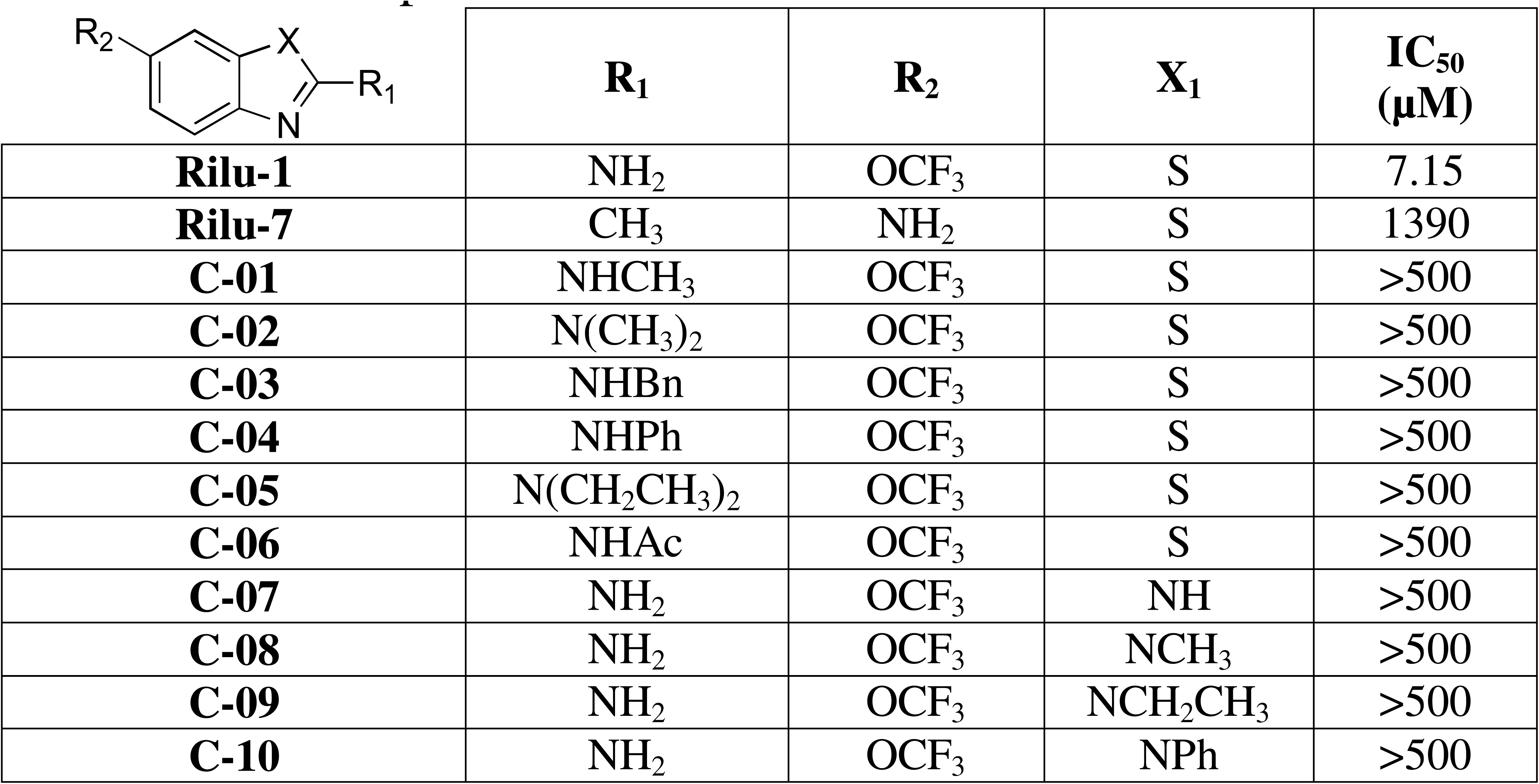
2-Aminobenzothiazole-based inhibitor *in vitro* IC_50_ values for modifications at the 2- amino and sulfur positions.

|  | R<sub>1</sub> | R<sub>2</sub> | X<sub>1</sub> | IC<sub>50</sub><br>(μM) |
| --- | --- | --- | --- | --- |
| **Rilu-1** | NH<sub>2</sub> | OCF<sub>3</sub> | S | 7.15 |
| **Rilu-7** | CH<sub>3</sub> | NH<sub>2</sub> | S | 1390 |
| **C-01** | NHCH<sub>3</sub> | OCF<sub>3</sub> | S | >500 |
| **C-02** | N(CH<sub>3</sub>)<sub>2</sub> | OCF<sub>3</sub> | S | >500 |
| **C-03** | NHBn | OCF<sub>3</sub> | S | >500 |
| **C-04** | NHPh | OCF<sub>3</sub> | S | >500 |
| **C-05** | N(CH<sub>2</sub>CH<sub>3</sub>)<sub>2</sub> | OCF<sub>3</sub> | S | >500 |
| **C-06** | NHAc | OCF<sub>3</sub> | S | >500 |
| **C-07** | NH<sub>2</sub> | OCF<sub>3</sub> | NH | >500 |
| **C-08** | NH<sub>2</sub> | OCF<sub>3</sub> | NCH<sub>3</sub> | >500 |
| **C-09** | NH<sub>2</sub> | OCF<sub>3</sub> | NCH<sub>2</sub>CH<sub>3</sub> | >500 |
| **C-10** | NH<sub>2</sub> | OCF<sub>3</sub> | NPh | >500 |

Our initial SAR work indicated that modifications at the six-position produce the most active compounds (**Figure 2A**). Inhibitors from our original series that were modified at this position, such as **Rilu-2** (IC_50_ = 1.21 µM), were potent in both *in vitro* and *in cellulo* assays.^35^ Some evidence suggests that modifications at other positions may not be favorable given the poor activity of several di-halogenated analogues.^35^ However, data from two new compounds, a derivative of **Rilu-1** in which C-4 is changed to a nitrogen atom (**C-12**, 29.0 µM; **Table 3**) and a molecule that is functionalized at the five-position with a 2-oxazolidinone (**C-13**, 43.4 µM; **Table 3**), indicate that moderate potency can be achieved by functionalization at alternative positions. Docking experiments suggest that the benzene ring in the core of these compounds aligns with Tyr384 and that the addition of an endocyclic nitrogen may impair π-π stacking (**SI Figure 5**).^69-70^

**Table 3.** 2-Aminobenzothiazole-based inhibitor *in vitro* IC_50_ values for modifications on ring other than 6-position. ⊗ indicates no confidence interval available for non-inhibitor. Data for molecules with dose-responsive inhibition are provided in **SI Figure 4**. *Deprotection of benzyl groups resulted in molecule degradation.

|  |  | IC<sub>50</sub><br>(μM) | Confidence<br>interval<br>(μM) |
| --- | --- | --- | --- |
| **C-11\*** |  | >500 | ⊗ |
| **C-12** |  | 29.0 | 17.6 – 49.7 |
| **C-13** |  | 43.4 | 24.7 – 80.3 |

Given the previous success, we focused our effects on the generation of a wide range of molecules with substituent groups at the six-position. We started with several smaller substituents such as trifluoromethyl sulfonyl (**C-14**; **Table 4**). This molecule showed promising *in vitro* activity, with an IC_50_ value against HK853 comparable to that of our original leads (IC_50_= 2.28 µM). The addition of a single NH group to yield the *N*-1,1,1-trifluoromethylsulfonamide derivative (**C-18,** IC_50_ = 21.0 µM) resulted in a 9-fold reduction in activity relative to **C-14**. Analogues with an acetamide moiety connected through an ether linkage also have reduced inhibitory activity with the unmethylated (**C-15,** IC_50_ = 46.8 µM), *N*-methyl (**C-16,** IC_50_ = 50.1 µM), and *N*,*N*-dimethyl functionalized derivatives (**C-17**, IC_50_ = 119 µM) showing a 6- to 16- fold drop in potency. Molecular docking experiments suggest that the unhindered acetamide (**C- 15**) may result in a “flipped” binding orientation (**SI Figure 6**) where the acetamide moiety interacts with the conserved aspartate (D411) instead of the exocyclic nitrogen from the 2- aminobenzothiazole core. The methylated versions (**C-16**, **C-17**) are presumably too bulky and remain in the expected orientation. A *N*-trifluoroethylated amide derivative, with the nitrogen placed closer to the aromatic ring (**C-19,** IC_50_ = 10.3 µM), is more potent than the related derivatives and almost equipotent to **Rilu-1**. Overall, we found that molecules with aliphatic groups at the six-position have reduced potency compared with aromatic analogs [**Rilu-3**, **-8**, **C- 15, ™16, ™17**, **C-20** (IC_50_ = 20.0-119 µM) versus **C-21, ™22, ™23, ™24** (IC_50_ = 1.68-12.2 µM)].

**Table 4.**
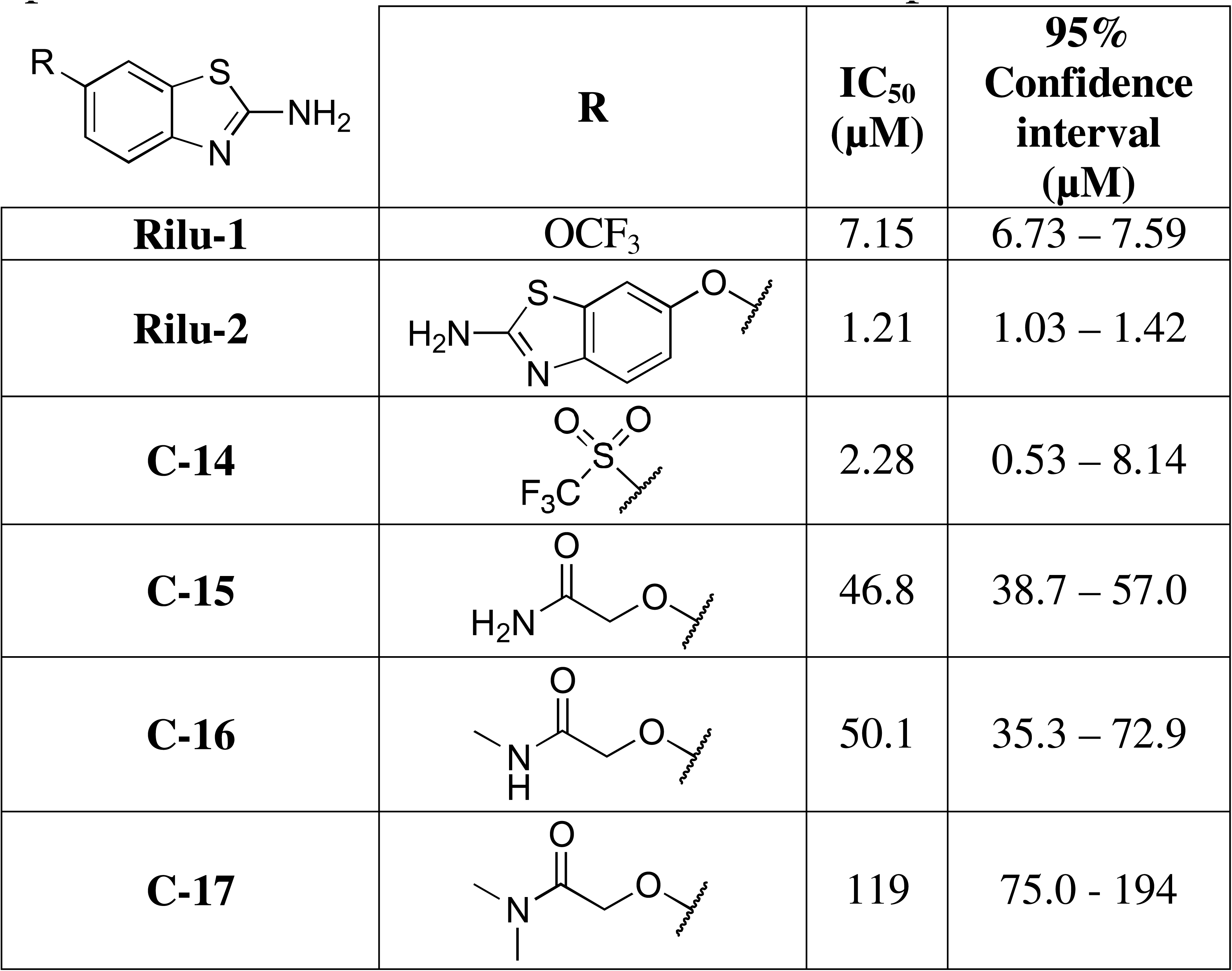
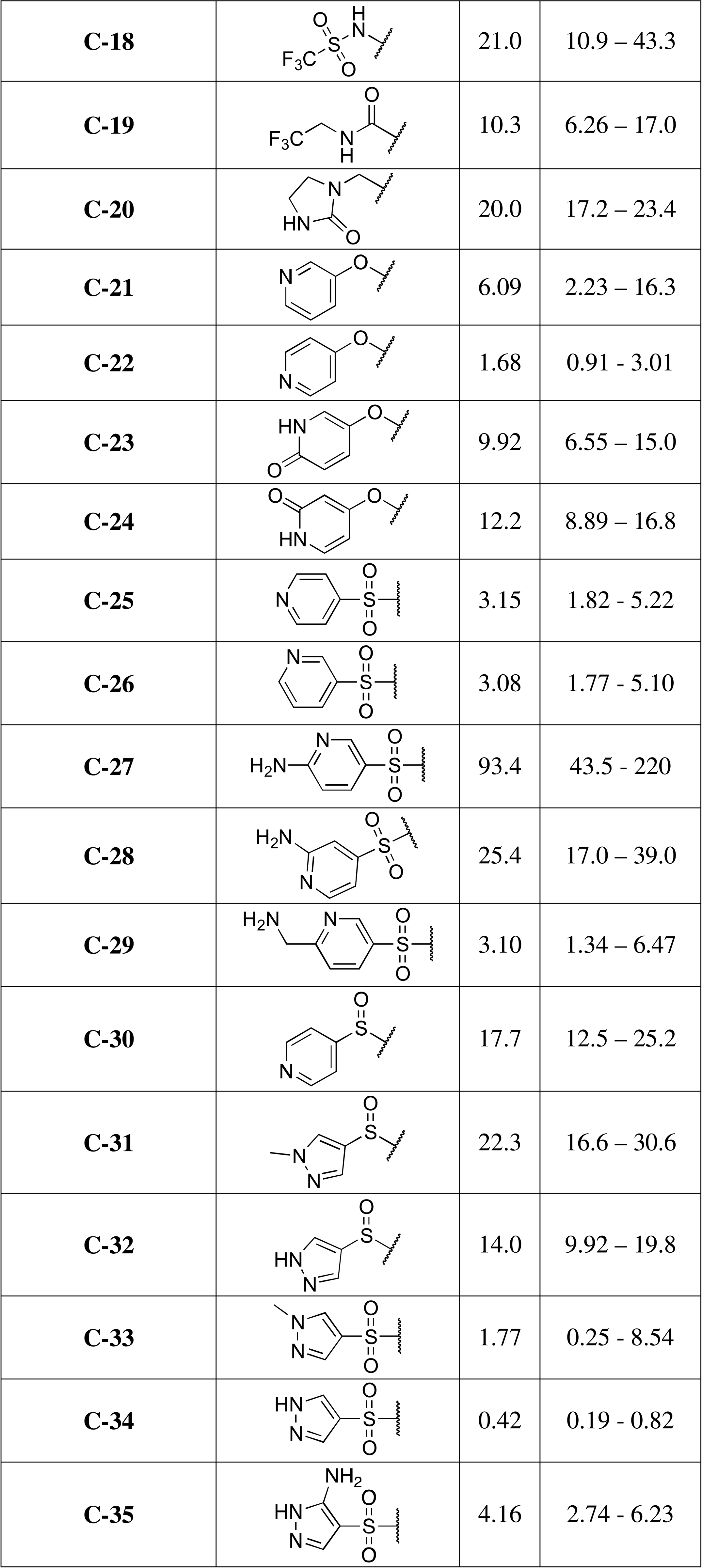
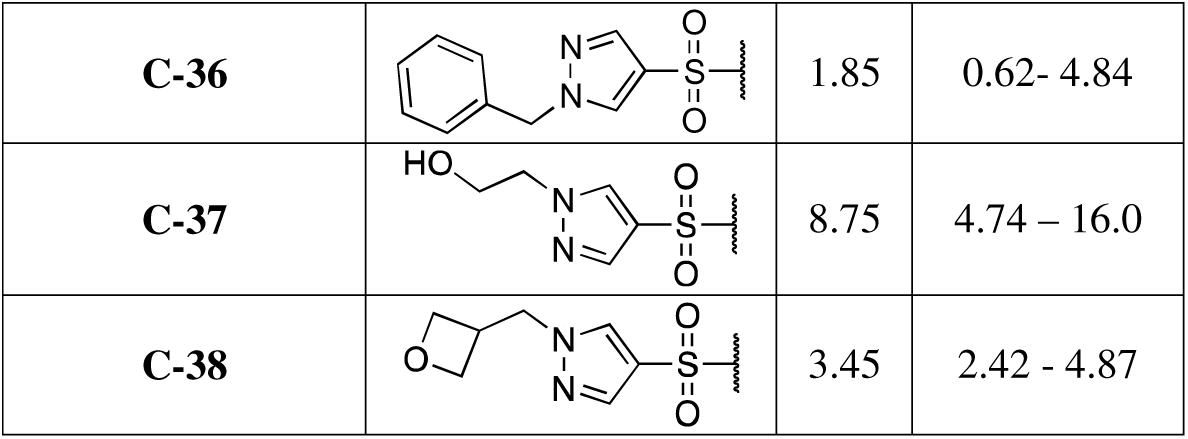
2-Aminobenzothiazole-based inhibitor *in vitro* IC_50_ values for modifications at 6- position. Data for molecules with dose-responsive inhibition are provided in SI Figure 4.

Our data also indicate that a large, heteroaromatic substituent on the six-position, as in **Rilu-2**, can yield more active molecules. Indeed, **Rilu-2** has an IC_50_ value that is six-fold more potent than that of **Rilu-1**, which may be attributed to an increased number of interactions within the binding pocket by the larger molecule, such as additional favorable π-π interactions. For example, docking studies indicate interactions with a tyrosine residue (Y429 PDB: 3DGE) in the flexible lid region (**Fig. 2B** and **2C**). Given this promising result, we sought to explore further analogues with larger substituents, including those with heteroaromatic moieties that vary in size, linker composition, heteroatom content, and substituents. Molecules with ether-linked oxypyridinyl substituents showed activity similar to **Rilu-2** (**C-21**, IC_50_ = 6.09 µM; **C-22**, IC_50_ = 1.68 µM). Addition of a carbonyl to increase polarity and hydrogen bond acceptor potential was found to be moderately disfavored (comparison of pyridinyl analogues, **C-21**, IC_50_ = 6.09 µM; **C-22**, IC_50_ = 1.68 µM; and pyridinone analogs, **C-23**, IC_50_ = 9.92 µM; **C-24**, IC_50_ = 12.2 µM). Analogues with sulfone and sulfoxide groups linked to heteroaromatic moieties were also evaluated. Sulfone-containing molecules (**C-25**, IC_50_ = 3.15 µM; **C-26**, IC_50_ = 3.08 µM) are comparably potent to their ether counterparts (**C-21**, IC_50_ = 6.09 µM; **C-22**, IC_50_ = 1.68 µM). The addition of an exocyclic amine to **C-25** and **C-26** caused a drop in potency (**C-27**, IC_50_ = 93.4 µM; **C-28**, IC_50_ = 25.4 µM). Sulfones are favored over the sulfoxides for inhibitory activity as **C-30 (**IC_50_ = 17.7 µM) is almost six-fold less potent than its sulfone counterpart in **C-25** (IC_50_ = 3.15). In addition, **C-31** (IC_50_ = 22.3 µM) and **C-32** (IC_50_ = 14.0 µM) are 10 to 20-fold less potent than the related sulfone-containing analogs **C-33 (**IC_50_ = 1.77 µM) and **C-34** (IC_50_ = 0.42 µM). **C-34** shows sub-micromolar activity, and additional modifications to the pyrazole ring, with small (e.g., methyl) or large (e.g., benzyl) groups all resulting in reduced potency to the single digit micromolar range (**C-33**, IC_50_ = 1.77 µM; **C-35**, IC_50_ = 4.16 µM; **C-36**, IC_50_ = 1.85 µM; **C-37**, IC_50_ = 8.75 µM; **C-38**, IC_50_ = 3.45 µM).

Overall, we confirmed our previous hypothesis that the heteroatom display within the 2- aminobenzothiazole core cannot be modified without affecting inhibition. The 2-amino group is crucial for *in vitro* activity, while the other two heteroatoms also appear to play significant roles. Our data also indicate that substitutions on this scaffold are most beneficial at the six-position, with the caveat that the five-position has not been extensively explored. To date, the most potent inhibitor is decorated with a pyrazole at this position. Finally, current evidence indicates that only one substitution on the benzyl ring is tolerated. However, the variety of substituents that are tolerated is wide and additional studies will be required to further assess functionalities that optimize both protein-inhibitor interactions and whole-cell activities (see below).

### Investigation of inhibitor binding using NMR-based methods

To gain further insights into the interactions between our inhibitors and the ATP-binding site within an HK, we employed NMR. Prior to initiating these experiments, several intrinsic parameters were assessed by 1D ^1^H to ensure protein suitability for binding studies including protein folding, stability over time, DMSO tolerance, and promiscuity (**SI Figure 7**).

### Inhibitor binding by ligand-detected NMR, 1D ^1^H/^19^F

Differential line-broadening (DLB), T_2_-Carr-Purcell-Meiboom-Gill (CPMG), and saturation transfer difference (STD) experiments were used to evaluate binding to HK853 with **C-12**, **14**, **17**, **30**, **34**, and **Rilu-1** as this molecule collection covers a range of potencies and functionalities (**Figure 3A**).^24, 71^ Free-state behaviors of each compound in buffer were also evaluated as a control to avoid false positive results due to compound misbehavior, such as aggregation, using 1D ^1^H and T_2_-CPMG.^72^ We found that the DLB experiments show line-broadening of the small molecule resonances upon the addition of HK853. In addition, T_2_- CPMG experiments indicate faster decay of the inhibitor resonances with increasing delay times when in the presence of HK853 (red traces) as compared to the free state (blue traces; **Figure 3B**). These data confirm inhibitor-HK853 interactions as the presence of the protein causes the small molecule to inherit its slow-tumbling behavior. STD experiments also confirm binding given that there is a measurable difference between the ON- and OFF-resonance saturation experiments (e.g., **Rilu-1** in **Figure 3B**). Three of the six molecules contain a CF_3_ group (**C-12**, **14**, **Rilu-1**), therefore ^19^F DLB was also used to evaluate binding and further indicated interactions (**Figure 3A**).

**Figure 3.**
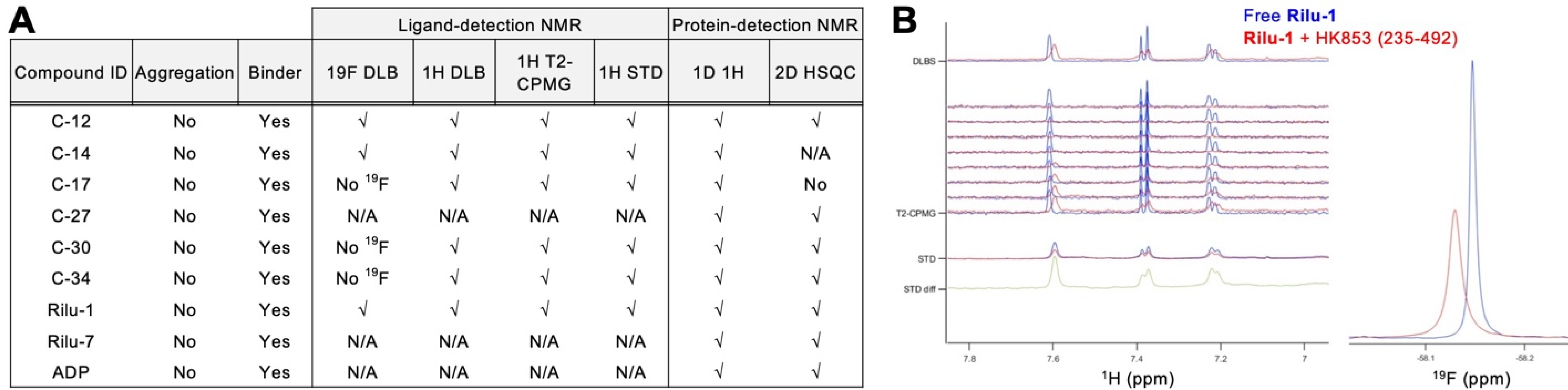
Data obtained for binding by ligand- and protein-detection NMR. **A**. Binding evaluated by ligand detection using ^1^H and ^19^F DLB, T_2_-CPMG and STD. 1D ^1^H and 2D HSQC were used to evaluate binding by protein detection. The symbol (√) is used when binding is observed by the selected method. N/A indicates that the experiment was not performed. **B**. Binding of **Rilu-1** to HK853 (residues 232-489). Differential line broadening and shifting (DLBS) and T_2_-CPMG experiments show the aromatic region of 1D ^1^H NMR spectra of 300 µM of **Rilu-1** in its free state (blue trace) and in presence of 15 µM of HK (red trace). STD indicates the ON and OFF resonance spectra of 300 µM of **Rilu-1** in presence of 15 µM of HK (blue and red traces, respectively), the differential spectra (STD diff) are represented in green. ^19^F Differential line broadening experiment shows ^19^F NMR spectra of 300 µM of **Rilu-1** in its free state (blue trace) and in presence of 15 µM of HK (red trace).

### Inhibitor binding by protein-detected NMR

Isotope-labeled HK853 (^15^N, residues 235-492) was used to measure chemical shift perturbations upon the addition of seven inhibitors (**C-12**, **17**, **27**, **30**, **34**, **Rilu-1**, **7**) and ADP by 1D ^1^H and 2D ^1^H-^15^N TROSY-HSQC. The HK is expected to be a dimer and the TROSY variant of the ^1^H-^15^N HSQC experiment was performed for improved signal-to-noise. Although the quality of the 2D spectrum of apo HK853 is limited in terms of peak count and resolution due to the high molecular weight of the dimer, it was possible to assess compound binding and evaluate the protein fingerprint upon the addition of compounds (**SI Figure 8**).

Combined NMR data obtained by 1D and 2D experiments show that ADP and the seven inhibitors bind to the protein. Analysis of the protein ^15^N-HSQC fingerprint shows that changes were observed with five of the seven compounds evaluated by 2D HSQC (**SI Figure 8**). As expected based on the IC_50_ value (1390 µM), **Rilu-7** only shows small changes by 1D ^1^H NMR and a very small change by 2D HSQC, suggesting very weak binding to HK853 (**Figure 3A**, **SI Figure 8**). The five potent inhibitors (IC_50_ < 100 µM) show similar chemical shift perturbation fingerprints, suggesting that they affect the same region of the protein via direct binding or allosteric changes. They also share similar chemical shift perturbation with that of ADP, which further suggests that they bind a similar region (**Figure 3A**, **SI Figure 8**).

### Competition studies: ADP versus inhibitors

Titration experiments using 1D ^19^F NMR were performed to determine if compounds **C- 14** and **Rilu-1** are ADP competitive in HK853 and could be monitored by their CF_3_ group. Spectra of these compounds free in buffer were compared to data acquired in the presence of protein to identify the chemical shift of their bound state (**Figure 4A and B**). The addition of incremental amounts of ADP to the inhibitor-protein complexes showed that ADP displaced these ligands, suggesting that **C-14** and **Rilu-1** are likely binding to the same or close site as ADP. Under the same ratio of ADP:compound:HK853, these data indicate that more **C-14** is displaced than **Rilu-1**, signifying that the latter may have a tighter binding affinity for HK, consistent with *in vitro* potency data.

**Figure 4.**
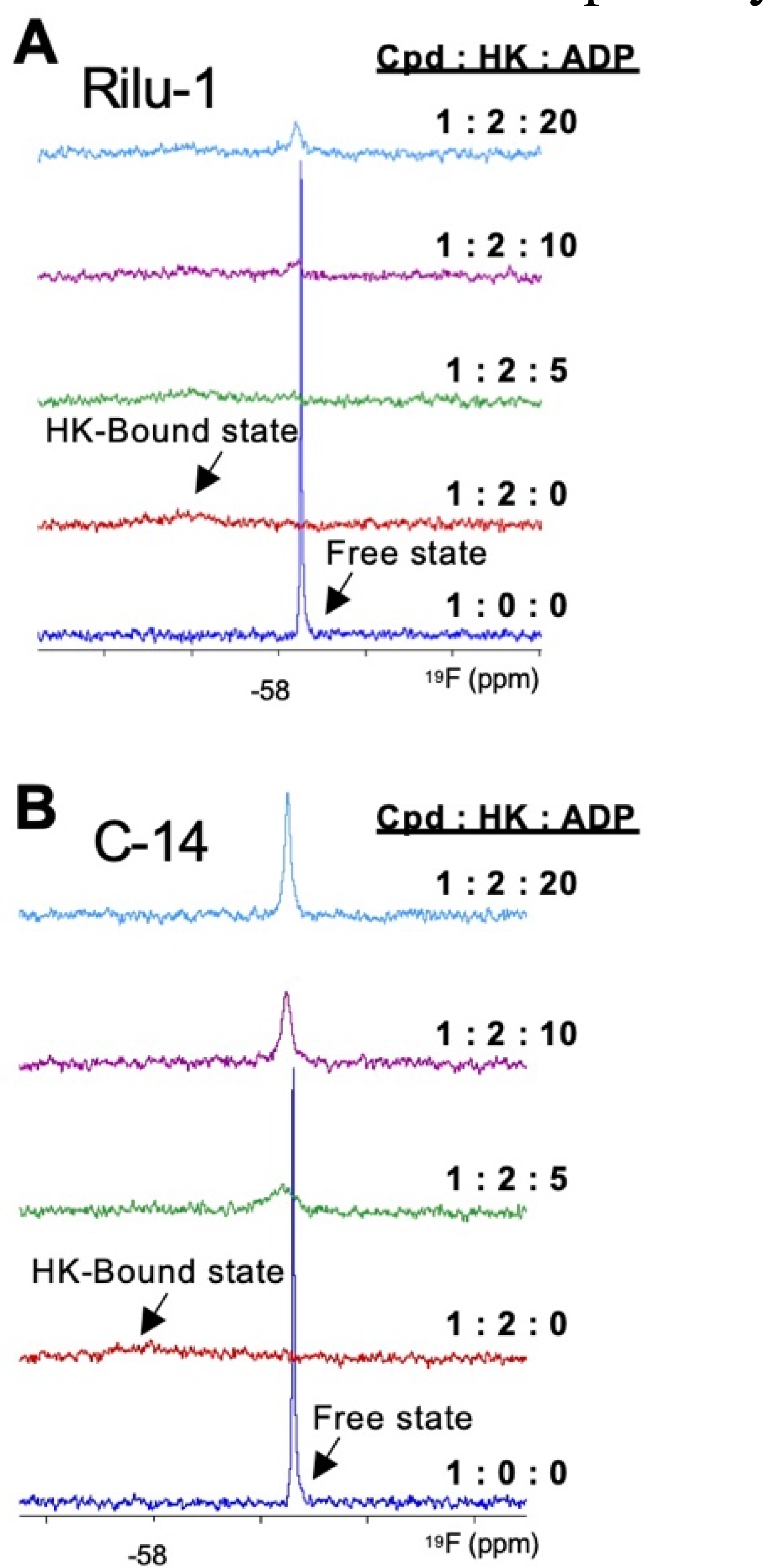
Competition studies with ADP for binding to HK by ^19^F NMR. Titration experiments performed by 1D ^19^F NMR with **Rilu-1** (**A**) and **C-14** (**B**) for binding to HK (232- 489) in the absence and in presence of ADP. For each titration, a 1D ^19^F spectrum of the compound was acquired free in buffer (blue), in presence of HK protein (red), and with increasing molar equivalent ratios of ADP (green to purple to light blue). The free and bound states of both compounds are identified on the spectra.

### Cell-Based Assays

#### Bacterial Growth

To evaluate the growth dynamics of *P. aeruginosa* treated with inhibitors, we diluted overnight cultures (1:100) into fresh media and a panel of inhibitors (200 μM) and measured growth spectrophotometrically (OD_600_) over 20 h. Average growth data from three biological replicates for all inhibitor-treated samples were compared to the DMSO control through a one-way ANOVA Brown and Forsythe test (GraphPad Prism 8.0). No statistically significant differences in the specific growth rates were observed between the inhibitors and DMSO control (**SI Table S1**). To more thoroughly evaluate potential differences in growth characteristics, we assessed the total area under the curve (AUC). The average AUC for each inhibitor (**Figure S9A**) was compared to the DMSO control on each plate. Statistically significant differences in AUC were obtained for inhibitors **C-14**, **C-29**, **C-30**, and **C-37**. Examination of the growth curves for **C-29** and **C-30** shows a lower final OD_600_ (**Figure S9B**) compared to the DMSO control, indicating some bactericidal activity. Conversely, **C-14** and **C-37** treatment result in a higher final OD_600_ indicating a higher final growth rate than death rate (**Figure S9C**). Phenotypic assay data were normalized to OD_600_ in all cases except the agar plate-based motility assay, which is not feasible.

#### Bacterial Mobility

Swarming is a complex motility phenotype that is associated with increased antibiotic resistance and production of virulence factors while also being correlated with poor patient outcomes.^63, 65-66^ Because of the pathogenic importance of *P. aeruginosa* swarming, a phenotype that is modulated by multiple HKs (**Fig. 1C**),^13, 61^ we tested the ability of our most potent inhibitors to affect this phenotype as one metric of their potential as anti-virulence agents.^35^ This coordinated, multicellular phenotype can be mimicked in a laboratory setting on low-nutrient and low-percentage agar plates, which provides a milieu resembling the pulmonary epithelial mucosa where *P. aeruginosa* infection occurs.^73^ There are multiple methods to quantify this phenotype, including thickness and number of tendrils, rhamnolipid quantification, and swarm speeds though these can be tedious to measure and can prove difficult to compare.^13, 61, 63-64, 74^ To incorporate these different aspects of motility into a single metric and enable simultaneous evaluation of multiple inhibitors, we quantified the area of overnight growth.^13, 56, 74^ Because quantities of inhibitor compounds were limited, each was examined at a single concentration of 200 µM. Given the challenges of effectively penetrating the cell envelope of Gram-negative organisms, such as *P. aeruginosa*, we hypothesized that even though most members of our compound library were effective protein inhibitors *in vitro*, there may be large disparities in cell-based assays related to the ability of each molecule to accumulate within the organism.^75-80^

From our initial list of hits, **Rilu-2** proved the most potent, reducing the swarm area by >90% relative to a DMSO control (**Table 5** and **Fig. 5**). Analogues with a smaller substituent at the six-position had more modest activity, with **Rilu-1** (36% reduction) and **C-15** (4.2%) causing reduction while **C-16** resulted in moderate activation (−25%). We found that the 2-amino group was critical for the inhibition of swarming activity, consistent with our previous hypothesis about the necessity of this moiety (e.g., **Rilu-7**). Beyond these results, concrete structural trends were less evident than for *in vitro* inhibition. For example, while the oxypyridine analogues (**C-21**, IC_50_ = 6.09 µM; **C-22**, IC_50_ = 1.68 µM) were slightly more potent *in vitro* than the oxypyridone analogues (**C-23**, IC_50_ = 9.92 µM; **C-24**, IC_50_ = 12.2 µM), **C-24** caused an 80% swarm reduction relative to only a 48% and 40% reduction observed with **C-21** and **C-22**, respectively (**Table 5; SI Figure 10** and **SI Figure 11**). Although **C-23** and **C-24** vary only in the orientation of the heteroatoms on the ring and have similar *in vitro* activity, **C-23** produced only a 26% reduction in swarm area. While **C-34** is the most potent *in vitro* inhibitor, its effect on this phenotype was modest, with a 26% reduction in swarm area. Moreover, although the addition of the exocyclic nitrogen to the pyrazole ring in **C-34** (IC_50_ = 0.42 µM) yielding **C-35** (IC_50_ = 4.16 µM) proved unfavorable *in vitro*, this modification improved its effectiveness in this assay, promoting an 87% reduction in swarming relative to the 26% reduction caused by **C-34** (**Table 5**). Finally, **C- 29** showed promise with a 92% reduction in swarm area, while several other similar analogues were largely inactive (e.g., **C-25-28**). While more work is required to fully understand trends of the phenotypic changes caused by different motifs in the inhibitor series, these data indicate that larger heteroaromatic groups will likely be the most effective. The most potent of these analogues showed effects similar to or were more potent than that of mutant strains of *P. aeruginosa* lacking HKs such as RetS, PilS, and PhoQ, which are important in cytotoxicity and infectivity in cell and murine models (**SI Figure 10** and **SI Figure 11**).^81-84^

**Figure 5.**
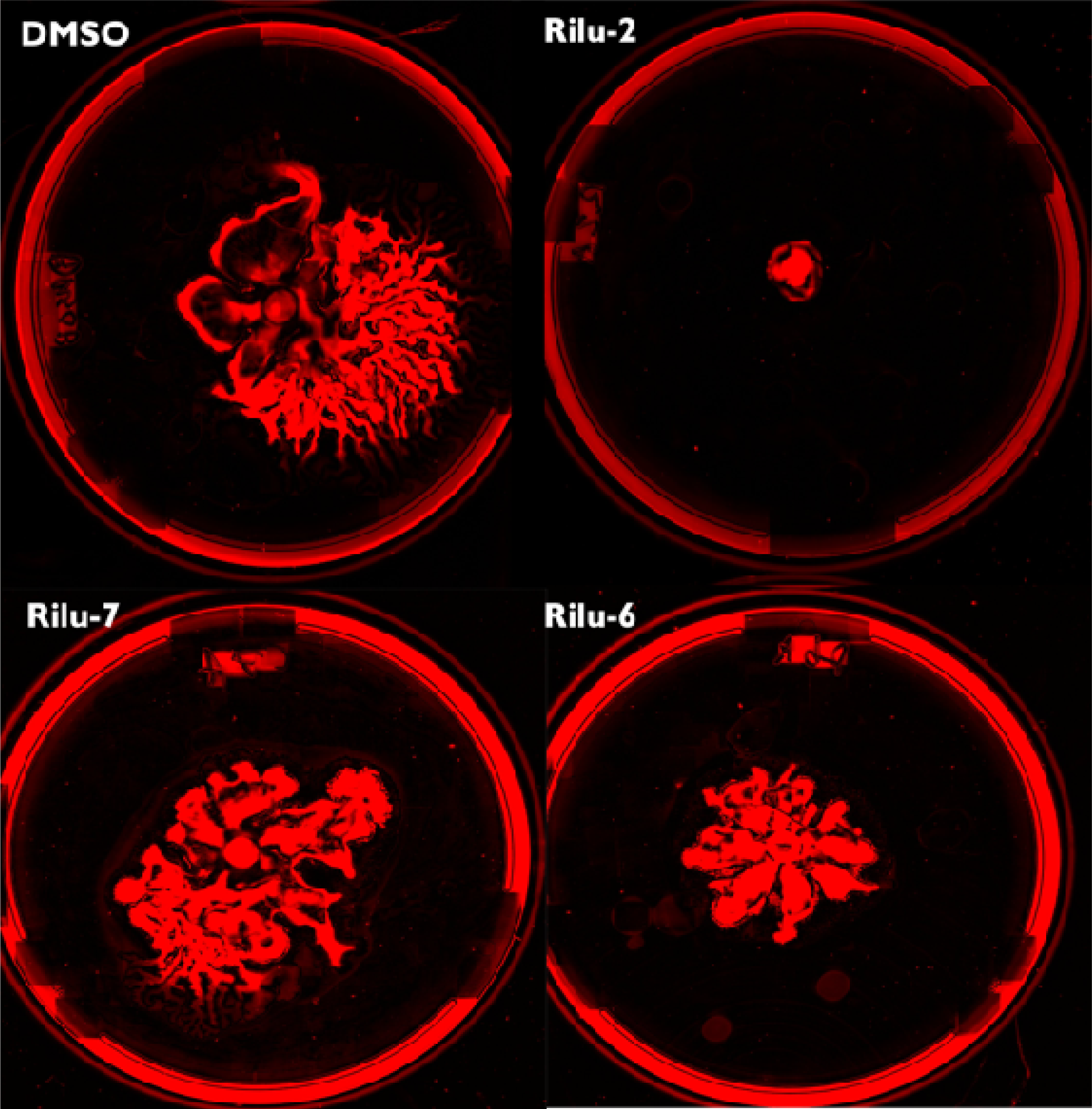
Swarming motifs of PA14 strain on modified fastidious anaerobe broth (FAB) agar. Comparison of DMSO vehicle-control to plates contain 100 µM of **Rilu-2**, **6** or **7**. **Rilu-2** resulted in a substantial decrease in the swarming capabilities of the bacteria, but did not prevent growth. Plates were imaged using a Typhoon FLA 9500 scanner (GE healthcare) on DY-520XL filter setting. Images were redshifted to improve contrast of bacteria tendrils from agar plate and analyzed using ImageJ (NIH).

**Table 5.**
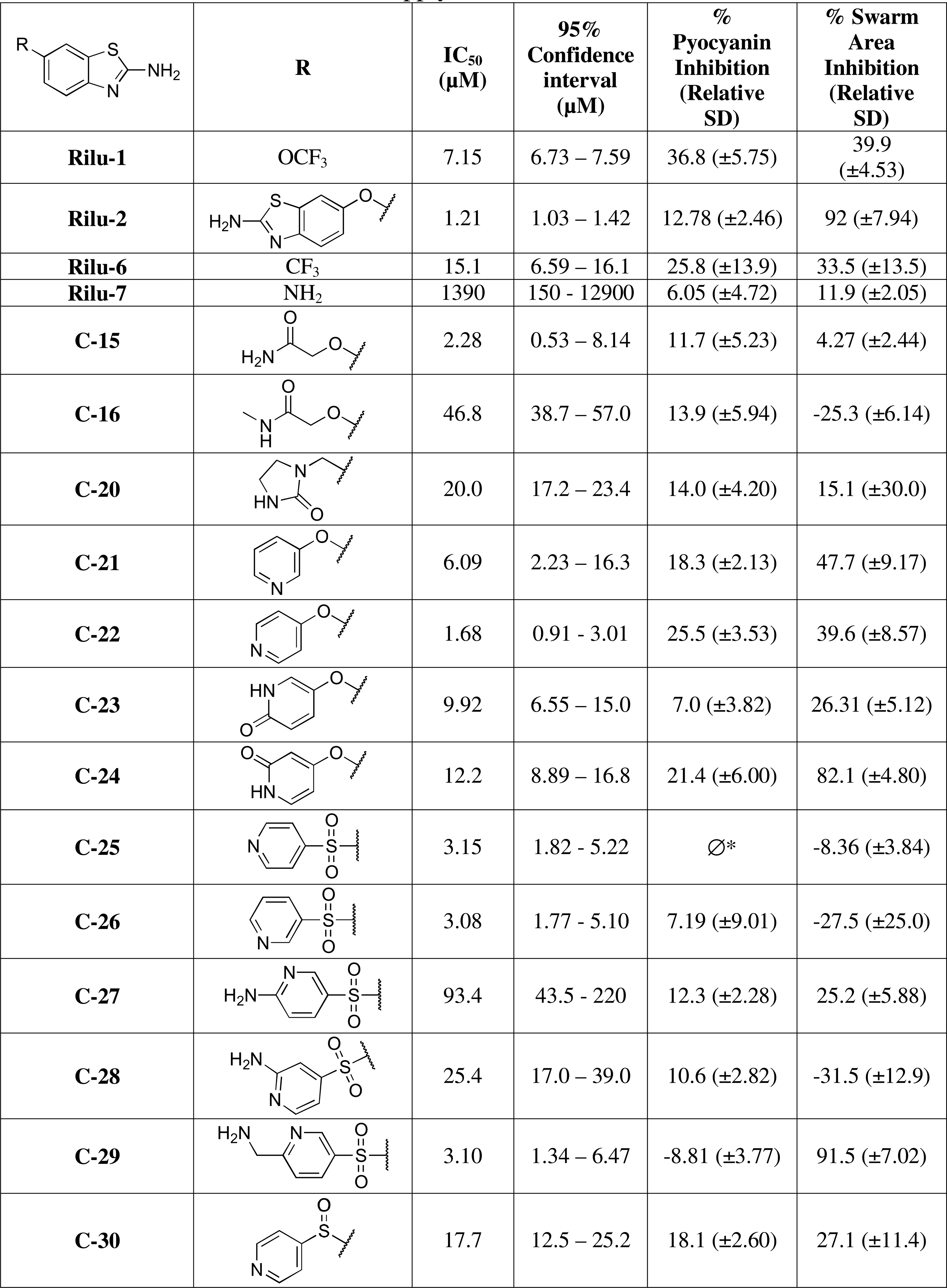
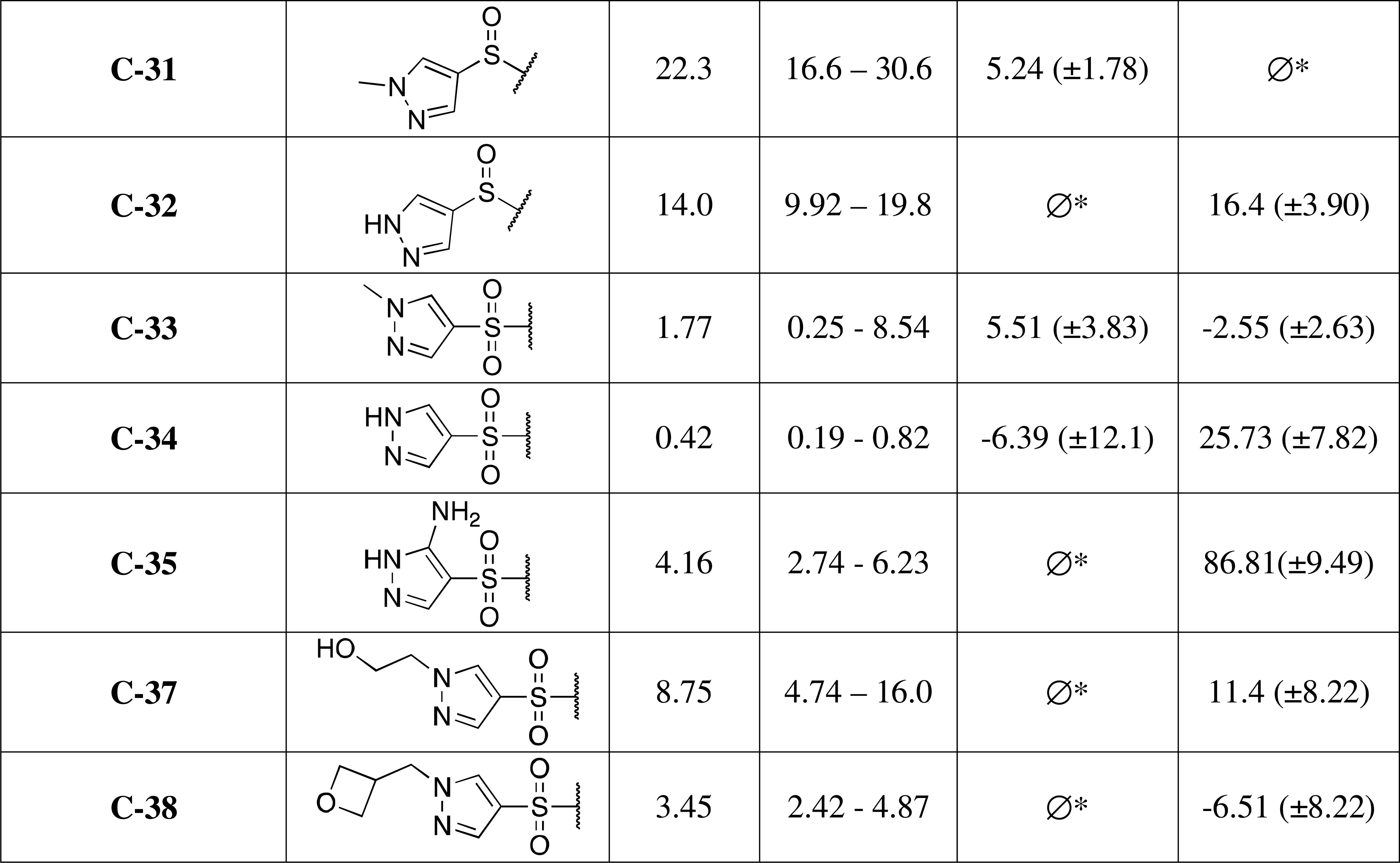
2-Aminobenzothiazole-based inhibitors *in vitro* IC_50_ values and *in cellulo* inhibition percent of pyocyanin production and swarm area relative to DMSO vehicle control. ∅* indicates data not available due to a limited supply of the inhibitor.

|  | R | IC<sub>50</sub><br>(μM) | 95%<br>Confidence<br>interval<br>(μM) | %<br>Pyocyanin<br>Inhibition<br>(Relative<br>SD) | % Swarm<br>Area<br>Inhibition<br>(Relative<br>SD) |
| --- | --- | --- | --- | --- | --- |
| **Rilu-1** | OCF<sub>3</sub> | 7.15 | 6.73 – 7.59 | 36.8 (±5.75) | 39.9<br>(±4.53) |
| **Rilu-2** |  | 1.21 | 1.03 – 1.42 | 12.78 (±2.46) | 92 (±7.94) |
| **Rilu-6** | CF<sub>3</sub> | 15.1 | 6.59 – 16.1 | 25.8 (±13.9) | 33.5 (±13.5) |
| **Rilu-7** | NH<sub>2</sub> | 1390 | 150 - 12900 | 6.05 (±4.72) | 11.9 (±2.05) |
| **C-15** |  | 2.28 | 0.53 – 8.14 | 11.7 (±5.23) | 4.27 (±2.44) |
| **C-16** |  | 46.8 | 38.7 – 57.0 | 13.9 (±5.94) | -25.3 (±6.14) |
| **C-20** |  | 20.0 | 17.2 – 23.4 | 14.0 (±4.20) | 15.1 (±30.0) |
| **C-21** |  | 6.09 | 2.23 – 16.3 | 18.3 (±2.13) | 47.7 (±9.17) |
| **C-22** |  | 1.68 | 0.91 - 3.01 | 25.5 (±3.53) | 39.6 (±8.57) |
| **C-23** |  | 9.92 | 6.55 – 15.0 | 7.0 (±3.82) | 26.31 (±5.12) |
| **C-24** |  | 12.2 | 8.89 – 16.8 | 21.4 (±6.00) | 82.1 (±4.80) |
| **C-25** |  | 3.15 | 1.82 - 5.22 | Ø* | -8.36 (±3.84) |
| **C-26** |  | 3.08 | 1.77 - 5.10 | 7.19 (±9.01) | -27.5 (±25.0) |
| **C-27** |  | 93.4 | 43.5 - 220 | 12.3 (±2.28) | 25.2 (±5.88) |
| **C-28** |  | 25.4 | 17.0 – 39.0 | 10.6 (±2.82) | -31.5 (±12.9) |
| **C-29** |  | 3.10 | 1.34 – 6.47 | -8.81 (±3.77) | 91.5 (±7.02) |
| **C-30** |  | 17.7 | 12.5 – 25.2 | 18.1 (±2.60) | 27.1 (±11.4) |
| **C-31** |  | 22.3 | 16.6 – 30.6 | 5.24 (±1.78) | Ø* |
| **C-32** |  | 14.0 | 9.92 – 19.8 | Ø* | 16.4 (±3.90) |
| **C-33** |  | 1.77 | 0.25 - 8.54 | 5.51 (±3.83) | -2.55 (±2.63) |
| **C-34** |  | 0.42 | 0.19 - 0.82 | -6.39 (±12.1) | 25.73 (±7.82) |
| **C-35** |  | 4.16 | 2.74 - 6.23 | Ø* | 86.81(±9.49) |
| **C-37** |  | 8.75 | 4.74 – 16.0 | Ø* | 11.4 (±8.22) |
| **C-38** |  | 3.45 | 2.42 - 4.87 | Ø* | -6.51 (±8.22) |

#### Pyocyanin Production

Upon treatment with our leads, we also observed major changes in the production of several secondary metabolites including pyocyanin (PYO), a redox-active toxin secreted by *P. aeruginosa*. This metabolite plays a role in signaling through the quorum sensing network, as well as crossing biological membranes to cause oxidative damage to the host.^57-59^ PYO is found in much higher concentrations in the lungs of cystic fibrosis patients with poorer clinical outcomes including declines in pulmonary function.^85^ As previously described, this phenotype is associated with multiple TCSs through control of various metabolic enzymes in *P. aeruginosa.*^14, 56, 60, 86^ We analyzed several of our most potent *in vitro* analogues following overnight treatment using an absorbance-based assay as PYO has inherent absorbance at 690 nm.^87-89^ Among our initial hit series **Rilu-2** was largely ineffective, while **Rilu-1** and **Rilu-6** had the greatest effect on PYO production with 37% and 26% reductions compared to DMSO, respectively (**Table 5**; **SI Figure 12**). None of the tested second-generation analogues exhibited greater or equal potency to that of **Rilu-1** with the most active being **C-21**, **C-22**, and **C-24** (18%, 26%, and 21% decreases respectively, **Table 5**). Next, we performed quantitative LC-MS analysis following longer treatments, 24 and 48 hr, and evaluated PYO, phenazine-1-carboxamide (PCN), and phenazine-1-carboxylic acid (PCA). These data show little to no effect on metabolite production with the exception of a dramatic decrease in PCN at 24 hr upon treatment with **Rilu-1** and **Rilu-2** (**SI Figure 13**). However, PCN production was completely or largely (>75%) recovered by 48 hr (**SI Figure 14**). These results indicate that while some of our inhibitors have an initial effect on metabolite production, the cells recover their ability to generate these molecules in longer exposure experiments.

#### Investigation of Cytotoxicity in Eukaryotic Cells

Our original series of molecules had modest cytotoxicity in Vero 76 cells at concentrations used in several of the assays described herein (200 µM).^40^ While our data indicate that this was not detrimental in initial animal studies, as treated mice remained healthy and we did not observe substantial effects on inflammatory response (**SI Figure 1**), improved eukaryotic tolerance to address potential safety concerns for these molecules is desirable. We employed the XTT cytotoxicity assay in Hep G2 and A549 cells to evaluate several molecules from the initial library, **Rilu-1**, **Rilu-2**, **Rilu-6**, and **Rilu-7**, as well as the three compounds that show the most promising activity in the swarming assay, **C-24**, **C-29**, and **C-35**. These cell lines were chosen for models of hepatic (Hep G2) and airway (A549) cytotoxicity.^90-91^

Data in Hep G2 for **Rilu-1** and **Rilu-2** are consistent with our previous results in Vero 76 cells, showing greater than fifty percent cytotoxicity at concentrations above 250 µM and almost complete cell death at 500 µM (**SI Figure 15**).^40^ In the A549 cell line, these two compounds showed only 30% cytotoxicity at 250 µM and were close to 80% at 500 µM. **Rilu-6** behaved comparably while **Rilu-7**, a similar structure with no substantive activity either *in vitro* or in our phenotypic assays, has modest toxicity only at 500 µM in both cell lines (**SI Figure 16**). Favorably, the second generation inhibitors that we assessed had little to no effect on cell viability at 250 µM and were substantially less cytotoxic at 500 µM, with none causing more than 50% cell death at this concentration. These data indicate that in addition to identifying molecules with equivalent or improved activity in anti-virulence assays, we have also reduced the potential of these molecules to negatively impact eukaryotic cells.

### Potent Analog Synthetic Details

The library of compounds designed for further investigation from the original benzothiazole core cover a wide span of chemical substituents and modifications. The synthetic routes for the most potent analogues in our *in vitro* assay, **C-14** (2.28 μM), **C-22** (1.68 μM), **C-25** (3.15 μM), **C-26** (3.08 μM), **C-33** (1.77 μM), **C-34** (0.42 μM), and **C-36** (1.85 μM), are described here (Scheme 1) in addition to the most potent analogues *in cellulo* **C-24**, **C-35**, and **C-29**. Lastly, synthesis of **C-12** is included given its important role in the NMR experiments. Additional details about all synthetic strategies are provided in the Supporting Information.

A close analogue of **Rilu-1**, alteration of the aromatic ring fused to the thiazole group to pyridine gives **C-12**. To synthesize **C-12,** 5-(trifluoromethoxy)pyridin-2-amine first underwent a standard radical bromination with NBS followed by the formation of the carbamothioylbenzamide through the addition of benzoyl isothiocyanate. The carbamothioylbenzamide was then hydrolyzed to the free thiourea and finally cyclized upon the addition of sodium hydride to afford the final product, **C-12** (Scheme 1A). Synthesis of **C-14** begins with *tert*-butyl *N*-[6-(trifluoromethylsulfanyl)-1,3-benzothiazol-2-yl]carbamate, which is then oxidized to the sulfone utilizing sodium periodate followed by deprotection of the Boc group yielding **C-14** (Scheme 1B). **C-22** and **C-24** are both synthesized by the one-step Hugerschoff reaction utilizing 4-(4-pyridyloxy)aniline and 4-(4-aminophenoxy)-1*H*-pyridin-2- one as their starting materials, respectively (Scheme 1C). **C-26** synthesis is more complex and begins with a cross-coupling of 4-nitrobenzenethiol and 3-bromopyridine with Pd_2_(dba)_3_ and Xantphos to yield the desired thioether, which was reduced with zero-valent iron, followed by the Hugerschoff reaction and a final oxidation step by sodium periodate to yield the desired product. Very similar to **C-26**, is the route for **C-29** with minor alterations (Scheme 1D). The initial Pd_2_(dba)_3_ and Xantphos cross coupling is performed with 4-nitrobenzenethiol and 5- bromopyridine-2-carbonnitrile to yield the desired thioether followed by the identical reduction and Hugerschoff reaction to form the benzothiazole core. Once formed, **C-29**’s route deviates and the oxidation occurs through oxone, followed by a final reduction with palladium on carbon to yield the product. To synthesize **C-33**, 4-iodo-1*H*-pyrazole was first methylated, and again underwent similar transformations to **C-26**, a cross coupling with Pd_2_(dba)_3_ and Xantphos, a reduction with zero valent iron, and the Hugerschoff reaction (Scheme 1D). Next, the benzathiazole intermediate was Boc protected to enable oxidation of the thiol linkage to a sulfone and deprotected under acidic conditions to yield **C-33**. To synthesize **C-35,** the cross-coupling substrate was prepared by radical bromination of 5-nitro-1*H*-pyrazole followed by trimethylsilylethoxymethyl (SEM) protection and then entered into the cross-coupling with *tert*- butyl *N*-(4-sulfanylphenyl)carbamate to form the *tert*-butyl *N*-[4-[5-nitro-1-(2- trimethylsilylethoxymethyl)pyrazol-4-yl]sulfanylphenyl] carbamate. The carbamate was deprotected to form the aniline and entered into the Hugerschoff reaction. The free nitro group was then reduced with Raney Nickel with subsequent oxidation of the thiol with oxone to form the sulfone bridge. Finally, the SEM group was removed to afford **C-35** as shown in Scheme 1E. The most active molecule, **C-34** was made by first protecting 4-iodo-1*H*-pyrazole with SEM, followed by cross-coupling of the protected aryl iodide and 4-nitrobenzenethiol with Pd_2_(dba)_3_ and Xantphos to yield the aryl thioether. This molecule was reduced to the free amine, which provides a readily accessible substrate for the Hugerschoff transformation, yielding the 6-((1-((2- (trimethylsilyl)ethoxy)methyl)-1***H***-pyrazol-4-yl)thio)benzo[***d***]thiazol-2-amine intermediate. From here, the free exocyclic amine was Boc protected and the thioether linkage oxidized with sodium periodate affording the sulfone followed by global deprotection to give **C-34** (Scheme 1F). **C-36** starts with a coupling of 1-benzyl-4-iodo-pyrazole and 4-nitrobenzenethiol. The nitro group in the resulting product is reduced to give the free amine, cyclization was performed through the Hugerschoff reaction, and oxidation of the sulfur linker was performed to yield **C-36** (Scheme 1G). **C-25** undergoes a standard cross-coupling between the 4-nitrobenzenethiol and 4- chloropyridine. The resulting nitro-functionalized intermediate is reduced to the free amine, followed by the formation of the benzothiazole ring, and Boc protection of the exocyclic amine. Hereafter, the thioether linkage was oxidized to the sulfone with sodium periodate and finally, deprotected with trifluoroacetic acid to yield **C-25** (Scheme 1H).

**Scheme 1:**
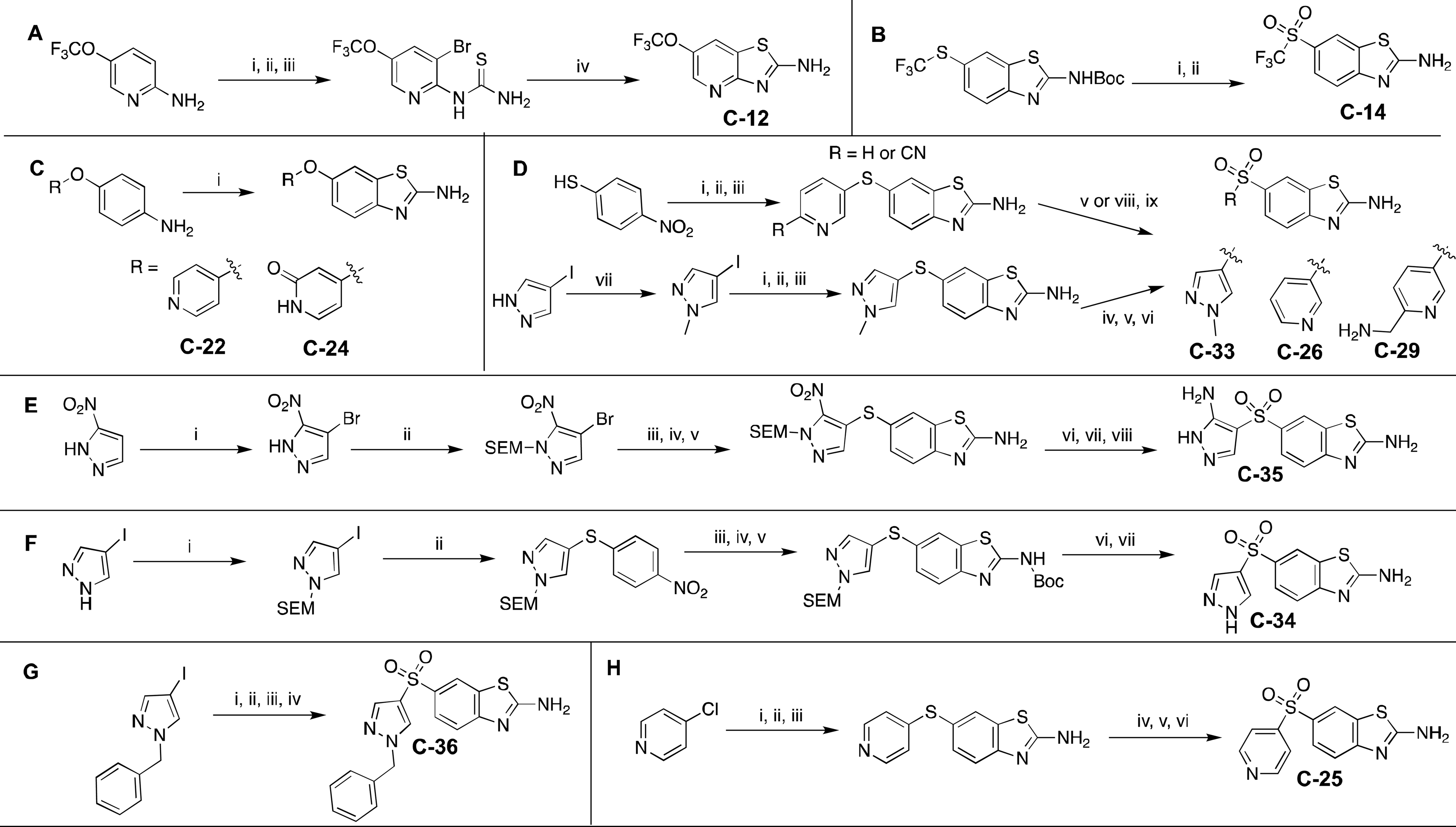
Synthesis of selected analogue set. **A**. i. NBS, CH_2_Cl_2_ ii. Benzoyl isothiocyanate, acetone iii. NaOH, H_2_O, CH_3_OH iv. NaH, DMF **B.** i. NaIO_4_, RuCl_3_, H_3_CCN, CCl_4_, H_2_O 20°C ii. CF_3_CO_2_H, CH_2_Cl_2_, 20°C **C.** i. KSCN, Br_2_, AcOH **D.** i. Pd_2_(dpa)_3_, Xantphos, Cs_2_CO_3_, dioxane ii. Fe, NH_4_Cl, CH_3_CH_2_OH, H_2_O, 80°C iii. KSCN, AcOH, Br_2_ iv. Boc_2_O, Et_3_N, CH_2_Cl_2_ v. NaIO_4_, RuCl_3_, H_3_CCN, CCl_4_, H_2_O 25°C vi. CF_3_CO_2_H, CH_2_Cl_2_ vii. CH_3_I, NaH, DMF. viii. Oxone, THF/H_2_O, 0-25°C ix. H_2_, Pd/C, NH_3_·H_2_O, CH_3_OH, 25°C **E.** i. NBS, DMF ii. NaH, SEMCl, DMF iii. *Tert*-butyl-*N*-(4-sulfanylphenyl)carbamate, Pd_2_(dpa)_3_, Xantphos, Cs_2_CO_3_, dioxane iv. Lutidine, TMSOTf, CH_2_Cl_2_ v. KSCN, Br_2_, AcOH vi. Raney Ni, H_2_, CH_3_OH vii. oxone, H_3_CCN, H_2_O viii. CF_3_CO_2_H, CH_2_Cl_2_ **F.** i. NaH, SEMCl, THF ii. 4-Nitrobenzenethiol, Pd_2_(dpa)_3_, Xantphos, Cs_2_CO_3_, dioxane iii. Fe, NH_4_Cl, CH_3_CH_2_OH, H_2_O, 80°C iv. KSCN, Br_2_, AcOH v. Boc_2_O, Et_3_N, CH_2_Cl_2_ vi. NaIO_4_, RuCl_3_, H_3_CCN, CCl_4_, H_2_O vii. CF_3_CO_2_H CH_2_Cl_2_ **G.** i. 4-Nitrobenzenethiol, CuI, Cs_2_CO_3_, DMF, 110°C ii. Fe, NH_4_Cl, CH_3_CH_2_OH, H_2_O, 80°C iii. KSCN, Br_2_, AcOH iv. Oxone, THF/H_2_O **H.** i. 4-Nitrobenzenethiol, K_2_CO_3_, DMF, 80°C ii. Fe, NH_4_Cl, CH_3_CH_2_OH, H_2_O, 80°C iii. KSCN, Br_2_, AcOH iv. Boc_2_O, N(CH_2_CH_3_)_3_, C_4_H_8_O v. NaIO_4_, RuCl_3_, H_3_CCN, CCl_4_, H_2_O, 25°C vi. CF_3_CO_2_H CH_2_Cl_2_, 20°C.

### Conclusions and Future Directions

We evaluated a series of putative HK inhibitors based on a 2-aminobenzothiazole scaffold. One of these compounds showed sub-micromolar activity *in vitro* against a model HK. We also observed strong inhibitory effects of the virulent-motility phenotype (swarming) with the most potent of these showing greater than a 90% reduction relative to a vehicle control without evidence of an effect on bacterial growth rates or cytotoxic effects at relevant concentrations. We identified several compounds in this series that inhibited another virulence phenotype, the production of a redox toxin (PYO), at early growth time points. Interestingly, the molecules that were the most potent in each assay were distinct from each other. This is likely explained by differences in the HK inhibition profile of these molecules, as well as potential non-HK inhibition events. We also anticipate that compound internalization and accumulation may affect their potency in cell-based assays.^75-76^ Promisingly, we were able to reduce cytotoxic effects from our original series of molecules without appreciably affecting the inhibition of virulence phenotypes. Future work will be aimed at further understanding of potential differences in the mechanism(s) of inhibition and intracellular accumulation. Overall, our data suggest that targeting the highly conserved CA domain within the HKs holds the promise of producing agents that interfere with the virulence properties of multi-drug resistant bacteria and could lead to clinically effective therapies.

## Materials and Methods

### Compound Purity

All tested molecules were >95% pure as judged by HPLC analysis. HPLC traces can be found in the SI.

### IC_50_ Activity Assay

HK853 (1 μM) in reaction buffer premixed with Triton X-100 (0.1% v/v) was incubated with each inhibitor at the following concentrations: 0 μM, 0.01 μM, 0.1 μM, 1 μM, 5 μM, 10 μM, 20 μM, 50 μM, 150 μM, 500 μM, 700 μM, 1000 μM, 1250 μM for a final volume of 25 μL for 30 min. B-ATPγS was added (50 μM) and incubated in the dark at RT for 1 h prior to quenching the reaction with 4X SDS-PAGE loading buffer (8.6 μL). Samples (15 μL) were loaded onto a 10% tris acrylamide gel and run at 180 V for 1 h in the dark on ice. Gel was removed from cassette, washed 3x with MilliQ water, and scanned in the BODIPY channel on a Typhoon gel scanner. Gel was stained was Coomassie (30 min), then washed 3x with MilliQ water and destained (40% MeOH, 10% AcOH) overnight and scanned in the Coomassie channel on a Typhoon gel scanner. Protein was prepared as previously described.^40, 46, 92^ Initially, inhibitors were screened at 3 concentrations, 10 μM, 100 μM, 500 μM, following the same procedure to determine if any activity was observed before moving to full dilution series for throughput purposes.

### Data analysis

Integrated density measurements of in-gel fluorescence and phosphorescence were performed in ImageJ.^93^ Data were prepared and analyzed in GraphPad Prism (version 9.0 for Windows, GraphPad Software, San Diego, California USA, www.graphpad.com). For all dose response curves, data were fit to a four-parameter logistic equation,

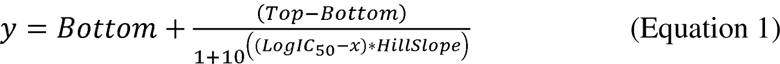

where *y* is the response, *Bottom* and *Top* are plateaus in the units of the y-axis, *x* is the log of the molar concentration of inhibitor, *HillSlope* is the slope of the curve, and IC_50_ is the concentration of compound required for 50% inhibition (a response half way between *Bottom* and *Top*).

### Aggregation Assay

Compounds were prepared in 25 mM stock concentrations in DMSO, and then further diluted with DMSO for a final concentration of 200 μM and incubated with 0.5 μM of HK853 in 20 mM HEPES buffer (25 μL). Samples were incubated at RT for 30 min, after which native loading buffer (8.6 μL) was added to each sample and then loaded onto a Native-PAGE gel and run on ice at 180 V for 1 hr. All Native-PAGE gels were silver stained with a Pierce Silver Stain Kit (ThermoFisher) according to manufacturer’s procedure. Gels were imaged on a white light illuminator with a camera. NH125 (Tocris Bioscience) was used as a positive aggregation control and DMSO was used as a negative control on each gel. Native-PAGE sample loading buffer recipe: 40 mM Tris, pH 7.5, 8% glycerol, and 0.08% bromophenol blue (w/v). Native-PAGE gels: 7.5% polyacrylamide Tris-glycine resolving gels. Native running Buffer recipe: 83 mM Tris, pH 9.4, and 33 mM glycine.

### Swarming Assay

Single colonies of *Pseudomonas aeruginosa* PA14-lux^94^ from blood agar plates were grown overnight in LB broth (Lennox – Sigma Aldrich) to an OD_600_ of 1.2. Culture media (5 µL) was pipetted in the center of freshly made modified FAB agar plates. Culture was allowed to dry in incubator (30°C) for 20 minutes upright before flipping. Incubation was carried out for 24 h before imaging plates on Typhoon FLA 9500 scanner (GE healthcare) using the DY-520XL filter setting. Images were analyzed using ImageJ (NIH) with the oval measurement tool and area was measured for furthest points of tendrils in quadruplicate. Measurements were averaged and taken as a percentage of a DMSO vehicle control. All experiments were carried out in biological duplicate due to limitations in material availability. Data were compared using a One-way ANOVA followed by Dunnett’s multiple comparisons test, which was performed using GraphPad Prism version 8.0.0 for Windows, GraphPad Software, San Diego, California USA, www.graphpad.com.

Modified Fastidious Anaerobe Broth (FAB) agar plates were made with the following recipe: Into 200 mL MilliQ H_2_O was added 1.8 g Na_2_HPO_4_ x 7H_2_O, 600 mg KH_2_PO_4_, 600 mg NaCl, 900 mg Bactoagar, and 200 mg Casamino acids. This was autoclaved at 121°C for 22 minutes. After cooling to 60°C, sterile filtered solutions of heat sensitive materials were added: 200 µL (198 g/L) MgCl_2_ x 7H_2_O, 200 µL (10.5 g/L) CaCl_2_, 2 mL (216 mg/ml) glucose, and 40 µL of a trace metal solution. The trace metal solution consists of CaSO_4_·2H_2_O (1 g/L), FeSO_4_·7H_2_O (1 g/L), CuSO_4_·6H_2_O (100 mg/L), MnSO_4_·H_2_O (100 mg/L), ZnSO_4_·7H_2_O (100mg/L), CuSO_4_·5H_2_O (42.5 mg/L), Na_2_MoO_4_·2H_2_O (50 mg/L), and boronic acid (25 mg/L). Inhibitors were added from 25 mM DMSO stocks (100 µM inhibitor concentration – 0.4% DMSO) before pouring 20 mL of agar into sterile petri dished (100 x 15 mm). Plates were dried in laminar flow hood for 45 min under UV.

### Pyocyanin Absorbance Assay

*Pseudomonas aeruginosa* PA14-lux^94^ was grown overnight in LB broth (20 g/L) in duplicate from colonies picked from blood agar plates and serially diluted, 1:100 in LB broth (15 g/L) with or without inhibitor (200 µM in 5 mL LB, 0.8% DMSO) and grown for 18 h at 37°C 220 rpm. Aliquots (200 µL x 4) were taken from each sample in the 96 well plate for OD_600_ measurement. Material (2 mL) was taken from each sample and centrifuged at 6,000 x *g* for 10 min at 4°C. Next, 150 µL x 3 from each sample was add to a well in Corning transparent 96-well plate (cat#CLS2585 from Millipore Sigma) and absorbance was measured at 690 nm in a Tecan Spark Plate Reader. For analysis, the average LB background was subtracted and technical replicates were averaged. Each technical replicate was adjusted to background subtracted OD600 (ABS 690/OD600) and then taken as a percentage of vehicle control before averaging biological replicates. Data were compared using a One-way ANOVA followed by Dunnett’s multiple comparisons test, which was performed using GraphPad Prism version 8.0.0 for Windows, GraphPad Software, San Diego, California USA, www.graphpad.com.

### Phenazine Quantification Assay

Post-incubation, media was carefully pipetted out of each well and transferred to 1.5 mL microfuge tubes. Culture aliquots were pelleted via centrifugation at 15,000 x *g* for 1 min. Supernatant was collected and subjected to filtration through a 0.22 mm PES centrifuge filter. Filtrate was transferred to 200 mL poly-spring inserts placed within 2 mL HPLC vials with pierceable septa caps. Vials were sealed prior to loading into the HPLC autosampler.

Pyocyanin (PYO), phenazine-1-carboxylic acid (PCA), and phenazine-1-carboxamide (PCN) were quantified using a Dionex UltiMate 3000 HPLC system operated by Chromeleon software (v.7.0, Thermo Fisher). This instrument is composed of compatible RS pump, autosampler, column oven, FLD detector, RS diode array, fraction collector and an Acclaim C18 3 mm 120A° 3.0 x 1500 mm column with accompanying guard column (Thermo Fisher). Analyte separation was achieved using a 30-min gradient method including a 5 min equilibration, and 0.1% trifluoroacetic acid (TFA) and 99.9% CH_3_CN mobile phases at a steady flow rate of 1 mL/min. 8 µL of each sample were used per run and injected into the system via the autosampler, with a brief washing of the injection needle with 10% CH_3_OH before and after each injection. Column oven temperature was maintained at 40°C. The RS diode array was configured to collect UV readings at the wavelengths of 370 nm with default frequency. Pure stocks of PYO, PCA, and PCN were used to generate standard curves.

Chromeleon software (v.7.0) was used to view and process raw data. Raw chromatograms were stacked and offsets removed to compare standard runs and perform an initial quality control of each run sequence. Quantitative processing methods were created for each target analyte. The integrated Cobra Wizard was used to gate and smooth peaks of interest on a single standard of intermediate concentration (1-10 mM) for a given analyte. To ensure gating accuracy, each sample was manually checked and the AUC recorded for later processing in Prism 9.0 (GraphPad). Cobra wizard was run with default settings.

### Bacterial Growth Assay

*Pseudomonas aeruginosa* PA14-lux^94^ was plated on LB agar from glycerol stocks and grown overnight at 37°C. Single colonies were selected and grown in LB media overnight (220 RPM, 37°C). Overnight cultures were diluted 1:100 into fresh LB media and incubated with selected inhibitors to a final concentration of 200 μM. The final concentrations of DMSO were 0.8% or below. Diluted cultures containing inhibitors were plated into a Corning transparent 96-well plate (cat#CLS2585 from Millipore Sigma) and shaken continuously (5 ms, orbital) for 20 hr in a Tecan Spark Plate Reader. OD values were measured at 600 nm at intervals of 20 min throughout the 20 hr run. All measurements were performed in biological triplicate and the specific growth rate (μ) was calculated for the average of each sample by the following equation:

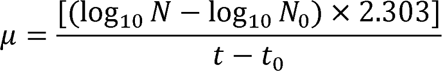

Where N is number of cells at any time (t) and the starting number of cells (N_0_) occurs at the beginning timepoint (t_0_). Total growth was assessed using Graphpad Prism v.10 by quantifying the area under curve (AUC) for each condition. All growth rates and AUC were then compared for statistical significance through a one-way ANOVA (α=0.05).

### Docking

Docking estimations of binding poses was carried out using Schrodinger Maestro software. HK853 crystal structure (3DGE) was imported from the RCSB repository and prepared with Schrödinger’s Protein Preparation Wizard suite adding missing hydrogens, assigning bond orders, and optimizing the H-bond network. Preprocessing was carried out using default methods, except removal of water molecules further than 3 Å away from heavy atoms. Localized clashes were removed through minimization and refinement before additional standard equilibration protocols. Glide was used to define the active site around the ATP ligand and subsequent ligand removal. Inhibitor structures were imported as .sdf files from ChemDraw and prepared for docking using LigPrep 5 with the OPLS_2005 force field. Ionization and tautomerization state of compounds at a pH range of 6–8, Epik v16207 was used, with a maximum number of 10 generated structures. The binding pose of the ligands was estimated using standard settings for extra precision Glide docking protocol.

### Cytotoxicity assay

Hep G2 cells (epithelial-like hepatocellular carcinoma; ATCC HB-8065, passages 16-19) or A549 cells (epithelial-like lung carcinoma; ATCC CCL-185) were grown in a T75 flask (Thermo 130190) incubated at 37°C, 5% CO_2_, and 21% O_2_ using 12 mL of 90% DMEM (v/v, Corning 10-101-CV), 10% HI-FBS (v/v, Gibco 10082147) 100 U/mL penicillin, 100 µg/mL streptomycin (Corning 30-002-CI), and 2 mM L-glutamine (Sigma G-1626) as media until they were 90% confluent. Media was replaced every other day until the intended confluency was reached. Cells were then detached using 5 mL of 0.25% trypsin, 0.53 mM EDTA solution (Gibco 25200056), quenched, centrifuged at 800 rcf for 5 min, and resuspended in fresh media. 50 µL of a 1.01 x 10^6^ cells/mL solution with 48 µL of media were then seeded in a 96- well plate (Sarstedt CLS3340) and incubated for 24 h. Cells were then challenged for 20 h in an incubator with 2 µL of a DMSO-solvated inhibitor solution. Following inhibitor challenge, 70 µL of XTT working solution (Invitrogen X12223) was added to each well and incubated for 4 h. For lysed cell controls, 15 µL of lysis solution (Promega G182A) was added 1 h prior to XTT working solution. DMSO controls, inhibitor-containing blanks, and growth controls (media only) were also included. After XTT working solution incubation, specific absorbance (450 nm) and nonspecific absorbance (660 nm) was measured using a BioTek Synergy HT plate reader. Blank specific absorbance and sample nonspecific absorbance values were subtracted from sample specific absorbances. Sample absorbances were then normalized to a 2% DMSO-containing cell control from the same 96-well plate. In GraphPad Prism, normalized values were plotted with respect to inhibitor concentration.

### Animal Experiments

Female BALB/c mice (Jackson Laboratories), aged 8 weeks, were anesthetized using isoflurane (3% at 3L min^−1^) and challenged intratracheally with 100 mL of 1% DMSO in phosphate buffered saline containing 100 µM, 200 µM, or 500 µM of **Rilu-2** or **Rilu-6**. DMSO alone was used as a vehicle control. Mice were monitored for signs of moribundity and sacrificed after 24 h via CO_2_ asphyxiation and cervical dislocation. Excised lungs were lavaged with 2 mL of sterile PBS, stained with 0.4% trypan blue, and total white blood cell count was determined on a Countess 3 (ThermoFisher) automated cell counter. Treatment groups were compared using a One-way ANOVA followed by Dunnett’s multiple comparisons test, which was performed using GraphPad Prism version 8.0.0 for Windows, GraphPad Software, San Diego, California USA, www.graphpad.com.

### Cloning, expression and purification

The codon optimized sequence for HK853 (235-492) was prepared and cloned into the pET-28a(+) plasmid at GenScript (https://www.genscript.com). Protein was expressed using *E.coli* BL21star (DE3) cells in ^15^N Minimal Media including ^15^NH_4_Cl as the sole nitrogen source (^15^N MM) and induced with 0.5 mM IPTG at 20°C overnight. Cells were harvested by centrifugation (5,000 x *g* for 20 min at 4°C) and resuspended in 50 mM Tris, pH 7.8, 500 mM NaCl, 10% glycerol, 10 mM Tris(2-carboxyethyl) phosphine hydrochloride (TCEP) containing Complete Ultra tablets, EDTA-free Protease inhibitor cocktail (Roche, USA). Cell lysis was accomplished by using a Fisherbrand^TM^ Model 505 Sonic Dismembrator (Fisher Scientific, USA). Cell resuspension, kept on ice, was lysed by sonication through four 45-sec cycles at 70% amplitude. Cell lysates were centrifuged at 20,000 x *g* for 30 min at 4°C and supernatant was diluted in TCEP-free lysis buffer to reduce TCEP concentration to 2 mM. The diluted solution was loaded onto a HisTrap crude column (Cytiva). The column was washed with buffer A (50 mM Tris, pH 7.8, 500 mM NaCl, 10% glycerol, 2 mM TCEP) and the bound proteins were eluted using a linear gradient of buffer B (50 mM Tris, pH 8, 500 mM NaCl, 10% glycerol, 2 mM TCEP, 500 mM imidazole). For NMR studies, HK853 N-terminal His-tag was cleaved by incubation with tobacco Etch virus protease (TEV) overnight at 4°C in a SnakeSkin Dialysis Tubing (ThermoScientific) to remove excess imidazole. His-tag and His-TEV were removed using a nickel column and HK853 was further purified on a size-exclusion (SEC) Superdex 75 column (Cytiva) in buffer C (50mM Tris, pH 8.0, 500 mM NaCl, 10% glycerol, 2 mM TCEP). Fractions containing the desired protein were pooled, concentrated using Amicon centrifugal filters (Millipore), flash frozen in liquid nitrogen and stored at ™80°C. The final purity was greater than 90% by SDS-PAGE.

### NMR sample preparation

All NMR samples were prepared in a final buffer containing 20 mM HEPES-d_18_ pH 8, 50 mM KCl, 10% D_2_O. DMSO-d_6_ content was kept constant at 3.6%. Samples containing compounds C-12, C-14, C-17, C-27, C-30, C-34, Rilu-1, Rilu-7 were prepared from a DMSO-d_6_ stock solution between 1.4 ™50 mM and diluted with buffer or protein solution to reach a concentration of 40-1000 µM. Samples containing adenosine diphosphate (ADP) were prepared from a stock solution in NMR buffer between 5-50 mM and diluted with buffer or protein solution to reach a concentration of 200-800 µM. Prior to preparing sample containing HK protein, the protein was buffer exchanged on a Zeba™ Spin Desalting Column, 7K MWCO (Thermo Scientific) and diluted in buffer to reach the final concentration: For ligand and protein detection 1D NMR, HK853 (232-489) was prepared at a final concentration of 15 µM and for 2D ^1^H-^15^N NMR experiments, ^15^N-labeled HK853 (235-492) was used at a final concentration of 100 µM.

### NMR experiments

NMR experiments were performed with the following hardware: 600 MHz Bruker Avance III NMR spectrometer equipped with a QCI HFCN helium cryoprobe, SampleJet, and ATMA autotune system. All samples were prepared in 3 mm Bruker SampleJet tubes to volumes of 180-200 µL with NMR buffer (20 mM HEPES-d_18_ pH 8, 50 mM KCl, 10% D_2_O). DMSO-d6 content was kept constant at 3.6%. The following NMR experiments were acquired for evaluation of compound binding: The 1D ^1^H used for differential line broadening (DLB) employed the standard Bruker 1D ^1^H sequence with excitation sculpting (zgesgp). Spectra were acquired with 16 scans. The T2-CPMG experiment employed is a modified version of the standard Bruker 1D ^1^H experiment with excitation sculpting (zgesgp) with the addition of a CPMG pulse train after the initial 90-deg excitation pulse. The total duration of each spin echo was fixed to 1 msec (τ=500 µsec) whereas the number of echoes in the pulse train was varied according to the total time (T), ranging from 0 to 800 ms. The number of scans for all spectra was 4. The sequence employed for saturation transfer difference (STD) is a modified version of standard zgesgp; with water suppression using excitation sculpting with gradients with saturation applied at 0 and 20 ppm for the ON and OFF-resonance experiments, respectively. 1D ^1^H Protein-Detected NMR was performed using the zgesgp pulse sequence.^95^ The 1D ^19^F used for differential line broadening (DLB) employed the standard ^19^F 1D NMR pulse sequence zgfhigqn.2 (1D sequence for F-19 observed with inverse gated H-1 decoupling). Spectra were acquired with 64 scans. Ligand and 1D protein detection experiment were performed on samples of 15 µM of HK853 (235-492) protein and 300 µM of compound.

2D ^1^H-^15^N NMR experiments were performed using the HSQC-TROSY sequence.^96^ HSQC-TROSY experiments were recorded at a protein concentration of 100 µM in absence or presence of compounds using ratios between 1:3.5 and 1:10. The weighted average changes in chemical shifts (CSP) were calculated according to the following equation^97^:

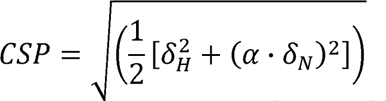

Where δ are the changes in chemical shift in ppm for ^1^H and ^15^N and the correction factor (α) was set at 0.15.

Competition experiments between ADP and compounds were performed using ^19^F 1D NMR with 128 scans. ADP was added at different ratio to samples containing fluorinated compound and HK853 (235-492). HK853 (235-492) protein and compound concentrations used were 40 and 80 µM respectively and ADP was used between 200 and 800 µM. Software: Topspin 3.5pl7 was used to control the spectrometer, for data acquisition and processing.

Figures were prepared with one or more of the following software: CCPNMR Analysis, Bruker Topspin, Microsoft Excel, and NMX’s in-house software packages. NMX’s custom software were employed in unsupervised and semi-supervised analysis of ^1^H/^19^F ligand-binding data and for 2D ^1^H-^15^N binding data.

### Inhibitor Synthesis

Abbreviations: Petroleum Ether: PE, ethyl Acetate: EA, methyl Acetate: MA, dichloromethane: DCM, trifluoroacetic acid: TFA, methanol: MeOH, ethanol: EtOH, formic acid: FA, triethylamine: TEA ***C-12***. To a solution of 5-(trifluoromethoxy)pyridin-2-amine (300 mg, 1.68 mmol) in DCM (4 mL) was added a solution of NBS (299 mg, 1.68 mmol) in DCM (2 mL). The mixture was stirred at 25°C for 12 hrs. On completion, the reaction mixture was concentrated and then purified by column chromatography (PE:EA = 10:1) to give 3-bromo-5- (trifluoromethoxy)pyridin-2-amine (250 mg, 57% yield) as a yellow solid. LCMS (MS+1)^+^: 257.0, 259.0. ^1^H NMR (400MHz, CDCl_3_) δ = 8.00 (d, *J* = 1.6 Hz, 1H), 7.61 (d, *J* = 1.6 Hz, 1H), 5.03 (br s, 2H).

3-Bromo-5-(trifluoromethoxy)pyridin-2-amine (250 mg, 972.73 μmol) was dissolved in acetone (5 mL) and benzoyl isothiocyanate (238 mg, 1.46 mmol) added at 0°C. The reaction mixture was stirred at 25°C for 12 hrs. On completion, the reaction was concentrated. The residue was triturated with PE (5 mL) to give N-((3-bromo-5-(trifluoromethoxy)pyridin-2- yl)carbamothioyl)benzamide (400 mg, 95% yield) as white solid. LCMS (MS+1)^+^: 419.8, 421.8. ^1^H NMR (400MHz, CDCl_3_) δ = 12.77 (br s, 1H), 9.35 (br s, 1H), 8.48 (s, 1H), 7.95 (s, 1H), 7.94 - 7.91 (m, 2H), 7.73 - 7.65 (m, 1H), 7.62 - 7.52 (m, 2H).

To a suspension of N-[[3-bromo-5-(trifluoromethoxy)-2- pyridyl]carbamothioyl]benzamide (400 mg, 951 μmol) in MeOH (0.6 mL) was added NaOH aqueous (2 M, 4.76 mL). The reaction mixture was stirred at 100°C for 1 hr. On completion, the reaction was adjusted pH to 9 with 1 N HCl aqueous and a white solid was precipitated. The mixture was filtered and the filter cake was dried in vacuo to give 1-(3-bromo-5-(trifluoromethoxy)pyridin-2-yl)thiourea (190 mg, 56% yield) as a white solid. LCMS (MS+1)^+^: 315.8, 417.8. ^1^H NMR (400MHz, CDCl_3_) δ = 10.47 (br s, 1H), 8.69 (br s, 1H), 8.18 (d, *J* = 2.0 Hz, 1H), 7.87 (d, *J* = 2.0 Hz, 1H), 7.07 (br s, 1H).

To a solution of [3-bromo-5-(trifluoromethoxy)-2-pyridyl]thiourea (190 mg, 601 μmol) in DMF (3 mL) was added NaH (72.1 mg, 1.80 mmol, 60% purity) at 0°C. The reaction mixture was stirred at 25°C for 30 min and then 80°C for 12 hrs. On completion, the reaction mixture was diluted with saturated NH_4_Cl aqueous solution. The mixture was extracted with EA (3 x 20 mL). The combined organic layer was washed with brine, dried over Na_2_SO_4_ and concentrated in vacuo. The residue was purified by Prep-HPLC (column: Phenomenex Gemini 150*25mm*10μm; mobile phase: [water (0.04%NH_3_ H_2_O + 10 mM NH_4_HCO_3_)-ACN]; B%: 23%-50%, 10 min) to give **C-12** (15.0 mg, 11% yield) as white solid. LCMS (MS+1)^+^: 236.0. ^1^H NMR (400MHz, DMSO-*d_6_*) δ = 8.25 (s, 1H), 8.14 (s, 1H). *Note ^13^C & ^19^F spectra not obtained due to lack of material.

***C-14***. A mixture of 4-(trifluoromethylsulfanyl)aniline (1.00 g, 5.18 mmol) and KSCN (2.01 g, 20.7 mmol) in HOAc (6 mL) was stirred at 20°C for 30 min, then a solution of Br_2_ (827 mg, 5.18 mmol) in HOAc (3 mL) was added drop-wise. The reaction mixture was stirred at 20°C for 20 hrs. On completion, the reaction was poured into ice-water (50 mL) and adjusted pH to 10 with 28% ammonium aqueous and a white solid was precipitated. The solid was collected and washed with water (10 mL) to give the 6-((trifluoromethyl)thio)benzo[d]thiazol-2-amine (1.00 g, 76% yield) as a yellow solid. ^1^H NMR (400MHz, DMSO-*d_6_*) δ = 7.78 (s, 1H), 7.33 (d, *J* = 8.4 Hz, 1H), 7.18 - 7.15 (m, 1H), 5.11 (s, 2H).

To a solution of 6-(trifluoromethylsulfanyl)-1,3-benzothiazol-2-amine (1.00 g, 4.00 mmol), TEA (1.21 g, 11.9 mmol) and DMAP (48.8 mg, 399 μmol) in THF (5 mL) was added Boc_2_O (1.31 g, 5.99 mmol). The reaction mixture was stirred at 20°C for 12 hrs. On completion, the reaction was diluted with water (50 mL), extracted with EA (3 x 20 mL). The combined organic layer was washed with 1 N HCl aqueous (20 mL), brine and dried in vacuo. The residue was purified by column chromatography (PE:EA = 20:1) to give the *tert*-butyl N-[6- (trifluoromethylsulfanyl) ™1,3-benzothiazol-2-yl]carbamate (0.7 g, 49% yield) as a white solid. LCMS (MS+1)^+^: 351.1. ^1^H NMR (400MHz, CDCl_3_) δ = 11.27 (s, 1H), 8.10 (d, *J* = 1.6 Hz, 1H), 7.94 (d, *J* = 8.4 Hz, 1H), 7.66 (dd, *J* = 1.6, 8.4 Hz, 1H), 1.61 (s, 9H).

To a solution of *tert*-butyl N-[6-(trifluoromethylsulfanyl) ™1,3-benzothiazol-2- yl]carbamate, 200 mg, 570 μmol) and ruthenium(III) chloride hydrate (6.43 mg, 28.54 μmol) in a mixed solvent of ACN (1 mL), CCl_4_ (1 mL) and H_2_O (2 mL) was added sodium periodate (488 mg, 2.28 mmol) in portions. The reaction mixture was stirred at 20°C for 16 hrs. On completion, the reaction was filtered and the filtrate was diluted with EA (50 mL) and water (50 mL). The mixture was extracted with EA (3 x 20 mL). The organic layer was washed with brine, dried over Na_2_SO_4_, filtered and concentrated in vacuo. The residue was purified by column chromatography (PE: EA = 20: 1) to give tert-butyl (6- ((trifluoromethyl)sulfonyl)benzo[d]thiazol-2-yl)carbamate (150 mg, 68% yield) as a yellow solid. LCMS (MS-56)^+^: 327.0. ^1^H NMR (400MHz, CDCl_3_) δ = 10.55 (br s, 1H), 8.50 (d, *J* = 0.8 Hz, 1H), 8.11 - 7.98 (m, 2H), 1.63 (s, 9H).

A solution of *tert*-butyl N-[6-(trifluoromethylsulfonyl)-1,3-benzothiazol-2-yl]carbamate (150 mg, 392 μmol) in DCM (2 mL) was added TFA (1 mL). The reaction mixture was stirred at 20°C for 4 hrs. On completion, the reaction was concentrated in vacuo. The residue was purified by Prep-HPLC (column: Phenomenex Synergi C18 150*25*10μm; mobile phase: [water (0.05% HCl)-ACN]; B%: 30%-50%, 9 min) to give **C-14** (54.0 mg, 43% yield, HCl salt) as a white solid. LCMS (MS+1)^+^: 283.0. ^1^H NMR (400MHz, DMSO-d_6_) δ = 8.65 (br s, 2H), 8.55 (d, *J* = 2.0 Hz, 1H), 7.87 (dd, *J* = 2.0, 8.8 Hz, 1H), 7.61 (d, *J* = 8.8 Hz, 1H). ^13^C NMR (101 MHz, DMSO-*d_6_*) δ 173.03, 158.31, 132.34, 129.19, 125.48, 120.10, 119.97, 118.05. ^19^F NMR (376 MHz, DMSO-*d_6_*) δ ™78.73.

***C-22***. 4-(4-Pyridyloxy)aniline (100 mg, 537 μmol) and KSCN (208 mg, 2.15 mmol) in HOAc (2 mL) was stirred at 20°C for 30 min, then Br_2_ (85.8 mg, 537 μmol) in HOAc (1 mL) was added. The reaction mixture was stirred at 20°C for 12 hrs. On completion, the reaction was adjusted pH to 8 with ammonium aqueous, extracted with EA (3 x 30 mL). The combined organic layer was washed with brine, dried over Na_2_SO_4_ and concentrated in vacuo. The residue was purified by Prep-HPLC (column: Phenomenex Gemini 150*25mm*10 μm; mobile phase: [water (0.04% NH_3_ H_2_O+10 mM NH_4_HCO_3_)-ACN]; B%: 27%-48%, 10 min) to give **C-22** (52.9 mg, 39% yield) as a white solid. LCMS (MS+1)^+^: 244.1. ^1^H NMR (400MHz, DMSO-*d_6_*) δ = 8.46 - 8.39 (m, 2H), 7.56 (d, *J* = 2.8 Hz, 1H), 7.50 (s, 2H), 7.38 (d, *J* = 8.4 Hz, 1H), 7.02 (dd, *J* = 2.4, 8.4 Hz, 1H), 6.91 - 6.86 (m, 2H). ^13^C NMR (101 MHz, DMSO-*d_6_*) δ 167.33, 165.34, 151.85, 151.11, 147.71, 132.75, 119.13, 118.95, 114.24, 112.04.

***C-24***. A mixture of 4-(4-aminophenoxy)-1H-pyridin-2-one (100 mg, 494 μmol) and KSCN (240 mg, 2.47 mmol) in HOAc (2 mL) was stirred at 25°C for 30 min, then Br_2_ (79.0 mg, 494 μmol) in HOAc (1 mL) was added. The reaction mixture was stirred at 25°C for 12 hrs. On completion, the reaction was diluted with ice-water (50 mL), adjusted pH to 10 with ammonium water and extracted with EA (3 x 20 mL). The combined organic layer was washed with brine, dried over Na_2_SO_4_ and concentrated in vacuo. The residue was purified by Prep-HPLC (column: Phenomenex Gemini 150*25mm*10 μm; mobile phase: [water (10 mM NH_4_HCO_3_)-ACN]; B%: 5%-35%, 10min) to give **C-24** (11.9 mg, 9% yield) as a white solid. LCMS (MS+1)^+^: 259.9. ^1^H NMR (400MHz, DMSO-*d_6_*) δ = 11.28 (br s, 1H), 7.61 - 7.47 (m, 3H), 7.39 - 7.33 (m, 2H), 6.99 (dd, *J* = 2.4, 8.8 Hz, 1H), 5.98 (dd, *J* = 2.4, 7.2 Hz, 1H), 5.32 (d, *J* = 2.0 Hz, 1H). ^13^C NMR (101 MHz, DMSO-*d_6_*) δ 168.94, 167.39, 164.23, 151.16, 147.27, 137.10, 132.55, 119.28, 118.75, 114.41, 100.53, 99.04.

***C-25***. A mixture of 4-nitrobenzenethiol (5.00 g, 32.2 mmol), 4-chloropyridine (5.32 g, 35.4 mmol, HCl salt) and K_2_CO_3_ (8.91 g, 64.4 mmol) in DMF (20 mL) was stirred at 80°C for 12 hrs. On completion, the reaction was poured into ice-water (400 mL) and a light yellow solid was precipitated. The solid was collected and dried in vacuo to give 4-((4-nitrophenyl)thio)pyridine (3) (6.00 g, 72% yield) as a yellow solid. ^1^H NMR (400MHz, CDCl_3_) δ = 8.51 (d, J = 6.0 Hz, 2H), 8.23 (d, J = 9.2 Hz, 2H), 7.57 (d, J = 9.2 Hz, 2H), 7.17 (d, J = 6.0 Hz, 2H). To a mixture of 4-(4-nitrophenyl)sulfanylpyridine (6.00 g, 25.83 mmol) and NH_4_Cl (967 mg, 18.0 mmol) in a mixed solvent of EtOH (120 mL) and H_2_O (20 mL) was added Fe (8.66 g, 155 mmol) in portions at 80°C. The reaction mixture was stirred at 80°C for 2 hrs. On completion, the reaction was filtered and the filtrate was concentrated in vacuo. The residue was diluted with water (100 mL), extracted with EA (3 x 50 ml). The combined organic layer was washed with brine, dried over Na_2_SO_4_, filtered and concentrated in vacuo. The residue was purified by column chromatography (PE: EA = 5: 1) to give 4-(pyridin-4-ylthio)aniline (3.00 g, 48% yield) as a yellow solid. ^1^H NMR (400MHz, DMSO-*d_6_*) δ = 8.28 (d, *J* = 5.2 Hz, 2H), 7.20 (d, *J* = 8.0 Hz, 2H), 6.90 (d, *J* = 5.2 Hz, 2H), 6.67 (d, *J* = 8.0 Hz, 2H), 5.67 (br s, 2H).

A mixture of 4-(4-pyridylsulfanyl)aniline (1.50 g, 7.42 mmol) and KSCN (2.88 g, 29.6 mmol) in HOAc (10 mL) was stirred at 20°C for 30 min. Then a solution of Br_2_ (1.19 g, 7.42 mmol) in HOAc (5 mL) was added drop-wise. The reaction mixture was stirred at 20°C for 12 hrs. On completion, the reaction was poured into 30% ammonium aqueous and a white solid was precipitated. The solid was collected and washed with water, dried in vacuo to give 4-(pyridin-4- ylthio)aniline (1.00 g, 47% yield) as a white solid. LCMS (MS+1)^+^: 259.9. ^1^H NMR (400 MHz, DMSO-*d_6_*) δ = 8.33 (d, *J* = 6.0 Hz, 2H), 7.95 (d, *J* = 1.6 Hz, 1H), 7.79 (s, 2H), 7.49 - 7.37 (m, 2H), 6.96 (d, *J* = 6.0 Hz, 2H).

To a solution of 6-(4-pyridylsulfanyl)-1,3-benzothiazol-2-amine (1.00 g, 3.86 mmol) and TEA (1.17 g, 11.5 mmol) in THF (15 mL) was added Boc_2_O (2.10 g, 9.64 mmol). The reaction mixture was stirred at 25°C for 12 hrs. On completion, the reaction was concentrated in vacuo. The residue was diluted with EA (50 mL) and acidified with 1% citric acid aqueous. The organic layer was washed with brine, dried over Na_2_SO_4_ and concentrated in vacuo. The residue was purified by column chromatography (PE: EA = 5: 1) to give tert-butyl N-tert-butoxycarbonyl-N- [6-(4-pyridylsulfanyl)-1,3-benzothiazol-2-yl]carbamate (1.00 g, 56% yield) as a white solid. LCMS (MS+1)^+^: 460.1. ^1^H NMR (400MHz, CDCl_3_) δ = 8.34 (d, *J* = 6.0 Hz, 2H), 8.01 (d, *J* = 1.6 Hz, 1H), 7.86 (d, *J* = 8.8 Hz, 1H), 7.57 (dd, *J* = 1.6, 8.4 Hz, 1H), 6.92 (d, *J* = 6.4 Hz, 2H), 1.62 (s, 18H).

To a solution of tert-butyl N-tert-butoxycarbonyl-N-[6-(4-pyridylsulfanyl)-1,3- benzothiazol-2-yl] carbamate (200 mg, 435 μmol) in a mixed solvent of ACN (1 mL), CCl_4_ (1 mL) and H_2_O (2 mL) was added NaIO_4_ (372 mg, 1.74 mmol) and RuCl_3_ (4.51 mg, 21.7 μmol) at 0°C. Then the mixture was stirred at 25°C for 3 hrs. On completion, the reaction was filtered, and the filtrate was concentrated in vacuo. The residue was purified by column chromatography (PE: EA = 10: 1) to give tert-butyl N-tert-butoxycarbonyl-N-[6-(4-pyridylsulfonyl)-1,3- benzothiazol-2-yl]carbamate (120 mg, 35% yield) as a white solid which was put into next directly. LCMS (MS-100)^+^: 392.1. To a solution of tert-butyl N-tert-butoxycarbonyl-N-[6-(4- pyridylsulfonyl)-1,3-benzothiazol-2-yl] carbamate (0.12 g, 244 μmol) in DCM (2 mL) was added TFA (1 mL). Then the mixture was stirred at 25°C for 30 min. On completion, the reaction was concentrated in vacuo. The residue was purified by Prep-HPLC (column: Waters Xbridge 150 mm*25 mm*5 uM; mobile phase: [water (10mM NH_4_HCO_3_)-ACN];B%: 13%-38%, 8 min) to give **C-25** (4.75 mg, 6.5% yield) as a white solid. LCMS (MS+1)^+^: 291.9. ^1^H NMR (400 MHz, DMSO-*d_6_*) δ = 8.85 (d, *J* = 6.0 Hz, 2H), 8.39 (d, *J* = 2.0 Hz, 1H), 8.10 (s, 2H), 7.86 (d, *J* = 6.0 Hz, 2H), 7.79 (dd, *J* = 2.0, 8.8 Hz, 1H), 7.46 (d, *J* = 8.8 Hz, 1H). ^13^C NMR (101 MHz, DMSO-*d_6_*) δ 171.46, 158.13, 151.92, 150.47, 132.64, 130.43, 126.30, 122.30, 120.60, 118.21.

***C-26***. A mixture of 4-nitrobenzenethiol (2.00 g, 13.3 mmol), 3-bromopyridine (3.10 g, 19.3 mmol), Xantphos (1.50 g, 2.50 mmol), Cs_2_CO_3_ (12.6 g, 38.7 mmol) and Pd_2_(dba)_3_ (2.40 g, 2.60 mmol) in dioxane (20 mL) was stirred at 90°C for 3 hrs. On completion, the reaction mixture was concentrated in vacuo. The residue was purified by silica gel chromatography (PE:MA = 5:1) to give 3-(4-nitrophenyl)sulfanylpyridine (1.70 g, 48% yield) as yellow solid. LCMS (M+1)^+^: 233.0. ^1^H NMR (400 MHz, DMSO-*d*_6_) δ = 8.74 - 8.70 (m, 2H), 8.16 - 8.14 (m, 2H), 8.04 (d, *J* = 8.0 Hz, 1H), 7.58 - 7.56 (m, 1H), 7.36 - 7.33 (m, 2H).

To a mixture of 3-(4-nitrophenyl)sulfanylpyridine (1.60 g, 6.20 mmol) and NH_4_Cl (217 mg, 4.10 mmol,) in a mixed solvent of EtOH (18 mL) and H_2_O (3 mL) was added Fe (1.90 g, 35.5 mmol) in portions at 80°C. The reaction mixture was stirred at 80°C for 4 hrs. On completion, the reaction mixture was concentrated in vacuo. The residue was purified by silica gel chromatography (PE:MA = 3:1) to give 4-(3-pyridylsulfanyl)aniline (1.00 g, 76% yield) as yellow solid. ^1^H NMR (400 MHz, DMSO-*d*_6_) δ= 8.31 - 8.29 (m, 1H), 8.26 (d, *J* =2.0 Hz, 1H), 7.38 - 7.36 (m, 1H), 7.28 - 7.26 (m, 1H), 7.19 (d, *J* = 8.4 Hz, 2H), 6.62 (d, *J* = 8.4 Hz, 2H), 5.67 (s, 2H).

A mixture of 4-(3-pyridylsulfanyl)aniline (800 mg, 3.60 mmol) and KSCN (1.40 g, 14.2 mmol) in AcOH (5 mL) was stirred at 20°C for 30 min. Then a solution of Br_2_ (568 mg, 3.00 mmol) in AcOH (2.5 mL) was added drop-wise. The reaction mixture was stirred at 20°C for 12 hrs. On completion, the reaction was poured into ice water, and basified with NaHCO_3_ pH=7, and a white solid was precipitated. The solid was collected by filtration and washed with water. The filter cake was dried in vacuo to give 6-(3-pyridylsulfanyl)-1,3-benzothiazol-2-amine (0.600 g, 65% yield) as a light yellow solid. LCMS (M+1)^+^: 260.1, ^1^H NMR (400 MHz, DMSO-*d_6_*) δ = 8.40 (m, 2H), 7.88 (s, 1H), 7.69 (s, 2H), 7.54 (d, *J* = 6.0 Hz, 1H), 7.36 - 7.32 (m, 3H).

To a solution of 6-(3-pyridylsulfanyl)-1,3-benzothiazol-2-amine (200 mg, 771 μmol) in H_2_O (2 mL), ACN (1 mL) and CCl_4_ (1 mL) was added NaIO_4_ (659 mg, 3.10 mmol) and RuCl_3_ (8.00 mg, 38.6 μmol) at 0°C. The reaction mixture was stirred at 25°C for 3 hrs. On completion, the reaction was concentrated in vacuo. The residue was purified by Prep-TLC (PE:EA **=** 1:1) to give **C-26** (23.4 mg, 10% yield) as a white solid. LCMS (M+1)^+^: 292.1. ^1^H NMR (400 MHz, DMSO-*d*_6_) δ = 9.10 (d, *J* = 2.0 Hz, 1H), 8.82 (dd, *J* = 4.8, 1.2 Hz, 1H), 8.40 (d, *J* = 1.6 Hz, 1H), 8.33 - 8.30 (m, 1H), 8.09 (s, 2 H), 7.80 (dd, *J* = 8.4, 2.0 Hz, 1H), 7.65 (dd, *J* = 8.0, 4.8 Hz, 1H), 7.43 (d, *J* = 8.4 Hz, 1H). ^13^C NMR (101 MHz, DMSO-*d_6_*) δ 171.26, 157.80, 154.20, 147.99, 139.15, 135.57, 132.57, 131.72, 125.90, 125.09, 121.92, 118.11.

***C-29***. A mixture of 4-nitrobenzenethiol (3.00 g, 15.5 mmol), 5-bromopyridine-2-carbonitrile (4.30 g, 23.2 mmol), Xantphos (894 mg, 1.6 mmol), Cs_2_CO_3_ (15.1 g, 46.4 mmol) and Pd_2_(dba)_3_ (1.30 g, 1.40 mmol) in dioxane (100 mL), then the mixture was stirred at 90°C for 3 hrs. On completion, the reaction mixture was concentrated in vacuo. The residue was purified by silica gel chromatography (PE:MA = 6:1) to give 5-(4-nitrophenyl)sulfanylpyridine-2-carbonitrile (2.80 g, 70% yield) as yellow solid. ^1^H NMR (400 MHz, DMSO-*d*_6_) δ = 8.78 (s, 1H), 8.23 - 8.21 (d, *J* = 8.8 Hz, 2H), 8.09 (s, 2H), 7.63 - 7.61 (d, *J* = 8.8 Hz, 2H).

To a mixture of 5-(4-nitrophenyl)sulfanyl pyridine-2-carbonitrile (2.80 g, 10.9 mmol) and NH_4_Cl (407 mg, 7.60 mmol) in EtOH (27 mL) and H_2_O (4.5 mL) was added Fe (3.70 g, 65.3 mmol) in portions at 80°C. The mixture was stirred at 80°C for 2 hrs. On completion, the reaction mixture was filtered and concentrated in vacuo. The residue was purified by silica gel chromatography (PE:MA = 4:1) to give 5-(4-aminophenyl)sulfanylpyridine-2-carbonitrile (1.80 g, 65% yield) as yellow solid. ^1^H NMR (400 MHz, DMSO-*d*_6_) δ = 8.36 (s, 1H), 7.85 - 7.83 (d, *J* = 8.4 Hz, 1H), 7.42 (d, *J* = 2.0 Hz, 1H), 7.24 - 7.22 (d, *J* = 8.4 Hz, 2H), 6.69 - 6.67 (d, *J* = 8.8 Hz, 2H), 5.73 (s, 2H).

To a mixture of 5-(4-aminophenyl)sulfanyl pyridine-2-carbonitrile (1.60 g, 7.0 mmol) in HOAc (40 mL) was added KSCN (2.70 g, 28.2 mmol). The mixture was stirred at 25°C for 0.5 hour, then a solution of Br_2_ (1.1 g, 7.0 mmol) in HOAc (2 mL) was added. The reaction mixture stirred at 25°C for 12 hrs. On completion, the reaction mixture was poured into ice water and basified with NaHCO_3_ solution till pH =7, and a white solid was precipitated. The solid was collected and washed with water, dried in vacuo to give 5-[(2-amino-1,3-benzothiazol-6- yl)sulfanyl]pyridine-2-carbonitrile (1.60 g, 80% yield) as a light yellow solid. LCMS (M+1)^+^: 285.0. To a solution of 5-[(2-amino-1,3-benzothiazol-6-yl)sulfanyl]pyridine-2-carbonitrile (0.900 g, 3.20 mmol) in H_2_O (20 mL) and THF (60 mL) add Oxone (3.90 g, 6.30 mmol) at 0°C. The mixture was stirred at 25°C for 4 hrs. On completion, the reaction mixture was quenched by addition H_2_O (150 mL) at 0°C, and then filtered and the filter cake was dried in vacuo to give 5- [(2-Amino-1,3-benzothiazol-6-yl)sulfonyl]pyridine-2-carbonitrile (0.800 g, crude) as a light yellow solid. ^1^H NMR (400 MHz, DMSO-*d*_6_) δ = 9.26 (s, 1H), 8.59 - 8.56 (d, *J* = 10.4 Hz, 1H), 8.44 (s, 1H), 8.27 - 8.24 (d, *J* = 8.4 Hz, 1H), 8.13 (s, 2H), 7.85 - 7.83 (d, *J* = 8.4 Hz, 1H), 7.47 ™7.45 (d, *J* = 8.4 Hz, 1H).

To a solution of 5-[(2-amino-1,3-benzothiazol-6-yl)sulfonyl]pyridine-2-carbonitrile (200 mg, 632 μmol) in MeOH (10 mL) was added Pd/C (2.00 g, 1.90 mmol) and NH_3_.H_2_O (3.4 g, 26.0 mmol). The reaction mixture was stirred at 25°C for 12 hr under H_2_ (15 psi). On completion, the reaction was filtered and concentrated in vacuo. The residue was purified by Prep-HPLC (column: Phenomenex Luna C18 200*40 mm*10 μm;mobile phase: [water (0.2% FA)-ACN]; B%: 1%-30%, 10 min) to give **C-29** (20.0 mg, 10% yield) as light yellow solid. LCMS (M+1)^+^: 321.0. ^1^H NMR (400 MHz, DMSO-*d*_6_) δ = 9.02 (s, 1H), 8.39 (s, 1H), 8.30 (d, *J* = 7.6 Hz, 2H), 8.07 (s, 2H), 7.80 (d, *J* = 8.0 Hz, 1H), 7.69 (d, *J* =8.4 Hz, 1H), 7.45 (d, *J*=8.4 Hz, 1H), 3.97 (m, 2H). ^13^C NMR (101 MHz, DMSO-*d_6_*) δ 185.18, 171.22, 157.75, 147.33, 136.19, 132.54, 132.00, 125.82, 122.94, 121.80, 118.13, 55.77, 45.38.

***C-33***. To a solution of 4-iodo-1*H*-pyrazole (5.00 g, 25.7 mmol) in DMF (20 mL) was added NaH (2.58 g, 64.4 mmol, 60% purity) at 0°C. The reaction mixture was stirred at 0°C for 30 min. Then MeI (7.32 g, 51.5 mmol) was added and the mixture was stirred at 25°C for 12 hrs. On completion, the reaction was poured into saturated NH_4_Cl aqueous at 0°C, and extracted with EA (3 x 20 mL). The combined organic layer was washed with brine, dried over Na_2_SO_4_, filtered and concentrated in vacuo. The residue was purified by column chromatography (PE:EA= 10:1) to give 4-iodo-1-methyl-1H-pyrazole (4.5 g, 84% yield) as white solid. ^1^H NMR (400 MHz, CDCl_3_) δ = 7.49 (s, 1H), 7.40 (s, 1H), 3.92 (s, 3H).

A mixture of 4-nitrobenzenethiol (2.61 g, 16.8 mmol), 4-iodo-1-methyl-pyrazole (3.50 g, 16.8 mmol), Pd_2_(dba)_3_ (770 mg, 841 μmol), Xantphos (973 mg, 1.68 mmol) and Cs_2_CO_3_ (10.9 g, 33.6 mmol) in dioxane (100 mL) was stirred at 90°C under nitrogen atmosphere for 12 hrs. On completion, the reaction was filtered and the filtrate was concentrated in vacuo. The residue was purified by column chromatography (PE: EA = 10: 1) to give 1-methyl-4-((4-nitrophenyl)thio)- 1H-pyrazole (2.50 g, 63% yield) as yellow solid. ^1^H NMR (400 MHz, CDCl_3_) δ = 8.05 (d, *J* = 8.8 Hz, 2H), 7.62 (s, 1H), 7.60 (s, 1H), 7.14 (d, *J* = 8.8 Hz, 2H), 4.00 (s, 3H).

A mixture of 1-methyl-4-(4-nitrophenyl)sulfanyl-pyrazole (3.00 g, 12.7 mmol), Fe (7.12 g, 127 mmol) and NH_4_Cl (477 mg, 8.93 mmol) in EtOH (40 mL) and H_2_O (4 mL) was stirred at 80°C for 1 hr. On completion, the reaction was filtered and the filtrate was concentrated in vacuo to give 4-((1-Methyl-1H-pyrazol-4-yl)thio)aniline (1.20 g, 90% yield) as brown oil. LCMS (MS+1)^+^: 206.1. A solution of 4-(1-methylpyrazol-4-yl)sulfanylaniline (0.9 g, 4.38 mmol) and KSCN (1.70 g, 17.5 mmol) in HOAc (10 mL) was stirred at 25°C for 30 min, then Br_2_ (700 mg, 4.38 mmol) in HOAc (5 mL) was added. The reaction mixture was stirred at 25°C for 12 hrs. On completion, the reaction was adjusted pH to 10 with 10% NH_3_•H_2_O aqueous at 0°C. The mixture was extracted with EA (3 x 50 mL). The combined organic layer was dried over Na_2_SO_4_, filtered and concentrated in vacuo. The residue was purified by column chromatography (PE:EA = 5:1) to give 6-((1-methyl-1H-pyrazol-4-yl)thio)benzo[d]thiazol-2-amine (700 mg, 49% yield) as yellow solid. LCMS (MS+1)^+^: 263.0. To a solution of 6-(1-methylpyrazol-4- yl)sulfanyl-1,3-benzothiazol-2-amine (700 mg, 2.67 mmol) and TEA (809 mg, 8.00 mmol) in DCM (20 mL) was added Boc_2_O (1.16 g, 5.34 mmol). Then the reaction mixture was stirred at 25°C for 4 hrs. On completion, the reaction was diluted with water (100 mL), and the aqueous phase was extracted with EA (3 x 30 mL). The combined organic layers were washed with 1% citric acid aqueous and brine, dried over Na_2_SO_4_, filtered and concentrated in vacuo. The residue was purified by column chromatography (PE: EA = 5: 1) to give *tert*-butyl (6-((1-methyl-1H- pyrazol-4-yl)thio)benzo[d]thiazol-2-yl)carbamate (600 mg, 58% yield) as yellow solid. ^1^H NMR (400 MHz, CDCl_3_) δ = 7.72 (d, *J* = 8.4 Hz, 1H), 7.62 (s, 1H), 7.56 (s, 1H), 7.51 (d, *J* = 2.0 Hz, 1H), 7.22 (dd, *J* = 2.0, 8.4 Hz, 1H), 3.97 (s, 3H), 1.57 (s, 9H).

To a solution of *tert*-butyl N-[6-(1-methylpyrazol-4-yl)sulfanyl-1,3-benzothiazol-2- yl]carbamate (200 mg, 551 μmol) in a mixed solvent of ACN (1 mL), CCl_4_ (1 mL) and H_2_O (2 mL) was added NaIO_4_ (472 mg, 2.21 mmol) and RuCl_3_ (5.72 mg, 27.5 μmol) at 0°C. Then the mixture was stirred at 25°C for 12 hrs. On completion, the reaction was filtered and the filtrate was extracted with EA (3 x 10 mL). The combined organic layer was washed with brine and dried over Na_2_SO_4_, filtered and concentrated in vacuo. The residue was purified by column chromatography (PE: EA = 5: 1) to give *tert*-butyl (6-((1-methyl-1H-pyrazol-4- yl)sulfonyl)benzo[d]thiazol-2-yl)carbamate (100 mg, 46% yield) as colorless gum. LCMS (MS+1)^+^: 395.1. A mixture of *tert*-butyl N-[6-(1-methylpyrazol-4-yl)sulfonyl-1,3-benzothiazol-2-yl]carbamate (100 mg, 253 μmol) and TFA (0.5 mL) in DCM (1 mL) was stirred at 20°C for 30 min. On completion the reaction was concentrated in vacuo. The residue was purified by Prep-HPLC (column: Waters Xbridge 150*25 5 μm; mobile phase: [water (10 mM NH_4_HCO_3_)- ACN]; B%: 10%-37%, 9 min) to give **C-33** (26.5 mg, 36% yield) as white solid. LCMS (MS+1)^+^: 295.0. ^1^H NMR (400 MHz, DMSO-*d6*) δ = 8.38 (s, 1H), 8.27 (s, 1H), 7.97 (s, 2H), 7.87 (s, 1H), 7.76 - 7.67 (m, 1H), 7.42 (d, *J* = 8.4 Hz, 1H), 3.85 (s, 3H). ^13^C NMR (101 MHz, DMSO-*d_6_*) δ 170.69, 157.02, 138.64, 134.71, 133.59, 132.12, 124.83, 124.64, 120.69, 117.91.

***C-34***. To a solution of 4-iodo-1H-pyrazole (5.00 g, 25.7 mmol) in THF (40 mL) was added NaH (1.55 g, 38.6 mmol, 60% purity) in portions. The reaction mixture was stirred at 25°C for 30 min. Then 2-(chloromethoxy)ethyl-trimethyl-silane (4.73 g, 28.3 mmol) was added dropwise. The reaction mixture was stirred at 25°C for 12 hrs. On completion, the reaction mixture was poured into saturated aqueous NH_4_Cl. The mixture was extracted with EA (3 x 50 mL) and washed with water, brine. The organic layer was dried over anhydrous sodium sulfate, filtered and concentrated in vacuo. The residue was purified by column chromatography (PE: EA = 100: 1) to give 4-iodo-1-((2-(trimethylsilyl)ethoxy)methyl)-1H-pyrazole (6.00 g, 72% yield) as light yellow oil. LCMS (M+1)^+^: 325.0. ^1^H NMR (400 MHz, CDCl_3_) δ = 7.62 (s, 1H), 7.54 (s, 1H), 5.41 (s, 2H), 3.55 (t, *J* = 8.0, 2H), 0.90 (t, *J* = 8.0, 2H), ™0.02 (s, 9H). A mixture of 4-nitrobenzenethiol (2.87 g, 18.5 mmol), 2-[(4-iodopyrazol-1-yl)methoxy]ethyl-trimethyl-silane (6.00 g, 18.5 mmol), Pd_2_(dba)_3_ (847 mg, 925 umol), Xantphos (1.07 g, 1.85 mmol) and Cs_2_CO_3_ (12.0 g, 37.0 mmol) in dioxane (100 mL) was stirred at 90°C for 12 hrs under N_2_ atmosphere. On completion, the reaction was filtered and the filtrate was concentrated in vacuo. The residue was purified by column chromatography (PE:EA = 10:1) to give 4-((4-nitrophenyl)thio)-1-((2- (trimethylsilyl)ethoxy)methyl)-1H-pyrazole (5.00 g, 54% yield) as brown oil. LCMS (M+1)^+^: 352.1. To a mixture of trimethyl-[2-[[4-(4-nitrophenyl)sulfanylpyrazol-1- yl]methoxy]ethyl]silane (5.00 g, 14.2 mmol) and NH_4_Cl (532 mg, 9.96 mmol) in a mixture of EtOH (100 mL) and H_2_O (10 mL) was added Fe (7.94 g, 142 mmol) at 80°C. The resulting mixture was stirred at 80°C for 12 hrs. On completion, the reaction mixture was filtered and the filtrate was concentrated in vacuo. The residue was purified by column chromatography (PE:EA = 10:1) to give 4-((1-((2-(trimethylsilyl)ethoxy)methyl)-1H-pyrazol-4-yl)thio)aniline (3.00 g, 55% yield) as a brown oil. LCMS (M+1)^+^: 322.0. ^1^H NMR (400 MHz, CDCl_3_) δ = 7.63 (s, 1H), 7.54 (s, 1H), 7.11 (d, *J* = 8.0 Hz, 2H), 6.59 (d, *J* = 8.0 Hz, 2H), 5.39 (s, 2H), 3.7 (br s, 2H), 3.56 (t, *J* = 8.0, 2H), 0.90 (t, *J* = 8.0, 2H), ™0.02 (s, 9H).

To a solution of 4-[1-(2-trimethylsilylethoxymethyl)pyrazol-4-yl]sulfanylaniline (3.00 g, 9.33 mmol) in HOAc (2 mL) was added KSCN (3.63 g, 37.3 mmol). The reaction mixture was stirred at 25°C for 30 min. Then a solution of Br_2_ (1.49 g, 9.33 mmol) in HOAc (1 mL) was added and the mixture was stirred at 25°C for 12 hrs. On completion, the reaction was concentrated in vacuo. The residue was purified by reverse phase to give 6-((1-((2- (trimethylsilyl)ethoxy)methyl)-1H-pyrazol-4-yl)thio)benzo[d]thiazol-2-amine (2.20 g, 42% yield) as brown oil. LCMS (M+1)^+^: 379.1. ^1^H NMR (400 MHz, CDCl_3_) δ = 7.73 (s, 1H), 7.62 (s, 1H), 7.42 - 7.37 (m, 2H), 7.21 - 7.15 (m, 1H), 5.44 (s, 2H), 3.60 (m, 2H), 0.96 - 0.91 (m, 2H), ™0.01 (s, 9H). To a solution of 6-[1-(2-trimethylsilylethoxymethyl)pyrazol-4-yl]sulfanyl-1,3- benzothiazol-2-amine (2.20 g, 5.81 mmol) and TEA (2.35 g, 23.2 mmol) in THF (100 mL) was added Boc_2_O (3.17 g, 14.5 mmol). Then the reaction mixture was stirred at 25°C for 12 hrs. On completion, the reaction was concentrated in vacuo. The residue was diluted with EA (100 mL), washed with 1% critic acid, brine and the organic layer was concentrated in vacuo. The residue was purified by column chromatography (PE: EA = 10: 1) to give *tert*-butyl (6-((1-((2- (trimethylsilyl)ethoxy)methyl)-1H-pyrazol-4-yl)thio)benzo[d]thiazol-2- yl)carbamate (1.80 g, 62% yield) as a brown oil. LCMS (M+1)^+^: 479.1. ^1^H NMR (400 MHz, CDCl3) δ = 7.76 (s, 1H), 7.72 (d, *J* = 8.4 Hz, 1H), 7.65 (s, 1H), 7.55 (d, *J* = 1.6 Hz, 1H), 7.23 (dd, *J* = 1.6, 8.4 Hz, 1H), 5.46 (s, 2H), 3.62 (t, *J* = 8.0 Hz, 2H), 1.57 (s, 9H), 0.93 (t, *J* = 8.0 Hz, 2H), 0.00 (s, 9H).

To a solution of *tert*-butyl N-[6-[1-(2-trimethylsilylethoxymethyl)pyrazol-4-yl]sulfanyl-1,3- benzothiazol-2-yl]carbamate (200 mg, 417 μmol) in a mixture of ACN (1 mL), CCl_4_ (1 mL) and H_2_O (2 mL) was added NaIO_4_ (357 mg, 1.67 mmol) and RuCl_3_ (4.33 mg, 20.8 μmol) at 0°C. Then the reaction mixture was stirred at 25°C for 12 hrs. On completion, the reaction mixture was filtered and the organic layer was separated. The aqueous phase was extracted with EA (3 x 10 mL). The combined organic layers were washed with brine, dried over Na_2_SO_4_, filtered and concentrated in vacuo The residue was purified by column chromatography (PE: EA = 5: 1) to give *tert*-butyl (6-((1-((2-(trimethylsilyl)ethoxy)methyl)-1H-pyrazol-4- yl)sulfonyl)benzo[d] thiazol-2-yl)carbamate (120 mg, 56% yield) as a colorless gum. LCMS (M+1)^+^: 511.2. To a solution of *tert*-butyl N-[6-[1-(2-trimethylsilylethoxymethyl)pyrazol-4- yl]sulfonyl-1,3- benzothiazol-2-yl]carbamate (100 mg, 195 μmol) in DCM (1 mL) was added TFA (1 mL). Then the mixture was stirred at 25°C for 1 hr. On completion, **t**he reaction mixture was concentrated in vacuo. The residue was purified by Prep-HPLC (column: Phenomenex Gemini 150*25 mm*10 μm; mobile phase: [water (0.04% NH_3_ H_2_O + 10 mM NH_4_HCO_3_)- ACN]; B%: 10%-37%, 10 min) to give **C-34** (17.8 mg, 32% yield) as a white solid. LCMS (M+1)^+^: 281.0. ^1^H NMR (400 MHz, DMSO-*d_6_*) δ = 13.66 (br s, 1H), 8.41 (br s, 1H), 8.25 (d, *J* = 2.0 Hz, 1H), 7.94 (s, 2H), 7.89 (br s, 1H), 7.70 (dd, *J* = 2.0, 8.4 Hz, 1H), 7.39 (d, *J* = 8.4 Hz, 1H). ^13^C NMR (101 MHz, DMSO-*d_6_*) δ 170.64, 156.95, 134.91, 132.07, 124.83, 124.56, 120.67, 117.88.

***C-35***. A mixture of 5-nitro-1*H*-pyrazole (5.00 g, 44.2 mmol) and NBS (8.66 g, 48.6 mmol) in DMF (90 mL) was stirred at 90°C for 12 hrs under N_2_ atmosphere. On completion, the reaction was concentrated in vacuo. The residue was diluted with water (150 mL) and the mixture was filtered. The filter cake was washed with water (50 mL) and PE (20 mL) to give 4-bromo-5- nitro-1H-pyrazole (6.00 g, 70% yield) as a white solid. ^1^H NMR (400 MHz, DMSO-*d*_6_) δ = 14.36 (br s, 1H), 8.34 (s, 1H). To a solution of 4-bromo-5-nitro-1*H*-pyrazole (5.00 g, 26 mmol) in DMF (15 mL) was added NaH (1.56 g, 39.1 mmol, 60% purity) at 0°C. The reaction mixture was stirred at 0°C for 30 min. Then 2-(chloromethoxy)ethyl-trimethyl-silane (4.78 g, 28.6 mmol) was added, and the mixture was stirred at 10°C for 12 hrs. On completion, the reaction was diluted with water (50 mL), extracted with EA (3 x 20 ml). The combined organic layer was washed with brine, dried over Na_2_SO_4_ and concentrated in vacuo. The residue was purified by column chromatography (PE: EA = 20: 1) to give 4-bromo-5-nitro-1-((2- (trimethylsilyl)ethoxy)methyl)-1H-pyrazole in one pure isomer (2.60 g, 31% yield, SEM- position on pyrazol was not confirmed by 2D NMR). ^1^H NMR (400 MHz, CDCl_3_) δ = 7.63 (s, 1H), 5.86 (s, 2H), 3.63 - 3.51 (m, 2H), 0.92 - 0.88 (m, 2H), ™0.02 (s, 9H).

To a solution of 4-aminobenzenethiol (5.00 g, 39.9 mmol) and TEA (8.08 g, 79.8 mmol, 11.1 mL) in MeOH (50 mL) was added Boc_2_O (9.59 g, 43.9 mmol) under 0 - 5°C. The reaction mixture was stirred at 25°C for 12 hours. On completion, the reaction mixture was diluted with water (20 mL) and filtered. The filtered cake was washed with MeOH to give the *tert*-butyl N-(4- sulfanylphenyl)carbamate (4.02 g, 44% yield) as white solid. ^1^H NMR (400 MHz, DMSO-*d*_6_) δ = 9.51 (s, 1H), 7.51 - 7.41 (m, 2H), 7.40 - 7.30 (m, 2H), 1.50 (s, 9H).

A mixture of 2-[(4-bromo-5-nitro-pyrazol-1-yl)methoxy]ethyl-trimethyl-silane (2.40 g, 7.45 mmol), *tert*-butyl N-(4-sulfanylphenyl)carbamate (1.68 g, 7.45 mmol), Pd_2_(dba)_3_ (682 mg, 745 μmol), Xantphos (862 mg, 1.49 mmol) and Cs_2_CO_3_ (7.28 g, 22.3 mmol) in dioxane (50 mL) was stirred at 110°C for 12 hrs under N_2_ atmosphere. On completion, the reaction was filtered. The filtrate was concentrated in vacuo. The residue was purified by column chromatography (PE: EA = 20: 1) to give *tert*-butyl (4-((5-nitro-1-((2-(trimethylsilyl)ethoxy)methyl)-1H-pyrazol-4-yl)thio)phenyl) carbamate (1.20 g, 34% yield) as brown oil. ^1^H NMR (400 MHz, CDCl_3_) δ = 7.57 - 7.51 (m, 2H), 7.49 - 7.42 (m, 2H), 6.92 (s, 1H), 6.59 (s, 1H), 5.34 (s, 2H), 3.61 - 3.54 (m, 2H), 1.54 (s, 9H), 0.91 - 0.86 (m, 2H), ™0.02 (s, 9H).

To a solution of *tert*-butyl N-[4-[5-nitro-1-(2-trimethylsilylethoxymethyl)pyrazol-4- yl]sulfanylphenyl] carbamate (800 mg, 1.71 mmol) in DCM (20 mL) was added 2,6- dimethylpyridine (1.47 g, 13.7 mmol) and TMSOTf (1.52 g, 6.86 mmol). Then the mixture was stirred at 15°C for 1 hr. On completion, the reaction mixture was concentrated in vacuo. The residue was purified by column chromatography (PE: EA = 10: 1) to give 4-((5-nitro-1-((2- (trimethylsilyl)ethoxy)methyl)-1H-pyrazol-4-yl)thio)aniline (500 mg, 79% yield) as red oil. ^1^H NMR (400 MHz, CDCl_3_) δ = 7.40 (d, *J* = 8.8 Hz, 2H), 6.85 (s, 1H), 6.72 (d, *J* = 8.8 Hz, 2H), 5.33 (s, 2H), 3.92 (s, 2H), 3.56 (t, *J* = 8.0 Hz, 2H), 0.88 (t, *J* = 8.0 Hz, 2H), ™0.02 (s, 9H).

To a solution of 4-[5-nitro-1-(2-trimethylsilylethoxymethyl)pyrazol-4-yl]sulfanylaniline (500 mg, 1.36 mmol) in HOAc (10 mL) was added KSCN (530 mg, 5.46 mmol). Then the reaction mixture was stirred at 15°C for 30 min, and Br_2_ (218 mg, 1.36 mmol) in HOAc (2 mL) was added. The reaction mixture was stirred at 15°C for 12 hrs. On completion, reaction was concentrated in vacuo to remove HOAc, the residue was diluted with water (50 mL), basified with saturated NaHCO_3_ till pH=8 and extracted with EA (3 x 50 mL). The combined organic layer was washed with brine, and dried over Na_2_SO_4_, filtered and concentrated in vacuo The residue was purified by column chromatography (PE: EA = 10: 1) to give 6-((5-nitro-1-((2- (trimethylsilyl)ethoxy)methyl)-1H-pyrazol-4-yl)thio)benzo[d]thiazol-2- amine (450 mg, 78% yield) as a yellow solid. ^1^H NMR (400 MHz, CDCl_3_) δ = 7.85 (d, *J* = 1.2 Hz, 1H), 7.62 - 7.51 (m, 2H), 6.93 (s, 1H), 5.41 (s, 2H), 5.34 (s, 2H), 3.57 (t, *J* = 8.0 Hz, 2H), 0.88 (t, *J* = 8.0 Hz, 2H), ™0.03 (s, 9H).

A mixture of 6-[5-nitro-1-(2-trimethylsilylethoxymethyl)pyrazol-4-yl]sulfanyl-1,3- benzothiazol ™2-amine (400 mg, 944 μmol) and Raney Ni (20 mg) in MeOH (10 mL) was stirred at 25°C for 12 hrs under H_2_ (15 psi) atmosphere. On completion, the reaction was filtered and the filtrate was concentrated in vacuo to give *tert*-butyl (6-((1-((2-(trimethylsilyl)ethoxy)methyl)- 1H-pyrazol-4-yl)sulfonyl)benzo[d] thiazol-2-yl)carbamate (350 mg, 40% yield) as a yellow solid. ^1^H NMR (400 MHz, CDCl3) δ = 7.56 (s, 1H), 7.41 (d, *J* = 8.4 Hz, 1H), 7.32 (d, *J* = 1.6 Hz, 1H), 7.14 (dd, *J* = 1.6, 8.4 Hz, 1H), 5.31 (br s, 2H), 5.25 (s, 2H), 3.96 (s, 2H), 3.60 (t, *J* = 8.0 Hz, 2H), 0.94 (t, *J* = 8.0 Hz, 2H), 0.01 (s, 9H).

To a solution of 6-[5-amino-1-(2-trimethylsilylethoxymethyl)pyrazol-4-yl]sulfanyl-1,3- benzothiazol-2-amine (100 mg, 254 μmol) in H_2_O (5 mL) and ACN (5 mL) was added Oxone (624 mg, 1.02 mmol) at 0°C. Then the reaction mixture was stirred at 15°C for 1.5 hrs. On completion, the reaction mixture was extracted with EA (3 x 20 mL). The combined organic layer was washed with saturated Na_2_SO_3_ solution, brine and dried over Na_2_SO_4_ to give a brown solid (100 mg, 80% yield). 10 mg was purified by Prep-HPLC (column: Phenomenex Synergi C18 150*30 mm*4 μm; mobile phase: [water (0.05% HCl)-ACN]; B%: 35%-55%, 12 min) to give 6-((5-amino-1-((2-(trimethylsilyl)ethoxy)methyl)-1H-pyrazol-4- yl)sulfonyl)benzo[d]thiazol-2-amine (6.90 mg, HCl salt, SEM-position on pyrazol was not confirmed) as a brown solid. LCMS (M+1)^+^: 426.1. ^1^H NMR (400 MHz, DMSO-*d*_6_) δ = 8.39 (s, 1H), 8.32 (s, 1H), 8.23 (br s, 2H), 7.83 (d, *J* = 8.8 Hz, 1H), 7.52 (d, *J* = 8.4 Hz, 1H), 5.23 (s, 2H), 3.59 - 3.54 (m, 2H), 0.91 - 0.85 (m, 2H), 0.00 (s, 9H). To a solution of 6-[5-amino-1-(2- trimethylsilylethoxymethyl)pyrazol-4-yl]sulfonyl-1,3-benzothiazol-2-amine (50.0 mg, 117 μmol) in DCM (1 mL) was added TFA (1 mL). Then the reaction mixture was stirred at 15°C for 30 min. On completion, the reaction was concentrated in vacuo. The residue was purified by Prep-HPLC (column: Phenomenex Synergi C18 150*30 mm*4 μm; mobile phase: [water (0.05% HCl)-ACN]; B%: 6%-26%, 12 min) to give **C-35** (3.16 mg, 9% yield, HCl salt) as colorless oil. LCMS (M+1)^+^: 296.0. ^1^H NMR (400 MHz, DMSO-*d*_6_) δ = 8.55 (br s, 2H), 8.33 (d, *J* = 2.0 Hz, 1H), 7.77 (dd, *J* = 2.0, 8.4 Hz, 1H), 7.64 (s, 1H), 7.46 (d, *J* = 8.4 Hz, 1H). ^13^C NMR (101 MHz, DMSO-*d_6_*) δ 170.66, 149.89, 138.93, 136.98, 127.13, 125.53, 121.59, 115.62, 102.97.

***C-36***. A mixture of 1-benzyl-4-iodo-pyrazole (2.00 g, 3.50 mmol), 4-nitrobenzenethiol (655 mg, 4.20 mmol), CuI (67.0 mg, 352 μmol), Cs_2_CO_3_ (2.30 g, 7.00 mmol) in DMF (30 mL) was stirred at 110°C for 12 hrs. On completion, the reaction mixture was concentrated in vacuo. The residue was purified by silica gel chromatography (PE:MA = 15: 1) to give 1-benzyl-4-(4- nitrophenyl)sulfanyl-pyrazole (0.800 g, 36% yield) as yellow solid. LCMS (MS+1)^+^: 312.0. To a mixture of 1-benzyl-4-(4-nitrophenyl) sulfanyl-pyrazole (0.800 g, 2.40 mmol) and NH_4_Cl (90.2 mg, 1.70 mmol) in EtOH (24 mL) and H_2_O (4 mL) was added Fe (807 mg, 14.5 mmol) slowly at 80°C. The reaction mixture was stirred at 80°C for 2 hrs. On completion, the reaction mixture was *concentrated* in vacuo. The residue was purified by silica gel chromatography (PE:MA = 5: 1) to give 4-(1-benzylpyrazol-4-yl)sulfanylaniline (0.600 g, 80% yield) as light yellow solid. ^1^H NMR (400 MHz, DMSO-*d*_6_) δ = 7.99 (s, 1H), 7.49 (s, 1H), 7.34 - 7.32 (m, 3H), 7.23 - 7.21 (m, 2H), 7.00 (d, *J* = 8.4 Hz, 2H), 6.50 (d, *J* = 8.8 Hz, 2H), 5.30 (s, 2H), 5.18 (s, 2H).

A mixture of 4-(1-benzylpyrazol-4-yl)sulfanylaniline (600 mg, 1.70 mmol) and KSCN (663 mg, 6.80 mmol) in AcOH (5 mL) was stirred at 20°C for 0.5 hr. Then a solution of Br_2_ (286 mg, 1.80 mmol) in AcOH (1 mL) was added dropwise. The reaction mixture was stirred at 25°C for 12.5 hrs. On completion, the reaction was poured into ice water and basified with NaHCO_3_ till pH = 7, and a white solid was precipitated. The mixture was filtered and the filter cake was washed with water, dried in vacuo to give 6-(1-benzylpyrazol-4-yl)sulfanyl-1,3- benzothiazol-2-amine (0.46 g, 64% yield) as a light yellow solid. ^1^H NMR (400 MHz, DMSO- *d*_6_) δ = 8.17 (s, 1H), 7.63 (s, 1H), 7.51 - 7.46 (m, 3H), 7.35 - 7.29 (m, 3H), 7.26 - 7.21 (m, 3 H), 7.06 - 7.04 (m, 1 H), 5.35 (s, 2 H).

To a mixture of 6-(1-benzylpyrazol-4-yl) sulfanyl-1,3-benzothiazol-2-amine (100 mg, 295 μmol) in H_2_O (1 mL) and THF (3 mL) was added Oxone (363 mg, 590 μmol) at 0°C for 4 hrs. On completion, the reaction mixture was quenched by addition H_2_O (50 mL), filtered and the filter cake was dried in vacuo. The residue was purified by Prep-TLC (DCM: MeOH**=** 30:1) to give **C-36** (60.0 mg, 53% yield) as a white solid. LCMS (MS+1)^+^: 371.0. ^1^H NMR (400 MHz, DMSO-*d*_6_) δ = 8.56 (s, 1 H), 8.28 (s, 1 H), 7.99 (s, 2 H), 7.93 (s, 1 H), 7.73 - 7.71 (m, 1 H), 7.43 - 7.41 (m, 1 H), 7.34 - 7.33 (m, 2 H), 7.31 - 7.29 (m, 1 H), 7.27 - 7.25 (m,2 H), 5.34 (s, 2 H). ^13^C NMR (101 MHz, DMSO-*d_6_*) δ 170.75, 157.11, 139.13, 136.63, 134.49, 133.11, 132.16, 129.13, 128.51, 128.43, 125.02, 124.90, 120.76, 117.95, 55.81.

## Supporting Information Statement

The Supporting Information is available free of charge at XXX.

Complete details on methods and additional figures for compound synthesis and spectra, biochemical assays, docking studies, bacterial phenotype assays, cellular and *in vivo* toxicity, cloning, protein expression and purification, and NMR studies. (PDF) Molecular formula strings (CSV)

## Acknowledgements

We gratefully acknowledge funding as follows: National Institutes of Health Biotechnology Training Grant 5T32GM008347 (H.K.L.)., University of Minnesota Department of Chemistry Excellence Fellowship (H.K.L.), National Institutes of Health grant GM134538-01A1, and the UMN Office of Academic Clinical Affairs. This work was also supported by funding provided under cooperative agreement #IDSEP160030-01-00 from Biomedical Advanced Research and Development Authority (BARDA). Its contents are solely the responsibility of the authors and do not necessarily represent the official views of The Assistant Secretary for Preparedness and Response. The authors acknowledge the Minnesota Supercomputing Institute (MSI) at the University of Minnesota for providing resources that contributed to the research results reported within this paper. BioRender was used for generation of some figures.

## Author Contribution Statement

The manuscript was written through the contributions of all authors. C. Fihn conceived of the idea, planned and carried out the experiments, data analysis, and interpretation except as indicated for other authors, as well as wrote the manuscript. H. Lembke assisted in writing, performed growth curve experiments and analysis, and assisted on *in vitro* activity and aggregation assays. J. Gaulin contributed to analogue design, synthesis was performed by contract with WuXi AppTec. P. Bouchard carried out NMR experimental design, sample preparation, acquisition, and analysis, along with writing for the NMR studies. A. Villarreal validated the effects of select inhibitors on *P. aeruginosa* physiology via growth assays, liquid chromatography-based phenazine quantification, and subsequent data analysis. M. Penningroth planned and carried out inhibitor cytotoxicity experiments and analysis with assistance from G. Vogt. A. Gilbertsen designed and executed animal experiments and data analysis. Y. Ayotte and L. Couthino de Oliveira assisted with sample preparation, acquisition, and data analysis for NMR studies. K. Crone assisted on *in vitro* activity assays. N. Drouin expressed and purified protein for NMR studies. M. Serrano-Wu assisted in manuscript drafting. D. Hung, R. Hunter, and E. Carlson directed study design, data analysis, manuscript preparation and editing, and obtained funding. All authors have given approval to the final version of the manuscript.

---

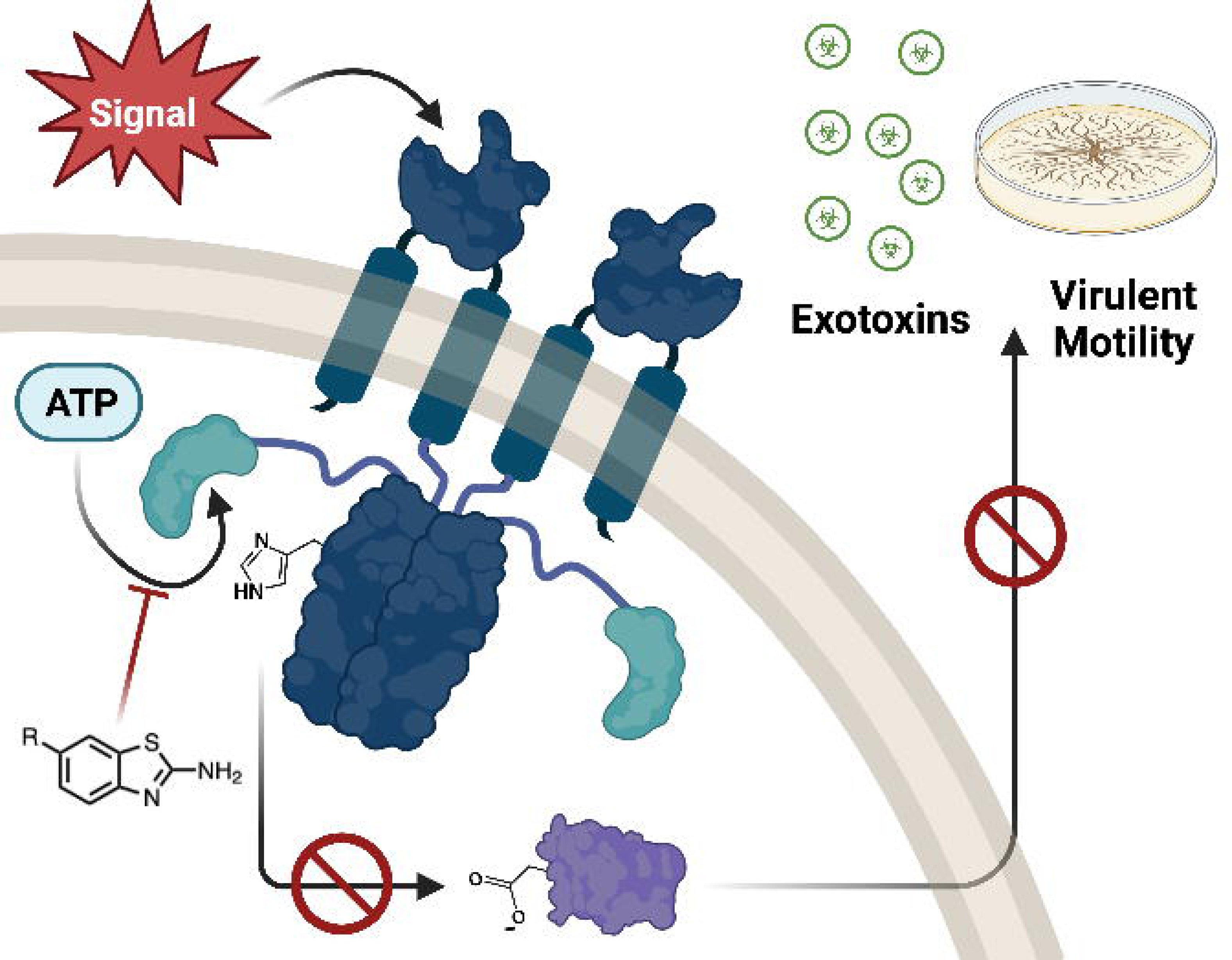

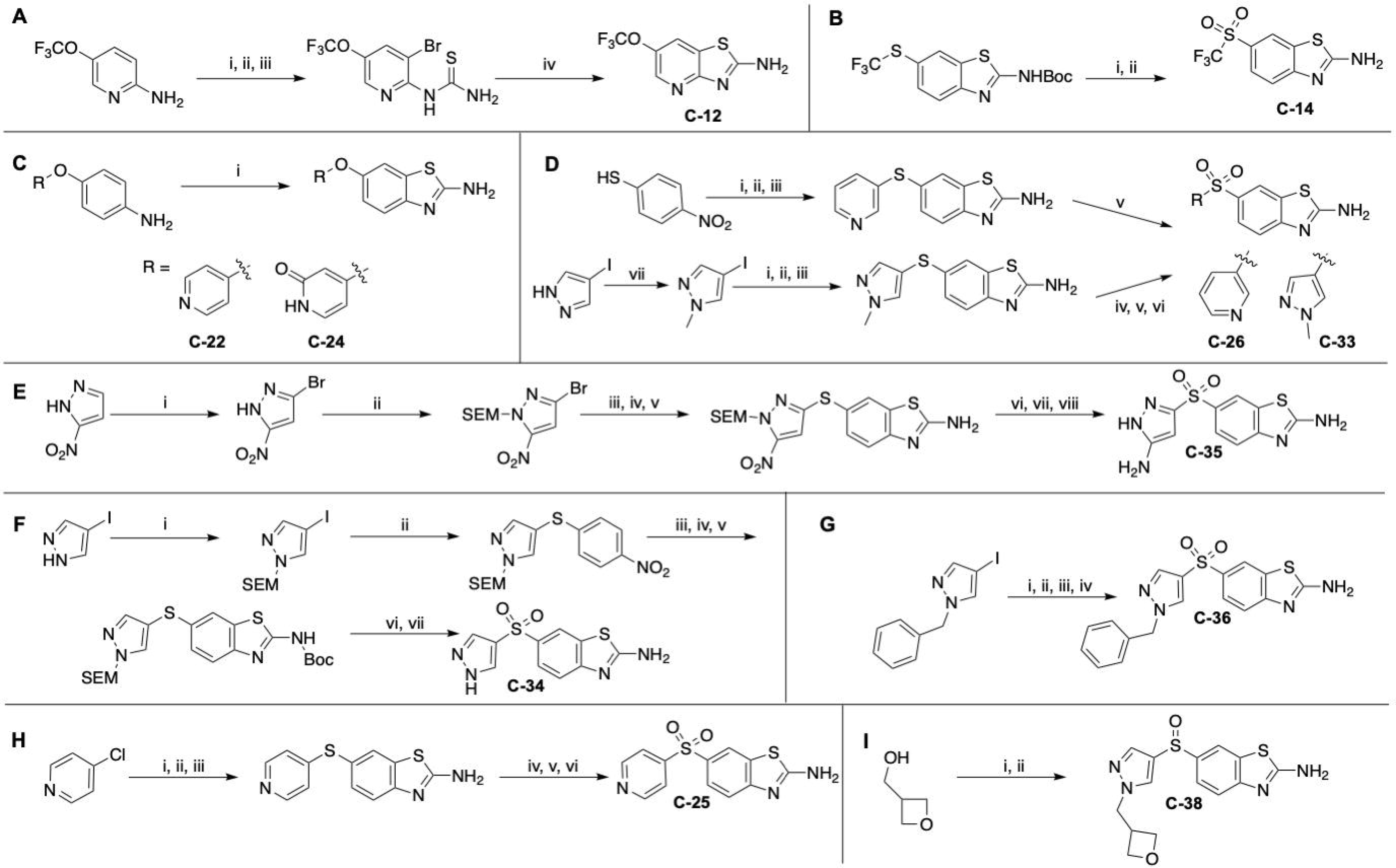

