## Supporting Information Document for "Evaluation of Expanded 2-Aminobenzothiazole Library for Inhibition of *Pseudomonas aeruginosa* Virulence Phenotypes"

[a]*Department of Medicinal Chemistry, University of Minnesota, 308 Harvard Street SE, Minneapolis, Minnesota 55455, United States*
[b]*Department of Chemistry, University of Minnesota, 207 Pleasant Street SE, Minneapolis, Minnesota 55454, United States*
[c]*The Broad Institute of MIT and Harvard, Cambridge, Massachusetts 02142, United States*
[d]*NMX Research and Solution Inc., 500 Cartier Boulevard W., Suite 6000, Laval, Quebec, Canada, H1Y 2R1*
[e]*Department of Microbiology & Immunology, University of Minnesota, 689 23rd Ave Se Minneapolis, Minnesota 55455, United States*
[f]*Department of Biochemistry, Molecular Biology and Biophysics, University of Minnesota, 321 Church Street SE, Minneapolis, Minnesota 55455, United States*
[g]*3 Point Bio LLC, 245 Main Street, Cambridge, Massachusetts 02142, United States*

**Table of Contents:**

**Supporting Figures and Tables**
SI Figure 1. **White blood cell counts in mice upon exposure to Rilu-compounds**
SI Figure 2. **Three concentration *in vitro* inhibitor screen**
SI Figure 3. **Single concentration aggregation test for inhibitors**
SI Figure 4. **Serial dilution series in gel activity assays**
SI Figure 5. **Inhibitor π-π stacking**
SI Figure 6. **Predicted inhibitor 'flipped' binding poses**
SI Figure 7. **Folding evaluation of HK (232-489) by 1D NMR**
SI Figure 8. **Evaluation of compound binding by 2D HSQC**
SI Table 1. **Specific growth rates**
SI Figure 9. **Area under the growth curve following inhibitor treatment**
SI Figure 10. **Swarm assay plate images**
SI Figure 11. **Swarm assay result bar graph**
SI Figure 12. **Pyocyanin assay result bar graph**
SI Figure 13. **Metabolite quantification at 24 hrs**
SI Figure 14. **Metabolite quantification at 48 hrs**
SI Figure 15. **Cytotoxicity of Rilu-series compounds in Hep G2 cells**
SI Figure 16. **Cytotoxicity of Rilu-series compounds in A549 cells**

**Synthesis Methods**
**Molecule Characterization**

#### Supplementary Figures

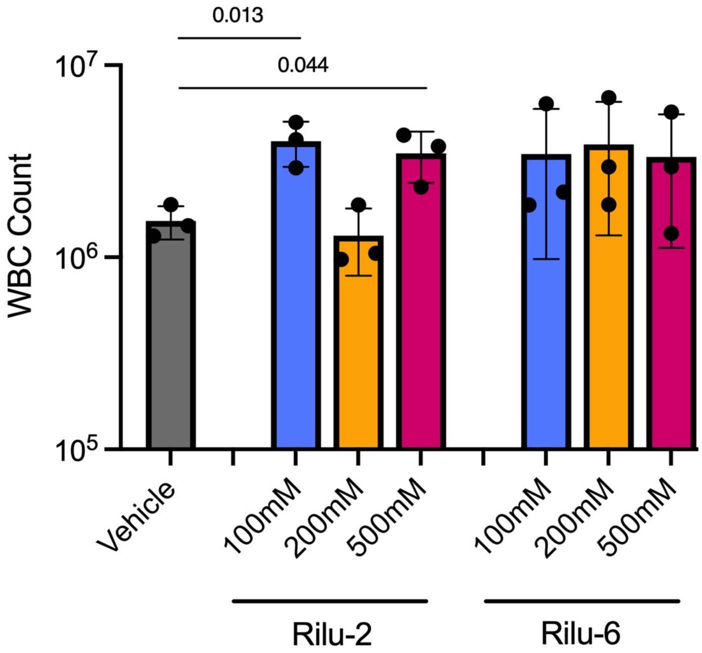


SI Figure 1. **White blood cell counts in mice upon exposure to Rilu-compounds (BALB/c).** Mice were challenged intratracheally with Rilu inhibitors in 1% DMSO in PBS (100 μL). Total white blood cell count was determined by bronchoalveolar lavage and trypan blue staining after 24 h. Data were compared using a one-way ANOVA with multiple comparisons.

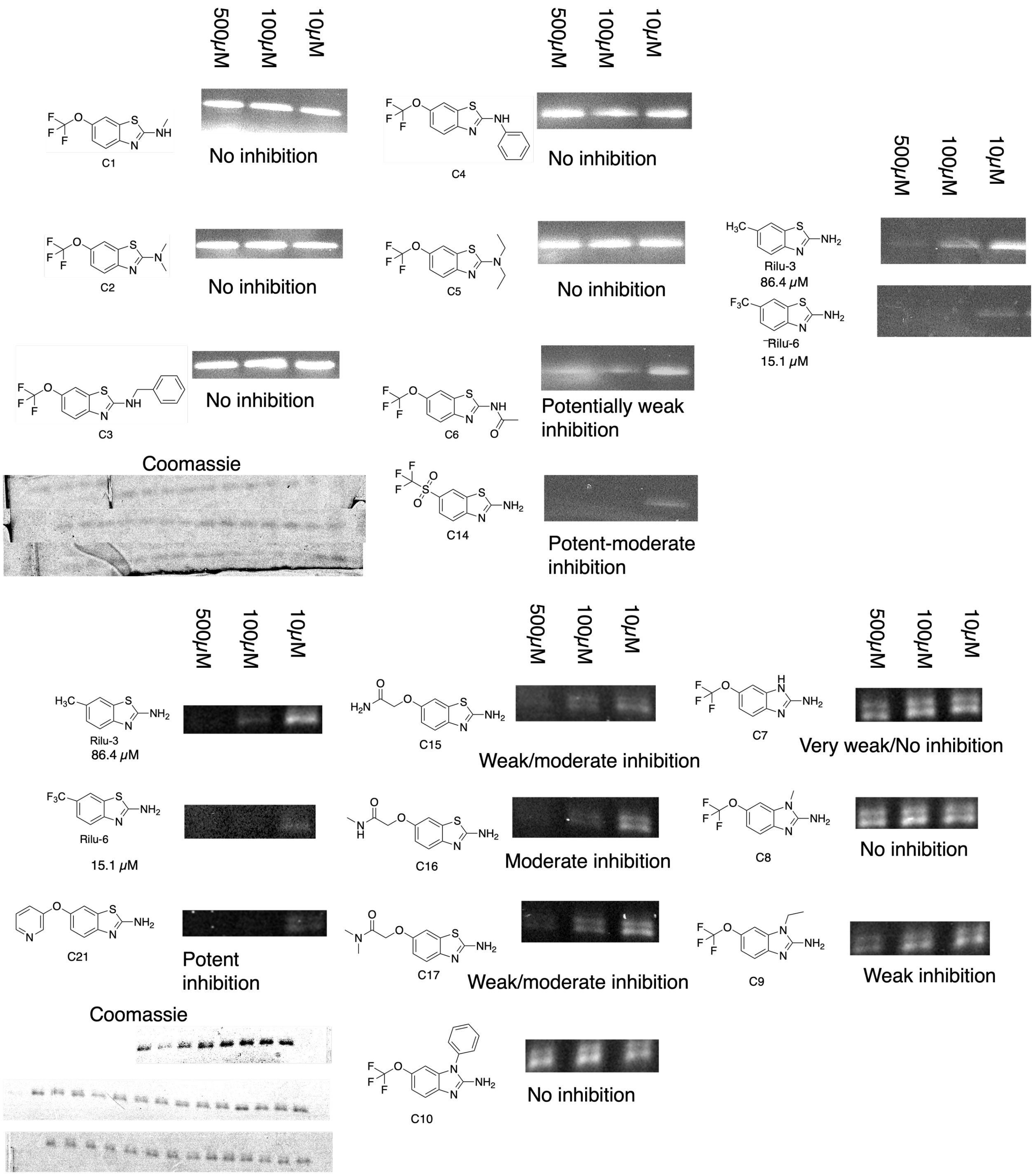
500μM
100μM
10μM
C1
No inhibition
C2
No inhibition
C3
No inhibition
C4
No inhibition
C5
No inhibition
C6
Potentially weak inhibition
C14
Potent-moderate inhibition
Coomassie
Rilu-3
86.4 μM
Rilu-6
15.1 μM
C21
Potent inhibition
Coomassie
C15
Weak/moderate inhibition
C16
Moderate inhibition
C17
Weak/moderate inhibition
C10
No inhibition
C7
Very weak/No inhibition
C8
No inhibition
C9
Weak inhibition

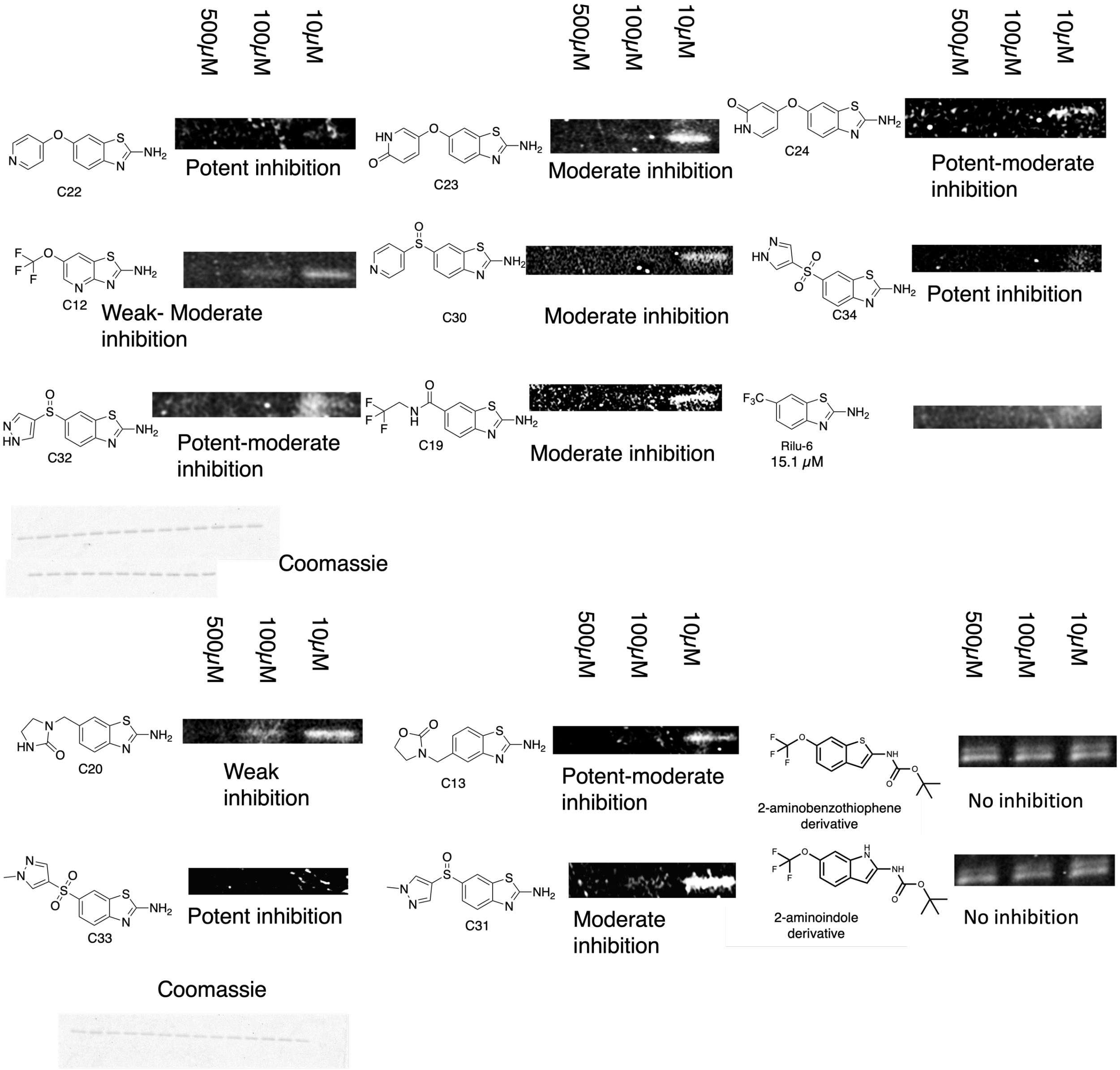


SI Figure 2. **Preliminary three concentration *in vitro* inhibitor screen:** Three concentration screen of 2-aminobenzothiazole analogues. Compounds were judged for inhibition levels and any compound that showed inhibition capacity was tested further in complete dilution series. Not all inhibitors were tested using this method, some were taken directly to full concentration range assay (**C-11**, **C-18**, **C25-29**). Total protein content was determined by Coomassie staining. Gels were scanned on Typhoon FLA 9500 scanner (GE healthcare) on BODIPY or Coomassie filter settings.

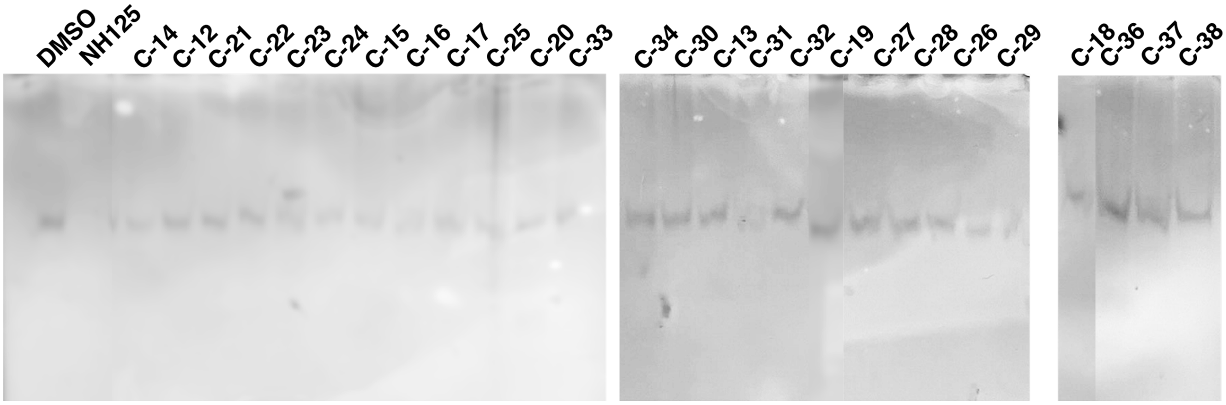


SI Figure 3. **Single concentration aggregation test for inhibitors:** Aggregation screen for inhibitors at 200 µM concentration (0.4% DMSO in all samples). Only molecules showing promising inhibition of HK activity were tested in this assay. Proteins were resolved by native-PAGE and silver stained. NH125, a non-specific aggregator[1], used as positive control. Disappearance of the dimeric band indicates formation of higher order aggregates, preventing the protein from migrating into the gel. None of the test compounds showed significant aggregatory effects.

C-12

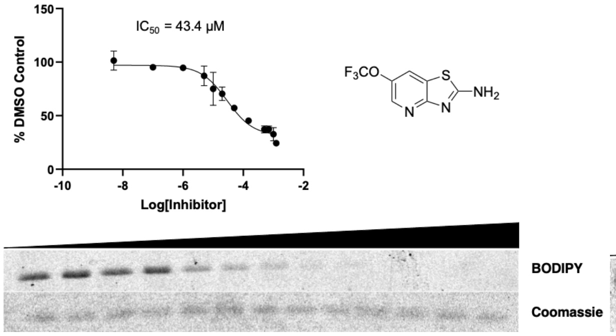


C-13

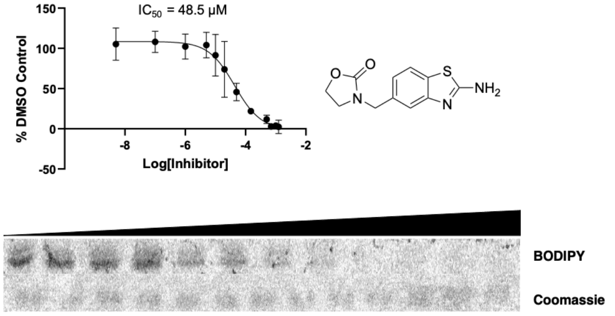


C-14

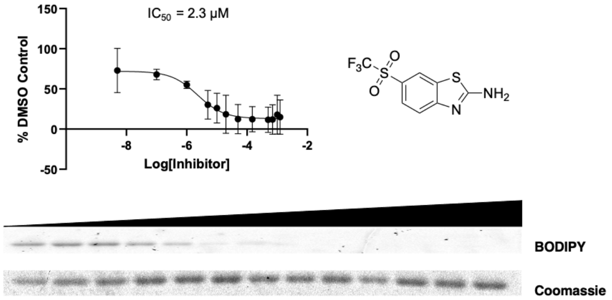


C-15

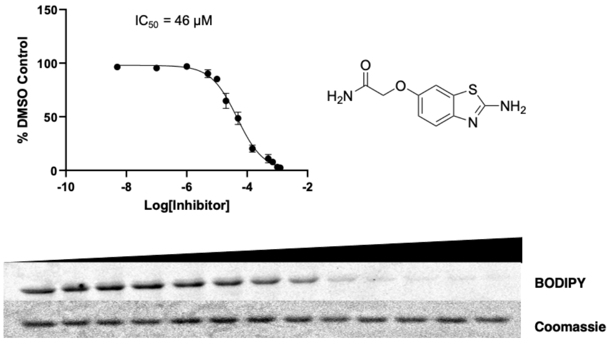


C-16

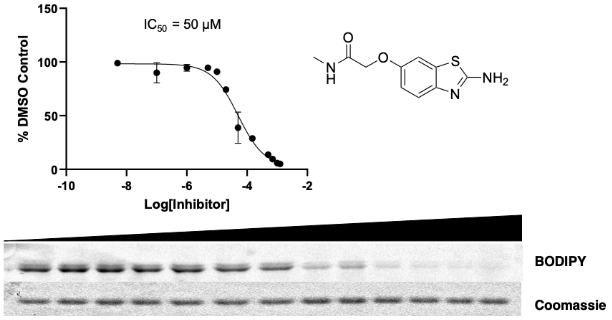


C-17

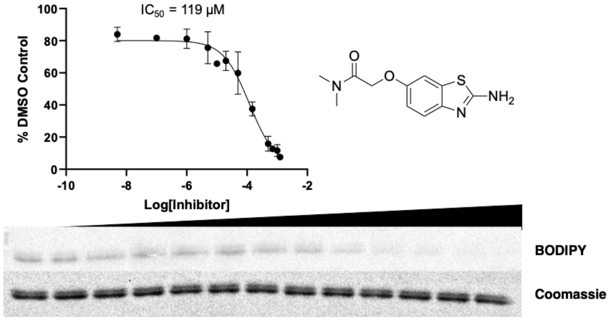


C-18

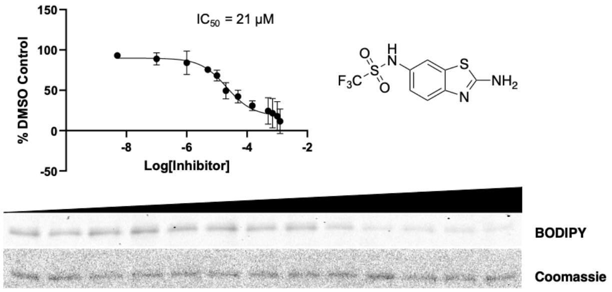


C-19

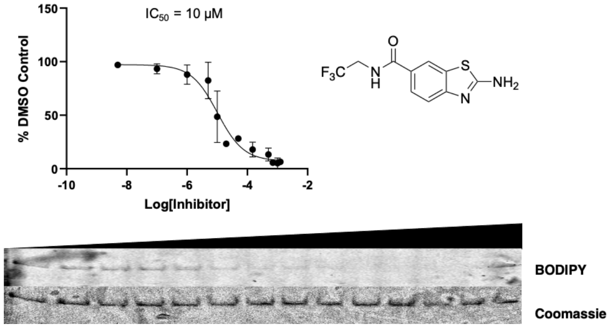

C-20

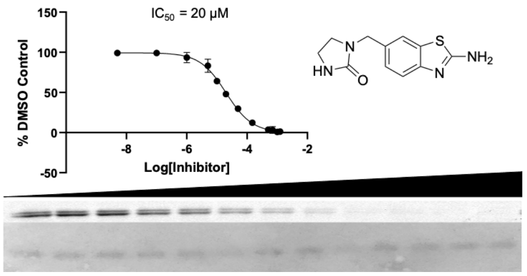


C-21

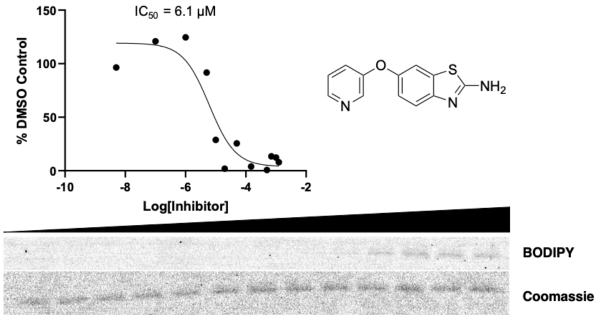


C-22

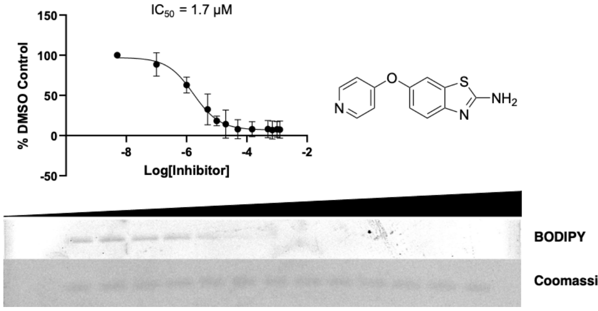


C-23

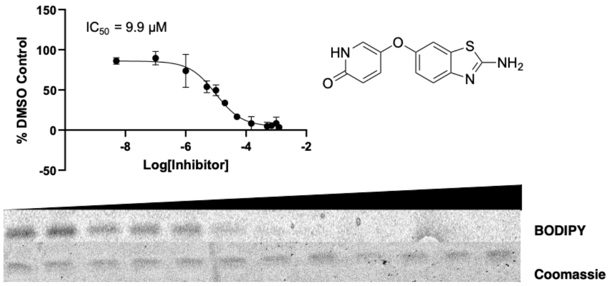


C-24

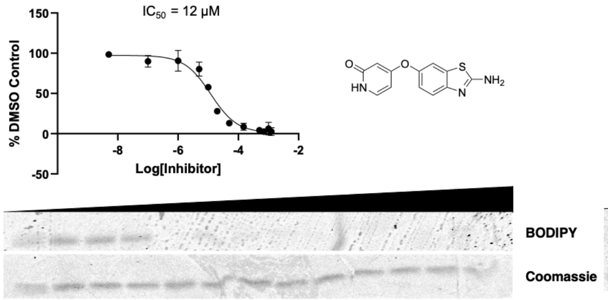


C-25

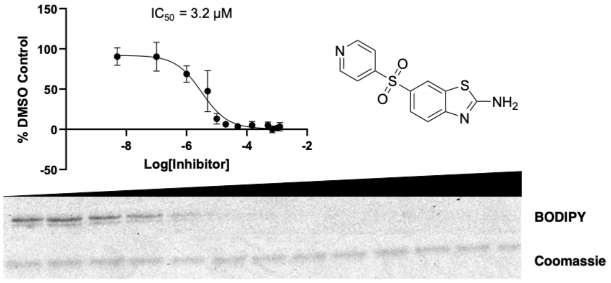


C-26

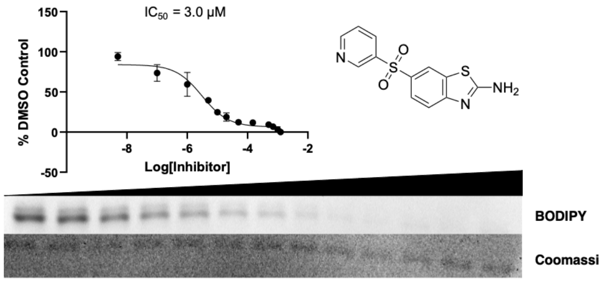


C-27

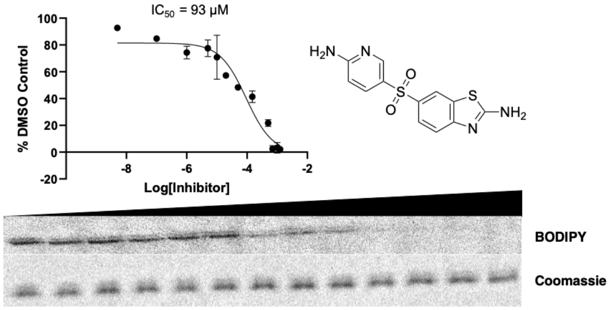


C-28

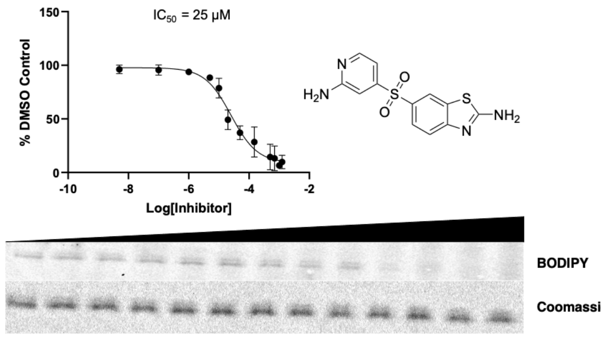


C-29

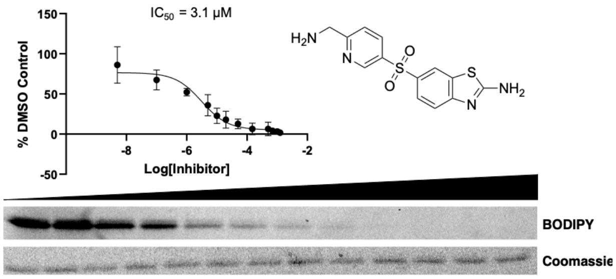

**C-30**

$IC_{50}$ = 18 µM

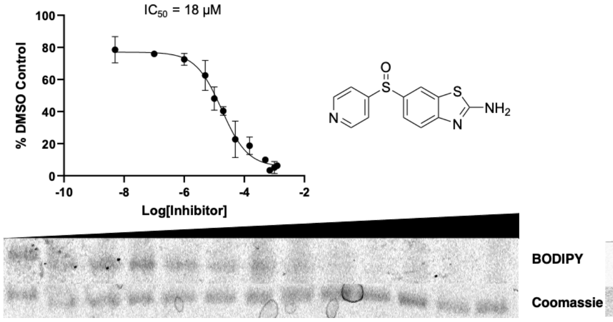


**C-31**

$IC_{50}$ = 22 µM

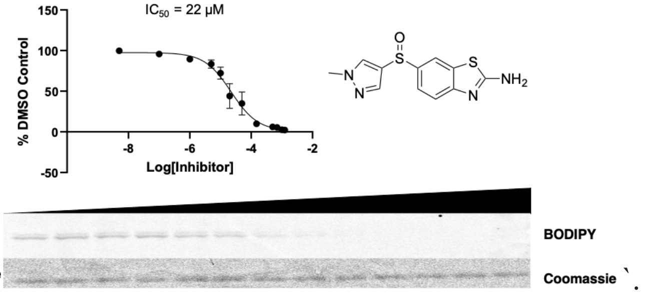


**C-32**

$IC_{50}$ = 14 µM

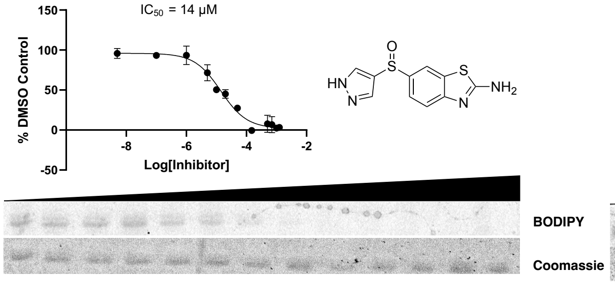


**C-33**

$IC_{50}$ = 1.7 µM

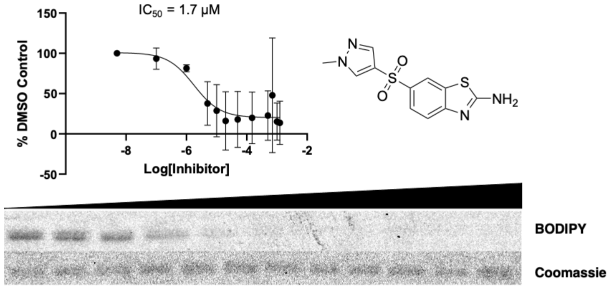


**C-34**

$IC_{50}$ = 0.42 µM

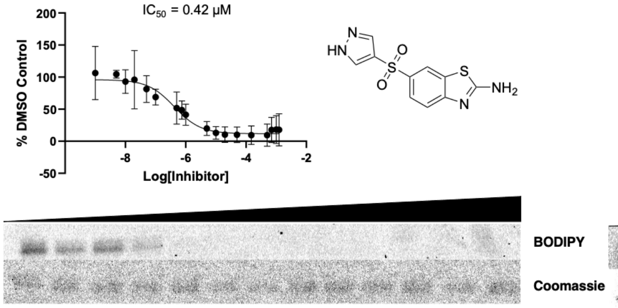


**C-35**

$IC_{50}$ = 4.2 µM

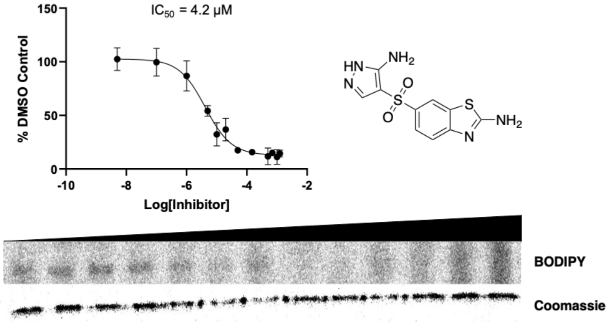


**C-36**

$IC_{50}$ = 1.9 µM

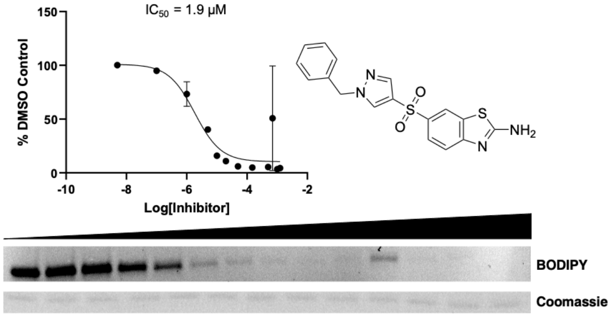


**C-37**

$IC_{50}$ = 8.8 µM

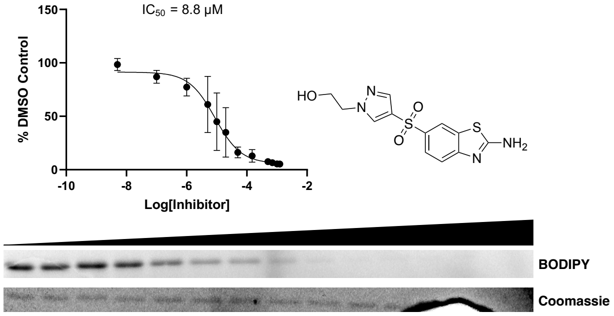


**C-38**

$IC_{50}$ = 3.4 µM

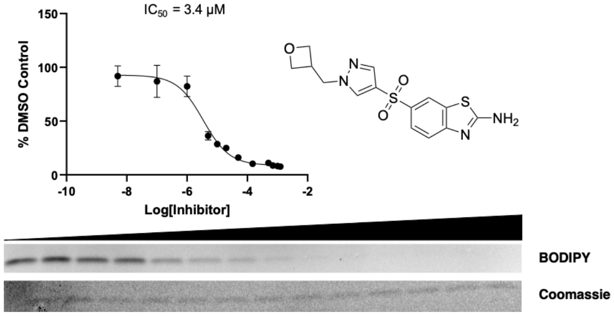

SI Figure 4. **Serial dilution series in gel-based activity assays:** In-gel activity assay measuring autophosphorylation of HK853 with varying concentrations of inhibitors from DMSO stocks (0 μM, 0.01 μM, 0.1 μM, 1 μM, 5 μM, 10 μM, 20 μM, 50 μM, 150 μM, 500 μM, 700 μM, 1000 μM, 1250 μM). Shown are representative gels scanned for BODIPY fluorescence, Coomassie stain for total protein, and $IC_{50}$ curves of replicate averages. Gels were scanned on Typhoon FLA 9500 scanner (GE healthcare) on BODIPY or Coomassie filter settings. Integrated density measurements of in-gel fluorescence and phosphorescence were performed in ImageJ.[2] Data were prepared and analyzed in GraphPad Prism (version 9.0 for Windows, GraphPad Software, San Diego, California USA, www.graphpad.com). For all dose response curves, data were fit to a four-parameter logistic equation.

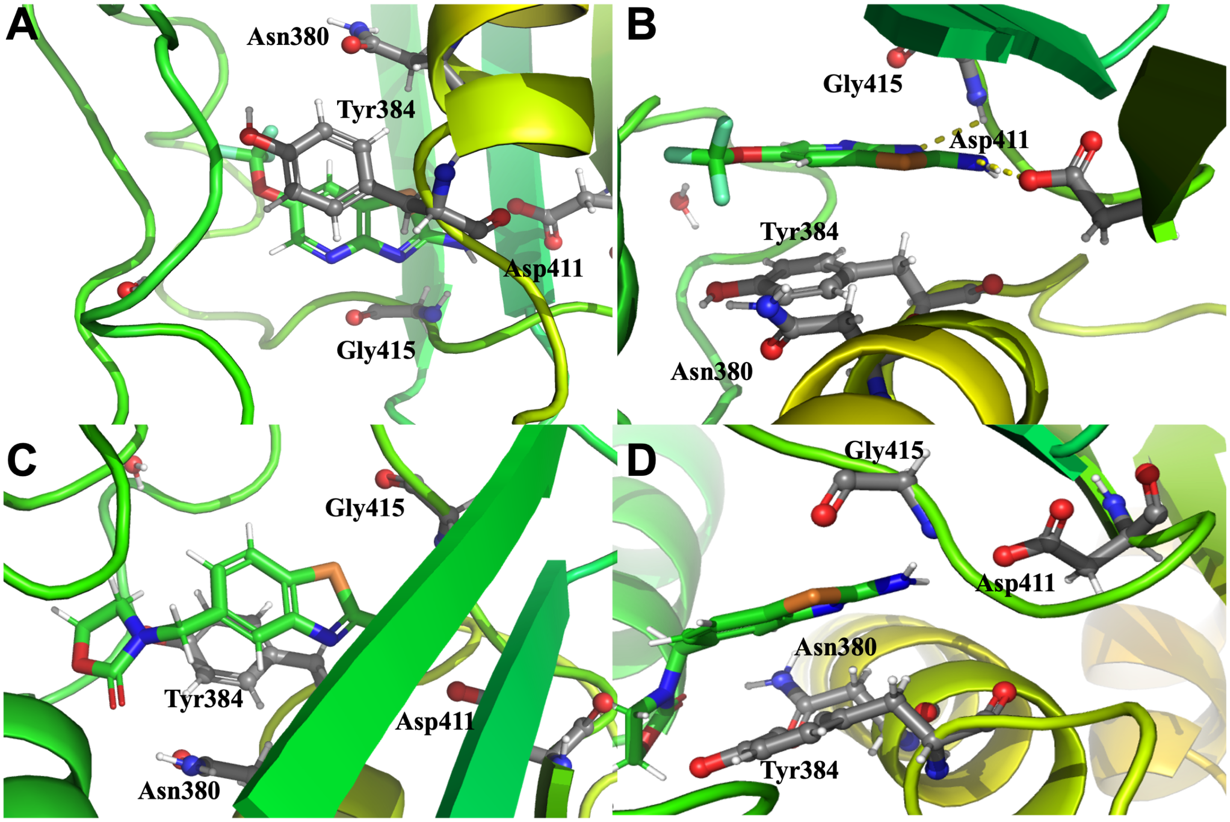


SI Figure 5. **Inhibitor π-π stacking:** Docking images showing potential binding poses of putative HK inhibitors **C-12** (**A** & **B**) and **C-13** (**C** & **D**) in CA domain active site of HK853 (PDB: 3DGE).[3] Both inhibitors, along with almost all other inhibitors that are not shown here, adopt a pose that places them in a potential parallel displaced or staggered stacking with tyrosine 384 (**A** & **C**) and put them in close enough proximity at approximately 3.5-4 Å from the residue to allow such an interaction (**B** & **D**). Docking was performed using Shrodinger Maestro and images created using Pymol.

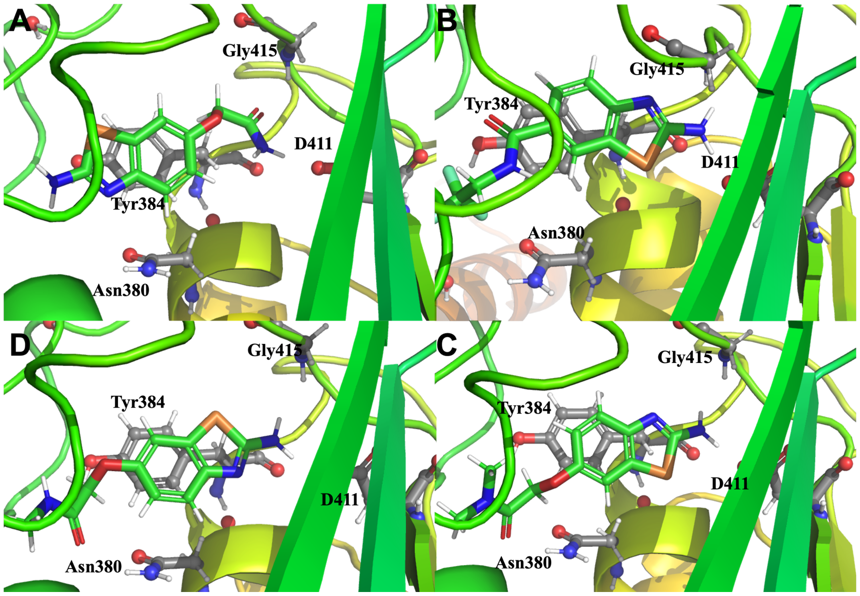


SI Figure 6. **Predicted inhibitor 'flipped' binding poses:** Docking images showing potential binding poses of inhibitors **C-15 (A)**, **C-16 (B)**, and **C-17 (C)** in CA domain (HK853; PDB: 3DGE). Clockwise from top left **C-15 (A)**, **C-16 (B)**, **C-17 (C)** and **C-19 (D)**. **C-15** adopted a flipped orientation relative to other inhibitors with the 2-aminothiazole moiety pointing towards the other side of the pocket while the unhindered acetamide moiety interacts with aspartate reside. Hindered acetamide groups were not able to adopt this flipped orientation. Docking was performed using Shrodinger Maestro and images created using Pymol.

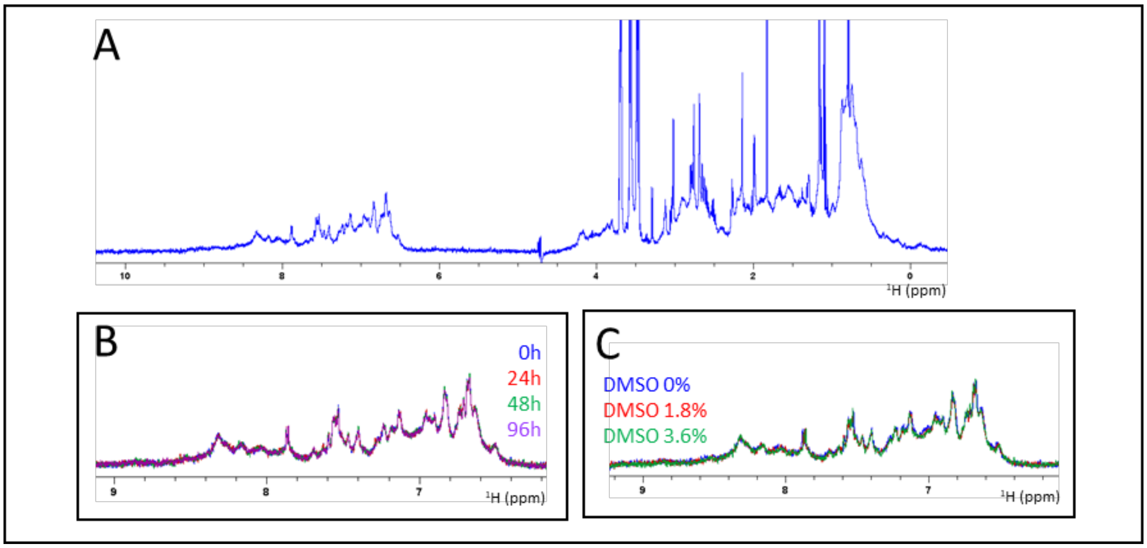


SI Figure 7. **Folding evaluation of HK853 (residues 232-489) by 1D NMR**: **A**) Full 1D $^{1}$H spectrum of HK853 (232-489). Spectra overlays zoomed in on the aromatic region showing **B**) the protein profile after various time points and **C**) the protein profile in presence of varying amounts of DMSO-$d_6$.

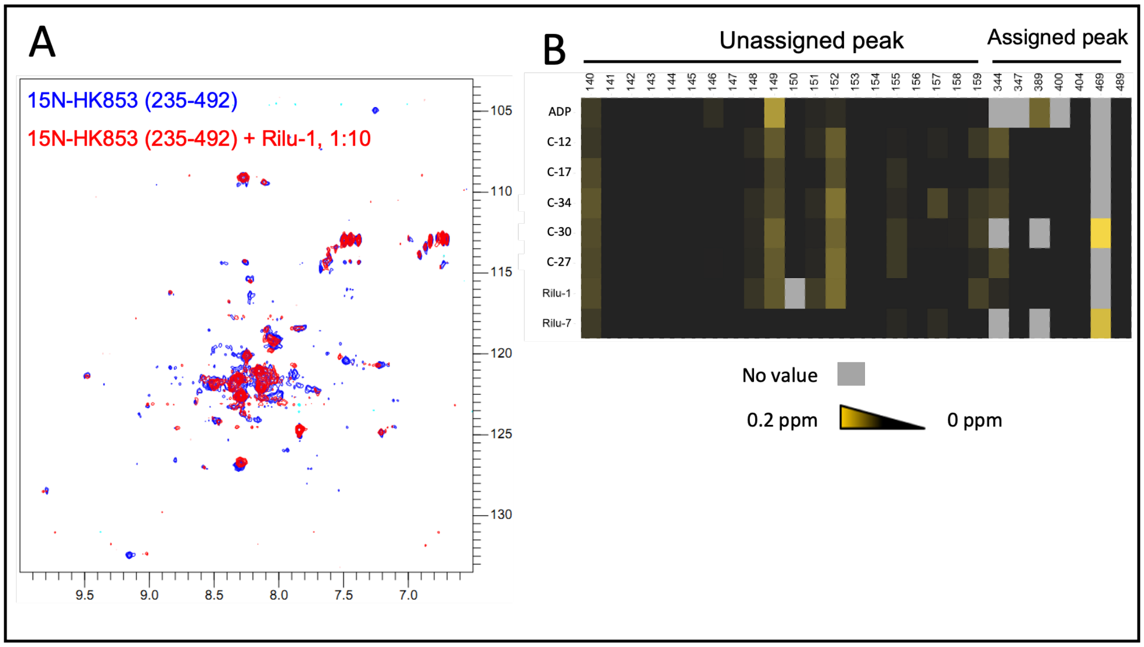


SI Figure 8. **Evaluation of compound binding by 2D HSQC: A)** 2D HSQC-TROSY spectra of $^{15}$N-labeled HK853 (235-492) in its apo state (in blue) and in presence of 10 molar equivalent of compound **Rilu-1** (in red). **B**) Heat map representations of the chemical shift perturbation (CSP) observed on HK853 (235-492) upon addition of different compounds to evaluate the similarities / differences in binding fingerprints. The color scale goes from black (no CSP) to yellow (in this case maximal CSP of 0.2 ppm). Most of the spectra was unassigned and a peak number was assigned to all peaks resolved enough for analysis. Only a few resonances were assigned from available literature[4], see assigned peak number. The grey color represents peak for which there is no value available (low signal to noise, overlap, etc).

**SI Table 1. Specific growth rates.** The maximum specific growth rate ($\mu_{max}$) for each sample with its standard deviation.

| Sample | $\mu_{max}$ |
|---|---|
| DMSO | 1.74±0.11 |
| R1 | 1.48±0.13 |
| R2 | 1.62±0.33 |
| R6 | 1.45±0.10 |
| R7 | 1.67±0.15 |
| C-12 | 1.45±0.09 |
| C-13 | 1.31±0.16 |
| C-15 | 1.76±0.15 |
| C-16 | 1.83±0.13 |
| C-18 | 1.36±0.10 |
| C-19 | 1.33±0.09 |
| C-20 | 2.15±0.12 |
| C-21 | 1.38±0.41 |
| C-22 | 1.46±0.07 |
| C-23 | 1.82±0.14 |
| C-24 | 1.62±0.14 |
| C-25 | 2.15±0.64 |
| C-27 | 1.49±0.26 |
| C-28 | 1.49±0.14 |
| C-29 | 1.16±0.23 |
| C-30 | 1.36±0.13 |
| C-31 | 1.47±7.72 |
| C-32 | 1.33±0.20 |
| C-33 | 1.98±0.11 |
| C-34 | 2.11±0.15 |
| C-37 | 1.29±0.19 |
| C-38 | 1.29±0.16 |

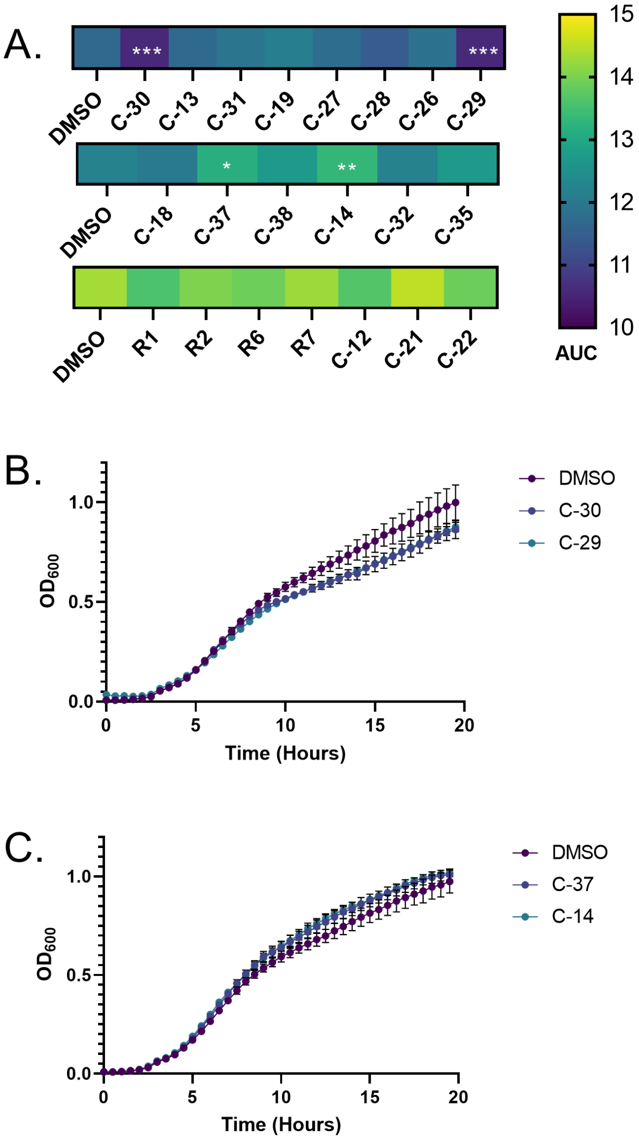


SI Figure 9. **Total area under the curve (AUC) measurement following inhibitor treatment.** **A.** Area under the curve (AUC) average measurement calculated for each inhibitor (200 µM) from three biological replicate growth curves. Due to the number of inhibitors, multiple plates were required and all inhibitors were compared to the DMSO control (0.4%) on that plate. One-Way ANOVA was completed with Prism 9.0 comparing each inhibitor to the DMSO control on its plate. Statistically significant differences are denoted as follows: p-value <0.05 *, <0.01 ** and <0.005 ***. **B**. Average growth curves of significantly different inhibitors compared to DMSO control for **C-29** and **C-30**. **C**. Average growth curves of significantly different inhibitors compared to DMSO control for **C-37** and **C-14**.

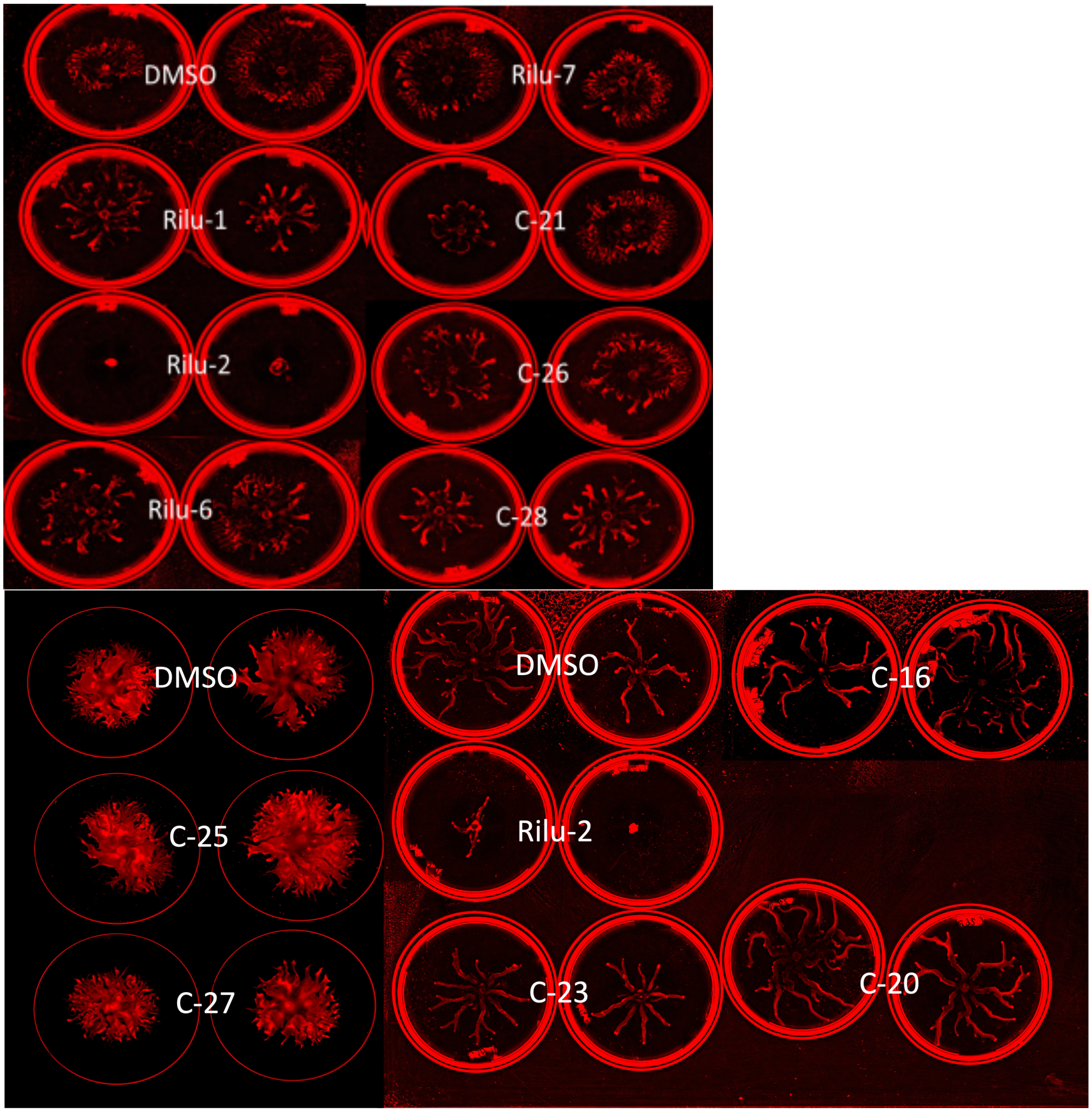
DMSO
Rilu-7
Rilu-1
C-21
Rilu-2
C-26
Rilu-6
C-28
DMSO
DMSO
C-16
C-25
Rilu-2
C-27
C-23
C-20

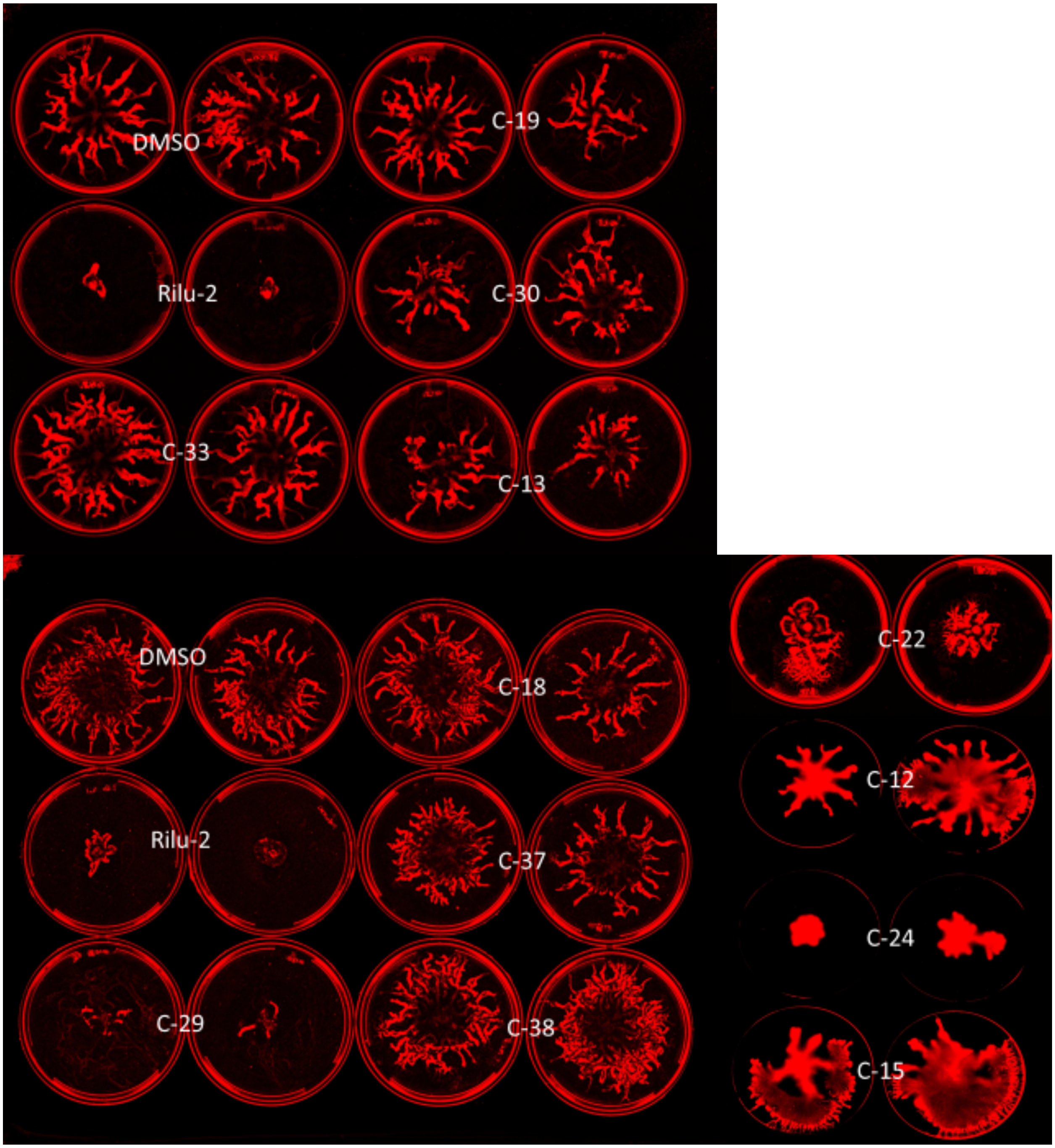
DMSO
C-19
Rilu-2
C-30
C-33
C-13
DMSO
C-18
Rilu-2
C-37
C-29
C-38
C-22
C-12
C-24
C-15

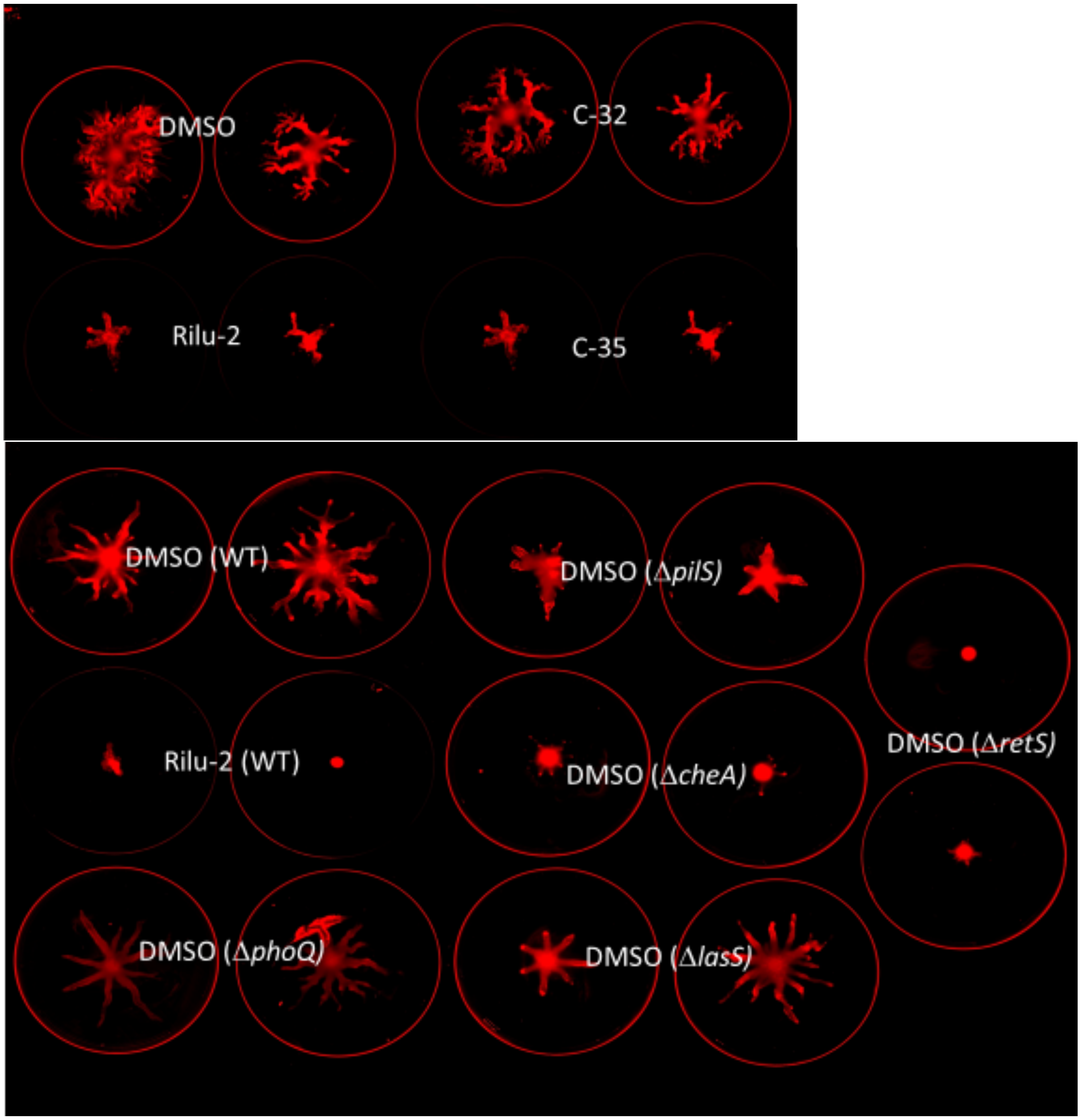


SI Figure 10. **Swarm assay plate images:** Modified FAB media swarm plates of PA14 when exposed to inhibitors at 100 μM. Inhibitors were compared to DMSO control within same experiment (0.4% final DMSO concentration). WT PA14 was compared to swarm of several knockout strains of HKs.

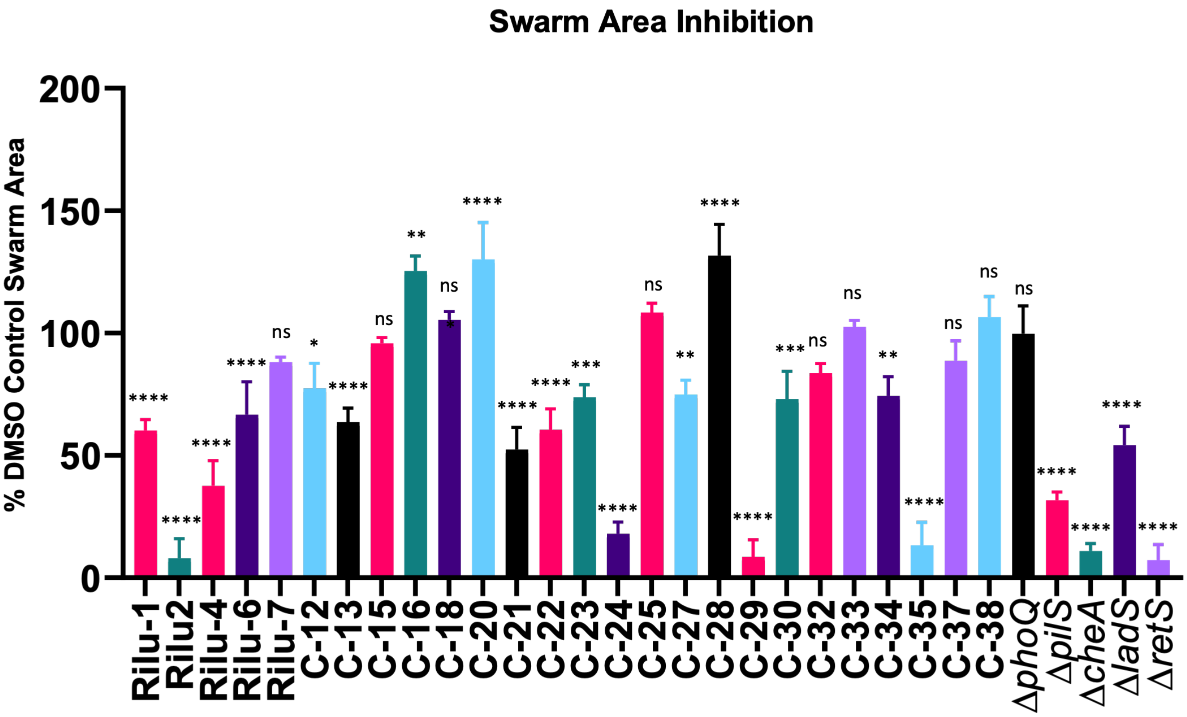


SI Figure 11. **Swarm assay result bar graph:** Graphical representation of swarm area as a percentage of DMSO control (0.4% final DMSO concentration). HK-deletion strains of PA14 included for comparison. Data were compared using a One-way ANOVA followed by Dunnett's multiple comparisons test, which was performed using GraphPad Prism with p-value <0.05 considered statistically significant. **** p-value < 0.0001, *** p-value < 0.0002, ** p-value < 0.0021, * p-value < 0.0332, and ns p-value > 0.1234 relative to DMSO control.

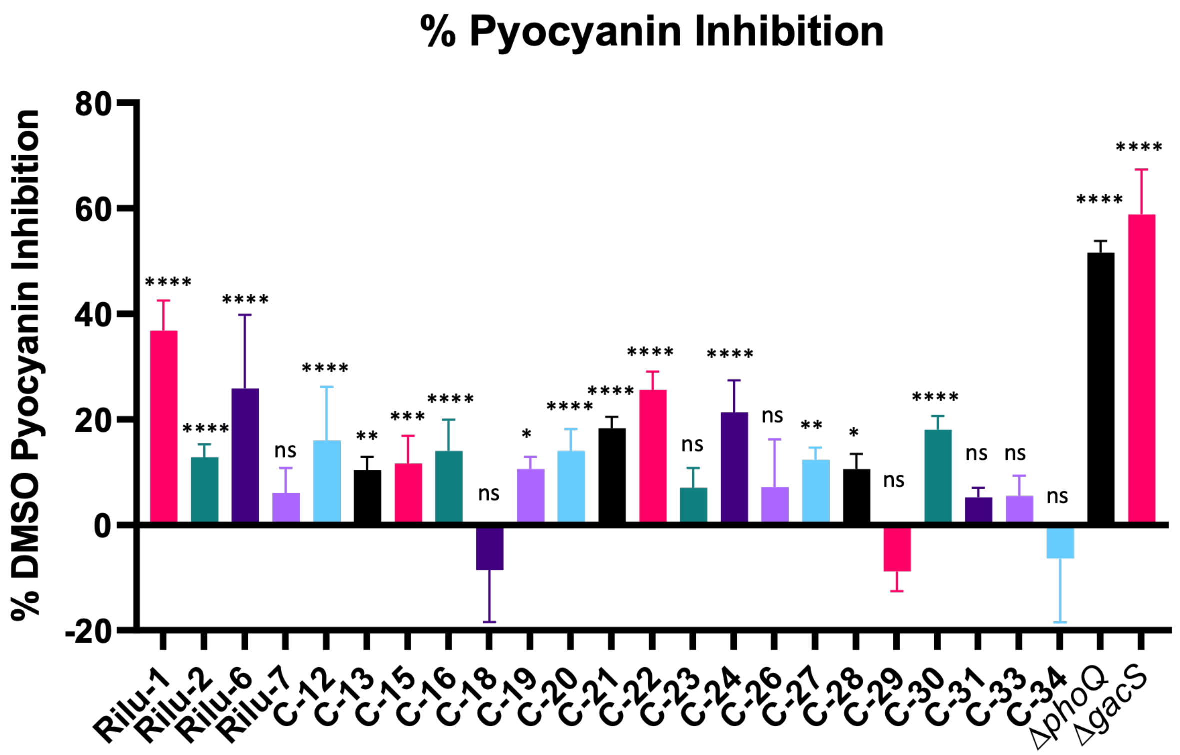


SI Figure 12. **Pyocyanin assay result bar graph:** Graphical representation of the inhibition of pyocyanin production as a percentage of DMSO control (absorbance at 690 nm; 0.8% final DMSO concentration). HK-deletion strains of PA14 included for comparison. Data were compared using a One-way ANOVA followed by Dunnett's multiple comparisons test which was performed using GraphPad Prism with p-value <0.05 considered statistically significant. **** p-value < 0.0001, *** p-value < 0.0002, ** p-value < 0.0021, * p-value < 0.0332, and ns p-value > 0.1234 relative to DMSO control.

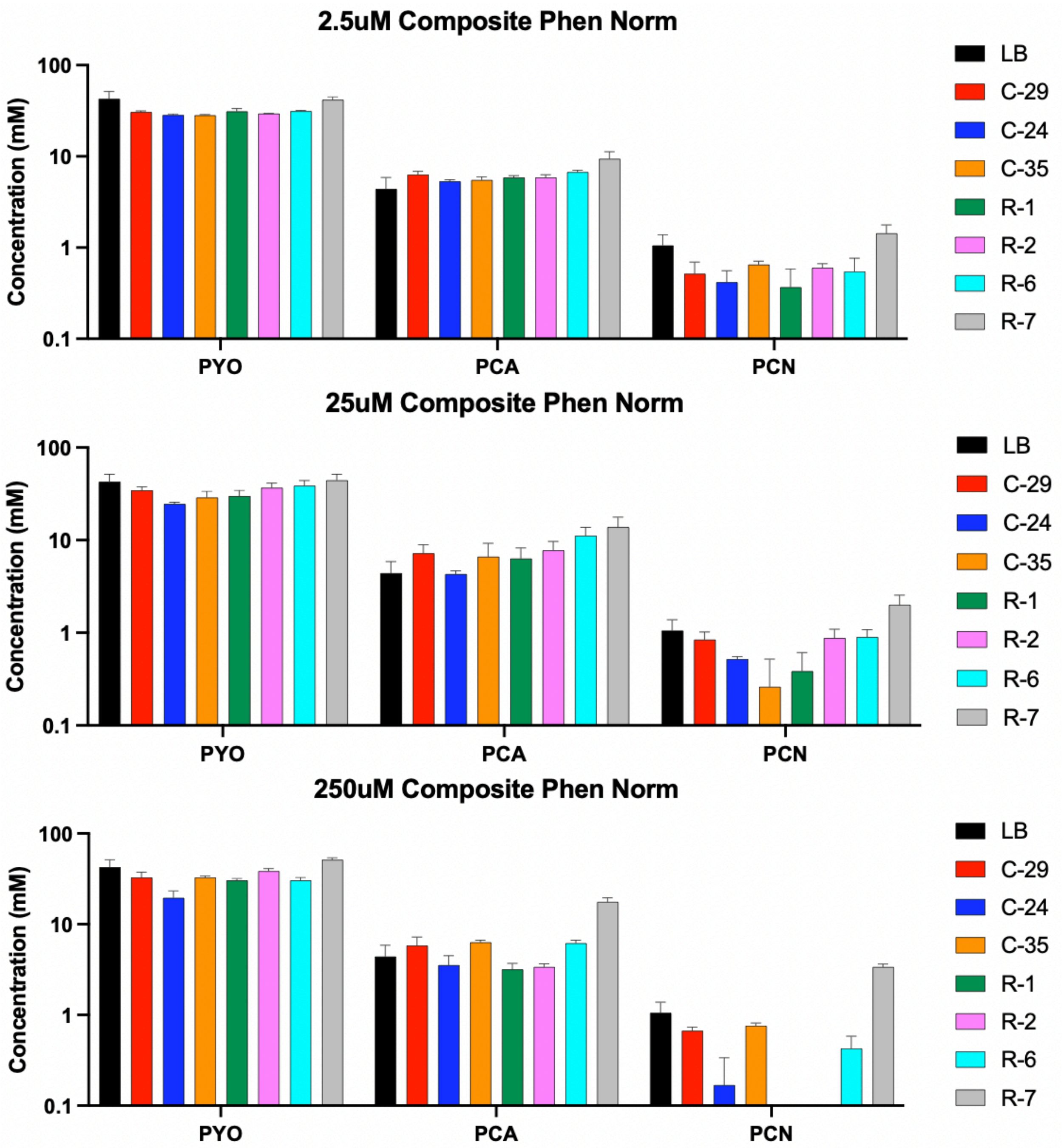


SI Figure 13. **Metabolite quantification after 24 h of inhibitor treatment.** *P. aeruginosa* cultures were treated with 2.5, 25, or 250 μM of inhibitor. Pyocyanin (PYO), phenazine-1-carboxylic acid (PCA), and phenazine-1-carboxamide (PCN) were quantified by HPLC and compared to an untreated control. Concentrations were determined by comparison to purified PYO, PCA, and PCN standards. No noteworthy differences were observed between treatments, with the exception of PCN production in response to inhibitors **C-24**, **R-1**, and **R-2**. Data shown are the mean +/- SD for three biological replicates.

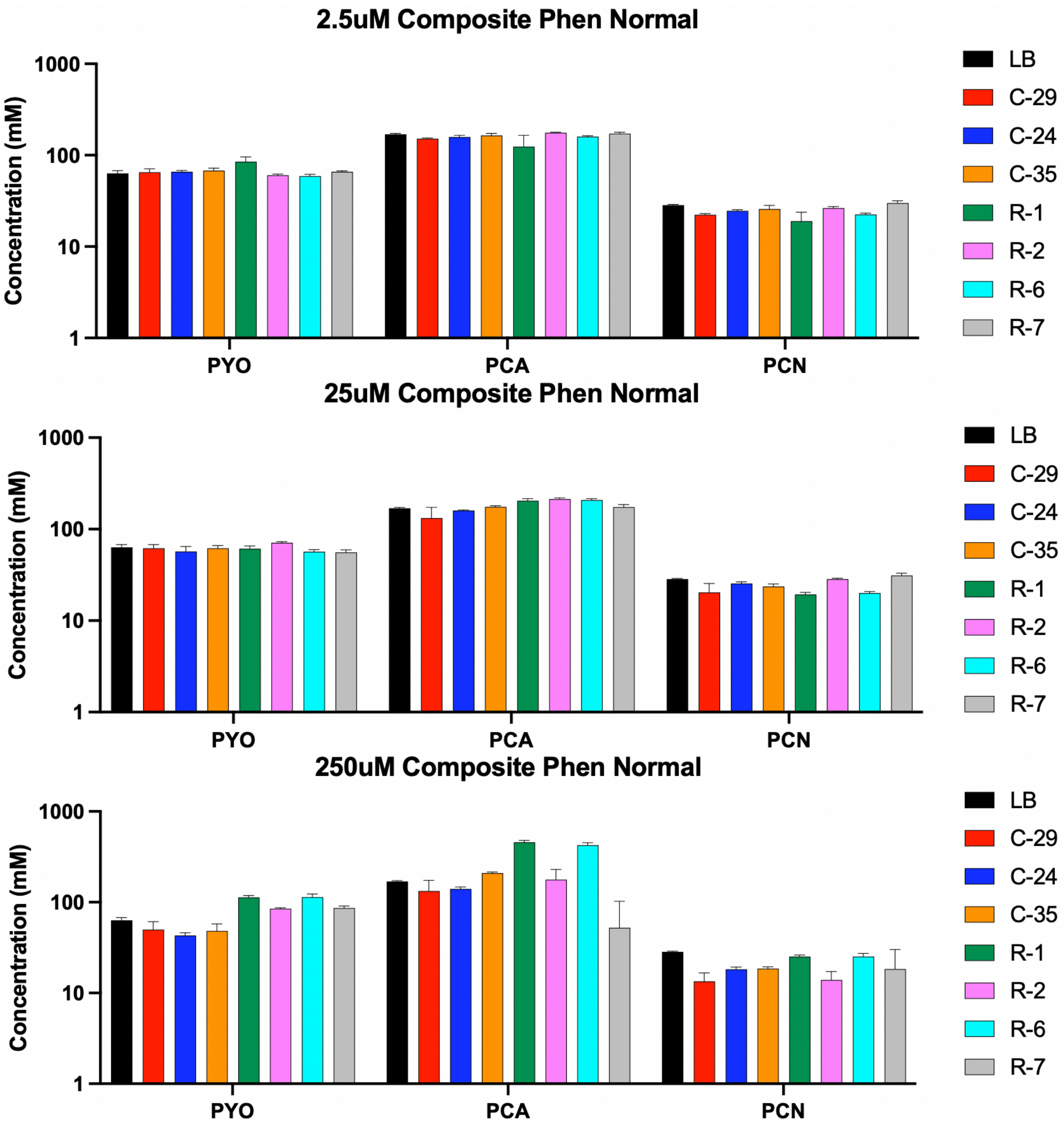


SI Figure 14. **Metabolite quantification after 48 h of inhibitor treatment.** *P. aeruginosa* cultures were treated with 2.5, 25, or 250 µM of inhibitor. Pyocyanin (PYO), phenazine-1-carboxylic acid (PCA), and phenazine-1-carboxamide (PCN) were quantified by HPLC and compared to an untreated control. Concentrations were determined by comparison to purified PYO, PCA, and PCN standards. No significant differences were observed between treatments, Data shown are the mean +/- SD for three biological replicates.

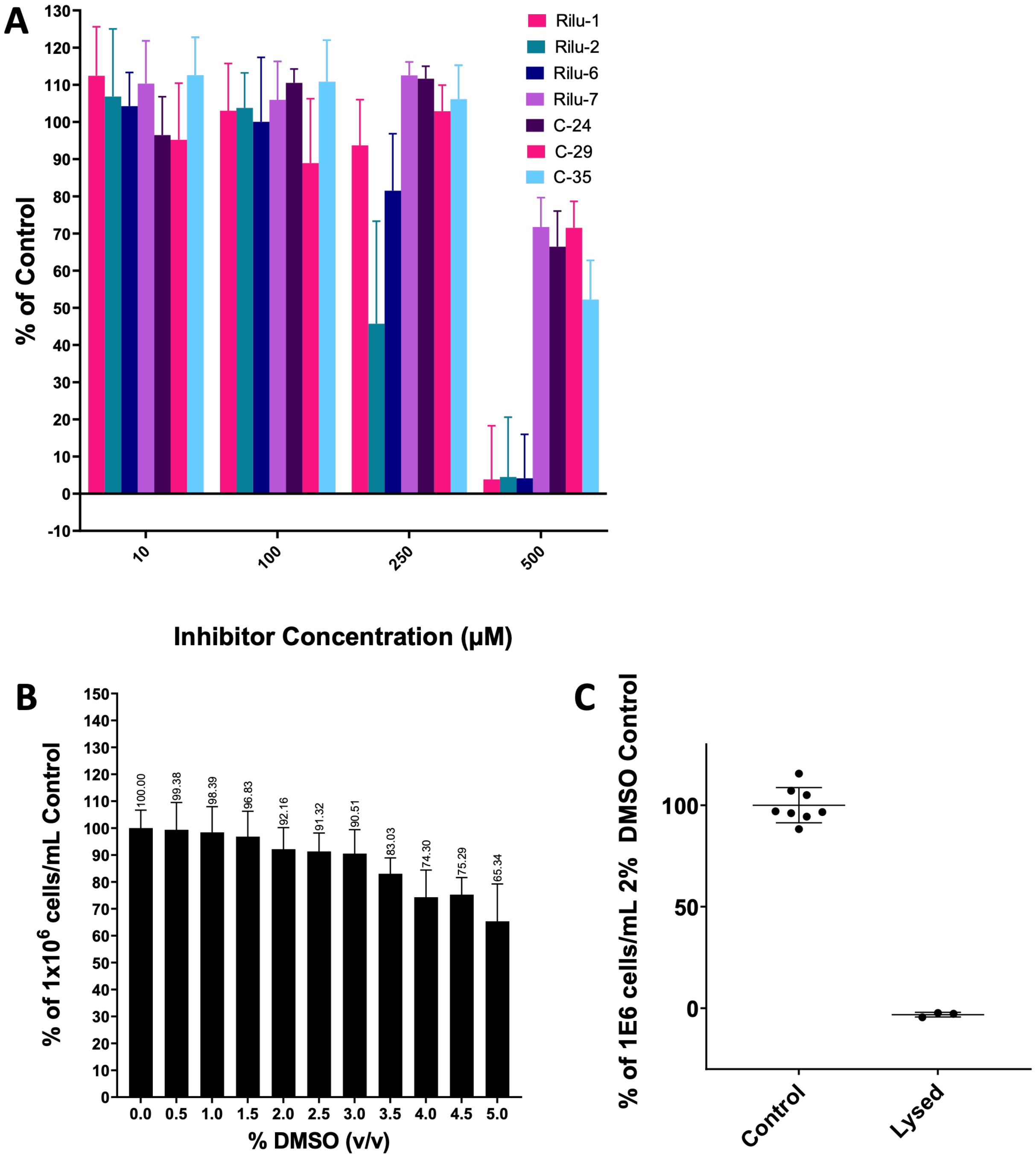


SI Figure 15. **Cytotoxicity of Rilu-series compounds in Hep G2 cells:** a) Cytotoxicity assessment of leads with Hep G2 cells. Percent viability was graphed as percent of a DMSO vehicle control. b) Hep G2 cell viability with 0.5–5.0% (v/v) DMSO, the concentration of DMSO used in testing lead compounds in part a was 2% as it allowed for compound dissolution with minimal effects on viability. c) A 1 h incubation with lysis solution (Promega G182A) served as a positive control of 100% cytotoxicity.

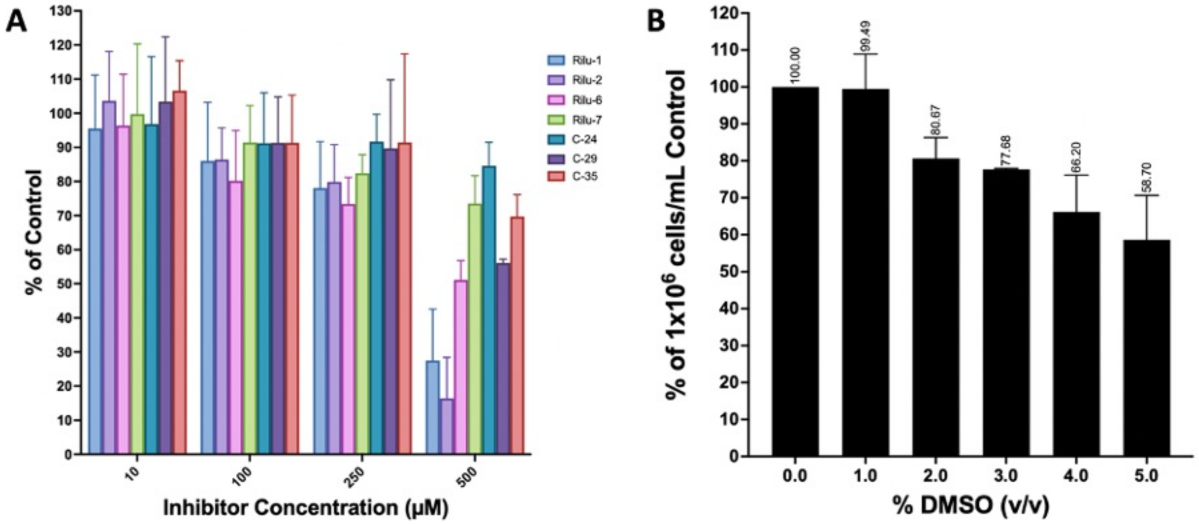


SI Figure 16. **Cytotoxicity of Rilu-series compounds in A549 cells:** A) Cytotoxicity assessment of leads with A549 cells. Percent viability was graphed as percent of a DMSO vehicle control. B) A549 cell viability with 0.5–5.0% (v/v) DMSO, the concentration of DMSO used in testing lead compounds in part a was 2% as it allowed for compound dissolution with minimal effects on viability. A 1 h incubation with lysis solution (Promega G182A) served as a positive control of 100% cytotoxicity.

**Molecule Synthesis**

Abbreviations: Petroleum Ether: PE, ethyl Acetate: EA, methyl Acetate: MA, dichloromethane: DCM, trifluoroacetic acid: TFA, methanol: MeOH, ethanol: EtOH, formic acid: FA, triethylamine: TEA, trifluoroacetic anhydride: TFAA.

Note: $^{13}$C NMR fluorine split quartet for compounds with $CF_3$ groups (C-1 to C-10, C-14, C-18, & C-19) were not observed. Compounds C-11 and C-12 do not have $^{13}$C NMR, & $^{19}$F NMR due to lack of material available.

***Synthesis of C-1***

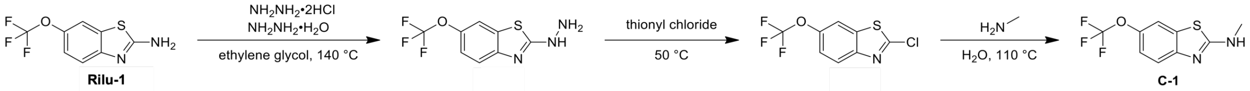


***2-Hydrazinyl-6-(trifluoromethoxy)benzo[d]thiazole***

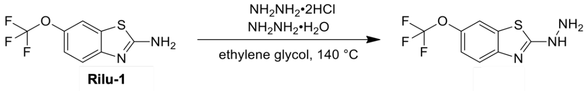


A mixture of 6-(trifluoromethoxy)-1,3-benzothiazol-2-amine (2 g, 8.54 mmol), hydrazine;hydrate (872.46 mg, 17.08 mmol) and hydrazine;dihydrochloride (896.39 mg, 8.54 mmol) in ethylene glycol (40 mL) was stirred at 140 °C for 2 hr. On completion, the reaction was cooled to room temperature and a white solid was precipitated. The solid was collected and washed with 10 ml of water to give the title compound (2.00 g, 93% yield) as a white solid. LCMS $(MS+1)^+$: 250.0. $^1$H NMR (400 MHz, DMSO-$d_6$) δ = 9.20 (br s, 1H), 7.78 (d, $J$ = 1.6 Hz, 1H), 7.35 (d, $J$ = 8.8 Hz, 1H), 7.21 - 7.11 (m, 1H), 5.11 (s, 2H).

***2-Chloro-6-(trifluoromethoxy)benzo[d]thiazole***

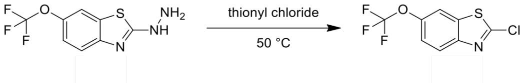


[6-(trifluoromethoxy)-1,3-benzothiazol-2-yl]hydrazine (1.50 g, 6.02 mmol) was added during 1 hr to thionyl chloride (2.86 g, 24.08 mmol) at 50 °C. The mixture was stirred at 50 °C for 1 hr. On completion, the reaction was poured into 10 mL of ice-water and a yellow solid was precipitated. The solid was collected and dissolved in 50 mL of PE. The mixture was washed with water (3 x 20 mL), dried over $Na_2SO_4$, filtered and concentrated in vacuo to give the title compound (1.10 g,

72% yield) as a yellow solid. LCMS (MS+1)$^+$: 254.0. $^1$H NMR (400 MHz, $CDCl_3$) δ = 7.97 (d, *J* = 8.8 Hz, 1H), 7.67 (d, *J* = 1.2 Hz, 1H), 7.37 (dd, *J* = 1.2, 8.8 Hz, 1H).

***N-methyl-6-(trifluoromethoxy)benzo[d]thiazol-2-amine (C-1)***

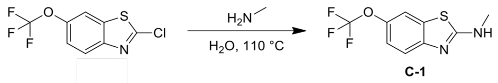


A mixture of 2-chloro-6-(trifluoromethoxy)-1,3-benzothiazole (100 mg, 394 μmol) in $MeNH_2$ (30% aqueous solution, 3 mL) was stirred at 110 °C for 12 hr in a sealed tube. On completion, the reaction was concentrated. The residue was purified by reverse phase to give the title compound (14.23 mg, 14.5% yield) as off-white solid. LCMS (MS+1)$^+$: 249.1. $^1$H NMR (400 MHz, DMSO-*d*$_6$) δ = 8.09 (br s, 1H), 7.78 (d, *J* = 1.6 Hz, 1H), 7.42 (d, *J* = 8.8 Hz, 1H), 7.19 (dd, *J* = 1.6, 8.8 Hz, 1H), 2.94 (d, *J* = 4.4 Hz, 3H). $^{13}$C NMR (101 MHz, DMSO-*d*$_6$) δ 168.42, 152.32, 142.54, 131.89, 119.52, 118.66, 114.97, 30.98 ($\underline{C}F_3$ not observed). $^{19}$F NMR (376 MHz, DMSO-*d*$_6$) δ -57.12.

***Synthesis of C-2***

***N,N-dimethyl-6-(trifluoromethoxy)benzo[d]thiazol-2-amine (C-2)***

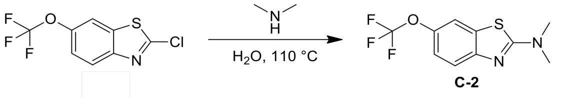


2-chloro-6-(trifluoromethoxy)-1,3-benzothiazole (0.200 g, 788 μmol, from **C-1**) in N-methylmethanamine (16.1 g, 33% aqueous solution) was stirred at 110 °C for 12 hr. On completion, the reaction was concentrated. The residue was purified by column chromatography to give the title compound (185 mg, 89% yield) as a off-white solid. LCMS (MS+1)$^+$: 263.0. $^1$H NMR (400 MHz, DMSO-*d*$_6$) δ = 7.89 (s, 1H), 7.48 (d, *J* = 8.8 Hz, 1H), 7.24 (d, *J* = 8.8 Hz, 1H), 3.15 (s, 6H). $^{13}$C NMR (101 MHz, DMSO-*d*$_6$) δ 169.60, 152.60, 142.35, 142.33, 132.10, 122.74 (q, *J* = 254 Hz), 119.95, 118.95, 115.09. $^{19}$F NMR (376 MHz, DMSO-*d*$_6$) δ -57.17.

***Synthesis of C-3***

***N-benzyl-6-(trifluoromethoxy)benzo[d]thiazol-2-amine (C-3)***

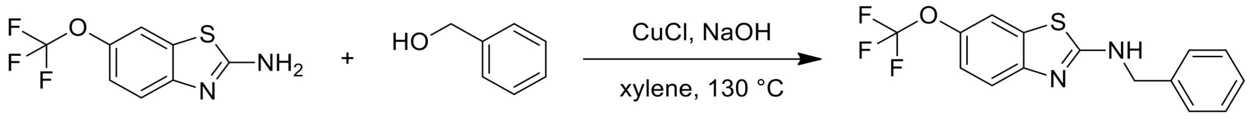

Rilu-1 C-3

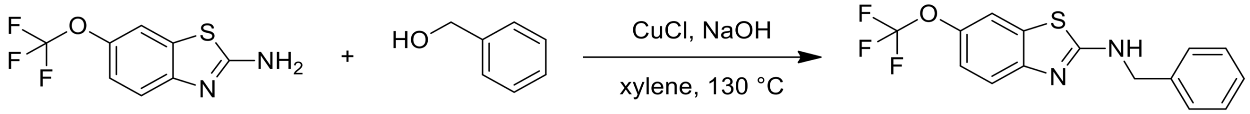

Rilu-1 C-3

A mixture of 6-(trifluoromethoxy)-1,3-benzothiazol-2-amine (100 mg, 427 µmol), phenylmethanol (55.4 mg, 512 µmol), CuCl (4.23 mg, 42.70 µmol) and NaOH (3.42 mg, 85.4 µmol) in xylene (3 mL) was stirred at 130 °C for 12 hr. On completion, the reaction was concentrated in vacuo. The residue was diluted with water (50 mL) and extracted with EA (3 x 20 mL). The combined organic layer was dried over $Na_2SO_4$, filtered and concentrated in vacuo. The residue was purified by reverse phase **t**o give the title compound (42.0 mg, 30% yield) as a white solid. LCMS $(MS+1)^+$: 325.1. $^1H$ NMR (400 MHz, DMSO-$d_6$) δ = 8.65 (t, $J$ = 6.0 Hz, 1H), 7.79 (s, 1H), 7.42 (d, $J$ = 8.8 Hz, 1H), 7.40 - 7.30 (m, 4H), 7.30 - 7.23 (m, 1H), 7.22 - 7.13 (m, 1H), 4.60 (d, $J$ = 6.0 Hz, 2H). $^{13}C$ NMR (101 MHz, DMSO-$d_6$) δ 167.86, 152.09, 142.69, 139.14, 132.00, 128.89, 127.87, 127.57, 120.93 (q, $J$ = 254 Hz), 119.54, 118.84, 115.00, 47.71. $^{19}F$ NMR (376 MHz, DMSO-$d_6$) δ -57.16.

***Synthesis of C-4***

***N-phenyl-6-(trifluoromethoxy)-1,3-benzothiazol-2-amine (C-4)***

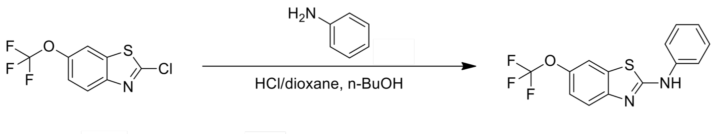

C-4

To a solution of 2-chloro-6-(trifluoromethoxy)-1,3-benzothiazole (100 mg, 394 µmol, **From C-1**) in n-BuOH (2 mL) was added a catalyst amount of HCl/dioxane (4 M, 20 uL) and aniline (44.0 mg, 473 µmol). The mixture was stirred at 90 °C for 12 hr. On completion, the reaction mixture was concentrated in vacuo to get a residue. The residue was diluted with ice water (10 mL) and extracted with DCM (3 X 6 mL). The organic layer was collected and dried over anhydrous $Na_2SO_4$, filtered and concentrated in vacuo. The residue was purified by column chromatography

(PE:EA = 6:1) to give the title compound (103 mg, 82% yield) as a white solid. LCMS $(M+1)^+$: 311.1. $^1H$ NMR (400MHz, $CDCl_3$) δ = 8.75 (br s, 1H), 7.57 - 7.50 (m, 4H), 7.48 - 7.42 (m, 2H), 7.27 - 7.19 (m, 2H). $^{13}C$ NMR (101 MHz, DMSO-$d_6$) δ 163.36, 151.69, 143.61, 140.78, 131.65, 129.52, 122.87, 120.72 (q, $J$ = 254 Hz), 120.18, 119.87, 118.42, 115.00. $^{19}F$ NMR (376 MHz, DMSO-$d_6$) δ -57.10.

***Synthesis of C-5***

***N,N-diethyl-6-(trifluoromethoxy)benzo[d]thiazol-2-amine (C-5)***

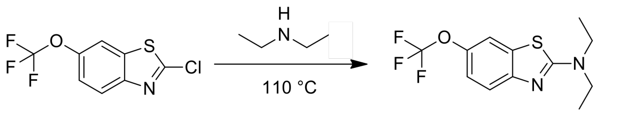


**C-5**

2-chloro-6-(trifluoromethoxy)-1,3-benzothiazole (200 mg, 788.55 μmol, from **C-1**) in N-ethylethanamine (5 mL) was stirred at 110 °C for 12 hr. On completion, the reaction was concentrated in vacuo. The residue was purified by column chromatography (PE:EA = 10:1) to give the title compound (218 mg, 95% yield) as a brown solid. LCMS $(MS+1)^+$: 291.0. $^1H$ NMR (400 MHz, $CDCl_3$) δ = 7.48 (d, $J$ = 8.8 Hz, 1H), 7.45 (d, $J$ = 1.6 Hz, 1H), 7.14 (dd, $J$ = 1.6, 8.8 Hz, 1H), 3.58 (q, $J$ = 7.2 Hz, 4H), 1.30 (t, $J$ = 7.2 Hz, 6H). $^{13}C$ NMR (101 MHz, DMSO-$d_6$) δ 167.99, 152.62, 142.29, 131.59, 120.74, 119.86, 118.79, 115.02, 45.65, 13.05. $^{19}F$ NMR (376 MHz, DMSO-$d_6$) δ -57.22.

***Synthesis of C-6***

***N-(6-(trifluoromethoxy)benzo[d]thiazol-2-yl)acetamide (C-6)***

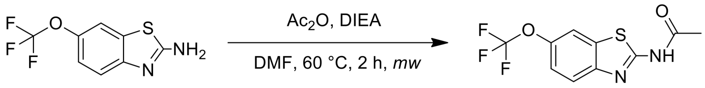


**Rilu-1** **C-6**

A mixture of 6-(trifluoromethoxy)-1,3-benzothiazol-2-amine (100 mg, 427 μmol), $Ac_2O$ (95.9 mg, 939 μmol) and DIEA (165 mg, 1.28 mmol) in DMF (4 mL) was stirred at 60 °C for 2 hr under microwave condition. On completion, the reaction was poured into 10 ml of saturated $NaHCO_3$ aqueous and a white solid was precipitated. The solid was collected and washed with 5 mL of water to give the title compound (71.73 mg, 58% yield) as a brown solid. LCMS $(MS+1)^+$: 277.0.

$^{1}H$ NMR (400 MHz, DMSO-$d_6$) δ = 8.05 (s, 1H), 7.76 (d, $J$ = 8.4 Hz, 1H), 7.38 (d, $J$ = 8.4 Hz, 1H), 2.18 (s, 3H). $^{13}C$ NMR (101 MHz, DMSO-$d_6$) δ 170.47, 160.38, 148.13, 144.36, 133.16, 121.77, 120.69 (q, $J$ = 254 Hz), 120.12, 115.35, 23.38. $^{19}F$ NMR (376 MHz, DMSO-$d_6$) δ -57.03.

***<u>Synthesis of C-7</u>***

***1. Synthetic Scheme:***

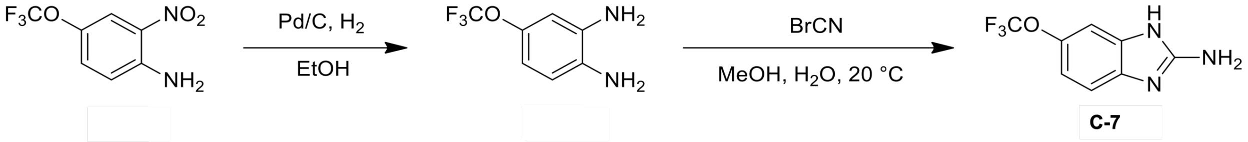


***<u>4-(Trifluoromethoxy)benzene-1,2-diamine</u>***

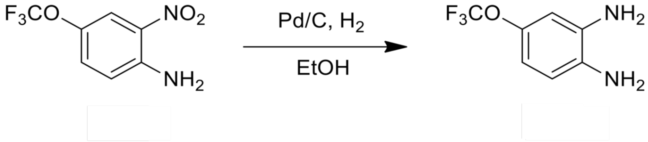


2-nitro-4-(trifluoromethoxy)aniline (2.00 g, 9.00 mmol) and Pd/C (0.2 g, 10% purity) in EtOH (25 mL) was stirred at 20 °C for 12 hr under $H_2$ (15 psi) atmosphere. On completion, the reaction was filtered and the filtrate was concentrated in vacuo to give the title compound (1.5 g, 80% yield) as red oil which was used for next step directly. LCMS $(MS+1)^+$: 193.1.

***<u>6-(Trifluoromethoxy)-1H-benzo[d]imidazol-2-amine (C-7)</u>***

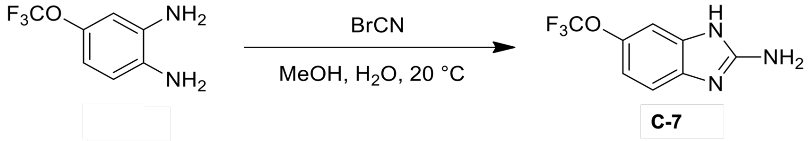


BrCN (1.10 g, 10.4 mmol) was added to a mixture of 4-(trifluoromethoxy)benzene-1,2-diamine (500 mg, 2.60 mmol) in MeOH (6 mL) and $H_2O$ (6 mL). The reaction mixture was stirred at 20 °C for 12 hr. On completion, the reaction was adjusted pH to 7 with 2 N NaOH aqueous. The mixture was concentrated in vacuo to remove MeOH and then extracted with EA (3 x 10 mL). The combined organic layer was dried over $Na_2SO_4$, filtered and concentrated in vacuo. The residue was purified by Prep-HPLC (column: Phenomenex Synergi C18 150mm*25mm*10μm; mobile phase: [water(0.225%FA)-ACN];B%: 8%-32%,8 min) to give the title compound (271 mg, 48% yield) as a white solid. LCMS $(MS+1)^+$: 218.0. $^{1}H$ NMR (400 MHz, DMSO-$d_6$) δ = 8.15 (s, 1H),

7.11 (d, $J$ = 8.4 Hz, 1H), 7.04 (d, $J$ = 1.2 Hz, 1H), 6.81 (dd, $J$ = 1.2, 8.4 Hz, 1H), 6.45 (s, 2H). $^{13}$C NMR (101 MHz, DMSO-$d_6$) δ 163.97, 142.54, 140.12, 136.47, 120.89 (q, $J$ = 253 Hz), 112.51, 111.28, 105.68. $^{19}$F NMR (376 MHz, DMSO-$d_6$) δ -56.99.

***Synthesis of C-8***

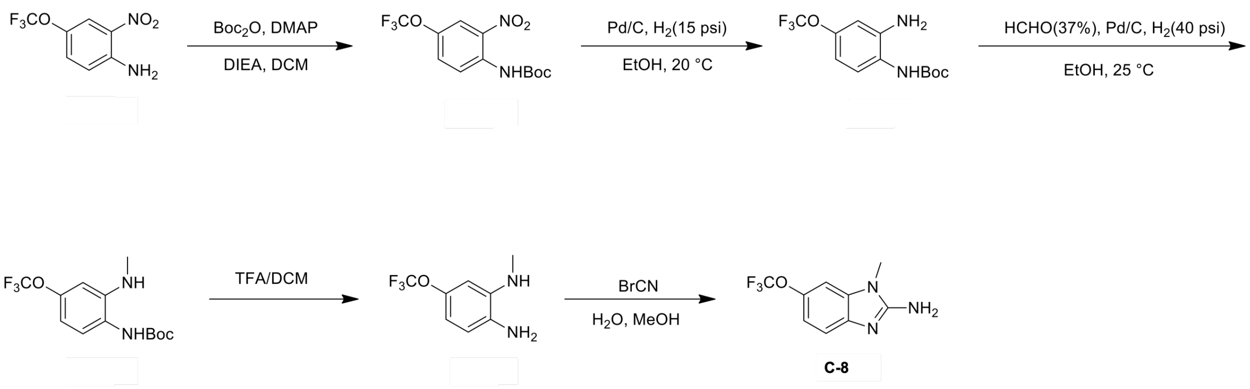


***Tert-butyl (2-nitro-4-(trifluoromethoxy)phenyl)carbamate***

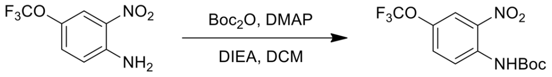


To a solution of 2-nitro-4-(trifluoromethoxy)aniline (10.0 g, 45.0 mmol), DIEA (5.82 g, 45.0 mmol,) and DMAP (550 mg, 4.50 mmol) in DCM (200 mL) was added $Boc_2O$ (14.7 g, 67.5 mmol) for 30 min in portions. Then the mixture was stirred at 20 °C for 12 hr. On completion, the reaction was washed with water, 5% citric acid aqueous and brine. The organic layer was concentrated in vacuo to give the title compound (6 g, 50% yield) as a white solid. $^{1}$H NMR (400MHz, $CDCl_3$) δ = 9.64 (br s, 1H), 8.68 (d, $J$ = 9.2 Hz, 1H), 8.09 (d, $J$ = 2.4 Hz, 1H), 7.49 (dd, $J$ = 2.4, 9.2 Hz, 1H), 1.55 (s, 9H).

***Tert-butyl (2-amino-4-(trifluoromethoxy)phenyl)carbamate***

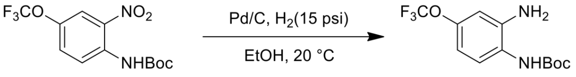


A mixture of *tert*-butyl N-[2-nitro-4-(trifluoromethoxy)phenyl]carbamate (5.00 g, 15.5 mmol) and Pd/C (0.5 g, 10% purity) in EtOH (50 mL) was stirred at 20 °C for 12 hr under $H_2$ (15 psi). On completion, the reaction was filtered and the filtrate was concentrated in vacuo to give the title compound (4.4 g, 90% yield) as a white solid. $^{1}$H NMR (400MHz, $CDCl_3$) δ = 7.24 (d, $J$ = 9.2 Hz, 1H), 6.68 - 6.58 (m, 2H), 6.13 (br s, 1H), 3.91 (br s, 2H), 1.52 (s, 9H).

***Tert-butyl (2-(methylamino)-4-(trifluoromethoxy)phenyl)carbamate***

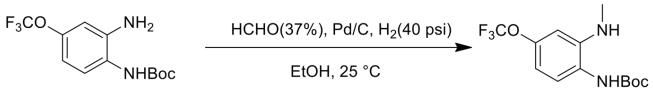


To a mixture of *tert*-butyl N-[2-amino-4-(trifluoromethoxy)phenyl]carbamate (500 mg, 1.71 mmol), Pd/C (20 mg, 10% purity) in EtOH (5 mL) was added Formaldehyde (37%, 138 mg, 1.71 mmol), then the mixture was stirred at 20 °C for 12 hr under $H_2$ (40 psi). On completion, the reaction was filtered and the filtrate was concentrated in vacuo. The residue was purified by column chromatography (PE:EA = 50:1) to give the title compound (350 mg, 66% yield) as a white solid. LCMS $(MS+1)^+$: 307.0. $^1H$ NMR (400MHz, $CDCl_3$) δ = 7.22 (d, $J$ = 8.8 Hz, 1H), 6.57 (d, $J$ = 8.8 Hz, 1H), 6.51 (s, 1H), 5.94 (br s, 1H), 2.85 (s, 3H), 1.51 (s, 9H).

***N1-methyl-5-(trifluoromethoxy)benzene-1,2-diamine***

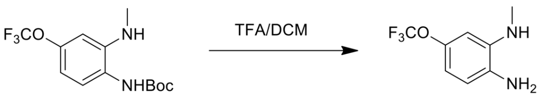


To a solution of *tert*-butyl N-[2-(methylamino)-4-(trifluoromethoxy)phenyl]carbamate (200 mg, 653 μmol) in DCM (4 mL) was added TFA (2 mL). Then the mixture was stirred at 25 °C for 30 min. On completion, the reaction was concentrated in vacuo, the residue was diluted with water (50 mL), adjusted pH=10 with ammonium aqueous. The mixture was extracted with EA (3 x 30 mL). The combined organic layer was washed with brine, dried over $Na_2SO_4$ and concentrated in vacuo to give the title compound (120 mg, 80% yield) as a brown oil. $^1H$ NMR (400MHz, DMSO-$d_6$) δ = 6.53 (d, $J$ = 8.4 Hz, 1H), 6.39 - 6.29 (m, 1H), 6.24 (d, $J$ = 2.0 Hz, 1H), 4.80 (br s, 3H), 2.70 (s, 3H).

***1-Methyl-6-(trifluoromethoxy)-1H-benzo[d]imidazol-2-amine***

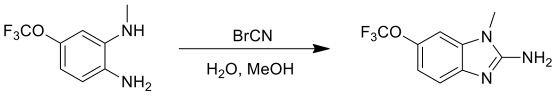


**C-8**

To a mixture of *N*2-methyl-4-(trifluoromethoxy)benzene-1,2-diamine (120 mg, 582 μmol) in $H_2O$ (3 mL) and MeOH (3 mL) was added BrCN (61.6 mg, 582 μmol). Then the mixture was stirred at 25 °C for 12 hr. On completion, the reaction was adjusted pH=9 with 2N NaOH aqueous and then concentrated in vacuo to remove MeOH. The residue was extracted with EA (3 x 20 mL). The combined organic layer was washed with brine, dried over $Na_2SO_4$ and concentrated in vacuo. The residue was triturated with PE (5 mL) to give the title compound (88.9 mg, 65% yield) as a brown solid. LCMS $(MS+1)^+$: 231.9. $^1H$ NMR (400MHz, DMSO-$d_6$) δ = 7.18 (d, $J$ = 1.2 Hz, 1H), 7.13 (d, $J$ = 8.4 Hz, 1H), 6.92 - 6.85 (m, 1H), 6.56 (s, 2H), 3.51 (s, 3H). $^{13}C$ NMR (101 MHz, DMSO-$d_6$) δ 157.26, 142.13, 141.38, 135.52, 120.92 (q, $J$ = 254 Hz), 114.91, 113.80, 101.83, 29.09. $^{19}F$ NMR (376 MHz, DMSO-$d_6$) δ -57.00.

***Synthesis of C-9***

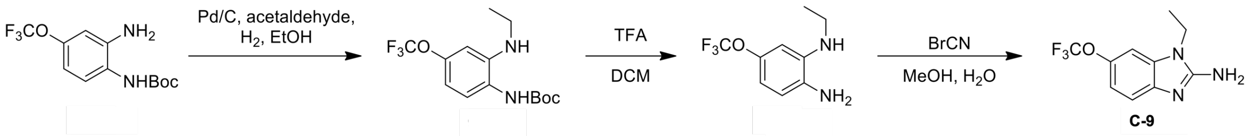


***Tert-butyl (2-(ethylamino)-4-(trifluoromethoxy)phenyl)carbamate***

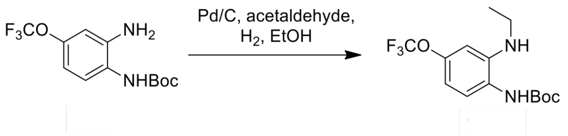


To a mixture of *tert*-butyl N-[2-amino-4-(trifluoromethoxy)phenyl]carbamate (500 mg, 1.71 mmol), Pd/C (20 mg, 10% purity) in EtOH (5 mL) was added acetaldehyde (40%, 188 mg, 1.71 mmol). The mixture was stirred at 20 °C for 12 hr under $H_2$ (40 psi) atmosphere. On completion, the reaction was filtered and the filtrate was concentrated in vacuo. The residue was purified by column chromatography (PE:EA = 50:1) to give the title compound (350 mg, 64% yield) as a white solid. $^1H$ NMR (400MHz, $CDCl_3$) δ = 7.23 (d, $J$ = 8.4 Hz, 1H), 6.56 (d, $J$ = 8.4 Hz, 1H), 6.52 (s, 1H), 5.94 (br s, 1H), 3.93 (br s, 1H), 3.16 - 3.10 (m, 2H), 1.52 (s, 9H), 1.30 (t, $J$ = 7.2 Hz, 3H).

***$N^1$-ethyl-5-(trifluoromethoxy)benzene-1,2-diamine***

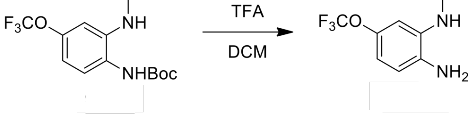

To a solution of *tert*-butyl N-[2-(ethylamino)-4-(trifluoromethoxy)phenyl]carbamate (350 mg, 1.09 mmol) in DCM (6 mL) was added TFA (2 mL). Then the mixture was stirred at 20 °C for 3 hr. On completion, the reaction was concentrated in vacuo. The residue was adjusted pH to 10 with ammonoim water, extracted with EA (3 x 20 mL). The combined organic layer was washed with brine, dried over $Na_2SO_4$, filtered and concentrated to give the title compound (250 mg, 90% yield) as a brown oil which was used for next step directly.

***1-Ethyl-6-(trifluoromethoxy)-1H-benzo[d]imidazol-2-amine (C-9)***

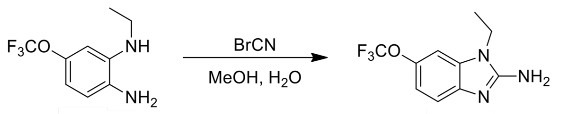

**C-9**

To a mixture of $N^2$-ethyl-4-(trifluoromethoxy)benzene-1,2-diamine (250 mg, 1.14 mmol) in MeOH (3 mL) and $H_2O$ (3 mL) was added BrCN (120 mg, 1.14 mmol). Then the mixture was stirred at 25 °C for 12 hr. On completion, the reaction was adjusted pH to 10 with ammonium water, concentrated in vacuo to remove MeOH. The residue was extracted with EA (3 x 20 mL). The combined organic layer was washed with brine, dried over $Na_2SO_4$, filtered and concentrated in vacuo. The residue was purified by column chromatography (PE:EA = 5:1) to give the title compound (104 mg, 37% yield) as a brown solid. LCMS $(MS+1)^+$: 245.9. $^1H$ NMR (400 MHz, DMSO-$d_6$) δ = 7.25 (s, 1H), 7.16 (d, $J$ = 8.4 Hz, 1H), 6.91 (d, $J$ = 8.4 Hz, 1H), 6.76 (br s, 2H), 4.07 - 4.01 (m, 2H), 1.19 (t, $J$ = 7.2 Hz, 3H). $^{13}C$ NMR (101 MHz, DMSO-$d_6$) δ 155.99, 141.67, 134.11, 120.89 (q, $J$ = 254 Hz), 114.76, 114.09, 102.10, 40.55, 37.03, 14.16. $^{19}F$ NMR (376 MHz, DMSO-$d_6$) δ -57.04.

***Synthesis of C-10***

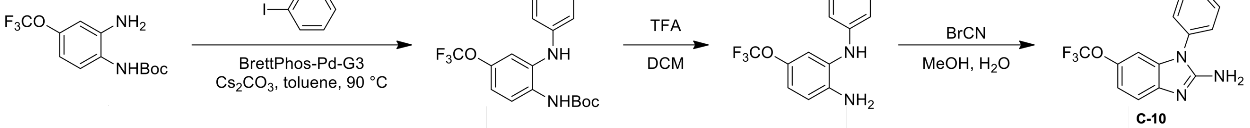

**C-10**

***Tert-butyl (2-(phenylamino)-4-(trifluoromethoxy)phenyl)carbamate***

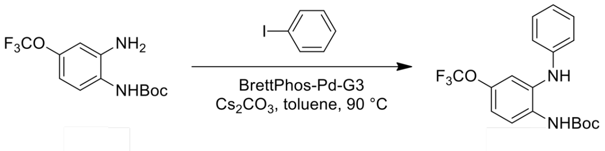

A mixture of *tert*-butyl N-[2-amino-4-(trifluoromethoxy)phenyl]carbamate (500 mg, 1.71 mmol, from **C-22**), iodobenzene (418 mg, 2.05 mmol), BrettPhos-Pd-G3 (155 mg, 171 µmol) and $Cs_2CO_3$ (1.11 g, 3.42 mmol) in toluene (20 mL) was stirred at 90 °C for 3 hr. On completion, the reaction was filtered and the filtrate was concentrated in vacuo. The residue was purified by column chromatography (PE:EA = 50:1) to give the title compound (350 mg, 55% yield) as a white solid. $^1$H NMR (400MHz, $CDCl_3$) δ = 7.76 (d, *J* = 8.8 Hz, 1H), 7.31 - 7.27 (m, 2H), 7.13 (d, *J* = 2.4 Hz, 1H), 7.01 - 6.92 (m, 2H), 6.89 (d, *J* = 7.6 Hz, 2H), 6.71 (br s, 1H), 1.53 (s, 9H).

***$N^1$-phenyl-5-(trifluoromethoxy)benzene-1,2-diamine***

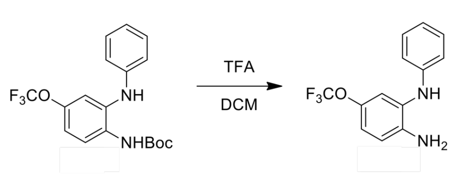


To a solution of *tert*-butyl N-[2-anilino-4-(trifluoromethoxy)phenyl]carbamate (300 mg, 814 µmol) in DCM (6 mL) was added TFA (3 mL). Then the mixture was stirred at 30 °C for 1 hr. On completion, the reaction was concentrated in vacuo. The residue was diluted with water (50 ml), adjusted pH to 10 with 28% ammonium water. The mixture was extracted with EA (3 x 30 mL). The combined organic layer was washed with brine, dried over $Na_2SO_4$ and concentrated in vacuo to give the title compound (200 mg, 92% yield) as a brown oil. $^1$H NMR (400MHz, DMSO-$d_6$) δ = 7.26 (s, 1H), 7.19 (dd, *J*=7.2, 8.4 Hz, 2H), 6.91 (s, 1H), 6.82 (dd, *J* = 1.2, 8.4 Hz, 2H), 6.79 - 6.72 (m, 3H), 4.98 (br s, 2H).

***1-Phenyl-6-(trifluoromethoxy)-1H-benzo[d]imidazol-2-amine (C-10)***

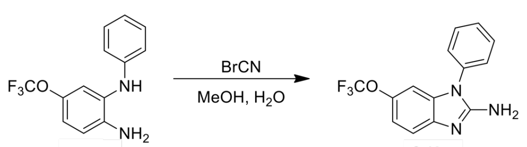

C-10

BrCN (78.9 mg, 745 µmol) was added to a solution of $N^2$-phenyl-4-(trifluoromethoxy)benzene - 1,2-diamine (200 mg, 745 µmol) in MeOH (3 mL) and $H_2O$ (3 mL). Then the mixture was stirred at 25 °C for 12 hr. On completion, the reaction was adjusted pH to 9 with 2 N NaOH aqueous and then concentrated in vacuo to remove MeOH. The residue was extracted with EA (3 x 20 mL). The combined organic layer was washed with brine, dried over $Na_2SO_4$, filtered and concentrated in vacuo. The residue was purified by column chromatography (PE:ethyl acetate = 5:1) to give the title compound (94.0 mg, 43% yield) as a brown solid. LCMS $(MS+1)^+$: 293.9. $^1$H NMR (400MHz,

DMSO-$d_6$) δ = 7.67 - 7.60 (m, 2H), 7.57 - 7.47 (m, 3H), 7.25 (d, *J* = 8.4 Hz, 1H), 7.02 - 6.95 (m, 1H), 6.73 (d, *J* = 1.2 Hz, 1H), 6.46 (s, 2H). $^{13}$C NMR (101 MHz, DMSO-$d_6$) δ 156.22, 142.53, 141.67, 135.53, 134.80, 130.74, 129.08, 127.15, 120.81 (q, *J* = 254 Hz), 115.56, 114.85, 101.69. $^{19}$F NMR (376 MHz, DMSO-$d_6$) δ -57.15.

***Synthesis of C-11***

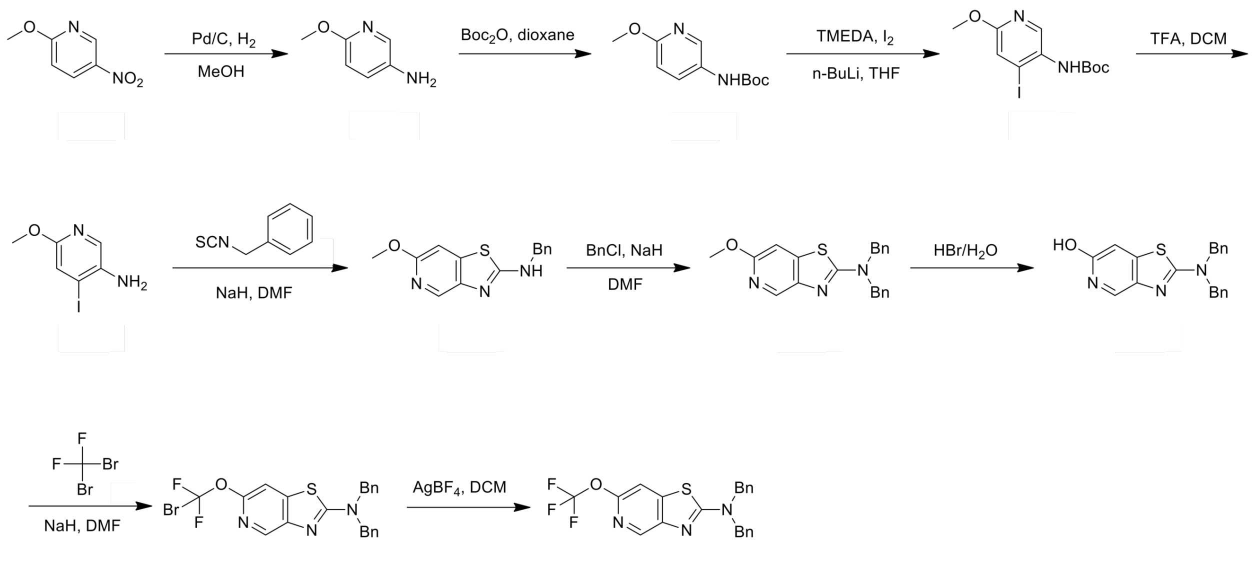


**C-11**

***6-Methoxypyridin-3-amine***

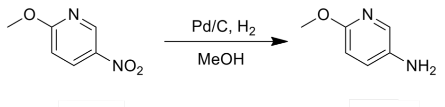


A mixture of 2-methoxy-5-nitro-pyridine (10.0 g, 64.9 mmol), Pd/C (200 mg, 10% purity) in MeOH (100 mL) was stirred at 25 °C for 12 hr under $H_2$ (15 psi) atmosphere. On completion, the reaction mixture was filtered and the filtrate was concentrated in vacuo to give the title compound (8.00 g, 99% yield) as colorless oil. LCMS $(M+1)^+$: 125.2.

***Tert-butyl (6-methoxypyridin-3-yl)carbamate***

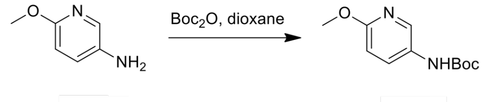

A mixture of 6-methoxypyridin-3-amine (5.00 g, 40.3 mmol) and $Boc_2O$ (11.4 g, 52.4 mmol) in dioxane (40 mL) was stirred at 75 °C for 16 hr. On completion, the reaction mixture was concentrated in vacuo, and the residue was diluted with EA (200 mL) and washed with water (150 mL). The organic layer was dried over sodium sulfate, filtered and evaporated in vacuo. The residue was purified by silica gel chromatography (PE:EA= 1:1) to give the title compound (8.00 g, 89% yield) as a white solid. $^1$H NMR (400 MHz, $CDCl_3$) δ = 8.00 (d, *J* = 2.8 Hz, 1H), 7.95 (br s, 1H), 6.71 (d, *J* = 8.8 Hz, 1H), 6.36 (s, 1H), 3.90 (s, 3H), 1.51 (s, 9H).

***Tert-butyl (4-iodo-6-methoxypyridin-3-yl)carbamate***

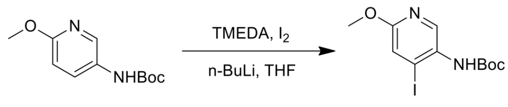


To a suspension of the tert-butyl N-(6-methoxy-3-pyridyl)carbamate (16.2 g, 72.2 mmol) and TMEDA (21.0 g, 181 mmol) in THF (300 mL) was added n-BuLi (2.5 M, 86.7 mL) slowly at -70 °C. The reaction mixture was stirred at -70 °C for 15 mins. After stirred for 2 hr at -10 °C, a white solid was precipitated slowly. Then the reaction mixture was cooled to -70 °C, and a solution of $I_2$ (45.8 g, 181 mmol) in THF (100 mL) was added. The reaction mixture was stirred at 25 °C for 16 hr. On completion, the excess iodine was quenched by the saturated potassium thiosulfate solution, and then extracted with EA (3 X 200 mL). The combined layer was dried over $Na_2SO_4$ and filtered and concentrated in vacuo to give a residue. The residue was purified by silica gel chromatography (PE:EA=5:1) to give the title compound (12 g, 45.1% yield) as a yellow solid. $^1$H NMR (400 MHz, $CDCl_3$) δ = 8.45 (s, 1H), 7.23 (s, 1H), 6.29 (s, 1H), 3.90 (s, 3H), 1.53 (s, 9H).

***4-Iodo-6-methoxypyridin-3-amine***

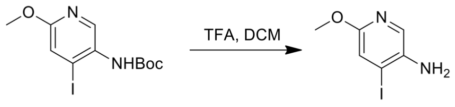


To a solution of tert-butyl N-(4-iodo-6-methoxy-3-pyridyl)carbamate (12.0 g, 34.3 mmol) in DCM (10 mL) was added TFA (66.0 g, 579 mmol). The reaction mixture was stirred at 25 °C for 0.5 hr. On completion, the reaction mixture was concentrated in vacuo to give a residue. The residue was re-dissolved into DCM (30 mL) and $H_2O$ (3 mL), and basified with $NaHCO_3$ solid to pH~9. Then the organic layer was dried over $Na_2SO_4$, filtered and concentrated in vacuo to give the title compound (8.8 g, 92% yield), which was used directly for next step without further purification. $^1$H NMR (400 MHz, $CDCl_3$) δ = 7.64 (s, 1H), 7.14 (s, 1H), 3.84 (s, 3H).

***N-benzyl-6-methoxythiazolo[4,5-c]pyridin-2-amine***

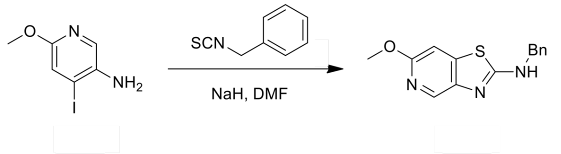


To a solution of 4-iodo-6-methoxy-pyridin-3-amine (8.8 g, 31.7 mmol) and isothiocyanatomethylbenzene (9.45 g, 63.4 mmol) in DMF (130 mL) was added NaH (3.04 g, 76.0 mmol, 60% purity) at 25 °C. The reaction mixture was stirred at 25 °C for 15 mins and then at 65 °C for 1 hr. On completion, the reaction mixture was diluted with EA (100 mL) and washed with the saturated NaCl aqueous (3 X 150 mL). The collected organic layer was dried over $Na_2SO_4$, filtered and concentrated in vacuo to give a residue. The residue was purified by silica gel chromatography (PE:EA=4:1) to give the title compound (7.10 g, 50% yield) as a yellow solid. LCMS $(M+1)^+$: 272.1.

***N,N-dibenzyl-6-methoxythiazolo[4,5-c]pyridin-2-amine***

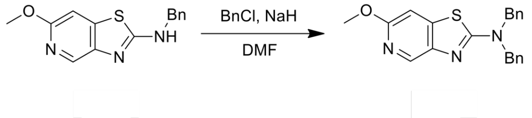


A mixture of N-benzyl-6-methoxy-thiazolo[4,5-c]pyridin-2-amine (6.7 g, 14.8 mmol) in DMF (130 mL) was added NaH (1.48 g, 37.0 mmol, 60% purity) . After being stirred for 30 mins, BnBr (3.04 g, 17.9 mmol) was added. The reaction mixture was stirred at 25 °C for 10 hr. On completion, the reaction mixture was diluted with $H_2O$ (20 mL) and extracted with EA (200 mL). The combined organic layers were washed with brine (3 X 100 mL), dried over $Na_2SO_4$, filtered and concentrated in vacuo to give a residue. The residue was purified by silica gel chromatography (PE:EA=4:1) to give the title compound (4.20 g, 74% yield) as a yellow oil. $^1$H NMR (400 MHz, $CDCl_3$) δ = 8.41 (s, 1H), 7.40 - 7.28 (m, 10H), 6.98 (s, 1H), 4.72 (s, 4H), 3.96 (s, 3H).

***2-(Dibenzylamino)thiazolo[4,5-c]pyridin-6-ol***

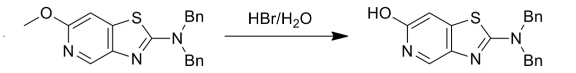

A mixture of N,N-dibenzyl-6-methoxy-thiazolo[4,5-c]pyridin-2-amine (1.00 g, 2.77 mmol) in HBr/$H_2O$ (2 M, 10 mL, 30% purity) was stirred at 80 °C for 12 hr. On completion, the reaction mixture was diluted with $H_2O$ (50 mL). The reaction mixture was basified with $NaHCO_3$ solid to pH~ 8 and extracted with EA (300 mL). The combined organic layers were washed with brine (3 X 30 mL), dried over $Na_2SO_4$, filtered and concentrated in vacuo to give the title compound (830 mg, 86% yield) as a black solid, which was used directly for next step without further purification. $^1$H NMR (400 MHz, DMSO-*d6*) δ = 11.3 (br s, 1H), 7.61 (s, 1H), 7.65 - 7.27 (m, 10H), 6.73 (s, 1H), 4.71 (s, 4H).

***<u>N,N-dibenzyl-6-(bromodifluoromethoxy)thiazolo[4,5-c]pyridin-2-amine</u>***

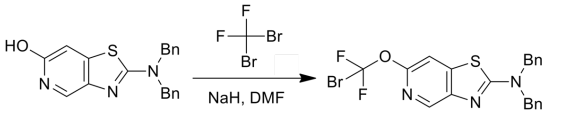

To a solution of 2-(dibenzylamino)thiazolo[4,5-c]pyridin-6-ol (400 mg, 1.15 mmol) in DMF (8 mL) was added NaH (207 mg, 5.18 mmol, 60% purity) at 0 °C. The reaction mixture was stirred at 50 °C for 30 mins. Then the reaction mixture was cooled to 0 °C, dibromo(difluoro)methane (725 mg, 3.45 mmol) was added, and the reaction mixture was stirred at 70 °C for 12 hr. On completion, the reaction mixture was diluted with $H_2O$ (20 mL) and extracted with EA (50 mL). The organic layer was washed with the saturated NaCl aqueous (3 X 30 mL). The organic layer was dried over $Na_2SO_4$, filtered and concentrated in vacuo to give a residue The residue was purified by silica gel chromatography (PE:EA=20:1) to give the title compound (100 mg, 18% yield) as a yellow oil. LCMS $(M+1)^+$ = 476.1, 478.0.

***<u>N,N-dibenzyl-6-(trifluoromethoxy)thiazolo[4,5-c]pyridin-2-amine (C-11)</u>***

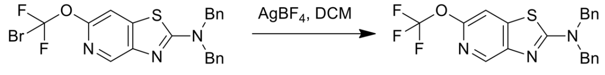

C-11

To a solution of N,N-dibenzyl-6-[bromo(difluoro)methoxy]thiazolo[4,5-c]pyridin-2-amine (100 mg, 210 μmol) in DCM (2.5 mL) was added $AgBF_4$ (100 mg, 514 μmol). The reaction mixture was stirred at 25 °C for 12 hr. On completion, the reaction mixture was filtered and concentrated in vacuo to give a residue. The residue was purified by silica gel chromatography (PE:EA=15:1) to give an impure product. The impure product was re-purified by Prep-HPLC (column: Shim-pack C18 150mm * 25mm * 10 µm; mobile phase: [water (0.225% FA)-ACN]; B%: 69%-99%, 10 min) to give the title compound (12.0 mg, 14% yield) as a white solid. LCMS $(M+1)^+$ = 416.1. $^1H$ NMR (400 MHz, $CDCl_3$) δ = 8.53 (s, 1H), 7.40 - 7.27(m, 11H), 4.76 (s, 4H). *Note $^{13}C$ & $^{19}F$ spectra not obtained due to lack of material.

***<u>Synthesis of C-13</u>***

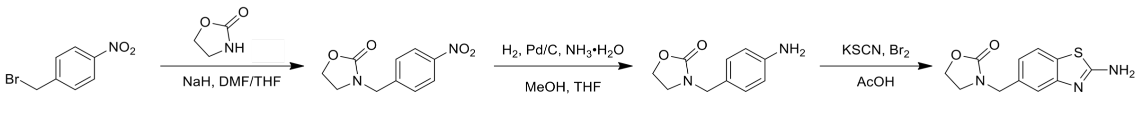


***<u>3-(4-Nitrobenzyl)oxazolidin-2-one</u>***

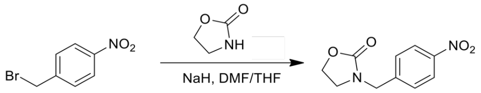


To a suspension of NaH (555 mg, 13.8 mmol, 60% purity) in THF (15 mL) was added oxazolidin-2-one (967 mg, 11.1 mmol) at 0 °C. The reaction mixture was stirred at 25 °C for 1 hr. Then a solution of 1-(bromomethyl)-4-nitro-benzene (2.00 g, 9.26 mmol) in THF/DMF=10/1 (60 mL) was added dropwise under 0 °C. The resulting mixture was stirred at 25 °C for 12 hr. On completion, the reaction mixture was poured into water (100 mL) and concentrated in vacuo to remove THF. The residue was diluted with water (100 mL) and extracted with DCM (2 x 100 mL). The combined organic layers were dried with $Na_2SO_4$, filtered and concentrated in vacuo to give the title compound (2.20 g, 87% purity). LCMS $(M+1)^+$ : 223.0, $^1H$ NMR (400MHz, MeOD-$d_4$) δ = 8.24 (d, $J$ = 8.8 Hz, 2H), 7.56 (d, $J$ = 8.8 Hz, 2H), 4.56 (s, 2H), 4.41 - 4.35 (m, 2H), 3.60 - 3.52 (m, 2H) ;

***<u>3-(4-Amiobenzyl)oxazolidin-2-one</u>***

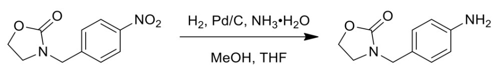


To a solution of 3-[(4-nitrophenyl)methyl]oxazolidin-2-one (2.20 g, 9.90 mmol) in MeOH (150 mL) and THF (30 mL) was added $NH_3{\cdot}H_2O$ (1.82 g, 2 mL, 30% purity) and Pd/C (0.1 g, 10% purity). The reaction mixture was stirred at 25 °C for 12 hr under $H_2$. On completion, the reaction mixture was filtered with celite and concentrated in vacuo to give a residue. The residue was purified by column chromatography (PE:EA = 1:2) to give the title compound (1.20 g, 63% yield) as gray solid. LCMS $(M+1)^+$ : 193.1, $^1H$ NMR (400 MHz, MeOD-$d_4$) δ = 7.03 (d, $J$ = 8.4 Hz, 2H), 6.70 (d, $J$ = 8.4 Hz, 2H), 4.32 - 4.26 (m, 2H), 4.26 (s, 2H), 3.46 - 3.40 (m, 2H).

***3-((2-Aminobenzo[d]thiazol-6-yl)methyl)oxazolidin-2-one (C-13)***

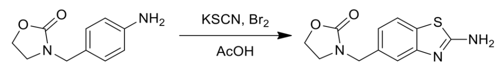


C-13

To a solution of 3-[(4-aminophenyl)methyl]oxazolidin-2-one (0.1 g, 520 µmol) in AcOH (2 mL) was added KSCN (202 mg, 2.08 mmol). The reaction mixture was stirred at 25 °C for 0.2 hr. Then a solution of $Br_2$ (83.0 mg, 520 µmol) in AcOH (1 mL) was added. The reaction mixture was stirred at 25 °C for 12 hr. On completion, the mixture was adjusted to pH= 8 with $NH_3{\cdot}H_2O$. The mixture was extracted with EA (3 x 20 mL). The combined organic layers were dried with $Na_2SO_4$, filtered and concentrated in vacuo to give yellow solid, which was purified by Prep-TLC and column chromatography (PE) to give the title compound (40.0 mg, 29% yield) as yellow solid. LCMS $(M+1)^+$ : 250.1, $^1H$ NMR (400 MHz, ,MeOD-$d_4$) δ = 7.55 (d, $J$ = 1.2 Hz, 1H), 7.37 (d, $J$ = 8.0 Hz, 1H), 7.20 (dd, $J$ = 1.2, 8.0 Hz, 1H), 4.43 (s, 2H), 4.35 - 4.29 (m, 2H), 3.51 - 3.45 (m, 2H). $^{13}C$ NMR (101 MHz, DMSO-$d_6$) δ 167.17, 158.44, 152.86, 131.75, 129.39, 125.95, 120.78, 118.14, 62.14, 47.71, 44.10.

#### ***Synthesis of C-15***

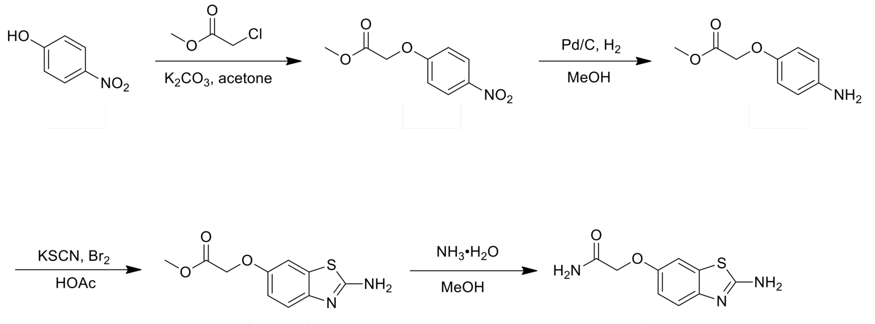


C-15

#### ***Methyl 2-(4-nitrophenoxy)acetate***

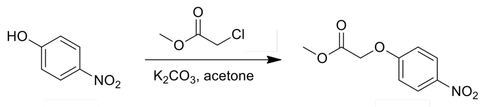


To a solution of 4-nitrophenol (3.00 g, 21.5 mmol) in acetone (50 mL) was added $K_2CO_3$ (4.47 g, 32.3 mmol) and methyl 2-chloroacetate (2.81 g, 25.8 mmol, 2.28 mL). The reaction mixture was stirred at 60 °C for 6 hr. On completion, the reaction mixture was concentrated in vacuo to give a residue. The residue was added ice water (100 mL) and a white solid was formed. The reaction was filtered and the filter cake was dried in vacuo to give the title compound (3.50 g, 76% yield) as a white solid. $^1$H NMR (400MHz, $CDCl_3$) δ = 8.24 (d, *J* = 9.2 Hz, 2H), 6.99 (d, *J* = 9.2 Hz, 2H), 4.76 (s, 2H), 3.85 (s, 3H).

#### ***Methyl 2-(4-aminophenoxy)acetate***

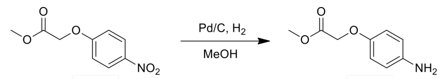


To a solution of methyl 2-(4-nitrophenoxy)acetate (3.50 g, 16.5 mmol) in MeOH (40 mL) was added Pd/C (350 mg, 10% purity) under $N_2$. The suspension was degassed under vacuum and purged with $H_2$ several times. The mixture was stirred under $H_2$ (15 psi) at 25 °C for 12 hr. On completion, the reaction mixture was filtered and the filtrate was concentrated in vacuo. The residue was purified by silica gel chromatography (PE:EA = 2:1) to give the title compound (2.00 g, 66% yield) as a light brown oil. $^1$H NMR (400MHz, $CDCl_3$) δ = 6.78 (d, *J* = 8.8 Hz, 2H), 6.64 (d, *J* = 8.8 Hz, 2H), 4.57 (s, 2H), 3.81 (s, 3H), 3.21 (br s, 2H).

***Methyl 2-[(2-amino-1,3-benzothiazol-6-yl)oxy]acetate***

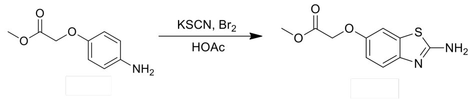

Methyl 2-(4-aminophenoxy)acetate (2.00 g, 11.0 mmol) and KSCN (4.29 g, 44.1 mmol) in AcOH (12 mL) was stirred at 25 °C for 30 mins. Then a solution of $Br_2$ (1.76 g, 11.0 mmol) in AcOH (6 mL) was added dropwise. The reaction mixture was stirred at 25 °C for 12 hr. On completion, the reaction mixture was quenched with ice water (60 mL), basified with saturated $NaHCO_3$ solution till the pH = 7 and a yellow solid was precipitated. The reaction mixture was filtered and the filter cake was dried in vacuo to give the title compound (2.00 g, 76% yield) as a yellow solid. $^1$H NMR (400MHz, $CDCl_3$) δ = 7.38 (d, $J$ = 8.8 Hz, 1H), 7.09 (br s, 1H), 6.87 (d, $J$ = 8.8 Hz, 1H), 5.09 (br s, 2H), 4.58 (s, 2H), 3.74 (s, 3H).

***2-[(2-Amino-1,3-benzothiazol-6-yl)oxy]acetamide (C-15)***

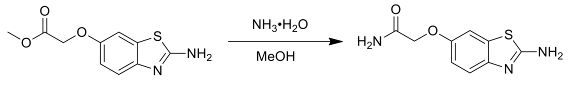
C-15

To a solution of methyl 2-[(2-amino-1,3-benzothiazol-6-yl)oxy]acetate (150 mg, 629 µmol) in MeOH (3 mL) was added $NH_3$•$H_2O$ (1.82 g, 2.0 mL, 27% purity). The mixture was stirred at 50 °C for 12 hr. On completion, the reaction mixture was concentrated in vacuo to get a residue. The residue was purified by Prep-HPLC (column: Xtimate C18 150*25mm*5um; mobile phase: [water (0.05% ammonia hydroxide v/v)-ACN]) to give the title compound (76.0 mg, 54% yield) as a white solid. LCMS $(M+1)^+$: 224.0. $^1$H NMR (400MHz, DMSO-$d_6$) δ = 7.50 (br s, 1H), 7.40 (br s, 1H), 7.33 - 7.20 (m, 4H), 6.87 (dd, $J$ = 2.4, 8.8 Hz, 1H), 4.39 (s, 2H). $^{13}$C NMR (101 MHz, DMSO-$d_6$) δ 170.64, 165.47, 153.18, 147.84, 132.22, 118.47, 114.14, 107.19, 68.03.

***Synthesis of C-16***

***2-[(2-Amino-1,3-benzothiazol-6-yl)oxy]-N-methyl-acetamide (C-16)***

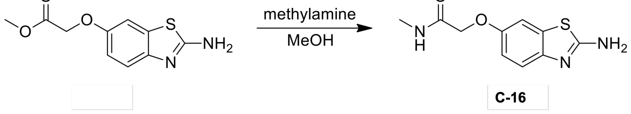
C-16

To a solution of methyl 2-[(2-amino-1,3-benzothiazol-6-yl)oxy]acetate (150 mg, 629 µmol) in MeOH (3 mL) was added methylamine (72.4 mg, 629 µmol, 27% purity) in MeOH (3 mL). The reaction mixture was stirred at 50 °C for 12 hr. On completion, the reaction mixture was concentrated in vacuo to get a residue. The residue was purified by Prep-HPLC (column: Xtimate C18 150*25mm*5um; mobile phase: [water (0.05% ammonia hydroxide v/v)-ACN]) to give the title compound (34.6 mg, 23% yield) as a white solid. LCMS $(M+1)^+$: 237.8. $^1$H NMR (400MHz, DMSO-$d_6$) δ = 8.01 (br s, 1H), 7.33 - 7.21 (m, 4H), 6.87 (dd, $J$ = 2.4, 8.8 Hz, 1H), 4.42 (s, 2H), 2.66 (d, $J$ = 4.4 Hz, 3H). $^{13}$C NMR (101 MHz, DMSO-$d_6$) δ 168.67, 165.52, 153.13, 147.93, 132.25, 118.49, 114.20, 107.28, 68.39, 25.83.

***Synthesis of C-17***

***2-[(2-Amino-1,3-benzothiazol-6-yl)oxy]-N,N-dimethyl-acetamide (C-17)***

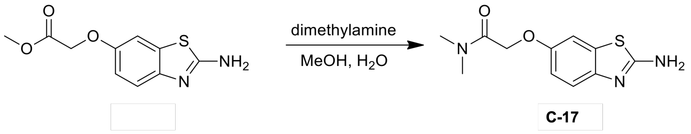


To a solution of methyl 2-[(2-amino-1,3-benzothiazol-6-yl)oxy]acetate (150 mg, 629 µmol) in MeOH (3 mL) was added a solution of dimethylamine (2.67 g, 19.5 mmol, 3.0 mL, 33% purity) in $H_2O$ (3 mL). The reaction mixture was stirred at 50 °C for 12 hr. On completion, the reaction mixture was concentrated in vacuo to get a residue. The residue was purified by Prep-HPLC (column: Xtimate C18 150*25mm*5um; mobile phase: [water (0.05% ammonia hydroxide v/v)-ACN]) to give the title compound (45.2 mg, 28% yield) as a white solid. LCMS $(M+1)^+$: 252.1. $^1$H NMR (400MHz, DMSO-$d_6$) δ = 7.31 - 7.19 (m, 4H), 6.82 (dd, $J$ = 2.4, 8.8 Hz, 1H), 4.75 (s, 2H), 3.00 (s, 3H), 2.84 (s, 3H). $^{13}$C NMR (101 MHz, DMSO-$d_6$) δ 167.78, 165.31, 153.55, 147.61, 132.17, 118.39, 114.01, 107.02, 67.03, 36.11, 35.42.

#### ***Synthesis of C-18***

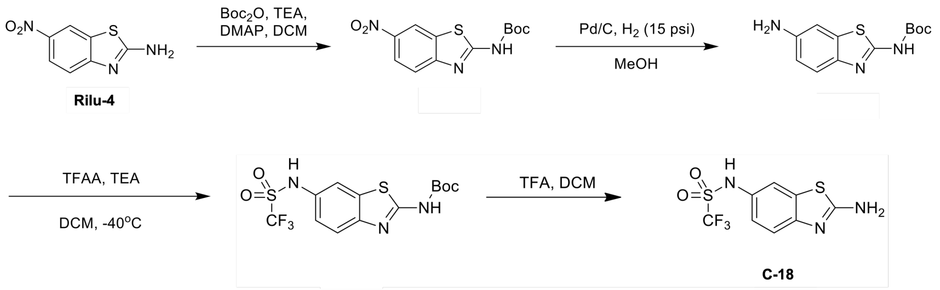


#### ***Tert-butyl (6-nitrobenzo[d]thiazol-2-yl)carbamate***

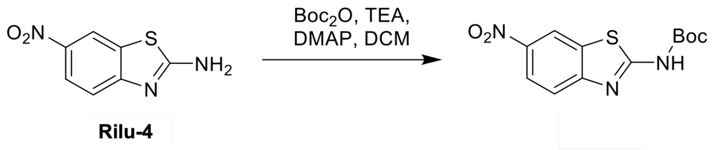


To a solution of 6-nitro-1,3-benzothiazol-2-amine (2.00 g, 10.2 mmol) and $Boc_2O$ (6.70 g, 30.7 mmol) in DCM (30 mL) was added TEA (3.10 g, 30.7 mmol) and DMAP (62.6 mg, 512 µmol). The reaction mixture was stirred at 40 °C for 16 hr. On completion, the reaction mixture was concentrated in vacuo to give the title compound (3.50 g, 95% yield) as a yellow solid. LCMS $(MS+1)^+$ : 296.0

#### ***Tert-butyl (6-aminobenzo[d]thiazol-2-yl)carbamate***

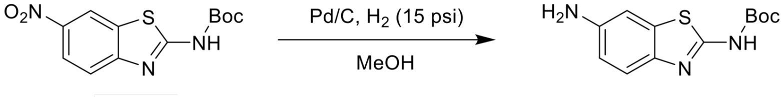


A mixture of tert-butyl N-tert-butoxycarbonyl-N-(6-nitro-1,3-benzothiazol-2-yl)carbamate (3.00 g, 7.59 mmol ) and Pd/C (3.00 g, 2.82 mmol, 10% purity) in MeOH (50 mL) was stirred at 35 °C for 2 hr under $H_2$ (15 psi). On completion, the reaction mixture was filtered and the filtrate was concentrated in vacuo. The residue was purified by column chromatography (PE:EA:DCM=5:1:1) to give the title compound (0.900 g, 39% yield) as a yellow solid. LCMS $(MS+1)^+$ : 266.0

#### ***Tert-butyl (6-(trifluoromethylsulfonamido)benzo[d]thiazol-2-yl)carbamate***

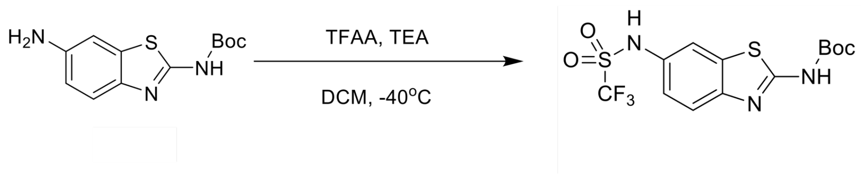

To a solution of tert-butyl N-(6-amino-1,3-benzothiazol-2-yl)carbamate (0.500 g, 1.88 mmol) in DCM (5 mL) was added TEA (381 mg, 3.80 mmol), and then a solution of TFAA (638 mg, 2.30 mmol ) in DCM (3 mL) was added at -40 °C . The mixture was stirred at -40 °C for 0.5 hr. On completion, the reaction mixture was quenched by $H_2O$ (10 mL) at 0 °C, and extracted with DCM (3 x 10 mL). The combined organic layers were washed with brine (3 x 10 mL), dried over $Na_2SO_4$, filtered and concentrated in vacuo to give a residue. The residue was purified by Prep-TLC (DCM:MeOH=25:1) to give the title compound (0.500 g, 43% yield) as a yellow solid. LCMS $(MS+1)^+$ : 398.0.

***N-(2-aminobenzo[d]thiazol-6-yl)-1,1,1-trifluoromethanesulfonamide (C-18)***

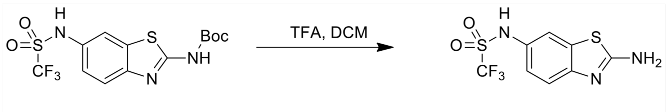

C-18

To a solution of tert-butyl N-[6-(trifluoromethylsulfonylamino)-1,3-benzothiazol-2-yl]carbamate (0.300 g, 755 µmol) in DCM (5 mL) was added TFA (1 mL) at 0 °C. The reaction mixture was stirred at 20 °C for 2 hr. On completion, the reaction mixture was concentrated in vacuo. The residue was diluted with DCM (10 mL) and basified with TEA (1 mL). The mixture was concentrated in vacuo to give a residue. The residue was purified by Prep-HPLC [water(0.225%FA)-ACN];B%: 10%-40%,10min) to give the title compound (50.0 mg, 19% yield, FA salt) as a yellow solid. LCMS $(MS+1)^+$ : 297.9, $^1$H NMR (400 MHz, DMSO-$d_6$) δ = 7.56 - 7.52 (m, 3H), 7.27 (d, $J$ = 8.4 Hz, 1H), 7.05 - 7.02 (m, 1H). $^{13}$C NMR (101 MHz, DMSO-$d_6$) δ 167.45, 151.33, 131.88, 129.87, 123.06, 120.70 (q, $J$ = 324 Hz), 118.07, 117.69. $^{19}$F NMR (376 MHz, DMSO-$d_6$) δ -75.18.

#### Synthesis of C-19

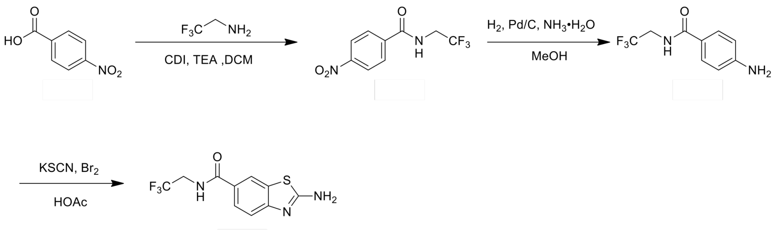

C-19

#### 4-Nitro-N-(2,2,2-trifluoroethyl)benzamide

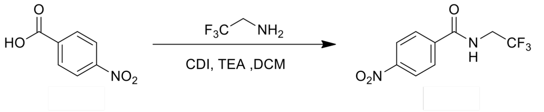

To a solution of 4-nitrobenzoic acid (2.00 g, 11.9 mmol) in DCM (20 mL) was added CDI (2.91 g, 17.9 mmol). The reaction mixture was stirred at 25 °C for 1 hr. Then a solution of 2,2,2-trifluoroethanamine (1.42 g, 14.3 mmol, 1.13 mL) and TEA (1.82 g, 17.9 mmol, 2.50 mL) in DCM (10 mL) was added. The reaction mixture was stirred at 25 °C for 1 hr. On completion, the reaction mixture was quenched with water (20 mL). The organic layer was separated and dried with $Na_2SO_4$, filtered and concentrated in vacuo to give the title compound (2.50 g, 55% yield) as yellow oil. LCMS $(MS+1)^+$ : 249.0.

#### 4-Amino-N-(2,2,2-trifluoroethyl)benzamide

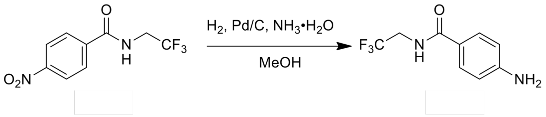

To a solution of 4-Nitro-N-(2,2,2-trifluoroethyl)benzamide (2.50 g, 6.65 mmol) in MeOH (40 mL) was added Pd/C (0.25 g, 10% purity) and $NH_3{\cdot}H_2O$ (330 uL, 30% purity). The reaction mixture was stirred at 25 °C for 12 hr under $H_2$ (15 psi). On completion, the reaction mixture was filtered through celite and concentrated in vacuo to give the brown oil, which was purified by column chromatography (PE:EA = 2:1) to get the title compound (1.10 g, 75% yield) as white solid. LCMS $(MS+1)^+$ : 219.1, $^1$H NMR (400 MHz, MeOD-$d_4$) δ = 7.66 - 7.60 (m, 2H), 6.69 - 6.65 (m, 2H), 4.03 (d, $J$ = 9.2 Hz, 2H).

#### *2-Amino-N-(2,2,2-trifluoroethyl)benzo[d]thiazole-6-carboxamide (C-19)*

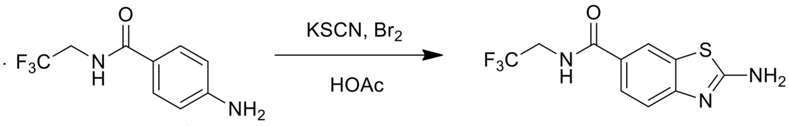

C-19

A solution of 4-amino-N-(2,2,2-trifluoroethyl)benzamide (0.200 g, 916 µmol) and KSCN (356 mg, 3.67 mmol) in AcOH (3 mL) was stirred at 25 °C for 0.2 hr. Then a solution of $Br_2$ (146 mg, 916 µmol) in AcOH (1 mL) was added. The reaction mixture was stirred at 25 °C for 12 hr. On completion, $NH_3$•$H_2O$ was added to adjust pH = 8. The mixture was extracted with EA (2 x 10 mL). The combined organic layers were dried with $Na_2SO_4$, filtered and concentrated in vacuo to give yellow solid, which was purified by Prep-HPLC (column: Waters Xbridge 150*25 5u;mobile phase: [water (0.05% ammonia hydroxide v/v)-ACN];B%: 10%-40%,10min) and re-purified by Prep-TLC (PE:EA=1:1) to give the title compound (38.3 mg, 15% yield) as yellow solid. LCMS $(MS+1)^+$ : 276.0, $^1$H NMR (300 MHz, MeOD-$d_4$) δ = 8.13 (d, *J* = 1.8 Hz, 1H), 7.78 (dd, *J* = 1.8, 8.7 Hz, 1H), 7.42 (d, *J* = 8.7 Hz, 1H), 4.15 - 4.03 (m, 2H). $^{13}$C NMR (101 MHz, DMSO-$d_6$) δ 169.35, 167.04, 156.21, 131.43, 126.04, 125.82, 125.37 (q, *J* = 278 Hz), 121.15, 117.37, 40.81. $^{19}$F NMR (376 MHz, DMSO-$d_6$) δ -70.43.

#### *Synthesis of C-20*

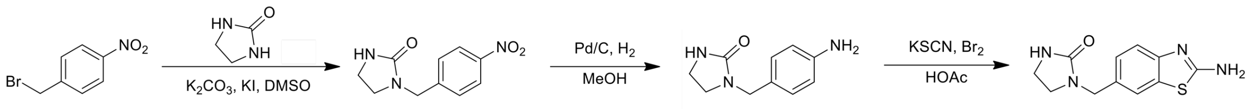

C-20

#### *1-(4-Nitrobenzyl)imidazolidin-2-one*

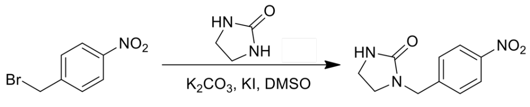

A mixture of imidazolidin-2-one (1.00 g , 11.6 mmol), 1-(bromomethyl)-4-nitro-benzene (2.51 g, 11.6 mmol), KI (964 mg, 5.81 mmol) and $K_2CO_3$ (1.61 g, 11.6 mmol) in DMSO (10 mL) was stirred at 105 °C for 2 hr. On completion, the reaction was poured into ice-water (150 mL), extracted with EA (3 x 50 mL). The combined organic layer was washed with brine, dried over $Na_2SO_4$, filtered and concentrated in vacuo. The residue was purified by column chromatography

(PE:EA = 2:1) to give the title compound (330 mg, 11% yield) as yellow solid. LCMS $(MS+1)^+$: 222.0. $^1$H NMR (400 MHz, $CDCl_3$) δ = 8.21 (d, *J* = 8.8 Hz, 2H), 7.47 (d, *J* = 8.8 Hz, 2H), 4.79 (br s, 1H), 4.48 (s, 2H), 3.52 - 3.43 (m, 2H), 3.38 - 3.21 (m, 2H).

***1-(4-Aminobenzyl)imidazolidin-2-one***

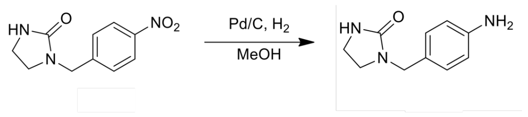


To a solution of 1-[(4-nitrophenyl)methyl]imidazolidin-2-one (330 mg, 1.49 mmol) in MeOH (10 mL) was added Pd/C (40 mg, 10% purity) under $N_2$. The suspension was degassed under vacuum and purged with $H_2$ several times. The mixture was stirred at 25°C for 12 hr under $H_2$ (15 psi). On completion, the reaction was filtered and the filtrate was concentrated in vacuo to give the title compound (260 mg, 85% yield) as yellow solid. LCMS $(MS+1)^+$: 192.0. $^1$H NMR (400 MHz, DMSO-*d*$_6$) δ = 6.94 (d, *J* = 8.4 Hz, 2H), 6.57 (d, *J* = 8.4 Hz, 2H), 6.01 (br s, 1H), 4.72 (br s, 2H), 3.25 - 3.21 (m, 2H), 3.16 - 3.13 (m, 2H).

***1-((2-Aminobenzo[d]thiazol-6-yl)methyl)imidazolidin-2-one (C-20)***

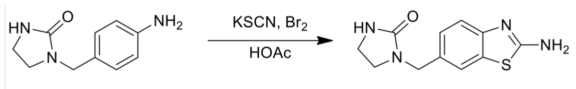


C-20

A mixture of 1-[(4-aminophenyl)methyl]imidazolidin-2-one (160 mg, 836 μmol) and KSCN (325 mg, 3.35 mmol) in HOAc (1 mL) was stirred at 25 °C for 30 min, then a solution of $Br_2$ (133.71 mg, 836 μmol) in HOAc (0.5 mL) was added. The reaction mixture was stirred at 25 °C for 12 hr. On completion, the reaction was poured into 28% ammonium aqueous (20 mL) at 0 °C, and extracted with DCM (5 x 20 mL). The combined organic layer was concentrated in vacuo. The residue was purified by Prep-HPLC (column: Phenomenex Gemini 150*25mm*10um;mobile phase: [water(0.04%NH3H2O+10mM NH4HCO3)-ACN];B%: 7%-34%,10min) to give the title compound (94.1 mg, 45% yield) as white solid. LCMS $(MS+1)^+$: 249.0. $^1$H NMR (400 MHz, DMSO-*d*$_6$) δ = 7.50 (d, *J* = 1.2 Hz, 1H), 7.43 (s, 2H), 7.28 (d, *J* = 8.0 Hz, 1H), 7.07 (dd, *J* = 1.6, 8.0 Hz, 1H), 6.38 (s, 1H), 4.22 (s, 2H), 3.25 - 3.13 (m, 4H). $^{13}$C NMR (101 MHz, DMSO-*d*$_6$) δ 166.90, 162.56, 152.53, 131.60, 130.88, 125.92, 120.62, 118.02, 47.14, 44.44, 37.77.

***Synthesis of C-21***

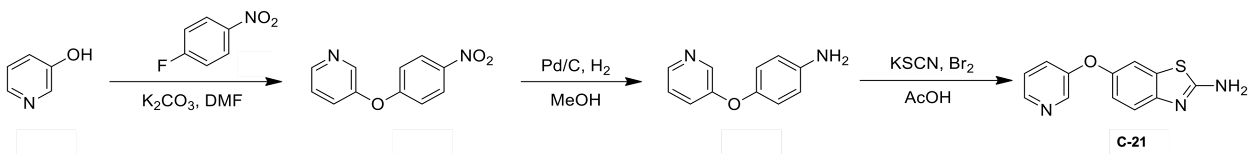


***3-(4-Nitrophenoxy)pyridine***

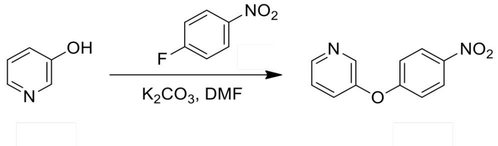


To a solution of pyridin-3-ol (1.00 g, 10.5 mmol) in DMF (15 mL) was added $K_2CO_3$ (2.18 g, 15.7 mmol) and 1-fluoro-4-nitro-benzene (1.48 g, 10.5 mmol). The reaction mixture was stirred at 120 °C for 3 hr. The reaction mixture was concentrated in vacuo to get a residue. The residue was diluted with water (30 mL) and EA (30 mL). The organic layer was collected, dried over anhydous $Na_2SO_4$, filtered and concentrated in vacuo to get a residue. The residue was purified by chromatography column (PE:EA = 2:1) to give the title compound (850 mg, 37% yield) as a yellow solid. $^1$H NMR (400MHz, $CDCl_3$) δ = 8.45 (dd, *J* = 1.2, 4.4 Hz, 1H), 8.41 (d, *J* = 2.4 Hz, 1H), 8.20 - 8.14 (m, 2H), 7.40 - 7.29 (m, 2H), 7.03 - 6.95 (m, 2H).

***4-(3-Pyridyloxy)aniline***

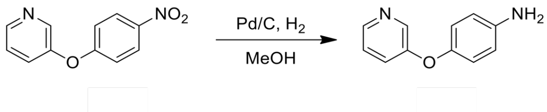


To a solution of 3-(4-nitrophenoxy)pyridine (850 mg, 3.93 mmol) in MeOH (40 mL) was added Pd/C (100 mg, 10% purity). The suspension was degassed under vacuum and purged with $H_2$ several times. The mixture was stirred under $H_2$ (15 psi) at 25 °C for 12 hr. The reaction mixture was filtered and the filtrate was concentrated in vacuo. The residue was purified by column chromatography (PE:EA = 1:1) to give the title compound (630 mg, 86% yield) as a white solid. $^1$H NMR (400MHz, $CDCl_3$) δ = 8.35 (d, *J* = 1.6 Hz, 1H), 8.27 (t, *J* = 3.2 Hz, 1H), 7.22 - 7.17 (m, 2H), 6.90 - 6.85 (m, 2H), 6.72 - 6.66 (m, 2H), 3.44 (br. s, 2H).

***6-(3-Pyridyloxy)-1,3-benzothiazol-2-amine (C-21)***

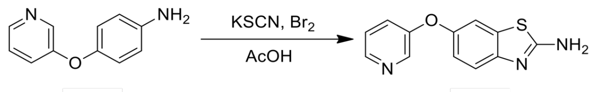


C-21

4-(3-pyridyloxy)aniline (300 mg, 1.61 mmol) and KSCN (626 mg, 6.44 mmol) in AcOH (5 mL) was stirred at 25 °C for 30 mins. Then a solution of $Br_2$ (257 mg, 1.61 mmol) in AcOH (2 mL) was added drop-wise. The reaction mixture was stirred at 25 °C for 1 hr. On completion, the reaction mixture was quenched with ice water (30 mL), basified with $NH_3$•$H_2O$ solution till the pH = 7 and then extracted with DCM (3 X 30 mL). The organic layer was collected, dried over anhydrous $Na_2SO_4$, filtered and concentrated in vacuo to get a residue. The residue was purified by chromatography column (PE:EA = 10:1) to give the title compound (220 mg, 56% yield) as a light yellow solid LCMS $(M+1)^+$: 244.1. $^1$H NMR (400MHz, DMSO-$d_6$) δ = 8.35 (d, $J$ = 2.4 Hz, 1H), 8.31 (dd, $J$ = 1.6, 4.4 Hz, 1H), 7.50 (d, $J$ = 2.4 Hz, 1H), 7.46 (s, 2H), 7.41 - 7.32 (m, 3H), 6.98 (dd, $J$ = 2.4, 8.8 Hz, 1H). $^{13}$C NMR (101 MHz, DMSO-$d_6$) δ 166.96, 155.07, 150.34, 149.95, 144.22, 140.45, 132.73, 125.02, 124.75, 118.90, 118.03, 112.90.

#### ***Synthesis of C-23***

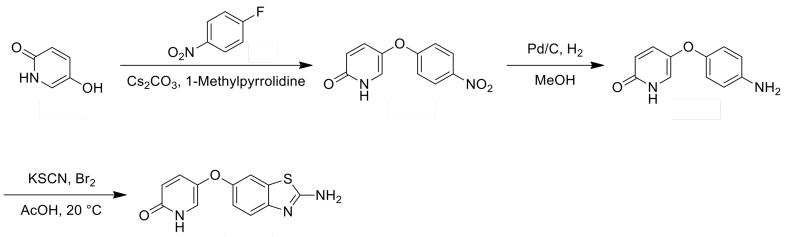


C-23

#### ***5-(4-Nitrophenoxy)pyridin-2(1H)-one***

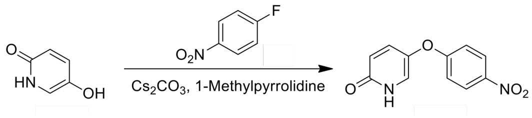


To a mixture of 5-hydroxy-1*H*-pyridin-2-one (3.15 g, 28.3 mmol) and 1-fluoro-4-nitro-benzene (2.00 g, 14.1 mmol) in 1-methylpyrrolidine (30 mL) was added $Cs_2CO_3$ (9.33 g, 28.6 mmol). The reaction mixture was stirred at 90 °C for 16 hr under nitrogen atmosphere. On completion, the reaction mixture was diluted with water (500 mL) and EA (500 mL). The organic layer was separated and dried over $Na_2SO_4$, filtered and concentrated in vacuo. The residue was purified by silica gel chromatography (DCM:MeOH=10:1) to give the title compound (60.0 mg, 1% yield) as yellow solid. LCMS $(MS+1)^+$: 233.1. $^1H$ NMR (400MHz, DMSO-$d_6$) δ = 11.59 (br s, 1H), 8.24 (d, *J* = 9.2 Hz, 2H), 7.59 (d, *J* = 3.2 Hz, 1H), 7.45 (dd, *J* = 3.2, 9.6 Hz, 1H), 7.15 (d, *J* = 9.2 Hz, 2H), 6.47 (d, *J* = 9.6 Hz, 1H).

***5-(4-Aminophenoxy)pyridin-2(1H)-one***

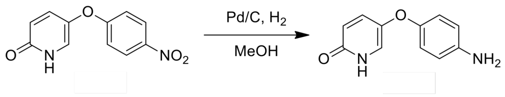


A mixture of 5-(4-nitrophenoxy)-1H-pyridin-2-one (60 mg, 258 µmol) and Pd/C (10 mg, 10% purity) in MeOH (5 mL) was stirred at 20 °C for 5 hr under $H_2$ (15 psi) atmosphere. On completion, the reaction was filtered and the filtrate was concentrated in vacuo to give the title compound (50.0 mg, 90% yield) as a brown solid. LCMS $(MS+1)^+$: 203.1.

***5-((2-Aminobenzo[d]thiazol-6-yl)oxy)pyridin-2(1H)-one (C-23)***

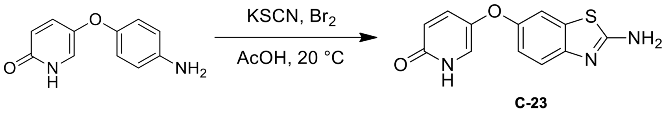


A mixture of 5-(4-aminophenoxy)-1*H*-pyridin-2-one (50.0 mg, 247 µmol) and KSCN (96.1 mg, 989 µmol) in HOAc (3 mL) was stirred at 20 °C for 30 min, then $Br_2$ (39.5 mg, 247 µmol) in HOAc (2 mL) was added. The reaction mixture was stirred at 20 °C for 12 hr. On completion, the reaction was adjusted pH to 10 with ammonium aqueous, extracted with EA (3 x 20 mL). The combined organic layer was washed with brine, dried over $Na_2SO_4$ and concentrated in vacuo. The residue was purified by Prep-HPLC (column: Phenomenex Gemini 150*25mm*10um;mobile phase: [water(0.04%NH3H2O+10mM NH4HCO3)-ACN];B%: 13%-40%,min) to give the title compound (11.9 mg, 18% yield) as a white solid. LCMS $(MS+1)^+$: 259.9. $^1H$ NMR (400MHz, DMSO-$d_6$) δ = 11.31 (br s, 1H), 7.40 - 7.30 (m, 5H), 7.27 (d, *J* = 8.8 Hz, 1H), 6.87 (dd, *J* = 2.4, 8.8 Hz, 1H), 6.40 (d, *J* = 9.6 Hz, 1H). Note $^{13}C$ spectra not obtained due to lack of material.

***Synthesis of C-27***

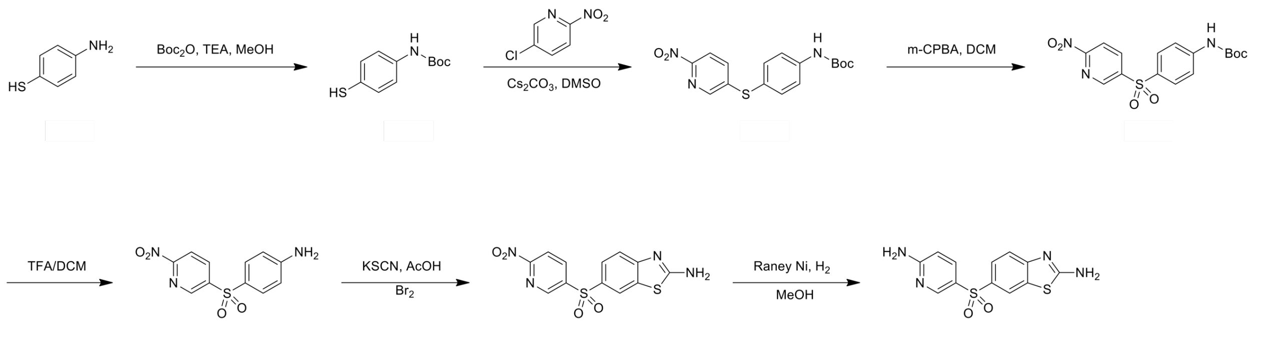


C-27

***Tert-butyl (4-mercaptophenyl)carbamate***

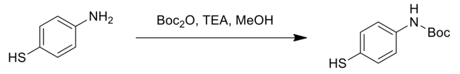


To a solution of 4-aminobenzenethiol (5.00 g, 39.9 mmol) and TEA (8.08 g, 79.8 mmol, 11.1 mL) in MeOH (50 mL) was added $Boc_2O$ (9.59 g, 43.9 mmol) under 0 - 5 °C. The reaction mixture was stirred at 25 °C for 12 hr. On completion, the reaction mixture was diluted with water (20 mL) and filtered. The filter cake was washed with MeOH to give the title compound (4.02 g, 44% yield) as white solid. $^1$H NMR (400 MHz, DMSO-$d_6$) δ = 9.51 (s, 1H), 7.51 - 7.41 (m, 2H), 7.40 - 7.30 (m, 2H), 1.50 (s, 9H).

***Tert-butyl (4-((6-nitropyridin-3-yl)thio)phenyl)carbamate***

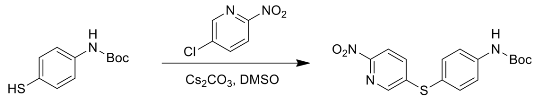


To a mixture of tert-butyl N-(4-sulfanylphenyl)carbamate (1.00 g, 4.44 mmol) and 5-chloro-2-nitro-pyridine (1.00 g, 6.31 mmol) in DMSO (20 mL) was added $Cs_2CO_3$ (2.89 g, 8.88 mmol,). The reaction mixture was stirred at 90 °C for 12 hr. On completion, the reaction mixture was diluted with water (200 mL). The mixture was extracted with EA (2 X 250 mL). The combined organic layers were dried over $Na_2SO_4$, filtered and concentrated in vacuo to give a brown residue. The residue was purified by column chromatography (PE) to give the title compound (1.32 g, 41% yield) as yellow sandy solid. LCMS (MS-56)$^+$ : 291.9, $^1$H NMR (400 MHz, $CDCl_3$) δ = 8.30 (d, $J$ = 2.4 Hz, 1H), 8.10 (d, $J$ = 8.4 Hz, 1H), 7.60 - 7.40 (m, 5H), 6.70 (br s, 1H), 1.52 (s, 9H).

***Tert-butyl (4-((6-nitropyridin-3-yl)sulfonyl)phenyl)carbamate***

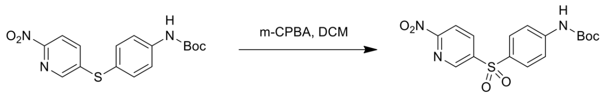


To a solution of tert-butyl N-[4-[(6-nitro-3-pyridyl)sulfanyl]phenyl]carbamate (1.30 g, 1.83 mmol) in DCM (100 mL) was added m-CPBA (1.05 g, 5.50 mmol, 90% purity). The reaction mixture was stirred at 25 °C for 2 hr. On completion, the reaction mixture was quenched by saturated $Na_2S_2O_3$ solution (50 mL). The organic layer was dried over $Na_2SO_4$, filtered and concentrated in vaucuo to give a brown residue. The residue was purified by column chromatography (PE:EA = 6:1) to give the title compound (480 mg, 53% yield) as yellow solid. LCMS $(MS-56)^+$ : 323.9, $^1$H NMR (400 MHz, $CDCl_3$) δ = 9.11 (d, *J* = 2.0 Hz, 1H), 8.51 (dd, *J* = 8.4, 2.0 Hz, 1H), 8.30 (d, *J* = 8.4 Hz, 1H), 7.91 (d, *J* = 8.8 Hz, 2H), 7.60 (d, *J* = 8.8 Hz, 2H), 6.90 (br s, 1H), 1.50 (s, 9H).

***4-((6-Nitropyridin-3-yl)sulfonyl)aniline***

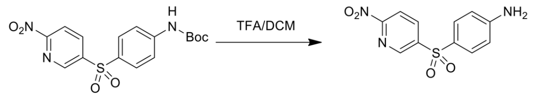


To a solution of tert-butyl N-[4-[(6-nitro-3-pyridyl)sulfonyl]phenyl]carbamate (480 mg, 1.27 mmol) in DCM (50 mL) was added TFA (8.03 g, 5.22 mL). The reaction mixture was stirred at 10 °C for 2 hr. On completion, the reaction mixture was concentrated in vacuo to give a yellow residue. The residue was purified by column chromatography (PE:EA = 2:1) to give the title compound (272 mg, 76% yield) as yellow solid. LCMS $(MS+1)^+$ : 279.9, $^1$H NMR (400 MHz, DMSO-$d_6$) δ = 9.11 (s, 1H), 8.61 (dd, *J* = 8.4, 2.0 Hz, 1H), 8.40 (d, *J* = 8.4 Hz, 1H), 7.70 (d, *J* = 8.8 Hz, 2H), 6.70 (d, *J* = 8.8 Hz, 2H), 6.40 (br s, 2H).

***6-((6-Nitropyridin-3-yl)sulfonyl)benzo[d]thiazol-2-amine***

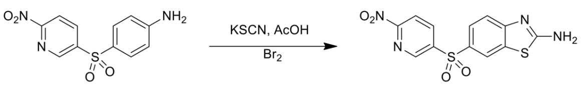


To a solution of 4-[(6-nitro-3-pyridyl)sulfonyl]aniline (100 mg, 358 µmol) in AcOH (2 mL) was added KSCN (139 mg, 1.43 mmol). The reaction mixture was stirred at 20 °C for 0.5 hr. Then a solution of $Br_2$ (62.9 mg, 393 µmol) in AcOH (0.5 mL) was added. The reaction mixture was stirred at 20 °C for 12 hr. On completion, the reaction mixture was diluted with water (20 mL) and basified with saturated $NaHCO_3$ solution (50 mL). The mixture was extracted with EA (2 X 30 mL). The combined organic layers were dried over $Na_2SO_4$, filtered and concentrated in vacuo to give the title compound (140 mg, 97% yield) as yellow solid. $^1$H NMR (400 MHz, DMSO-$d_6$) δ = 9.21 (d, $J$ = 2.0 Hz, 1H), 8.71 (dd, $J$ = 8.4, 2.0 Hz, 1H), 8.42 (d, $J$ = 8.4 Hz, 1H), 8.11 (d, $J$ = 2.0 Hz, 1H), 7.80 (dd, $J$ = 8.8, 2.0 Hz, 1H), 7.20 (br s, 2H), 6.90 (d, $J$ = 8.8 Hz, 1H).

##### ***6-((6-Aminopyridin-3-yl)sulfonyl)benzo[d]thiazol-2-amine (C-27)***

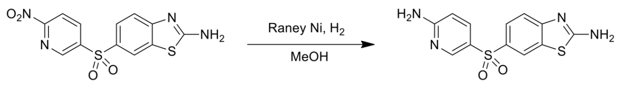


C-27

To a solution of 6-[(6-nitro-3-pyridyl)sulfonyl]-1,3-benzothiazol-2-amine (120 mg, 356 μmol) in MeOH (10 mL) was added Raney Ni (245 mg) at 25 °C. The reaction mixture was stirred for 17 hr under $H_2$ (15 psi) atmosphere. On completion, the reaction mixture was diluted with MeOH and filtered. The filtrate was concentrated in vacuo to give a green residue. The residue was purified by Prep-HPLC (column: Phenomenex Gemini 150*25mm*10um; mobile phase: [water (0.225%FA)-ACN]; B%: 1%-30%, 10min) to give the title compound (17.0 mg, 16% yield) as white solid. LCMS $(M+1)^+$ : 306.9, $^1$H NMR (400 MHz, DMSO-$d_6$) δ = 8.40 (d, $J$ = 2.4 Hz, 1H), 8.25 (d, $J$ = 2.0 Hz, 1H), 7.96 (s, 2H), 7.79 - 7.64 (m, 2H), 7.40 (d, $J$ = 8.4 Hz, 1H), 6.99 (s, 2H), 6.47 (d, $J$ = 8.8 Hz, 1H). $^{13}$C NMR (101 MHz, DMSO-$d_6$) δ 170.63, 162.67, 156.88, 149.10, 136.26, 134.47, 132.22, 125.31, 125.00, 120.88, 117.95, 108.11.

##### ***Synthesis of C-28***

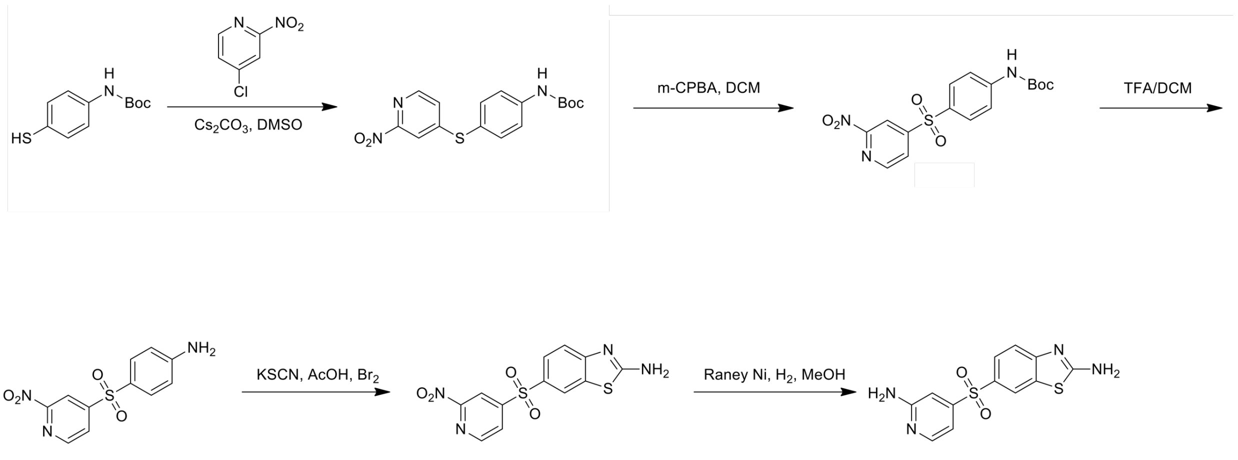


C-28

***Tert-butyl (4-((2-nitropyridin-4-yl)thio)phenyl)carbamate***

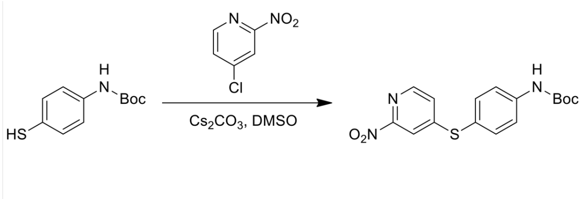


A mixture of tert-butyl N-(4-sulfanylphenyl)carbamate (500 mg, 2.22 mmol), 4-chloro-2-nitro-pyridine (527 mg, 3.33 mmol) and $Cs_2CO_3$ (1.45 g, 4.44 mmol) in DMSO (10 mL) was stirred at 90 °C for 18 hr under nitrogen atmosphere. On completion, the reaction mixture was poured into water (100 mL). The mixture was extracted wit EA (3 X 40 mL). The combined organic layers were concentrated in vacuo to give brown residue. The residue was purified by silica gel chromatography (PE:EA =5:1) to give the title compound (335 mg, 43% yield) as yellow solid. LCMS $(MS+1)^+$ : 348.0. $^1H$ NMR (400 MHz, $CDCl_3$) δ = 8.31 (d, $J$ = 5.2 Hz, 1H), 7.81 (d, $J$ = 1.2 Hz, 1H), 7.61 - 7.51 (m, 2H), 7.51 - 7.49 (m, 2H), 7.31 (s, 1H), 6.70 (s, 1H), 1.51 (s, 9H).

***Tert-butyl (4-((2-nitropyridin-4-yl)sulfonyl)phenyl)carbamate***

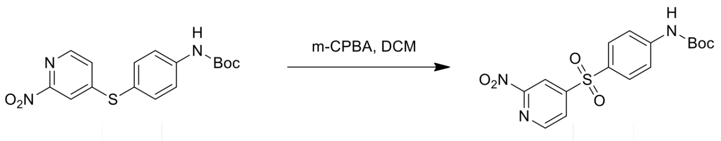


To a mixture of tert-butyl N-[4-[(2-nitro-4-pyridyl)sulfanyl]phenyl]carbamate (330 mg, 949 μmol) in DCM (10 mL) was added m-CPBA (546 mg, 2.85 mmol, 90% purity). The reaction mixture was stirred at 15 °C for 2 hr. On completion, the reaction mixture was poured into saturated $Na_2S_2O_3$ aqueous (20 mL) and stirred for 0.5 hr. The mixture was extracted with DCM (3 X 30 mL). The combined organic layers were washed with saturated $NaHCO_3$ aqueous (30 mL). The organic layer was concentrated in vacuo to give yellow residue. The residue was purified by silica gel chromatography (PE:EA =4:1) to give the title compound (350 mg, 97% yield) as yellow solid. $^1H$ NMR (400 MHz, $CDCl_3$) δ = 8.80 (d, $J$ = 4.8 Hz, 1H) 8.60 - 8.50 (m, 1H) 8.10 (dd, $J$ = 4.8, 1.2 Hz, 1H) 7.90 (s, 1H) 7.90 (s, 1H) 7.62 (s, 1 H) 7.60 (s, 1H) 6.82 (s, 1H) 1.52 (s, 9H).

***4-((6-Nitropyridin-3-yl)sulfonyl)aniline***

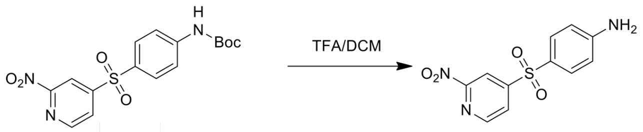


To a solution of tert-butyl N-[4-[(2-nitro-4-pyridyl)sulfonyl]phenyl]carbamate (350 mg, 922 µmol) in DCM (8 mL) was added TFA (1.54 g, 13.5 mmol, 1 mL). The reaction mixture was stirred at 15 °C for 2 hr. On completion, the reaction mixture was concentrated in vacuo to give brown residue. The residue was purified by silica gel chromatography (PE:EA=3:1) to give the title compound (160 mg, 62% yield) as orange solid. LCMS $(MS+1)^+$ : 280.1, $^1$H NMR (400 MHz, DMSO-$d_6$) δ = 8.89 (d, $J$ = 5.2 Hz, 1H), 8.46 (d, $J$ = 0.4 Hz, 1H), 8.29 - 8.26 (m, 1H), 7.69 (d, $J$ = 8.8 Hz, 2H), 6.67 (d, $J$ = 8.8 Hz, 2H), 6.47 (br s, 2H).

***6-((2-Nitropyridin-4-yl)sulfonyl)benzo[d]thiazol-2-amine***

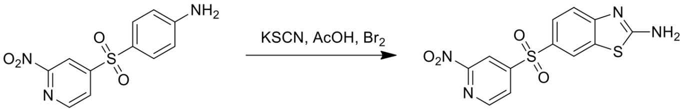


A mixture of 4-[(2-nitro-4-pyridyl)sulfonyl]aniline (80.0 mg, 286 µmol) and KSCN (111 mg, 1.15 mmol) in AcOH (2 mL) was stirred at 15 °C for 0.5 hr. Then a solution of $Br_2$ (45.7 mg, 286 µmol) in AcOH (1 mL) was added. On completion, the reaction mixture was stirred at 25 °C for 17 hr. The reaction mixture was poured into water (20 mL). The mixture was extracted with EA (3 X 50 mL). The combined organic layers were dried over $Na_2SO_4$, filtered and concentrated in vacuo to give the title compound (90.0 mg, 93 % yield) as yellow solid. LCMS $(MS+1)^+$ : 337.0, $^1$H NMR (400 MHz, DMSO-$d_6$) δ = 8.92 (d, $J$ = 4.4 Hz, 1H), 8.62 - 8.54 (m, 1H), 8.52 - 8.46 (m, 1H), 8.38 (d, $J$ = 4.4 Hz, 1H), 8.22 (s, 1H), 7.92 - 7.83 (m, 1H).

***6-((2-Aminopyridin-4-yl)sulfonyl)benzo[d]thiazol-2-amine (C-28)***

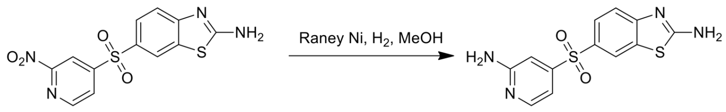


C-28

A mixture of 6-[(2-nitro-4-pyridyl)sulfonyl]-1,3-benzothiazol-2-amine (300 mg, 891 µmol) and Raney Ni (200 mg, 3.41 mmol) in MeOH (15 mL) was degassed under $H_2$ (15 Psi) three times. The mixture was stirred at 25 °C for 1 hr. On completion, the reaction mixture was filtered with celite and the filtrate was concentrated in vacuo to give a residue. The residue was purified by Prep-HPLC (column: Phenomenex Synergi C18 150*25*10um;mobile phase: [water(0.225%FA)-ACN];B%: 1%-28%,10min) to give an impure product (80.0 mg) as light green solid. The solid was re-purified by Prep-TLC (DCM:MeOH = 10:1) to give an off-white solid (60.0 mg). The off-white solid was washed with DCM (5 mL) to give the title compound (42.0 mg, 52% yield) as off-white solid. LCMS $(MS+1)^+$ : 307.1, $^1$H NMR (400 MHz, DMSO-$d_6$) δ = 8.31 (d, $J$ = 2.0 Hz, 1H), 8.10 - 8.00 (m, 3H), 7.70 (dd, $J$ = 8.4, 2.0 Hz, 1H), 7.50 (d, $J$ = 8.4 Hz, 1H), 6.90 (d, $J$ = 1.0 Hz, 1H), 6.80 (dd, $J$ = 5.2, 1.6 Hz, 1H), 6.50 (s, 2H). $^{13}$C NMR (101 MHz, DMSO-$d_6$) δ 171.30, 160.97, 157.85, 151.27, 150.40, 132.50, 131.04, 126.01, 121.96, 118.10, 107.68, 104.71.

***Synthesis of C-30***

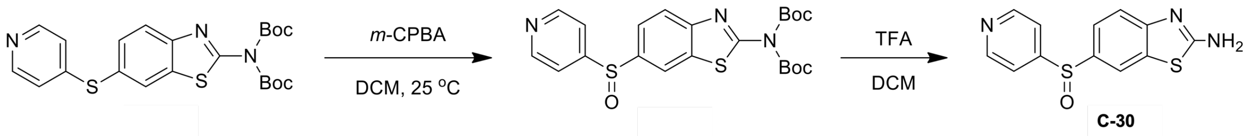


***Tert-butyl N-tert-butoxycarbonyl-N-[6-(4-pyridylsulfinyl)-1,3-benzothiazol-2-yl]carbamate***

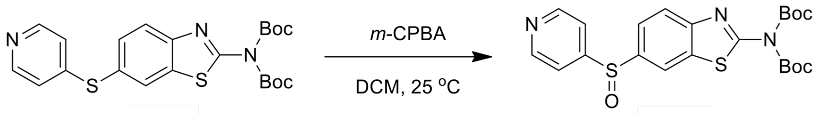


To a solution of tert-butyl N-tert-butoxycarbonyl-N-[6-(4-pyridylsulfanyl)-1,3-benzothiazol-2-yl] carbamate (200 mg, 435 µmol, from **C-25**) in DCM (4 mL) was added *m*-CPBA (93.8 mg, 435 µmol, 80% purity) at 0 °C. Then the mixture was stirred at 25 °C for 3 hr. On completion, the reaction was poured into saturated $Na_2S_2O_3$ aqueous. The organic layer was separated and the aqueous phase was extracted with EA (3 x 20 mL). The combined organic layers were dried over $Na_2SO_4$, filtered and concentrated in vacuo. The residue was purified by column chromatography (PE:EA = 10:1) to give the title compound (160 mg, 70% yield) as a white solid. LCMS $(MS+1)^+$: 476.1. $^1$H NMR (400MHz, $CDCl_3$) δ = 8.72 (d, $J$ = 6.0 Hz, 2H), 8.17 (d, $J$ = 1.6 Hz, 1H), 7.86 (d, $J$ = 8.4 Hz, 1H), 7.63 (dd, $J$ = 1.6, 8.4 Hz, 1H), 7.56 (d, $J$ = 6.0 Hz, 2H), 1.61 (s, 18H).

***6-(4-Pyridylsulfinyl)-1,3-benzothiazol-2-amine (C-30)***

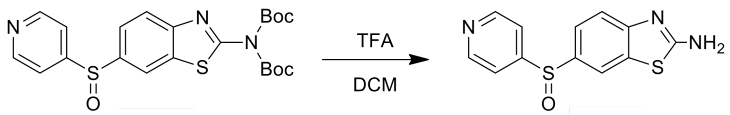


C-30

To a solution of tert-butyl N-tert-butoxycarbonyl-N-[6-(4-pyridylsulfinyl)-1,3-benzothiazol-2-yl] carbamate (130 mg, 273 µmol) in DCM (2 mL) was added TFA (1 mL). Then the mixture was stirred at 25 °C for 0.5 hr. On completion, the reaction was concentrated in vacuo and diluted with water (50 mL). The mixture was adjusted pH to 10 with 30% $NH_3$•$H_2O$ aqueous and extracted with EA (3 x 20 mL). The combined organic layer was concentrated in vacuo. The residue was triturated with EtOH (10 mL) to give the title compound (46.3 mg, 59% yield) as off-white solid. LCMS $(MS+1)^+$: 275.9. $^1$H NMR (400 MHz, DMSO-*d*$_6$) δ = 8.71 (d, *J* = 6.0 Hz, 2H), 8.11 (d, *J* = 2.0 Hz, 1H), 7.86 (s, 2H), 7.69 - 7.63 (d, *J* = 6.0 Hz, 2H), 7.55 (dd, *J* = 2.0, 8.4 Hz, 1H), 7.42 (d, *J* = 8.4 Hz, 1H). $^{13}$C NMR not obtained due to lack of material.

***Synthesis of C-31***

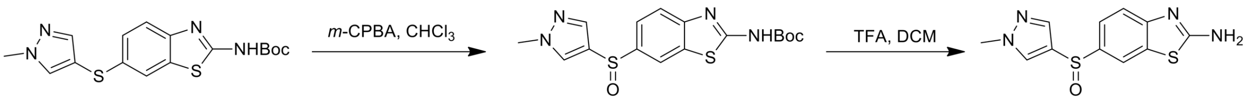


C-31

***Tert-butyl (6-((1-methyl-1H-pyrazol-4-yl)sulfinyl)benzo[d]thiazol-2-yl)carbamate***

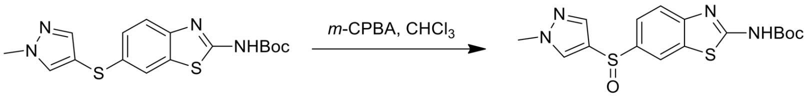


To a solution of tert-butyl N-[6-(1-methylpyrazol-4-yl)sulfanyl-1,3-benzothiazol-2-yl]carbamate (300 mg, 827 µmol, from **C-27**) in $CHCl_3$ (3 mL) was added *m*-CPBA (178 mg, 827 µmol, 80% purity) at 0 °C. The reaction mixture was stirred at 0 °C for 2 hr. On completion, the reaction mixture was diluted with water (10 mL) and extracted with EA (3 x 10 mL). The combined organic layers were washed with brine, dried over $Na_2SO_4$, filtered and concentrated in vacuo. The residue was purified by column chromatography (PE:EA = 2:1) to give the title compound (180 mg, 57% yield) as a yellow solid. LCMS $(MS+1)^+$: 379.0.

***6-((1-Methyl-1H-pyrazol-4-yl)sulfinyl)benzo[d]thiazol-2-amine (C-31)***

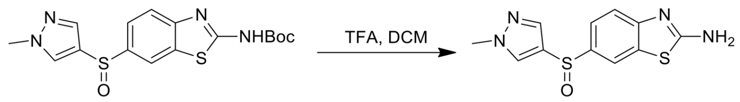

C-31

To a mixture of tert-butyl N-[6-(1-methylpyrazol-4-yl)sulfinyl-1,3-benzothiazol-2-yl]carbamate (180 mg, 475 µmol) in DCM (3 mL) was added TFA (1 mL). The reaction mixture was stirred at 25 °C for 30 min. On completion, the reaction was concentrated in vacuo. The residue was purified by Prep-HPLC (column: Waters Xbridge 150*25 5u;mobile phase: [water(10mM NH4HCO3)-ACN];B%: 7%-30%,7min) to give the title compound (69.6 mg, 53% yield) as a white solid. LCMS $(MS+1)^+$: 278.9. $^1$H NMR (400 MHz, DMSO-$d_6$) δ = 8.07 (s, 1H), 8.01 (s, 1H), 7.78 (s, 2H), 7.60 (d, $J$ = 0.4 Hz, 1H), 7.49 - 7.41 (m, 2H), 3.83 (s, 3H). $^{13}$C NMR (101 MHz, DMSO-$d_6$) δ 169.07, 155.48, 137.80, 137.06, 132.38, 132.16, 126.64, 122.38, 118.34, 117.94, 39.42.

***Synthesis of C-32***

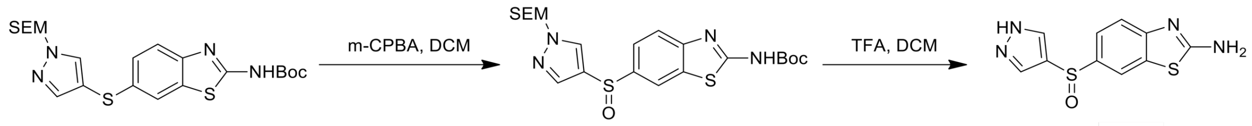

C-32

***Tert-butyl (6-((1-((2-(trimethylsilyl)ethoxy)methyl)-1H-pyrazol-4-yl)sulfinyl)benzo[d] thiazol-2-yl)carbamate***

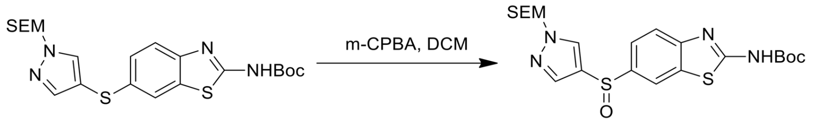

To a solution of tert-butyl N-[6-[1-(2-trimethylsilylethoxymethyl)pyrazol-4-yl]sulfanyl-1,3-benzothiazol-2-yl]carbamate (200 mg, 417 µmol) in DCM (4 mL) was added *m*-CPBA (90.1 mg, 417 µmol, 80% purity) at 0 °C. The reaction mixture was stirred at 25 °C for 3 hr. On completion, the reaction mixture was concentrated in vacuo to give a residue. The residue was purified by reverse phase to give the title compound (160 mg, 77% yield) as a colorless gum. $^1$H NMR (400 MHz, $CDCl_3$) δ = 8.25 (d, $J$ = 1.6 Hz, 1H), 7.88 (d, $J$ = 8.8 Hz, 1H), 7.80 (s, 1H), 7.59 - 7.54 (m, 2H), 5.40 (s, 2H), 3.56 (t, $J$ = 8.0 Hz, 2H), 1.62 (s, 9H), 0.90 (t, $J$ = 8.0 Hz, 2H), -0.03 (s, 9H)

***6-((1H-pyrazol-4-yl)sulfinyl)benzo[d]thiazol-2-amine (C-32)***

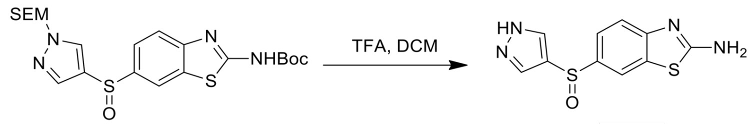

C-32

To a solution of tert-butyl N-[6-[1-(2-trimethylsilylethoxymethyl)pyrazol-4-yl]sulfinyl-1,3-benzothiazol-2-yl]carbamate (110 mg, 222 µmol) in DCM (2 mL) was added TFA (2 mL). The reaction mixture was stirred at 25 °C for 1 hr. On completion, the reaction mixture was concentrated in vacuo. The residue was purified by Prep-HPLC (column: Phenomenex Gemini 150*25mm*10um;mobile phase: [water(0.04%NH3H2O+10mM NH4HCO3)-ACN];B%: 8%-22%,10min) to give the title compound (29.8 mg, 50% yield) as a white solid. LCMS $(MS+1)^+$: 265.1. $^1$H NMR (400 MHz, DMSO-$d_6$) δ = 8.01 (s, 1H), 7.90 (s, 2H), 7.76 (s, 2H), 7.50 - 7.38 (m, 2H). $^{13}$C NMR (101 MHz, DMSO-$d_6$) δ 169.04, 155.41, 137.34, 132.35, 126.24, 122.39, 118.31, 117.94.

***Synthesis of C-37***

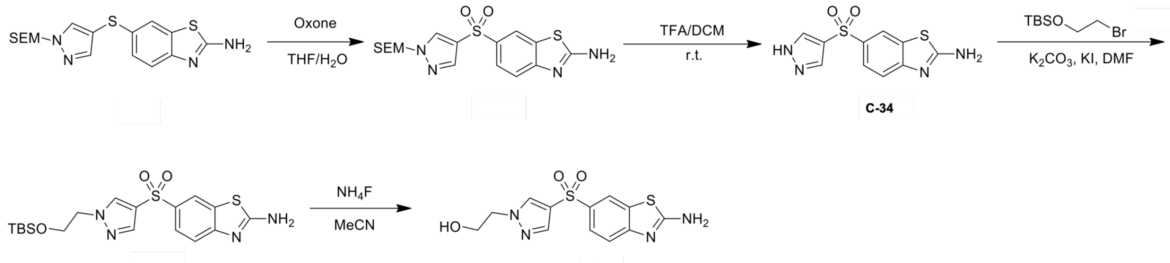

C-34

C-37

***6-[1-(2-Trimethylsilylethoxymethyl)pyrazol-4-yl]sulfonyl-1,3-benzothiazol-2-amine***

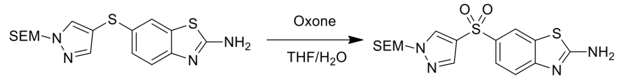

To a solution of 6-[1-(2-trimethylsilylethoxymethyl)pyrazol-4-yl]sulfanyl-1,3-benzothiazol-2-amine (1.90 g, 4.30 mmol) in THF (48 mL) and $H_2O$ (16 mL) was added Oxone (5.30 g, 8.6 mmol) at 0 °C. The mixture was stirred at 25 °C for 4 hr. On completion, the reaction mixture was extracted with EA (3 X 100 mL). The combined organic layer was washed with $H_2O$ (2 X 100 mL) and brine(2 X 100 mL), dried over anhydrous $Na_2SO_4$, filtered and concentrated in vacuo. The residue was purified by silica gel chromatography (PE:MA= 1:1) to give the title compound

(1.50 g, 78% yield) as yellow oil. LCMS $(M+1)^+$: 411.1. $^1H$ NMR (400 MHz, DMSO-$d_6$) δ = 8.63 (s, 1H), 8.33 (s, 1H), 7.99 (s, 1H), 7.77 (d, *J* = 2.0 Hz, 1H), 7.47 (d, *J* = 8.4 Hz, 1H), 5.42 (s, 2H), 3.53 - 3.49 (m, 2H), 0.79 - 0.75 (m, 2H), 0.11 (s, 9H).

***6-(1H-pyrazol-4-ylsulfonyl)-1,3-benzothiazol-2-amine***

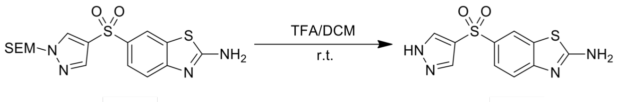


C-34

To the solution of 6-[1-(2-trimethylsilylethoxymethyl)pyrazol-4-yl]sulfonyl-1,3-benzothiazol-2-amine (1.00 g, 2.20 mmol) in DCM (10 mL) was added TFA (5 mL). The reaction mixture was stirred at 25 °C for 2 hr. On completion, add the reaction mixture was poured into ice water, and basified with $Na_2CO_3$ till pH =8. The mixture was extracted with DCM (3 X 30 mL). The organic layer was washed with **$H_2O$** (2 X 30 mL) and brine (2 X 30 mL), dried over **anhydrous $Na_2SO_4$,** filtered and concentrated in vacuo. The residue was purified by gel chromatography (MA) to give the title compound (370 mg, 52% yield) as brown solid. LCMS $(M+1)^+$: 281.1.

***6-[1-[2-[Tert-butyl(dimethyl)silyl]oxyethyl]pyrazol-4-yl]sulfonyl-1,3-benzothiazol-2-amine***

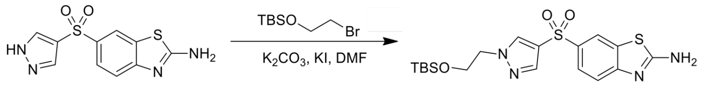


C-34

A mixture of 6-(1H-pyrazol-4-ylsulfonyl)-1,3-benzothiazol-2-amine (170 mg, 594 µmol), 2-bromoethoxy-tert-butyl-dimethyl-silane (170 mg, 713 µmol), $K_2CO_3$ (164 mg, 1.20 mmol) and KI (9.90 mg, 59.4 µmol) in DMF (2 mL) was stirred at 50 °C for 3 hr. On completion, the reaction mixture was diluted with water (10 mL) and extracted with EA (3 X 20 mL). The organic layer was washed with $H_2O$ (2 X 20 mL) and brine (2 X 20 mL), dried over anhydrous $Na_2SO_4$, filtered and concentrated in vacuo. The residue was purified by Prep-TLC (PE:EA = 1:1) to give the title compound (200 mg, 73% yield) as brown solid. LCMS $(M+1)^+$: 439.0.

***2-[4-[(2-Amino-1,3-benzothiazol-6-yl)sulfonyl]pyrazol-1-yl]ethanol (C-37)***

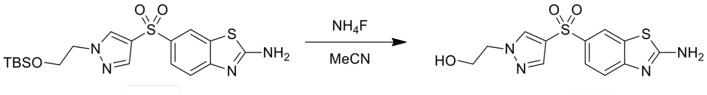


C-37

To a solution of 6-[1-[2-[tert-butyl(dimethyl)silyl]oxyethyl]pyrazol-4-yl]sulfonyl-1,3-benzothiazol -2-amine (180 mg, 385 μmol) in MeCN (5 mL) was add $NH_4F$ (142 mg, 3.80 μmol). The mixture was stirred at 80 °C for 6 hr. On completion, the reaction mixture was diluted with water (10 mL) and extracted with EA (3 X 20 mL). The organic layer was washed with $H_2O$ (2 X 20 mL) and brine (2 X 20 mL), dried over anhydrous $Na_2SO_4$, filtered and concentrated in vacuo. The residue was purified by Prep-TLC (DCM:MeOH = 10:1) to give the title compound (34.3 mg, 27% yield) as light yellow solid. LCMS $(M+1)^+$: 324.9. $^1H$ NMR (400 MHz, DMSO-$d_6$) δ = 8.35 (s, 1H), 8.28 (d, $J$ = 2.0 Hz, 1H), 7.98 (s, 2H), 7.90 (s, 1H), 7.73 (d, $J$ = 8.4 Hz, 1H), 7.42 (d, $J$ = 8.4 Hz, 1H), 4.94 (t, $J$ = 5.2 Hz, 1H), 4.15 (t, $J$ = 5.2 Hz, 2H), 3.71 (d, $J$ = 5.2 Hz, 2H). $^{13}C$ NMR (101 MHz, DMSO-$d_6$) δ 170.68, 157.02, 138.67, 134.73, 133.42, 132.11, 124.88, 124.35, 120.72, 117.91, 59.81, 55.16.

***Synthesis of C-38***

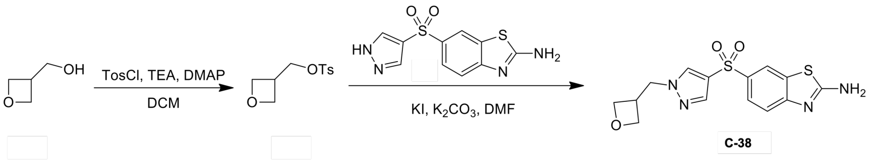


***Oxetan-3-ylmethyl 4-methylbenzenesulfonate***

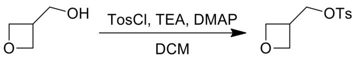


To a solution of oxetan-3-ylmethanol (2.00 g, 22.7 mmol), TosCl (4.30 g, 22.7 mmol) in DCM (30 mL) was added DMAP (277 mg, 2.30 mmol) and TEA (6.90 g, 68.1 mmol). The reaction mixture was stirred at 25 °C for 3 hr. On completion, the reaction mixture was diluted with DCM (3 X 200 mL) and $H_2O$ (2 X 200 mL). The organic layer was washed with brine (2 X 200 mL), dried over $Na_2SO_4$, filtered and concentrated in vacuo. The residue was purified by gel chromatography (PE :EA= 5:1) to give the title compound (3.20 g, 58% yield) as light yellow oil. $^1H$ NMR (400 MHz, DMSO-$d_6$) δ = 7.80 (d, $J$ = 8.4 Hz, 2H), 7.49 (d, $J$ = 8.0 Hz, 2H), 4.58 - 4.54 (m, 2H), 4.24 - 4.18 (m, 4H), 3.33 - 3.21 (m, 1H), 2.43 (s, 3H).

***6-[1-(Oxetan-3-ylmethyl)pyrazol-4-yl]sulfonyl-1,3-benzothiazol-2-amine (C-38)***

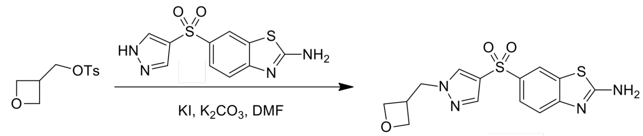


C-38

A mixture of 6-(1H-pyrazol-4-ylsulfonyl)-1,3-benzothiazol-2-amine (from **C-51**, 98.9 mg, 328 µmol), oxetan-3-ylmethyl 4-methylbenzenesulfonate (95.1 mg, 361.0 µmol), $K_2CO_3$ (90.7 mg, 656 µmol), KI (5.50 mg, 32.8 µmol) in DMF (2 mL) was stirred at 50 °C for 3 hr. On completion, the reaction mixture was concentrated in vacuo. The residue was purified by Prep-HPLC (column: Waters Xbridge BEH C18 100*25mm*5um; mobile phase: [water (10 mM $NH_4HCO_3$)-ACN];B%: 5%-35%,8min) to give the title compound (56.4 mg, 49% yield) as light yellow solid. LCMS $(M+1)^+$: 351.1. $^1H$ NMR (400 MHz, DMSO-$d_6$) δ = 8.47 (s, 1H), 8.27 (s, 1H), 7.99 (s, 2 H), 7.91 (s, 1H), 7.73 (d, $J$ = 10.4 Hz, 1H), 7.43 (d, $J$ = 8.4 Hz, 1H), 4.62 - 4.59 (m, 2H), 4.44 (d, $J$ = 7.6 Hz, 2H), 4.38 - 4.35 (m, 2H), 3.38 - 3.35 (m, 1 H). $^{13}C$ NMR (101 MHz, DMSO-$d_6$) δ 170.72, 157.07, 139.03, 134.55, 133.10, 132.12, 124.88, 124.66, 120.73, 117.94, 73.94, 54.30, 35.04.

**Compound ID: C-1**

EW8457-344-P1A DMSO Bruker_B_400MHz

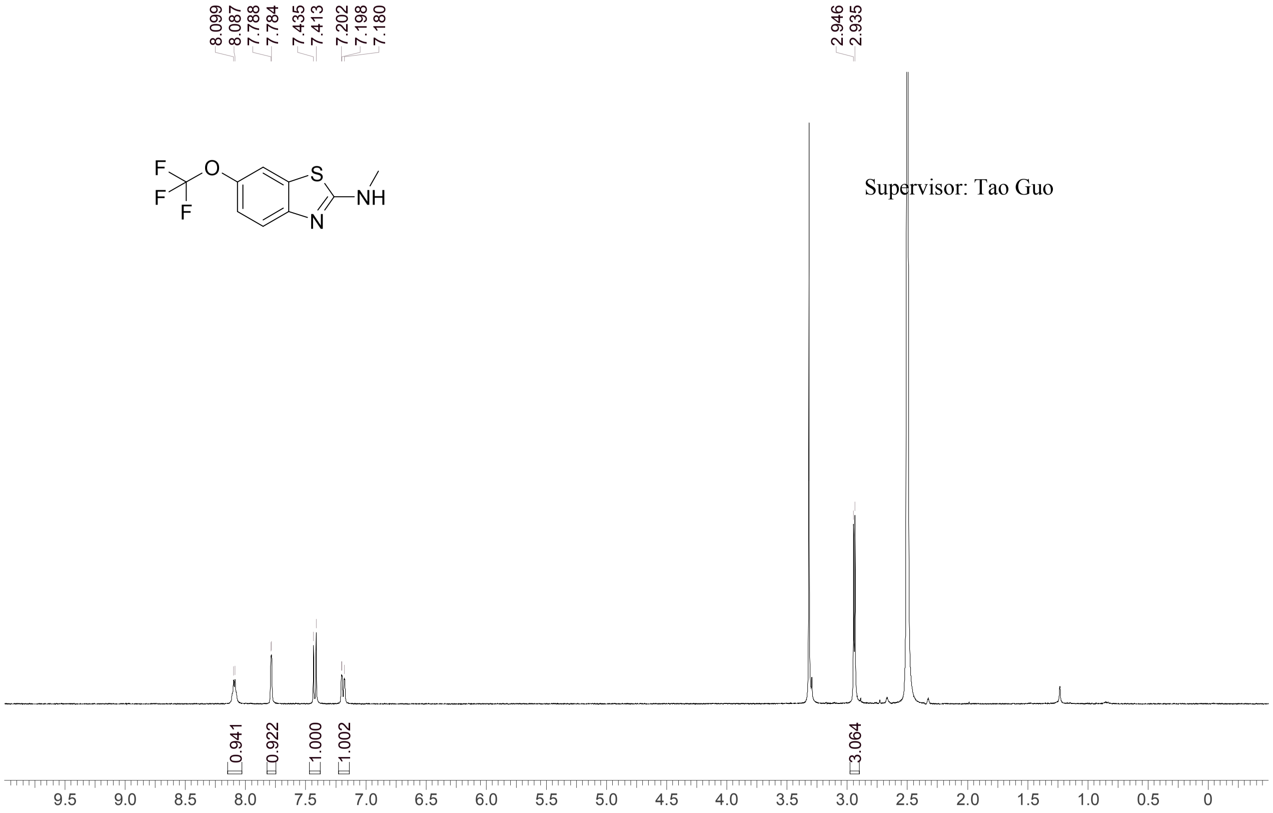


| | |
|---|---|
| Acquisition Time (sec) | 3.0671 |
| Comment | EW8457-344-P1A DMSO Bruker_B_400MHz |
| Date | 10 Apr 2018 15:19:24 |
| Frequency (MHz) | 400.1300 |
| Nucleus | 1H |
| Number of Transients | 8 |
| Origin | spect |
| Original Points Count | 24576 |
| Owner | nmrsu |
| Points Count | 65536 |
| Pulse Sequence | zg30 |
| Receiver Gain | 190.49 |
| SW(cyclical) (Hz) | 8012.82 |
| Solvent | DMSO-d6 |
| Spectrum Offset (Hz) | 2397.5386 |
| Spectrum Type | standard |
| Sweep Width (Hz) | 8012.70 |
| Temperature (degree C) | 25.352 |

Operator: Date:

Confidential, for research only not for regulatory filing

Confidential, for research only not for regulatory filing

### LCMS REPORT

Compound ID : C-1
Sample ID : EW8457-344-P1B
Injection Vol : 1ul
Location : vial87
Acq Method : d:\method\5-95AB_R_220&254.lcm
Org DataFile : D:\DATA\1804\180410\EW8457-344-P1B.lcd
Injection Date : 4/10/2018 3:23:46 PM
Instrument : LCMS-H 15-105

Chromatogram

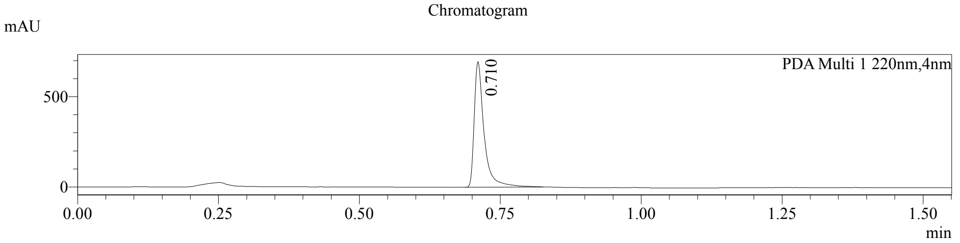


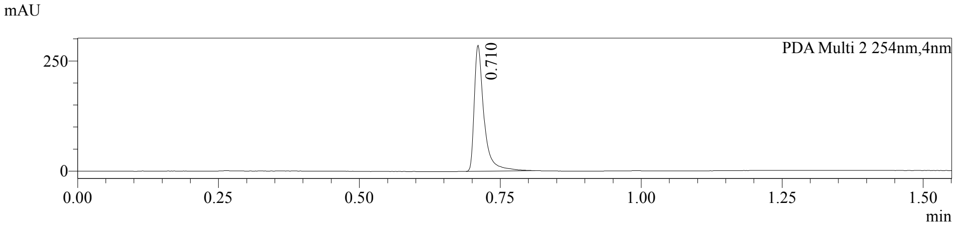


1 PDA Multi 1 / 220nm,4nm
2 PDA Multi 2 / 254nm,4nm

MS Chromatogram

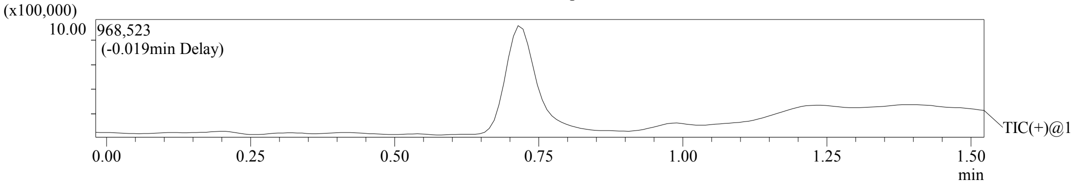


Integration Result

Peak Table

PDA Ch1 220nm

| Peak# | Ret. Time | Height | Height% | USP Width | Area | Area% |
|---|---|---|---|---|---|---|
| 1 | 0.710 | 696147 | 100.000 | 0.031 | 851197 | 100.000 |

Peak Table

PDA Ch2 254nm

| Peak# | Ret. Time | Height | Height% | USP Width | Area | Area% |
|---|---|---|---|---|---|---|
| 1 | 0.710 | 286276 | 100.000 | 0.031 | 345107 | 100.000 |

Confidential, for research only not for regulatory filing

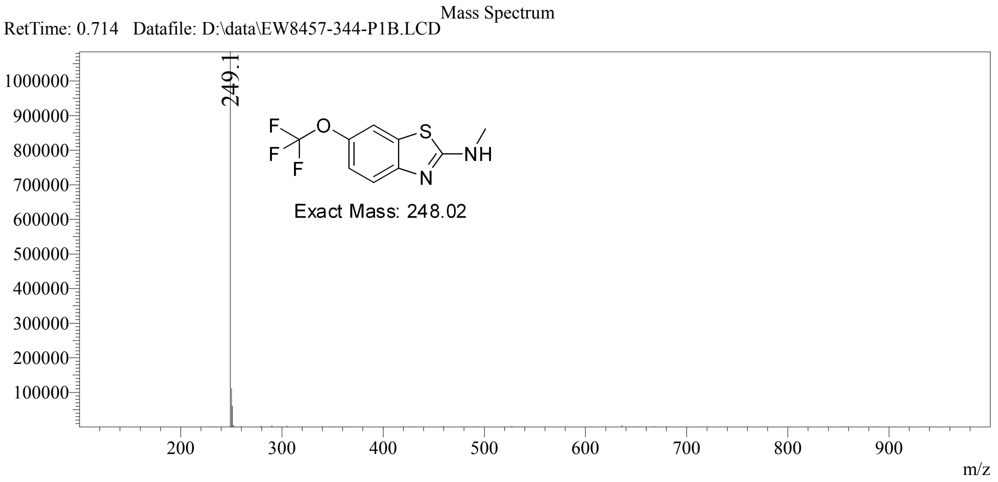

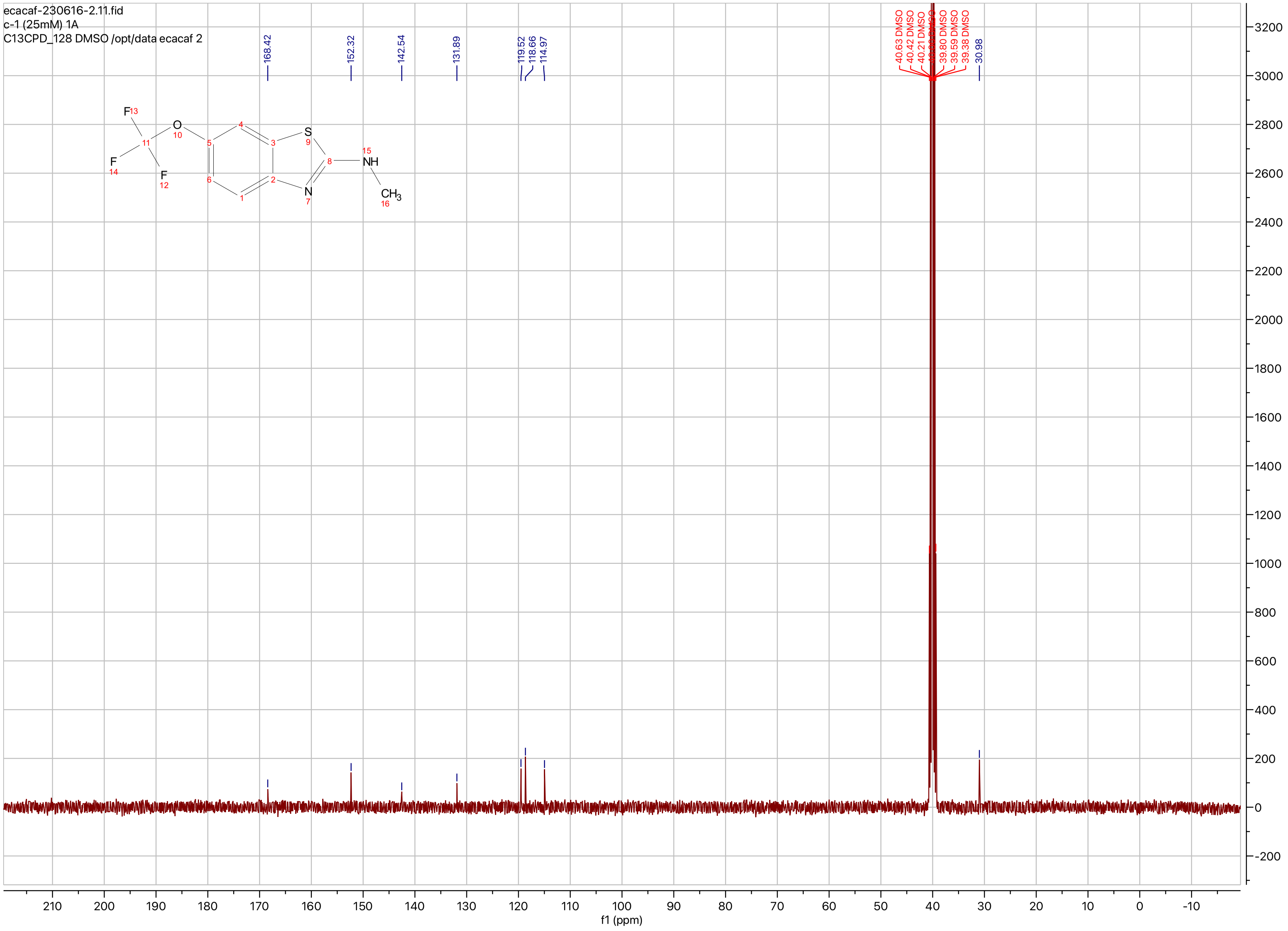

ecacaf-230616-2.11.fid
c-1 (25mM) 1A
C13CPD_128 DMSO /opt/data ecacaf 2
168.42
152.32
142.54
131.89
119.52
118.66
114.97
40.63 DMSO
40.42 DMSO
40.21 DMSO
39.80 DMSO
39.59 DMSO
39.38 DMSO
30.98
f1 (ppm)

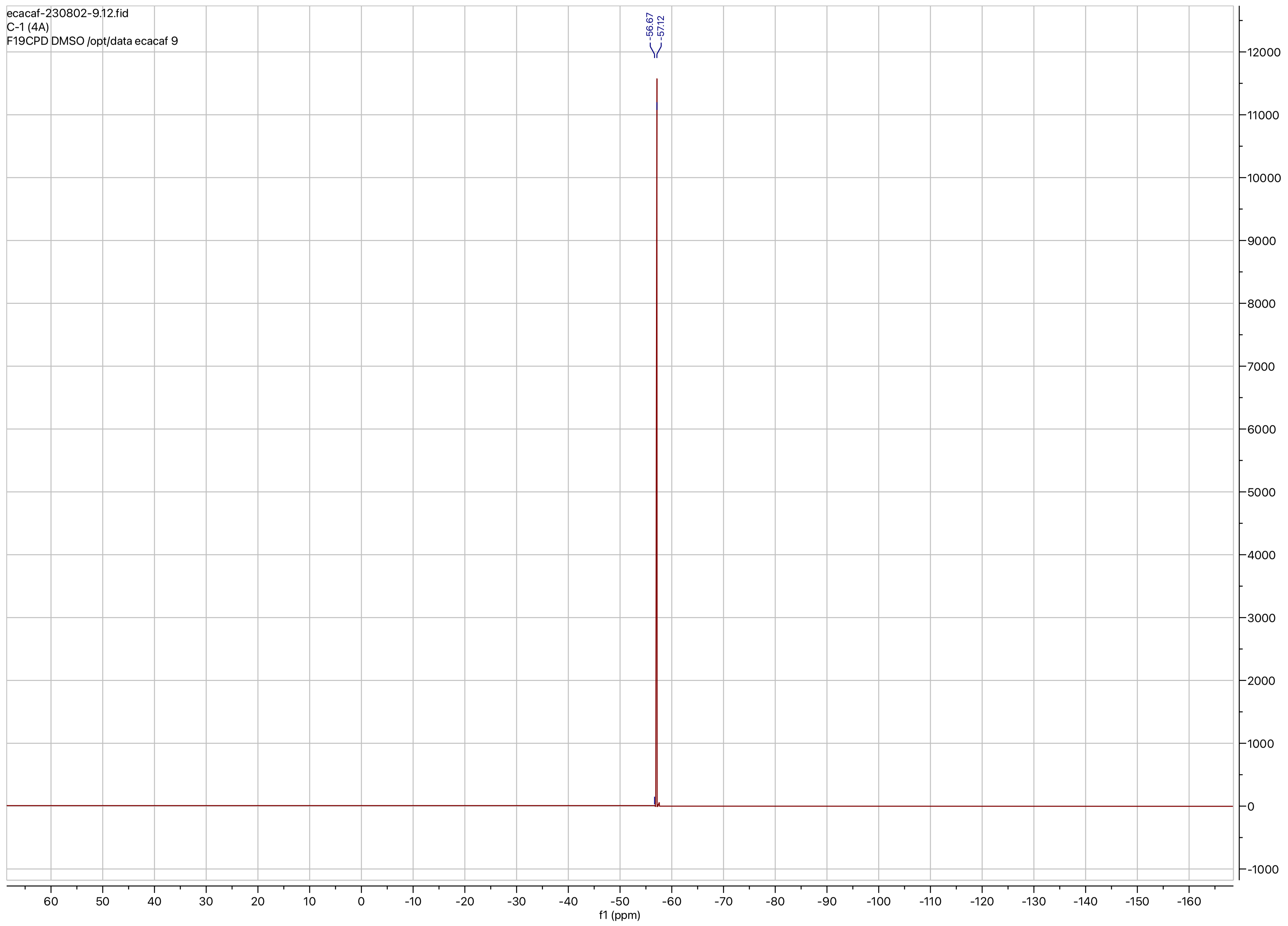

ecacaf-230802-9.12.fid
C-1 (4A)
F19CPD DMSO /opt/data ecacaf 9
-56.67
-57.12
f1 (ppm)

**Compound ID: C-2**

EW8457-352-P1C DMSO Bruker_B_400MHz

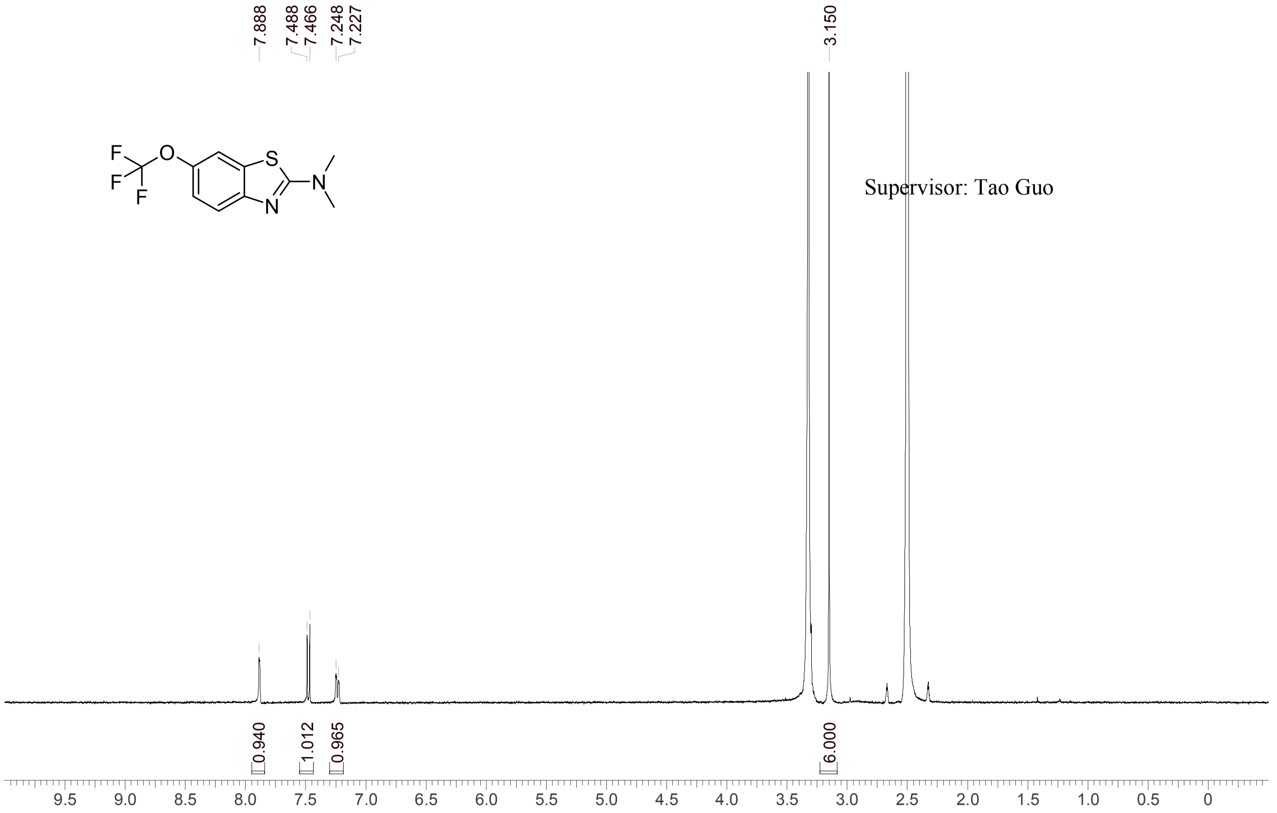


Supervisor: Tao Guo

| | |
|---|---|
| Acquisition Time (sec) | 3.0671 |
| Comment | EW8457-352-P1C DMSO Bruker_B_400MHz |
| Date | 12 Apr 2018 16:36:33 |
| Frequency (MHz) | 400.1300 |
| Nucleus | 1H |
| Number of Transients | 8 |
| Origin | spect |
| Original Points Count | 24576 |
| Owner | nmrsu |
| Points Count | 65536 |
| Pulse Sequence | zg30 |
| Receiver Gain | 190.49 |
| SW(cyclical) (Hz) | 8012.82 |
| Solvent | DMSO-d6 |
| Spectrum Offset (Hz) | 2397.8987 |
| Spectrum Type | standard |
| Sweep Width (Hz) | 8012.70 |
| Temperature (degree C) | 26.475 |

Operator: Date:

Confidential, for research only not for regulatory filing

Confidential, for research only not for regulatory filing

### LCMS REPORT

Compound ID : C-2
Sample ID : EW8457-352-P1B
Injection Vol : 1ul
Location : vial64
Acq Method : d:\method\5-95AB_R_220&254_50.lcm
Org DataFile : D:\DATA\1804\180413\EW8457-352-P1B.lcd
Injection Date : 2018-04-13 9:21:39
Instrument : LCMS-I

Chromatogram

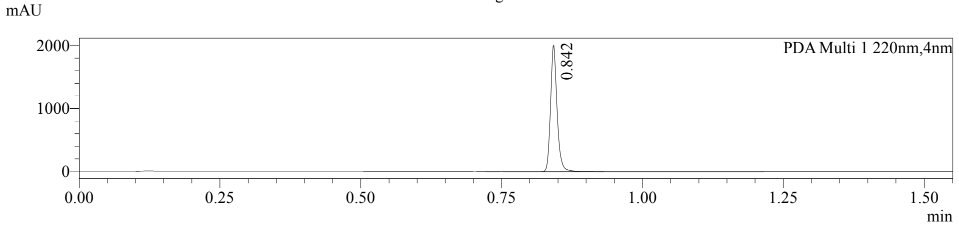


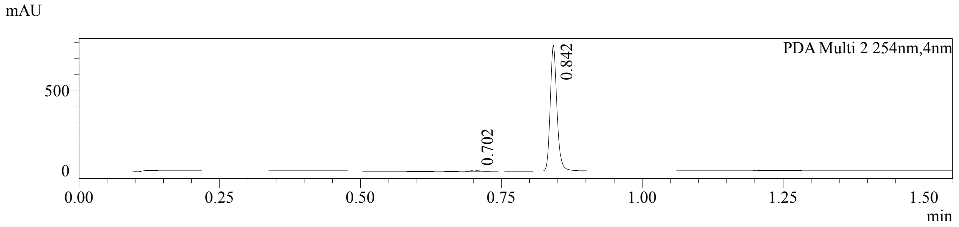


1 PDA Multi 1 / 220nm,4nm
2 PDA Multi 2 / 254nm,4nm

MS Chromatogram

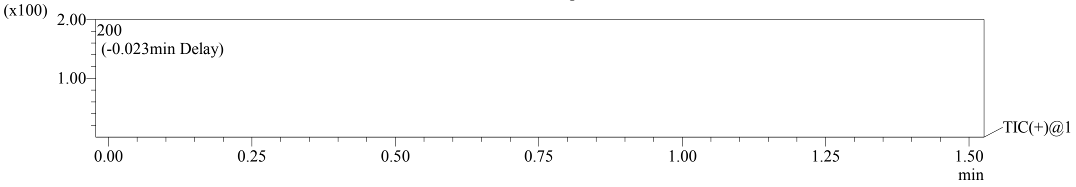


Integration Result

Peak Table

PDA Ch1 220nm

| Peak# | Ret. Time | Height | Height% | USP Width | Area | Area% |
|---|---|---|---|---|---|---|
| 1 | 0.842 | 2009584 | 100.000 | 0.023 | 1635809 | 100.000 |

Peak Table

PDA Ch2 254nm

| Peak# | Ret. Time | Height | Height% | USP Width | Area | Area% |
|---|---|---|---|---|---|---|
| 1 | 0.702 | 8428 | 1.067 | 0.022 | 6658 | 1.008 |
| 2 | 0.842 | 781437 | 98.933 | 0.024 | 653840 | 98.992 |

Operator:______________

PDF created with pdfFactory Pro trial version www.pdffactory.com

Confidential, for research only not for regulatory filing

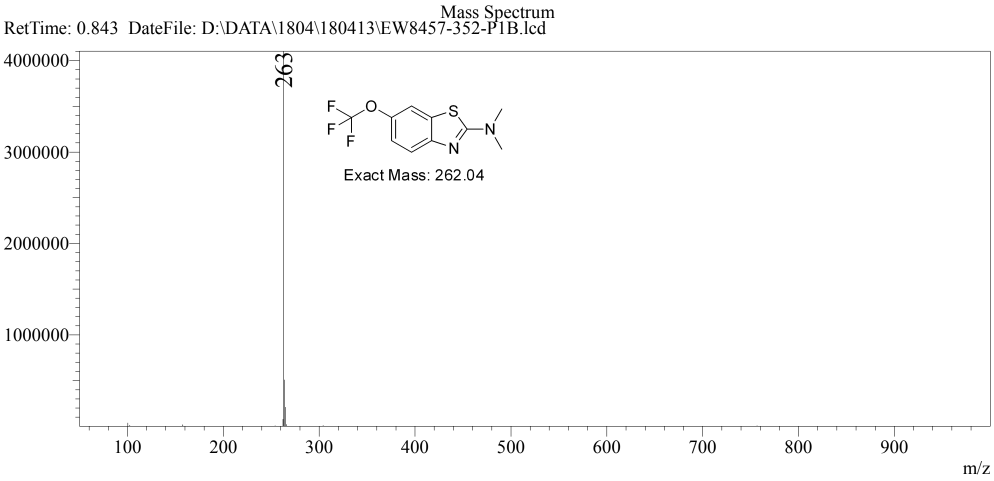


PDF created with pdfFactory Pro trial version www.pdffactory.com

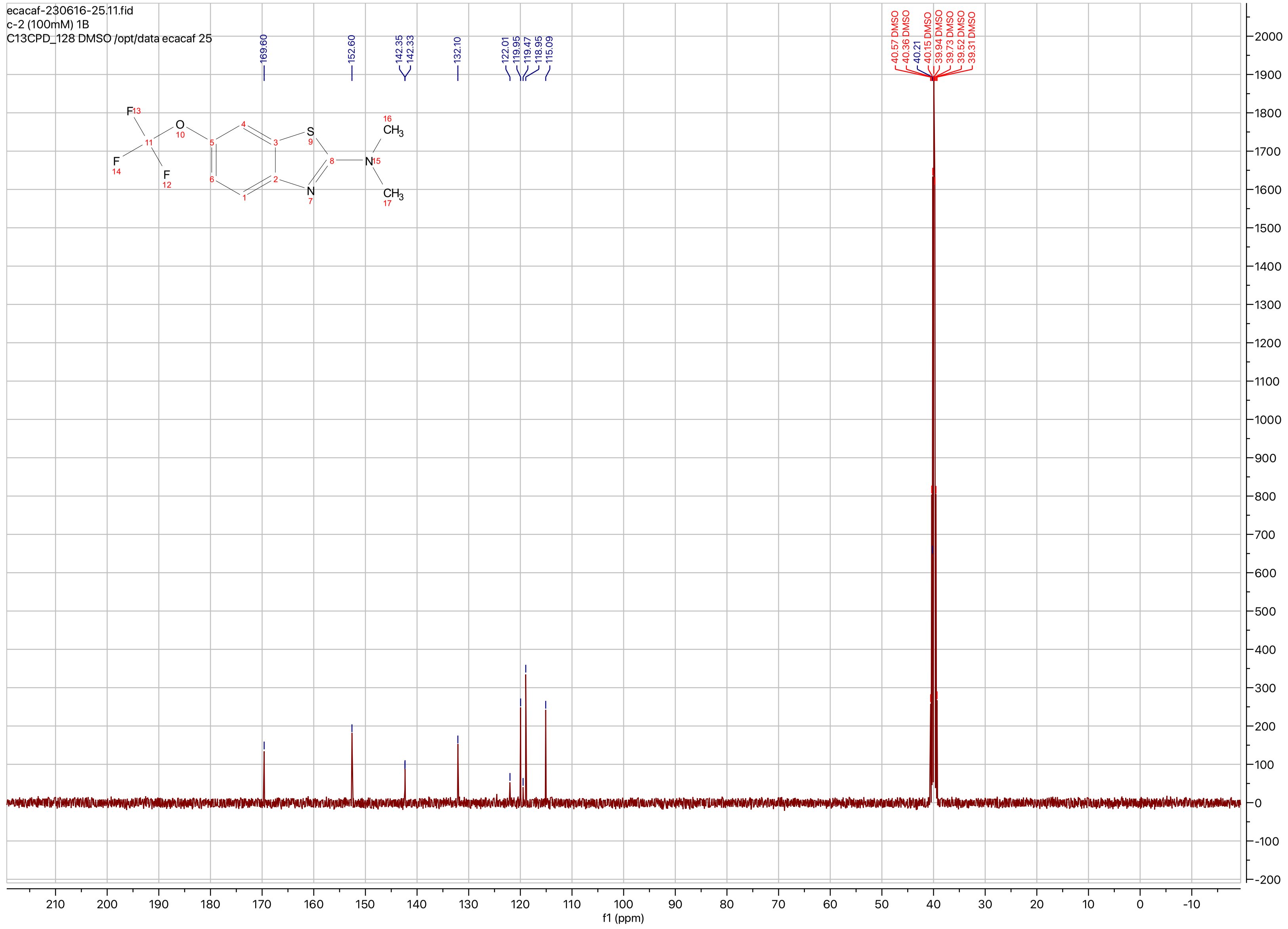

ecacaf-230616-25.11.fid
c-2 (100mM) 1B
C13CPD_128 DMSO /opt/data ecacaf 25
169.60
152.60
142.35
142.33
132.10
122.01
119.95
119.47
118.95
115.09
40.57 DMSO
40.36 DMSO
40.21
40.15 DMSO
39.94 DMSO
39.73 DMSO
39.52 DMSO
39.31 DMSO
f1 (ppm)

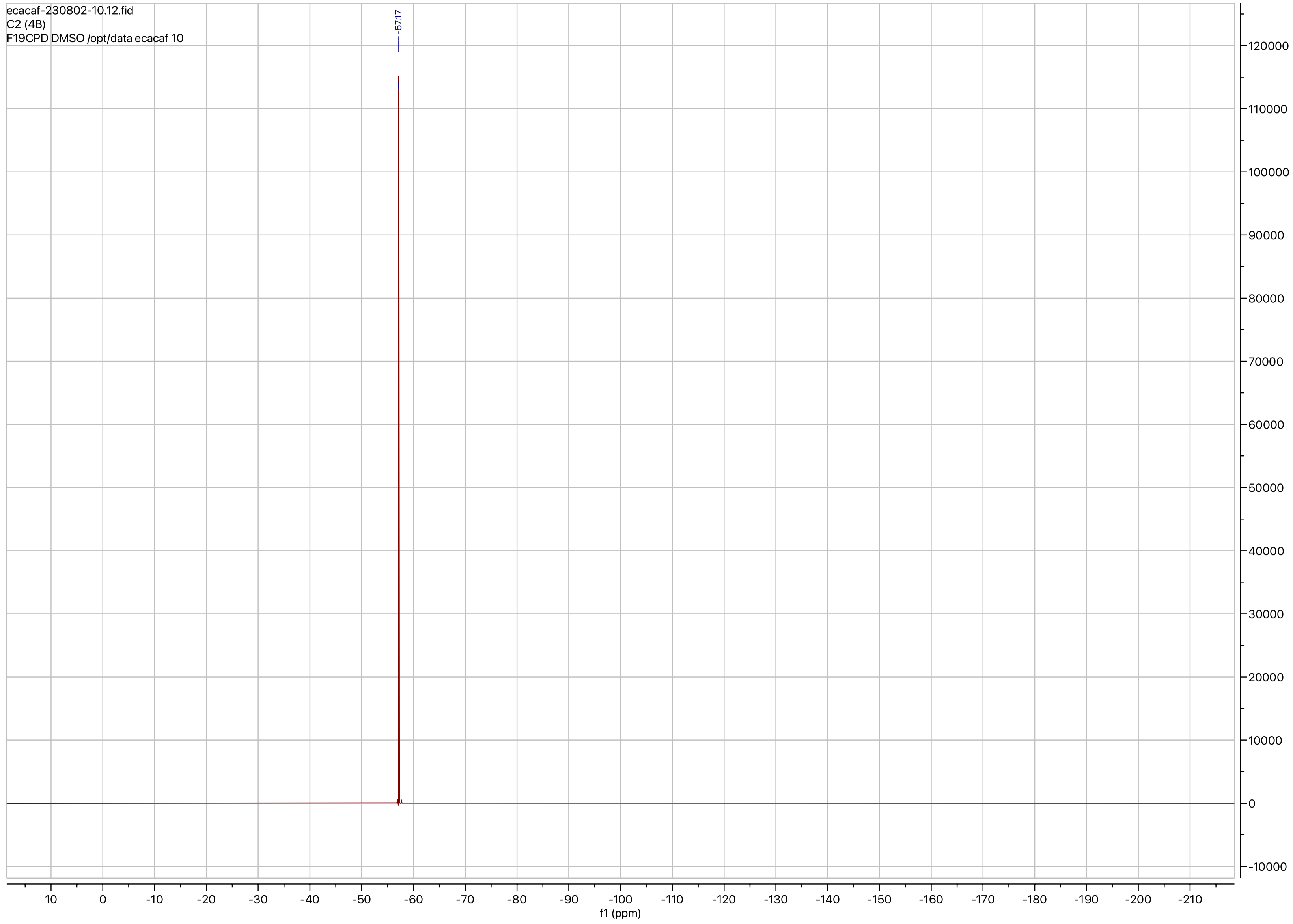
ecacaf-230802-10.12.fid
C2 (4B)
F19CPD DMSO /opt/data ecacaf 10
-57.17
f1 (ppm)

**Compound ID: C-3**

EW8457-336-P1A DMSO Bruker_A_400MHz

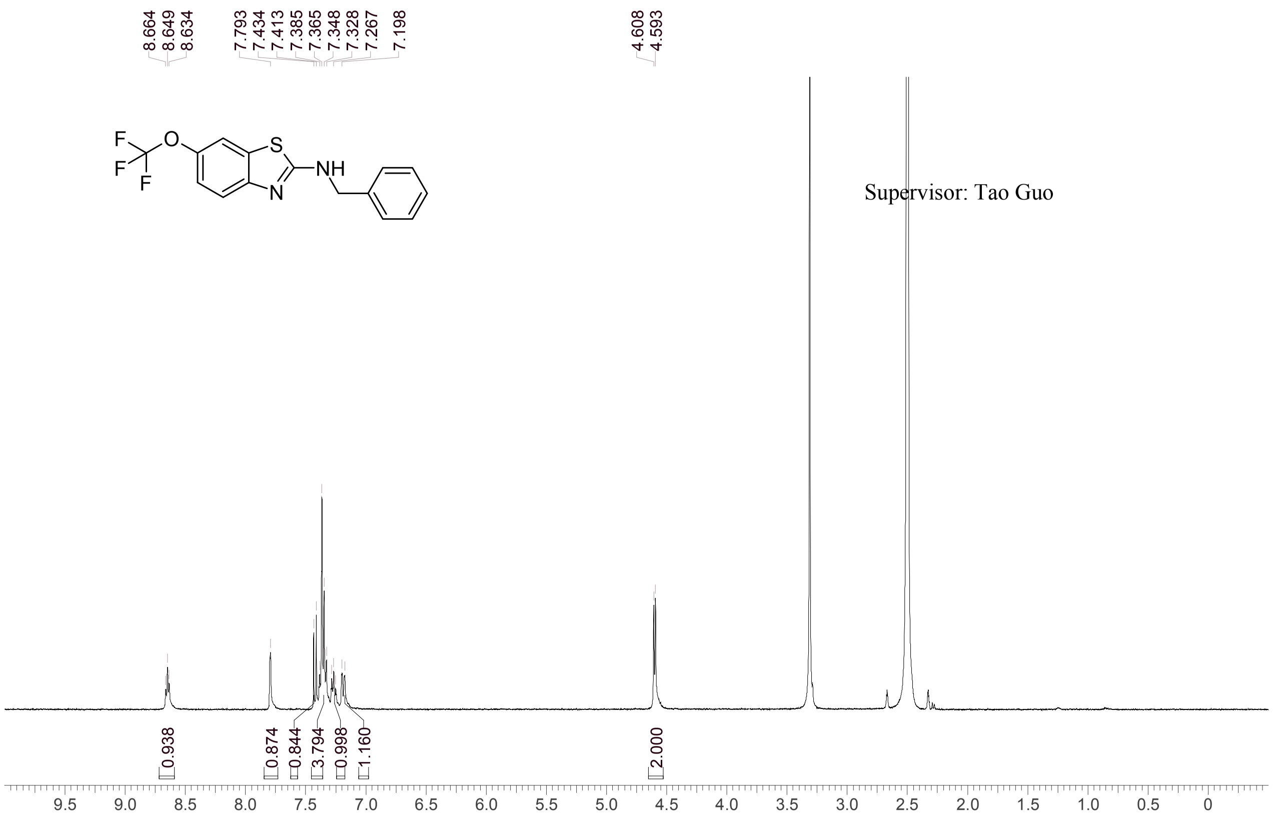


| | |
|---|---|
| Acquisition Time (sec) | 3.0671 |
| Comment | EW8457-336-P1A DMSO Bruker_A_400MHz |
| Date | 09 Apr 2018 17:26:42 |
| Frequency (MHz) | 400.1300 |
| Nucleus | 1H |
| Number of Transients | 8 |
| Origin | spect |
| Original Points Count | 24576 |
| Owner | nmrsu |
| Points Count | 65536 |
| Pulse Sequence | zg30 |
| Receiver Gain | 175.36 |
| SW(cyclical) (Hz) | 8012.82 |
| Solvent | DMSO-d6 |
| Spectrum Offset (Hz) | 2397.2585 |
| Spectrum Type | standard |
| Sweep Width (Hz) | 8012.70 |
| Temperature (degree C) | 27.157 |

Operator:

Date:

Confidential, for research only not for regulatory filing

Confidential, for research only not for regulatory filing

### LCMS REPORT

Compound ID : C-3
Sample ID : EW8457-336-P1B
Injection Vol : 2ul
Location : vial68
Acq Method : d:\method\5-95AB_R_220&254.lcm
Org DataFile : D:\DATA\1804\180409\EW8457-336-P1B.lcd
Injection Date : 2018-04-09 18:15:55
Instrument : LCMS-H 15-105

Chromatogram

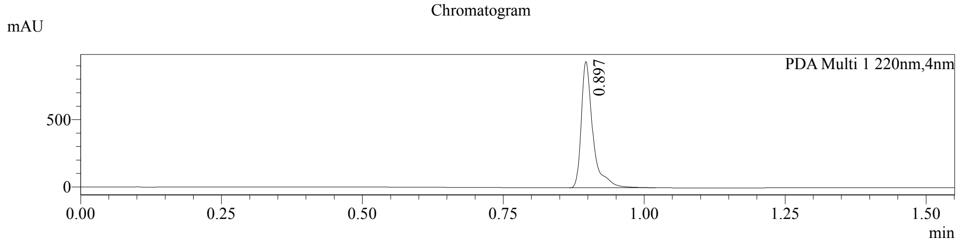


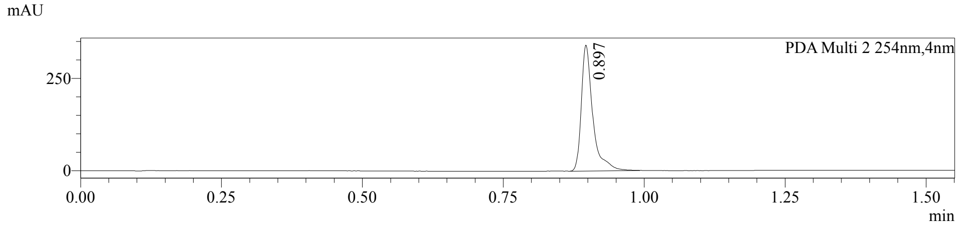


1 PDA Multi 1 / 220nm,4nm
2 PDA Multi 2 / 254nm,4nm

MS Chromatogram

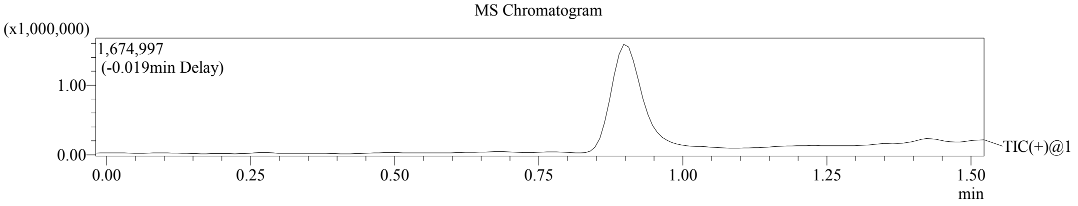


Integration Result

Peak Table

PDA Ch1 220nm

| Peak# | Ret. Time | Height | Height% | USP Width | Area | Area% |
|---|---|---|---|---|---|---|
| 1 | 0.897 | 936947 | 100.000 | 0.036 | 1348187 | 100.000 |

Peak Table

PDA Ch2 254nm

| Peak# | Ret. Time | Height | Height% | USP Width | Area | Area% |
|---|---|---|---|---|---|---|
| 1 | 0.897 | 341173 | 100.000 | 0.036 | 483143 | 100.000 |

Confidential, for research only not for regulatory filing

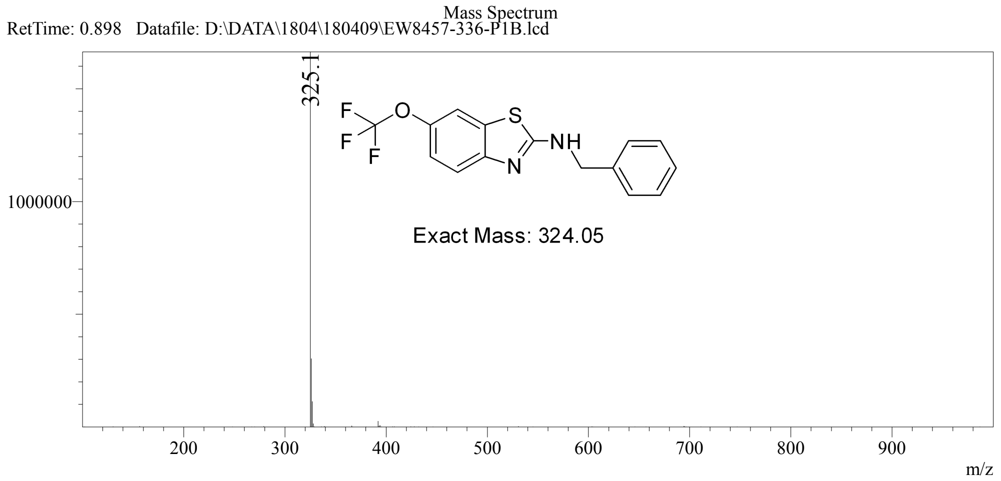

ecacaf-230616-26.11.fid
c-3 (100mM) 1C
C13CPD_128 DMSO /opt/data ecacaf 26

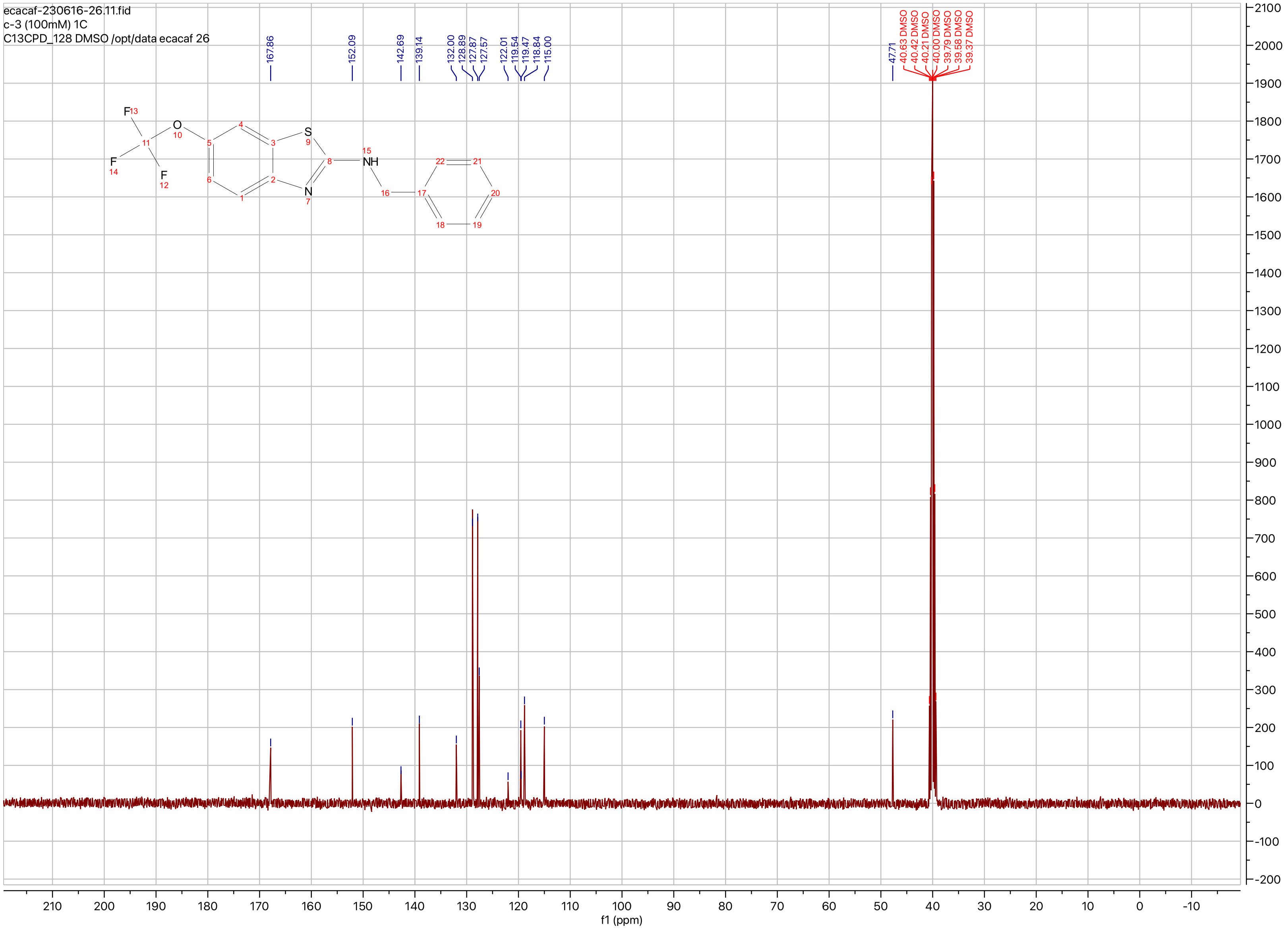

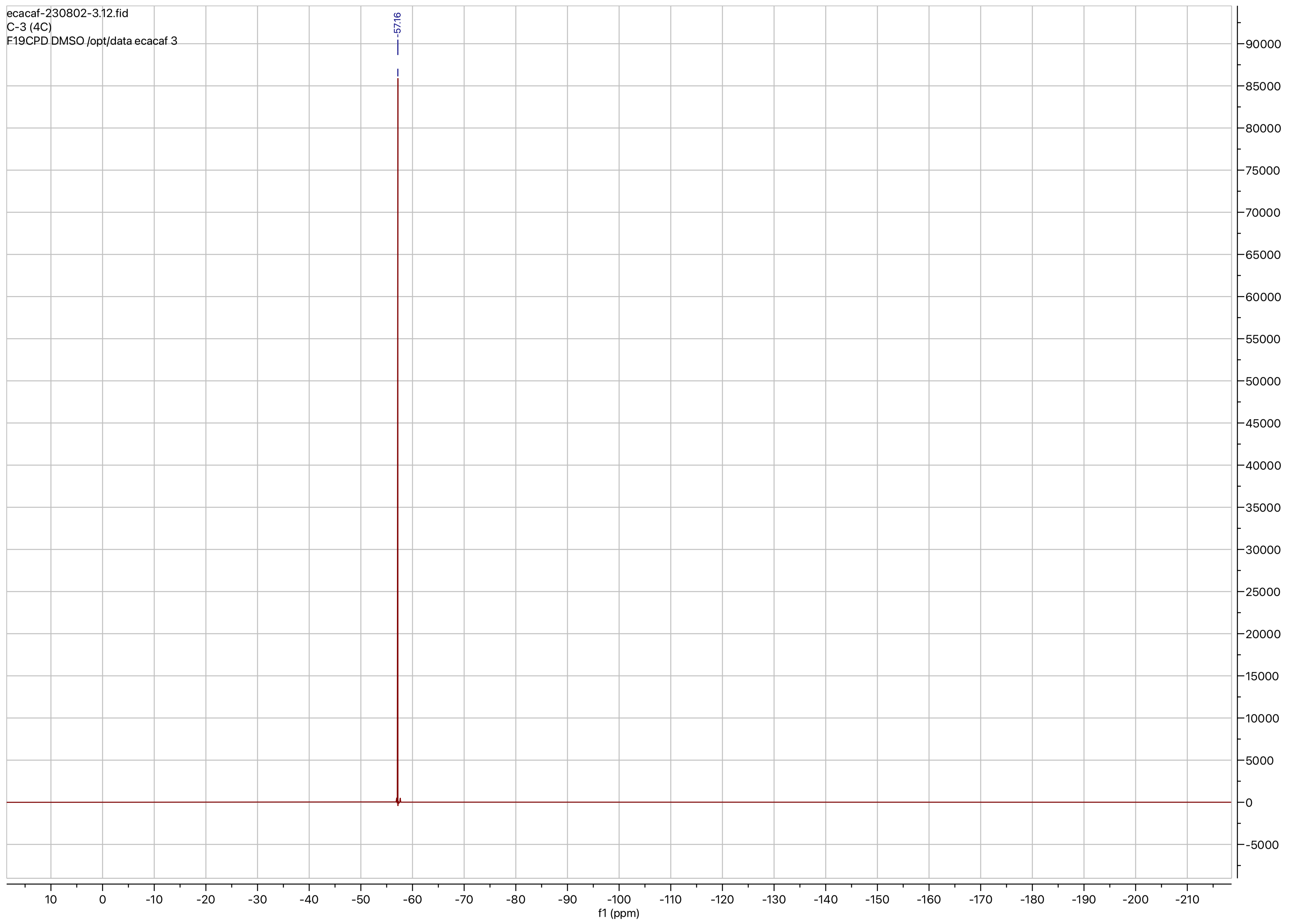
ecacaf-230802-3.12.fid
C-3 (4C)
F19CPD DMSO /opt/data ecacaf 3
-57.16
f1 (ppm)

**Compound ID: C-4**

EW8397-322-P1A CDCl3 Bruker_B_400MHz

Supervisor: Tao Guo

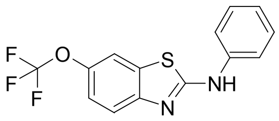


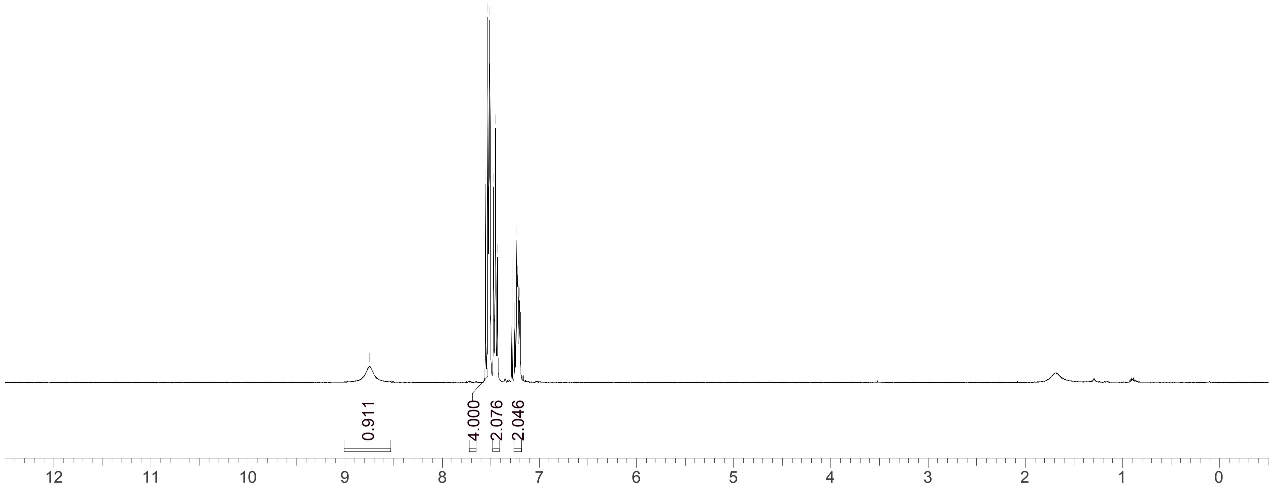


| | |
|---|---|
| Acquisition Time (sec) | 3.0671 |
| Comment | EW8397-322-P1A CDCl3 Bruker_B_400MHz |
| Date | 24 Apr 2018 14:04:16 |
| Frequency (MHz) | 400.1300 |
| Nucleus | 1H |
| Number of Transients | 8 |
| Origin | spect |
| Original Points Count | 24576 |
| Owner | nmrsu |
| Points Count | 65536 |
| Pulse Sequence | zg30 |
| Receiver Gain | 190.49 |
| SW(cyclical) (Hz) | 8012.82 |
| Solvent | CHLOROFORM-d |
| Spectrum Offset (Hz) | 2400.7795 |
| Spectrum Type | standard |
| Sweep Width (Hz) | 8012.70 |
| Temperature (degree C) | 24.610 |

Operator:

Date:

Confidential, for research only not for regulatory filing

Confidential, for research only not for regulatory filing

### LCMS REPORT

Compound ID : C-4
Sample ID : EW8397-322-P1C
Injection Vol : 1ul
Location : vial100
Acq Method : d:\method\5-95CD_R_220&254_POS.lcm
Org DataFile : D:\DATA\1804\180425\EW8397-322-P1C.lcd
Injection Date : 2018-04-25 10:28:24
Instrument :LCMS-O 15-105

Chromatogram

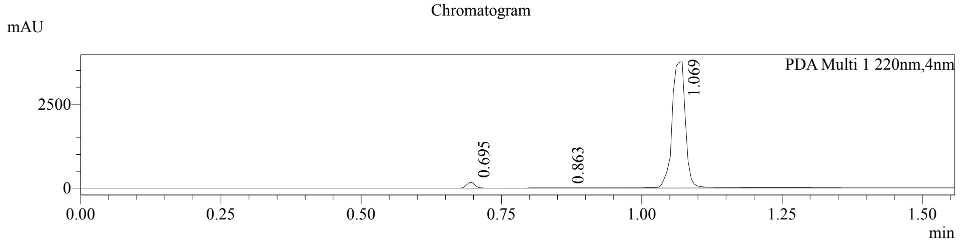


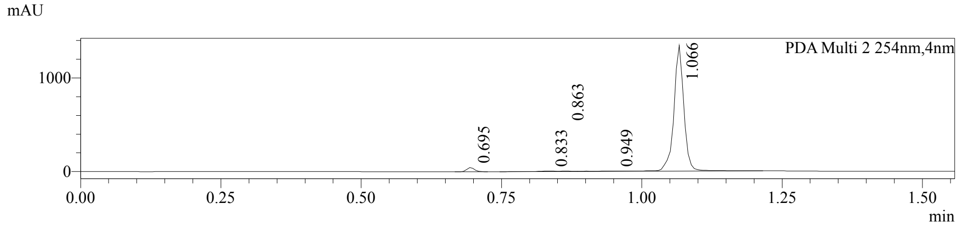


1 PDA Multi 1 / 220nm,4nm
2 PDA Multi 2 / 254nm,4nm

MS Chromatogram

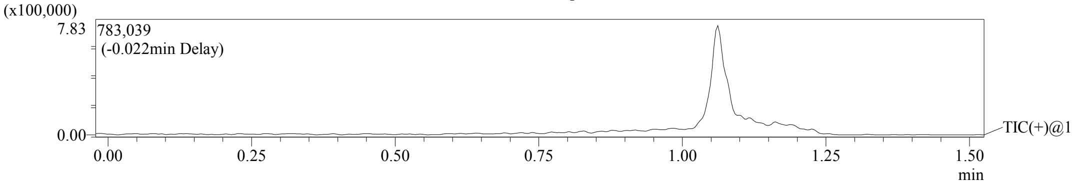


Integration Result

Peak Table

PDA Ch1 220nm

| Peak# | Ret. Time | Height | Height% | USP Width | Area | Area% |
|---|---|---|---|---|---|---|
| 1 | 0.695 | 179676 | 4.558 | 0.029 | 173382 | 2.653 |
| 2 | 0.863 | 6474 | 0.164 | 0.132 | 27267 | 0.417 |
| 3 | 1.069 | 3755807 | 95.278 | 0.308 | 6333535 | 96.929 |

Peak Table

PDA Ch2 254nm

| Peak# | Ret. Time | Height | Height% | USP Width | Area | Area% |
|---|---|---|---|---|---|---|
| 1 | 0.695 | 43201 | 3.090 | 0.029 | 41675 | 2.396 |
| 2 | 0.833 | 6119 | 0.438 | 0.049 | 12799 | 0.736 |
| 3 | 0.863 | 4589 | 0.328 | 0.084 | 7673 | 0.441 |
| 4 | 0.949 | 2274 | 0.163 | 0.214 | 12917 | 0.743 |
| 5 | 1.066 | 1341981 | 95.982 | 0.036 | 1664064 | 95.684 |

Confidential, for research only not for regulatory filing

Mass Spectrum

RetTime: 0.695 Datafile: D:\DATA\1804\180425\EW8397-322-P1C.lcd

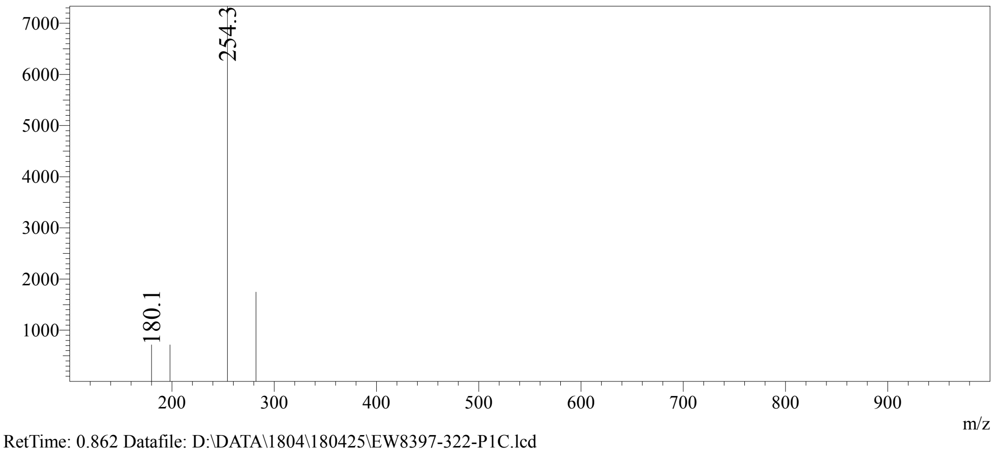


RetTime: 0.862 Datafile: D:\DATA\1804\180425\EW8397-322-P1C.lcd

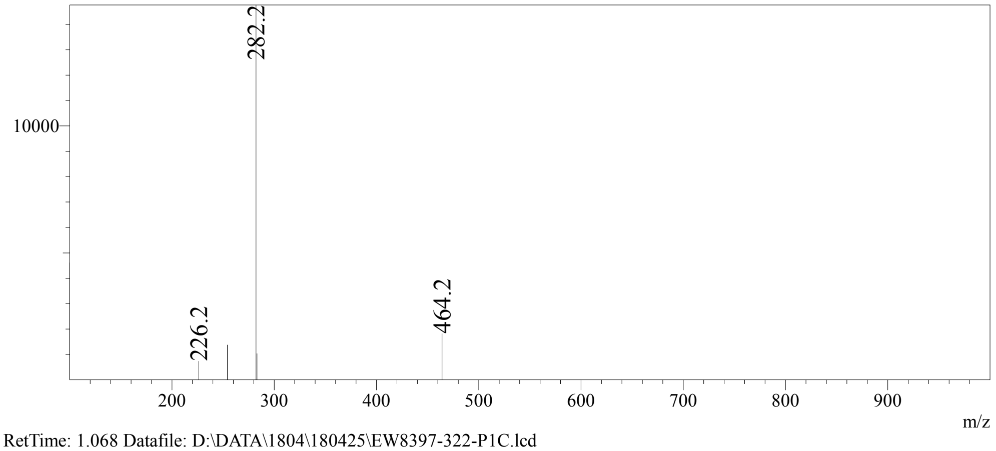


RetTime: 1.068 Datafile: D:\DATA\1804\180425\EW8397-322-P1C.lcd

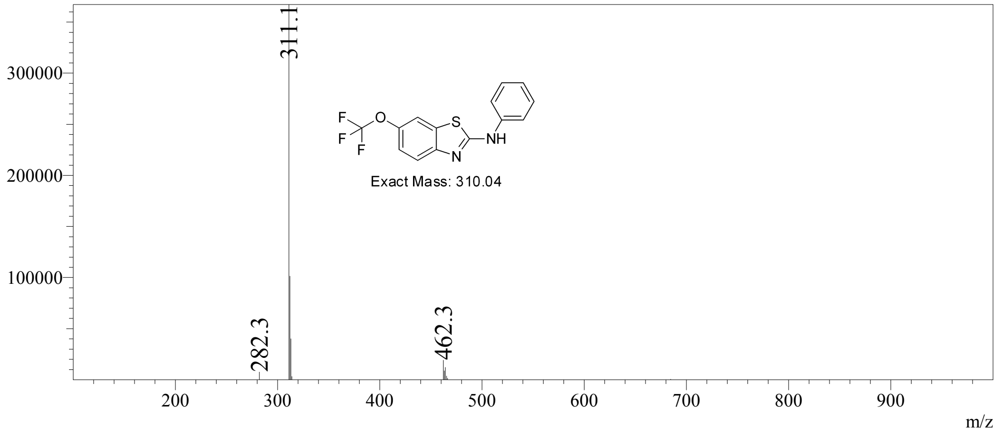

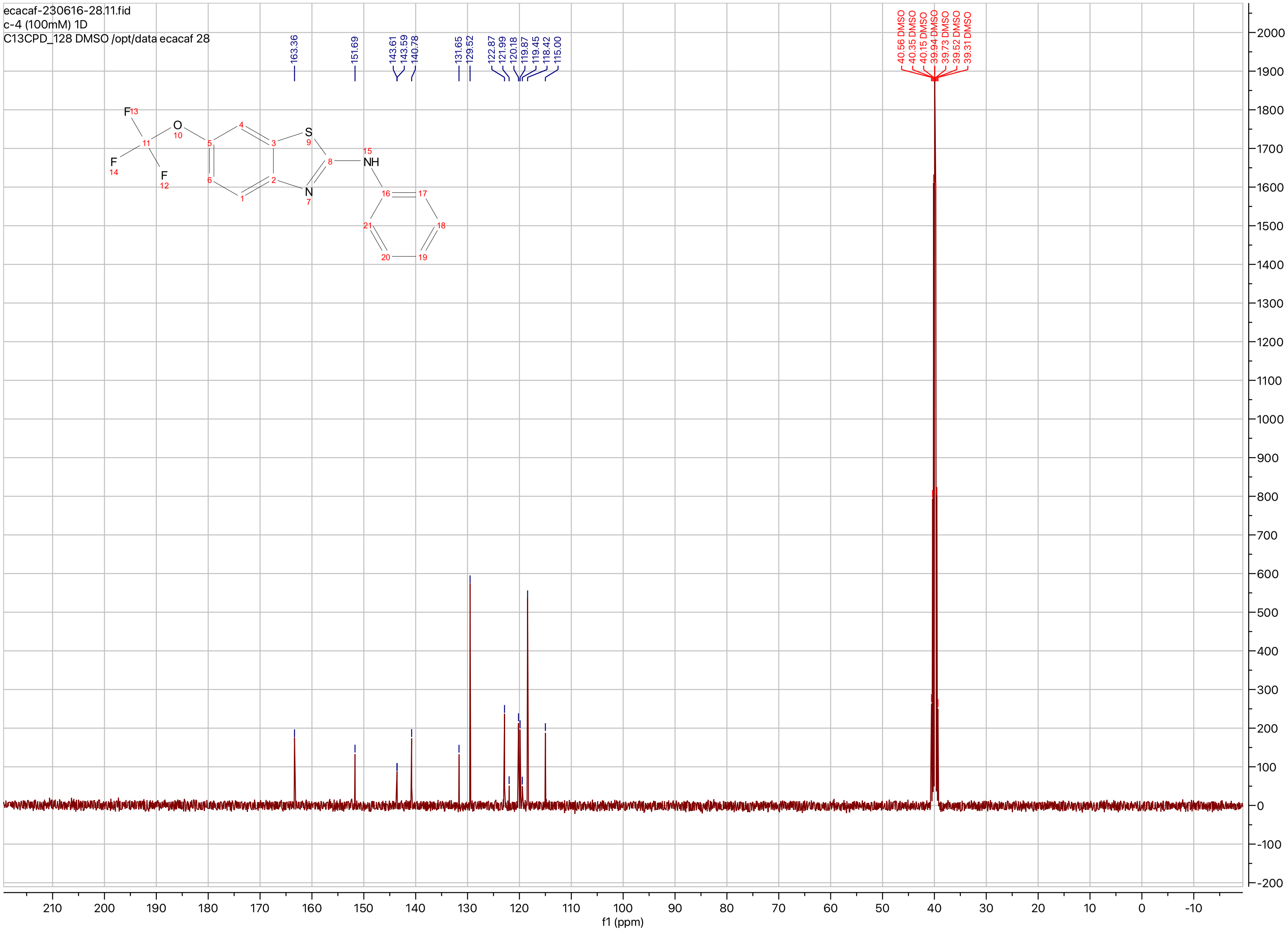
ecacaf-230616-28.11.fid
c-4 (100mM) 1D
C13CPD_128 DMSO /opt/data ecacaf 28
163.36
151.69
143.61
143.59
140.78
131.65
129.52
122.87
121.99
120.18
119.87
119.45
118.42
115.00
40.56 DMSO
40.35 DMSO
40.15 DMSO
39.94 DMSO
39.73 DMSO
39.52 DMSO
39.31 DMSO
f1 (ppm)

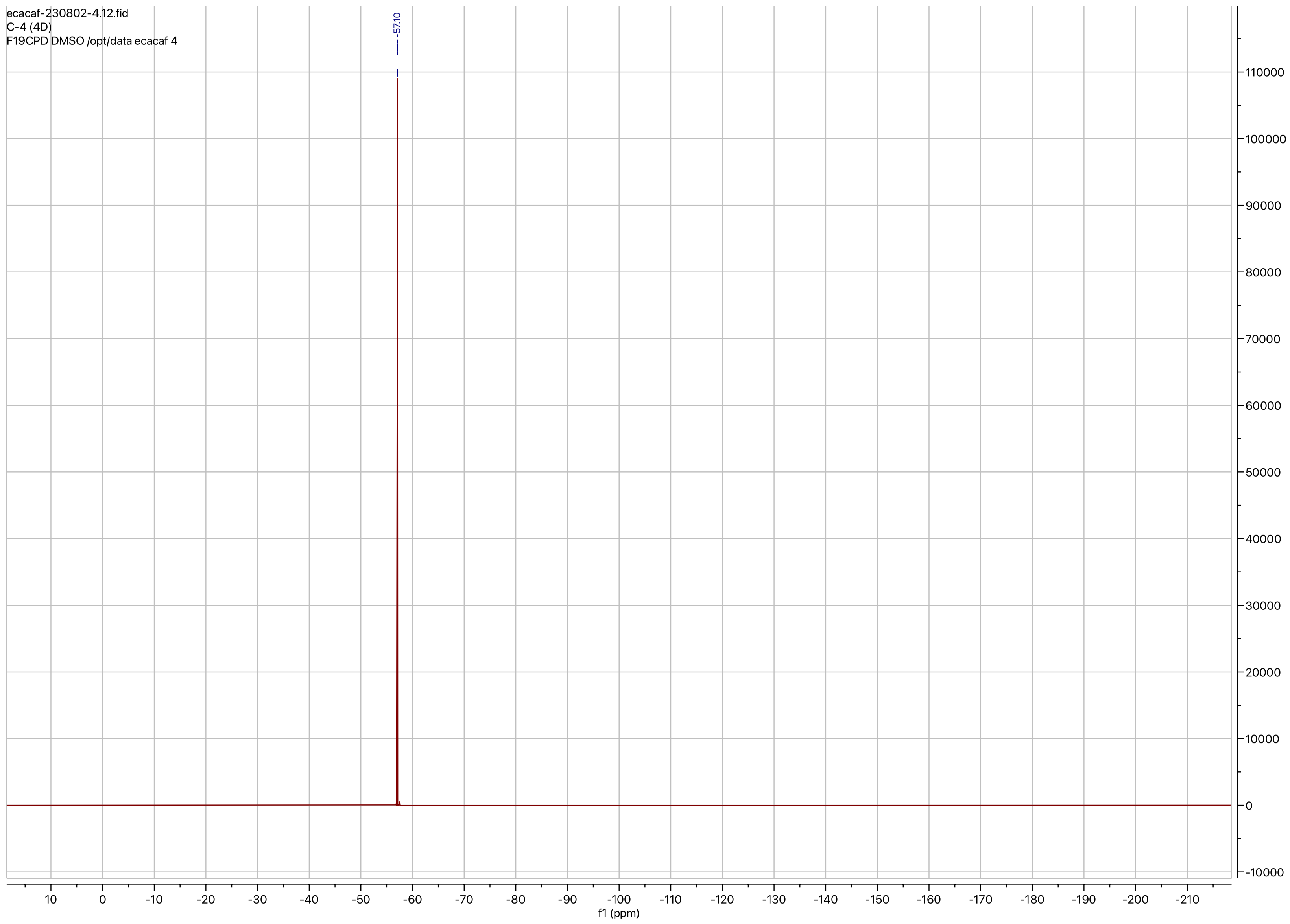

ecacaf-230802-4.12.fid
C-4 (4D)
F19CPD DMSO /opt/data ecacaf 4
-57.10
110000
100000
90000
80000
70000
60000
50000
40000
30000
20000
10000
0
-10000
10
0
-10
-20
-30
-40
-50
-60
-70
-80
-90
-100
-110
-120
-130
-140
-150
-160
-170
-180
-190
-200
-210
f1 (ppm)

**Compound ID: C-5**

EW8457-367-P1A CDCl3 Bruker_A_400MHz

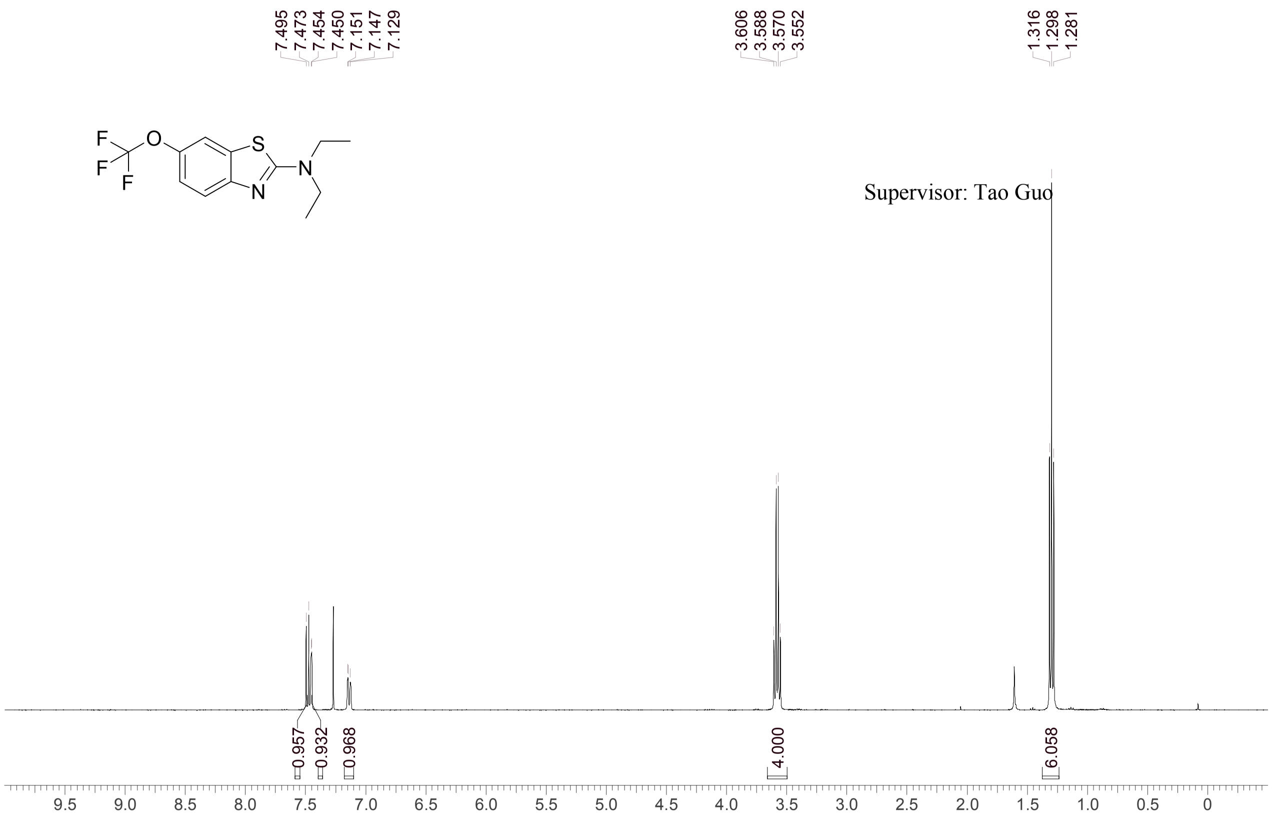


Supervisor: Tao Guo

| | |
|---|---|
| Acquisition Time (sec) | 3.0671 |
| Comment | EW8457-367-P1A CDCl3 Bruker_A_400MHz |
| Date | 19 Apr 2018 15:14:10 |
| Frequency (MHz) | 400.1300 |
| Nucleus | 1H |
| Number of Transients | 8 |
| Origin | spect |
| Original Points Count | 24576 |
| Owner | nmrsu |
| Points Count | 65536 |
| Pulse Sequence | zg30 |
| Receiver Gain | 195.78 |
| SW(cyclical) (Hz) | 8012.82 |
| Solvent | CHLOROFORM-d |
| Spectrum Offset (Hz) | 2394.8623 |
| Spectrum Type | standard |
| Sweep Width (Hz) | 8012.70 |
| Temperature (degree C) | 24.857 |

Operator: Date:

Confidential, for research only not for regulatory filing

Confidential, for research only not for regulatory filing

### LCMS REPORT

Compound ID : C-5
Sample ID : EW8457-367-P1B
Injection Vol : 1ul
Location : vial8
Acq Method : d:\method\5-95AB_R_220&254.lcm
Org DataFile : D:\DATA\1804\180419\EW8457-367-P1B.lcd
Injection Date : 2018-04-19 15:50:23
Instrument : LCMS-I

Chromatogram

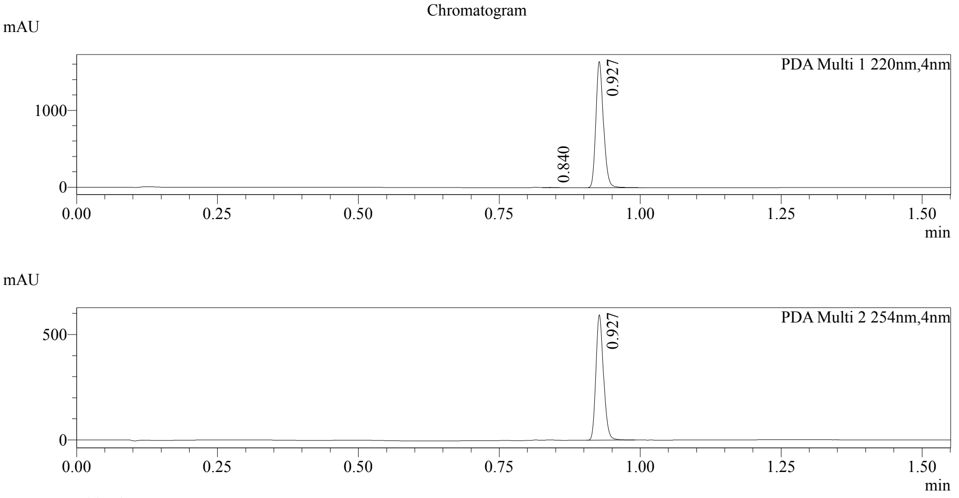


1 PDA Multi 1 / 220nm,4nm
2 PDA Multi 2 / 254nm,4nm

MS Chromatogram

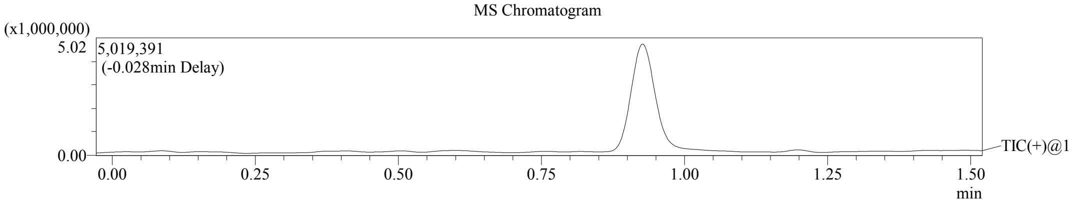


#### Integration Result

Peak Table

PDA Ch1 220nm

| Peak# | Ret. Time | Height | Height% | USP Width | Area | Area% |
|---|---|---|---|---|---|---|
| 1 | 0.840 | 4245 | 0.258 | 0.040 | 5370 | 0.348 |
| 2 | 0.927 | 1640734 | 99.742 | 0.027 | 1539003 | 99.652 |

Peak Table

PDA Ch2 254nm

| Peak# | Ret. Time | Height | Height% | USP Width | Area | Area% |
|---|---|---|---|---|---|---|
| 1 | 0.927 | 594864 | 100.000 | 0.027 | 562460 | 100.000 |

PDF created with pdfFactory Pro trial version www.pdffactory.com

Confidential, for research only not for regulatory filing

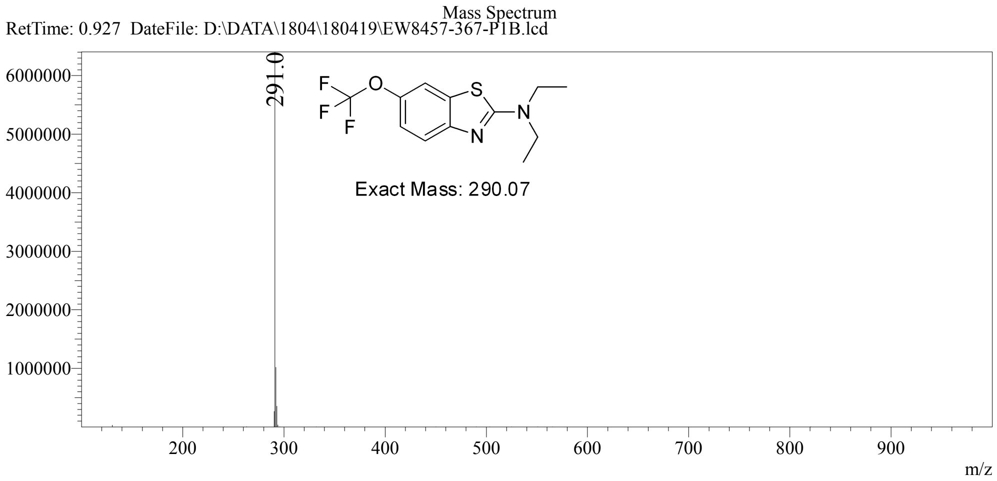


PDF created with pdfFactory Pro trial version www.pdffactory.com

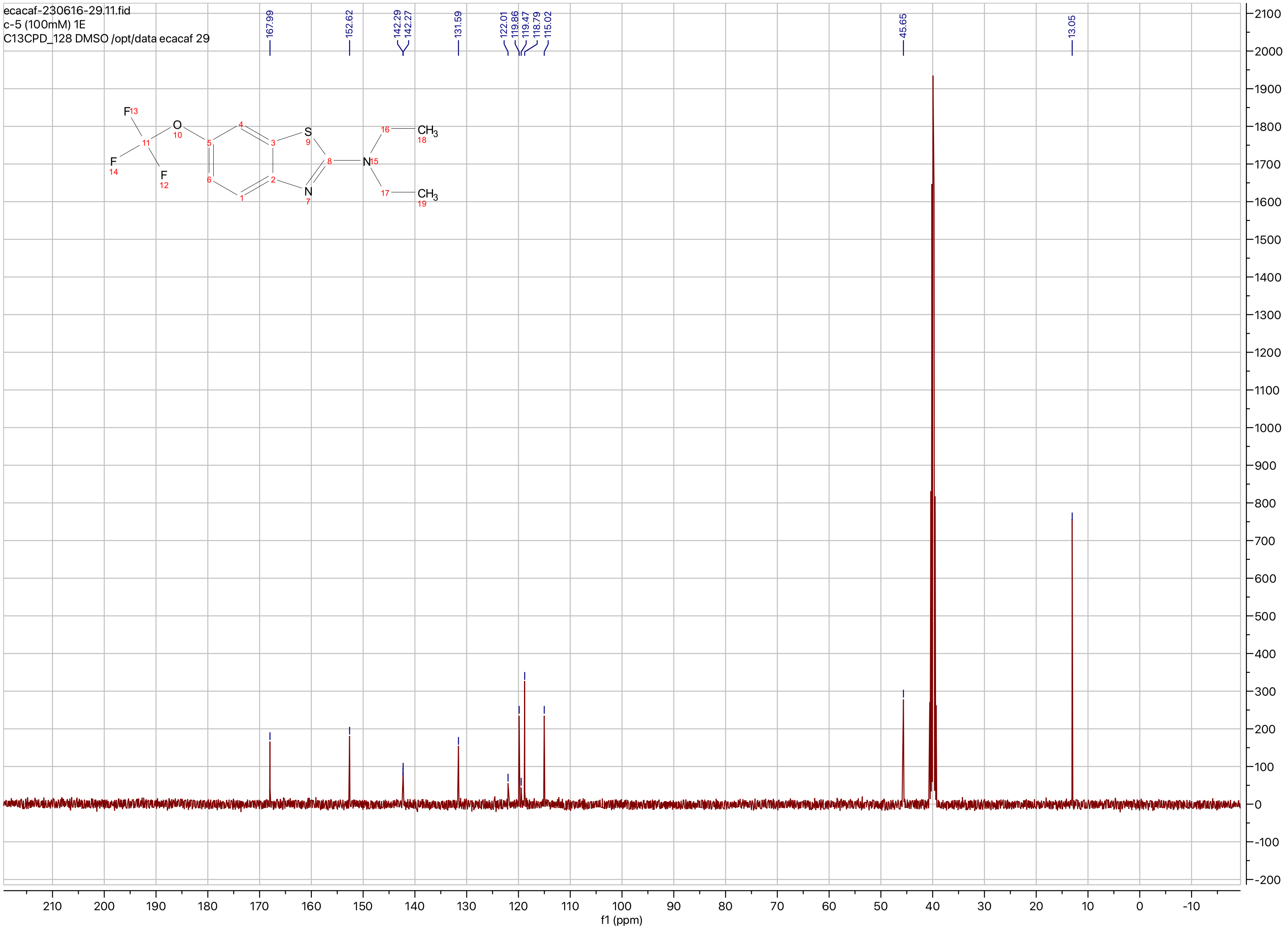
ecacaf-230616-29.11.fid
c-5 (100mM) 1E
C13CPD_128 DMSO /opt/data ecacaf 29
167.99
152.62
142.29
142.27
131.59
122.01
119.86
119.47
118.79
115.02
45.65
13.05
f1 (ppm)

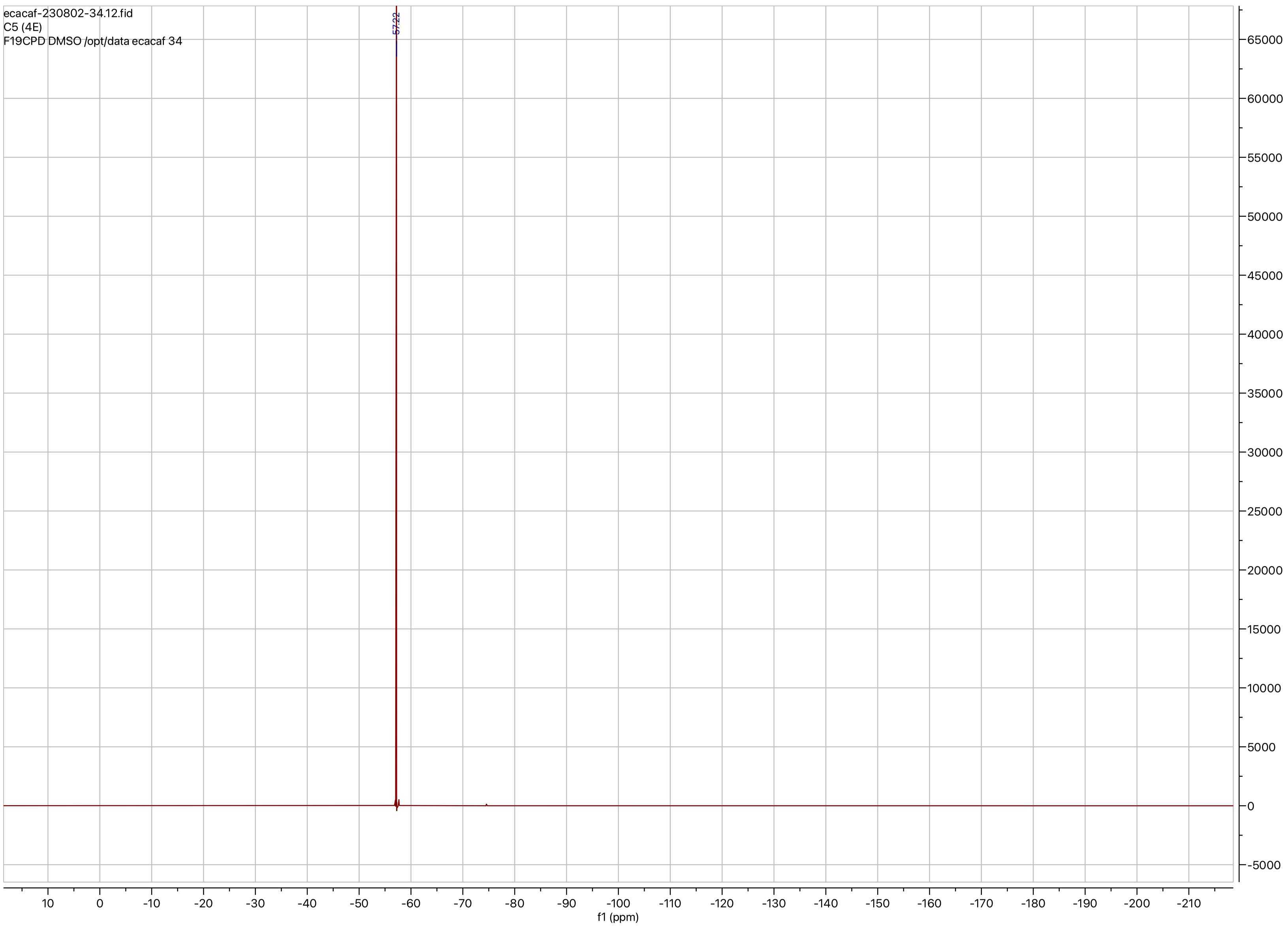
ecacaf-230802-34.12.fid
C5 (4E)
F19CPD DMSO /opt/data ecacaf 34
-57.22
f1 (ppm)

**Compound ID: C-6**

EW8457-335-P1A DMSO Bruker_B_400MHz

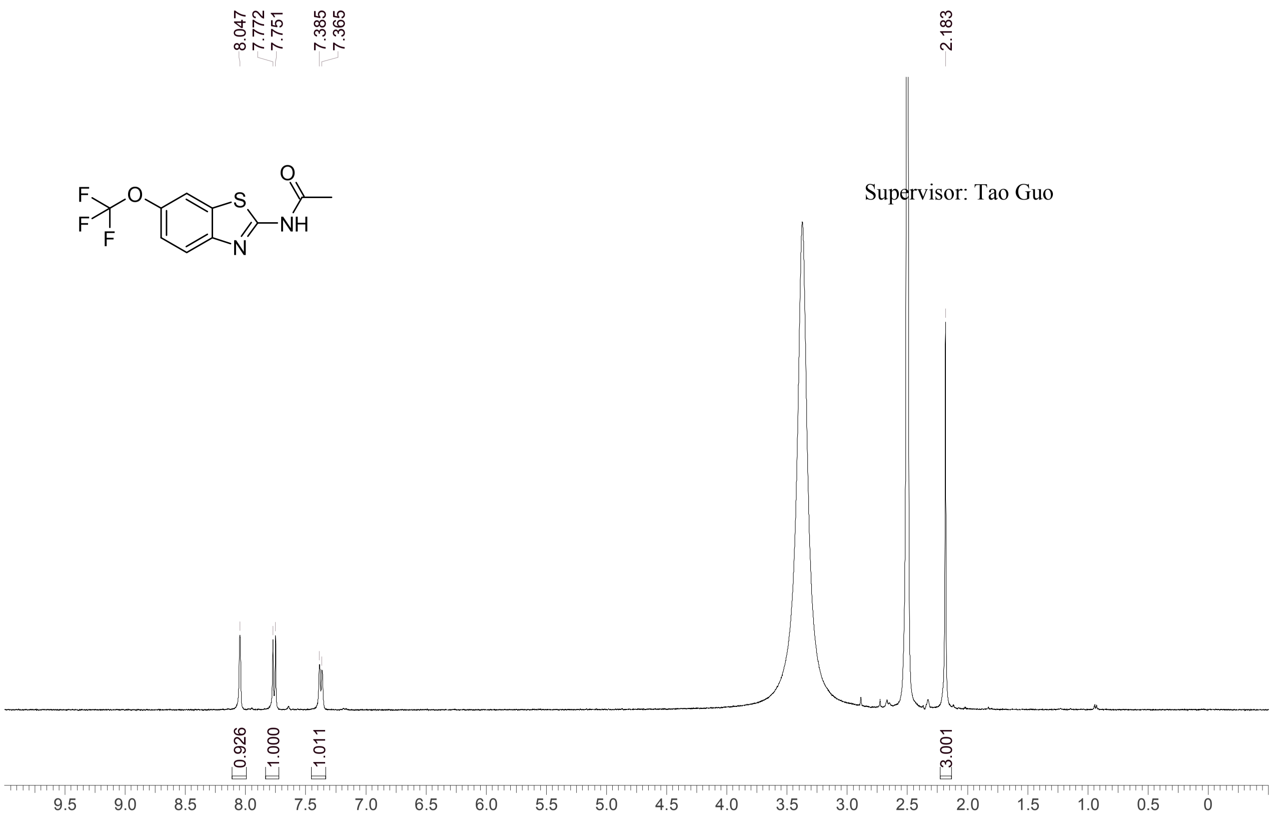


Supervisor: Tao Guo

| | |
|---|---|
| Acquisition Time (sec) | 3.0671 |
| Comment | EW8457-335-P1A DMSO Bruker_B_400MHz |
| Date | 08 Apr 2018 12:13:00 |
| Frequency (MHz) | 400.1300 |
| Nucleus | 1H |
| Number of Transients | 8 |
| Origin | spect |
| Original Points Count | 24576 |
| Owner | nmrsu |
| Points Count | 65536 |
| Pulse Sequence | zg30 |
| Receiver Gain | 190.49 |
| SW(cyclical) (Hz) | 8012.82 |
| Solvent | DMSO-d6 |
| Spectrum Offset (Hz) | 2397.7786 |
| Spectrum Type | standard |
| Sweep Width (Hz) | 8012.70 |
| Temperature (degree C) | 21.917 |

Operator: Date:

Confidential, for research only not for regulatory filing

Confidential, for research only not for regulatory filing

### LCMS REPORT

Compound ID : C-6
Sample ID : EW8457-335-P1B
Injection Vol : 1ul
Location : vial76
Acq Method : d:\method\5-95AB_R_220&254.lcm
Org DataFile : D:\DATA\1804\180408\EW8457-335-P1B.lcd
Injection Date : 2018-04-08 11:01:59
Instrument : LCMS-H 15-105

Chromatogram

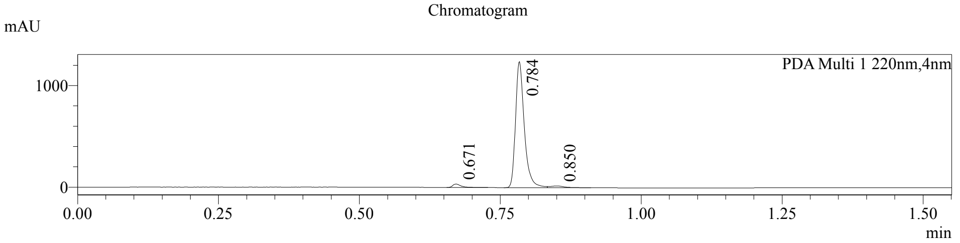


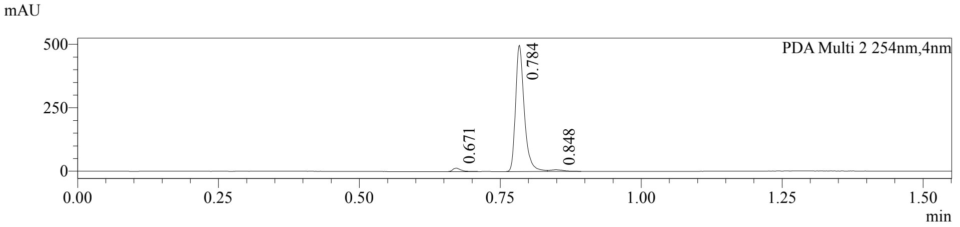


1 PDA Multi 1 / 220nm,4nm
2 PDA Multi 2 / 254nm,4nm

MS Chromatogram

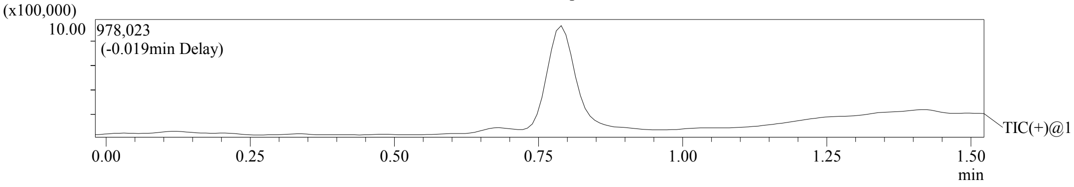


Integration Result

Peak Table

PDA Ch1 220nm

| Peak# | Ret. Time | Height | Height% | USP Width | Area | Area% |
|---|---|---|---|---|---|---|
| 1 | 0.671 | 34166 | 2.644 | 0.027 | 34915 | 2.473 |
| 2 | 0.784 | 1241044 | 96.036 | 0.029 | 1346696 | 95.367 |
| 3 | 0.850 | 17065 | 1.321 | 0.057 | 30515 | 2.161 |

Peak Table

PDA Ch2 254nm

| Peak# | Ret. Time | Height | Height% | USP Width | Area | Area% |
|---|---|---|---|---|---|---|
| 1 | 0.671 | 13690 | 2.643 | 0.027 | 13665 | 2.438 |
| 2 | 0.784 | 497634 | 96.090 | 0.029 | 535979 | 95.644 |
| 3 | 0.848 | 6558 | 1.266 | 0.054 | 10747 | 1.918 |

Confidential, for research only not for regulatory filing

Mass Spectrum

RetTime: 0.673 Datafile: D:\DATA\1804\180408\EW8457-335-P1B.lcd

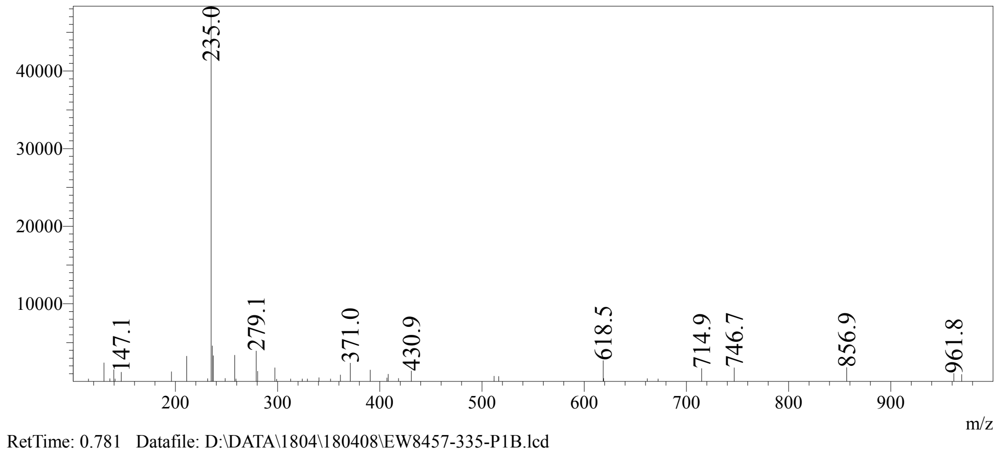


RetTime: 0.781 Datafile: D:\DATA\1804\180408\EW8457-335-P1B.lcd

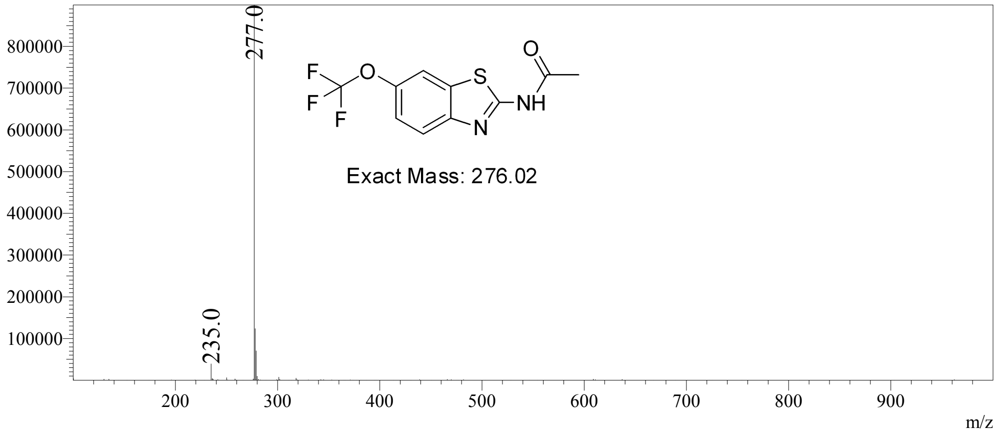


RetTime: 0.848 Datafile: D:\DATA\1804\180408\EW8457-335-P1B.lcd

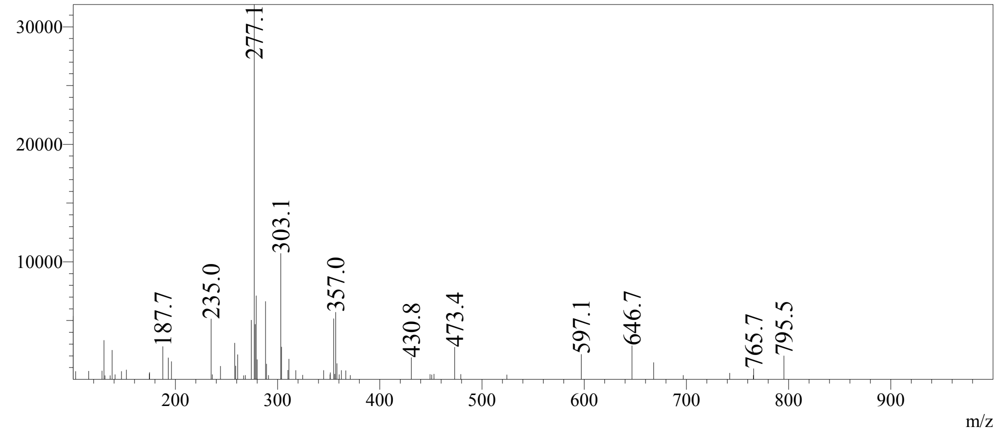

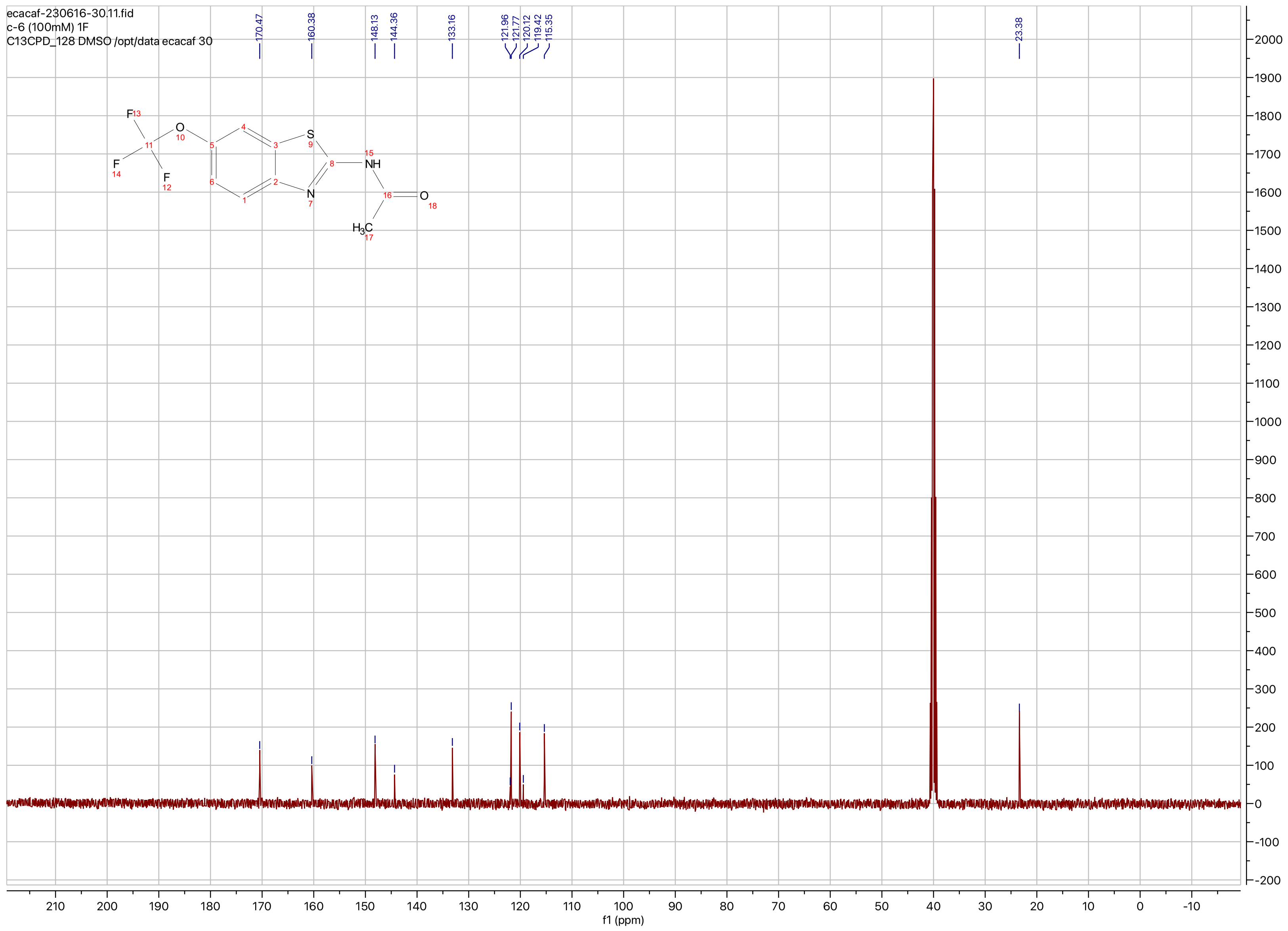
ecacaf-230616-30.11.fid
c-6 (100mM) 1F
C13CPD_128 DMSO /opt/data ecacaf 30
170.47
160.38
148.13
144.36
133.16
121.96
121.77
120.12
119.42
115.35
23.38
f1 (ppm)

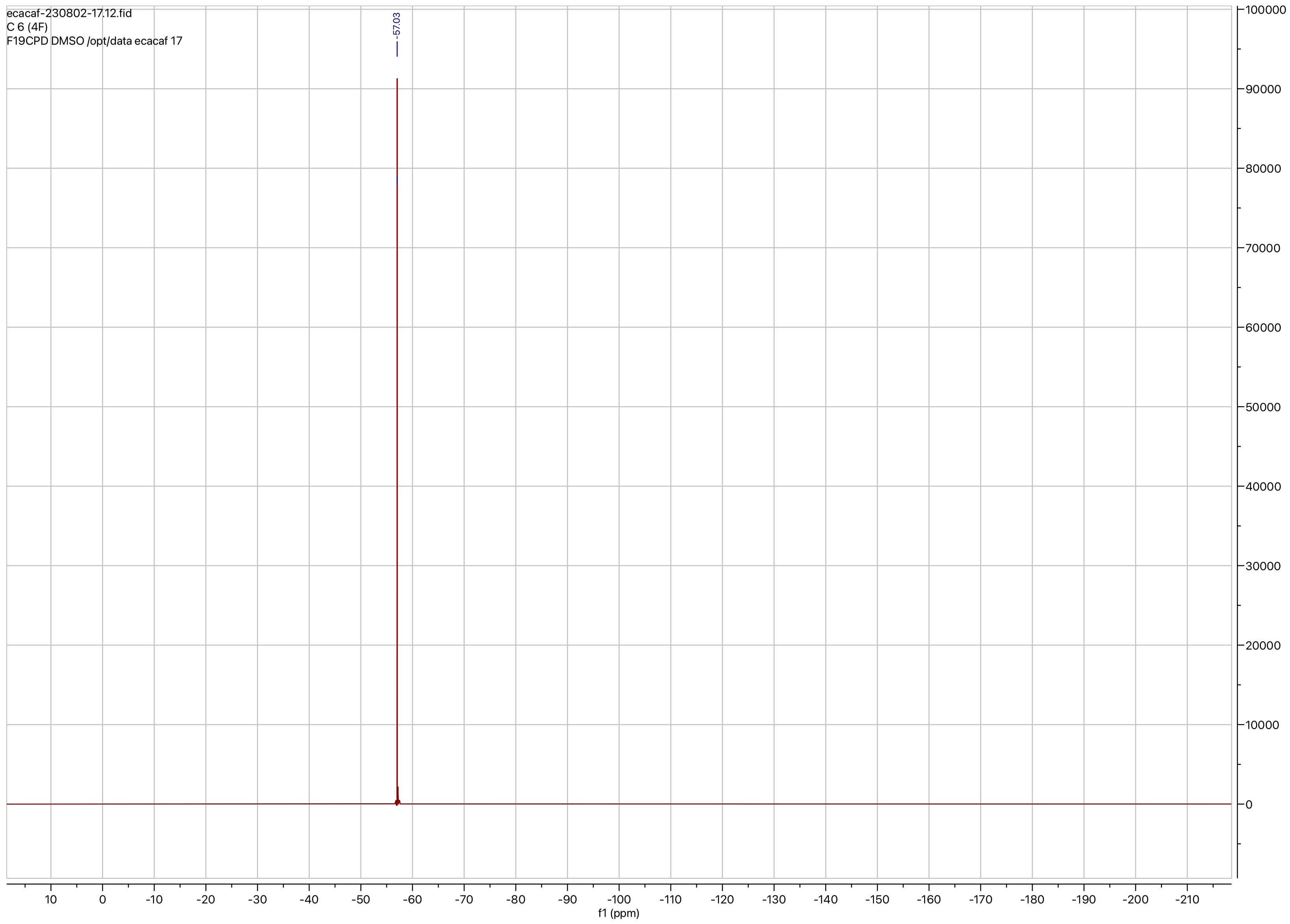

ecacaf-230802-17.12.fid
C 6 (4F)
F19CPD DMSO /opt/data ecacaf 17
-57.03
f1 (ppm)

**Compound ID: C-7**

EW8457-386-P1A DMSO Bruker_B_400MHz

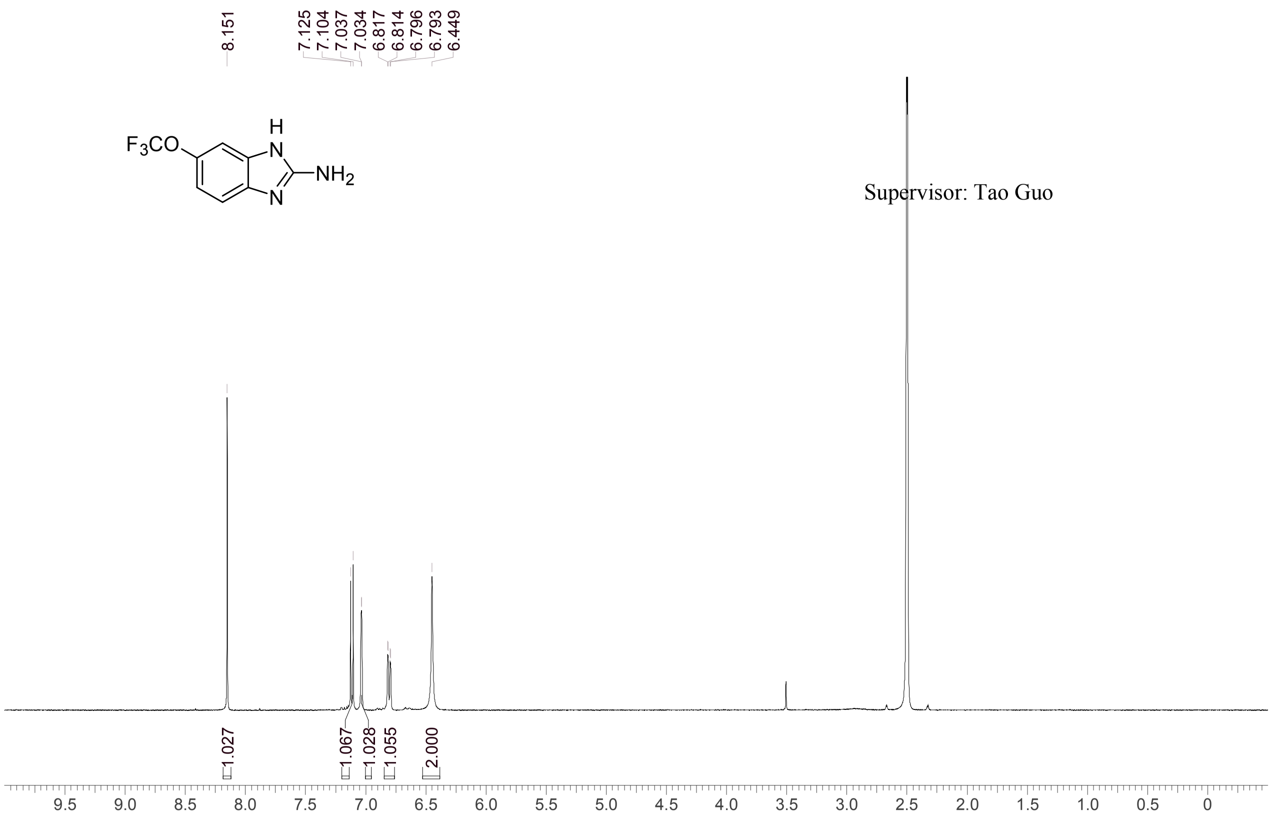


| | |
|---|---|
| Acquisition Time (sec) | 3.0671 |
| Comment | EW8457-386-P1A DMSO Bruker_B_400MHz |
| Date | 11 May 2018 08:27:55 |
| Frequency (MHz) | 400.1300 |
| Nucleus | 1H |
| Number of Transients | 8 |
| Origin | spect |
| Original Points Count | 24576 |
| Owner | nmrsu |
| Points Count | 65536 |
| Pulse Sequence | zg30 |
| Receiver Gain | 190.49 |
| SW(cyclical) (Hz) | 8012.82 |
| Solvent | DMSO-d6 |
| Spectrum Offset (Hz) | 2397.5386 |
| Spectrum Type | standard |
| Sweep Width (Hz) | 8012.70 |
| Temperature (degree C) | -273.000 |

Operator: Date:

Confidential, for research only not for regulatory filing

Confidential, for research only not for regulatory filing

### LCMS REPORT

Compound ID : C-7
Sample ID : EW8457-386-P1B
Injection Vol : 3ul
Location : vial27
Acq Method : d:\method\5-95AB_R_220&254.lcm
Org DataFile : D:\DATA\1805\180510\EW8457-386-P1B.lcd
Injection Date : 2018-05-10 16:56:35
Instrument : LCMS-I

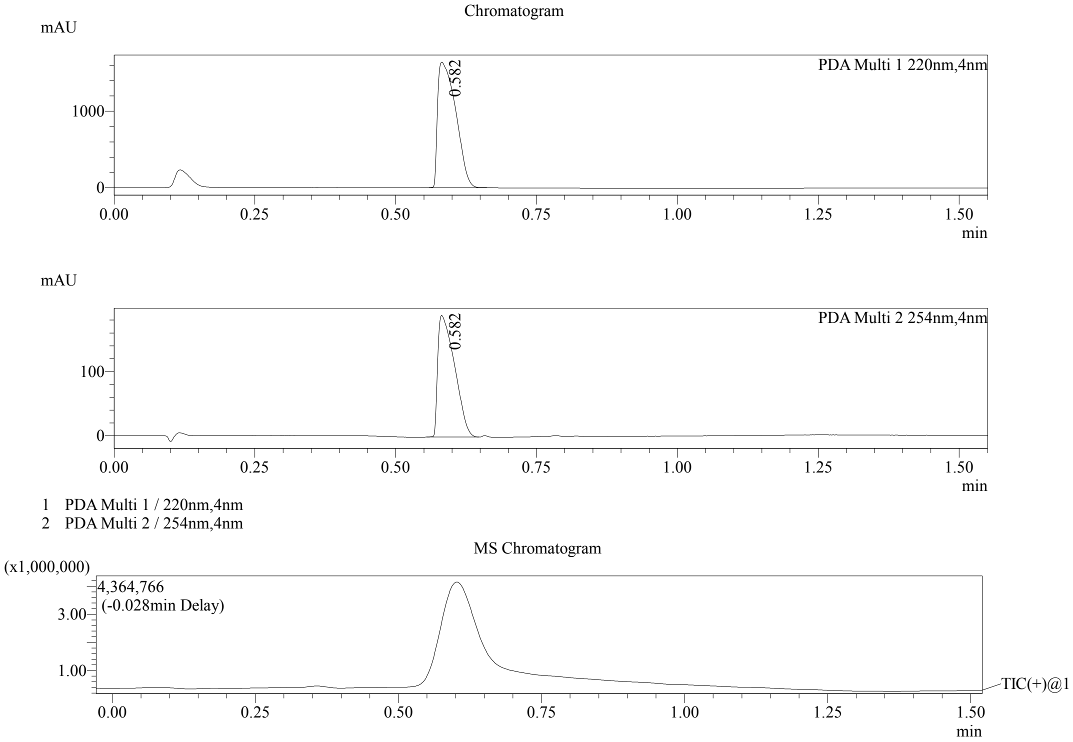


Integration Result

Peak Table

PDA Ch1 220nm

| Peak# | Ret. Time | Height | Height% | USP Width | Area | Area% |
|---|---|---|---|---|---|---|
| 1 | 0.582 | 1645607 | 100.000 | 0.060 | 3686092 | 100.000 |

Peak Table

PDA Ch2 254nm

| Peak# | Ret. Time | Height | Height% | USP Width | Area | Area% |
|---|---|---|---|---|---|---|
| 1 | 0.582 | 189394 | 100.000 | 0.059 | 392977 | 100.000 |

PDF created with pdfFactory Pro trial version www.pdffactory.com

Confidential, for research only not for regulatory filing

Mass Spectrum

RetTime: 0.582 DateFile: D:\DATA\1805\180510\EW8457-386-P1B.lcd

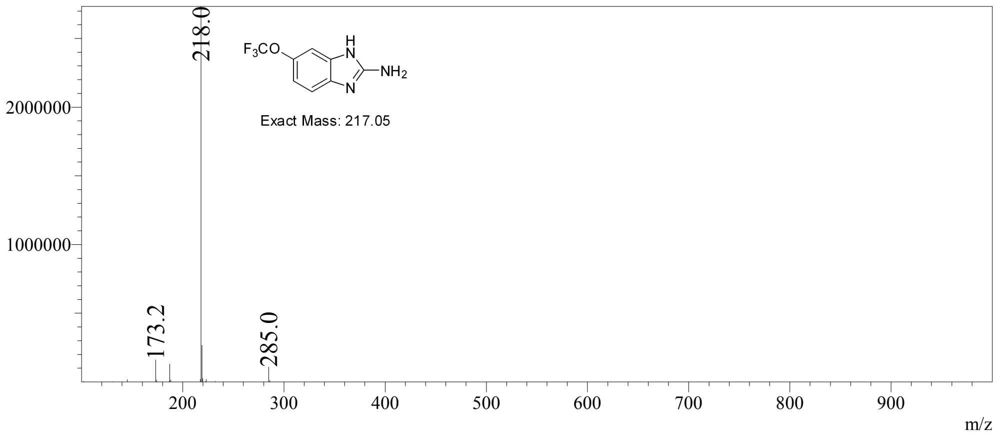


PDF created with pdfFactory Pro trial version www.pdffactory.com

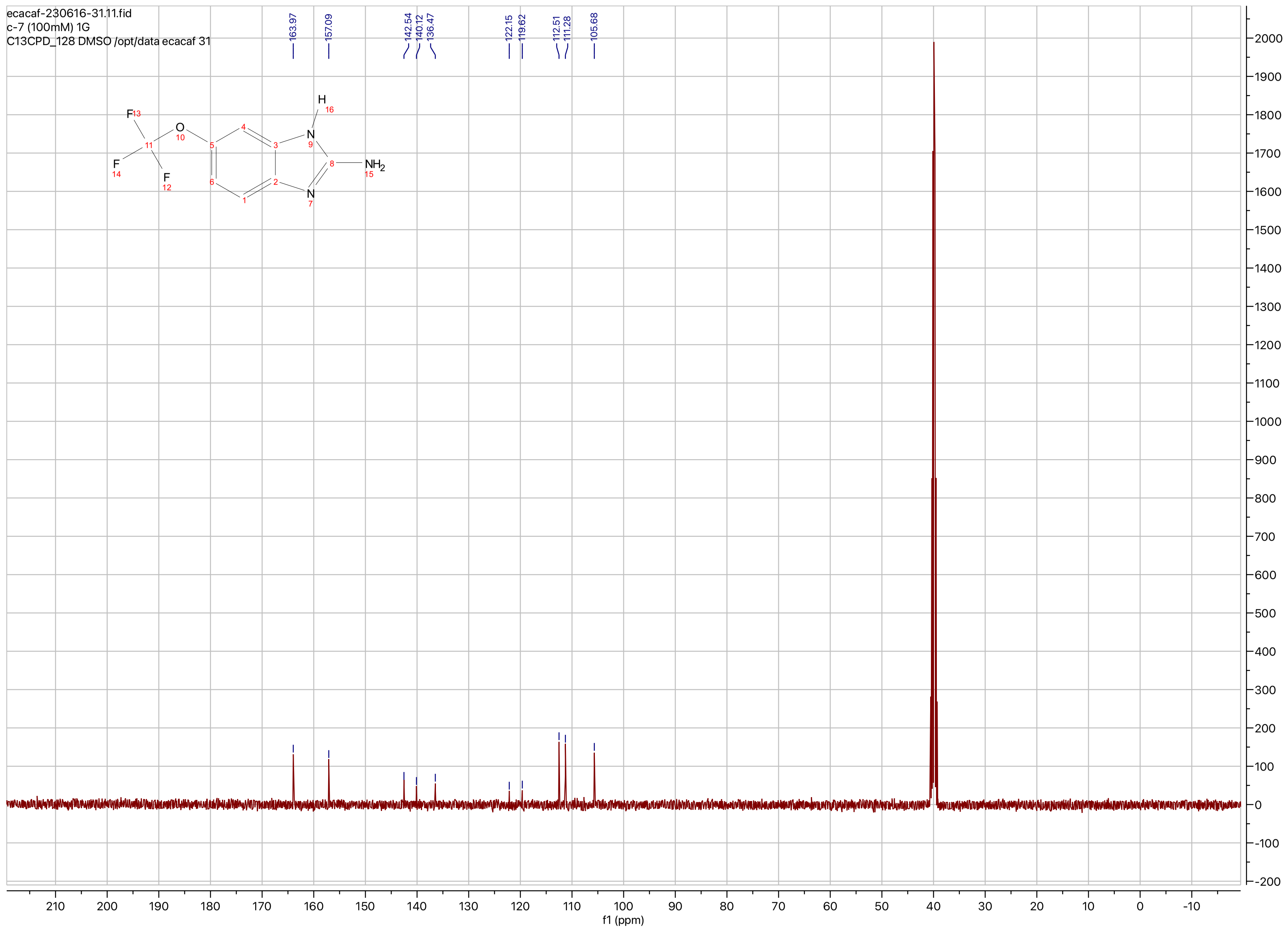
ecacaf-230616-31.11.fid
c-7 (100mM) 1G
C13CPD_128 DMSO /opt/data ecacaf 31
163.97
157.09
142.54
140.12
136.47
122.15
119.62
112.51
111.28
105.68
F
O
N
H
NH2
N
F
F
f1 (ppm)

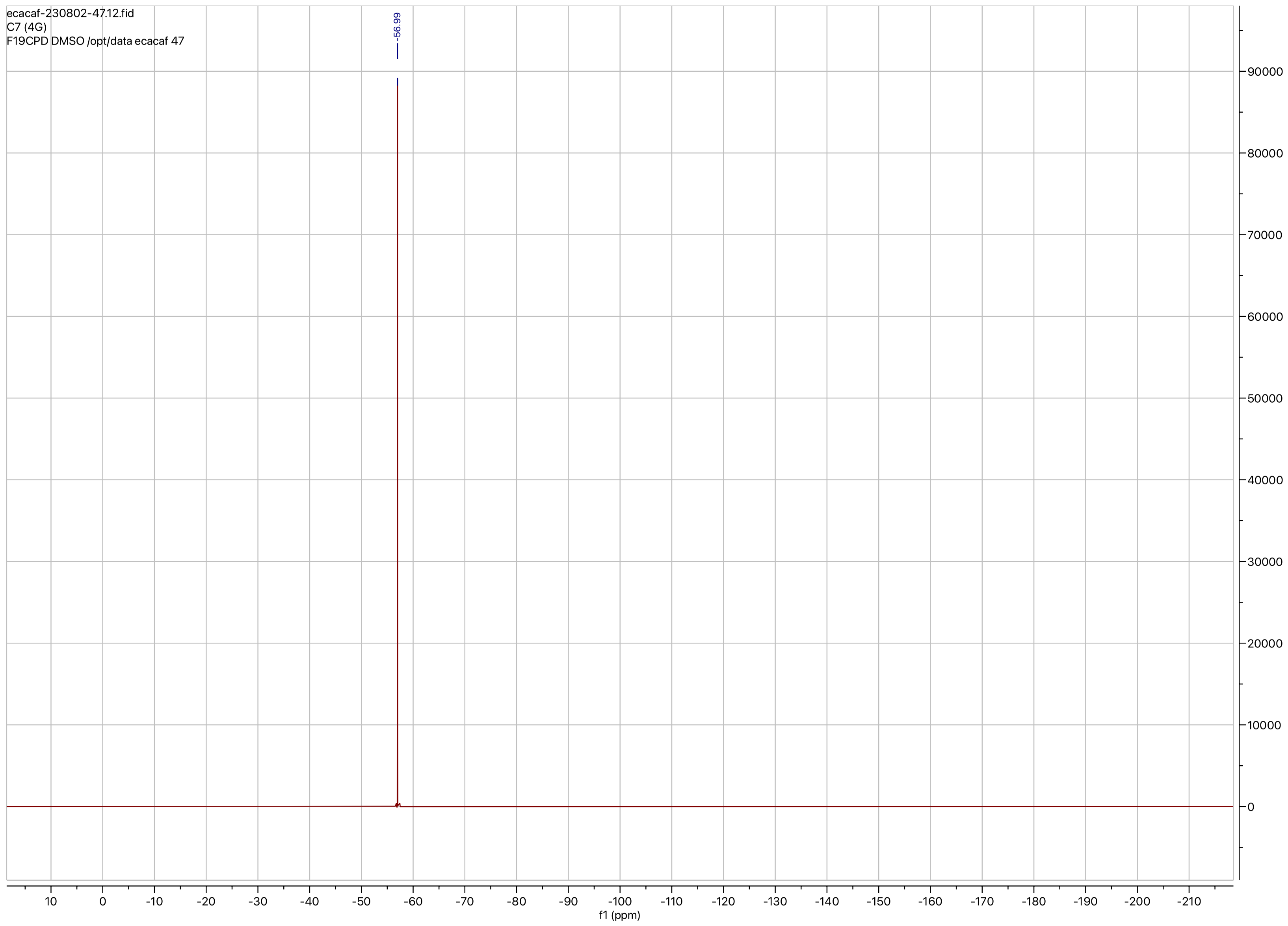

ecacaf-230802-47.12.fid
C7 (4G)
F19CPD DMSO /opt/data ecacaf 47
-56.99
f1 (ppm)

**Compound ID: C-8**

EW8457-412-P1A DMSO Bruker_G_400MHz

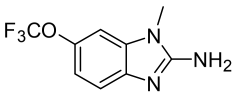


Supervisor: Tao Guo

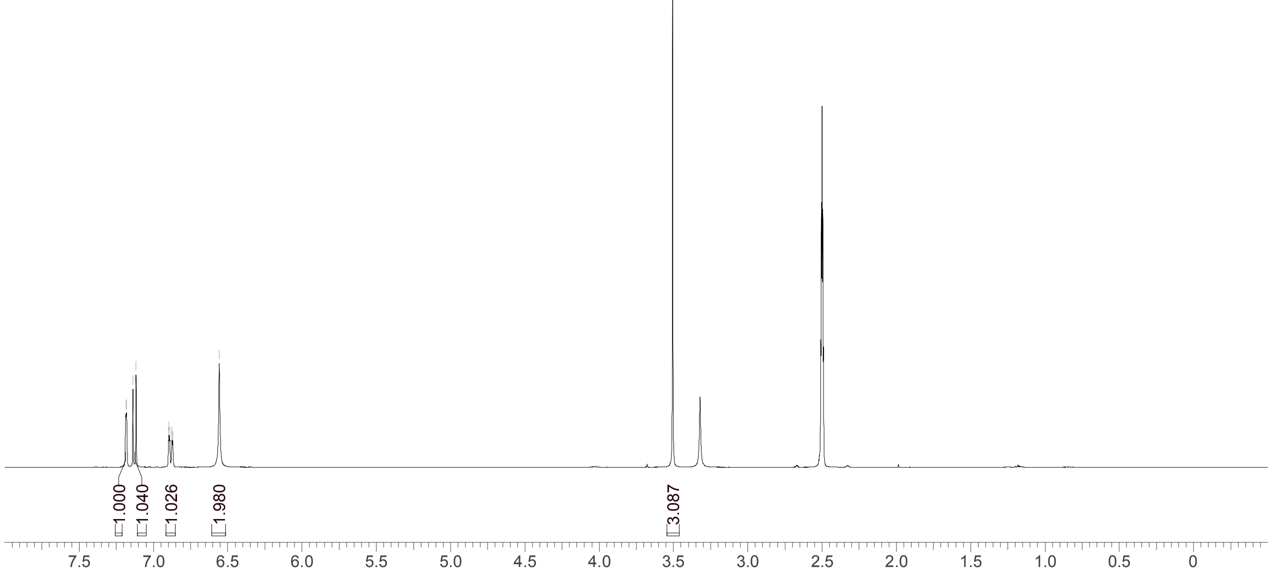


| | |
|---|---|
| Acquisition Time (sec) | 3.9977 |
| Comment | EW8457-412-P1A DMSO Bruker_G_400MHz |
| Date | 22 May 2018 03:50:50 |
| Frequency (MHz) | 400.1900 |
| Nucleus | 1H |
| Number of Transients | 8 |
| Origin | Avance |
| Original Points Count | 32768 |
| Owner | nmrsu |
| Points Count | 65536 |
| Pulse Sequence | zg30 |
| Receiver Gain | 101.00 |
| SW(cyclical) (Hz) | 8196.72 |
| Solvent | DMSO-d6 |
| Spectrum Offset (Hz) | 2468.0127 |
| Spectrum Type | standard |
| Sweep Width (Hz) | 8196.60 |
| Temperature (degree C) | 27.018 |

Operator:

Date:

Confidential, for research only not for regulatory filing

Confidential, for research only not for regulatory filing

### LCMS REPORT

Compound ID : C-8
Sample ID : EW8457-412-P1B
Injection Vol : 2ul
Location : vial102
Acq Method : d:\method\5-95CD_R_220&254_POS_50.lcm
Org DataFile : D:\DATA\1805\180522\EW8457-412-P1B.lcd
Injection Date : 2018-05-22 11:25:36
Instrument :LCMS-O 15-105

Chromatogram

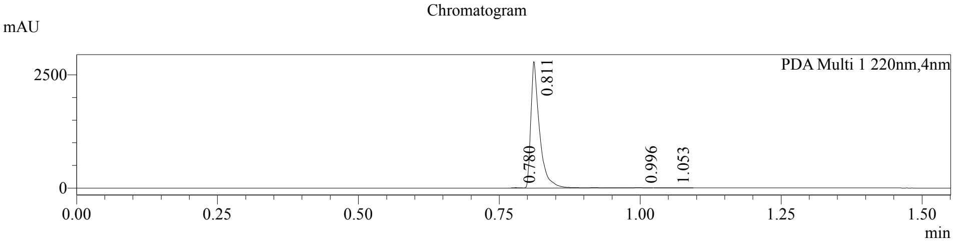


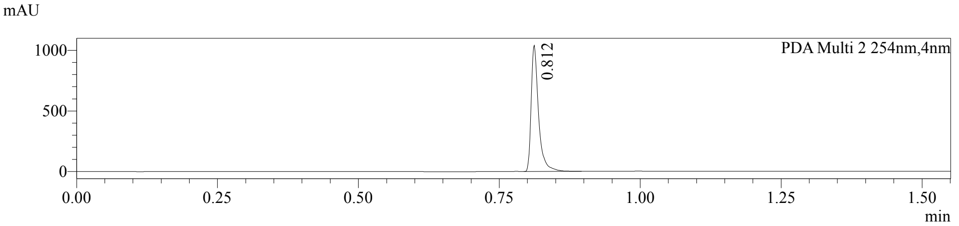


1 PDA Multi 1 / 220nm,4nm
2 PDA Multi 2 / 254nm,4nm

MS Chromatogram

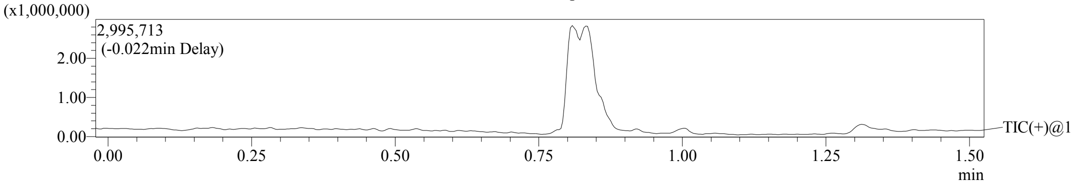


Integration Result

Peak Table

PDA Ch1 220nm

| Peak# | Ret. Time | Height | Height% | USP Width | Area | Area% |
|---|---|---|---|---|---|---|
| 1 | 0.780 | 7689 | 0.273 | 0.026 | 6675 | 0.219 |
| 2 | 0.811 | 2791700 | 99.284 | 0.029 | 3027320 | 99.313 |
| 3 | 0.996 | 9638 | 0.343 | 0.025 | 8340 | 0.274 |
| 4 | 1.053 | 2819 | 0.100 | 0.056 | 5912 | 0.194 |

Peak Table

PDA Ch2 254nm

| Peak# | Ret. Time | Height | Height% | USP Width | Area | Area% |
|---|---|---|---|---|---|---|
| 1 | 0.812 | 1041595 | 100.000 | 0.025 | 959660 | 100.000 |

Confidential, for research only not for regulatory filing

Mass Spectrum

RetTime:0.778 Datafile: D:\DATA\1805\180522\EW8457-412-P1B.lcd

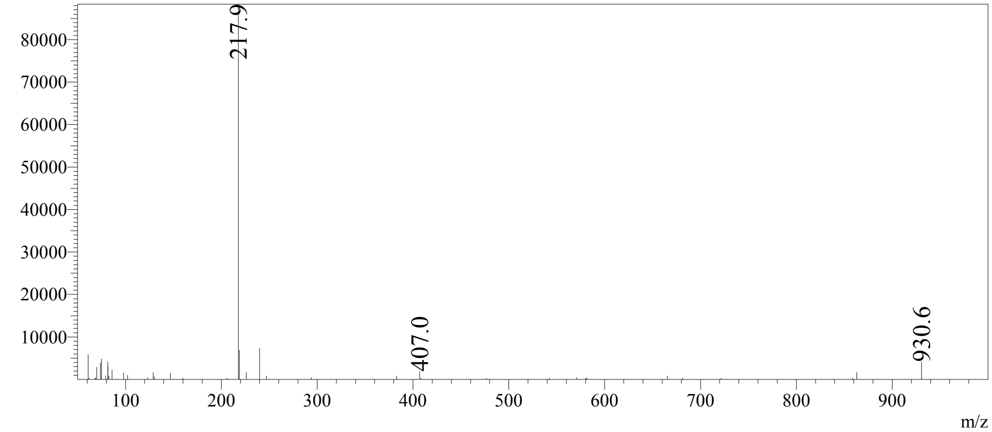


RetTime:0.812 Datafile: D:\DATA\1805\180522\EW8457-412-P1B.lcd

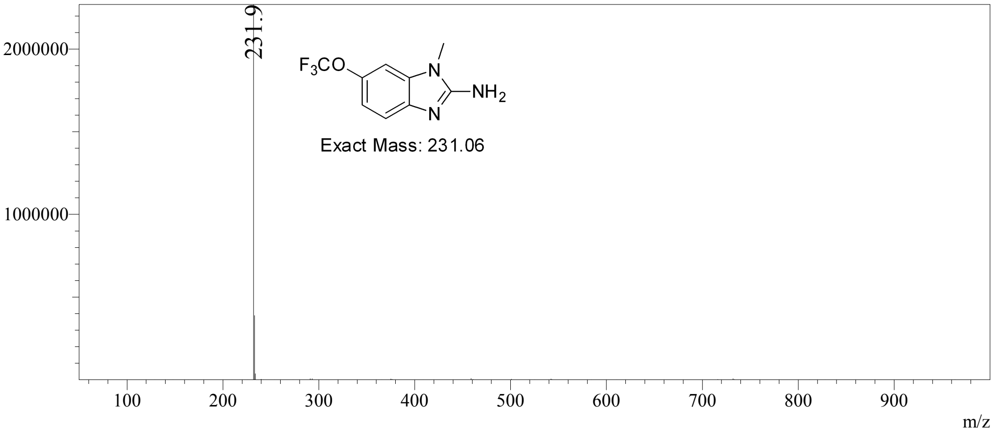


RetTime:0.995 Datafile: D:\DATA\1805\180522\EW8457-412-P1B.lcd

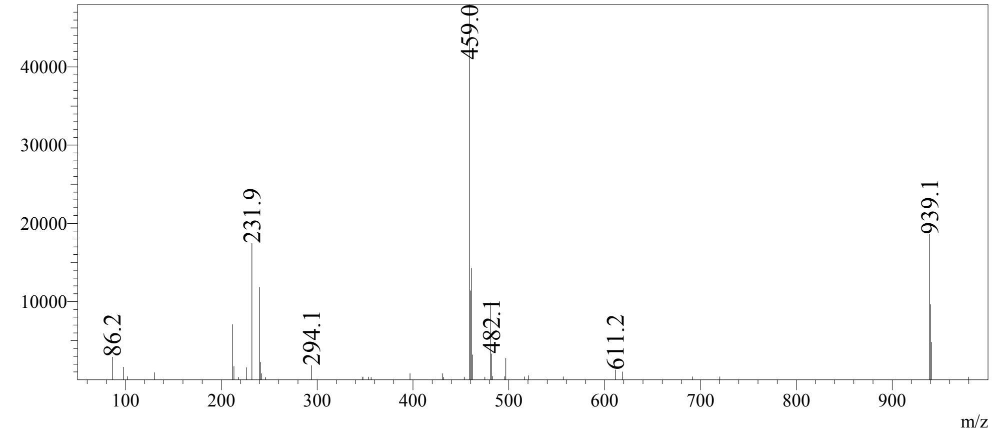

Confidential, for research only not for regulatory filing

RetTime:1.052 Datafile: D:\DATA\1805\180522\EW8457-412-P1B.lcd

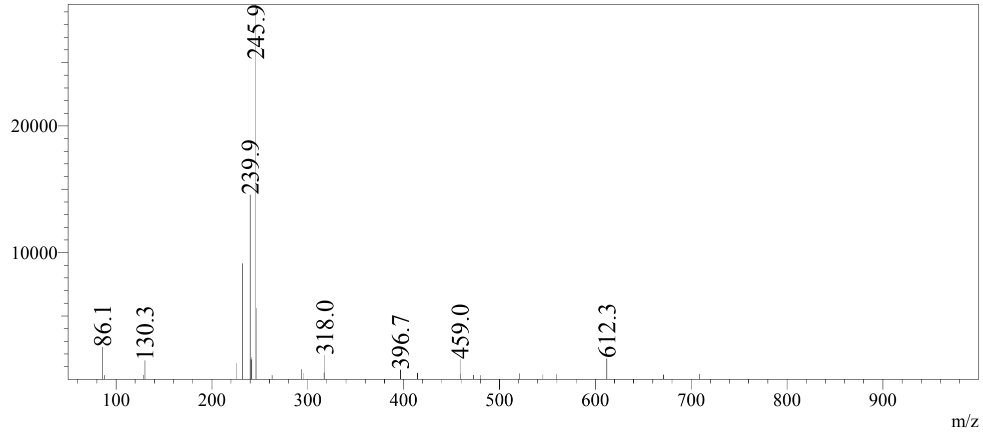

ecacaf-230616-32.11.fid
c-8 (100mM) 1l
C13CPD_128 DMSO /opt/data ecacaf 32

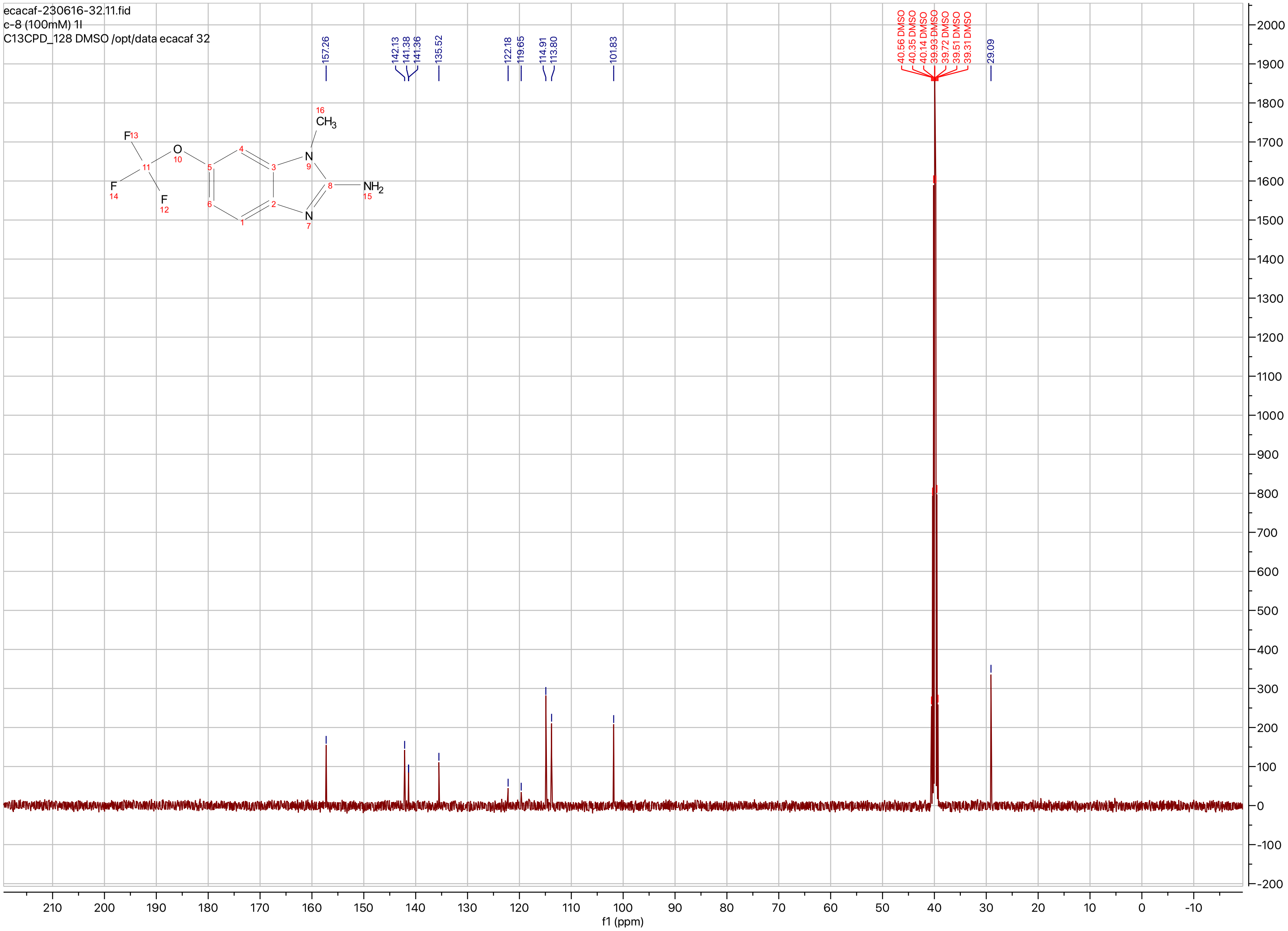

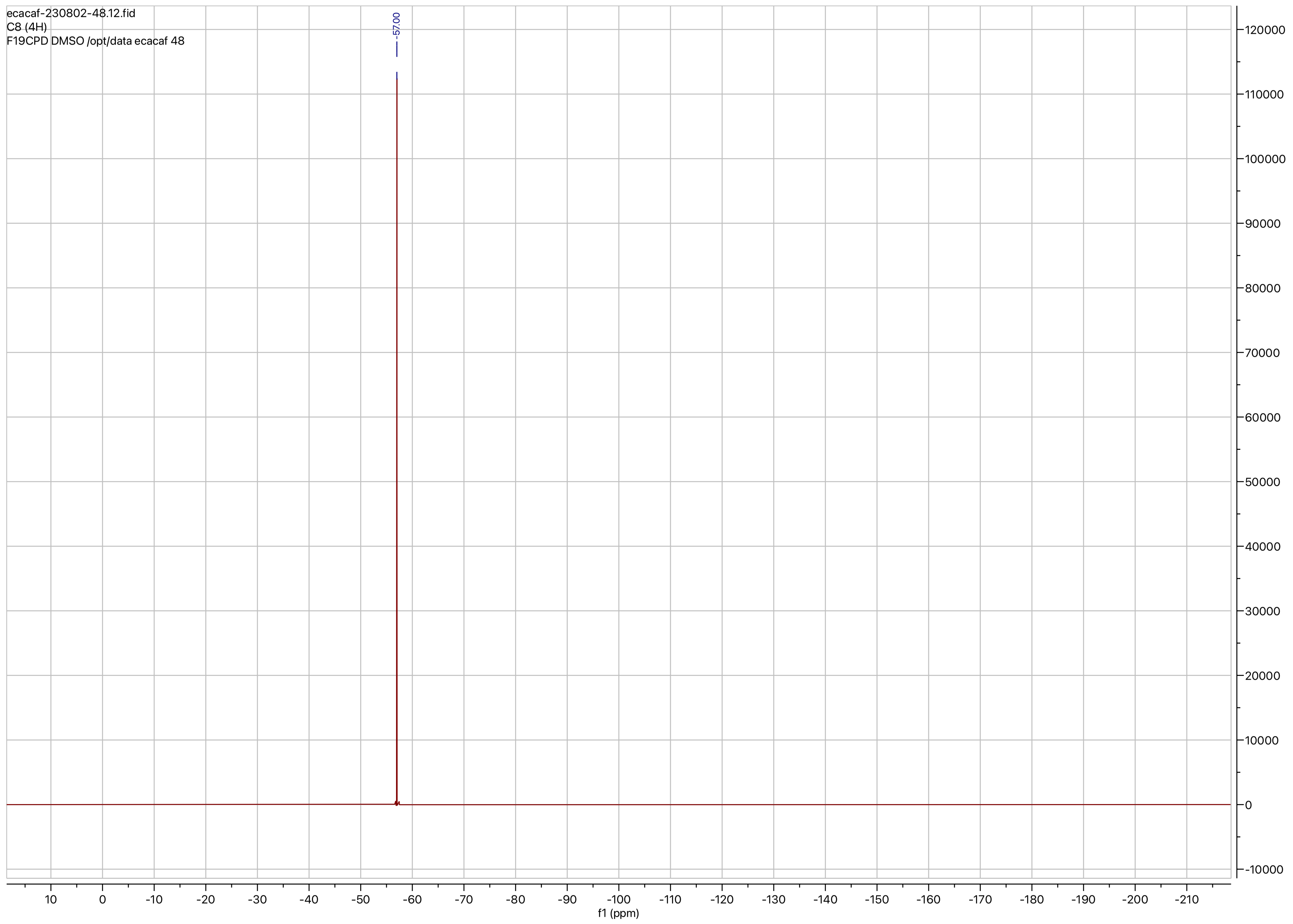
ecacaf-230802-48.12.fid
C8 (4H)
F19CPD DMSO /opt/data ecacaf 48
-57.00
f1 (ppm)

**Compound ID: C-9**

EW8457-417-P1D DMSO Bruker_E_400MHz

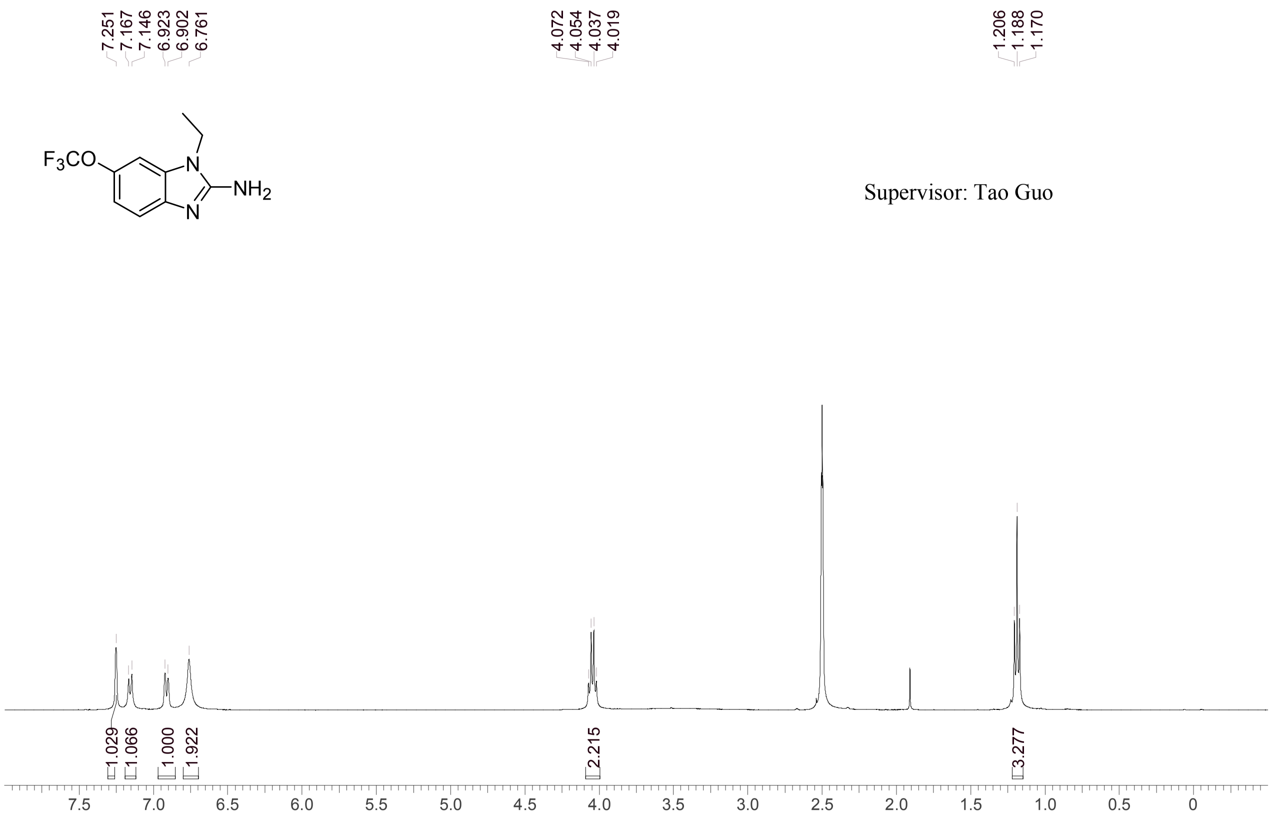


Supervisor: Tao Guo

| | |
|---|---|
| Acquisition Time (sec) | 3.0671 |
| Comment | EW8457-417-P1D DMSO Bruker_E_400MHz |
| Date | 27 May 2018 09:33:02 |
| Frequency (MHz) | 400.1500 |
| Nucleus | 1H |
| Number of Transients | 8 |
| Origin | spect |
| Original Points Count | 24576 |
| Owner | nmrsu |
| Points Count | 65536 |
| Pulse Sequence | zg30 |
| Receiver Gain | 160.91 |
| SW(cyclical) (Hz) | 8012.82 |
| Solvent | DMSO-d6 |
| Spectrum Offset (Hz) | 2397.3147 |
| Spectrum Type | standard |
| Sweep Width (Hz) | 8012.70 |
| Temperature (degree C) | 27.143 |

Operator:

Date:

Confidential, for research only not for regulatory filing

Confidential, for research only not for regulatory filing

### LCMS REPORT

Compound ID : C-9
Sample ID : EW8457-417-P1B
Injection Vol : 3ul
Location : vial20
Acq Method : d:\method\5-95CD_R_220&254_POS_50.lcm
Org DataFile : D:\DATA\1805\180524\EW8457-417-P1B.lcd
Injection Date : 2018-05-24 15:04:23
Instrument :LCMS-O 15-105

Chromatogram

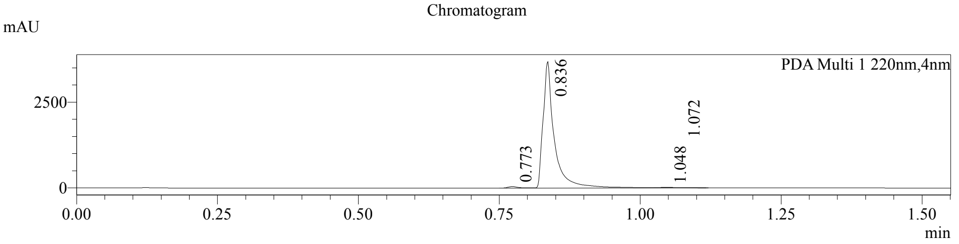


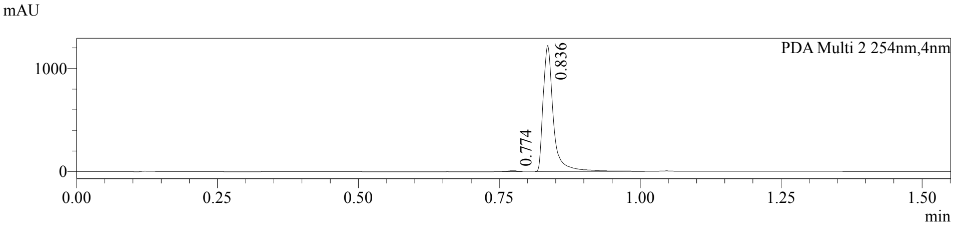


1 PDA Multi 1 / 220nm,4nm
2 PDA Multi 2 / 254nm,4nm

MS Chromatogram

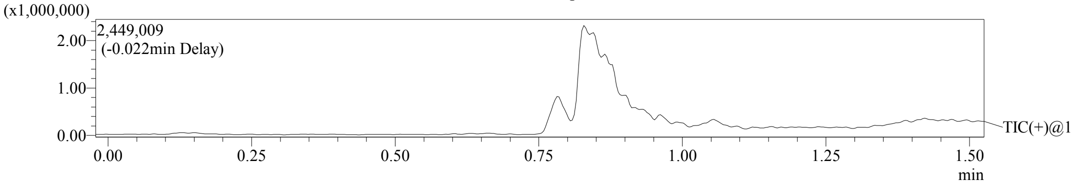


Integration Result

Peak Table

PDA Ch1 220nm

| Peak# | Ret. Time | Height | Height% | USP Width | Area | Area% |
|---|---|---|---|---|---|---|
| 1 | 0.773 | 42531 | 1.137 | 0.031 | 51698 | 1.000 |
| 2 | 0.836 | 3680342 | 98.405 | 0.033 | 5095157 | 98.550 |
| 3 | 1.048 | 11518 | 0.308 | 0.040 | 14782 | 0.286 |
| 4 | 1.072 | 5587 | 0.149 | 0.161 | 8465 | 0.164 |

Peak Table

PDA Ch2 254nm

| Peak# | Ret. Time | Height | Height% | USP Width | Area | Area% |
|---|---|---|---|---|---|---|
| 1 | 0.774 | 9004 | 0.730 | 0.031 | 9898 | 0.638 |
| 2 | 0.836 | 1225325 | 99.270 | 0.032 | 1541120 | 99.362 |

Confidential, for research only not for regulatory filing

Mass Spectrum

RetTime:0.775 Datafile: D:\DATA\1805\180524\EW8457-417-P1B.lcd

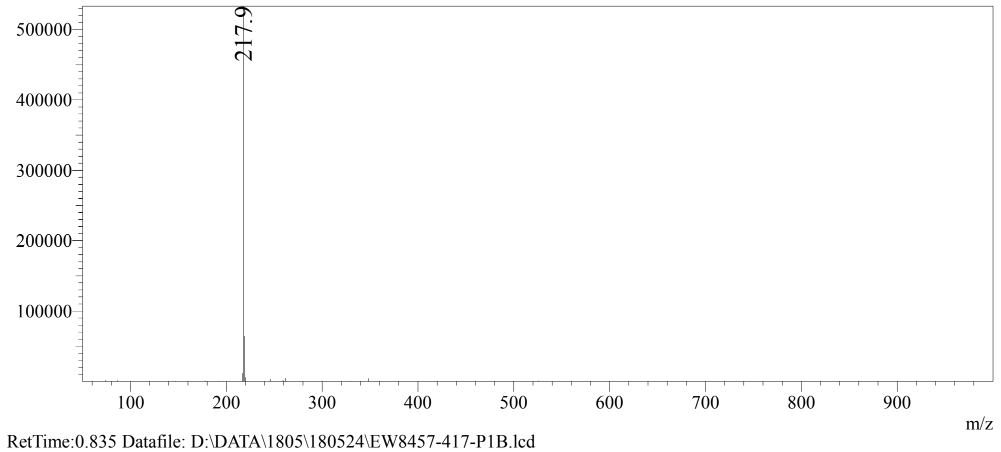


RetTime:0.835 Datafile: D:\DATA\1805\180524\EW8457-417-P1B.lcd

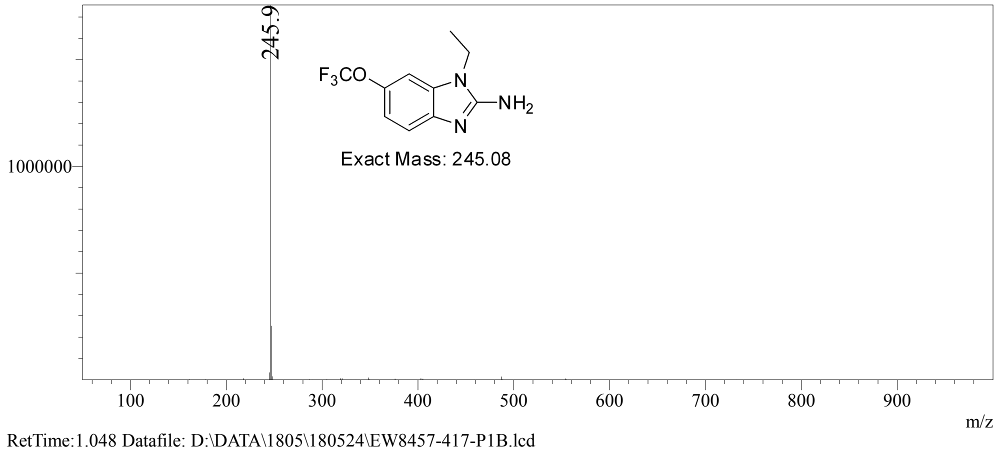


RetTime:1.048 Datafile: D:\DATA\1805\180524\EW8457-417-P1B.lcd

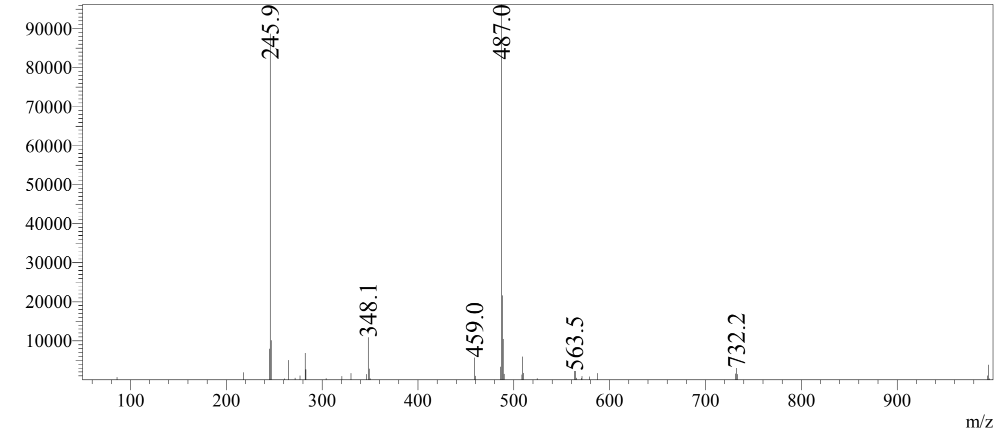

Confidential, for research only not for regulatory filing

RetTime:1.072 Datafile: D:\DATA\1805\180524\EW8457-417-P1B.lcd

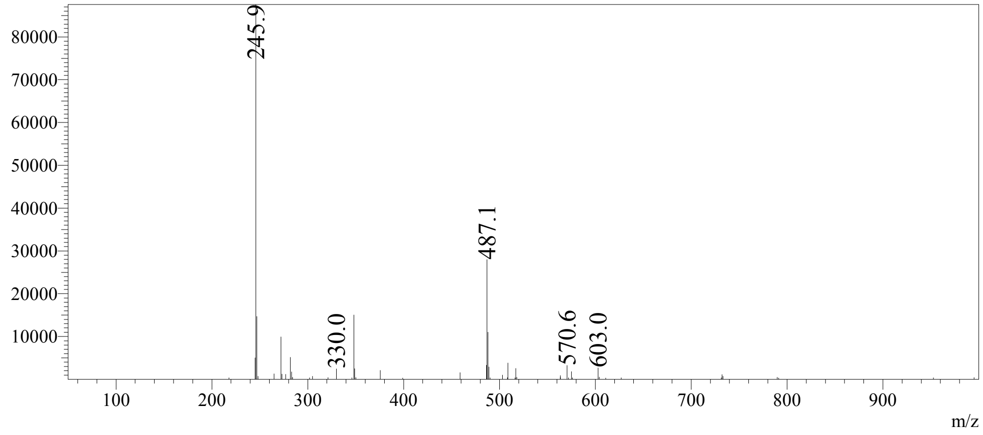

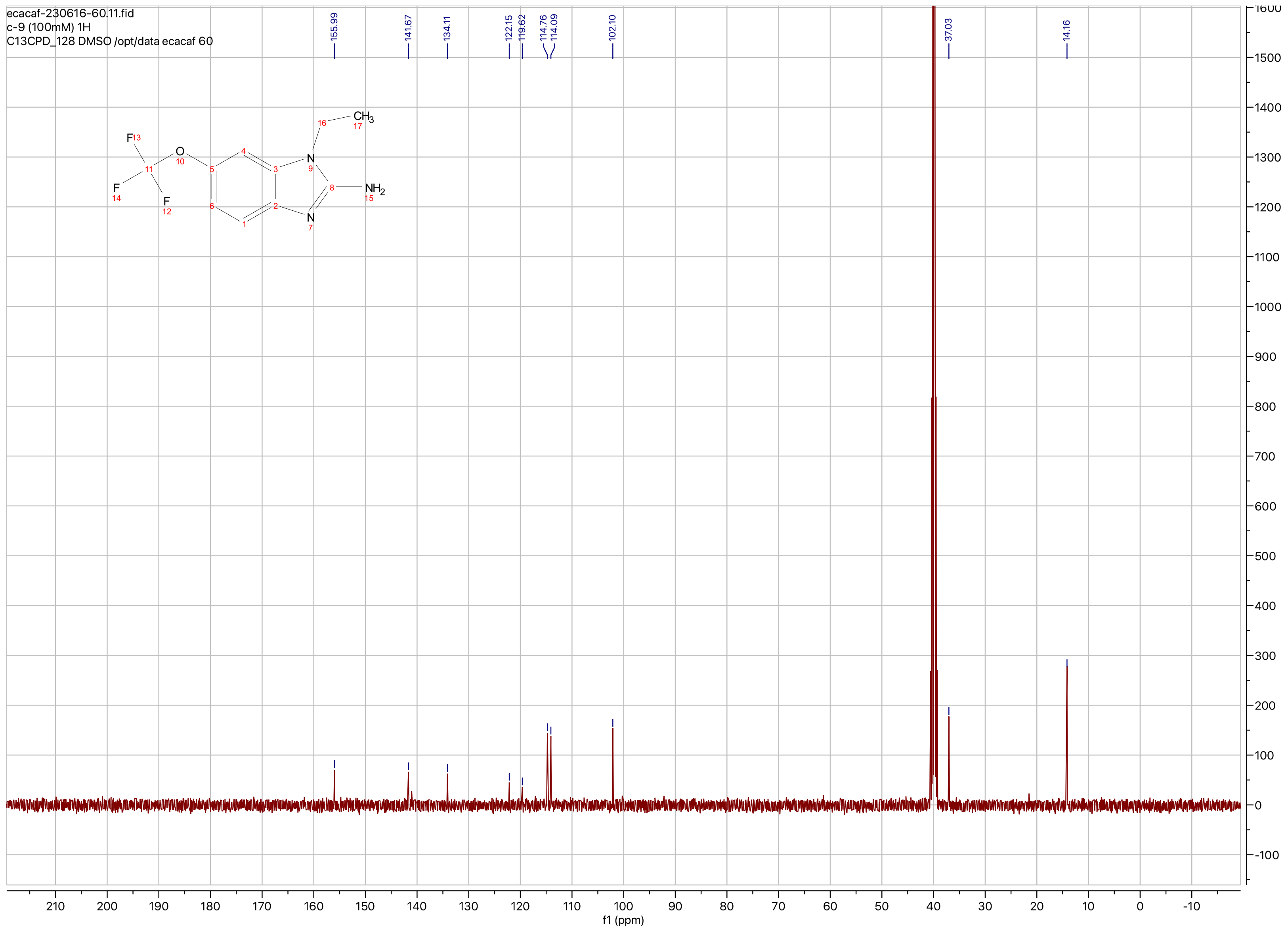
ecacaf-230616-60.11.fid
c-9 (100mM) 1H
C13CPD_128 DMSO /opt/data ecacaf 60
155.99
141.67
134.11
122.15
119.62
114.76
114.09
102.10
37.03
14.16
f1 (ppm)

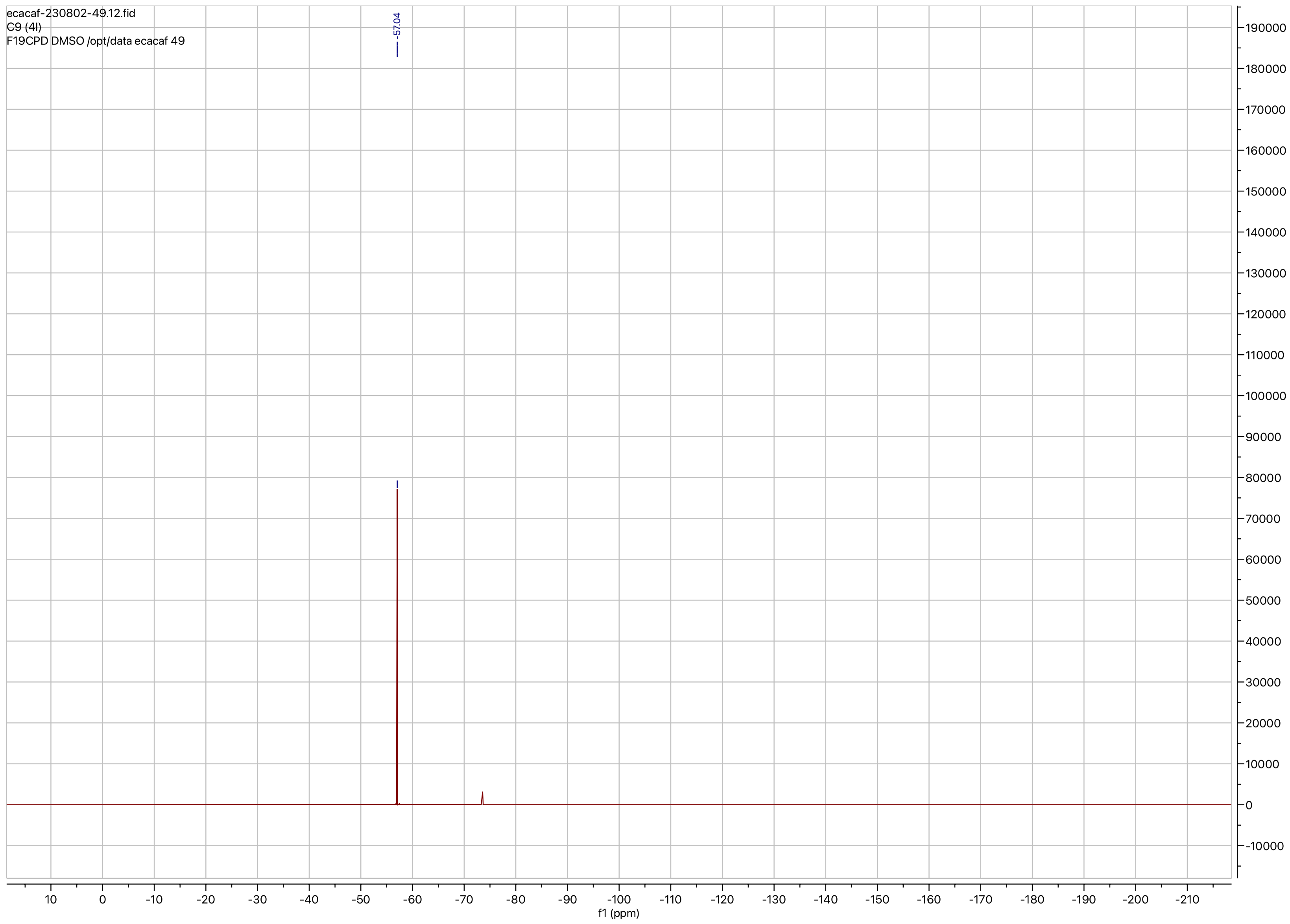
ecacaf-230802-49.12.fid
C9 (4l)
F19CPD DMSO /opt/data ecacaf 49
-57.04
f1 (ppm)

**Compound ID: C-10**

EW8457-411-P1C DMSO Bruker_G_400MHz

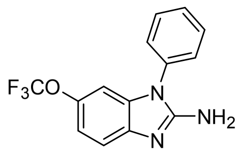

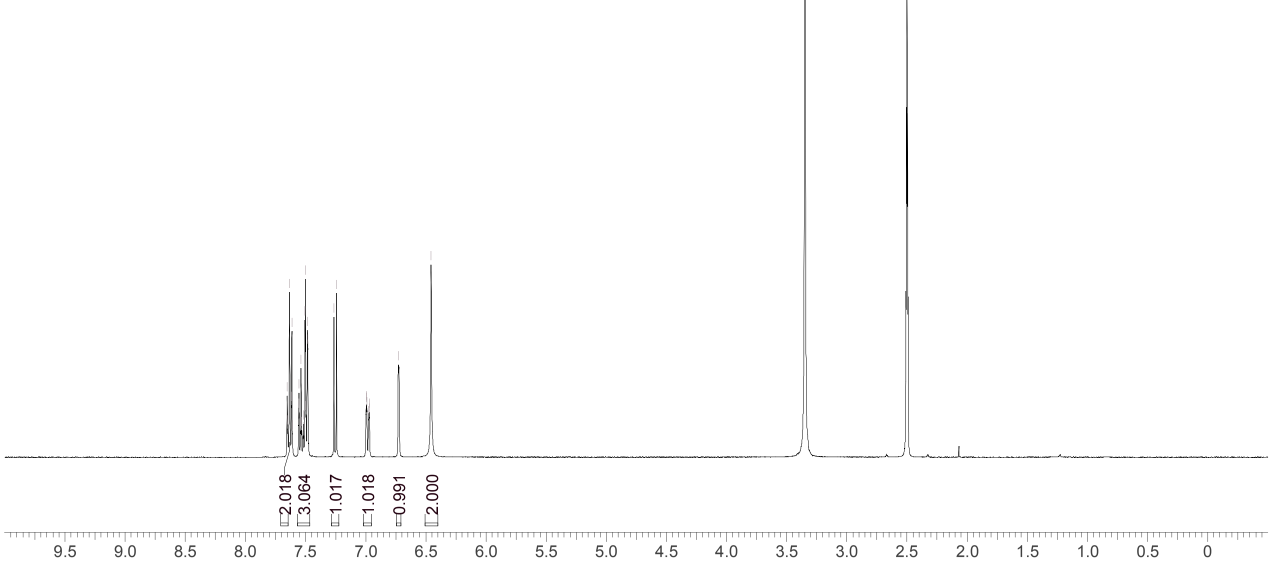


| | |
|---|---|
| Acquisition Time (sec) | 3.9977 |
| Comment | EW8457-411-P1C DMSO Bruker_G_400MHz |
| Date | 23 May 2018 09:28:07 |
| Frequency (MHz) | 400.1900 |
| Nucleus | 1H |
| Number of Transients | 8 |
| Origin | Avance |
| Original Points Count | 32768 |
| Owner | nmrsu |
| Points Count | 65536 |
| Pulse Sequence | zg30 |
| Receiver Gain | 101.00 |
| SW(cyclical) (Hz) | 8196.72 |
| Solvent | DMSO-d6 |
| Spectrum Offset (Hz) | 2468.0127 |
| Spectrum Type | standard |
| Sweep Width (Hz) | 8196.60 |
| Temperature (degree C) | 27.009 |

Operator: Date:

Confidential, for research only not for regulatory filing

Confidential, for research only not for regulatory filing

### LCMS REPORT

Compound ID : C-10
Sample ID : EW8457-411-P1B
Injection Vol : 1ul
Location : vial101
Acq Method : d:\method\5-95CD_R_220&254_POS_50.lcm
Org DataFile : D:\DATA\1805\180522\EW8457-411-P1B.lcd
Injection Date : 2018-05-22 11:23:31
Instrument :LCMS-O 15-105

Chromatogram

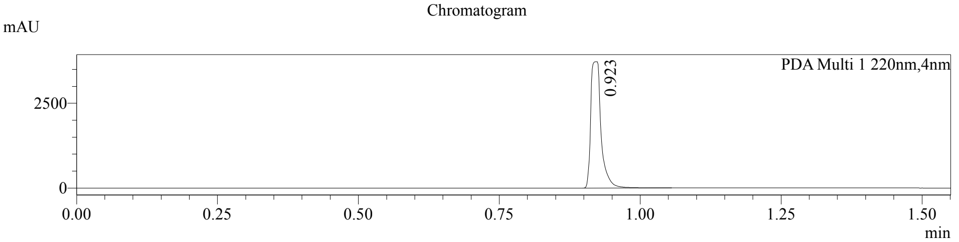


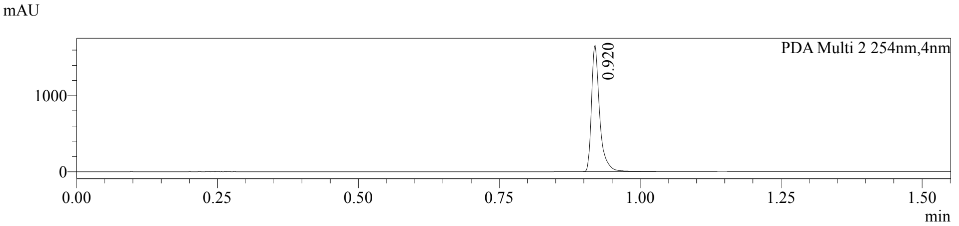


1 PDA Multi 1 / 220nm,4nm
2 PDA Multi 2 / 254nm,4nm

MS Chromatogram

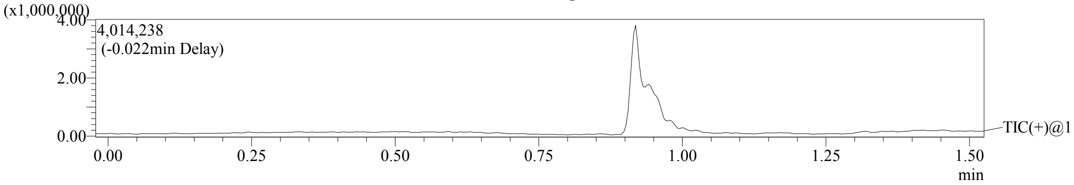


Integration Result

Peak Table

PDA Ch1 220nm

| Peak# | Ret. Time | Height | Height% | USP Width | Area | Area% |
|---|---|---|---|---|---|---|
| 1 | 0.923 | 3723548 | 100.000 | 0.029 | 4788279 | 100.000 |

Peak Table

PDA Ch2 254nm

| Peak# | Ret. Time | Height | Height% | USP Width | Area | Area% |
|---|---|---|---|---|---|---|
| 1 | 0.920 | 1662467 | 100.000 | 0.027 | 1681087 | 100.000 |

Confidential, for research only not for regulatory filing

Mass Spectrum
RetTime:0.922 Datafile: D:\DATA\1805\180522\EW8457-411-P1B.lcd

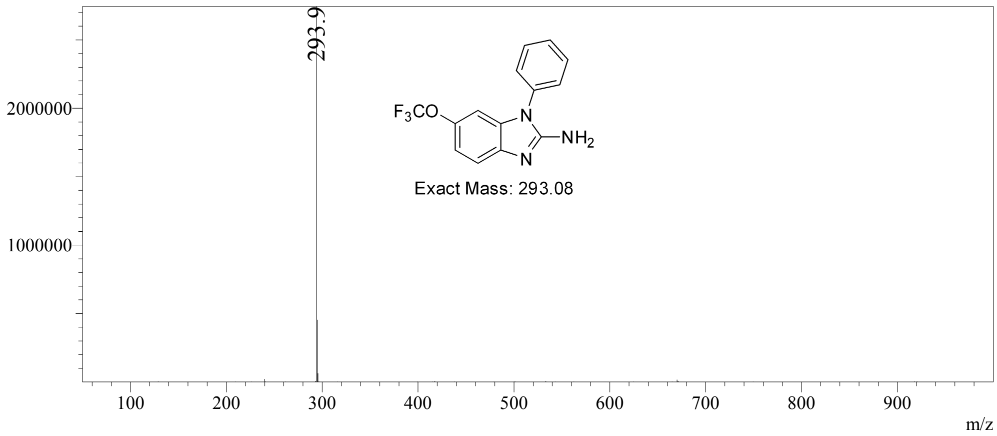

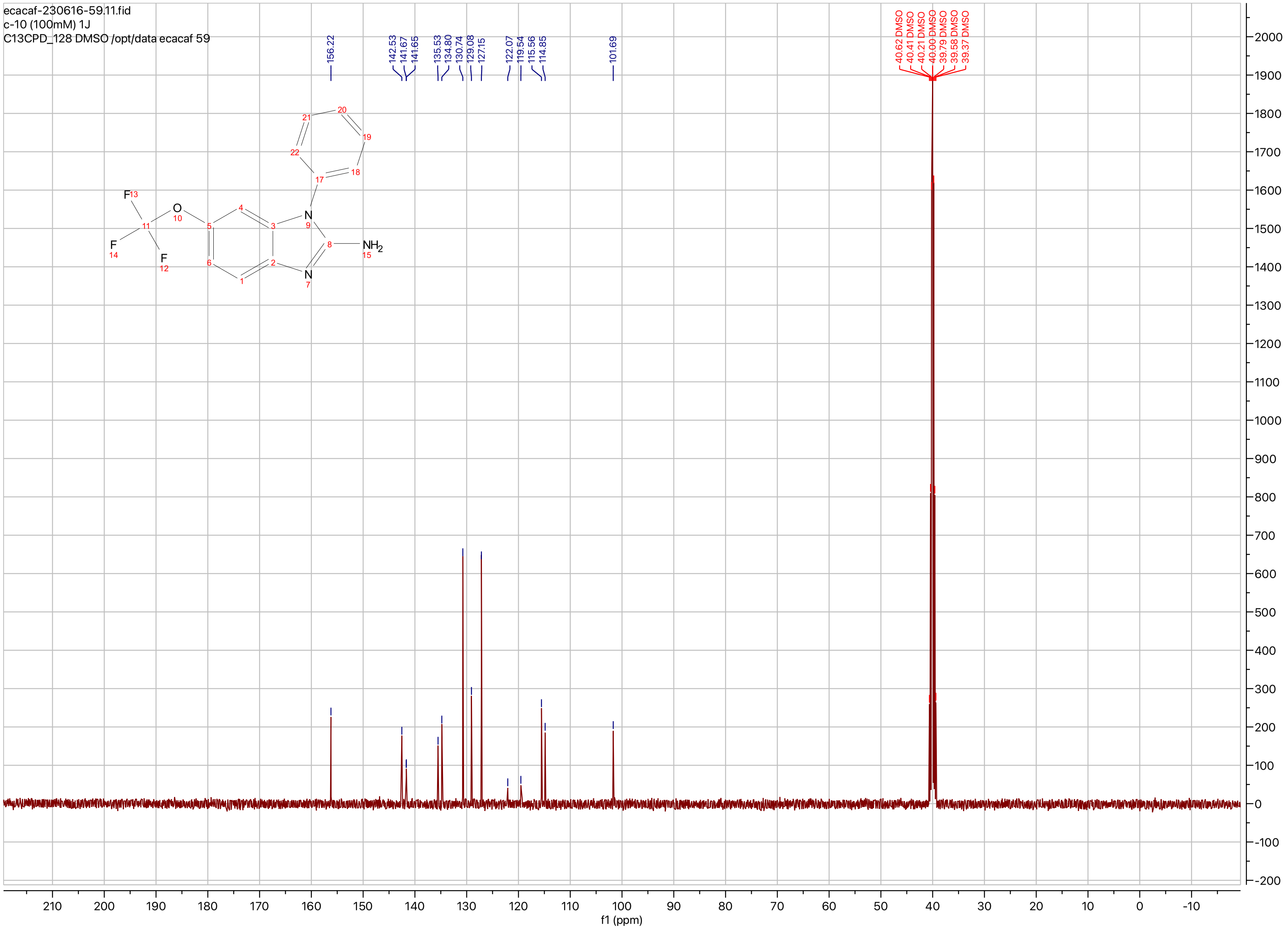

ecacaf-230616-59.11.fid
c-10 (100mM) 1J
C13CPD_128 DMSO /opt/data ecacaf 59
156.22
142.53
141.67
141.65
135.53
134.80
130.74
129.08
127.15
122.07
119.54
115.56
114.85
101.69
40.62 DMSO
40.41 DMSO
40.21 DMSO
40.00 DMSO
39.79 DMSO
39.58 DMSO
39.37 DMSO
F
O
N
NH2
f1 (ppm)

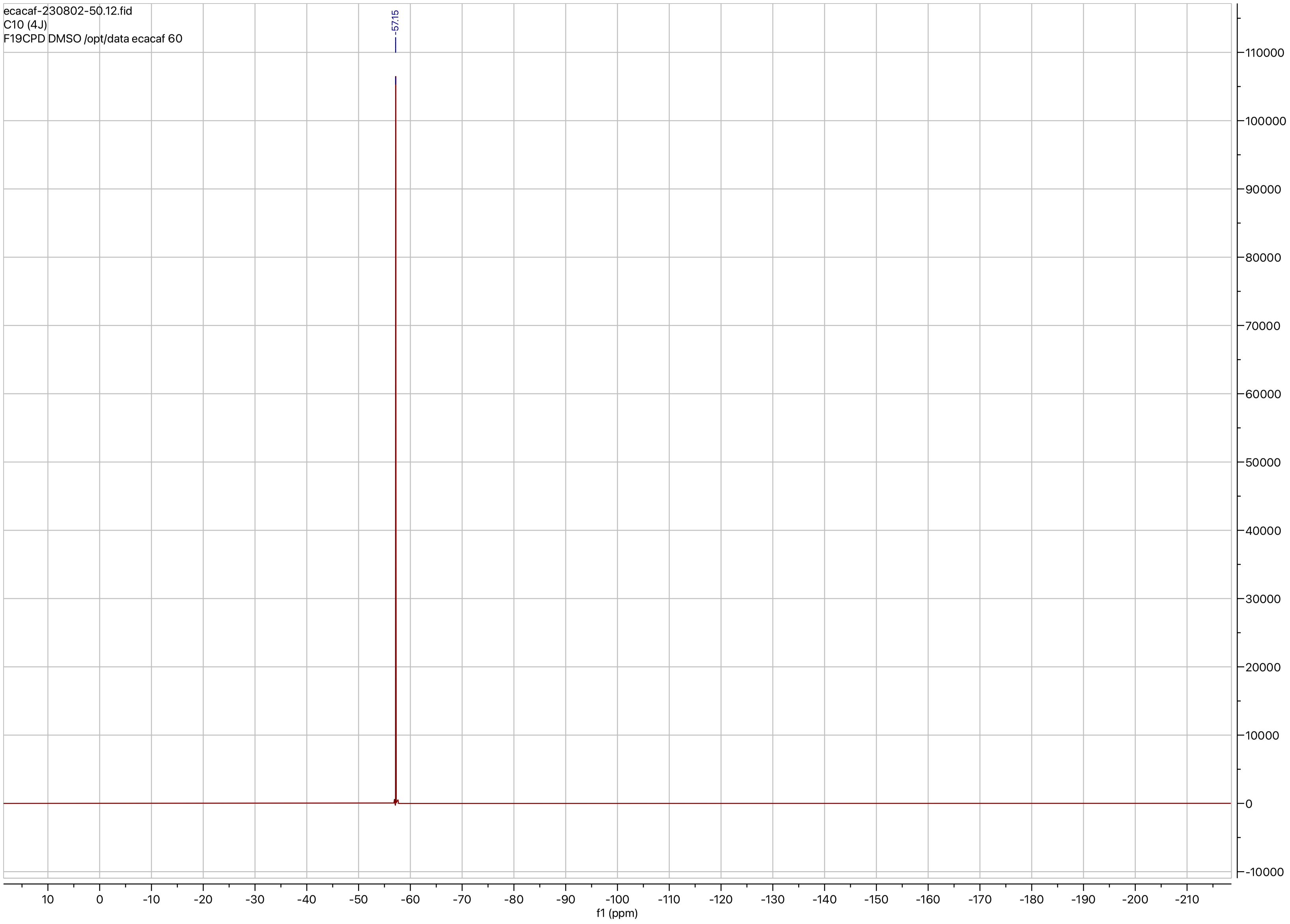
ecacaf-230802-50.12.fid
C10 (4J)
F19CPD DMSO /opt/data ecacaf 60
-57.15
f1 (ppm)

**Compound ID: C-11**

EW8457-643-P1B CDCl3 Bruker_A_400MHz

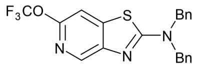

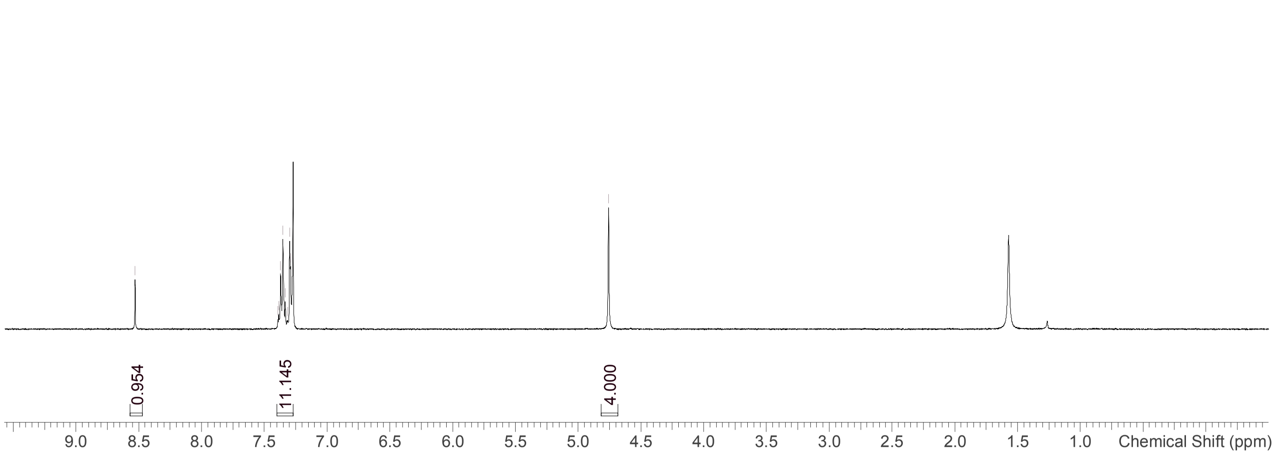


Supervisor: Tao Guo

| | |
|---|---|
| Acquisition Time (sec) | 3.0671 |
| Comment | EW8457-643-P1B CDCl3 Bruker_A_400MHz |
| Date | 14 Sep 2018 08:56:11 |
| Frequency (MHz) | 400.1300 |
| Nucleus | 1H |
| Number of Transients | 8 |
| Origin | spect |
| Original Points Count | 24576 |
| Owner | nmrsu |
| Points Count | 65536 |
| Pulse Sequence | zg30 |
| Receiver Gain | 195.78 |
| SW(cyclical) (Hz) | 8012.82 |
| Solvent | CHLOROFORM-d |
| Spectrum Offset (Hz) | 2395.1377 |
| Spectrum Type | standard |
| Sweep Width (Hz) | 8012.70 |
| Temperature (degree C) | 25.146 |

Operator:

Date:

Confidential, for research only not for regulatory filing

Confidential, for research only not for regulatory filing

LCMS REPORT

Compound ID : C - 11 ->
Sample ID : EW8457-643-P1B
Injection Date : 2018-09-14 09:40:03
Location : P2-B-02
Injection volume : 2.000
Acq Method : D:\DATA\1809\180914 7\5-95AB_R_220&254.M
Data Filename : D:\DATA\1809\180914 7\EW8457-643-P1B.D
Instrument : LCMS-B

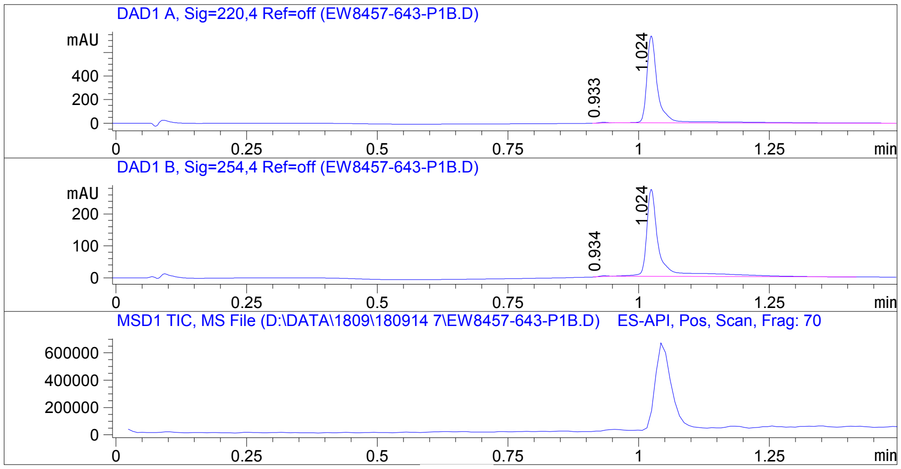


Report

Signal 1 : DAD1 A, Sig=220,4 Ref=off

| # | Meas. Ret. | Height | Width | Area | Area % |
|---|---|---|---|---|---|
| 1 | 0.933 | 6.888 | 0.017 | 7.212 | 0.641 |
| 2 | 1.024 | 739.280 | 0.022 | 1117.227 | 99.359 |

Signal 2 : DAD1 B, Sig=254,4 Ref=off

| # | Meas. Ret. | Height | Width | Area | Area % |
|---|---|---|---|---|---|
| 1 | 0.934 | 2.433 | 0.016 | 2.479 | 0.519 |
| 2 | 1.024 | 272.945 | 0.025 | 475.224 | 99.481 |

Operator:___________________

Date:____________________

Confidential, for research only not for regulatory filing

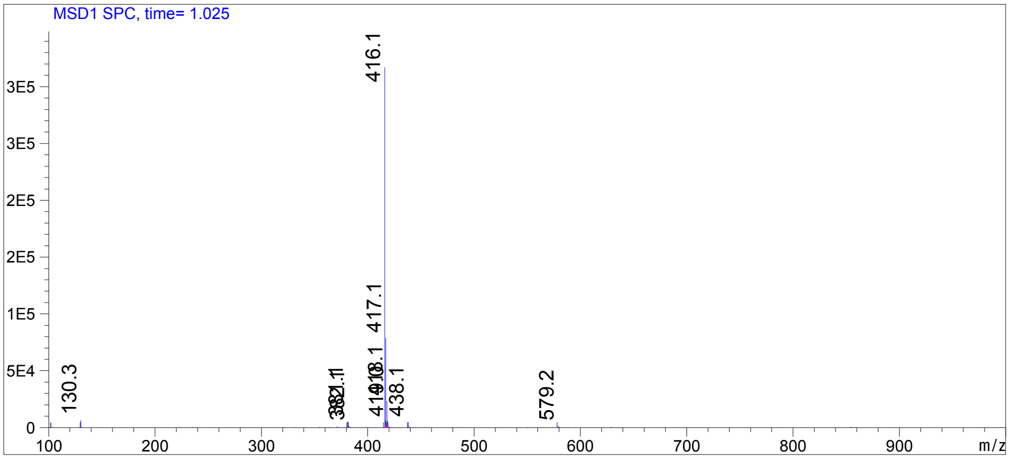

**Compound ID: C-12**

EW8457-471-P1A DMSO Bruker_A_400MHz

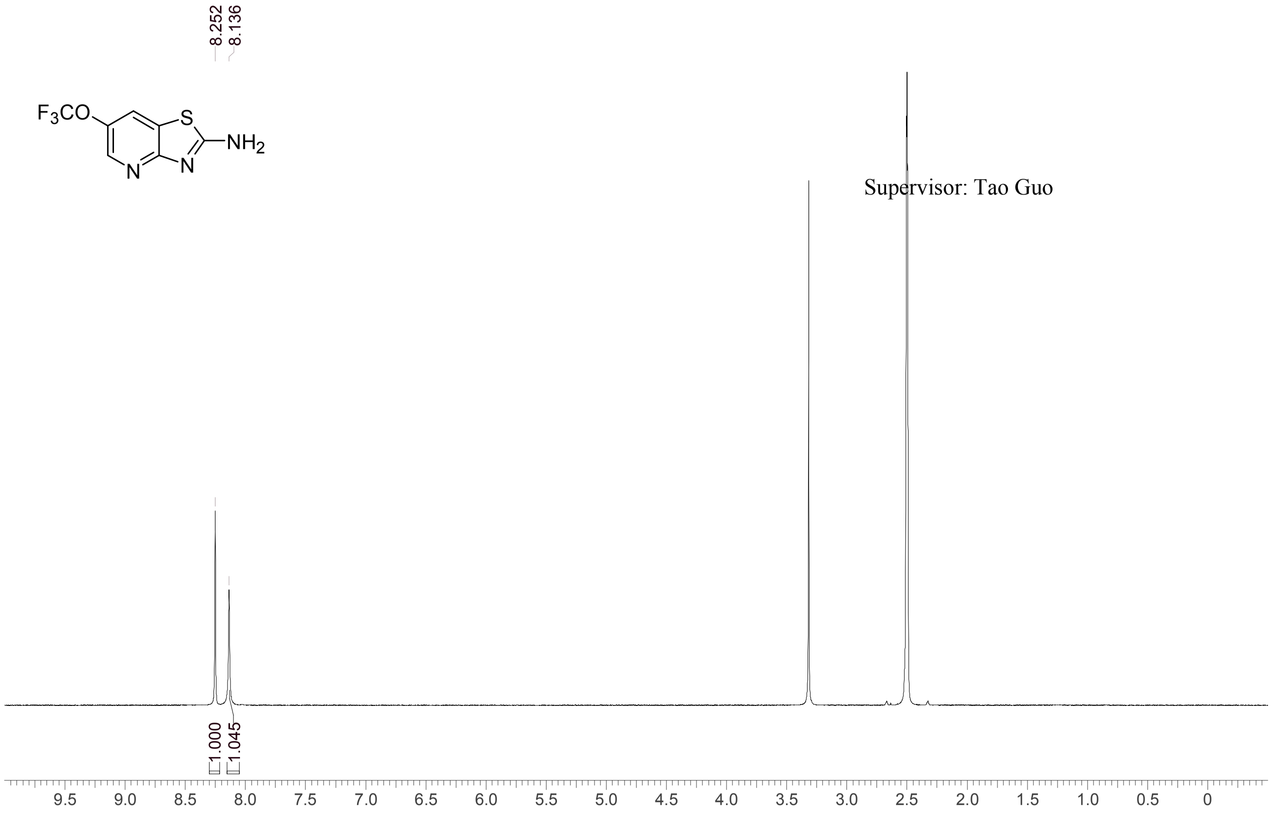


Supervisor: Tao Guo

| | |
|---|---|
| Acquisition Time (sec) | 3.0671 |
| Comment | EW8457-471-P1A DMSO Bruker_A_400MHz |
| Date | 05 Jul 2018 09:21:56 |
| Frequency (MHz) | 400.1300 |
| Nucleus | 1H |
| Number of Transients | 8 |
| Origin | spect |
| Original Points Count | 24576 |
| Owner | nmrsu |
| Points Count | 65536 |
| Pulse Sequence | zg30 |
| Receiver Gain | 175.36 |
| SW(cyclical) (Hz) | 8012.82 |
| Solvent | DMSO-d6 |
| Spectrum Offset (Hz) | 2397.5386 |
| Spectrum Type | standard |
| Sweep Width (Hz) | 8012.70 |
| Temperature (degree C) | 27.150 |

Operator:

Date:

Confidential, for research only not for regulatory filing

Confidential, for research only not for regulatory filing

### LCMS REPORT

Compound ID : C-12
Sample ID : EW8457-471-P1C
Injection Vol : 2ul
Location : vial83
Acq Method : d:\method\5-95AB_R_220&254_50.lcm
Org DataFile : D:\DATA\1807\180704\EW8457-471-P1C.lcd
Injection Date : 2018-07-04 15:04:10
Instrument : LCMS-P 15-105

Chromatogram

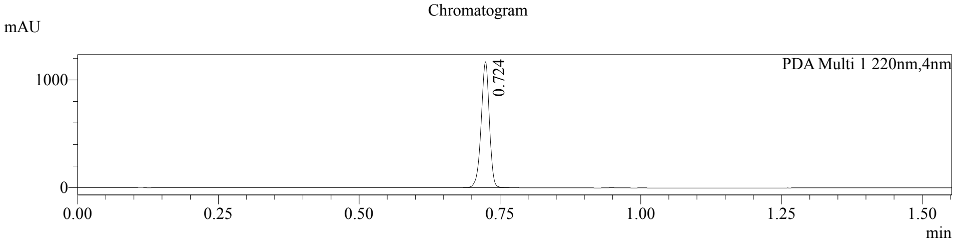


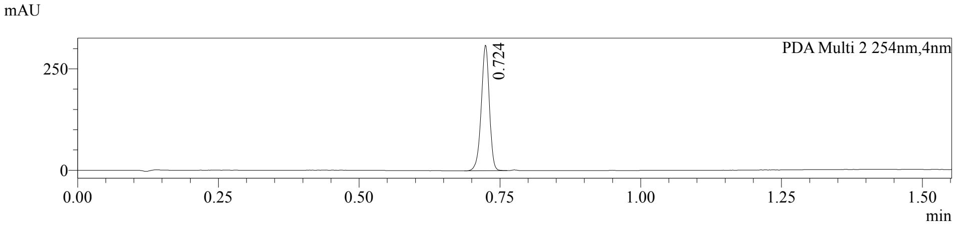


1 PDA Multi 1 / 220nm,4nm
2 PDA Multi 2 / 254nm,4nm

MS Chromatogram

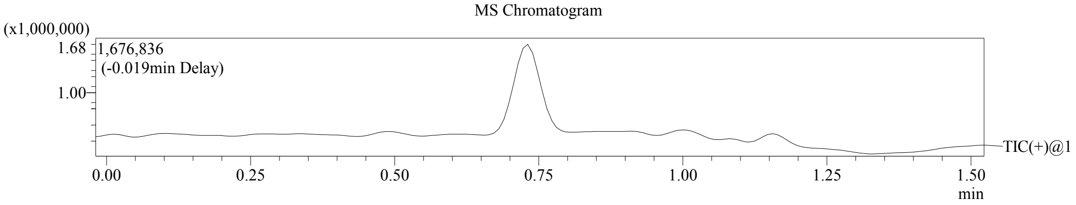


Integration Result

Peak Table

PDA Ch1 220nm

| Peak# | Ret. Time | Height | Height% | USP Width | Area | Area% |
|---|---|---|---|---|---|---|
| 1 | 0.724 | 1172193 | 100.000 | 0.029 | 1190012 | 100.000 |

Peak Table

PDA Ch2 254nm

| Peak# | Ret. Time | Height | Height% | USP Width | Area | Area% |
|---|---|---|---|---|---|---|
| 1 | 0.724 | 309330 | 100.000 | 0.028 | 310241 | 100.000 |

Confidential, for research only not for regulatory filing

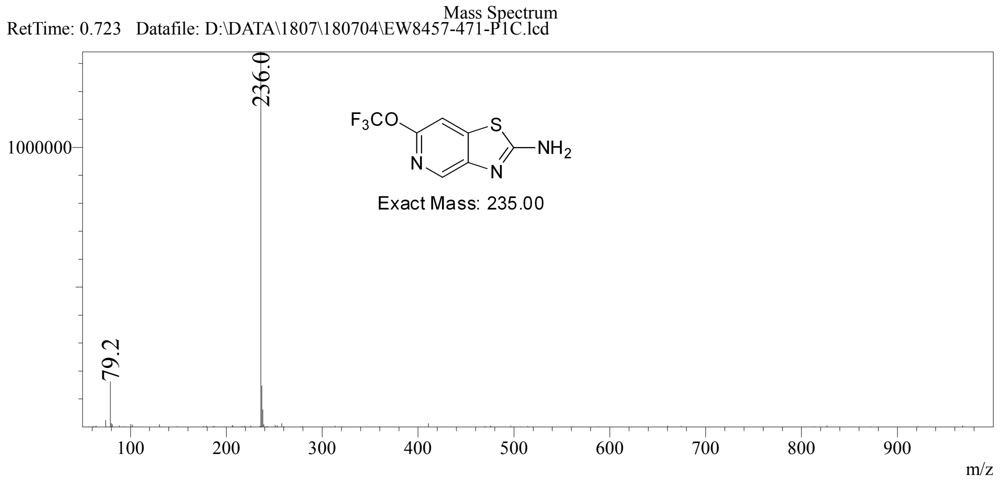

**Compound ID: C-13**

EW12557-76-P1E MeOD Bruker_E_400MHz

Supervisor: Tao Guo

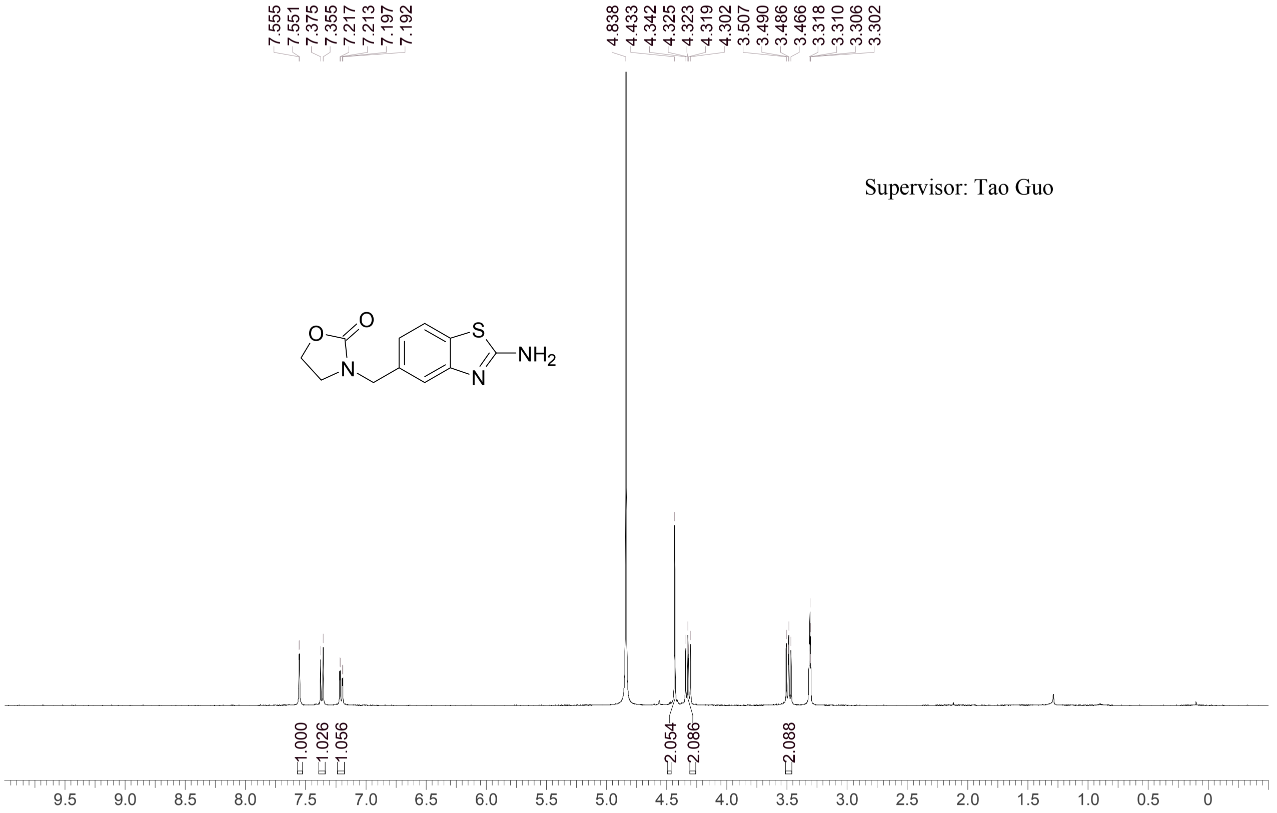


| | |
|---|---|
| Acquisition Time (sec) | 3.0671 |
| Comment | EW12557-76-P1E MeOD Bruker_E_400MHz |
| Date | 13 Aug 2018 13:24:39 |
| Frequency (MHz) | 400.1501 |
| Nucleus | 1H |
| Number of Transients | 8 |
| Origin | spect |
| Original Points Count | 24576 |
| Owner | nmrsu |
| Points Count | 65536 |
| Pulse Sequence | zg30 |
| Receiver Gain | 137.93 |
| SW(cyclical) (Hz) | 8012.82 |
| Solvent | METHANOL-d4 |
| Spectrum Offset (Hz) | 2393.3018 |
| Spectrum Type | standard |
| Sweep Width (Hz) | 8012.70 |
| Temperature (degree C) | 27.153 |

Operator: Date:

Confidential, for research only not for regulatory filing

Confidential, for research only not for regulatory filing

LCMS REPORT

Compound ID : C-13 ->
Sample ID : EW12557-76-P1P
Injection Date : 9-Aug-2018 14:01:03
Location : P2-A-07
Injection volume : 3.000
Acq Method : D:\DATA\1808\180809 18\10-80CD_2MIN_220&25
Data Filename : D:\DATA\1808\180809 18\EW12557-76-P1P.D
Instrument : LCMS-C

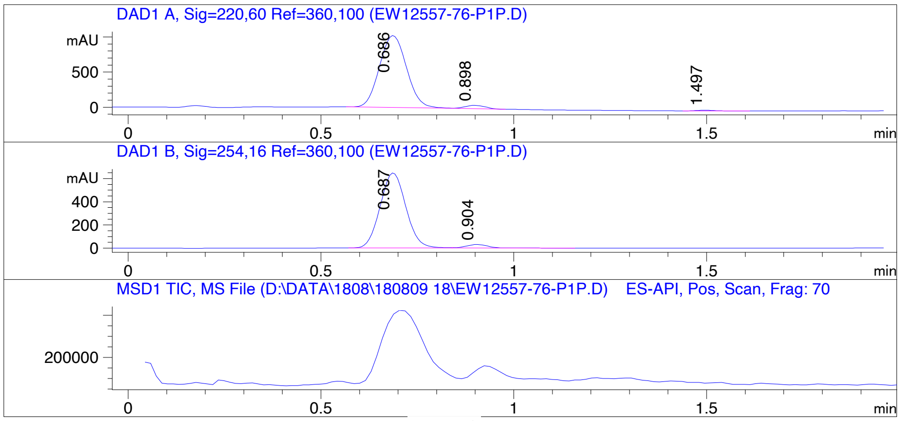


Report

==================================================================

Signal 1 : DAD1 A, Sig=220,60 Ref=360,100

| # | Meas. Ret. | Height | Width | Area | Area % |
|---|---|---|---|---|---|
| 1 | 0.686 | 1026.288 | 0.079 | 5000.127 | 96.126 |
| 2 | 0.898 | 47.699 | 0.059 | 167.289 | 3.216 |
| 3 | 1.497 | 10.018 | 0.052 | 34.239 | 0.658 |

Signal 2 : DAD1 B, Sig=254,16 Ref=360,100

| # | Meas. Ret. | Height | Width | Area | Area % |
|---|---|---|---|---|---|
| 1 | 0.687 | 647.622 | 0.072 | 3003.982 | 96.163 |
| 2 | 0.904 | 30.338 | 0.062 | 119.849 | 3.837 |

Operator:____________________

Date:____________________

Confidential, for research only not for regulatory filing

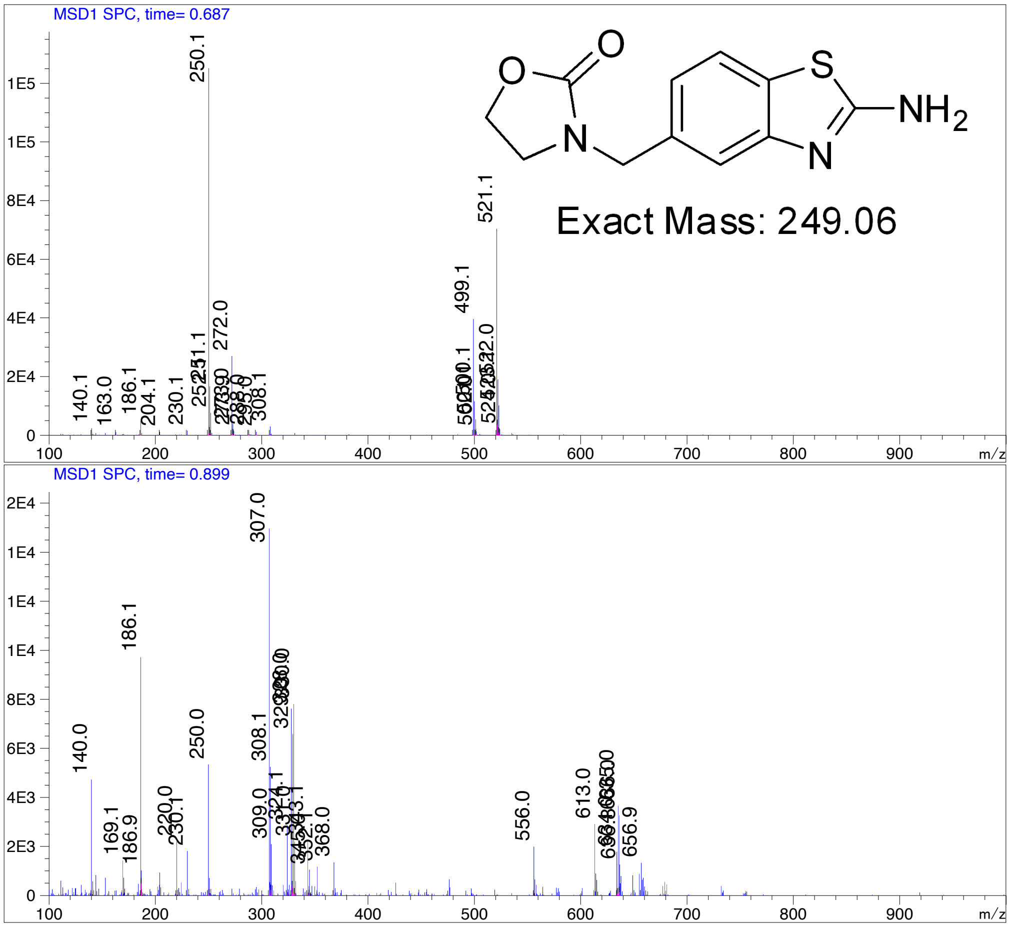

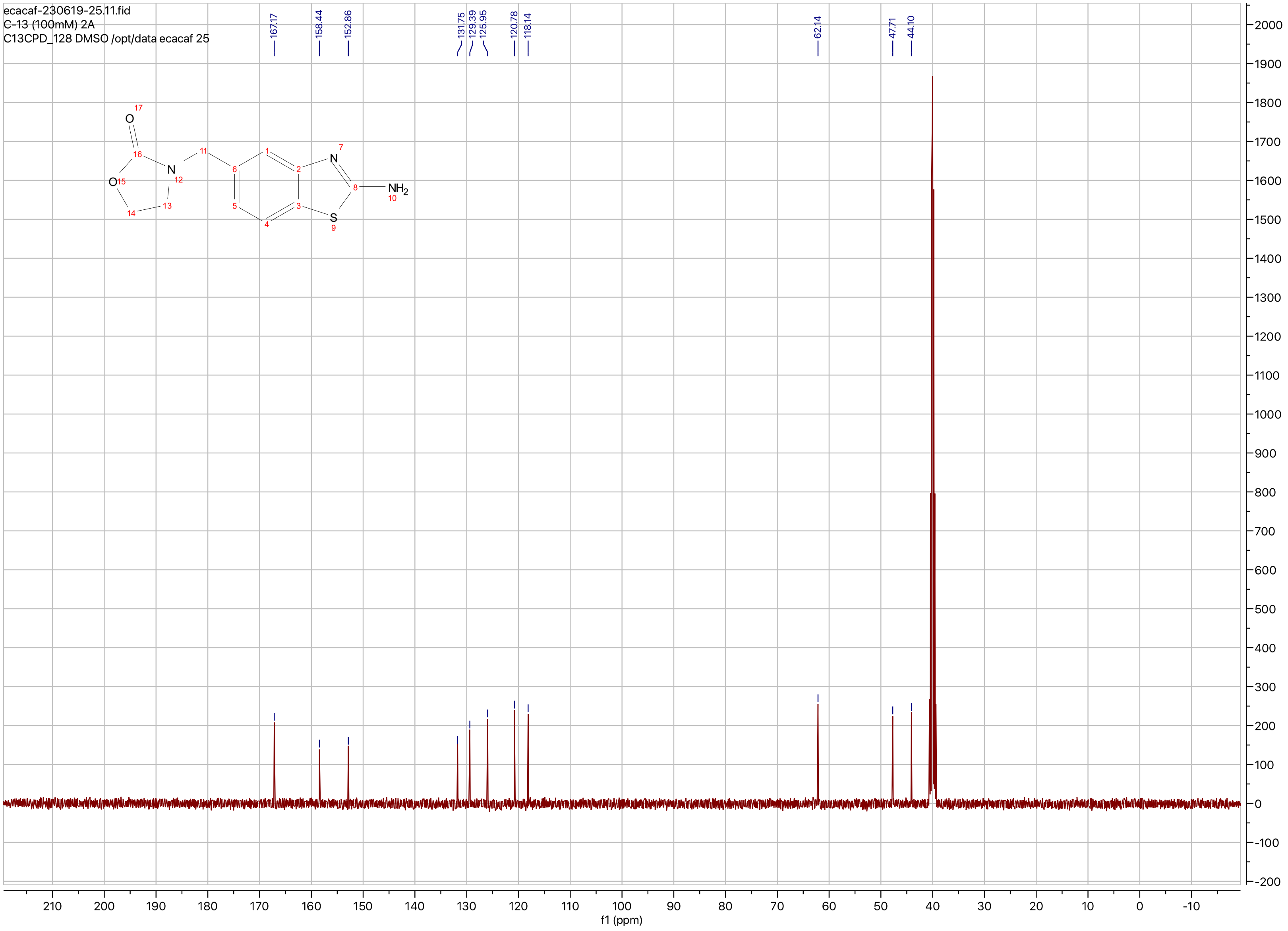
ecacaf-230619-25.11.fid
C-13 (100mM) 2A
C13CPD_128 DMSO /opt/data ecacaf 25
167.17
158.44
152.86
131.75
129.39
125.95
120.78
118.14
62.14
47.71
44.10
O
N
NH2
S
f1 (ppm)

**Compound ID: C-14**

EW8457-372-P1A DMSO Bruker_B_400MHz

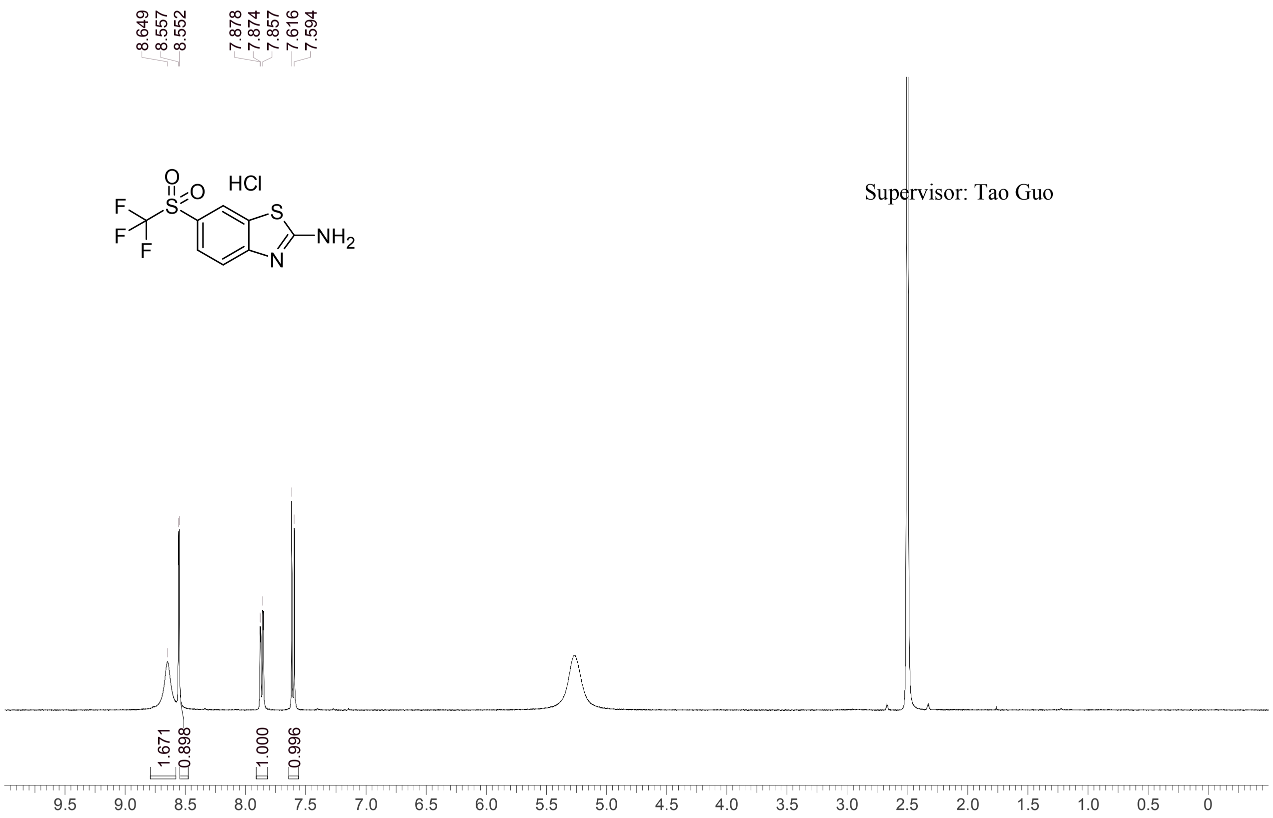


| | |
|---|---|
| Acquisition Time (sec) | 3.0671 |
| Comment | EW8457-372-P1A DMSO Bruker_B_400MHz |
| Date | 23 Apr 2018 08:43:01 |
| Frequency (MHz) | 400.1300 |
| Nucleus | 1H |
| Number of Transients | 8 |
| Origin | spect |
| Original Points Count | 24576 |
| Owner | nmrsu |
| Points Count | 65536 |
| Pulse Sequence | zg30 |
| Receiver Gain | 190.49 |
| SW(cyclical) (Hz) | 8012.82 |
| Solvent | DMSO-d6 |
| Spectrum Offset (Hz) | 2397.3784 |
| Spectrum Type | standard |
| Sweep Width (Hz) | 8012.70 |
| Temperature (degree C) | 21.206 |

Operator: Date:

Confidential, for research only not for regulatory filing

### LCMS REPORT

Compound ID : C-14
Sample ID : EW8457-372-P1B
Injection Vol : 3ul
Location : vial37
Acq Method : d:\method\5-95AB_R_220&254.lcm
Org DataFile : D:\data\1804\180422\EW8457-372-P1B.lcd
Injection Date : 2018/4/22 8:53:56
Instrument : LCMS-J

Chromatogram

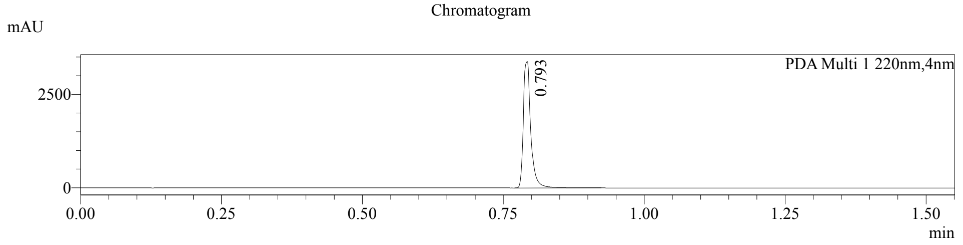


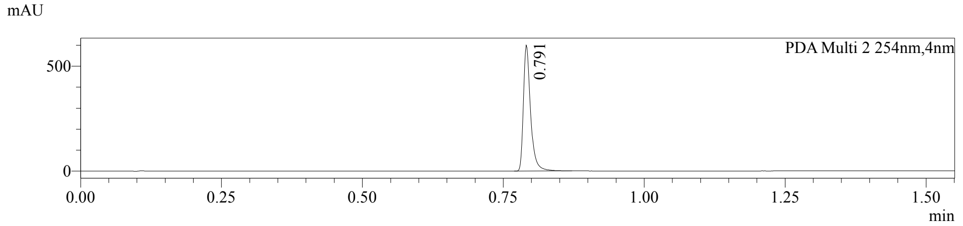


1 PDA Multi 1 / 220nm,4nm
2 PDA Multi 2 / 254nm,4nm

MS Chromatogram

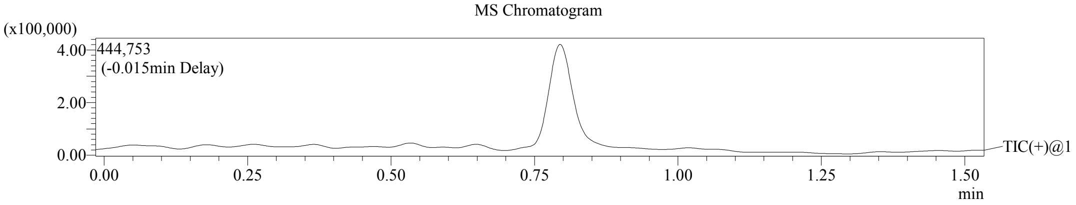


Integration Result

Peak Table

PDA Ch1 220nm

| Peak# | Ret. Time | Height | Height% | USP Width | Area | Area% |
|---|---|---|---|---|---|---|
| 1 | 0.793 | 3377402 | 100.000 | 0.023 | 3132328 | 100.000 |

PDA Ch2 254nm

| Peak# | Ret. Time | Height | Height% | USP Width | Area | Area% |
|---|---|---|---|---|---|---|
| 1 | 0.791 | 599474 | 100.000 | 0.025 | 533419 | 100.000 |

Operator:______________

Date:_________________

Confidential, for research only not for regulatory filing

Mass Spectrum

RetTime: 0.793  DateFile: D:\data\1804\180422\EW8457-372-P1B.lcd

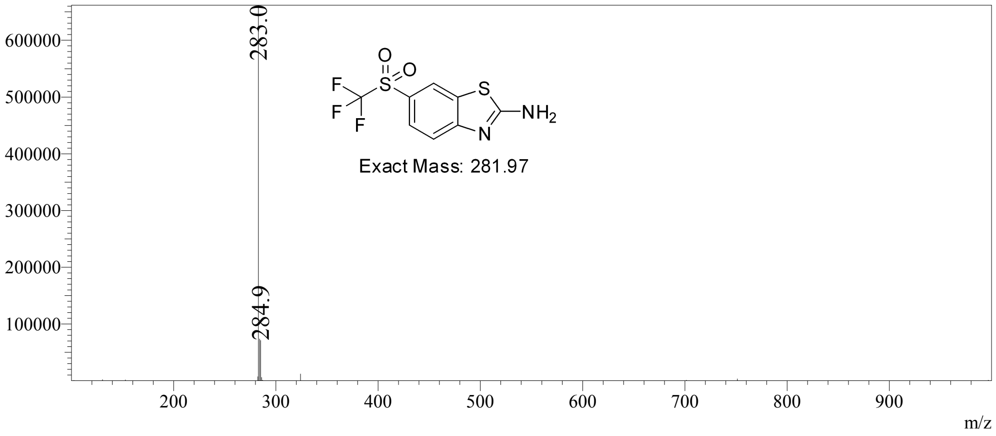


Confidential, for research only not for regulatory filing

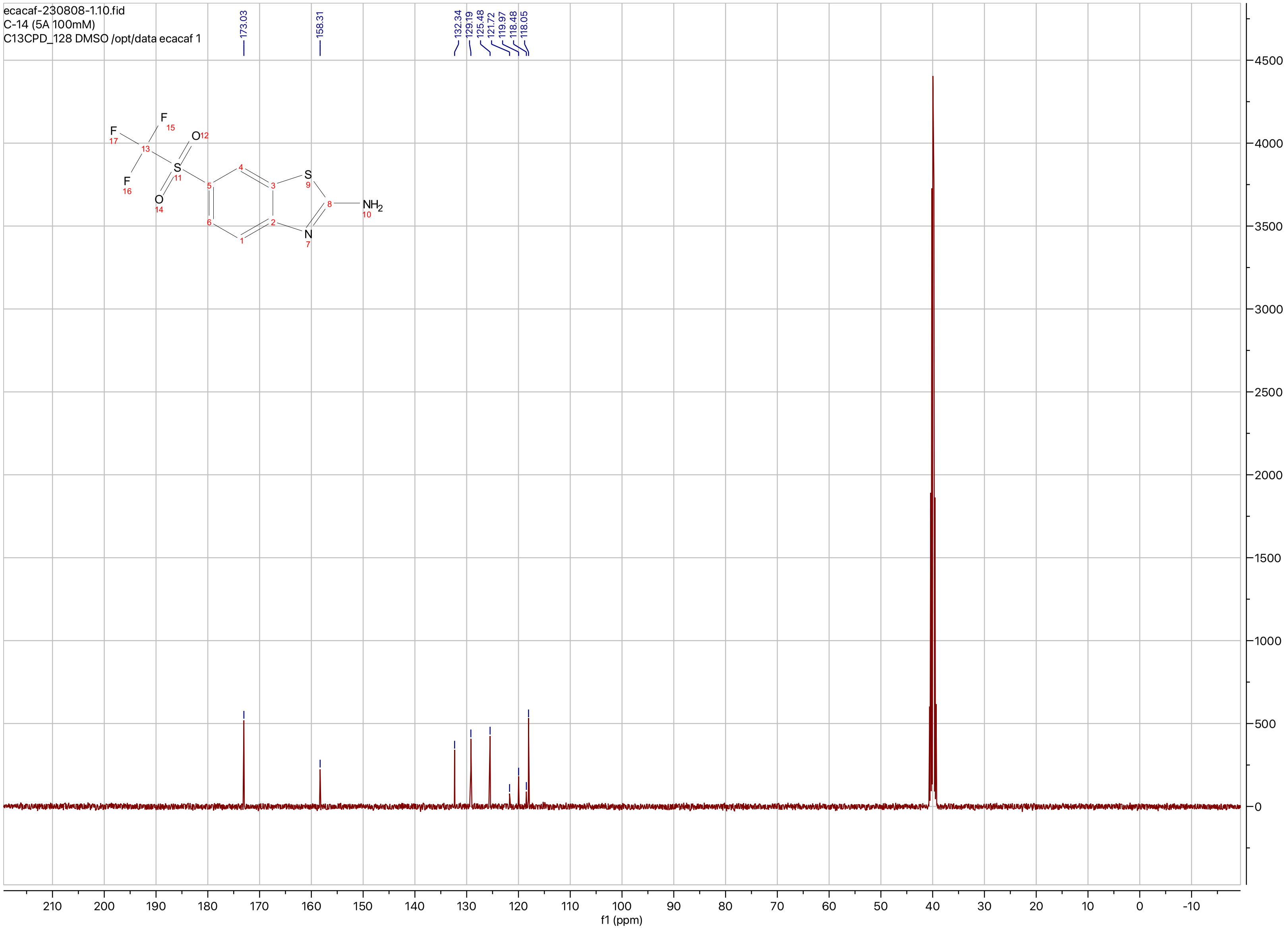
ecacaf-230808-1.10.fid
C-14 (5A 100mM)
C13CPD_128 DMSO /opt/data ecacaf 1
173.03
158.31
132.34
129.19
125.48
121.72
119.97
118.48
118.05
NH2
f1 (ppm)

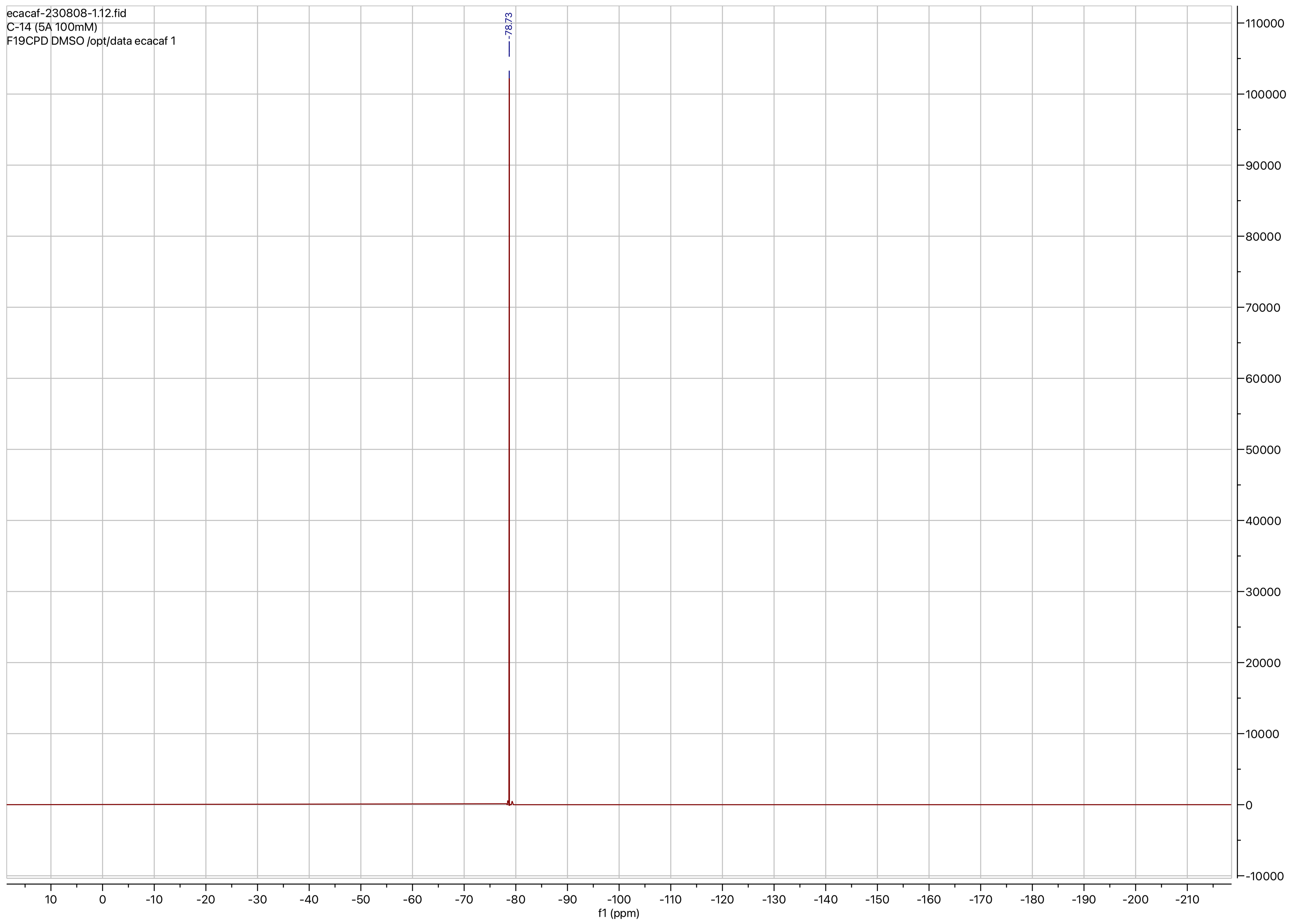
ecacaf-230808-1.12.fid
C-14 (5A 100mM)
F19CPD DMSO /opt/data ecacaf 1
-78.73
f1 (ppm)

**Compound ID: C-15**

EW8397-340-P1A DMSO Bruker_B_400MHz

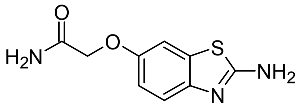


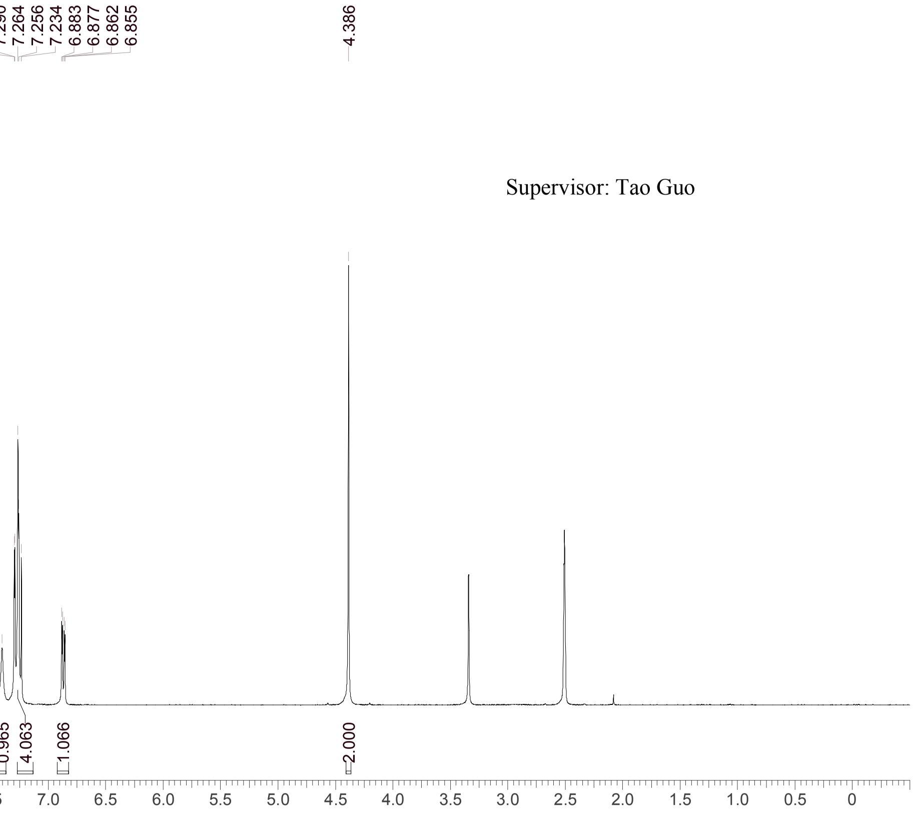


Supervisor: Tao Guo

| | |
|---|---|
| Acquisition Time (sec) | 3.0671 |
| Comment | EW8397-340-P1A DMSO Bruker_B_400MHz |
| Date | 29 May 2018 08:43:24 |
| Frequency (MHz) | 400.1300 |
| Nucleus | 1H |
| Number of Transients | 8 |
| Origin | spect |
| Original Points Count | 24576 |
| Owner | nmrsu |
| Points Count | 65536 |
| Pulse Sequence | zg30 |
| Receiver Gain | 190.49 |
| SW(cyclical) (Hz) | 8012.82 |
| Solvent | DMSO-d6 |
| Spectrum Offset (Hz) | 2400.7795 |
| Spectrum Type | standard |
| Sweep Width (Hz) | 8012.70 |
| Temperature (degree C) | 22.533 |

Operator:

Date:

Confidential, for research only not for regulatory filing

Confidential, for research only not for regulatory filing

### LCMS REPORT

Compound ID : C-15
Sample ID : EW8397-340-P1A1
Injection Vol : 2ul
Location : vial30
Acq Method : d:\method\5-95CD_R_220&254_POS.lcm
Org DataFile : D:\DATA\1805\180529\EW8397-340-P1A1.lcd
Injection Date : 29/05/2018 09:32:49
Instrument :LCMS-K 15-105

Chromatogram

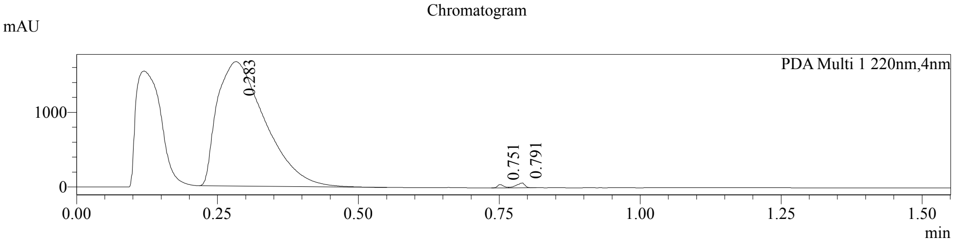


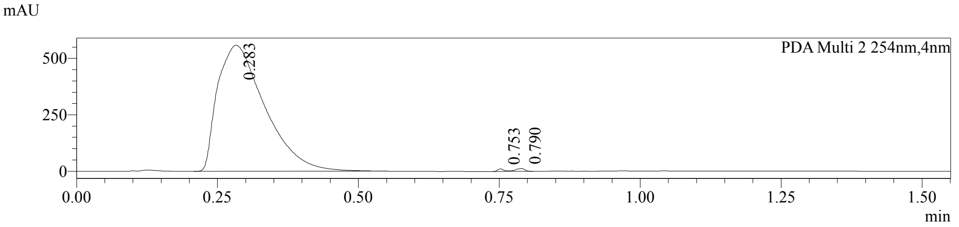


1 PDA Multi 1 / 220nm,4nm
2 PDA Multi 2 / 254nm,4nm

MS Chromatogram

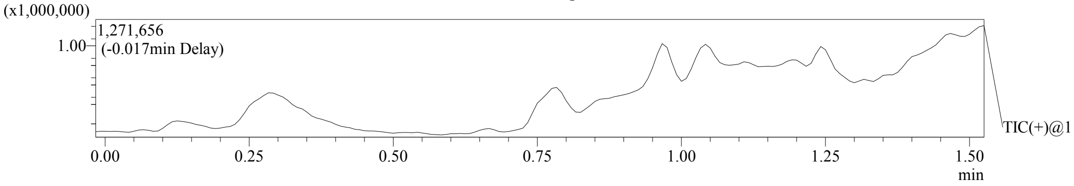


Integration Result

Peak Table

PDA Ch1 220nm

| Peak# | Ret. Time | Height | Height% | USP Width | Area | Area% |
|---|---|---|---|---|---|---|
| 1 | 0.283 | 1668549 | 93.941 | 0.153 | 9856257 | 98.981 |
| 2 | 0.751 | 43053 | 2.424 | 0.021 | 33589 | 0.337 |
| 3 | 0.791 | 64556 | 3.635 | 0.031 | 67880 | 0.682 |

Peak Table

PDA Ch2 254nm

| Peak# | Ret. Time | Height | Height% | USP Width | Area | Area% |
|---|---|---|---|---|---|---|
| 1 | 0.283 | 558815 | 96.032 | 0.150 | 3263986 | 99.301 |
| 2 | 0.753 | 11008 | 1.892 | 0.020 | 8586 | 0.261 |
| 3 | 0.790 | 12080 | 2.076 | 0.032 | 14405 | 0.438 |

Confidential, for research only not for regulatory filing

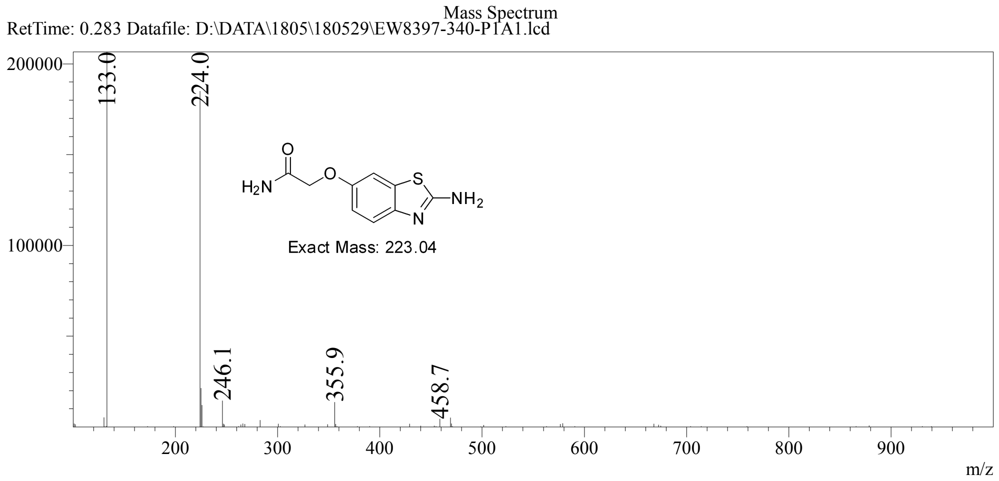

ecacaf-230619-20.11.fid
C-15 (100mM) 2C
C13CPD_128 DMSO /opt/data ecacaf 20

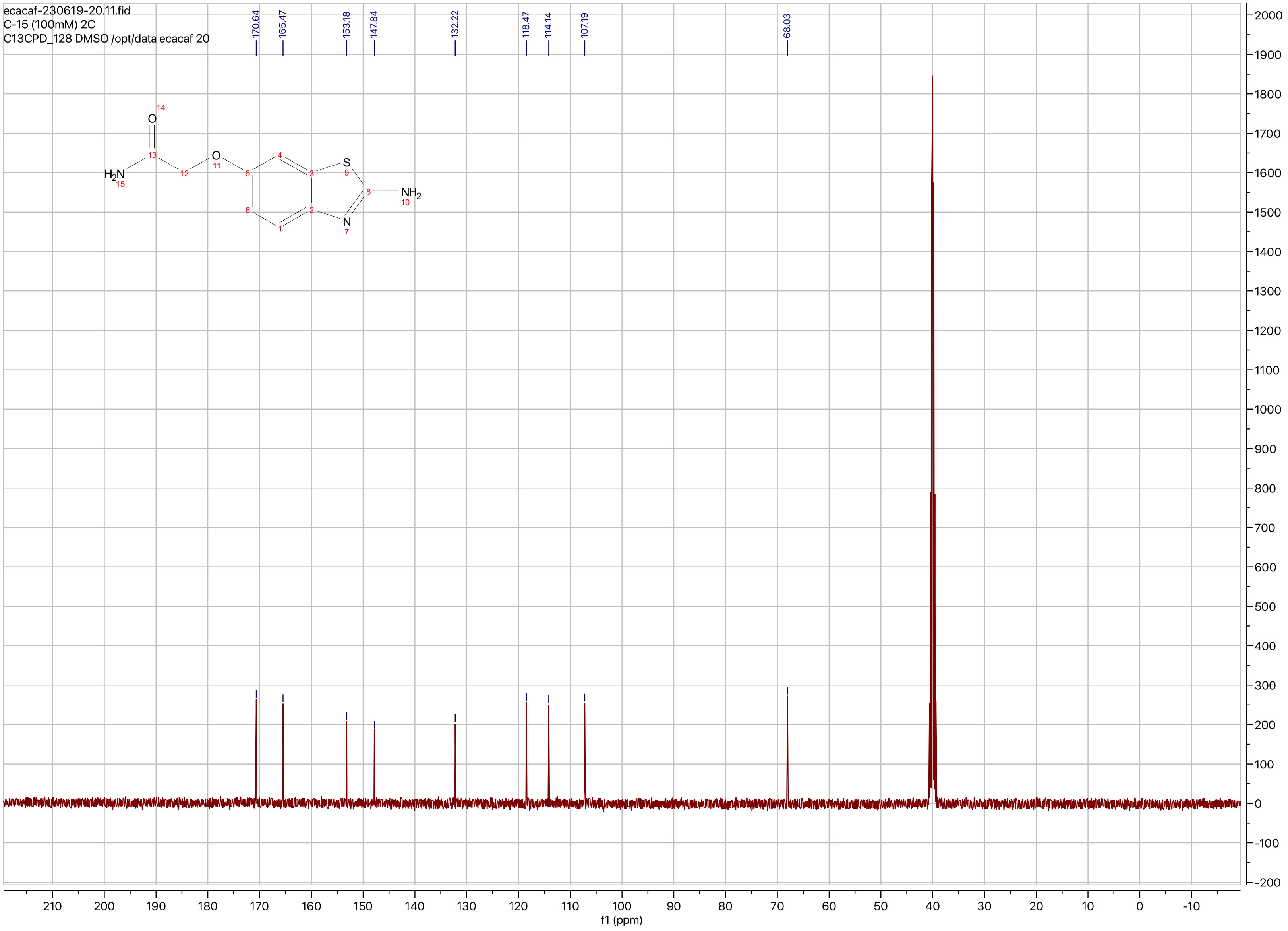

**Compound ID: C-16**

EW8397-341-P1A DMSO Bruker_B_400MHz

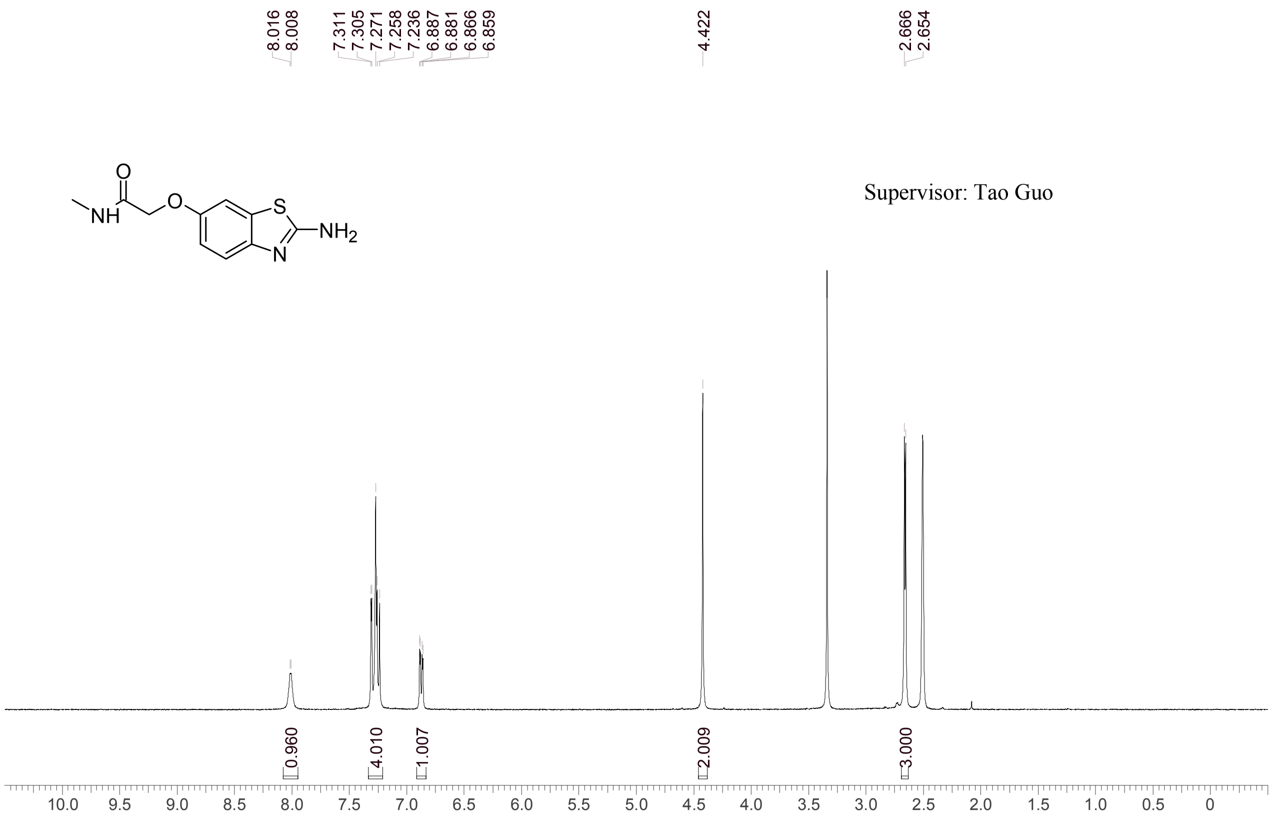


Supervisor: Tao Guo

| | |
|---|---|
| Acquisition Time (sec) | 3.0671 |
| Comment | EW8397-341-P1A DMSO Bruker_B_400MHz |
| Date | 29 May 2018 08:49:06 |
| Frequency (MHz) | 400.1300 |
| Nucleus | 1H |
| Number of Transients | 8 |
| Origin | spect |
| Original Points Count | 24576 |
| Owner | nmrsu |
| Points Count | 65536 |
| Pulse Sequence | zg30 |
| Receiver Gain | 190.49 |
| SW(cyclical) (Hz) | 8012.82 |
| Solvent | DMSO-d6 |
| Spectrum Offset (Hz) | 2400.7795 |
| Spectrum Type | standard |
| Sweep Width (Hz) | 8012.70 |
| Temperature (degree C) | 22.510 |

Operator: Date:

Confidential, for research only not for regulatory filing

Report(Report Editor)  Status:Manual Integration/Temporary
Confidential, for research only not for regulatory filing

### LCMS REPORT

Compound ID : C-16
Sample ID : EW8397-341-P1A1
Injection Vol : 0.5ul
Location : vial43
Acq Method : D:\METHOD\XBridge_C18_5cm\WUXICD05.lcm
Org DateFile : D:\2018\05\0529\EW8397-341-P1A1.lcd
Injection Date : 5/29/2018 11:18:37 AM
Instrument : CAS-WH-LCMS-S
operator : yu_xiujuan

Chromatogram

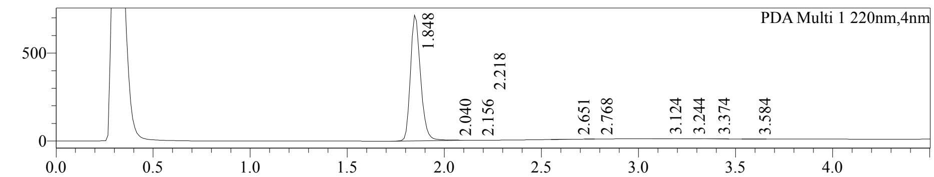


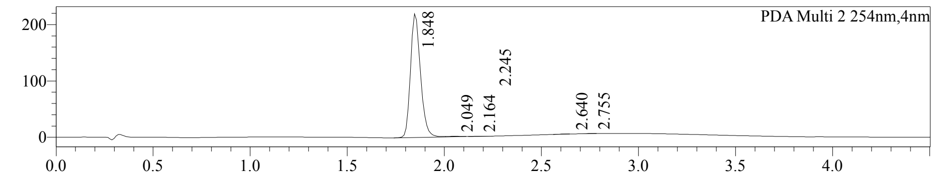


1 PDA Multi 1 / 220nm,4nm
2 PDA Multi 2 / 254nm,4nm

MS Chromatogram

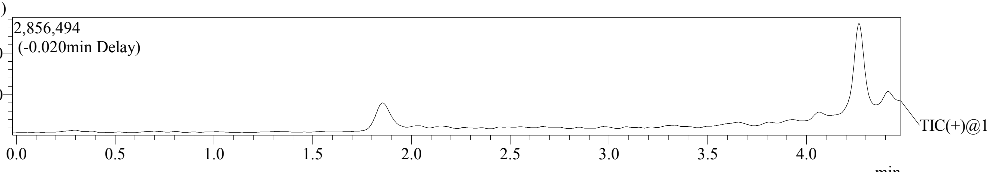


Integration Result

Peak Table

PDA Ch1 220nm

| Peak# | Ret. Time | Height | Tailing Factor | Resolution | Separation F. | Area | Area% |
|---|---|---|---|---|---|---|---|
| 1 | 1.848 | 714812 | 1.223 | -- | -- | 2585957 | 99.387 |
| 2 | 2.040 | 2931 | -- | -- | -- | 9602 | 0.369 |
| 3 | 2.156 | 152 | 1.006 | -- | 1.611 | 252 | 0.010 |
| 4 | 2.218 | 142 | 0.834 | 1.251 | 1.201 | 351 | 0.013 |
| 5 | 2.651 | -18 | -- | 5.708 | 2.169 | 1671 | 0.064 |
| 6 | 2.768 | 26 | -- | -- | 1.146 | 553 | 0.021 |
| 7 | 3.124 | 531 | 1.044 | -- | 1.387 | 1554 | 0.060 |
| 8 | 3.244 | 298 | 1.744 | 1.239 | 1.095 | 925 | 0.036 |
| 9 | 3.374 | 222 | 2.430 | 1.208 | 1.093 | 644 | 0.025 |

| Peak# | Ret. Time | Height | Tailing Factor | Resolution | Separation F. | Area | Area% |
|---|---|---|---|---|---|---|---|
| 10 | 3.584 | 135 | 1.040 | 2.023 | 1.138 | 391 | 0.015 |

Peak Table

PDA Ch2 254nm

| Peak# | Ret. Time | Height | Tailing Factor | Resolution | Separation F. | Area | Area% |
|---|---|---|---|---|---|---|---|
| 1 | 1.848 | 219556 | 1.215 | -- | -- | 786449 | 99.230 |
| 2 | 2.049 | 1068 | -- | 1.634 | -- | 4015 | 0.507 |
| 3 | 2.164 | 135 | -- | 1.234 | 1.572 | 379 | 0.048 |
| 4 | 2.245 | -44 | -- | 1.286 | 1.256 | 287 | 0.036 |
| 5 | 2.640 | -0 | -- | 4.843 | 1.994 | 867 | 0.109 |
| 6 | 2.755 | 172 | 0.828 | 1.514 | 1.145 | 553 | 0.070 |

Operator:______________

Date:_________________

Report(Report Editor)    Status:Manual Integration/Temporary
Confidential, for research only not for regulatory filing

Mass Spectrum
RetTime: 1.847 Datafile: D:\2018\05\0529\EW8397-341-P1A1.lcd

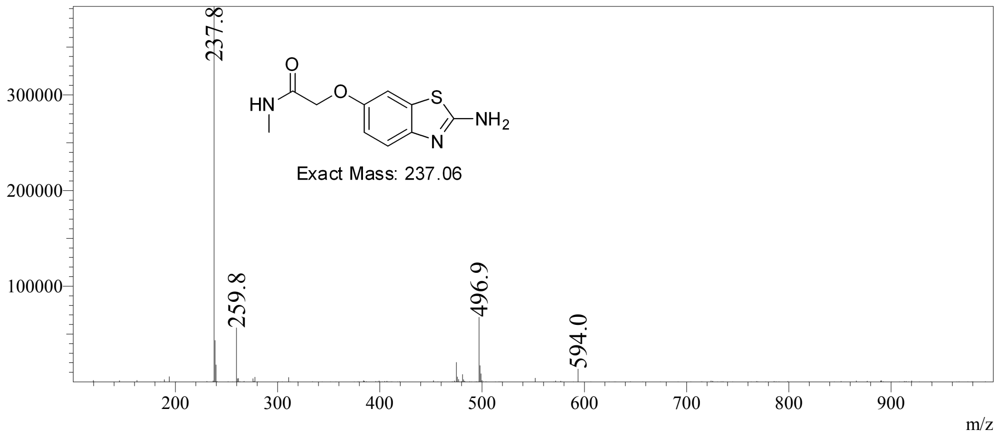

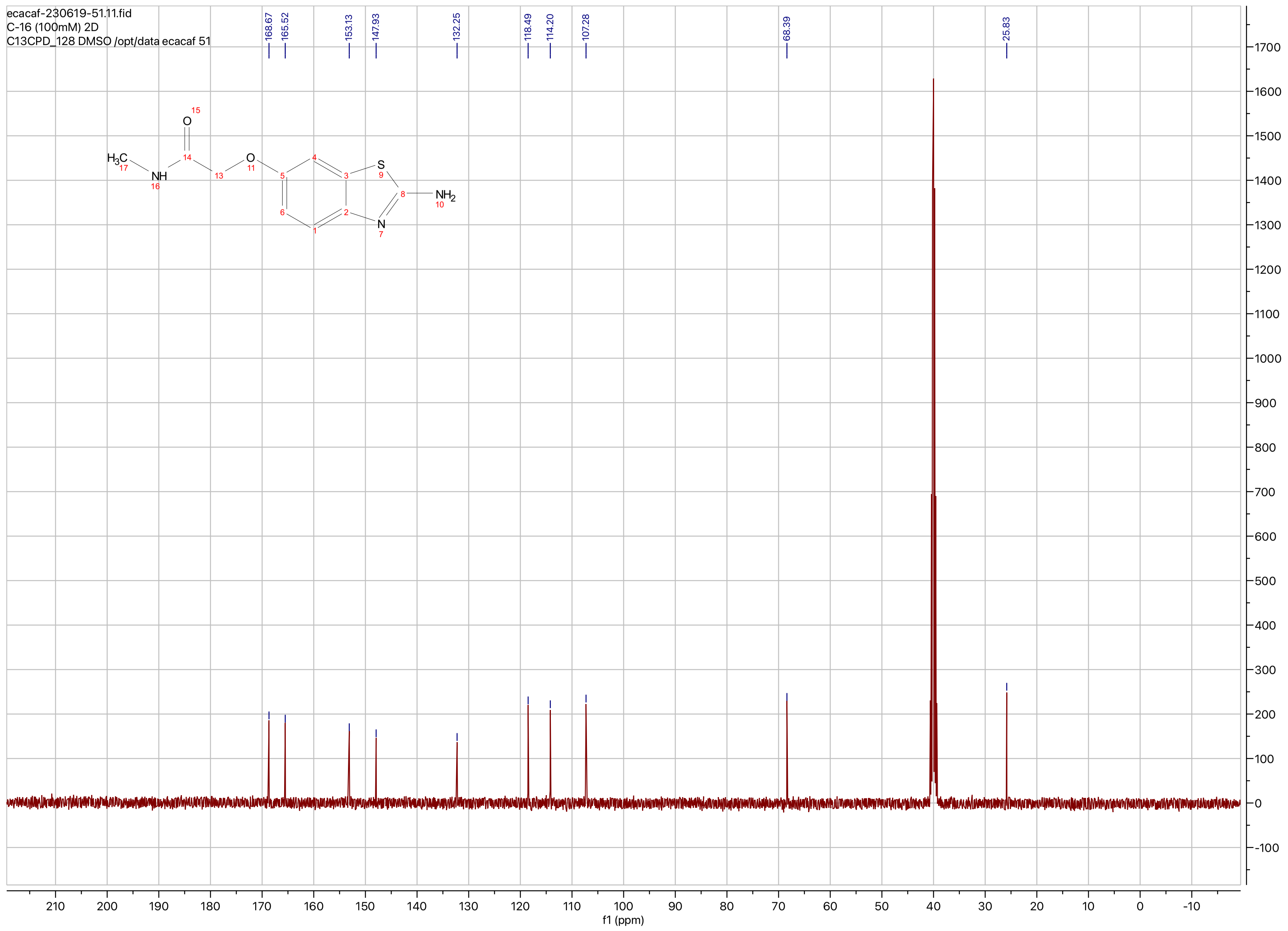
ecacaf-230619-51.11.fid
C-16 (100mM) 2D
C13CPD_128 DMSO /opt/data ecacaf 51
168.67
165.52
153.13
147.93
132.25
118.49
114.20
107.28
68.39
25.83
H3C
NH
O
O
S
N
NH2
f1 (ppm)

**Compound ID: C-17**

EW8397-342-P1A DMSO Bruker_B_400MHz

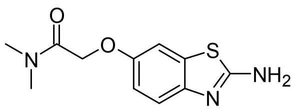

Supervisor: Tao Guo

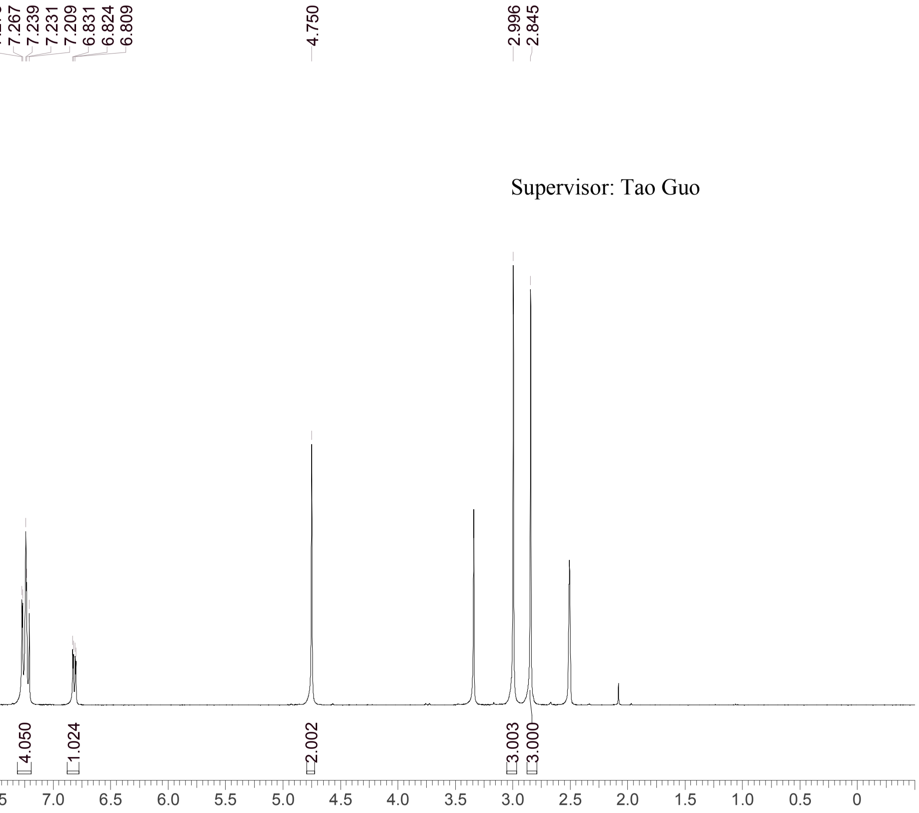


| | |
|---|---|
| Acquisition Time (sec) | 3.0671 |
| Comment | EW8397-342-P1A DMSO Bruker_B_400MHz |
| Date | 29 May 2018 08:51:22 |
| Frequency (MHz) | 400.1300 |
| Nucleus | 1H |
| Number of Transients | 8 |
| Origin | spect |
| Original Points Count | 24576 |
| Owner | nmrsu |
| Points Count | 65536 |
| Pulse Sequence | zg30 |
| Receiver Gain | 190.49 |
| SW(cyclical) (Hz) | 8012.82 |
| Solvent | DMSO-d6 |
| Spectrum Offset (Hz) | 2400.7795 |
| Spectrum Type | standard |
| Sweep Width (Hz) | 8012.70 |
| Temperature (degree C) | 22.519 |

Operator: Date:

Confidential, for research only not for regulatory filing

Confidential, for research only not for regulatory filing

### LCMS REPORT

Compound ID : C-17
Sample ID : EW8397-342-P1A1
Injection Vol : 2ul
Location : vial32
Acq Method : d:\method\5-95CD_R_220&254_POS.lcm
Org DataFile : D:\DATA\1805\180529\EW8397-342-P1A1.lcd
Injection Date : 29/05/2018 09:36:46
Instrument :LCMS-K 15-105

Chromatogram

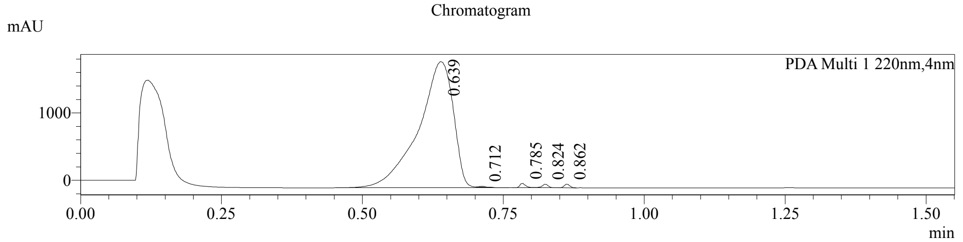


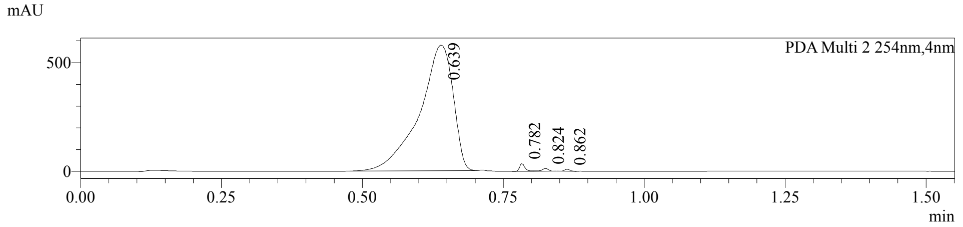


1 PDA Multi 1 / 220nm,4nm
2 PDA Multi 2 / 254nm,4nm

MS Chromatogram

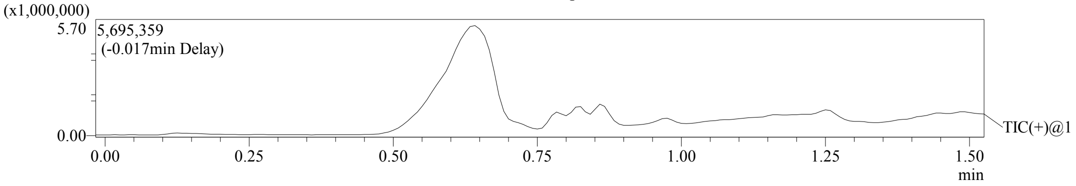


Integration Result

Peak Table

PDA Ch1 220nm

| Peak# | Ret. Time | Height | Height% | USP Width | Area | Area% |
|---|---|---|---|---|---|---|
| 1 | 0.639 | 1872834 | 91.156 | 0.101 | 8052386 | 98.298 |
| 2 | 0.712 | 14073 | 0.685 | 0.024 | 12225 | 0.149 |
| 3 | 0.785 | 62115 | 3.023 | 0.022 | 48290 | 0.589 |
| 4 | 0.824 | 51913 | 2.527 | 0.023 | 42020 | 0.513 |
| 5 | 0.862 | 53604 | 2.609 | 0.021 | 36919 | 0.451 |

Peak Table

PDA Ch2 254nm

| Peak# | Ret. Time | Height | Height% | USP Width | Area | Area% |
|---|---|---|---|---|---|---|
| 1 | 0.639 | 577382 | 91.123 | 0.097 | 2396138 | 98.350 |
| 2 | 0.782 | 34802 | 5.492 | 0.020 | 23417 | 0.961 |
| 3 | 0.824 | 12782 | 2.017 | 0.023 | 10668 | 0.438 |
| 4 | 0.862 | 8664 | 1.367 | 0.022 | 6110 | 0.251 |

Confidential, for research only not for regulatory filing

Mass Spectrum

RetTime: 0.642 Datafile: D:\DATA\1805\180529\EW8397-342-P1A1.lcd

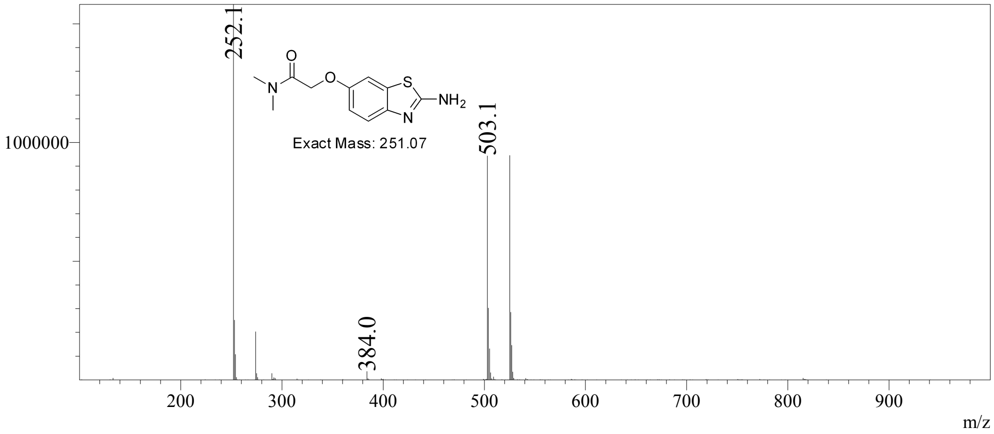

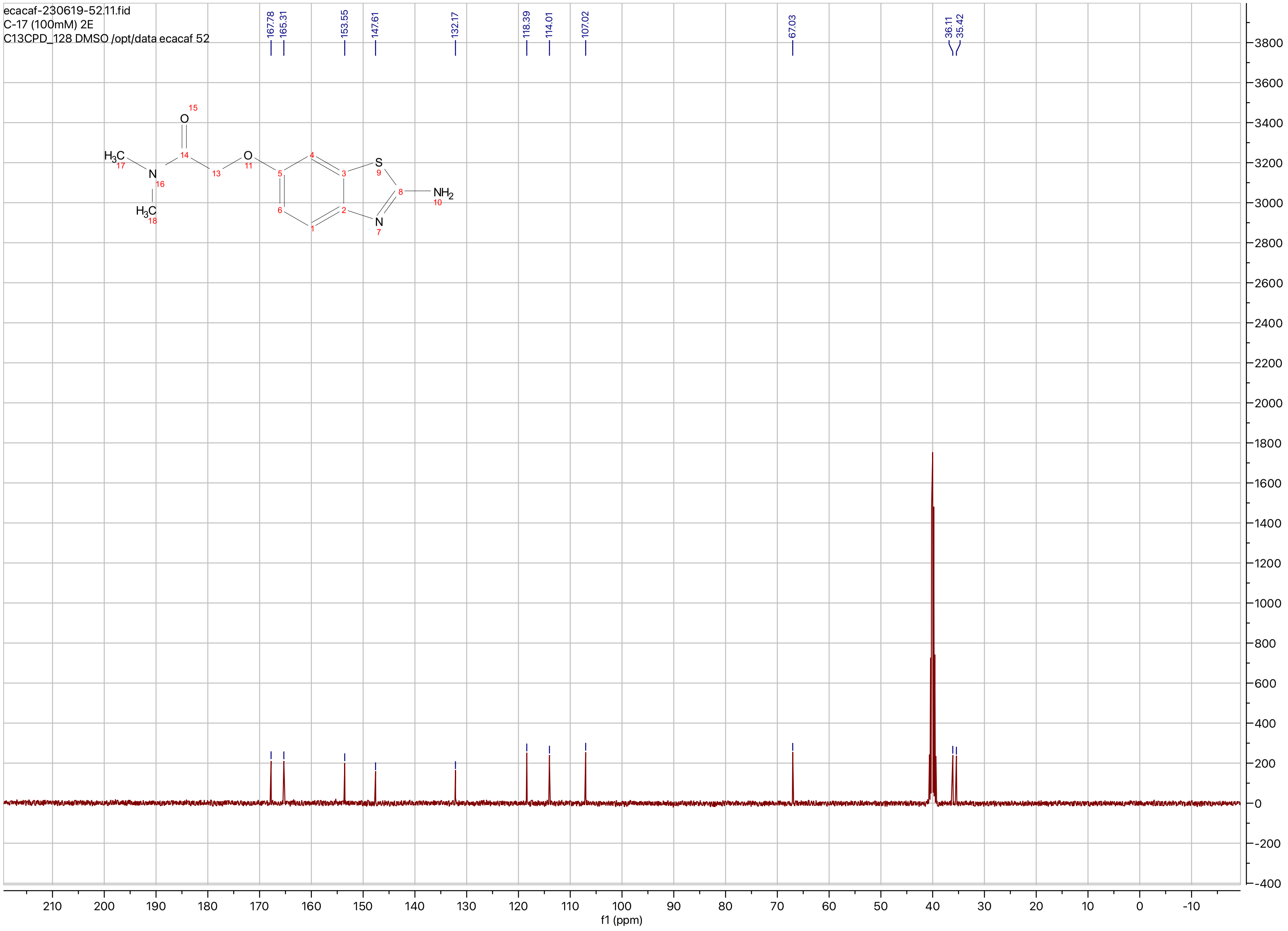

ecacaf-230619-52.11.fid
C-17 (100mM) 2E
C13CPD_128 DMSO /opt/data ecacaf 52
167.78
165.31
153.55
147.61
132.17
118.39
114.01
107.02
67.03
36.11
35.42
$H_3C$
N
O
S
$NH_2$
f1 (ppm)

**Compound ID: C-18**

ET28001-42-P1C DMSO Bruker_K_400MHz

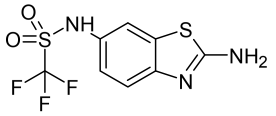

FA

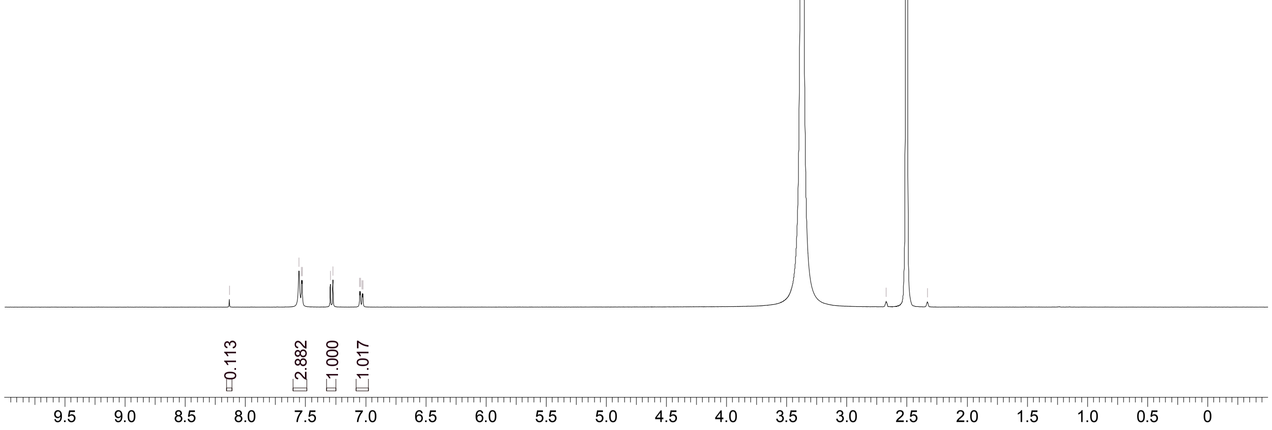


Supervisor: Tao Guo

| | |
|---|---|
| Acquisition Time (sec) | 1.9988 |
| Comment | ET28001-42-P1C DMSO Bruker_K_400MHz |
| Date | 21 Jun 2019 02:46:56 |
| Frequency (MHz) | 400.1300 |
| Nucleus | 1H |
| Number of Transients | 8 |
| Origin | Avance |
| Original Points Count | 16384 |
| Owner | nmr |
| Points Count | 65536 |
| Pulse Sequence | zg30 |
| Receiver Gain | 101.00 |
| SW(cyclical) (Hz) | 8196.72 |
| Solvent | DMSO-d6 |
| Spectrum Offset (Hz) | 2465.1196 |
| Spectrum Type | standard |
| Sweep Width (Hz) | 8196.60 |
| Temperature (degree C) | 22.959 |

Confidential. For research only Not for regulatory filing

Operator:

Date:

LCMS REPORT

Compound ID : C - 18 ->
Sample ID : ET28001-42-P1C
Injection Date : 21. Jun. 2019
Inj. Vol. : 0.50 ul
Location : P2-B-04
Acq Method : D:\DATA\190621-HD 12\5_95AB_6min-220.M
Data Filename : D:\DATA\190621-HD 12\ET28001-42-P1C.D
Instrument : H

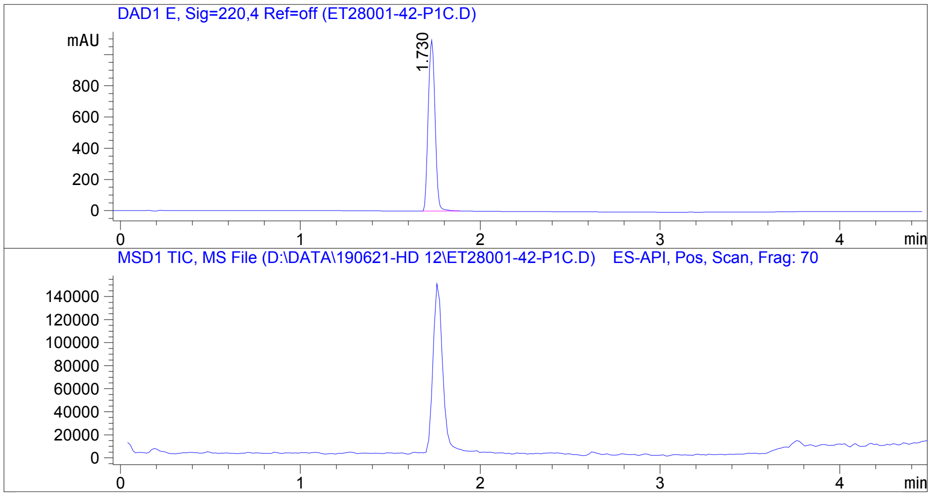


Integration Result

Signal 1 : DAD1 E, Sig=220,4 Ref=off

| Peak # | RT [min] | Area | Height | Height % | Width [min] | Area % |
|---|---|---|---|---|---|---|
| 1 | 1.730 | 2909.633 | 1096.893 | 100.000 | 0.043 | 100.000 |

Operator:_____________

Date:_________________

Confidential. For research information only.

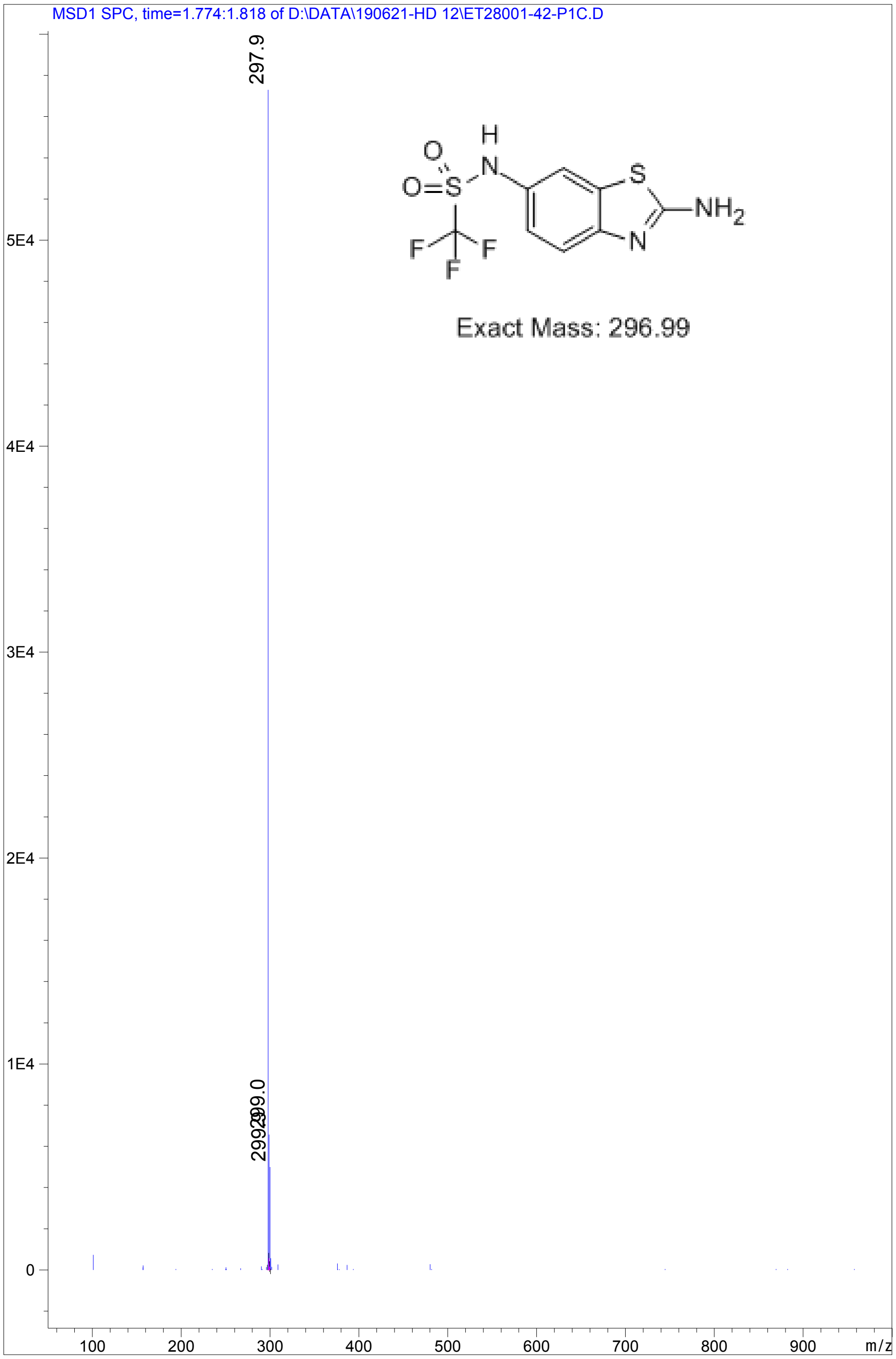


Confidential. For research information only.

ecacaf-230619-39.11.fid
C-18 (100mM) 2F
C13CPD_128 DMSO /opt/data ecacaf 39

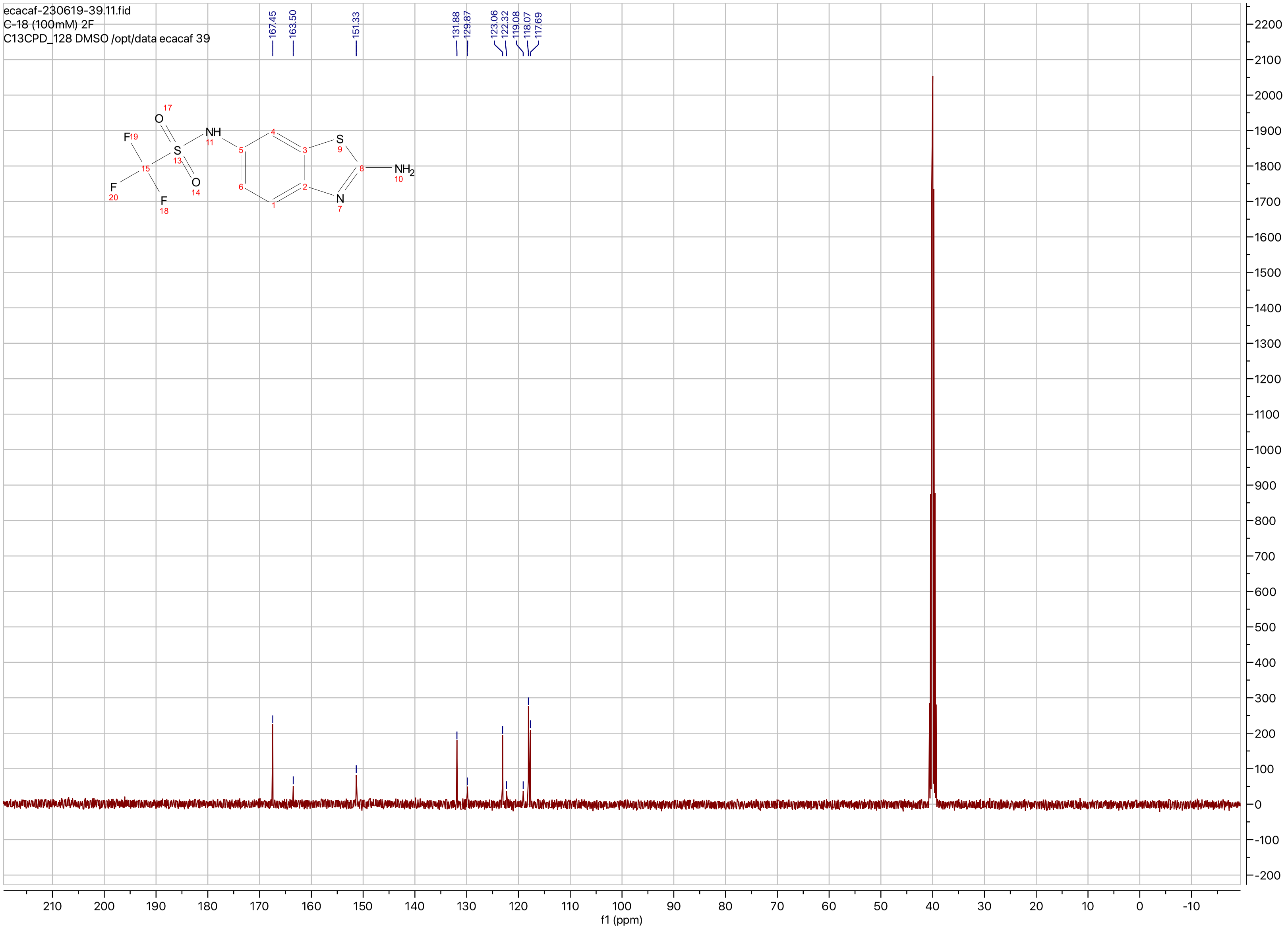

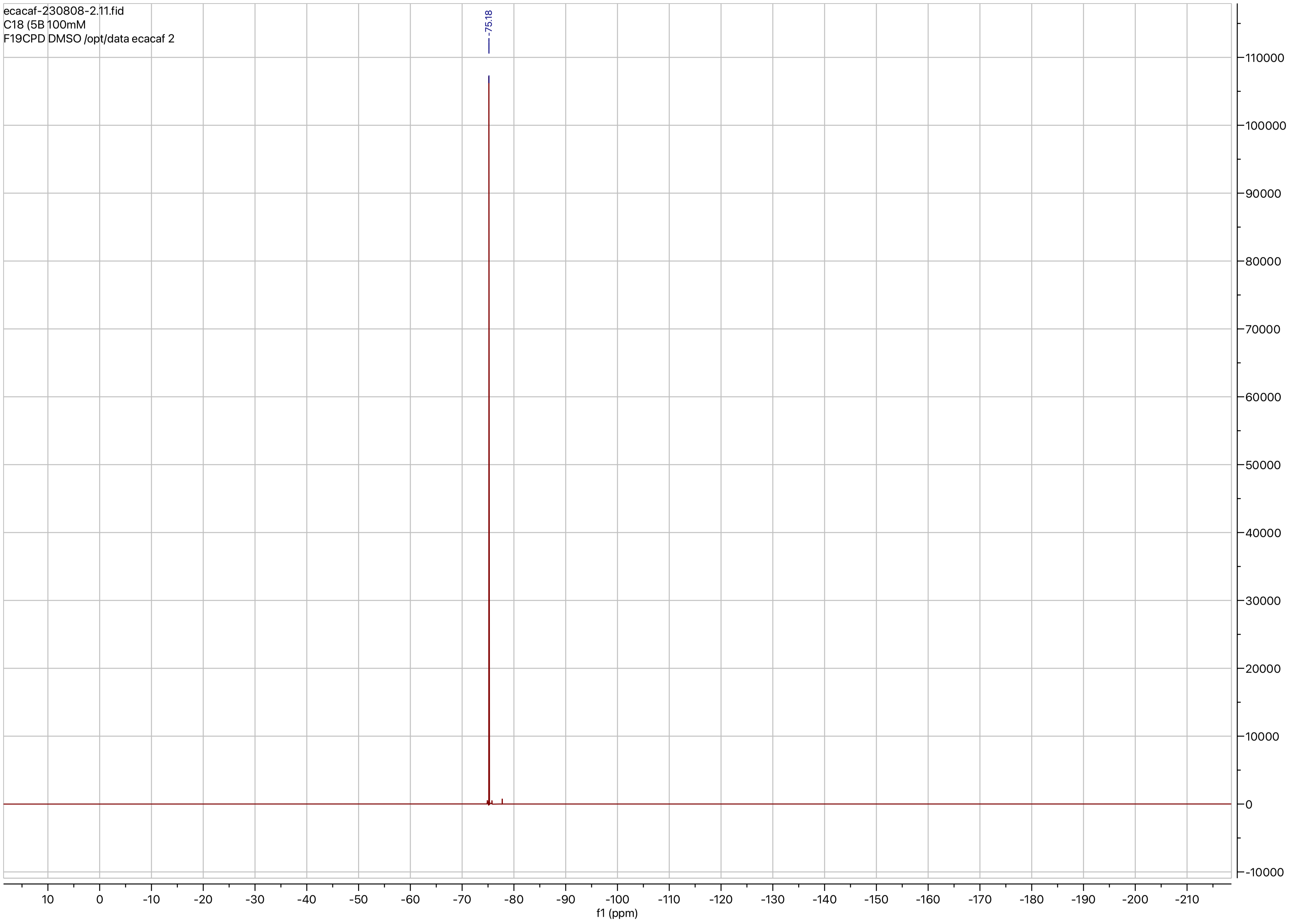
ecacaf-230808-2.11.fid
C18 (5B 100mM
F19CPD DMSO /opt/data ecacaf 2
-75.18
f1 (ppm)

**Compound ID: C-19**

EW12557-58-P1B MeOD Bruker_C_300MHz

Supervisor: Tao Guo

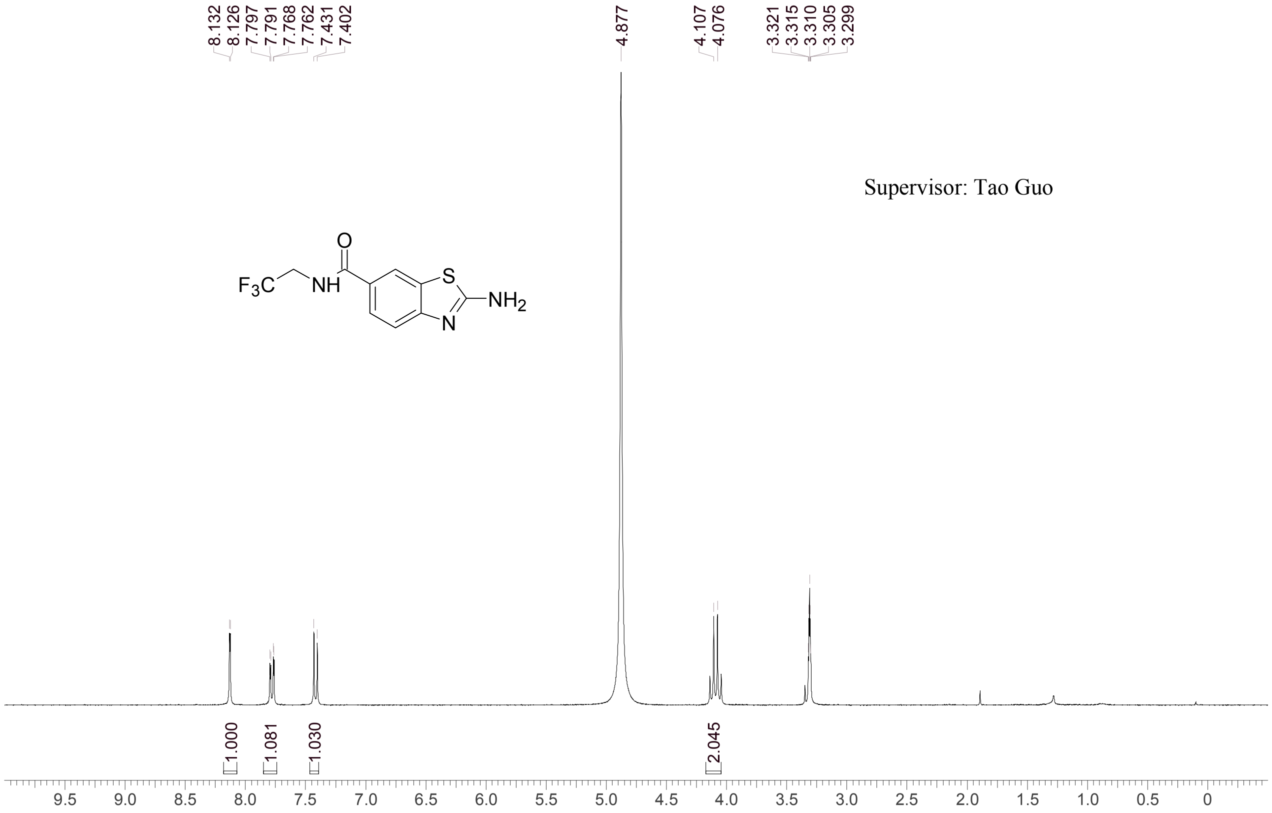


| | |
|---|---|
| Acquisition Time (sec) | 4.0993 |
| Comment | EW12557-58-P1B MeOD Bruker_C_300MHz |
| Date | 25 Jul 2018 13:35:36 |
| Frequency (MHz) | 300.1300 |
| Nucleus | 1H |
| Number of Transients | 8 |
| Origin | spect |
| Original Points Count | 24576 |
| Owner | nmrsu |
| Points Count | 65536 |
| Pulse Sequence | zg30 |
| Receiver Gain | 256.00 |
| SW(cyclical) (Hz) | 5995.20 |
| Solvent | METHANOL-d4 |
| Spectrum Offset (Hz) | 1796.1171 |
| Spectrum Type | standard |
| Sweep Width (Hz) | 5995.11 |
| Temperature (degree C) | 22.860 |

Operator: Date:

Confidential, for research only not for regulatory filing

LCMS REPORT

Compound ID : C-19 ->
Sample ID : EW12557-58-P1P
Injection Date : 24-Jul-2018 13:28:46
Location : P2-A-02
Injection volume : 2.000
Acq Method : D:\DATA\1807\180724 22\10-80CD_2MIN_220&25
Data Filename : D:\DATA\1807\180724 22\EW12557-58-P1P.D
Instrument : LCMS-C

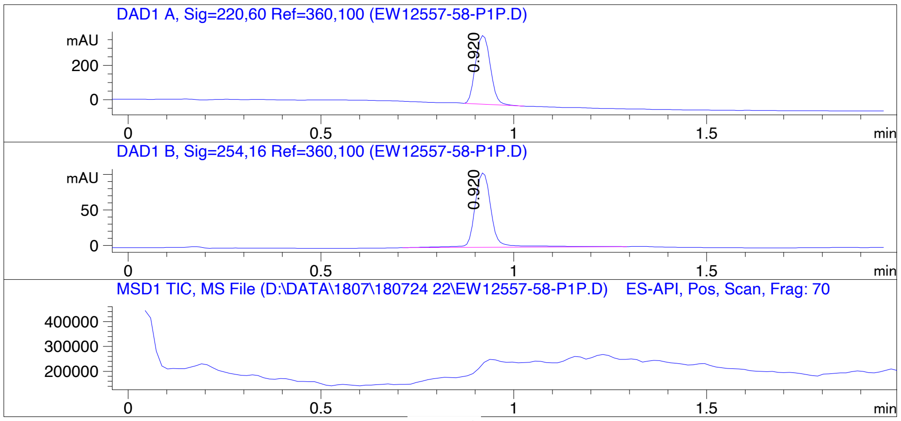


Report

Signal 1 : DAD1 A, Sig=220,60 Ref=360,100

| # | Meas. Ret. | Height | Width | Area | Area % |
|---|---|---|---|---|---|
| 1 | 0.920 | 401.501 | 0.045 | 1069.022 | 100.000 |

Signal 2 : DAD1 B, Sig=254,16 Ref=360,100

| # | Meas. Ret. | Height | Width | Area | Area % |
|---|---|---|---|---|---|
| 1 | 0.920 | 104.884 | 0.049 | 313.791 | 100.000 |

Operator:__________________

Date:___________________

Confidential, for research only not for regulatory filing

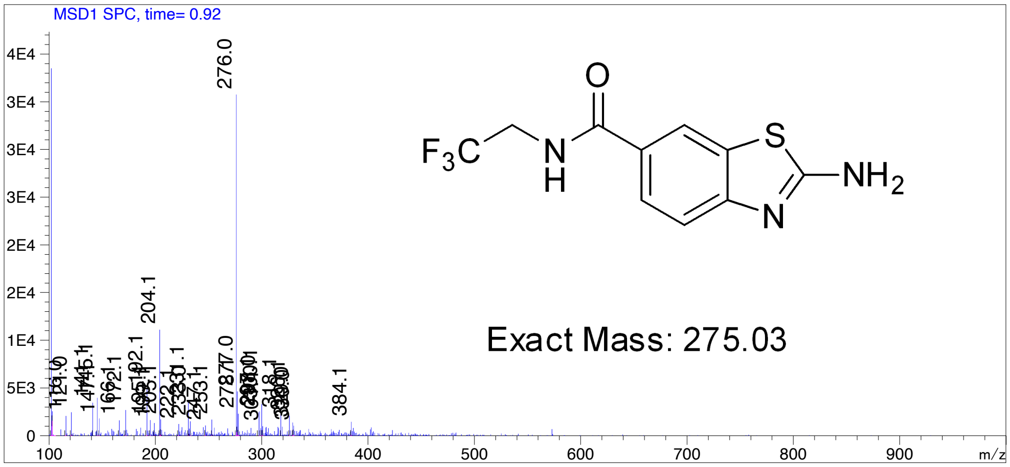

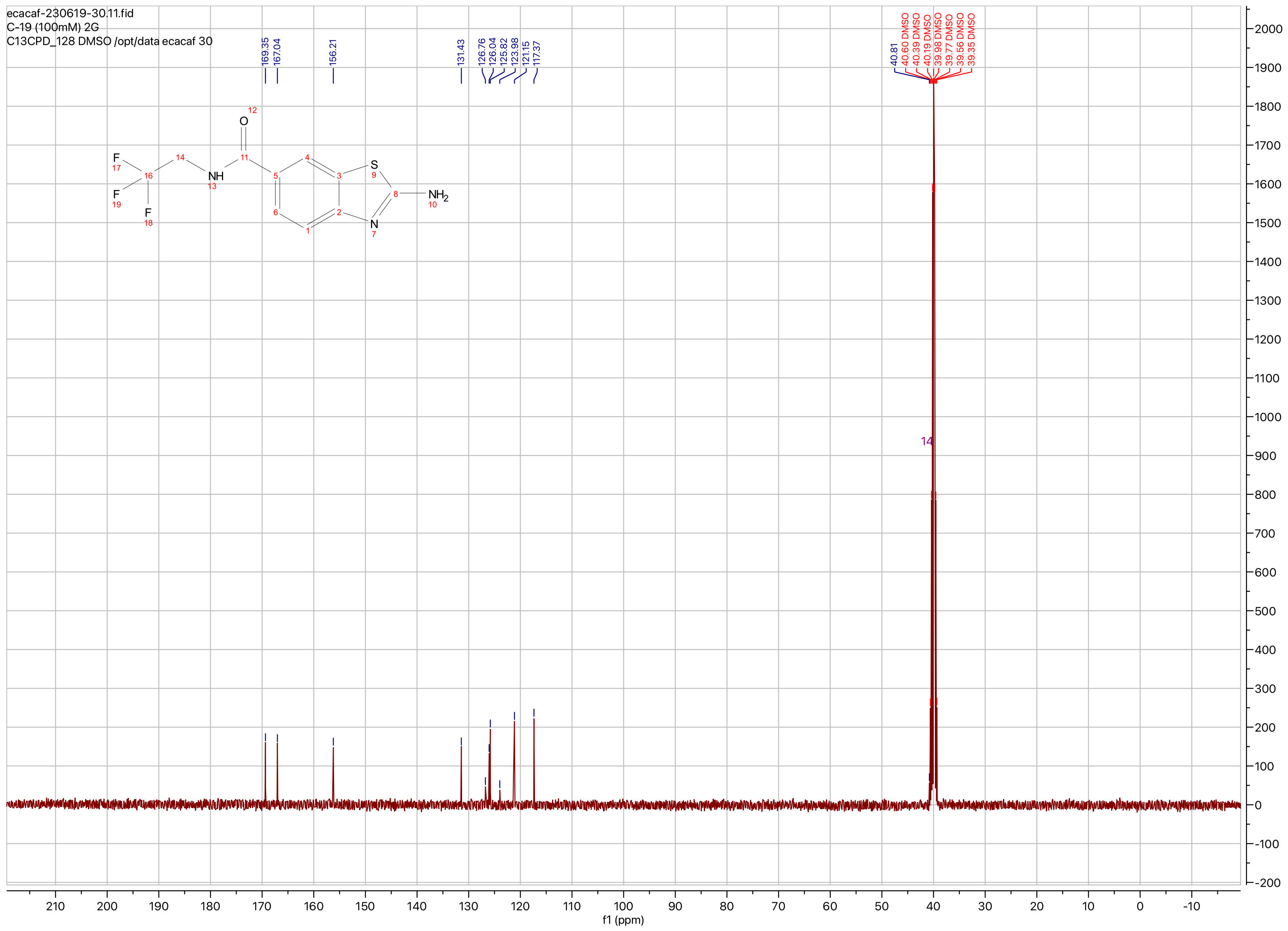

ecacaf-230619-30.11.fid
C-19 (100mM) 2G
C13CPD_128 DMSO /opt/data ecacaf 30
169.35
167.04
156.21
131.43
126.76
126.04
125.82
123.98
121.15
117.37
40.81
40.60 DMSO
40.39 DMSO
40.19 DMSO
39.98 DMSO
39.77 DMSO
39.56 DMSO
39.35 DMSO
14
f1 (ppm)

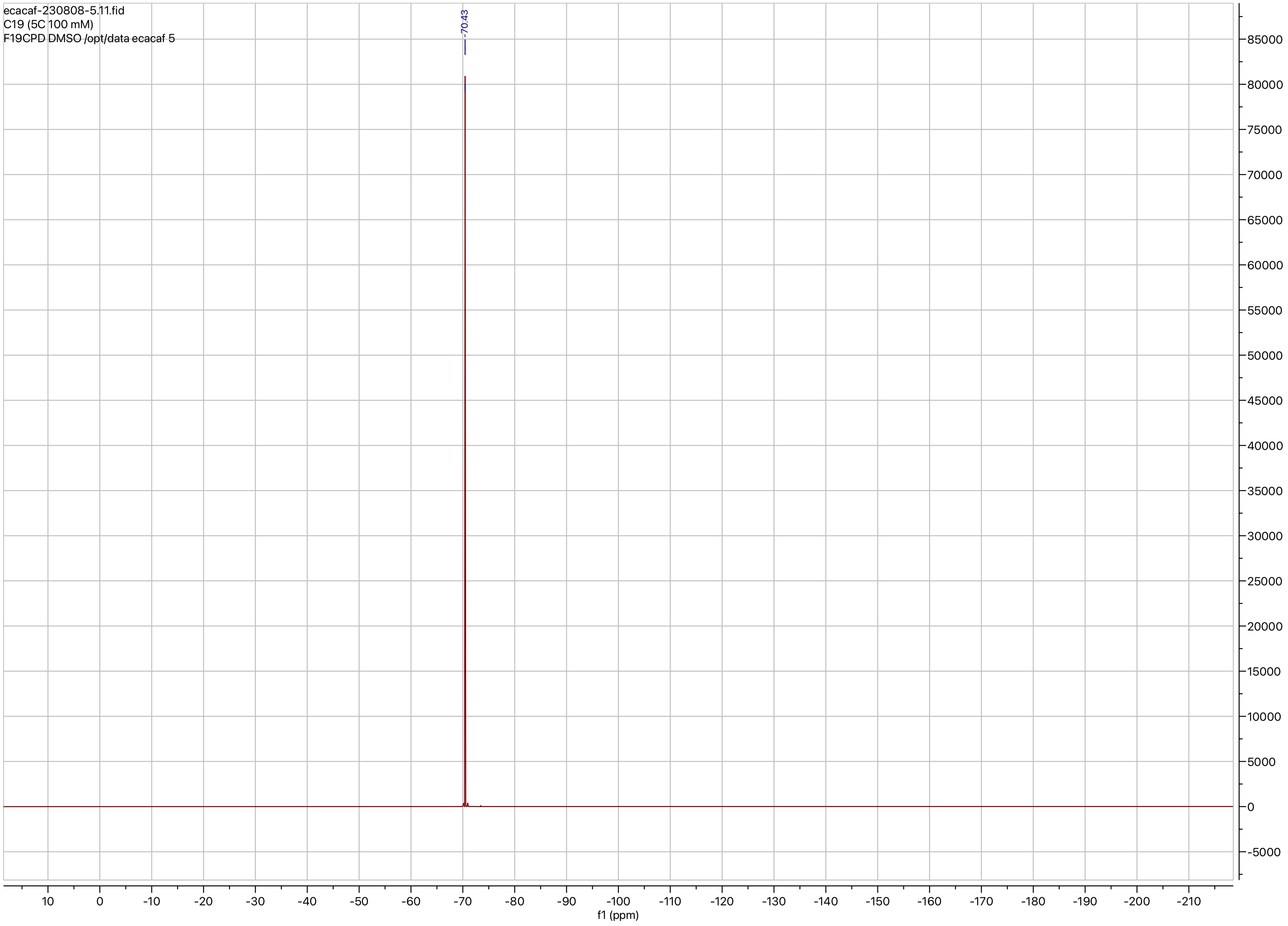
ecacaf-230808-5.11.fid
C19 (5C 100 mM)
F19CPD DMSO /opt/data ecacaf 5
-70.43
10
0
-10
-20
-30
-40
-50
-60
-70
-80
-90
-100
-110
-120
-130
-140
-150
-160
-170
-180
-190
-200
-210
f1 (ppm)
85000
80000
75000
70000
65000
60000
55000
50000
45000
40000
35000
30000
25000
20000
15000
10000
5000
0
-5000

**Compound ID: C-20**

EW8457-542-P1A DMSO Bruker_B_400MHz

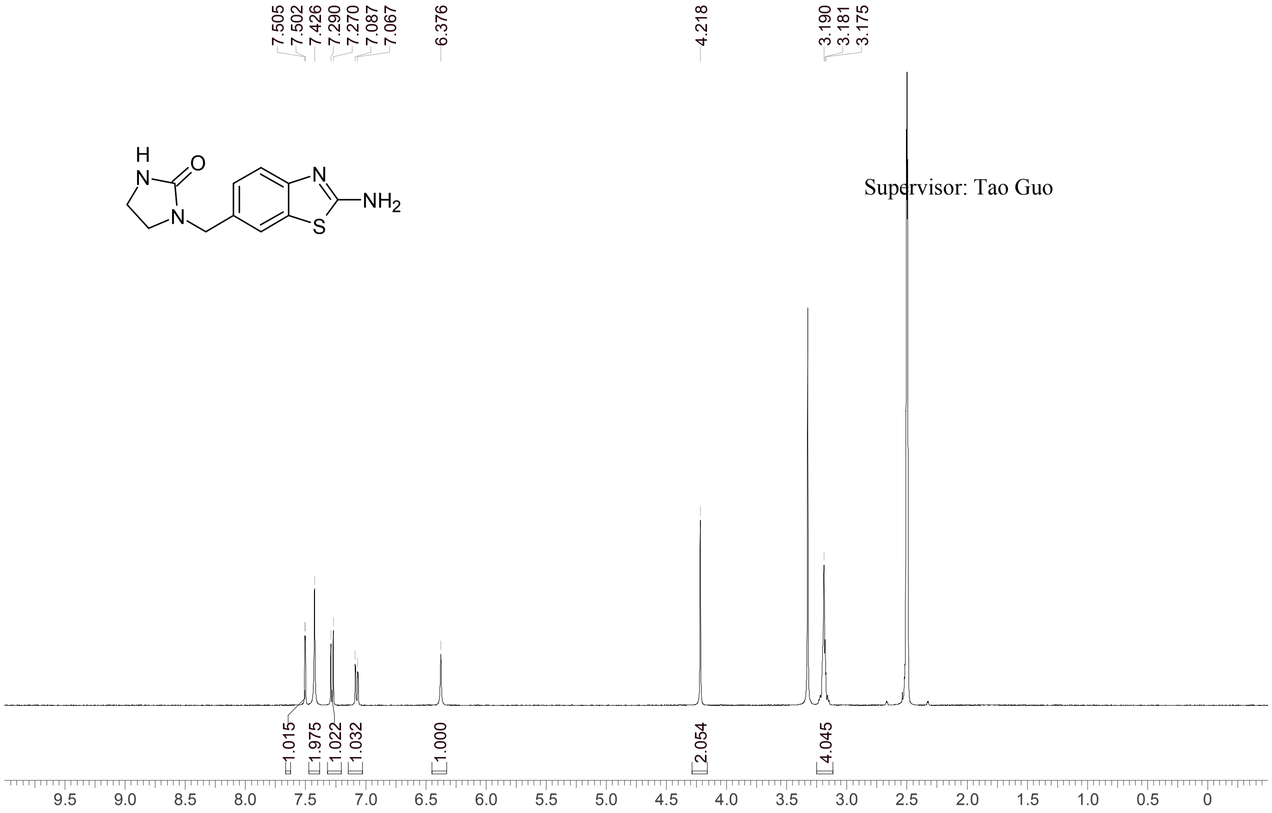


Supervisor: Tao Guo

| | |
|---|---|
| Acquisition Time (sec) | 3.0671 |
| Comment | EW8457-542-P1A DMSO Bruker_B_400MHz |
| Date | 09 Aug 2018 08:24:20 |
| Frequency (MHz) | 400.1300 |
| Nucleus | 1H |
| Number of Transients | 8 |
| Origin | spect |
| Original Points Count | 24576 |
| Owner | nmrsu |
| Points Count | 65536 |
| Pulse Sequence | zg30 |
| Receiver Gain | 190.49 |
| SW(cyclical) (Hz) | 8012.82 |
| Solvent | DMSO-d6 |
| Spectrum Offset (Hz) | 2397.1384 |
| Spectrum Type | standard |
| Sweep Width (Hz) | 8012.70 |
| Temperature (degree C) | 25.145 |

Operator: Date:

Confidential, for research only not for regulatory filing

Confidential, for research only not for regulatory filing

### LCMS REPORT

Compound ID : C-20
Sample ID : EW8457-542-P1D
Injection Vol : 2ul
Location : vial77
Acq Method : d:\method\5-95CD_R_220&254_POS_50.lcm
Org DataFile : D:\DATA\1808\180809\EW8457-542-P1D.lcd
Injection Date : 2018-08-09 11:09:10
Instrument :LCMS-O 15-105

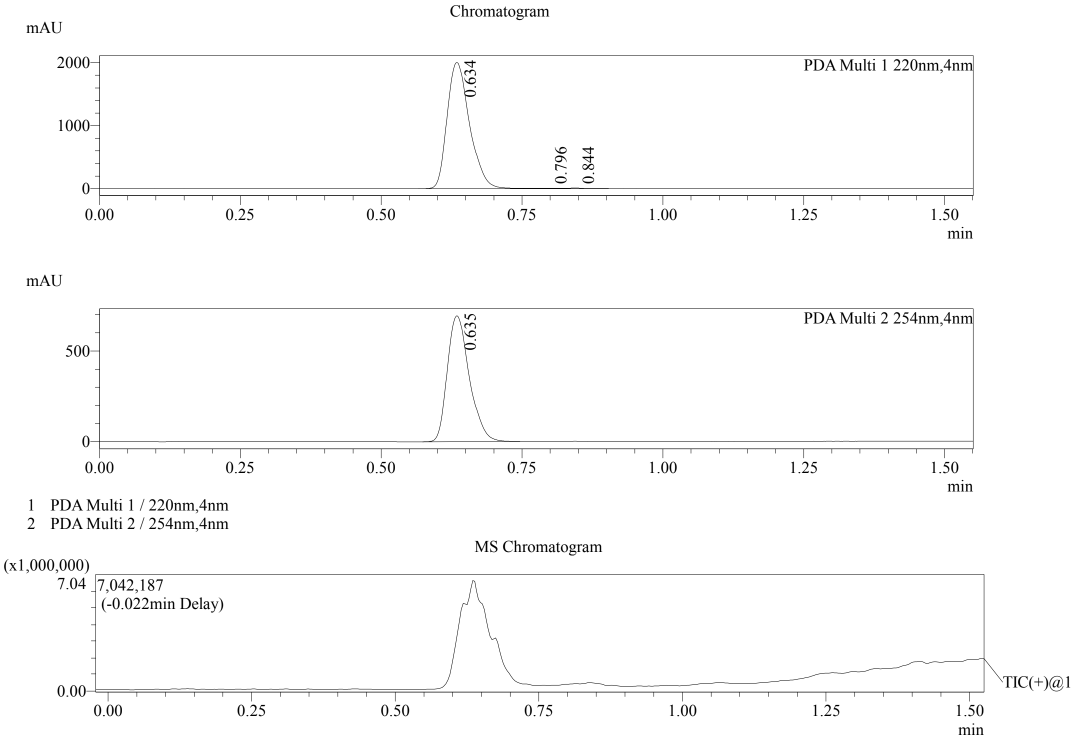


Integration Result

Peak Table

PDA Ch1 220nm

| Peak# | Ret. Time | Height | Height% | USP Width | Area | Area% |
|---|---|---|---|---|---|---|
| 1 | 0.634 | 2001501 | 99.145 | 0.071 | 5554850 | 99.250 |
| 2 | 0.796 | 6699 | 0.332 | 0.108 | 24956 | 0.446 |
| 3 | 0.844 | 10559 | 0.523 | 0.044 | 17012 | 0.304 |

Peak Table

PDA Ch2 254nm

| Peak# | Ret. Time | Height | Height% | USP Width | Area | Area% |
|---|---|---|---|---|---|---|
| 1 | 0.635 | 693594 | 100.000 | 0.070 | 1885450 | 100.000 |

Mass Spectrum

RetTime:0.635 Datafile: D:\DATA\1808\180809\EW8457-542-P1D.lcd

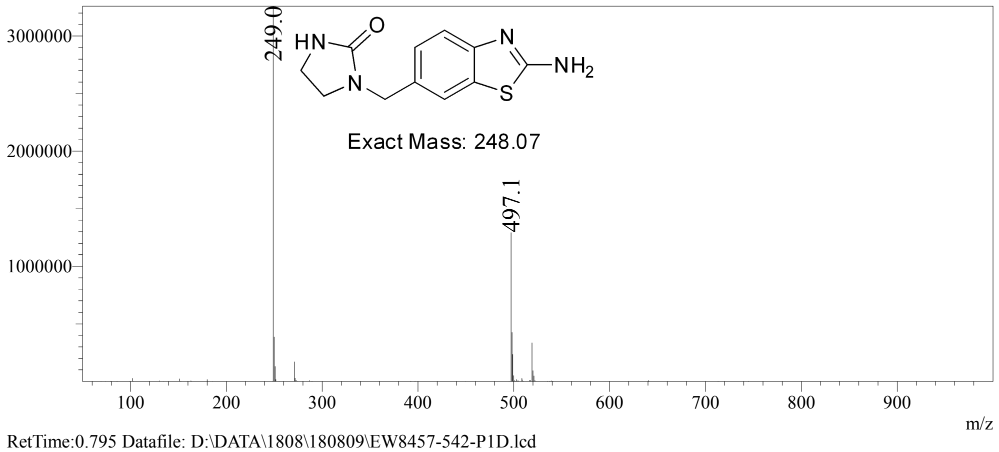


RetTime:0.795 Datafile: D:\DATA\1808\180809\EW8457-542-P1D.lcd

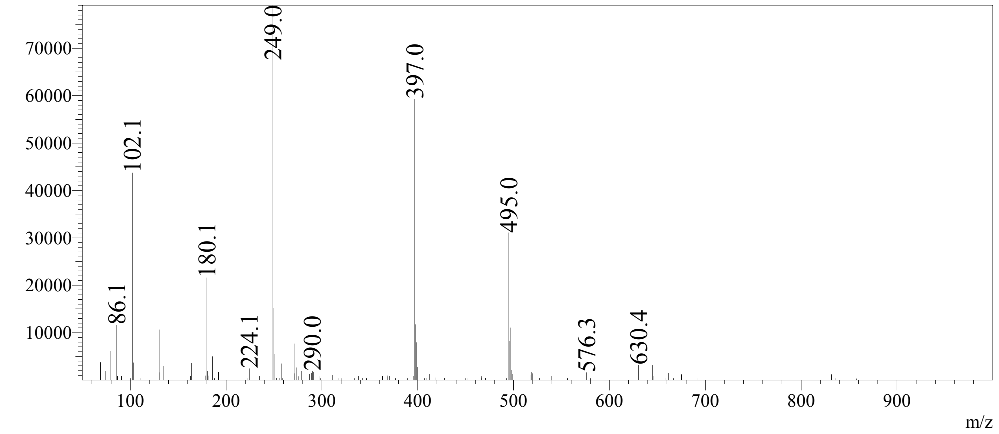


RetTime:0.845 Datafile: D:\DATA\1808\180809\EW8457-542-P1D.lcd

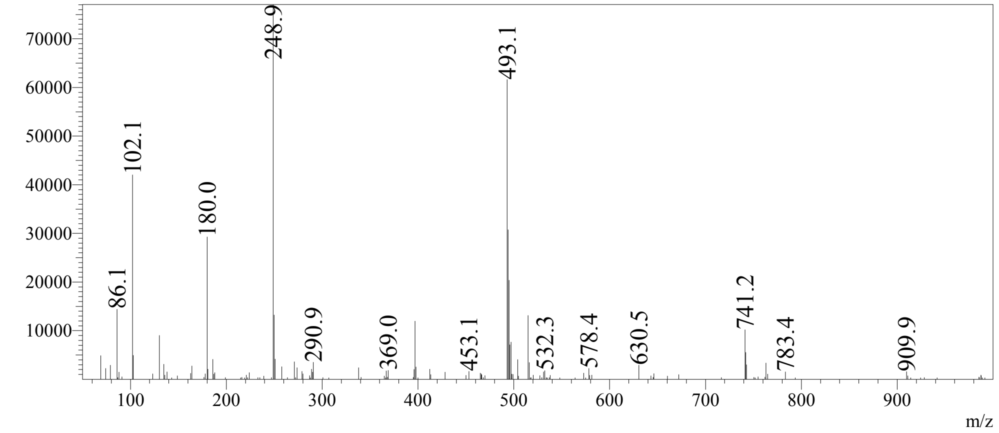

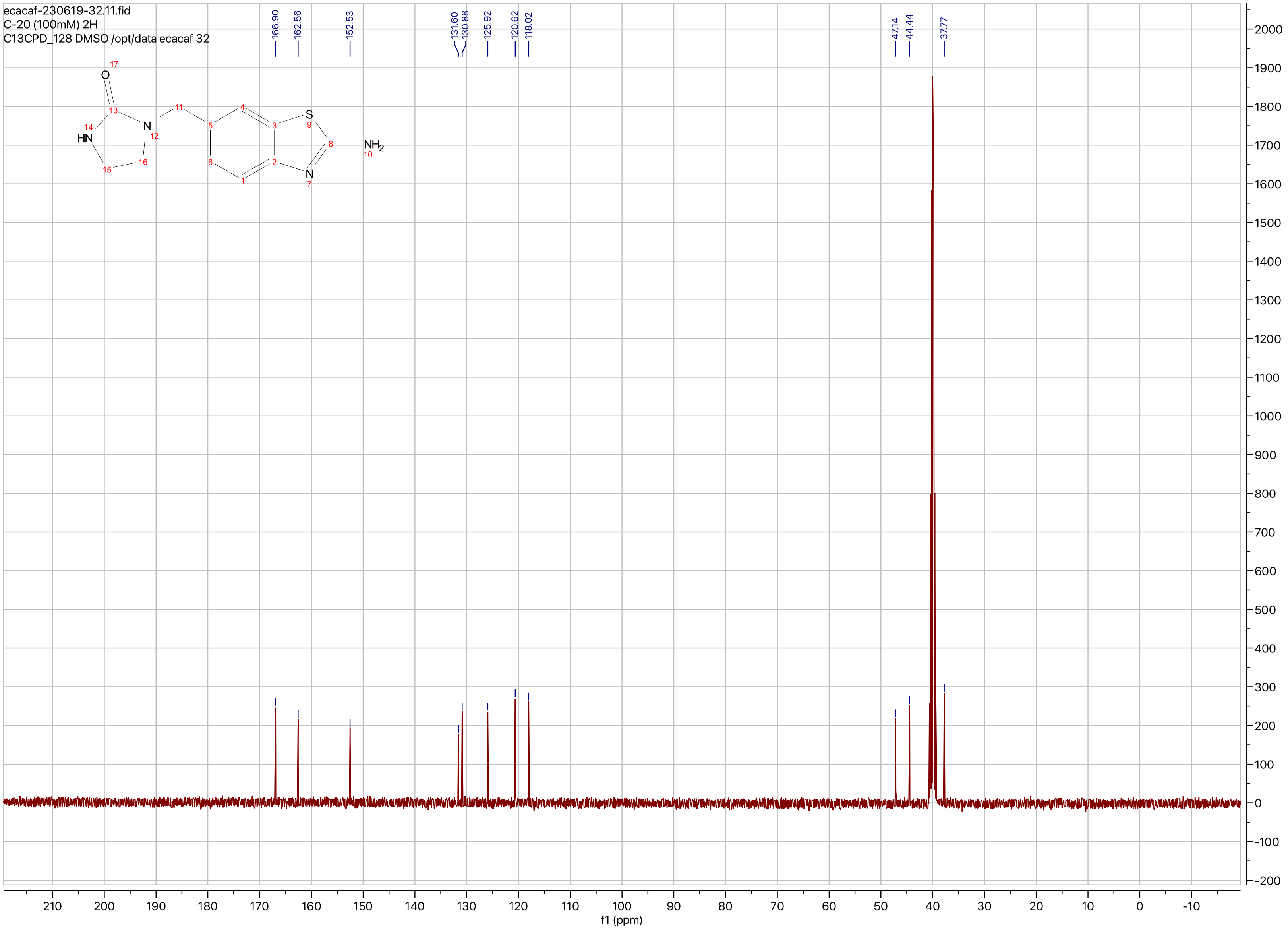
ecacaf-230619-32.11.fid
C-20 (100mM) 2H
C13CPD_128 DMSO /opt/data ecacaf 32
166.90
162.56
152.53
131.60
130.88
125.92
120.62
118.02
47.14
44.44
37.77
O
HN
N
S
N
$NH_2$
f1 (ppm)

**Compound ID: C-21**

EW8397-344-P1A DMSO Bruker_B_400MHz

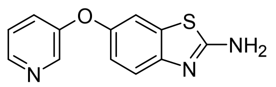


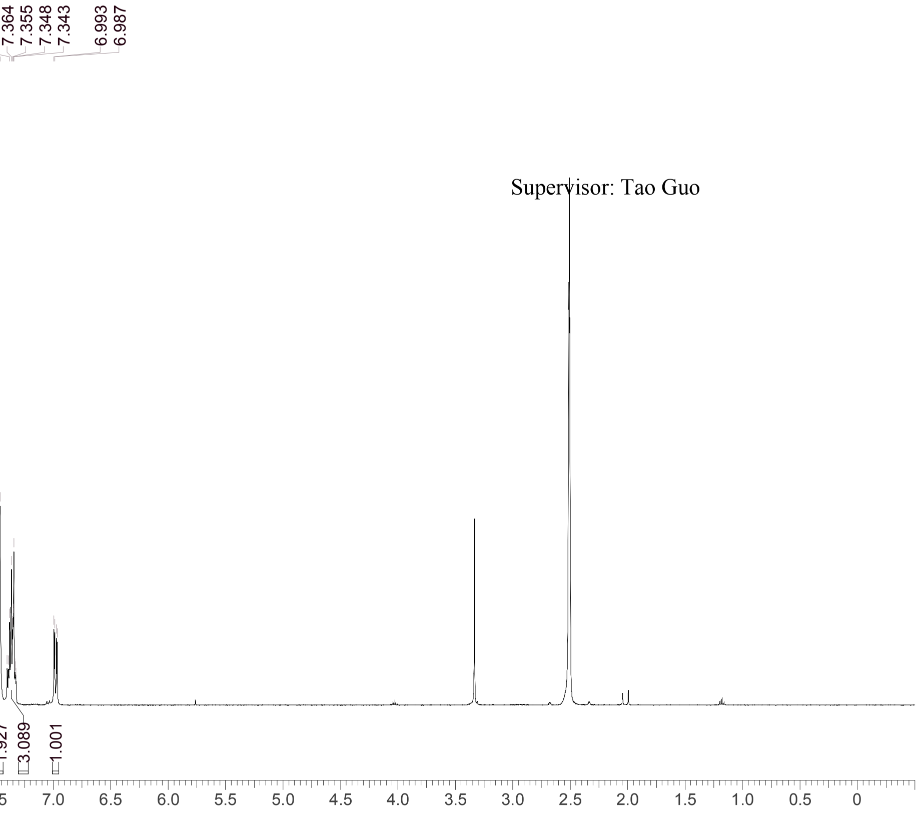


Supervisor: Tao Guo

| | |
|---|---|
| Acquisition Time (sec) | 3.0671 |
| Comment | EW8397-344-P1A DMSO Bruker_B_400MHz |
| Date | 25 May 2018 13:32:32 |
| Frequency (MHz) | 400.1300 |
| Nucleus | 1H |
| Number of Transients | 8 |
| Origin | spect |
| Original Points Count | 24576 |
| Owner | nmrsu |
| Points Count | 65536 |
| Pulse Sequence | zg30 |
| Receiver Gain | 190.49 |
| SW(cyclical) (Hz) | 8012.82 |
| Solvent | DMSO-d6 |
| Spectrum Offset (Hz) | 2400.7795 |
| Spectrum Type | standard |
| Sweep Width (Hz) | 8012.70 |
| Temperature (degree C) | 24.412 |

Operator: Date:

Confidential, for research only not for regulatory filing

Confidential, for research only not for regulatory filing

### LCMS REPORT

Compound ID : C-21
Sample ID : EW8397-344-P1A1
Injection Vol : 0.4ul
Location : vial33
Acq Method : d:\method\5-95CD_R_220&254_POS.lcm
Org DataFile : D:\DATA\1805\180529\EW8397-344-P1A1.lcd
Injection Date : 29/05/2018 09:38:47
Instrument :LCMS-K 15-105

Chromatogram

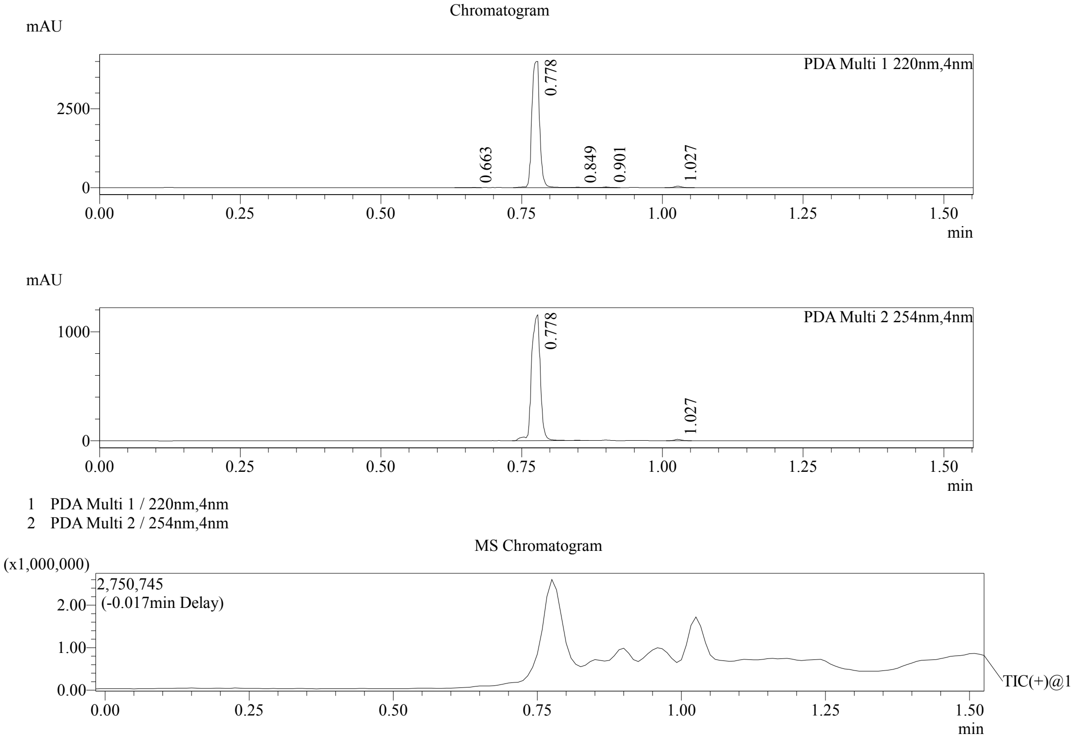


Integration Result

Peak Table

PDA Ch1 220nm

| Peak# | Ret. Time | Height | Height% | USP Width | Area | Area% |
|---|---|---|---|---|---|---|
| 1 | 0.663 | 2726 | 0.067 | 0.083 | 5234 | 0.132 |
| 2 | 0.778 | 3998437 | 98.205 | 0.023 | 3890497 | 98.407 |
| 3 | 0.849 | 7897 | 0.194 | 0.022 | 6262 | 0.158 |
| 4 | 0.901 | 16732 | 0.411 | 0.023 | 14299 | 0.362 |
| 5 | 1.027 | 45724 | 1.123 | 0.025 | 37185 | 0.941 |

Peak Table

PDA Ch2 254nm

| Peak# | Ret. Time | Height | Height% | USP Width | Area | Area% |
|---|---|---|---|---|---|---|
| 1 | 0.778 | 1153737 | 98.987 | 0.028 | 1252003 | 99.237 |
| 2 | 1.027 | 11807 | 1.013 | 0.024 | 9630 | 0.763 |

Confidential, for research only not for regulatory filing

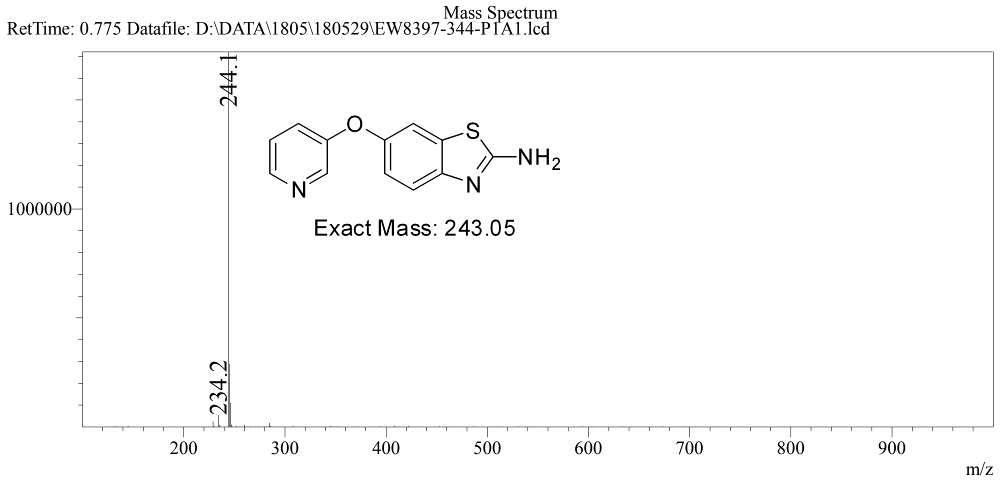

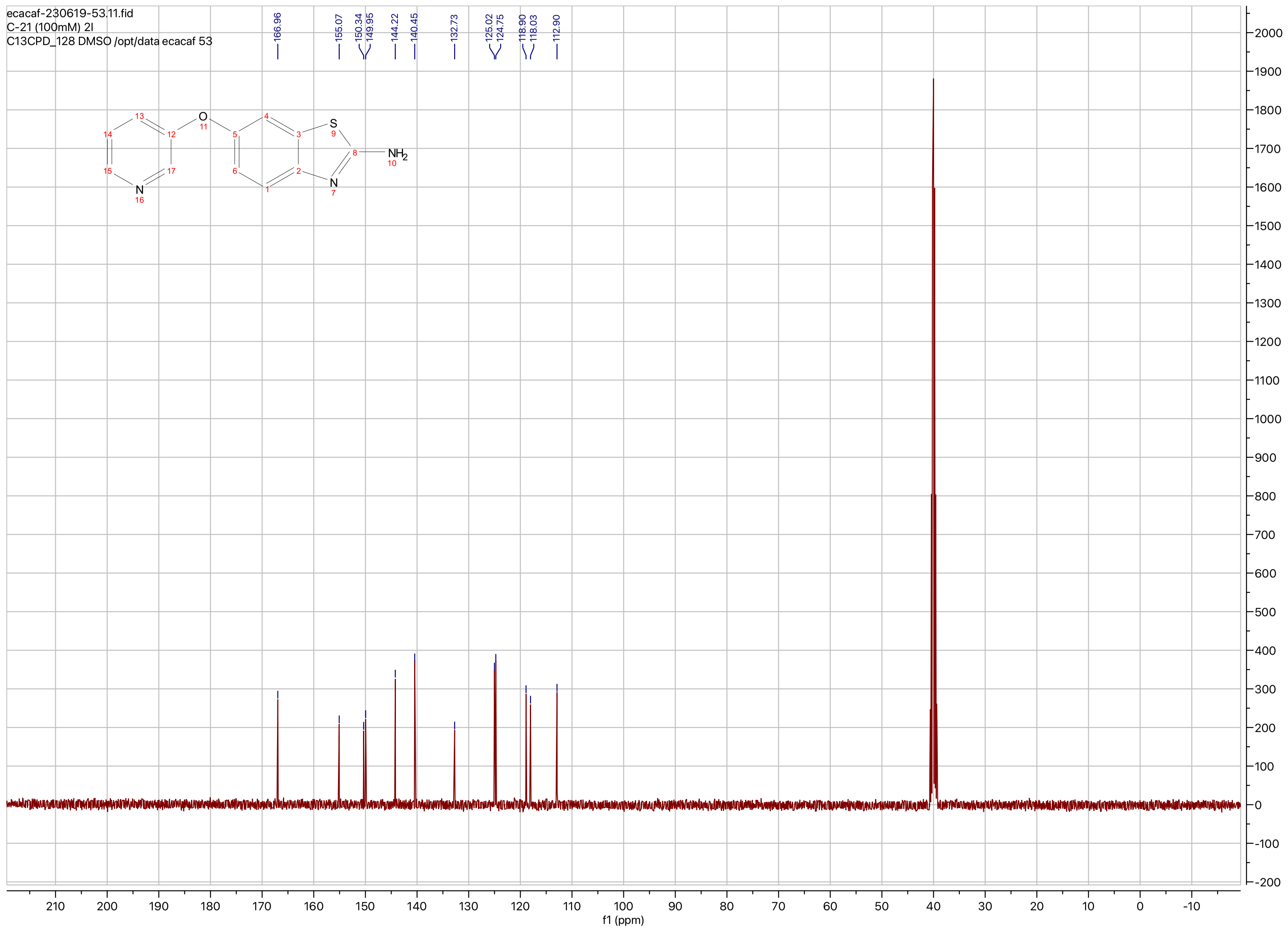

ecacaf-230619-53.11.fid
C-21 (100mM) 2l
C13CPD_128 DMSO /opt/data ecacaf 53
166.96
155.07
150.34
149.95
144.22
140.45
132.73
125.02
124.75
118.90
118.03
112.90
O
S
N
NH2
N
f1 (ppm)

**Compound ID: C-22**

EW8457-427-P1A DMSO Bruker_A_400MHz

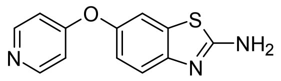

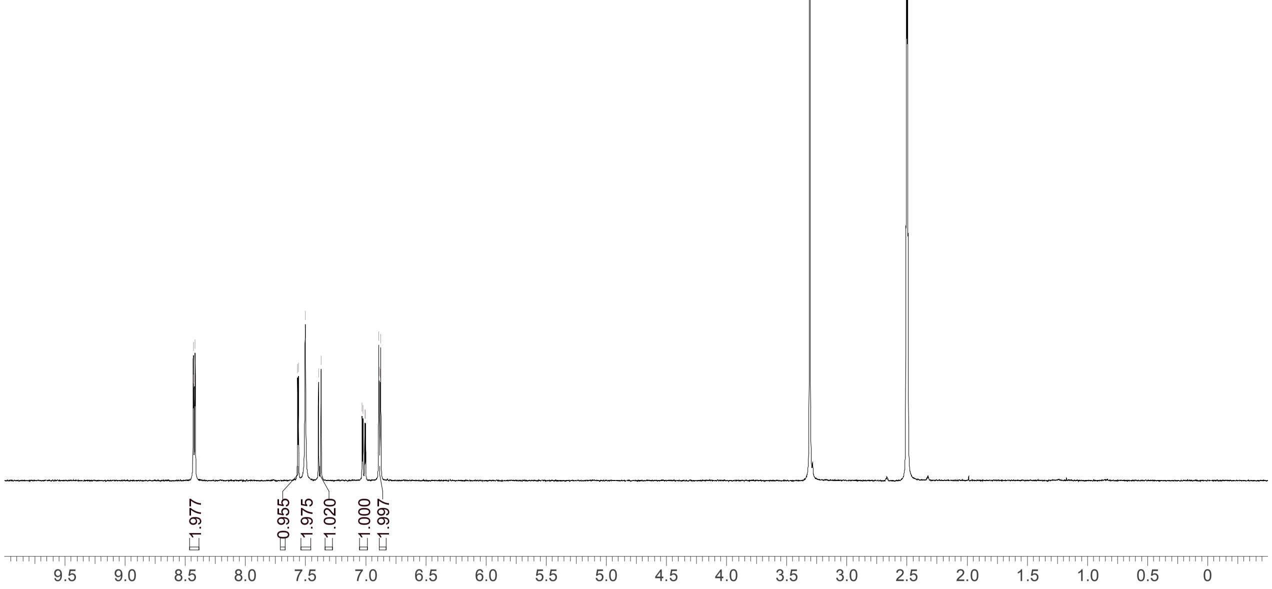


Supervisor: Tao Guo

| | |
|---|---|
| Acquisition Time (sec) | 3.0671 |
| Comment | EW8457-427-P1A DMSO Bruker_A_400MHz |
| Date | 03 Jun 2018 09:33:59 |
| Frequency (MHz) | 400.1300 |
| Nucleus | 1H |
| Number of Transients | 8 |
| Origin | spect |
| Original Points Count | 24576 |
| Owner | nmrsu |
| Points Count | 65536 |
| Pulse Sequence | zg30 |
| Receiver Gain | 175.36 |
| SW(cyclical) (Hz) | 8012.82 |
| Solvent | DMSO-d6 |
| Spectrum Offset (Hz) | 2397.5386 |
| Spectrum Type | standard |
| Sweep Width (Hz) | 8012.70 |
| Temperature (degree C) | 27.149 |

Operator:

Date:

Confidential, for research only not for regulatory filing

Confidential, for research only not for regulatory filing

### LCMS REPORT

Compound ID : C-22
Sample ID : EW8457-427-P1C
Injection Vol : 1ul
Location : vial11
Acq Method : d:\method\5-95CD_R_210&254_POS.lcm
Org DataFile : D:\DATA\1806\180603\EW8457-427-P1C.lcd
Injection Date : 03/06/2018 09:36:38
Instrument :LCMS-K 15-105

Chromatogram

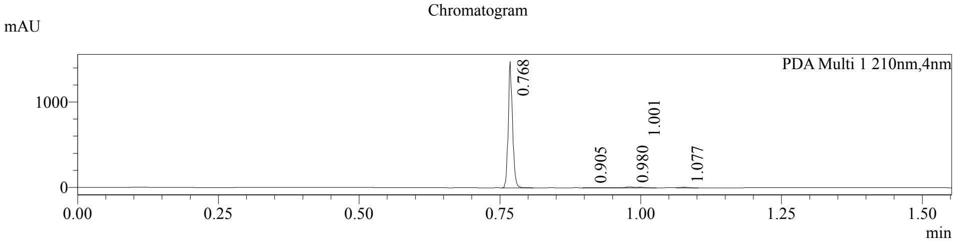


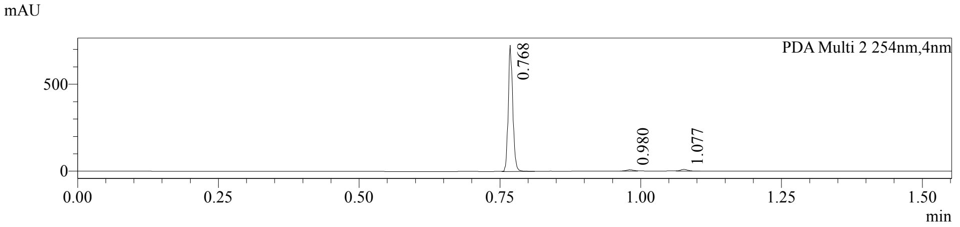


1 PDA Multi 1 / 210nm,4nm
2 PDA Multi 2 / 254nm,4nm

MS Chromatogram

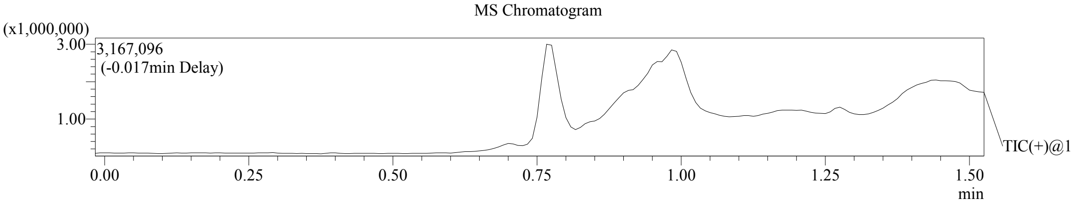


Integration Result

Peak Table

PDA Ch1 210nm

| Peak# | Ret. Time | Height | Height% | USP Width | Area | Area% |
|---|---|---|---|---|---|---|
| 1 | 0.768 | 1481776 | 98.107 | 0.016 | 784353 | 96.186 |
| 2 | 0.905 | 1675 | 0.111 | 0.428 | 6913 | 0.848 |
| 3 | 0.980 | 11739 | 0.777 | 0.028 | 10527 | 1.291 |
| 4 | 1.001 | 7543 | 0.499 | 0.032 | 6709 | 0.823 |
| 5 | 1.077 | 7632 | 0.505 | 0.027 | 6952 | 0.853 |

Peak Table

PDA Ch2 254nm

| Peak# | Ret. Time | Height | Height% | USP Width | Area | Area% |
|---|---|---|---|---|---|---|
| 1 | 0.768 | 725761 | 97.607 | 0.017 | 407572 | 96.243 |
| 2 | 0.980 | 7618 | 1.025 | 0.026 | 6624 | 1.564 |
| 3 | 1.077 | 10172 | 1.368 | 0.026 | 9288 | 2.193 |

Confidential, for research only not for regulatory filing

Mass Spectrum

RetTime: 0.767 Datafile: D:\DATA\1806\180603\EW8457-427-P1C.lcd

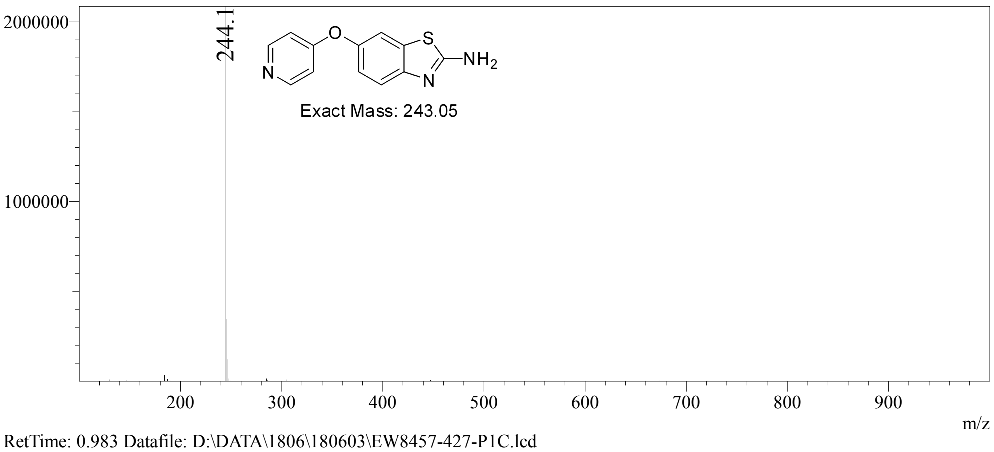


RetTime: 0.983 Datafile: D:\DATA\1806\180603\EW8457-427-P1C.lcd

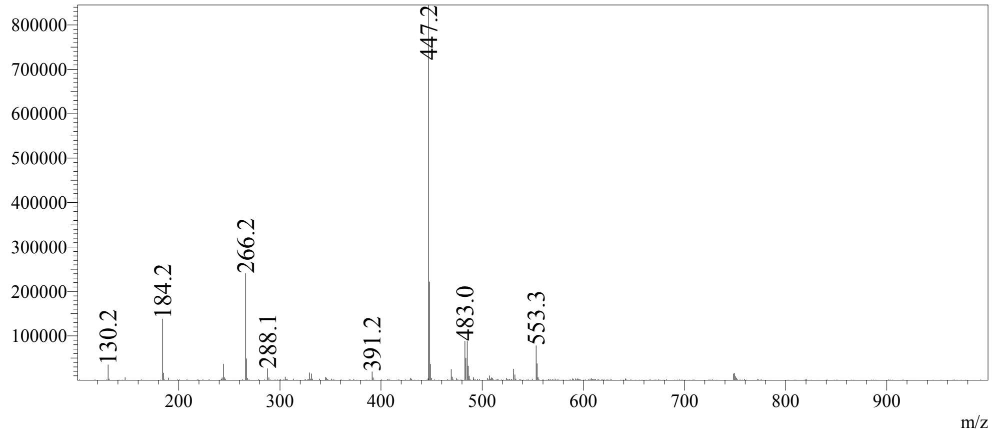

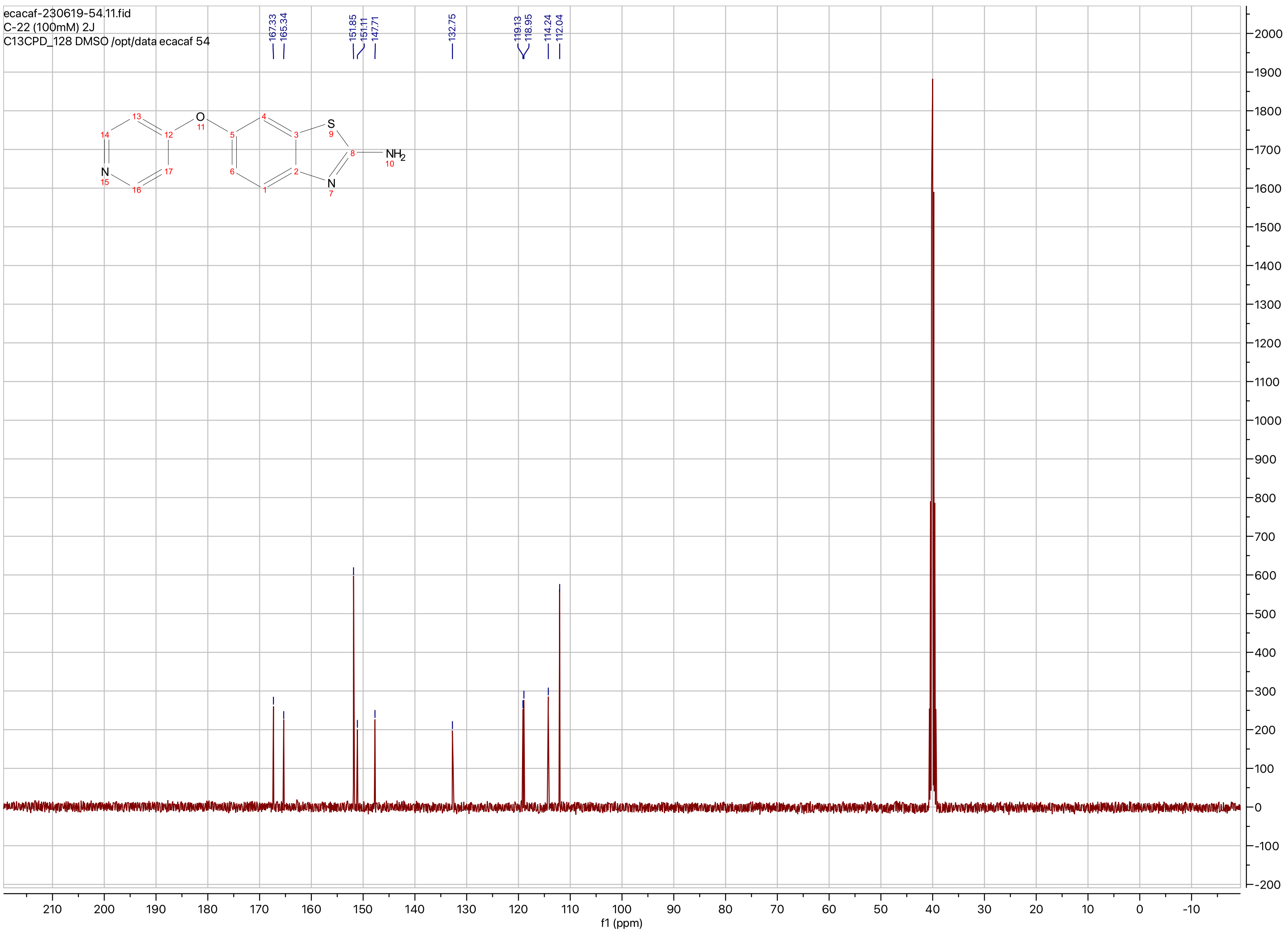

ecacaf-230619-54.11.fid
C-22 (100mM) 2J
C13CPD_128 DMSO /opt/data ecacaf 54
167.33
165.34
151.85
151.11
147.71
132.75
119.13
118.95
114.24
112.04
O
S
N
NH2
f1 (ppm)

**Compound ID: C-23**

EW8457-437-P1A DMSO Bruker_A_400MHz

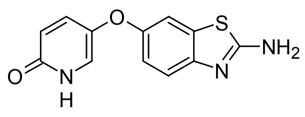

Supervisor: Tao Guo

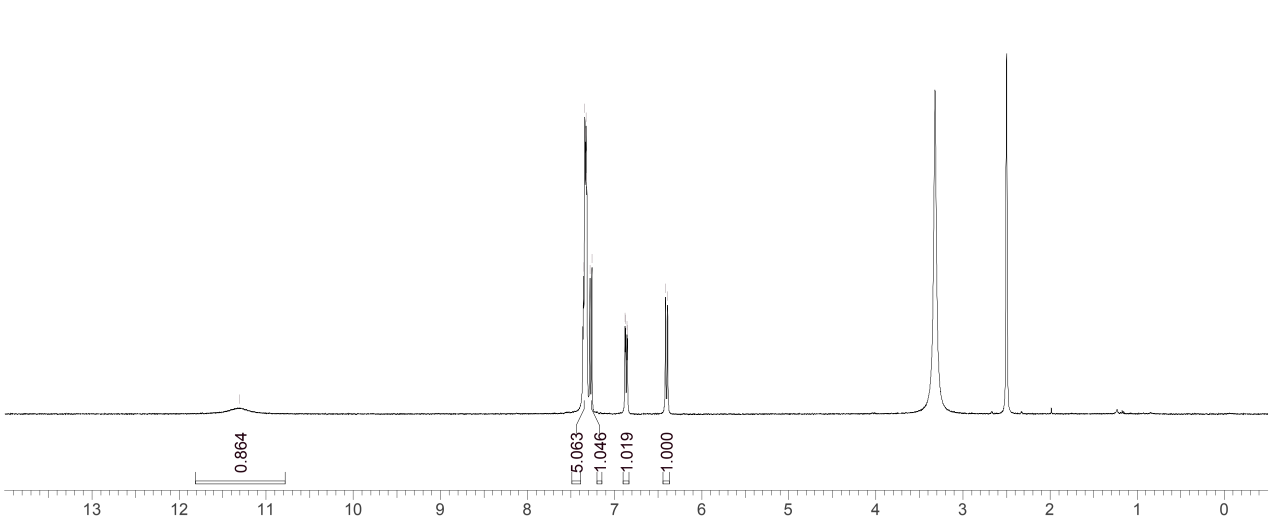


| | |
|---|---|
| Acquisition Time (sec) | 3.0671 |
| Comment | EW8457-437-P1A DMSO Bruker_A_400MHz |
| Date | 11 Jun 2018 08:32:09 |
| Frequency (MHz) | 400.1300 |
| Nucleus | 1H |
| Number of Transients | 8 |
| Origin | spect |
| Original Points Count | 24576 |
| Owner | nmrsu |
| Points Count | 65536 |
| Pulse Sequence | zg30 |
| Receiver Gain | 175.36 |
| SW(cyclical) (Hz) | 8012.82 |
| Solvent | DMSO-d6 |
| Spectrum Offset (Hz) | 2397.6584 |
| Spectrum Type | standard |
| Sweep Width (Hz) | 8012.70 |
| Temperature (degree C) | 27.152 |

Operator: Date:

Confidential, for research only not for regulatory filing

### LCMS REPORT

Compound ID : C-23
Sample ID : EW8457-437-P1B
Injection Vol : 2ul
Location : vial98
Acq Method : d:\method\5-95CD_R_220&254_POS.lcm
Org DataFile : D:\DATA\1806\180608\EW8457-437-P1B.lcd
Injection Date : 2018-06-08 16:22:05
Instrument :LCMS-O 15-105

Chromatogram

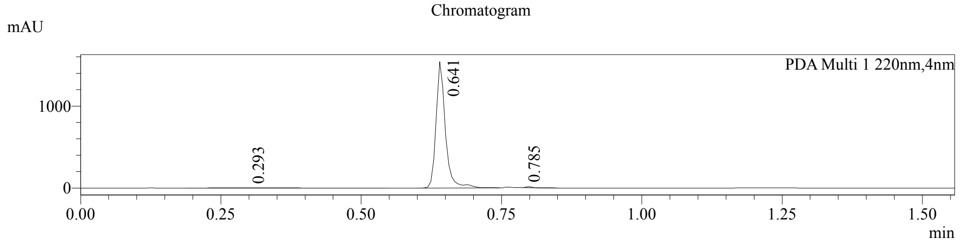


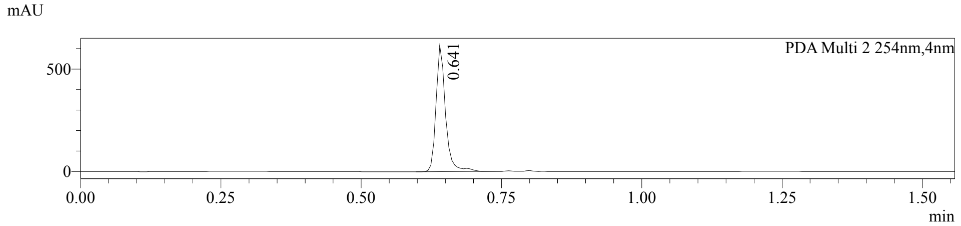


1 PDA Multi 1 / 220nm,4nm
2 PDA Multi 2 / 254nm,4nm

MS Chromatogram

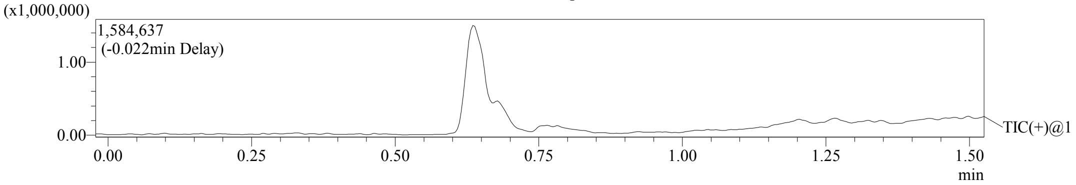


Integration Result

Peak Table

PDA Ch1 220nm

| Peak# | Ret. Time | Height | Height% | USP Width | Area | Area% |
|---|---|---|---|---|---|---|
| 1 | 0.293 | 3284 | 0.213 | 0.226 | 31395 | 1.716 |
| 2 | 0.641 | 1540638 | 99.686 | 0.034 | 1787380 | 97.695 |
| 3 | 0.785 | 1562 | 0.101 | 0.037 | 10785 | 0.589 |

Peak Table

PDA Ch2 254nm

| Peak# | Ret. Time | Height | Height% | USP Width | Area | Area% |
|---|---|---|---|---|---|---|
| 1 | 0.641 | 616918 | 100.000 | 0.034 | 730450 | 100.000 |

Confidential, for research only not for regulatory filing

Mass Spectrum

RetTime: 0.292 Datafile: D:\DATA\1806\180608\EW8457-437-P1B.lcd

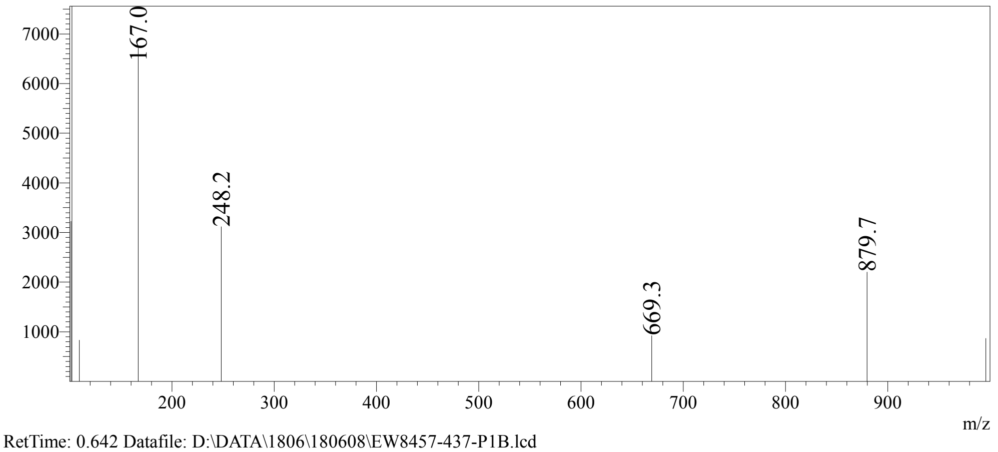


RetTime: 0.642 Datafile: D:\DATA\1806\180608\EW8457-437-P1B.lcd

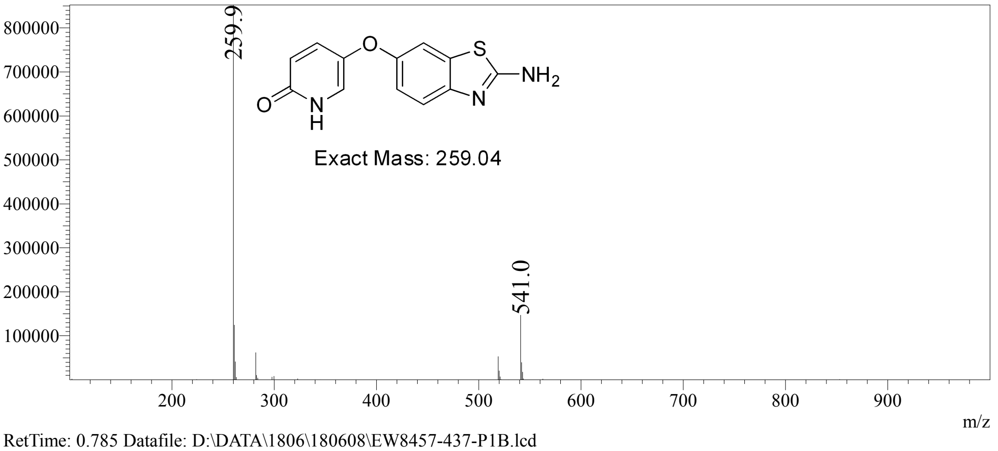


RetTime: 0.785 Datafile: D:\DATA\1806\180608\EW8457-437-P1B.lcd

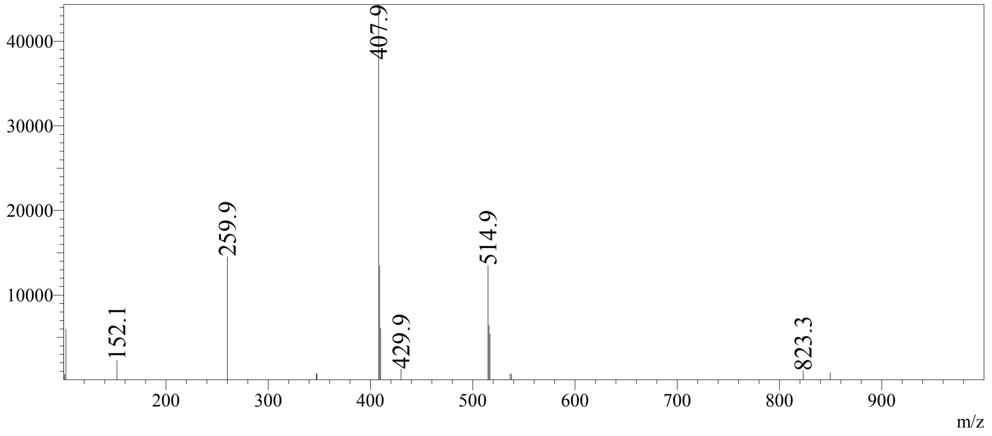

**Compound ID: C-24**

EW8457-440-P1A DMSO Bruker_B_400MHz

Supervisor: Tao Guo

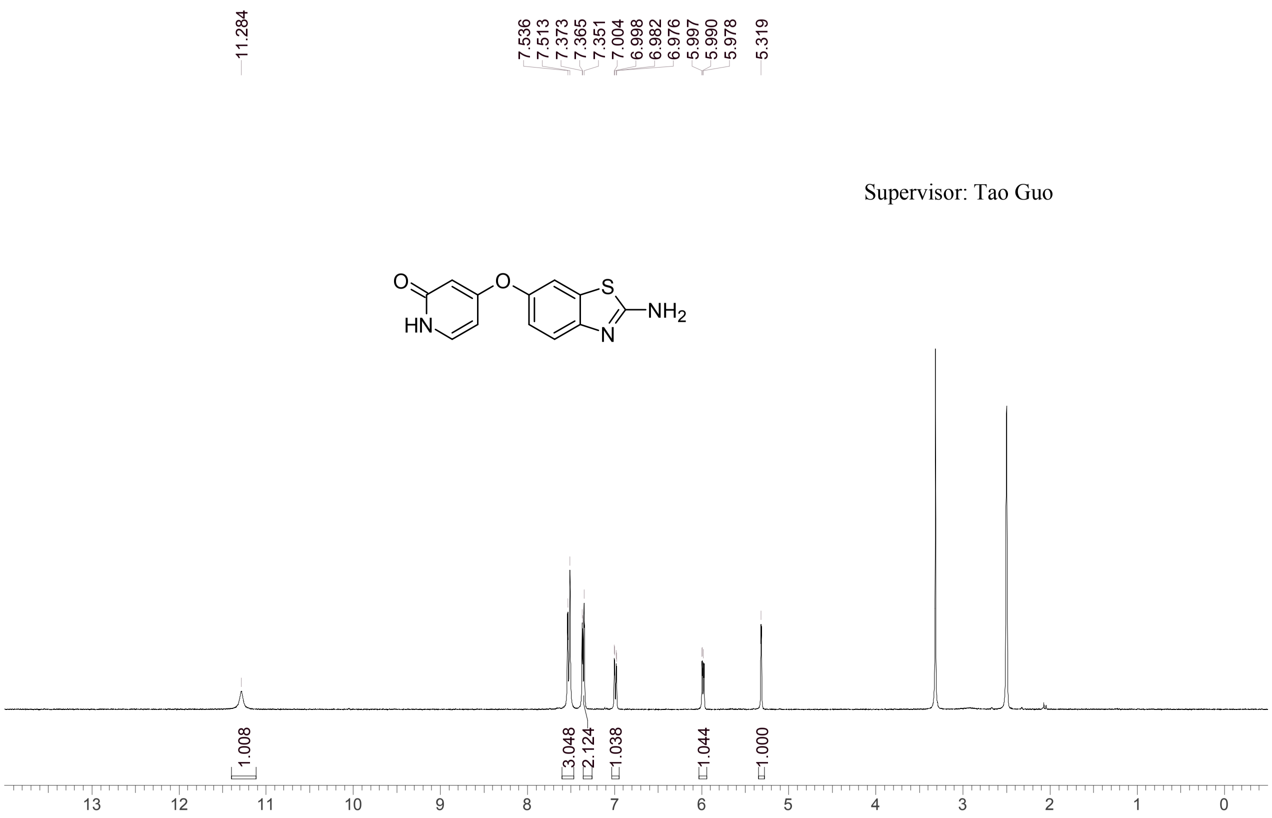


| | |
|---|---|
| Acquisition Time (sec) | 3.0671 |
| Comment | EW8457-440-P1A DMSO Bruker_B_400MHz |
| Date | 15 Jun 2018 08:31:09 |
| Frequency (MHz) | 400.1300 |
| Nucleus | 1H |
| Number of Transients | 8 |
| Origin | spect |
| Original Points Count | 24576 |
| Owner | nmrsu |
| Points Count | 65536 |
| Pulse Sequence | zg30 |
| Receiver Gain | 190.49 |
| SW(cyclical) (Hz) | 8012.82 |
| Solvent | DMSO-d6 |
| Spectrum Offset (Hz) | 2397.2583 |
| Spectrum Type | standard |
| Sweep Width (Hz) | 8012.70 |
| Temperature (degree C) | 22.902 |

Operator:

Date:

Confidential, for research only not for regulatory filing

Confidential, for research only not for regulatory filing

### LCMS REPORT

Compound ID : C-24
Sample ID : EW8457-440-P1B
Injection Vol : 3ul
Location : vial60
Acq Method : d:\method\5-95CD_R_220&254_POS_50.lcm
Org DataFile : D:\DATA\1806\180613\EW8457-440-P1B.lcd
Injection Date : 2018-06-13 14:20:32
Instrument :LCMS-O 15-105

Chromatogram

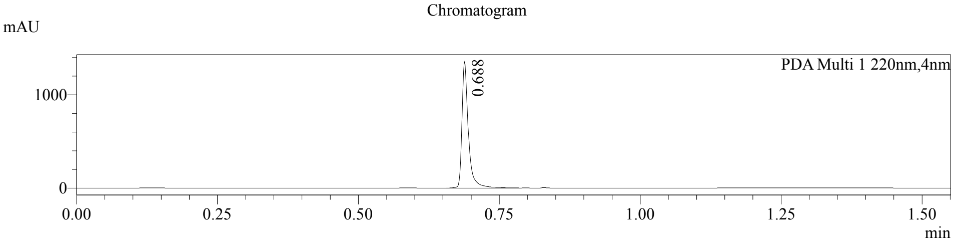


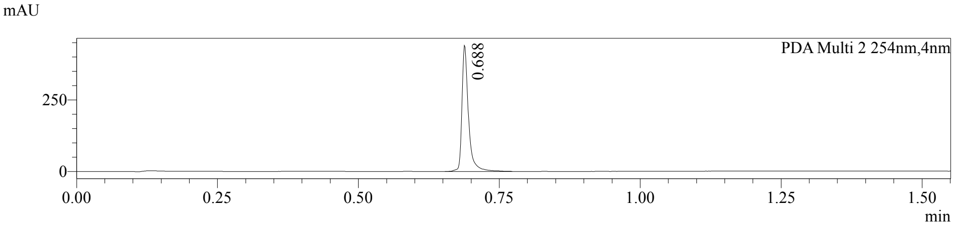


1 PDA Multi 1 / 220nm,4nm
2 PDA Multi 2 / 254nm,4nm

MS Chromatogram

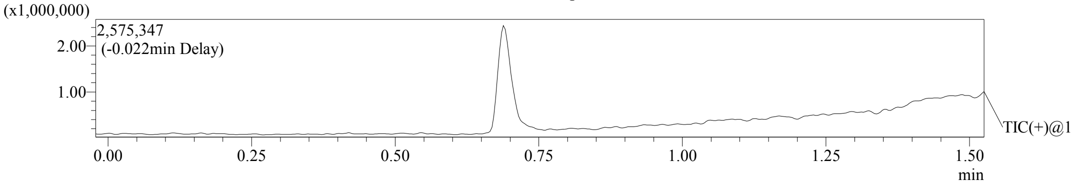


Integration Result

Peak Table

PDA Ch1 220nm

| Peak# | Ret. Time | Height | Height% | USP Width | Area | Area% |
|---|---|---|---|---|---|---|
| 1 | 0.688 | 1354261 | 100.000 | 0.022 | 1079938 | 100.000 |

Peak Table

PDA Ch2 254nm

| Peak# | Ret. Time | Height | Height% | USP Width | Area | Area% |
|---|---|---|---|---|---|---|
| 1 | 0.688 | 441003 | 100.000 | 0.022 | 356168 | 100.000 |

Confidential, for research only not for regulatory filing

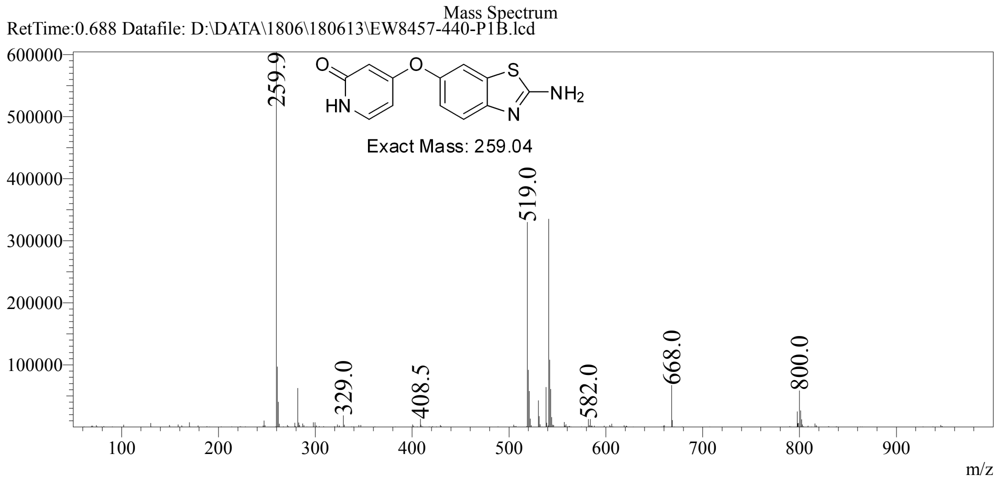

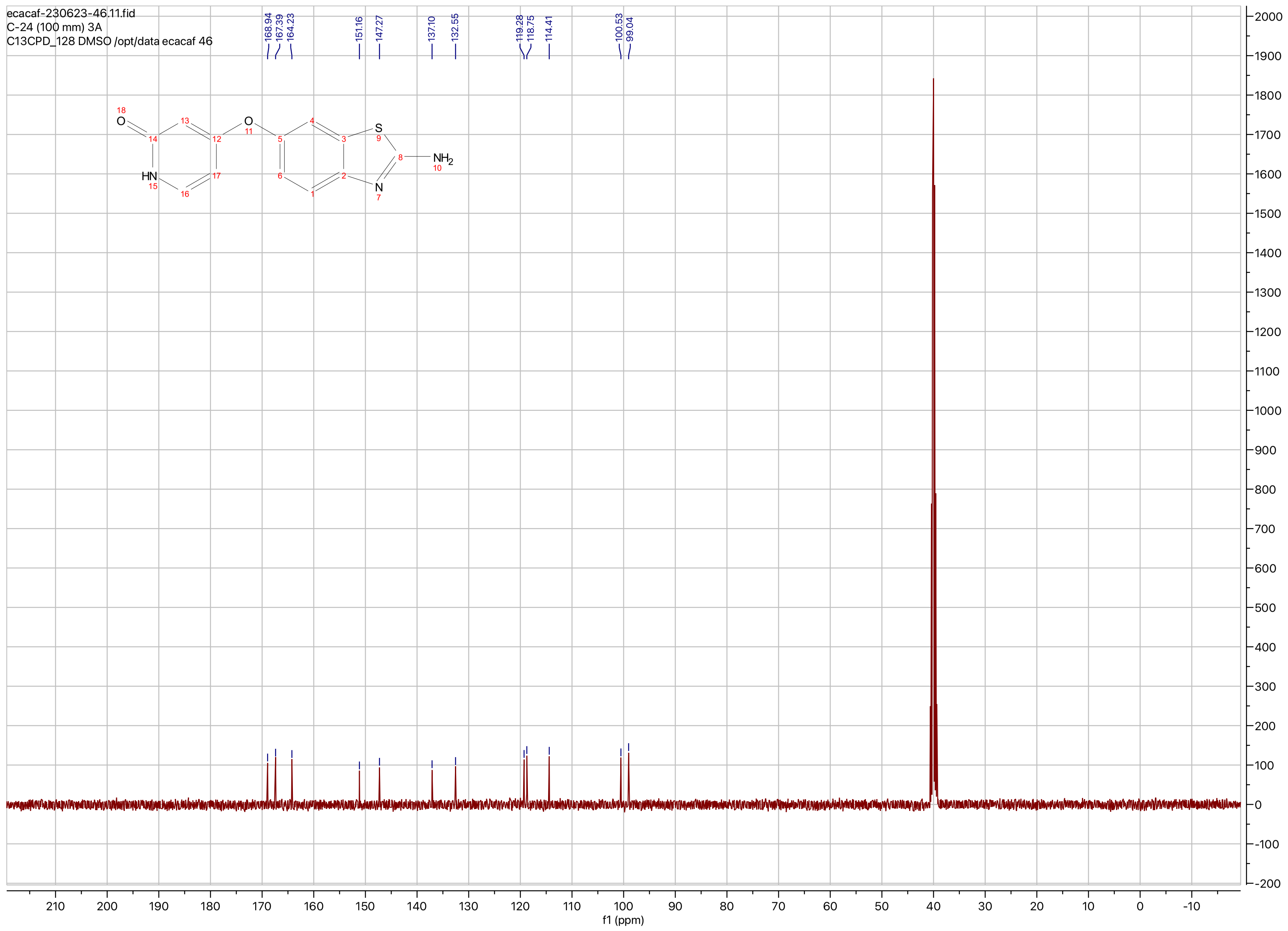
ecacaf-230623-46.11.fid
C-24 (100 mm) 3A
C13CPD_128 DMSO /opt/data ecacaf 46
168.94
167.39
164.23
151.16
147.27
137.10
132.55
119.28
118.75
114.41
100.53
99.04
O
HN
O
S
N
NH2
f1 (ppm)

**Compound ID: C-25**

EW8457-506-P1A DMSO Bruker_A_400MHz

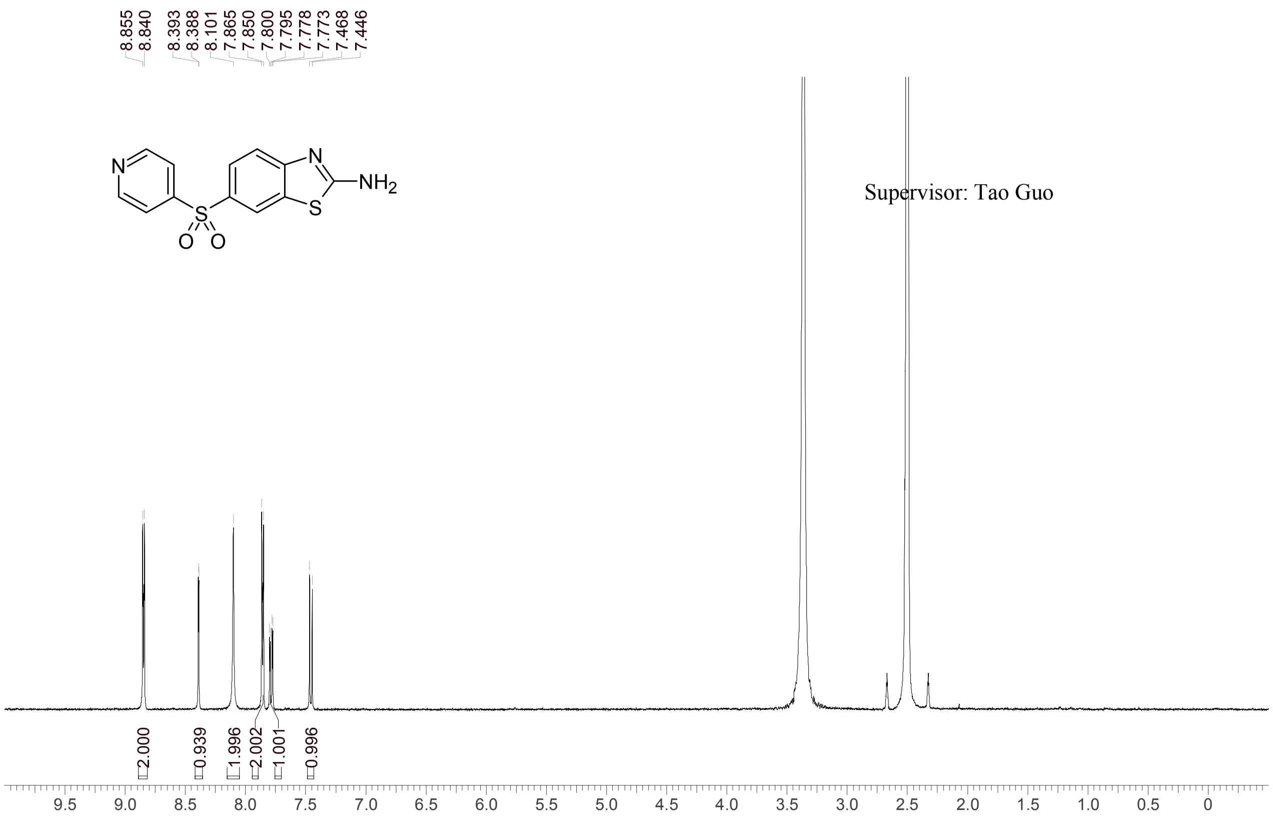


Supervisor: Tao Guo

| | |
|---|---|
| Acquisition Time (sec) | 3.0671 |
| Comment | EW8457-506-P1A DMSO Bruker_A_400MHz |
| Date | 25 Jul 2018 08:33:50 |
| Frequency (MHz) | 400.1300 |
| Nucleus | 1H |
| Number of Transients | 8 |
| Origin | spect |
| Original Points Count | 24576 |
| Owner | nmrsu |
| Points Count | 65536 |
| Pulse Sequence | zg30 |
| Receiver Gain | 120.01 |
| SW(cyclical) (Hz) | 8012.82 |
| Solvent | DMSO-d6 |
| Spectrum Offset (Hz) | 2397.5386 |
| Spectrum Type | standard |
| Sweep Width (Hz) | 8012.70 |
| Temperature (degree C) | 25.147 |

Operator:

Date:

Confidential, for research only not for regulatory filing

Confidential, for research only not for regulatory filing

### LCMS REPORT

Compound ID : C-25
Sample ID : EW8457-506-P1B
Injection Vol : 10ul
Location : vial104
Acq Method : d:\method\5-95CD_R_220&254_POS.lcm
Org DataFile : D:\DATA\1807\180724\EW8457-506-P1B.lcd
Injection Date : 2018-07-24 14:29:59
Instrument :LCMS-O 15-105

Chromatogram

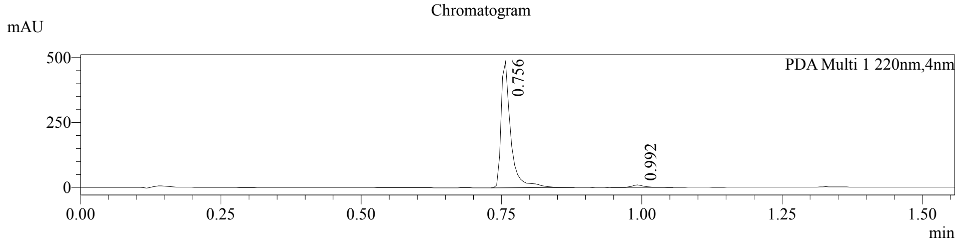


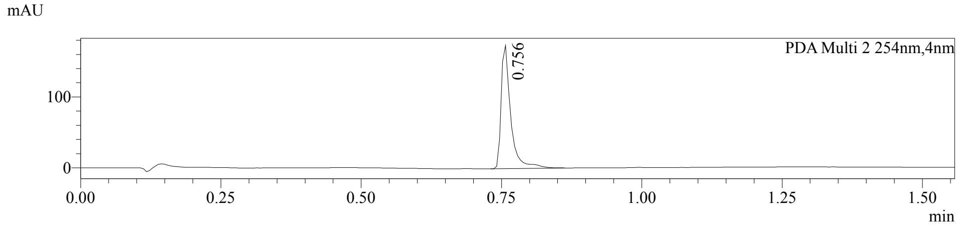


1 PDA Multi 1 / 220nm,4nm
2 PDA Multi 2 / 254nm,4nm

MS Chromatogram

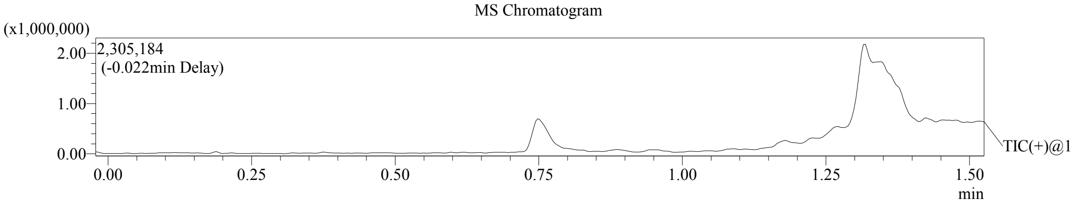


Integration Result

Peak Table

PDA Ch1 220nm

| Peak# | Ret. Time | Height | Height% | USP Width | Area | Area% |
|---|---|---|---|---|---|---|
| 1 | 0.756 | 485414 | 98.097 | 0.100 | 576676 | 97.598 |
| 2 | 0.992 | 9416 | 1.903 | 0.044 | 14193 | 2.402 |

Peak Table

PDA Ch2 254nm

| Peak# | Ret. Time | Height | Height% | USP Width | Area | Area% |
|---|---|---|---|---|---|---|
| 1 | 0.756 | 173755 | 100.000 | 0.099 | 205034 | 100.000 |

Confidential, for research only not for regulatory filing

Mass Spectrum

RetTime: 0.755 Datafile: D:\DATA\1807\180724\EW8457-506-P1B.lcd

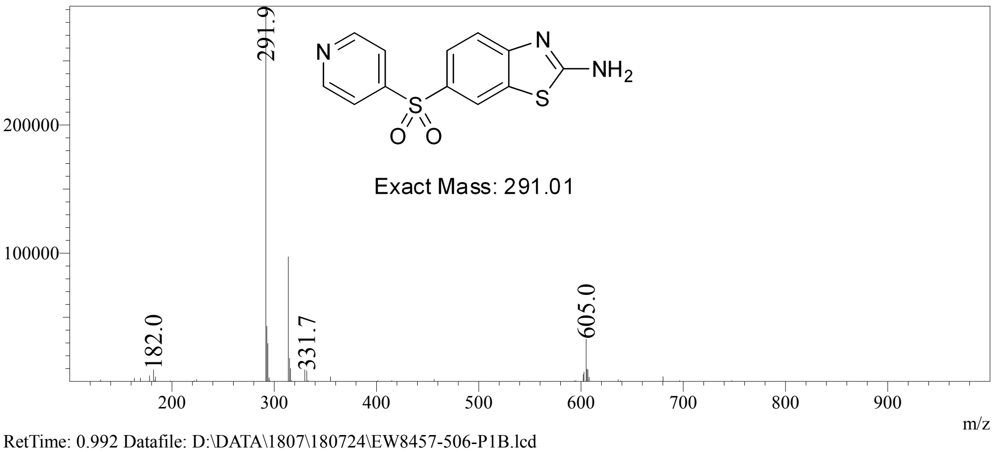


RetTime: 0.992 Datafile: D:\DATA\1807\180724\EW8457-506-P1B.lcd

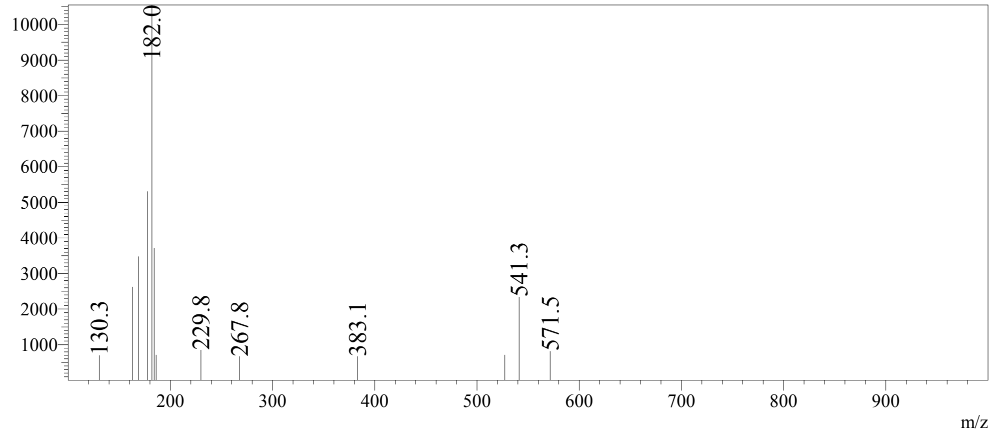

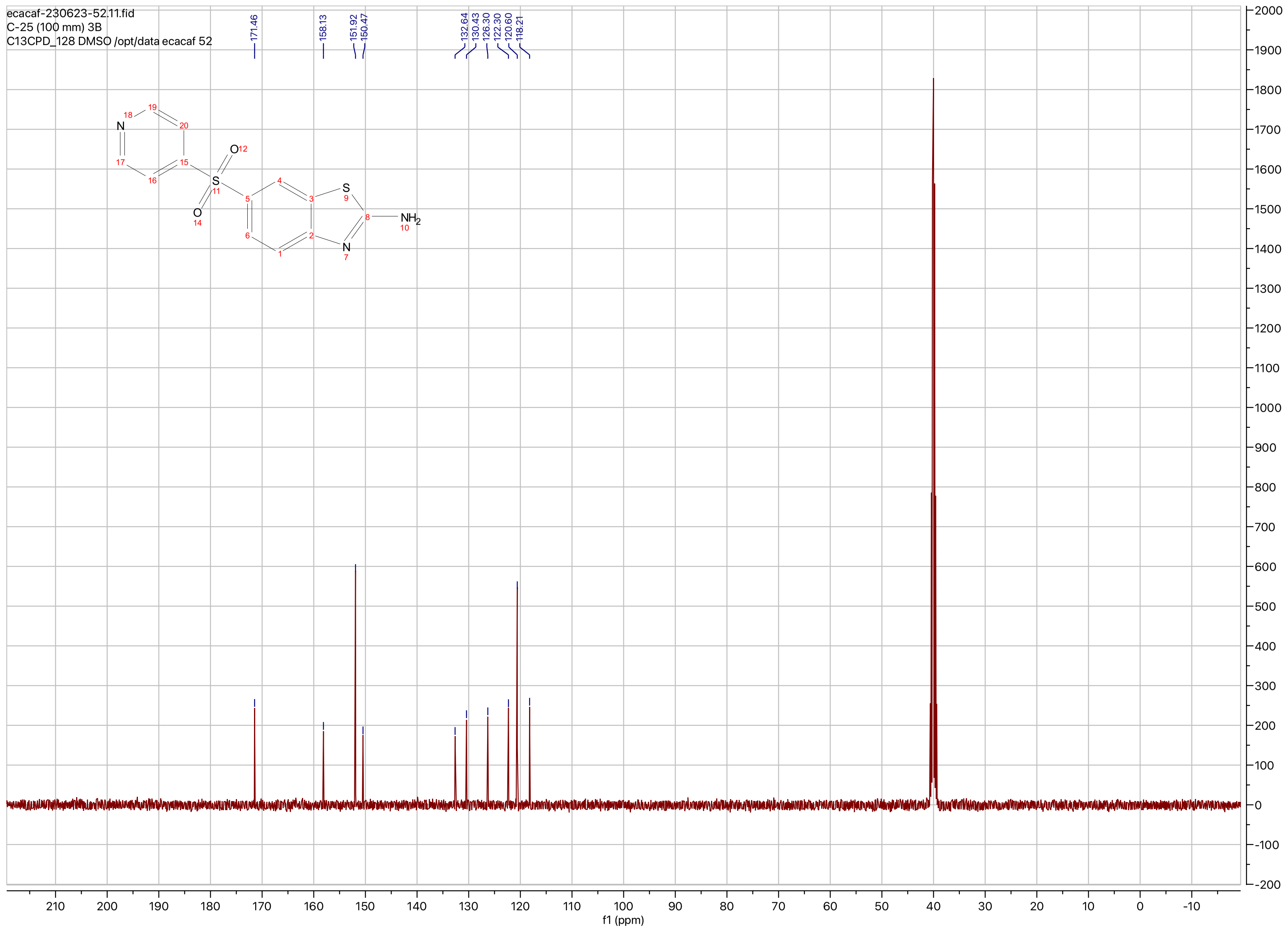
ecacaf-230623-52.11.fid
C-25 (100 mm) 3B
C13CPD_128 DMSO /opt/data ecacaf 52
171.46
158.13
151.92
150.47
132.64
130.43
126.30
122.30
120.60
118.21
f1 (ppm)

**Compound ID: C-26**

ET27821-71-P1H1 DMSO Bruker_J_400MHz

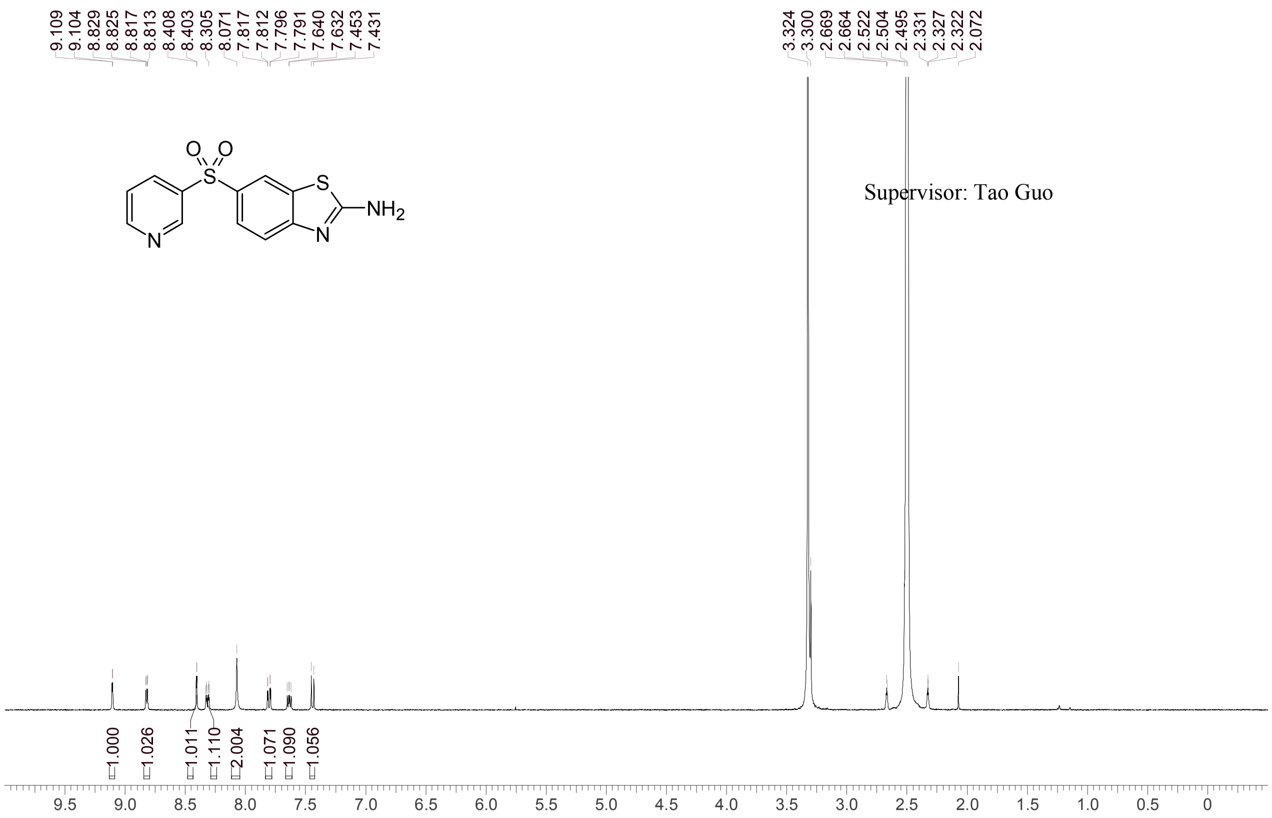


Supervisor: Tao Guo

| | |
|---|---|
| Acquisition Time (sec) | 1.9988 |
| Comment | ET27821-71-P1H1 DMSO Bruker_J_400MHz |
| Date | 19 Jun 2019 02:42:57 |
| Frequency (MHz) | 400.1500 |
| Nucleus | 1H |
| Number of Transients | 8 |
| Origin | Avance |
| Original Points Count | 16384 |
| Owner | nmr |
| Points Count | 65536 |
| Pulse Sequence | zg30 |
| Receiver Gain | 101.00 |
| SW(cyclical) (Hz) | 8196.72 |
| Solvent | DMSO-d6 |
| Spectrum Offset (Hz) | 2469.1653 |
| Spectrum Type | standard |
| Sweep Width (Hz) | 8196.60 |
| Temperature (degree C) | 25.115 |

Operator: Date:

Confidential, for research only not for regulatory filing

LCMS REPORT

Compound ID : C-26 ->
Sample ID : ET27821-71-P1C
Injection Date : 18. Jun. 2019
Inj. Vol. : 1.00 ul
Location : P2-E-03
Acq Method : D:\DATA\190618-SD 18\5_95CD_2MIN-220-254
Data Filename : D:\DATA\190618-SD 18\ET27821-71-P1C.D
Instrument : S

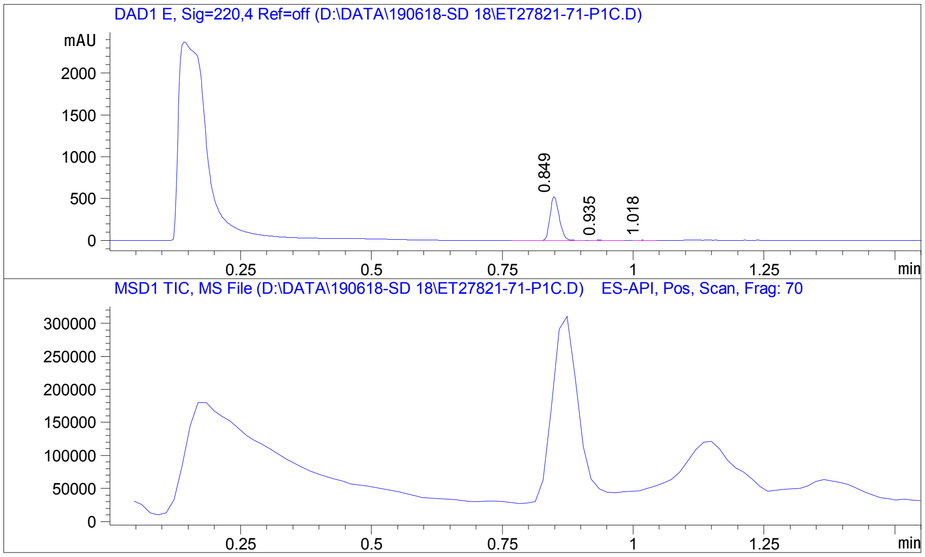


Integration Result

Signal 1 : DAD1 E, Sig=220,4 Ref=off

| Peak # | RT [min] | Area | Height | Height % | Width [min] | Area % |
|---|---|---|---|---|---|---|
| 1 | 0.849 | 642.595 | 520.126 | 99.033 | 0.019 | 98.746 |
| 2 | 0.935 | 5.429 | 3.089 | 0.588 | 0.025 | 0.834 |
| 3 | 1.018 | 2.734 | 1.989 | 0.379 | 0.020 | 0.420 |

Operator:_____________

Date:_________________

Confidential. For research information only
PDF created with pdfFactory Pro trial version www.pdffactory.com

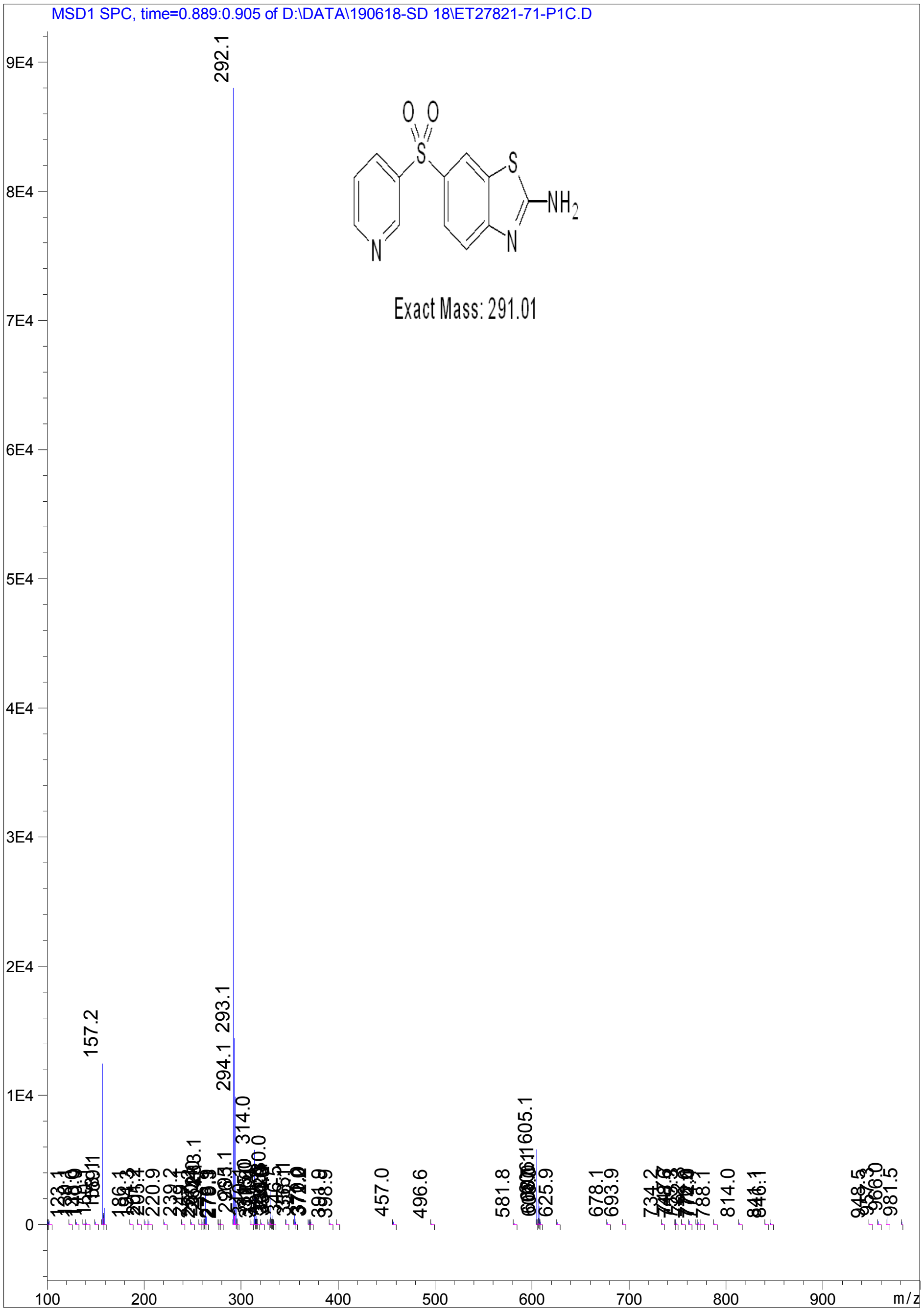


Confidential. For research information only

PDF created with pdfFactory Pro trial version www.pdffactory.com

Print of window 80: MS Spectrum

MS Spectrum

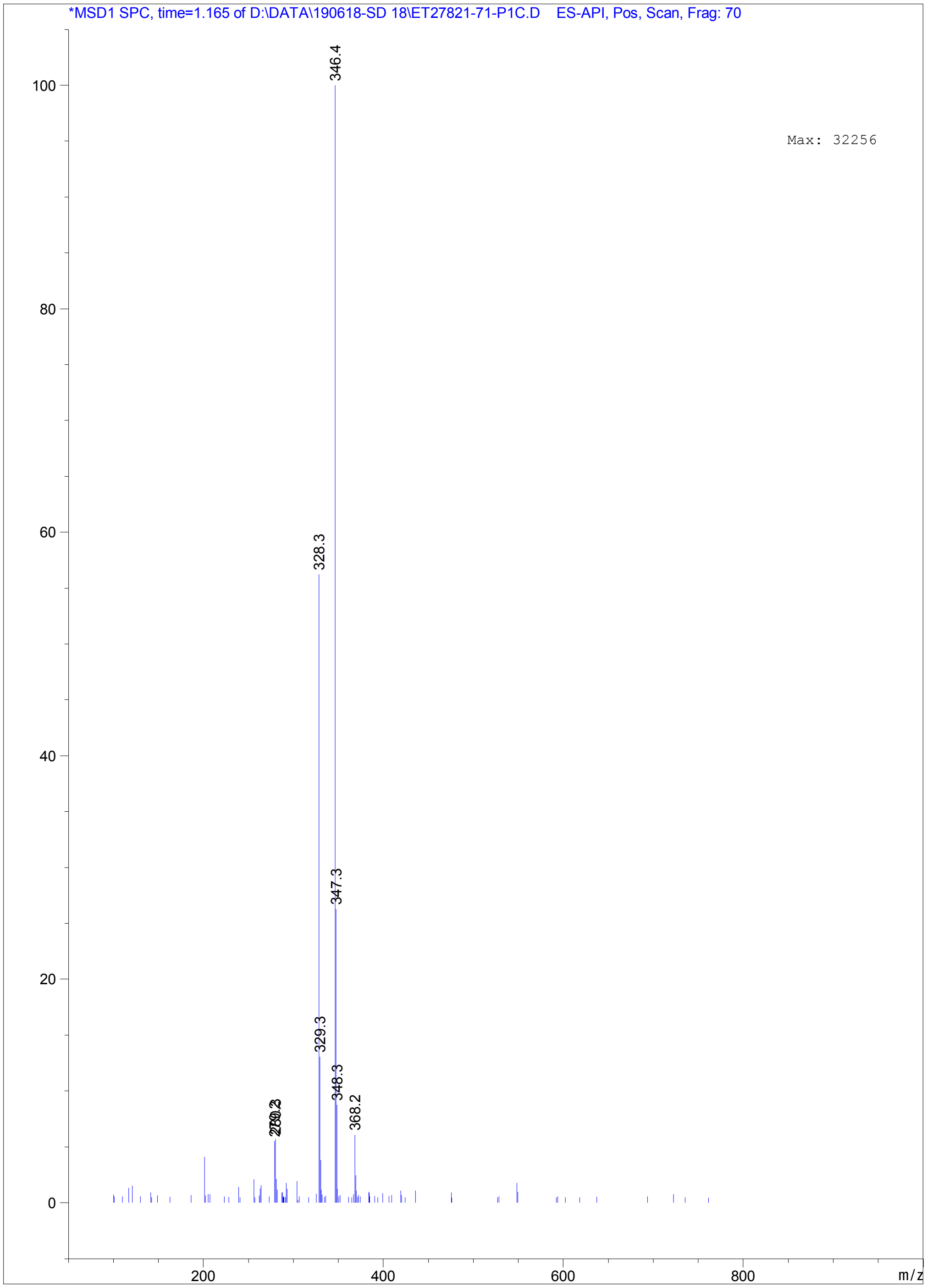




PDF created with pdfFactory Pro trial version www.pdffactory.com

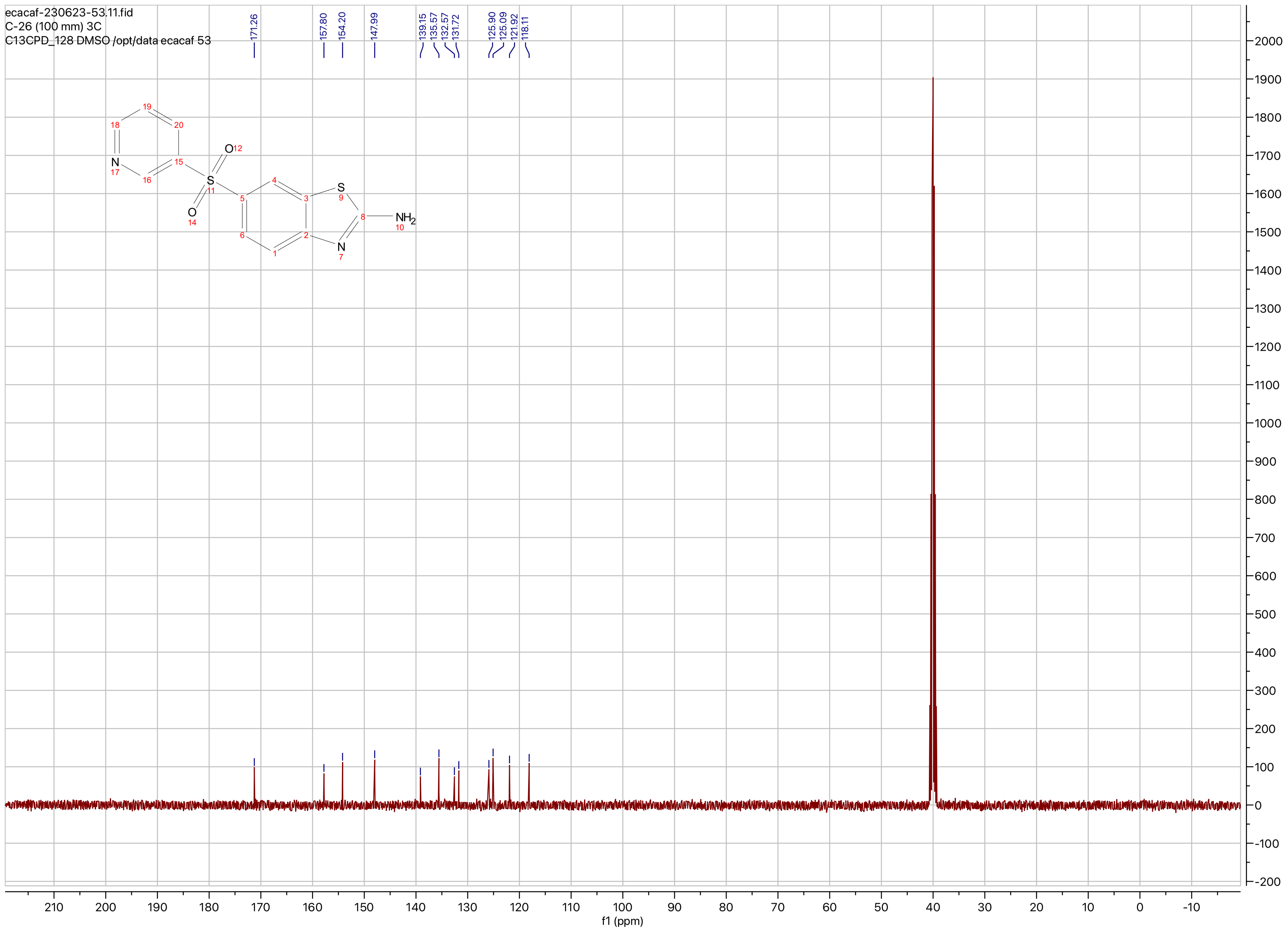
ecacaf-230623-53.11.fid
C-26 (100 mm) 3C
C13CPD_128 DMSO /opt/data ecacaf 53
171.26
157.80
154.20
147.99
139.15
135.57
132.57
131.72
125.90
125.09
121.92
118.11
NH2
f1 (ppm)

**Compound ID:C-27**

EW8457-830-P1F2 DMSO Bruker_A_400MHz

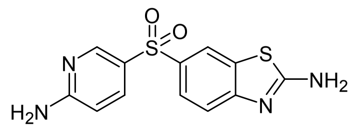


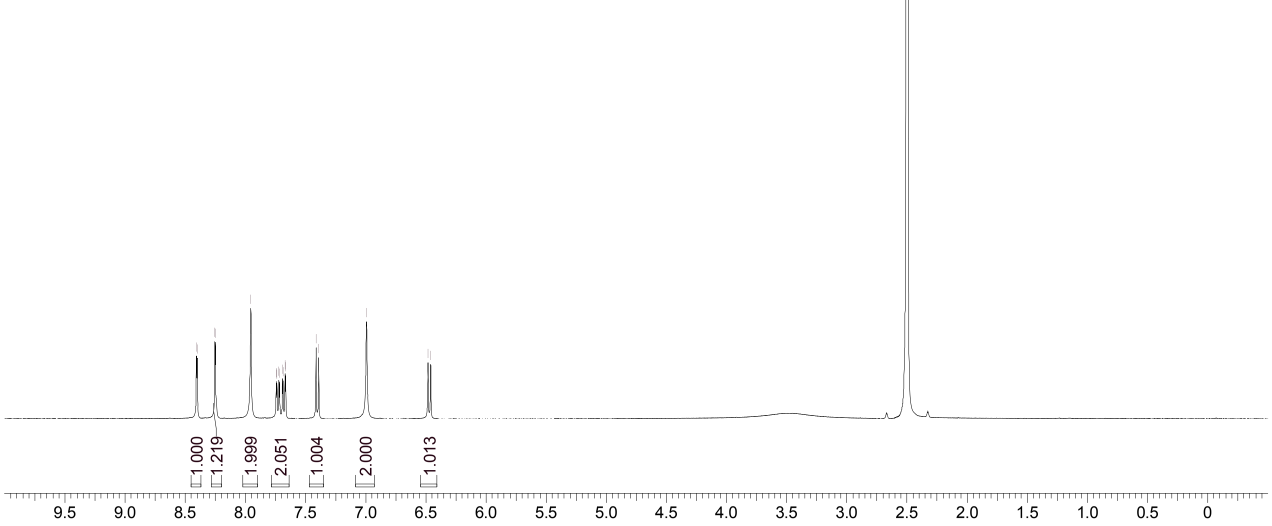


Supervisor: Tao Guo

| | |
|---|---|
| Acquisition Time (sec) | 3.0671 |
| Comment | EW8457-830-P1F2 DMSO Bruker_A_400MHz |
| Date | 07 Jan 2019 09:41:24 |
| Frequency (MHz) | 400.1300 |
| Nucleus | 1H |
| Number of Transients | 8 |
| Origin | spect |
| Original Points Count | 24576 |
| Owner | nmrsu |
| Points Count | 65536 |
| Pulse Sequence | zg30 |
| Receiver Gain | 175.36 |
| SW(cyclical) (Hz) | 8012.82 |
| Solvent | DMSO-d6 |
| Spectrum Offset (Hz) | 2397.5386 |
| Spectrum Type | standard |
| Sweep Width (Hz) | 8012.70 |
| Temperature (degree C) | 27.156 |

Operator:

Date:

Confidential, for research only not for regulatory filing

Confidential, for research only not for regulatory filing

### LCMS REPORT

Compound ID : C-27
Sample ID : EW8457-830-P1F
Injection Vol : 5ul
Location : vial13
Acq Method : d:\method\5-95CD_R_220&254_POS.lcm
Org DataFile : D:\DATA\1901\190107\EW8457-830-P1F.lcd
Injection Date : 2019-01-07 9:38:30
Instrument :LCMS-O 15-105

Chromatogram

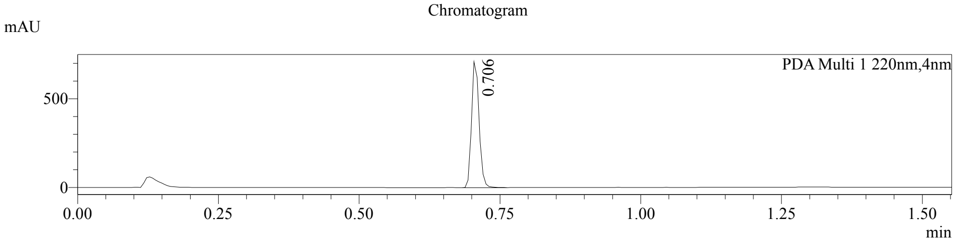


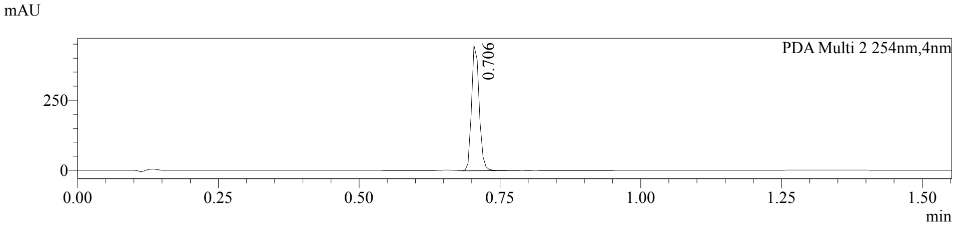


1 PDA Multi 1 / 220nm,4nm
2 PDA Multi 2 / 254nm,4nm

MS Chromatogram

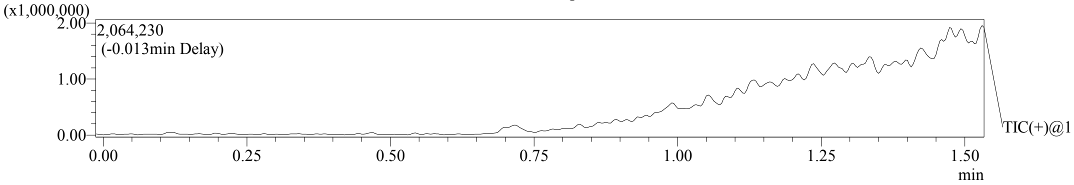


Integration Result

Peak Table

PDA Ch1 220nm

| Peak# | Ret. Time | Height | Height% | USP Width | Area | Area% |
|---|---|---|---|---|---|---|
| 1 | 0.706 | 710719 | 100.000 | 0.028 | 657252 | 100.000 |

Peak Table

PDA Ch2 254nm

| Peak# | Ret. Time | Height | Height% | USP Width | Area | Area% |
|---|---|---|---|---|---|---|
| 1 | 0.706 | 446814 | 100.000 | 0.028 | 416228 | 100.000 |

Confidential, for research only not for regulatory filing

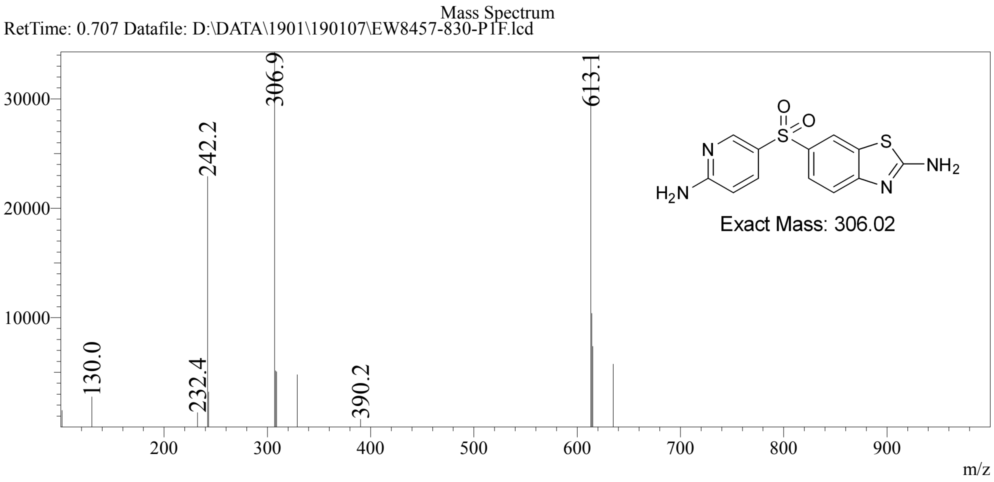

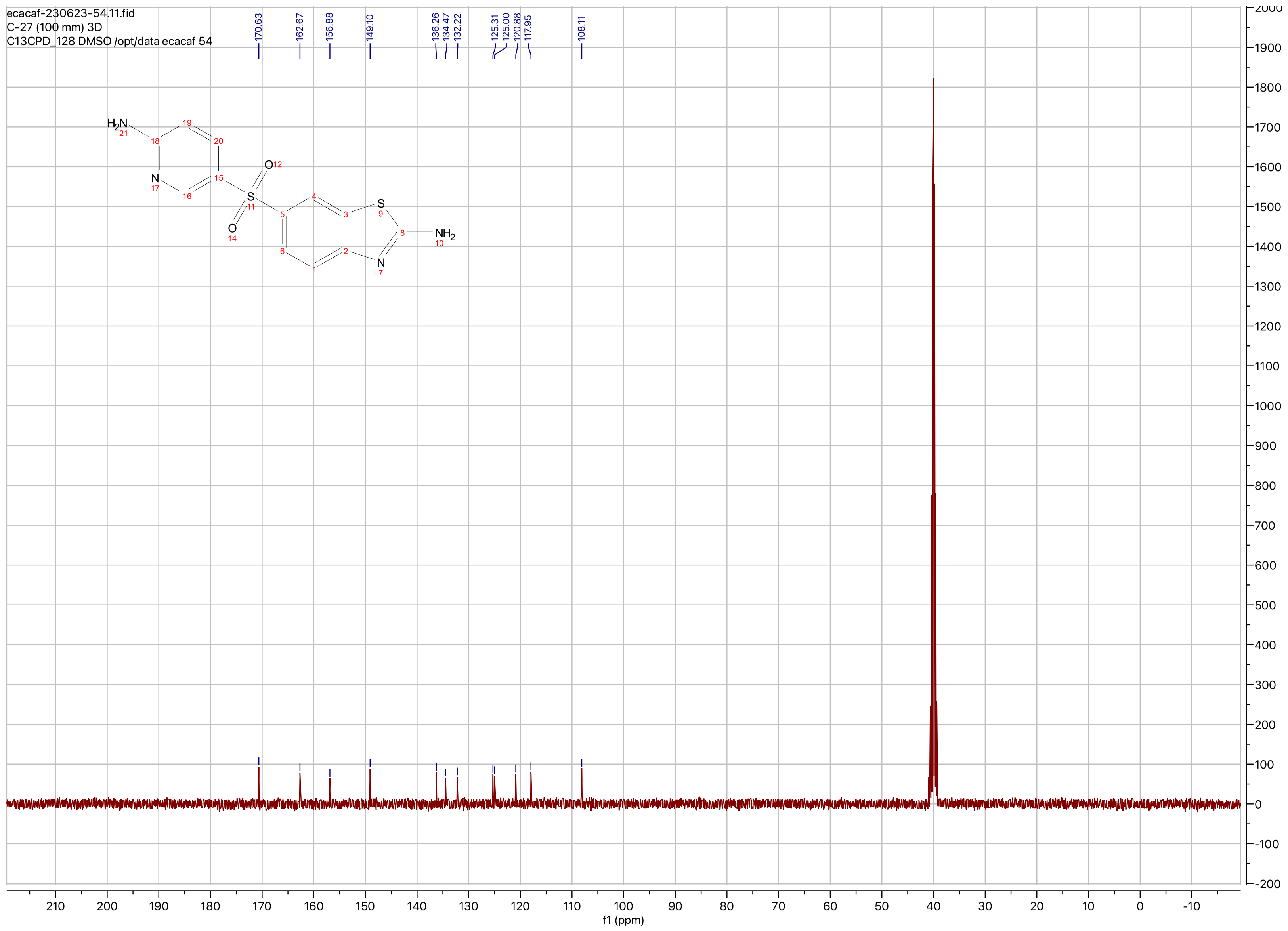
ecacaf-230623-54.11.fid
C-27 (100 mm) 3D
C13CPD_128 DMSO /opt/data ecacaf 54
170.63
162.67
156.88
149.10
136.26
134.47
132.22
125.31
125.00
120.88
117.95
108.11
f1 (ppm)

**Compound ID: C-28**

EW12557-369-p1a DMSO Bruker_A_400MHz

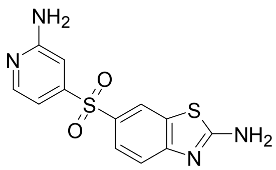

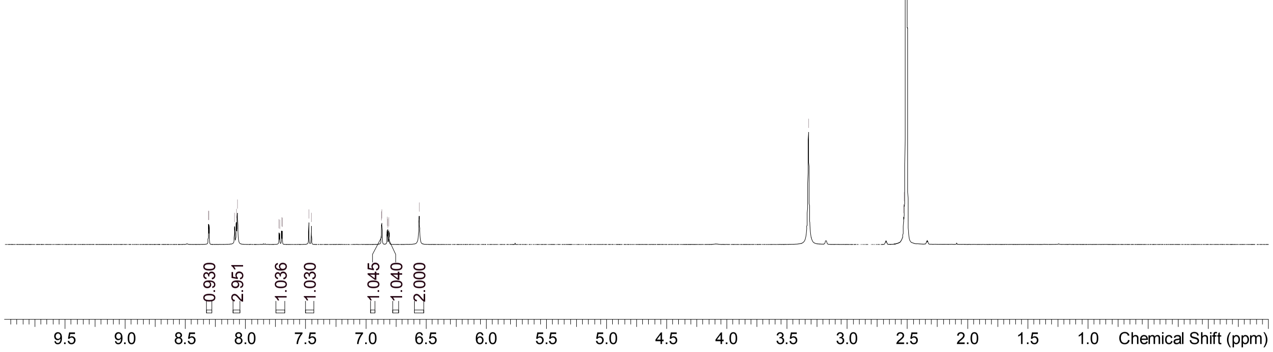


Supervisor: Tao Guo

| | |
|---|---|
| Acquisition Time (sec) | 3.0671 |
| Comment | EW12557-369-p1a DMSO Bruker_A_400MHz |
| Date | 15 Jan 2019 10:23:37 |
| Frequency (MHz) | 400.1300 |
| Nucleus | 1H |
| Number of Transients | 8 |
| Origin | spect |
| Original Points Count | 24576 |
| Owner | nmrsu |
| Points Count | 65536 |
| Pulse Sequence | zg30 |
| Receiver Gain | 175.36 |
| SW(cyclical) (Hz) | 8012.82 |
| Solvent | DMSO-d6 |
| Spectrum Offset (Hz) | 2400.7795 |
| Spectrum Type | standard |
| Sweep Width (Hz) | 8012.70 |
| Temperature (degree C) | 27.149 |

Operator:

Date:

Confidential, for research only not for regulatory filing

Confidential, for research only not for regulatory filing

### LCMS REPORT

Compound ID : C-28
Sample ID : EW12557-369-P1B
Injection Vol : 5ul
Location : vial10
Acq Method : d:\method\5-95CD_R_220&254_POS.lcm
Org DataFile : D:\DATA\1901\190116\EW12557-369-P1B.lcd
Injection Date : 16/01/2019 08:39:10
Instrument :LCMS-K 15-105

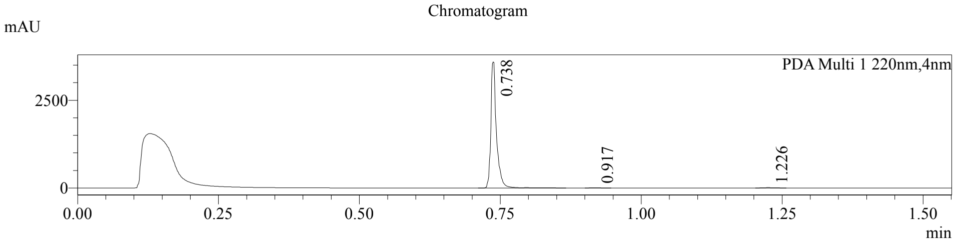


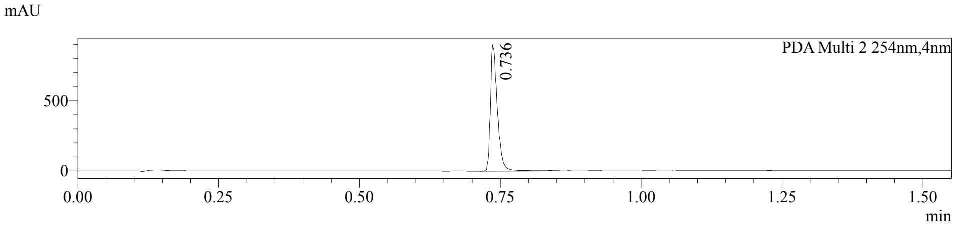


1 PDA Multi 1 / 220nm,4nm
2 PDA Multi 2 / 254nm,4nm

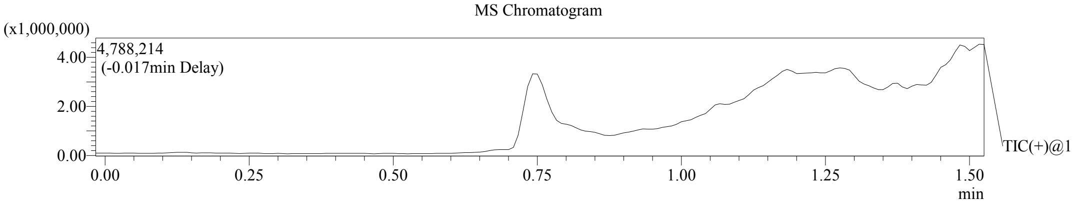


Integration Result

Peak Table

PDA Ch1 220nm

| Peak# | Ret. Time | Height | Height% | USP Width | Area | Area% |
|---|---|---|---|---|---|---|
| 1 | 0.738 | 3592181 | 99.566 | 0.018 | 2521936 | 99.384 |
| 2 | 0.917 | 7141 | 0.198 | 0.027 | 6523 | 0.257 |
| 3 | 1.226 | 8504 | 0.236 | 0.030 | 9114 | 0.359 |

Peak Table

PDA Ch2 254nm

| Peak# | Ret. Time | Height | Height% | USP Width | Area | Area% |
|---|---|---|---|---|---|---|
| 1 | 0.736 | 895063 | 100.000 | 0.023 | 772948 | 100.000 |

Confidential, for research only not for regulatory filing

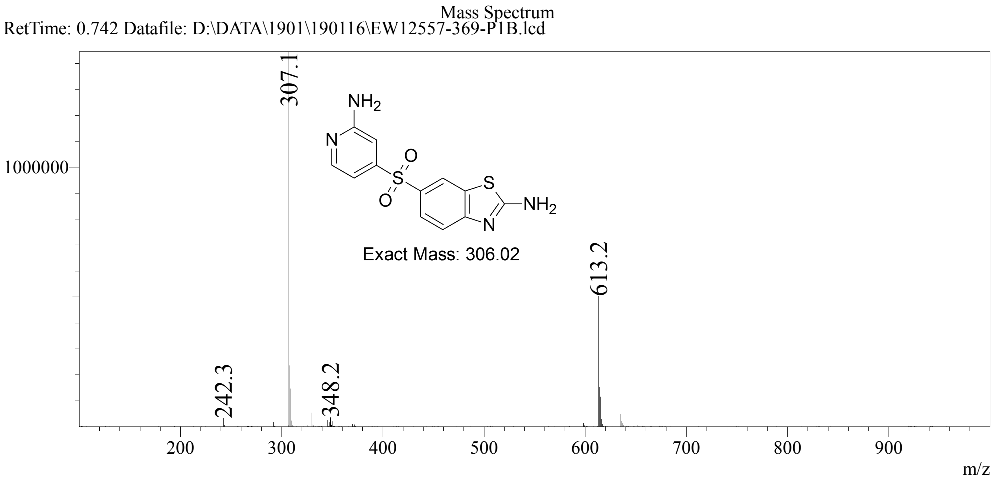

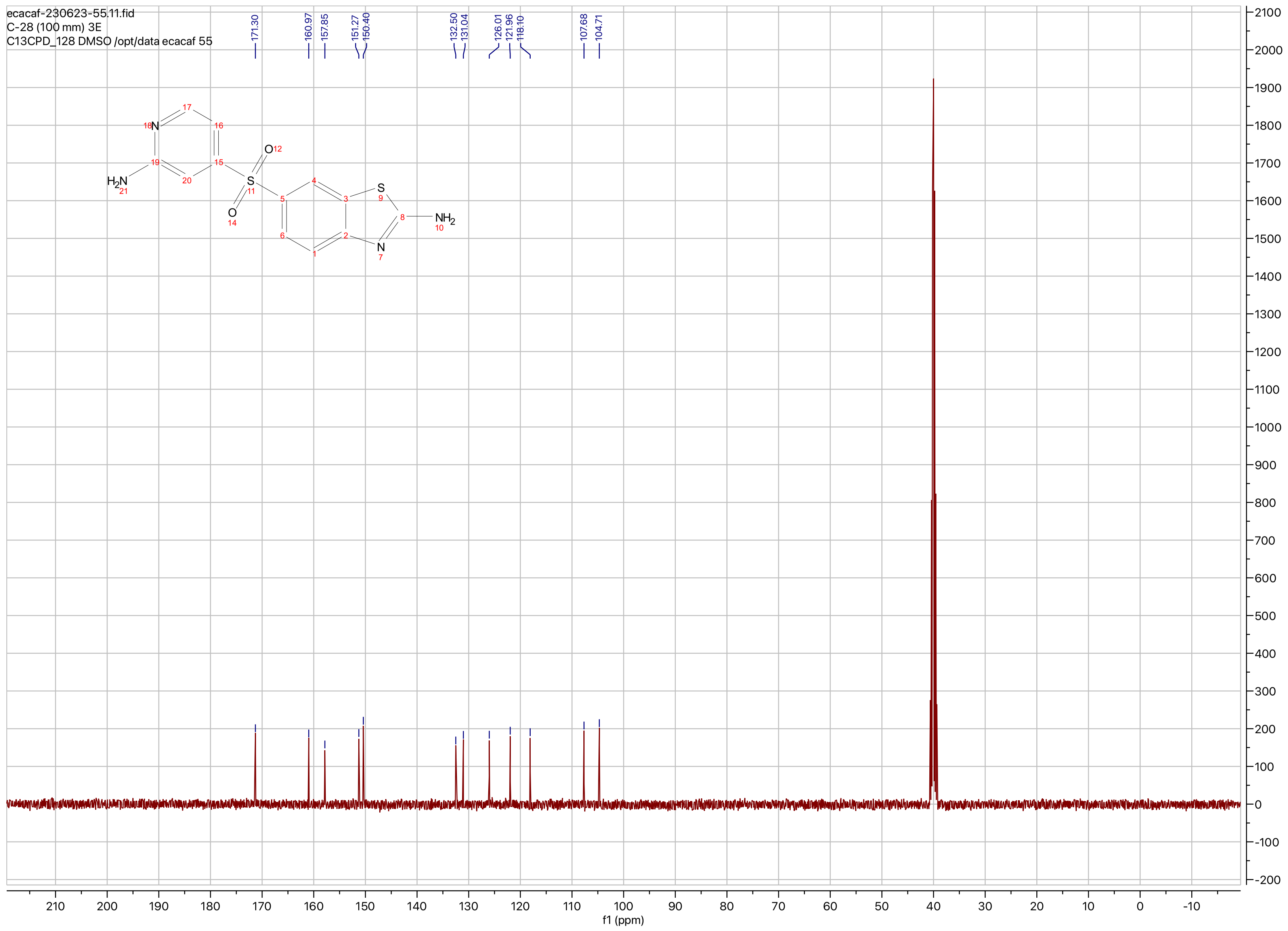

ecacaf-230623-55.11.fid
C-28 (100 mm) 3E
C13CPD_128 DMSO /opt/data ecacaf 55
171.30
160.97
157.85
151.27
150.40
132.50
131.04
126.01
121.96
118.10
107.68
104.71
f1 (ppm)

**Compound ID: C-29**

ET27821-90-P1H1 DMSO Bruker_K_400MHz

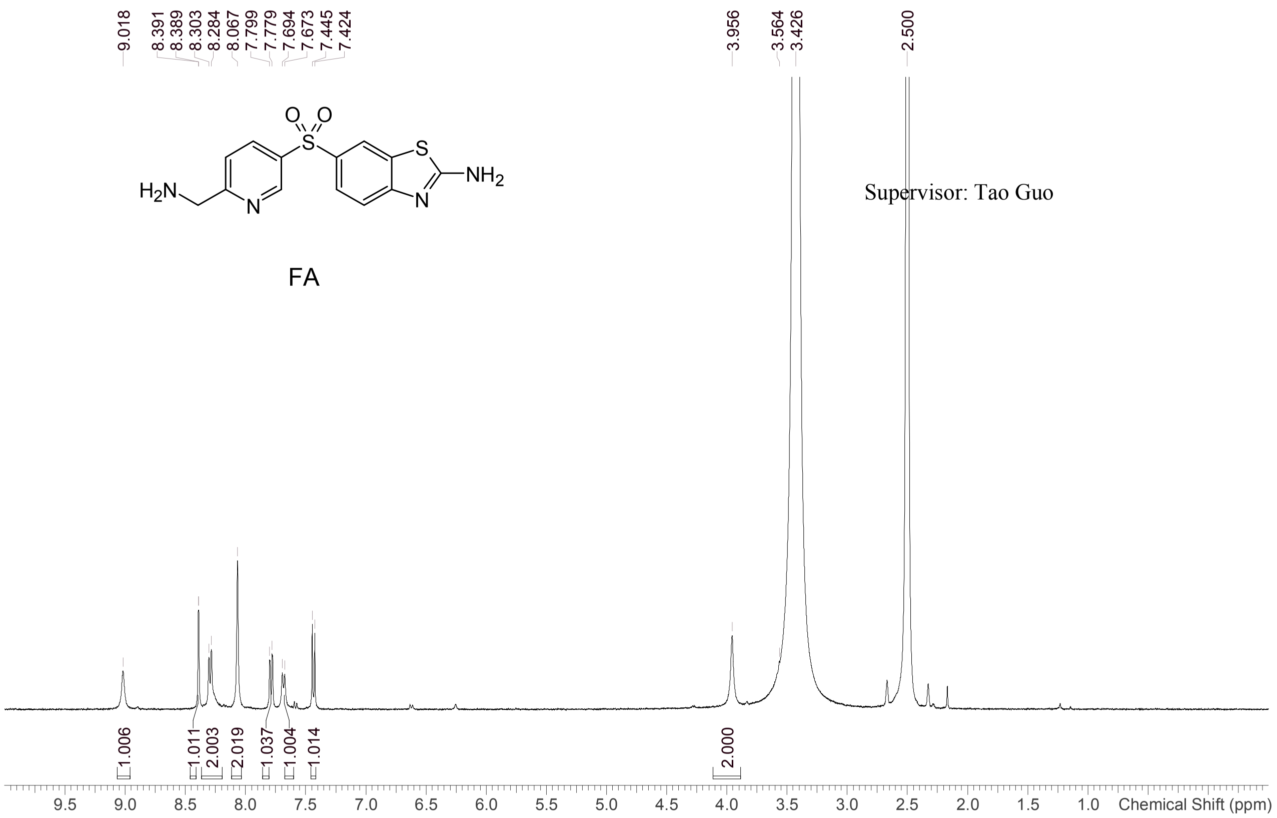


Supervisor: Tao Guo

| | |
|---|---|
| Acquisition Time (sec) | 1.9988 |
| Comment | ET27821-90-P1H1 DMSO Bruker_K_400MHz |
| Date | 28 Jun 2019 05:03:50 |
| Frequency (MHz) | 400.1300 |
| Nucleus | 1H |
| Number of Transients | 8 |
| Origin | Avance |
| Original Points Count | 16384 |
| Owner | nmr |
| Points Count | 65536 |
| Pulse Sequence | zg30 |
| Receiver Gain | 101.00 |
| SW(cyclical) (Hz) | 8196.72 |
| Solvent | DMSO-d6 |
| Spectrum Offset (Hz) | 2463.4790 |
| Spectrum Type | standard |
| Sweep Width (Hz) | 8196.60 |
| Temperature (degree C) | 23.287 |

Operator:

Date:

Confidential, for research only not for regulatory filing

LCMS REPORT

Compound ID : C - 29 ->
Sample ID : ET27821-90-P1H
Injection Date : 28. Jun. 2019
Inj. Vol. : 0.5 ul
Location : P2-C-03
Acq Method : D:\DATA\1906\190628-KD 9\5_95AB_6min-220.M
Data Filename : D:\DATA\1906\190628-KD 9\ET27821-90-P1H.D
Instrument : K

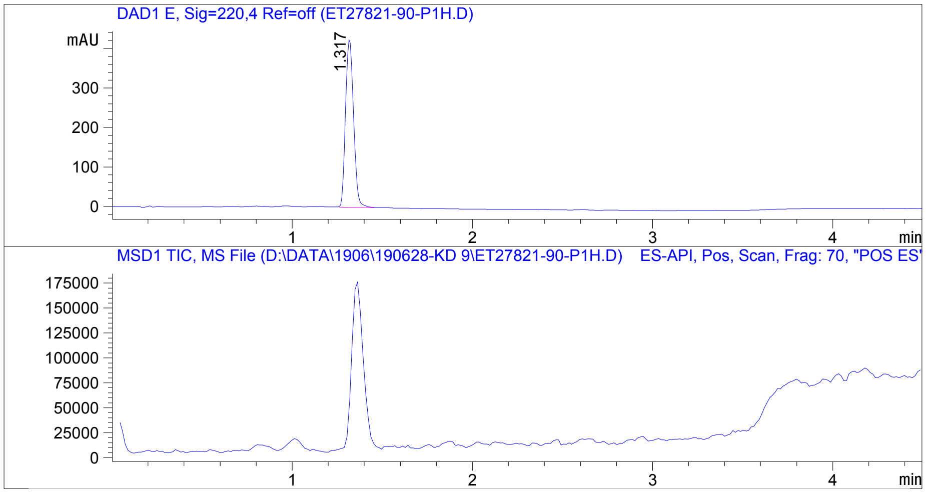


Integration Result

Signal 1 : DAD1 E, Sig=220,4 Ref=off

| Peak # | RT [min] | Area | Height | Height % | Width [min] | Area % |
|---|---|---|---|---|---|---|
| 1 | 1.317 | 1309.157 | 426.079 | 100.000 | 0.050 | 100.000 |

Operator:_____________

Date:_________________

Confidential. For research information only.

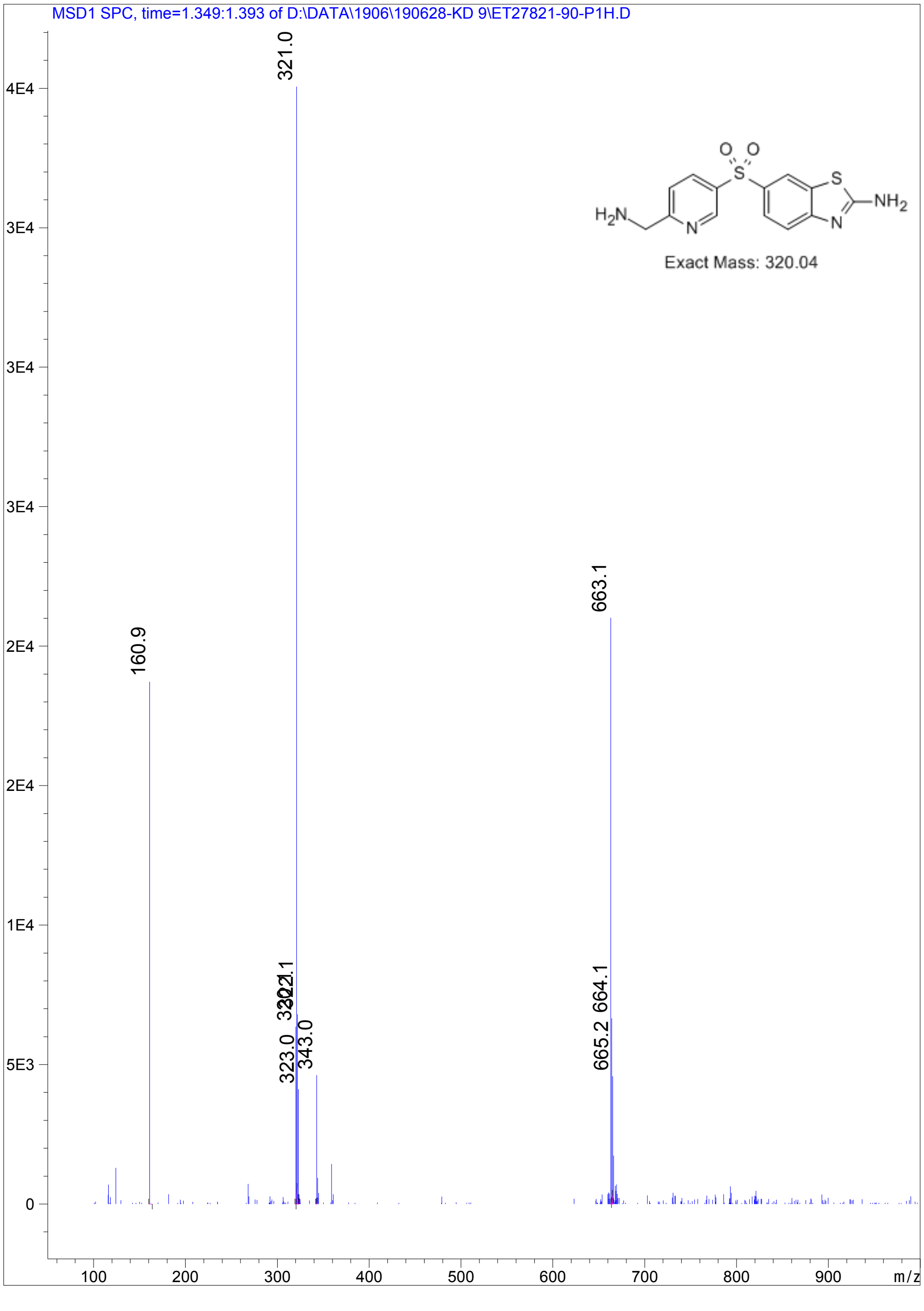


Confidential. For research information only.

ecacaf-230623-56.11.fid
C-29 (100 mm) 3F
C13CPD_128 DMSO /opt/data ecacaf 56

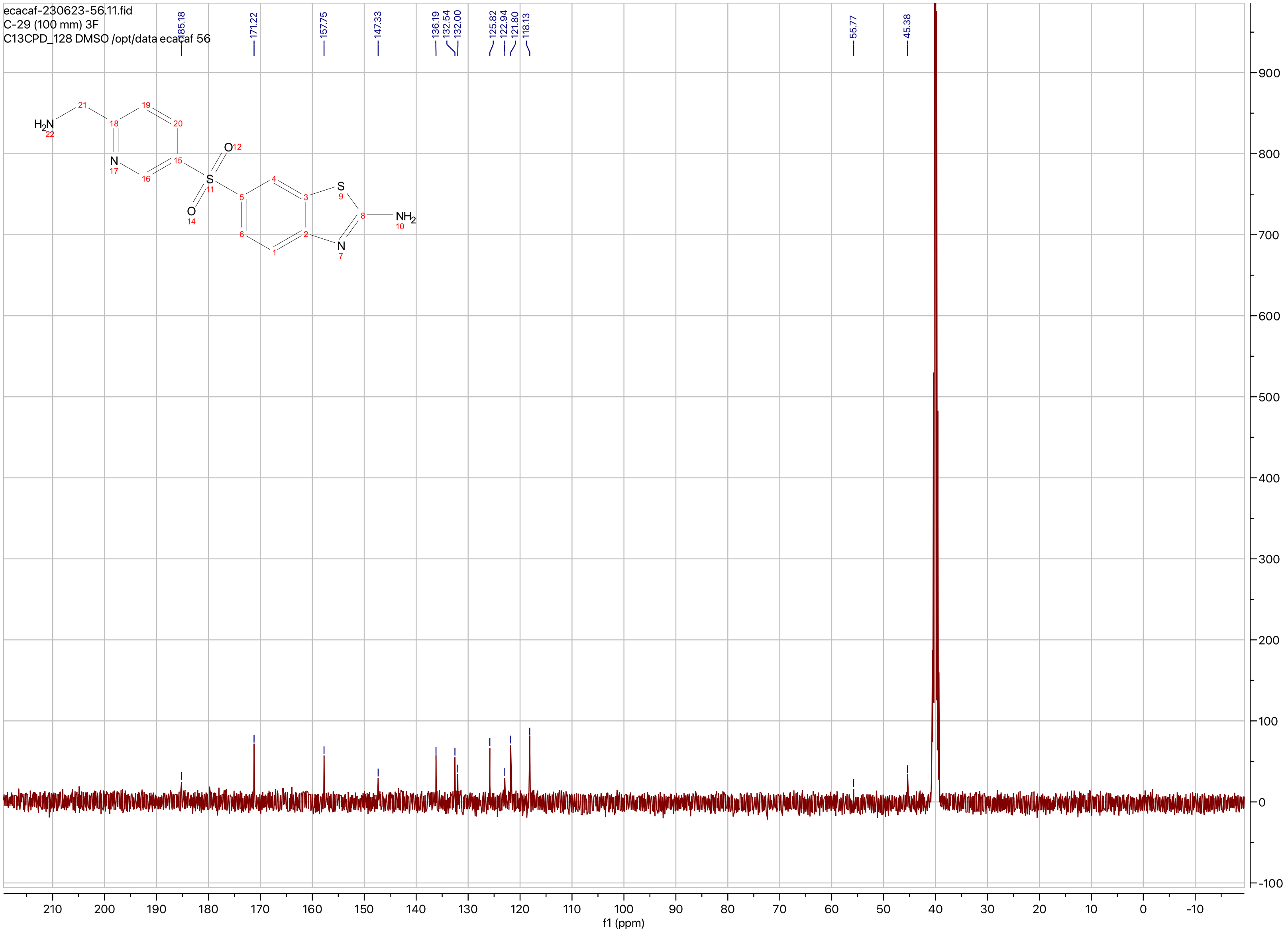

**Compound ID: C-30**

EW8457-501-P1A DMSO Bruker_B_400MHz

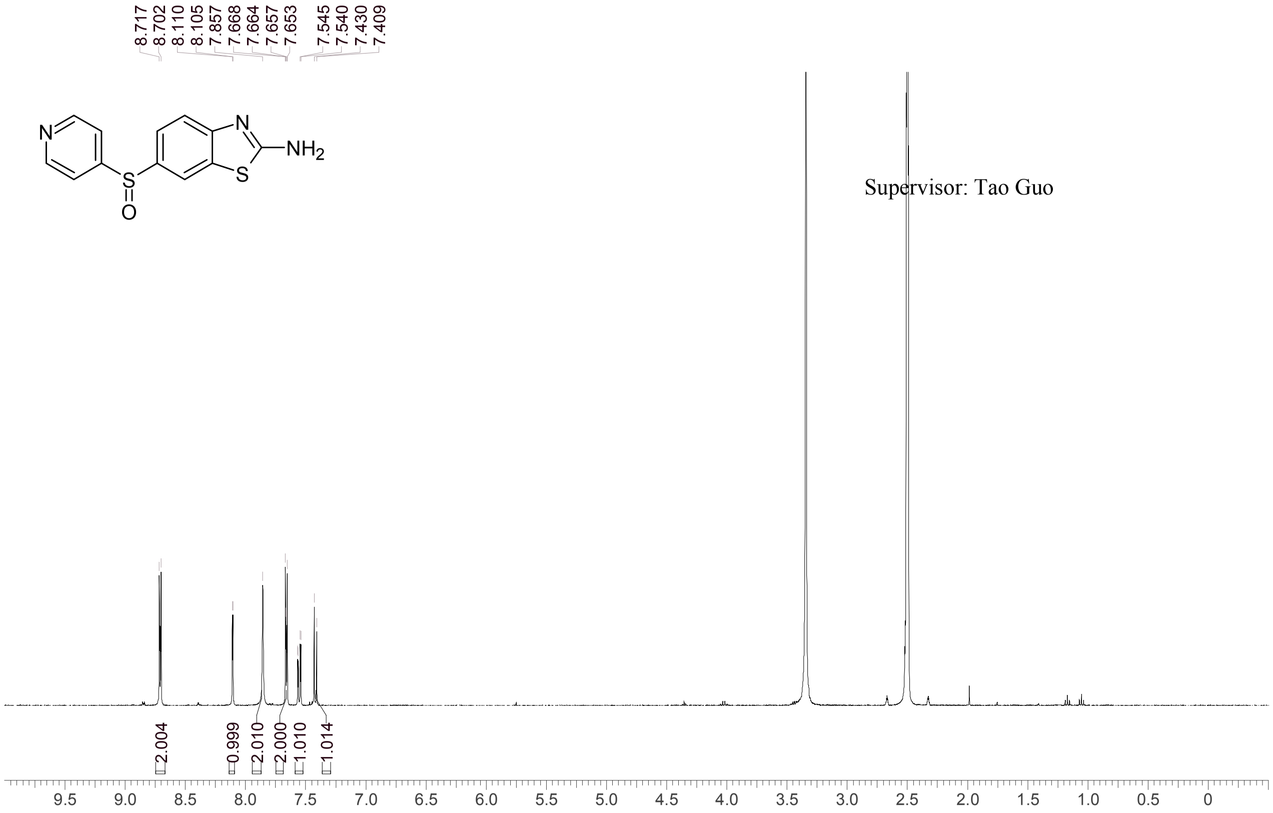


Supervisor: Tao Guo

| | |
|---|---|
| Acquisition Time (sec) | 3.0671 |
| Comment | EW8457-501-P1A DMSO Bruker_B_400MHz |
| Date | 20 Jul 2018 19:35:13 |
| Frequency (MHz) | 400.1300 |
| Nucleus | 1H |
| Number of Transients | 8 |
| Origin | spect |
| Original Points Count | 24576 |
| Owner | nmrsu |
| Points Count | 65536 |
| Pulse Sequence | zg30 |
| Receiver Gain | 190.49 |
| SW(cyclical) (Hz) | 8012.82 |
| Solvent | DMSO-d6 |
| Spectrum Offset (Hz) | 2397.3787 |
| Spectrum Type | standard |
| Sweep Width (Hz) | 8012.70 |
| Temperature (degree C) | 25.148 |

Operator: Date:

Confidential, for research only not for regulatory filing

Confidential, for research only not for regulatory filing

### LCMS REPORT

Compound ID : C-30
Sample ID : EW8457-501-P1D
Injection Vol : 2ul
Location : vial16
Acq Method : d:\method\5-95CD_R_220&254_POS.lcm
Org DataFile : D:\DATA\1807\180723\EW8457-501-P1D.lcd
Injection Date : 2018-07-23 14:18:07
Instrument :LCMS-O 15-105

Chromatogram

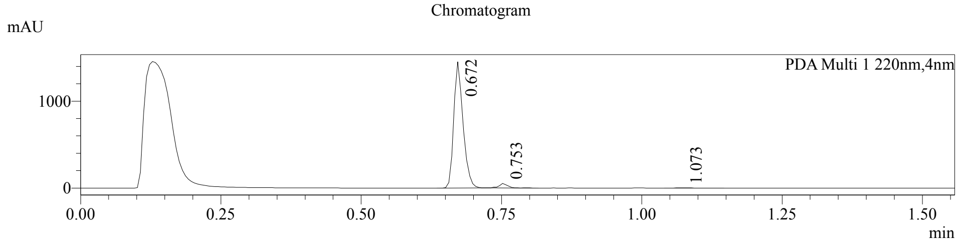


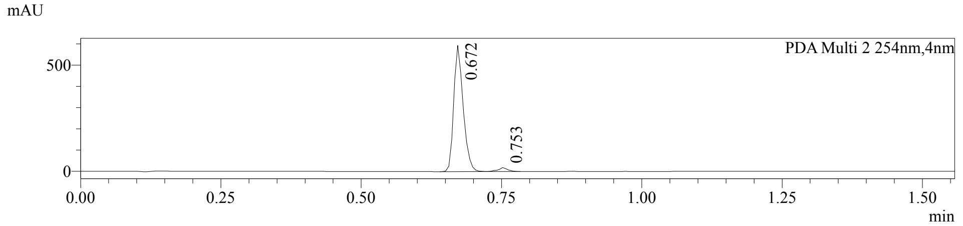


1 PDA Multi 1 / 220nm,4nm
2 PDA Multi 2 / 254nm,4nm

MS Chromatogram

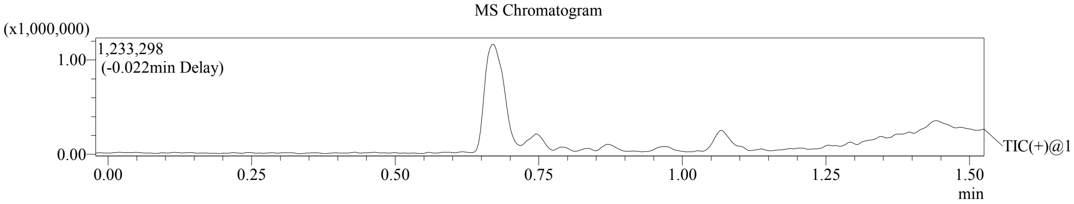


Integration Result

Peak Table

PDA Ch1 220nm

| Peak# | Ret. Time | Height | Height% | USP Width | Area | Area% |
|---|---|---|---|---|---|---|
| 1 | 0.672 | 1453120 | 96.289 | 0.037 | 1684000 | 96.314 |
| 2 | 0.753 | 52554 | 3.482 | 0.031 | 58103 | 3.323 |
| 3 | 1.073 | 3449 | 0.229 | 0.054 | 6338 | 0.362 |

Peak Table

PDA Ch2 254nm

| Peak# | Ret. Time | Height | Height% | USP Width | Area | Area% |
|---|---|---|---|---|---|---|
| 1 | 0.672 | 593605 | 96.913 | 0.037 | 697884 | 97.048 |
| 2 | 0.753 | 18911 | 3.087 | 0.032 | 21229 | 2.952 |

Confidential, for research only not for regulatory filing

Mass Spectrum

RetTime: 0.672 Datafile: D:\DATA\1807\180723\EW8457-501-P1D.lcd

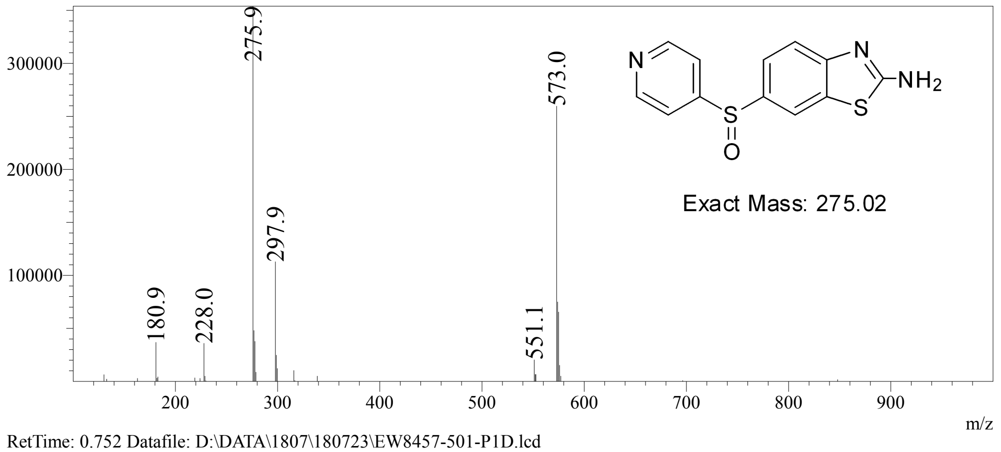


RetTime: 0.752 Datafile: D:\DATA\1807\180723\EW8457-501-P1D.lcd

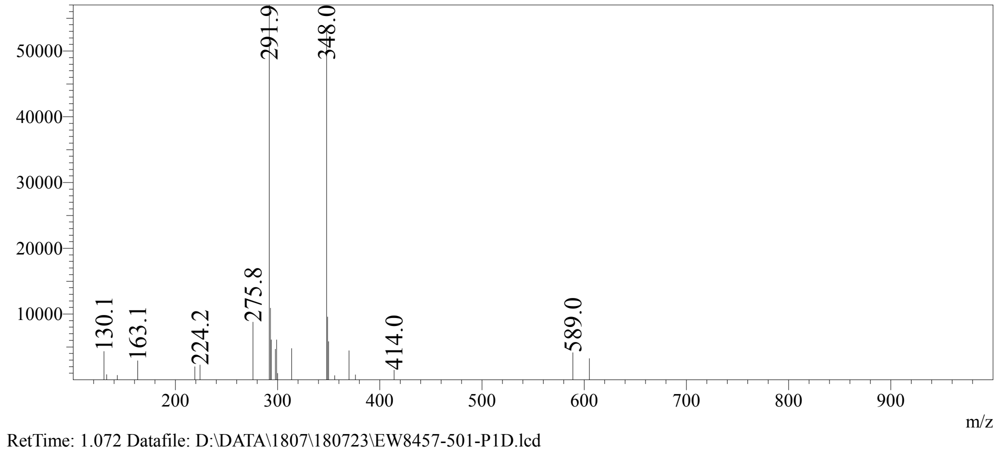


RetTime: 1.072 Datafile: D:\DATA\1807\180723\EW8457-501-P1D.lcd

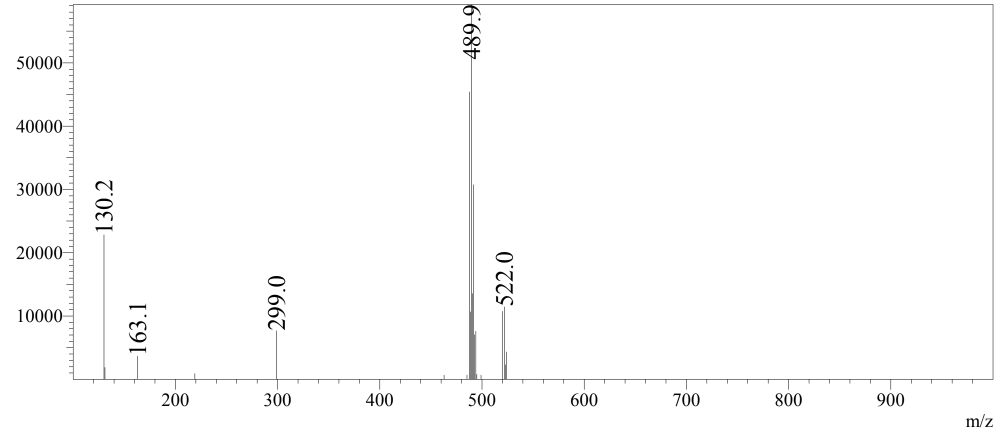

**Compound ID: C-31**

EW8457-583-P1A DMSO Bruker_A_400MHz

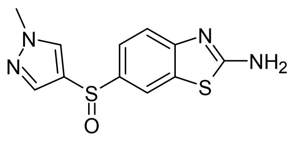


Supervisor: Tao Guo

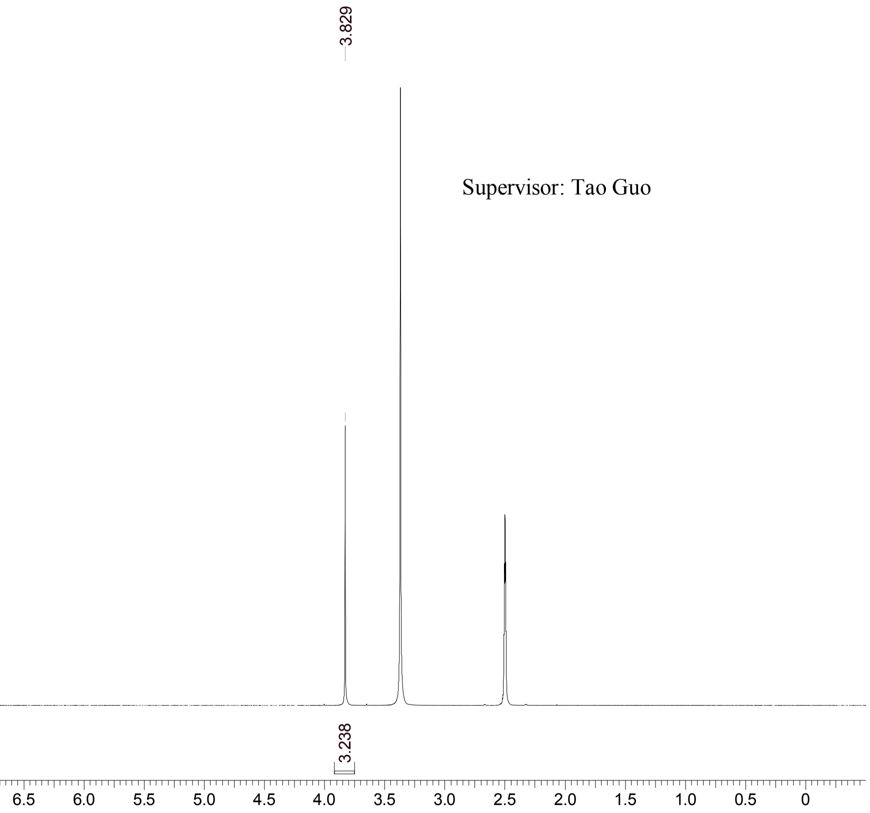


| | |
|---|---|
| Acquisition Time (sec) | 3.0671 |
| Comment | EW8457-583-P1A DMSO Bruker_A_400MHz |
| Date | 28 Aug 2018 08:49:06 |
| Frequency (MHz) | 400.1300 |
| Nucleus | 1H |
| Number of Transients | 8 |
| Origin | spect |
| Original Points Count | 24576 |
| Owner | nmrsu |
| Points Count | 65536 |
| Pulse Sequence | zg30 |
| Receiver Gain | 112.05 |
| SW(cyclical) (Hz) | 8012.82 |
| Solvent | DMSO-d6 |
| Spectrum Offset (Hz) | 2397.1384 |
| Spectrum Type | standard |
| Sweep Width (Hz) | 8012.70 |
| Temperature (degree C) | 25.144 |

Operator:

Date:

Confidential, for research only not for regulatory filing

Confidential, for research only not for regulatory filing

### LCMS REPORT

Compound ID : C-31
Sample ID : EW8457-583-P1C
Injection Vol : 0.5ul
Location : vial94
Acq Method : d:\method\5-95CD_R_220&254_POS_50.lcm
Org DataFile : D:\DATA\1808\180828\EW8457-583-P1C.lcd
Injection Date : 2018-08-28 11:14:06
Instrument :LCMS-O 15-105

Chromatogram

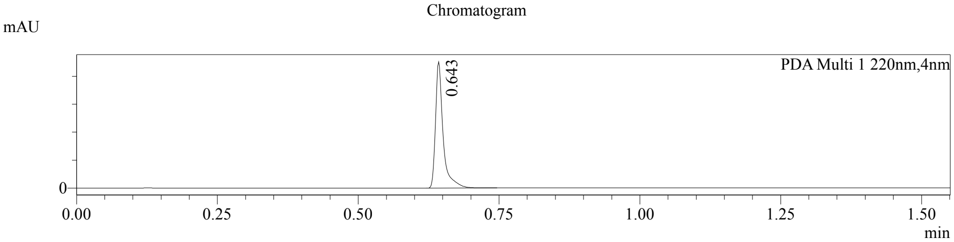


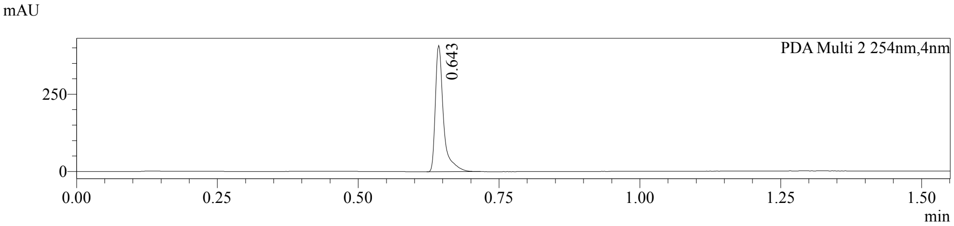


1 PDA Multi 1 / 220nm,4nm
2 PDA Multi 2 / 254nm,4nm

MS Chromatogram

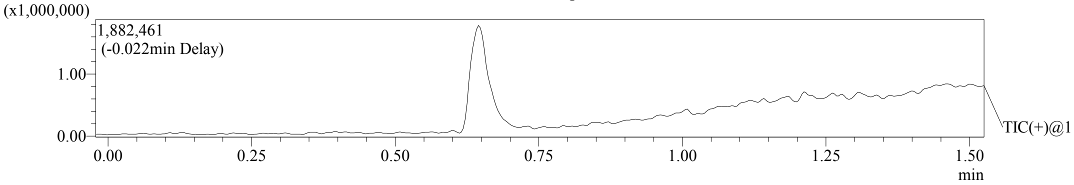


Integration Result

Peak Table

PDA Ch1 220nm

| Peak# | Ret. Time | Height | Height% | USP Width | Area | Area% |
|---|---|---|---|---|---|---|
| 1 | 0.643 | 2259314 | 100.000 | 0.025 | 2086155 | 100.000 |

Peak Table

PDA Ch2 254nm

| Peak# | Ret. Time | Height | Height% | USP Width | Area | Area% |
|---|---|---|---|---|---|---|
| 1 | 0.643 | 409018 | 100.000 | 0.026 | 405501 | 100.000 |

Confidential, for research only not for regulatory filing

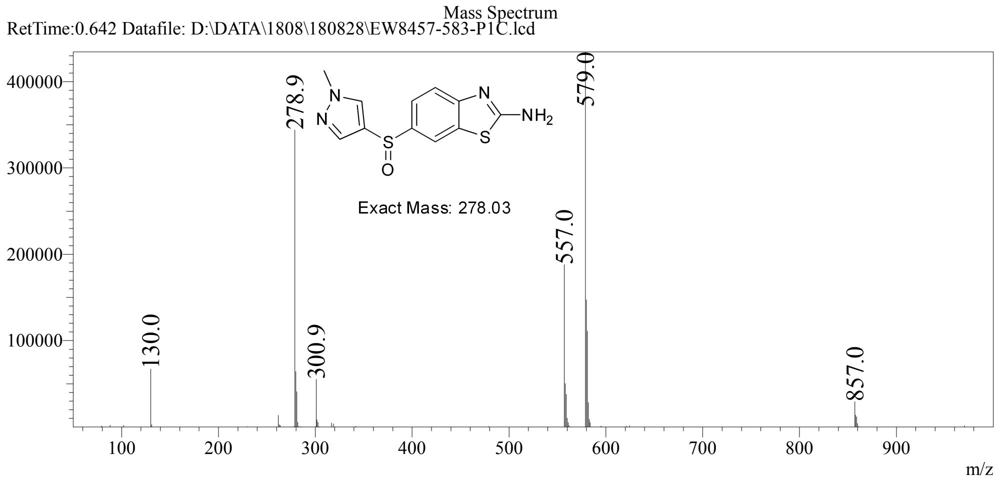

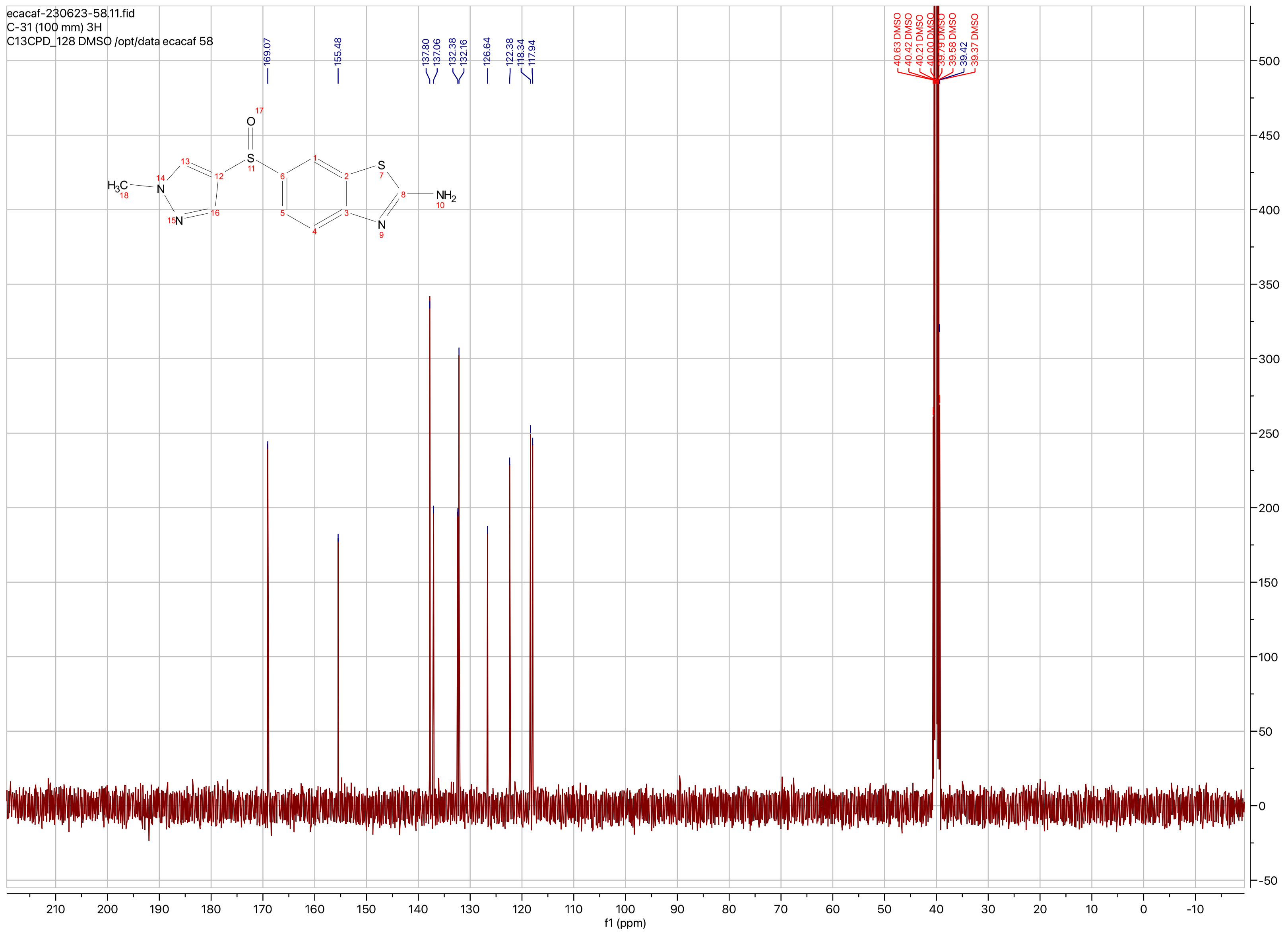

ecacaf-230623-58.11.fid
C-31 (100 mm) 3H
C13CPD_128 DMSO /opt/data ecacaf 58
169.07
155.48
137.80
137.06
132.38
132.16
126.64
122.38
118.34
117.94
40.63 DMSO
40.42 DMSO
40.21 DMSO
40.00 DMSO
39.79 DMSO
39.58 DMSO
39.42
39.37 DMSO
$H_3C$
$NH_2$
f1 (ppm)

**Compound ID: C-32**

EW8457-539-P1A DMSO Bruker_A_400MHz

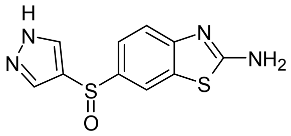

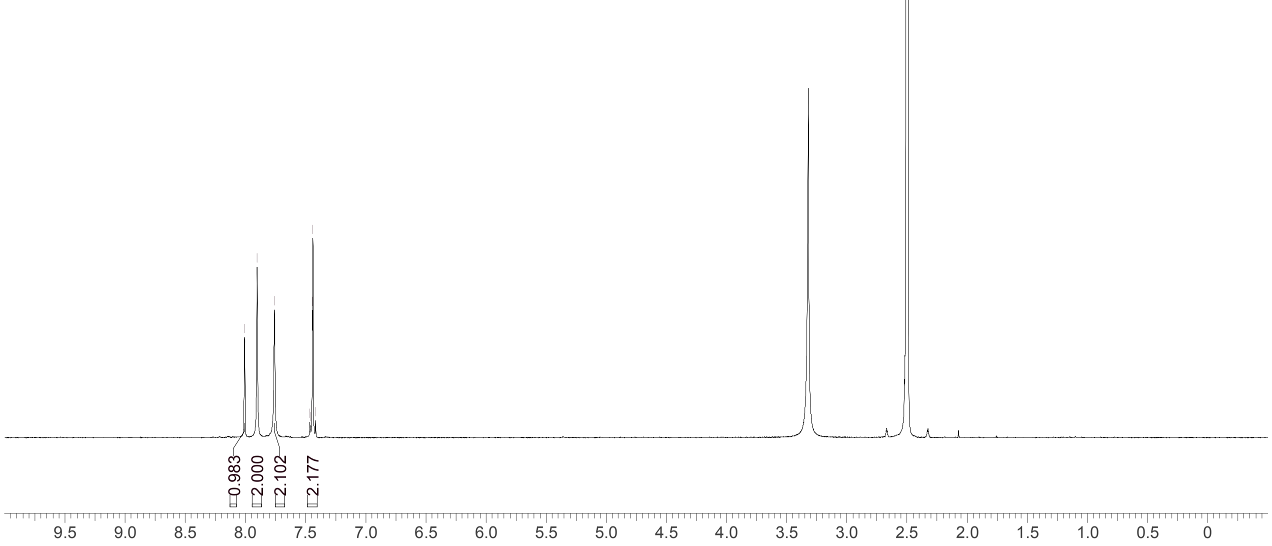


Supervisor: Tao Guo

| | |
|---|---|
| Acquisition Time (sec) | 3.0671 |
| Comment | EW8457-539-P1A DMSO Bruker_A_400MHz |
| Date | 08 Aug 2018 08:24:49 |
| Frequency (MHz) | 400.1300 |
| Nucleus | 1H |
| Number of Transients | 8 |
| Origin | spect |
| Original Points Count | 24576 |
| Owner | nmrsu |
| Points Count | 65536 |
| Pulse Sequence | zg30 |
| Receiver Gain | 175.36 |
| SW(cyclical) (Hz) | 8012.82 |
| Solvent | DMSO-d6 |
| Spectrum Offset (Hz) | 2396.4182 |
| Spectrum Type | standard |
| Sweep Width (Hz) | 8012.70 |
| Temperature (degree C) | 26.153 |

Operator: Date:

Confidential, for research only not for regulatory filing

Confidential, for research only not for regulatory filing

### LCMS REPORT

Compound ID : C-32
Sample ID : EW8457-539-P1B
Injection Vol : 3ul
Location : vial17
Acq Method : d:\method\5-95CD_R_220&254_POS.lcm
Org DataFile : D:\DATA\1808\180808\EW8457-539-P1B.lcd
Injection Date : 08/08/2018 08:07:08
Instrument :LCMS-K 15-105

Chromatogram

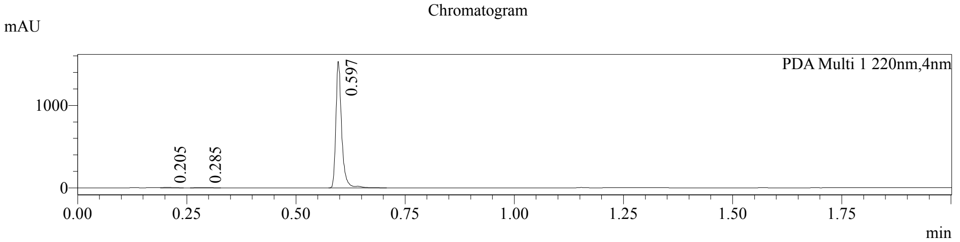


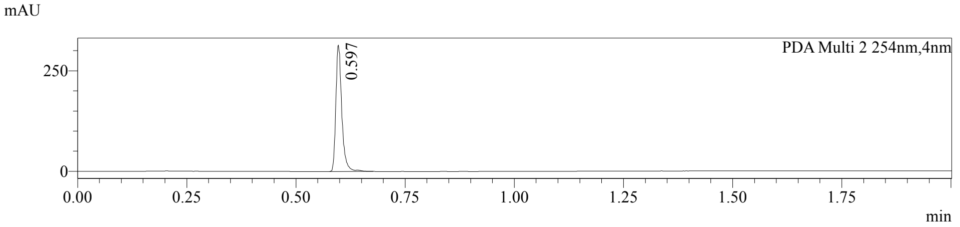


1 PDA Multi 1 / 220nm,4nm
2 PDA Multi 2 / 254nm,4nm

MS Chromatogram

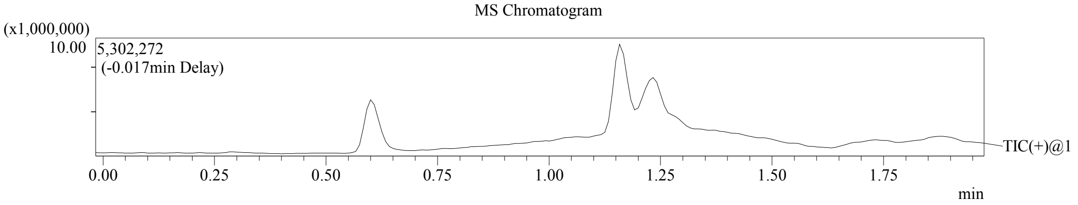


Integration Result

Peak Table

PDA Ch1 220nm

| Peak# | Ret. Time | Height | Height% | USP Width | Area | Area% |
|---|---|---|---|---|---|---|
| 1 | 0.205 | 5601 | 0.363 | 0.037 | 7514 | 0.531 |
| 2 | 0.285 | 3287 | 0.213 | 0.049 | 5913 | 0.418 |
| 3 | 0.597 | 1532561 | 99.423 | 0.025 | 1401970 | 99.051 |

Peak Table

PDA Ch2 254nm

| Peak# | Ret. Time | Height | Height% | USP Width | Area | Area% |
|---|---|---|---|---|---|---|
| 1 | 0.597 | 314085 | 100.000 | 0.026 | 297104 | 100.000 |

Confidential, for research only not for regulatory filing

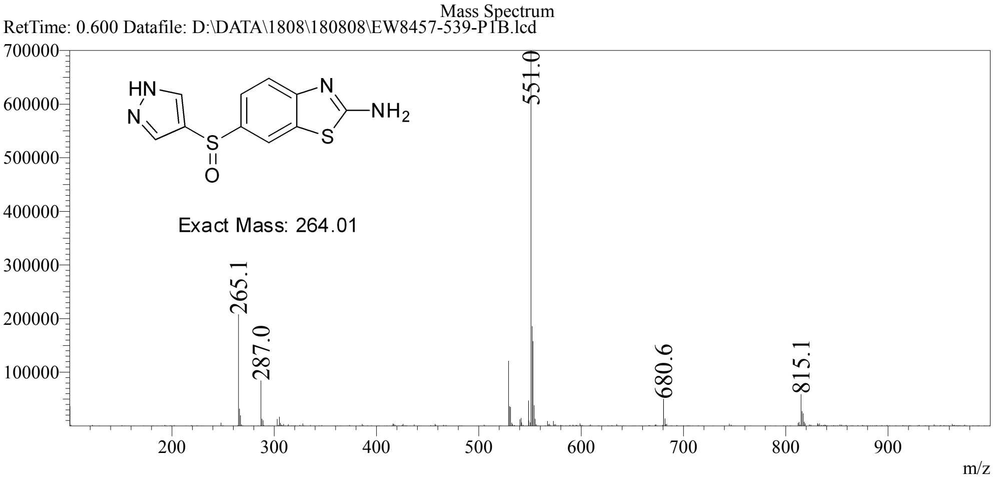

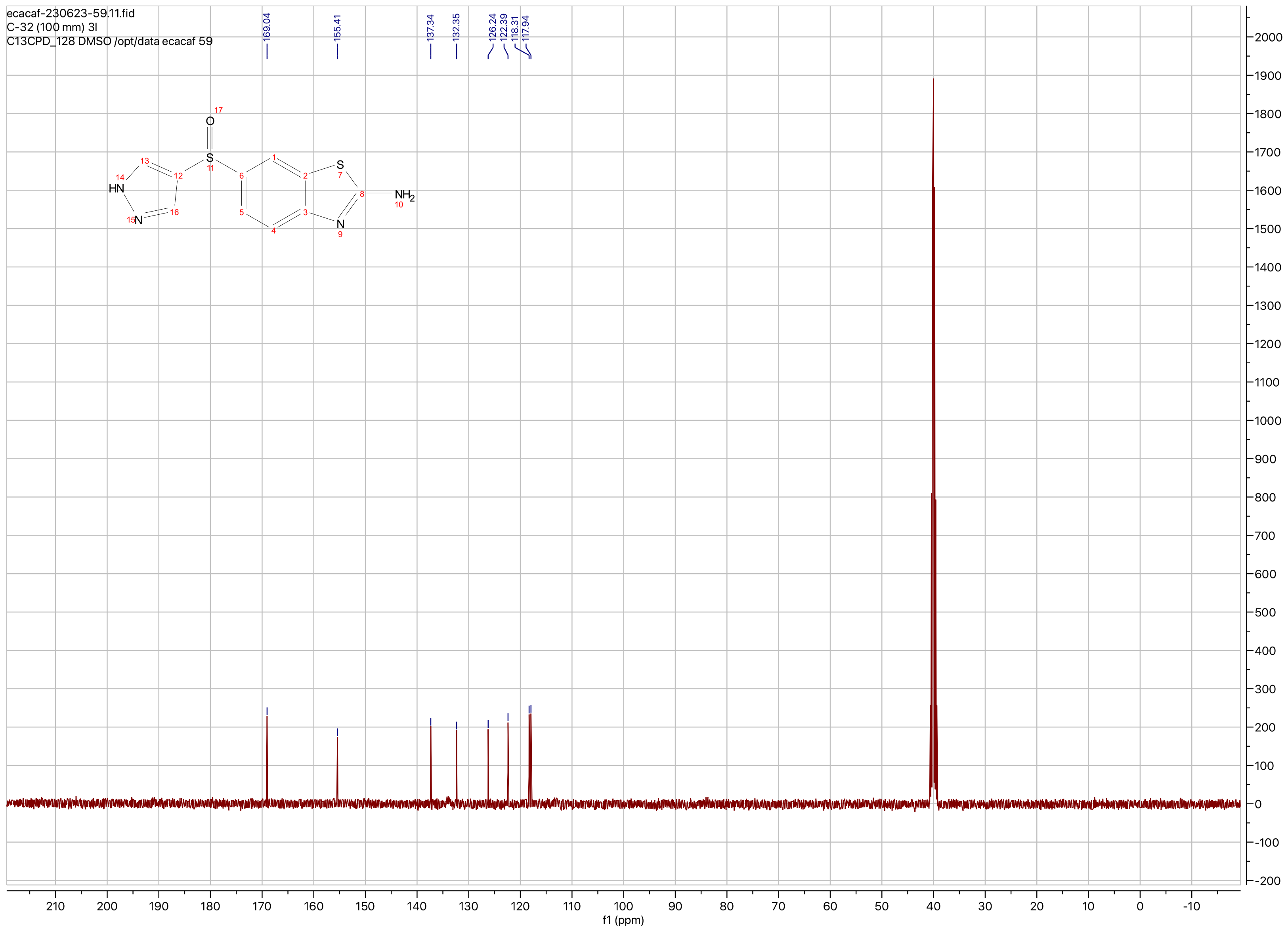

ecacaf-230623-59.11.fid
C-32 (100 mm) 3l
C13CPD_128 DMSO /opt/data ecacaf 59
169.04
155.41
137.34
132.35
126.24
122.39
118.31
117.94
O
S
HN
N
NH2
f1 (ppm)

**Compound ID: C-33**

EW8457-579-P1A DMSO Bruker_E_400MHz

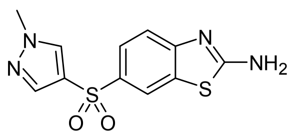


Supervisor: Tao Guo

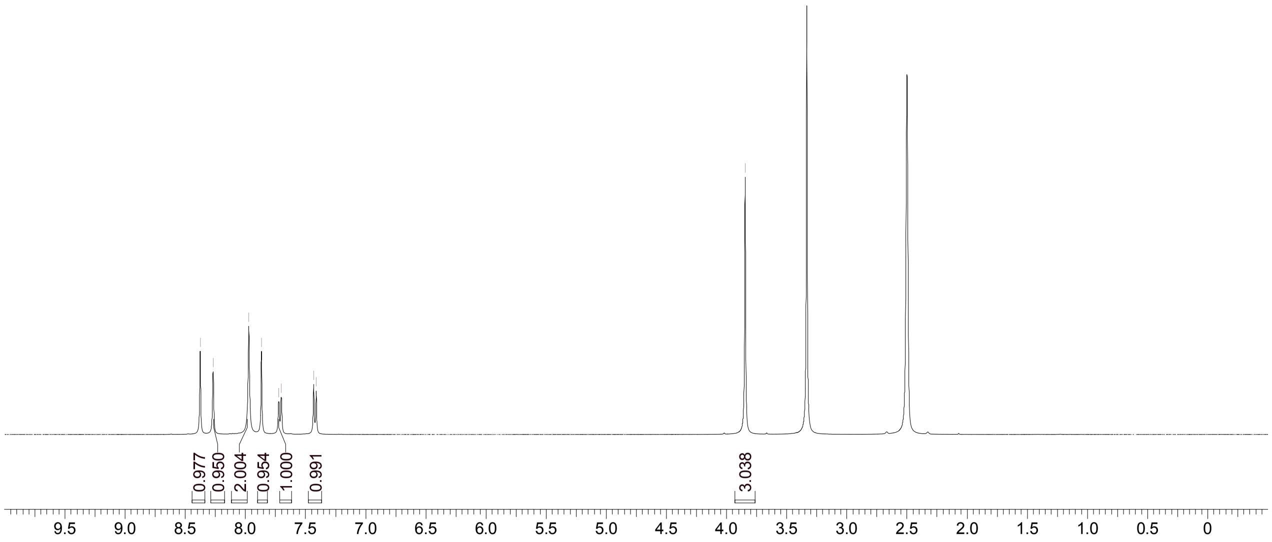


| | |
|---|---|
| Acquisition Time (sec) | 3.0671 |
| Comment | EW8457-579-P1A DMSO Bruker_E_400MHz |
| Date | 27 Aug 2018 08:22:08 |
| Frequency (MHz) | 400.1500 |
| Nucleus | 1H |
| Number of Transients | 8 |
| Origin | spect |
| Original Points Count | 24576 |
| Owner | nmrsu |
| Points Count | 65536 |
| Pulse Sequence | zg30 |
| Receiver Gain | 90.22 |
| SW(cyclical) (Hz) | 8012.82 |
| Solvent | DMSO-d6 |
| Spectrum Offset (Hz) | 2397.6985 |
| Spectrum Type | standard |
| Sweep Width (Hz) | 8012.70 |
| Temperature (degree C) | 27.165 |

Operator:

Date:

Confidential, for research only not for regulatory filing

Confidential, for research only not for regulatory filing

### LCMS REPORT

Compound ID : C-33
Sample ID : EW8457-579-P1B
Injection Vol : 2ul
Location : vial67
Acq Method : d:\method\5-95AB_R_220&254_50.lcm
Org DataFile : D:\DATA\1808\180827\EW8457-579-P1B.lcd
Injection Date : 2018-08-27 8:36:04
Instrument : LCMS-Y 15-105

Chromatogram

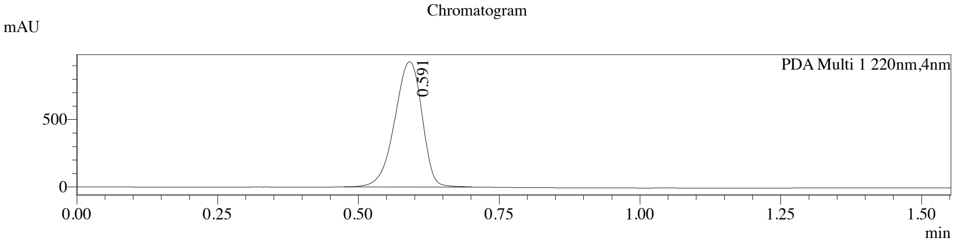


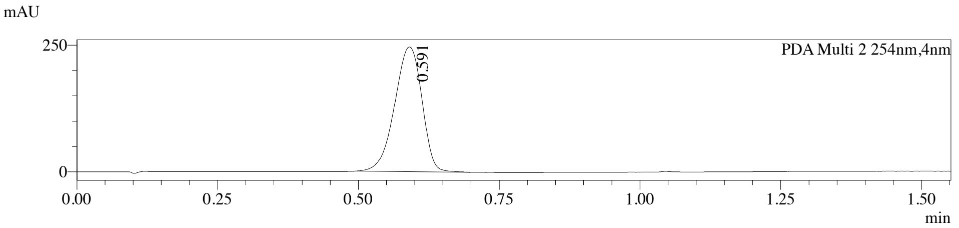


1 PDA Multi 1 / 220nm,4nm
2 PDA Multi 2 / 254nm,4nm

MS Chromatogram

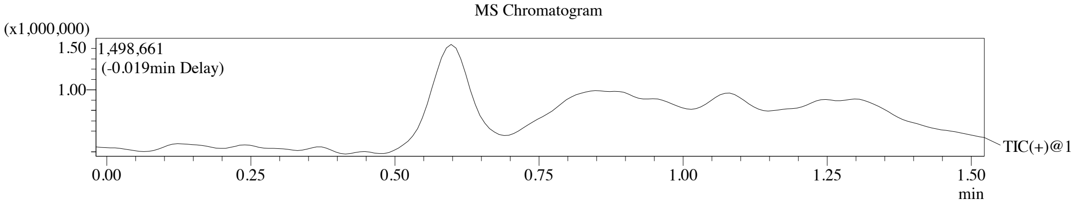


Integration Result

Peak Table

PDA Ch1 220nm

| Peak# | Ret. Time | Height | Height% | USP Width | Area | Area% |
|---|---|---|---|---|---|---|
| 1 | 0.591 | 929512 | 100.000 | 0.088 | 3149294 | 100.000 |

Peak Table

PDA Ch2 254nm

| Peak# | Ret. Time | Height | Height% | USP Width | Area | Area% |
|---|---|---|---|---|---|---|
| 1 | 0.591 | 246462 | 100.000 | 0.088 | 830297 | 100.000 |

Confidential, for research only not for regulatory filing

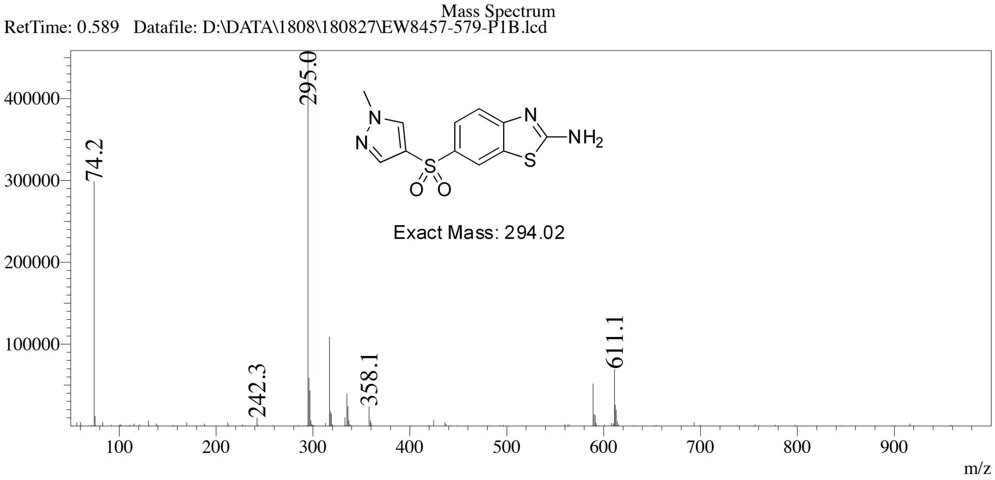

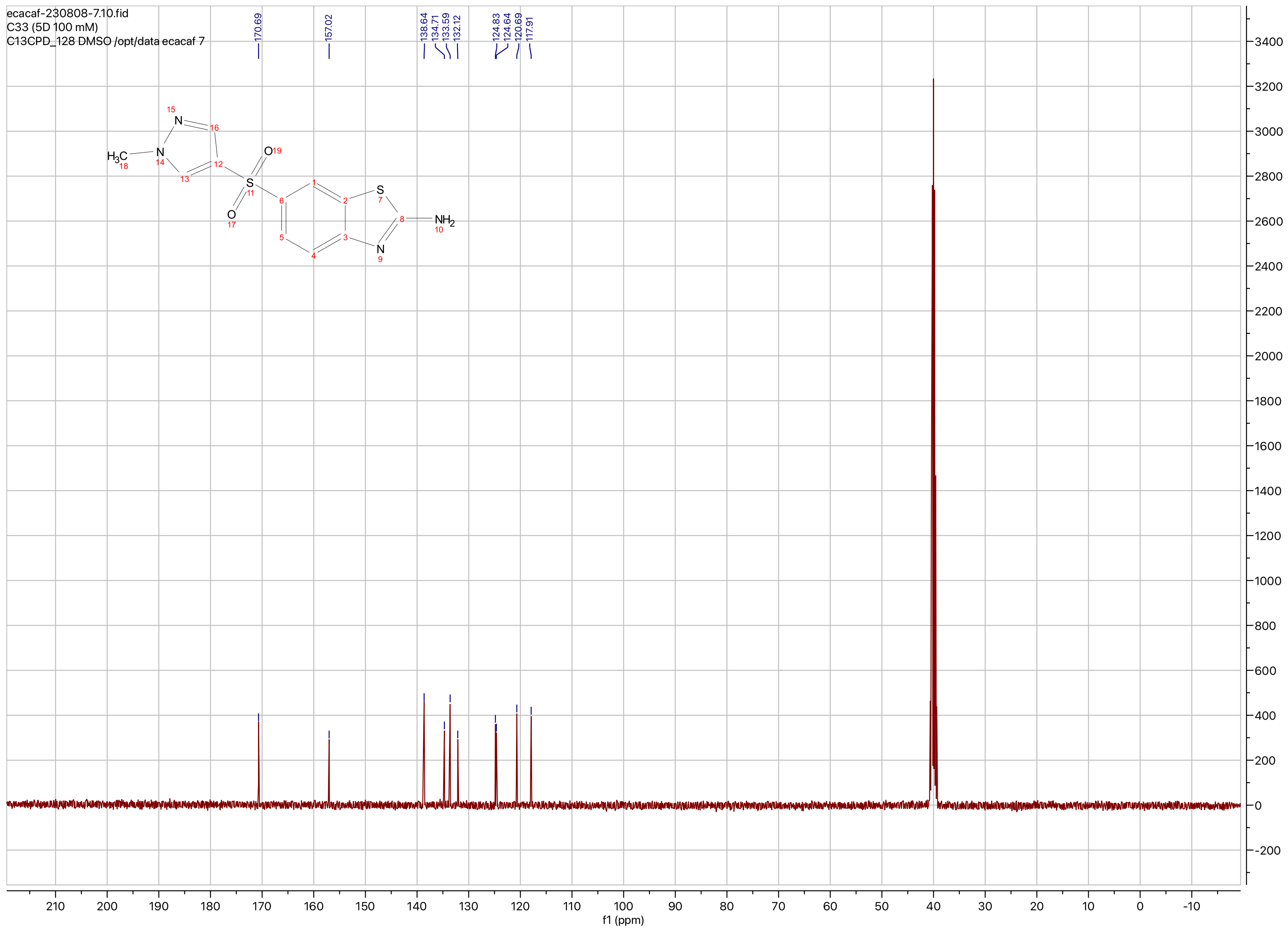

ecacaf-230808-7.10.fid
C33 (5D 100 mM)
C13CPD_128 DMSO /opt/data ecacaf 7
170.69
157.02
138.64
134.71
133.59
132.12
124.83
124.64
120.69
117.91
$H_3C$
N
N
S
O
O
S
N
$NH_2$
f1 (ppm)

**Compound ID: C-34**

EW8457-538-P1A DMSO Bruker_A_400MHz

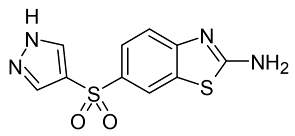

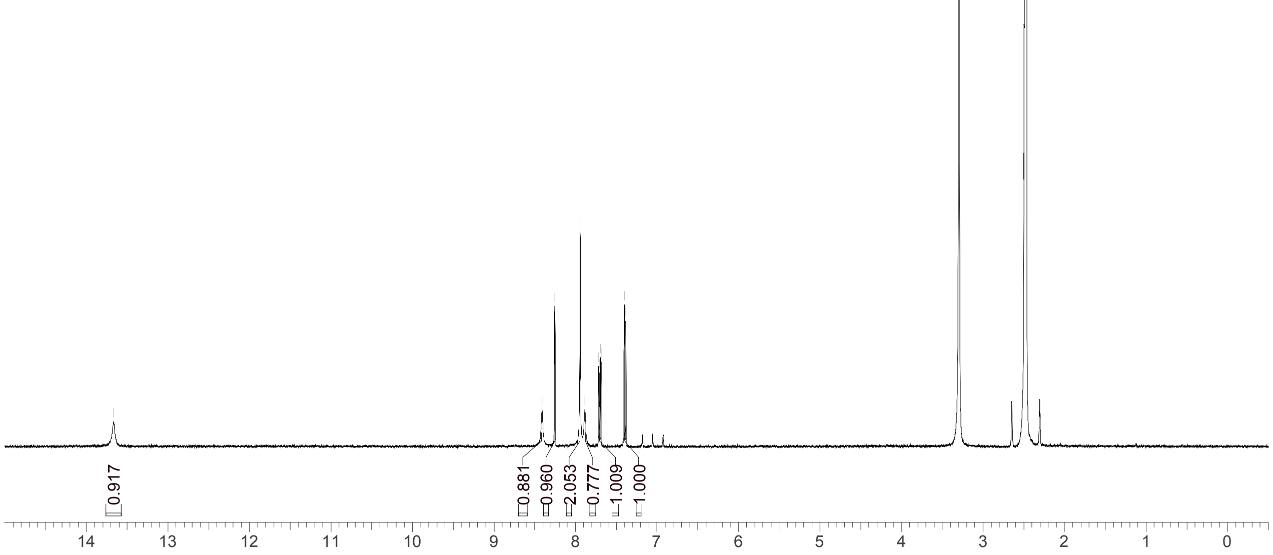


Supervisor: Tao Guo

| | |
|---|---|
| Acquisition Time (sec) | 3.0671 |
| Comment | EW8457-538-P1A DMSO Bruker_A_400MHz |
| Date | 08 Aug 2018 08:22:32 |
| Frequency (MHz) | 400.1300 |
| Nucleus | 1H |
| Number of Transients | 8 |
| Origin | spect |
| Original Points Count | 24576 |
| Owner | nmrsu |
| Points Count | 65536 |
| Pulse Sequence | zg30 |
| Receiver Gain | 175.36 |
| SW(cyclical) (Hz) | 8012.82 |
| Solvent | DMSO-d6 |
| Spectrum Offset (Hz) | 2387.4951 |
| Spectrum Type | standard |
| Sweep Width (Hz) | 8012.70 |
| Temperature (degree C) | 26.150 |

Operator:

Date:

Confidential, for research only not for regulatory filing

### LCMS REPORT

Compound ID : C-34
Sample ID : EW8457-538-P1B
Injection Vol : 3ul
Location : vial16
Acq Method : d:\method\5-95CD_R_220&254_POS.lcm
Org DataFile : D:\DATA\1808\180808\EW8457-538-P1B.lcd
Injection Date : 08/08/2018 08:04:35
Instrument :LCMS-K 15-105

Chromatogram

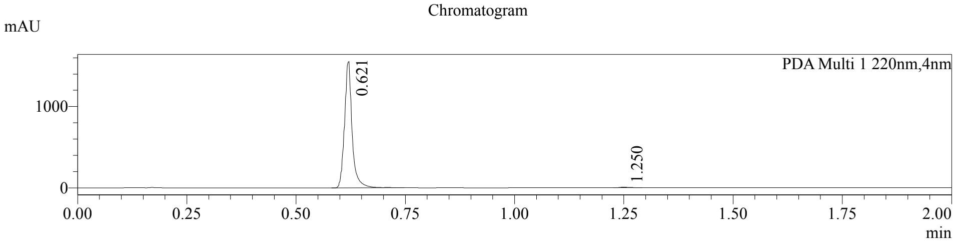


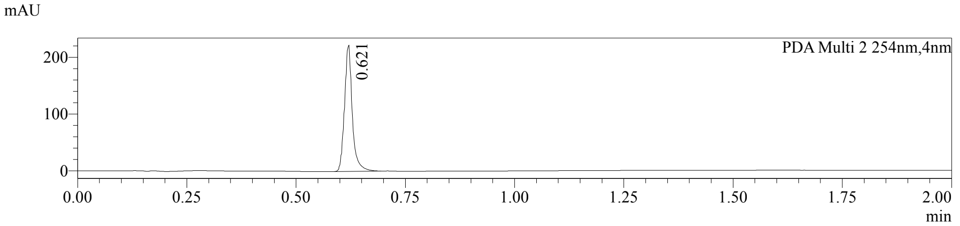


1 PDA Multi 1 / 220nm,4nm
2 PDA Multi 2 / 254nm,4nm

MS Chromatogram

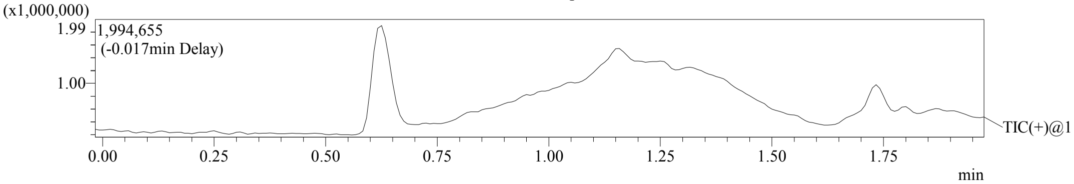


Integration Result

Peak Table

PDA Ch1 220nm

| Peak# | Ret. Time | Height | Height% | USP Width | Area | Area% |
|---|---|---|---|---|---|---|
| 1 | 0.621 | 1554268 | 99.621 | 0.031 | 1873811 | 99.549 |
| 2 | 1.250 | 5909 | 0.379 | 0.037 | 8493 | 0.451 |

Peak Table

PDA Ch2 254nm

| Peak# | Ret. Time | Height | Height% | USP Width | Area | Area% |
|---|---|---|---|---|---|---|
| 1 | 0.621 | 222304 | 100.000 | 0.033 | 274646 | 100.000 |

Confidential, for research only not for regulatory filing

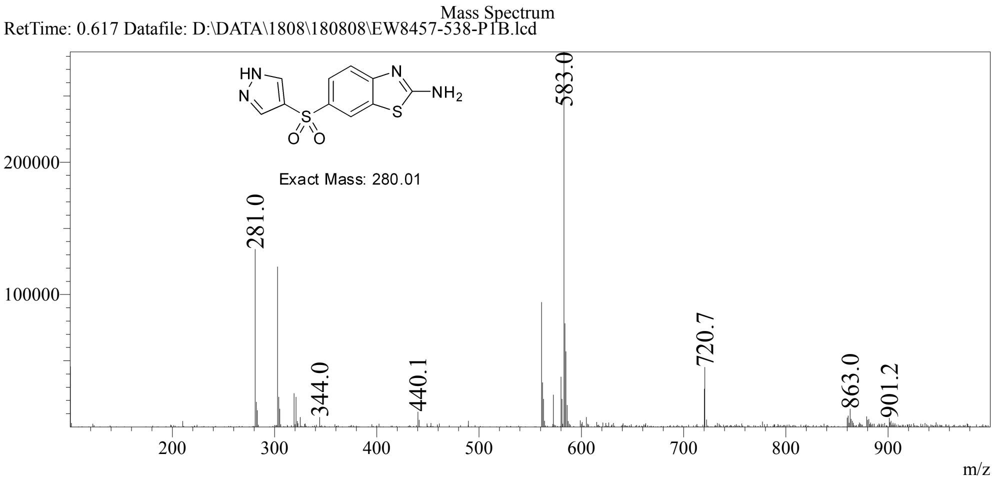

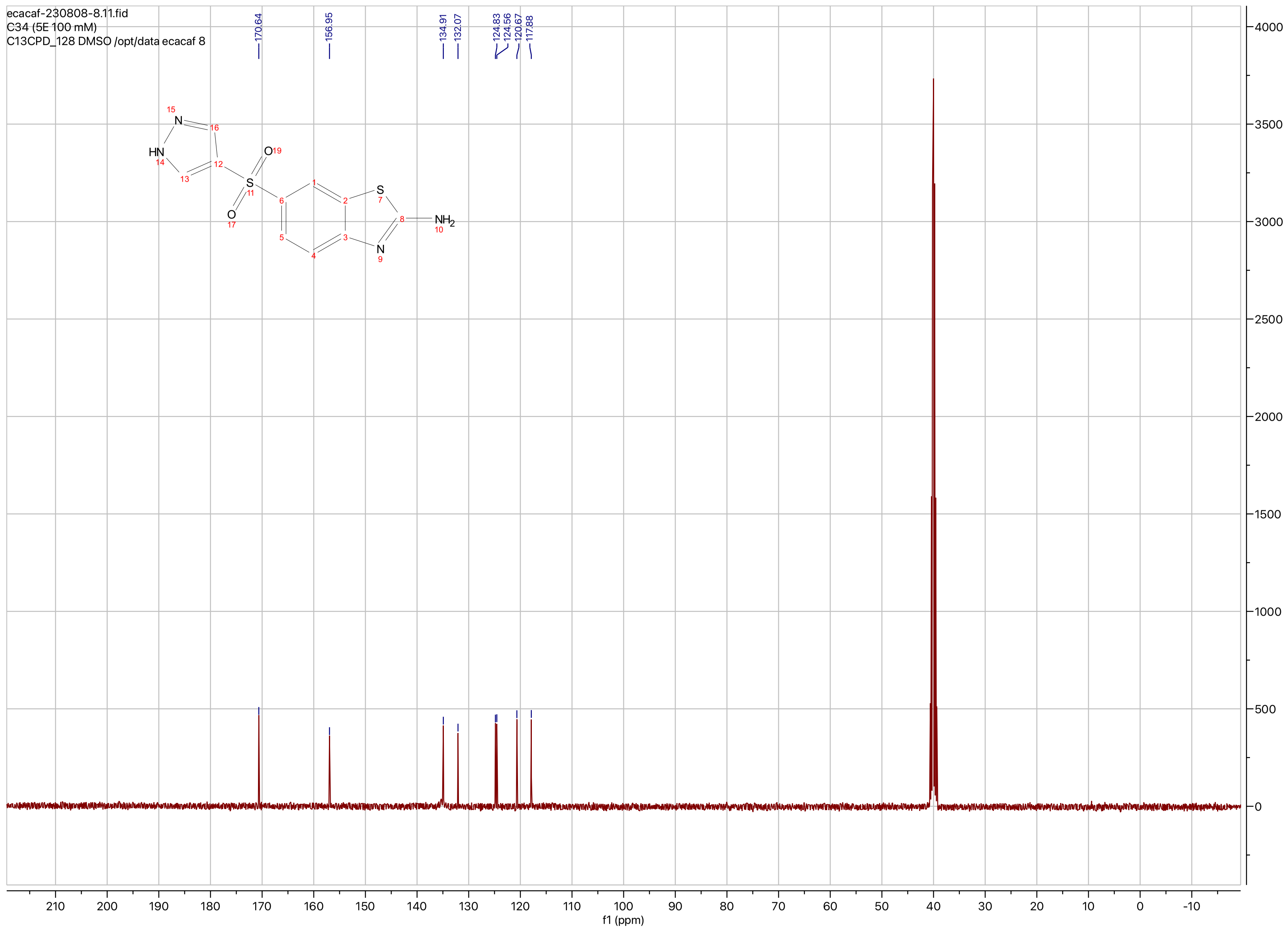

ecacaf-230808-8.11.fid
C34 (5E 100 mM)
C13CPD_128 DMSO /opt/data ecacaf 8
170.64
156.95
134.91
132.07
124.83
124.56
120.67
117.88
f1 (ppm)

**Compound ID: C-35**

EW8457-867-P1A DMSO Bruker_H_400MHz

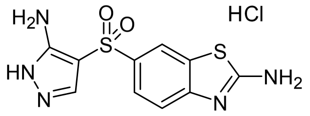


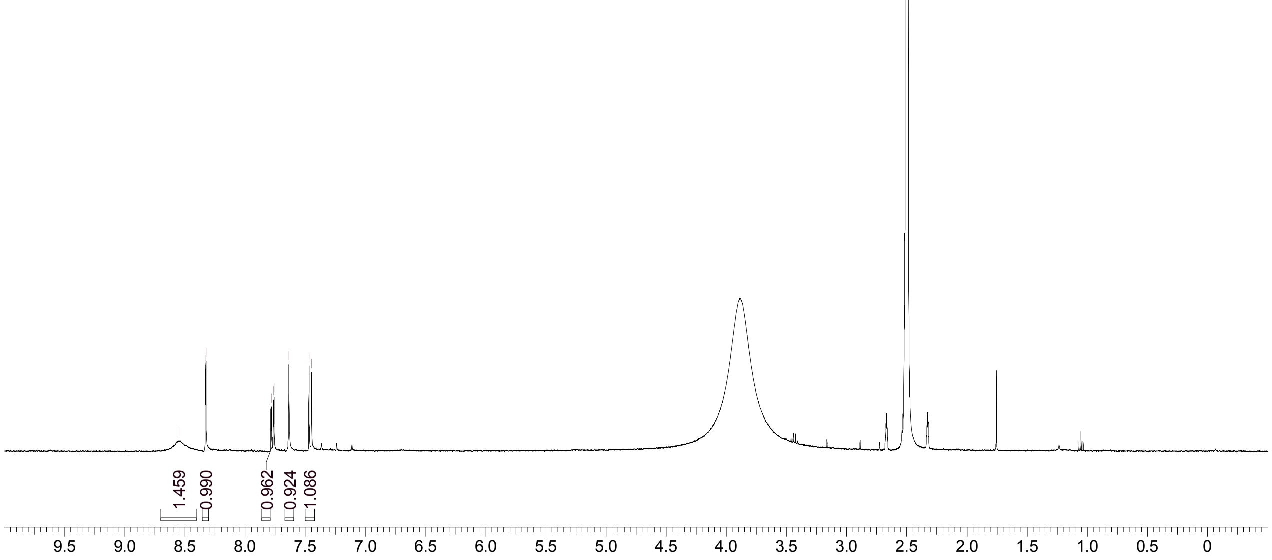


Supervisor: Tao Guo

| | |
|---|---|
| Acquisition Time (sec) | 3.9977 |
| Comment | EW8457-867-P1A DMSO Bruker_H_400MHz |
| Date | 24 Jan 2019 00:47:54 |
| Frequency (MHz) | 400.1300 |
| Nucleus | 1H |
| Number of Transients | 8 |
| Origin | Avance |
| Original Points Count | 32768 |
| Owner | nmrsu |
| Points Count | 65536 |
| Pulse Sequence | zg30 |
| Receiver Gain | 32.00 |
| SW(cyclical) (Hz) | 8196.72 |
| Solvent | DMSO-d6 |
| Spectrum Offset (Hz) | 2467.2405 |
| Spectrum Type | standard |
| Sweep Width (Hz) | 8196.60 |
| Temperature (degree C) | 23.710 |

Operator:

Date:

Confidential, for research only not for regulatory filing

Confidential, for research only not for regulatory filing

### LCMS REPORT

Compound ID : C-35
Sample ID : EW8457-867-P1B
Injection Vol : 5ul
Location : vial45
Acq Method : d:\method\0-60AB_0_R_220&254.lcm
Org DataFile : D:\DATA\1901\190123\EW8457-867-P1B.lcd
Injection Date : 23/01/2019 14:21:34
Instrument : LCMS-AA 15-105

Chromatogram

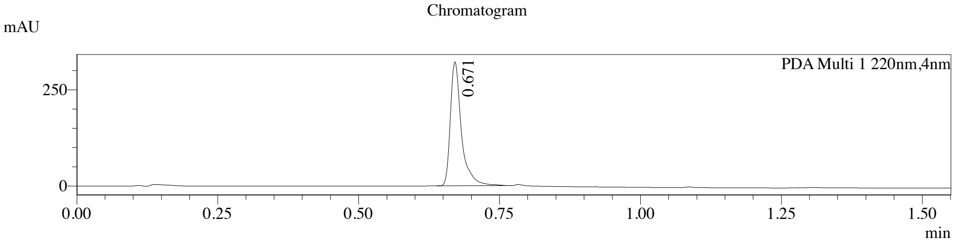


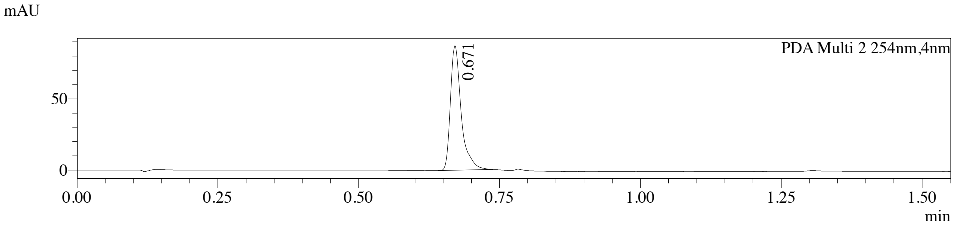


1 PDA Multi 1 / 220nm,4nm
2 PDA Multi 2 / 254nm,4nm

MS Chromatogram

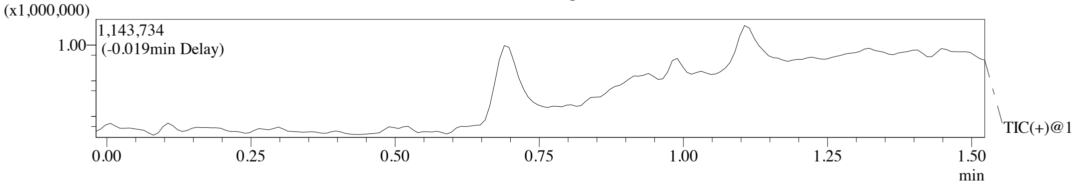


Integration Result

Peak Table

PDA Ch1 220nm

| Peak# | Ret. Time | Height | Height% | USP Width | Area | Area% |
|---|---|---|---|---|---|---|
| 1 | 0.671 | 322125 | 100.000 | 0.034 | 437401 | 100.000 |

Peak Table

PDA Ch2 254nm

| Peak# | Ret. Time | Height | Height% | USP Width | Area | Area% |
|---|---|---|---|---|---|---|
| 1 | 0.671 | 87501 | 100.000 | 0.034 | 116140 | 100.000 |

Confidential, for research only not for regulatory filing

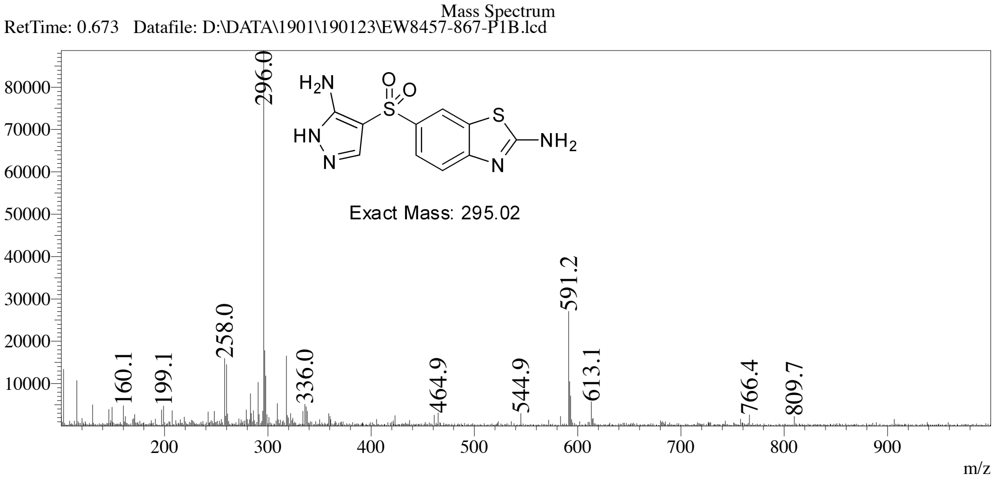

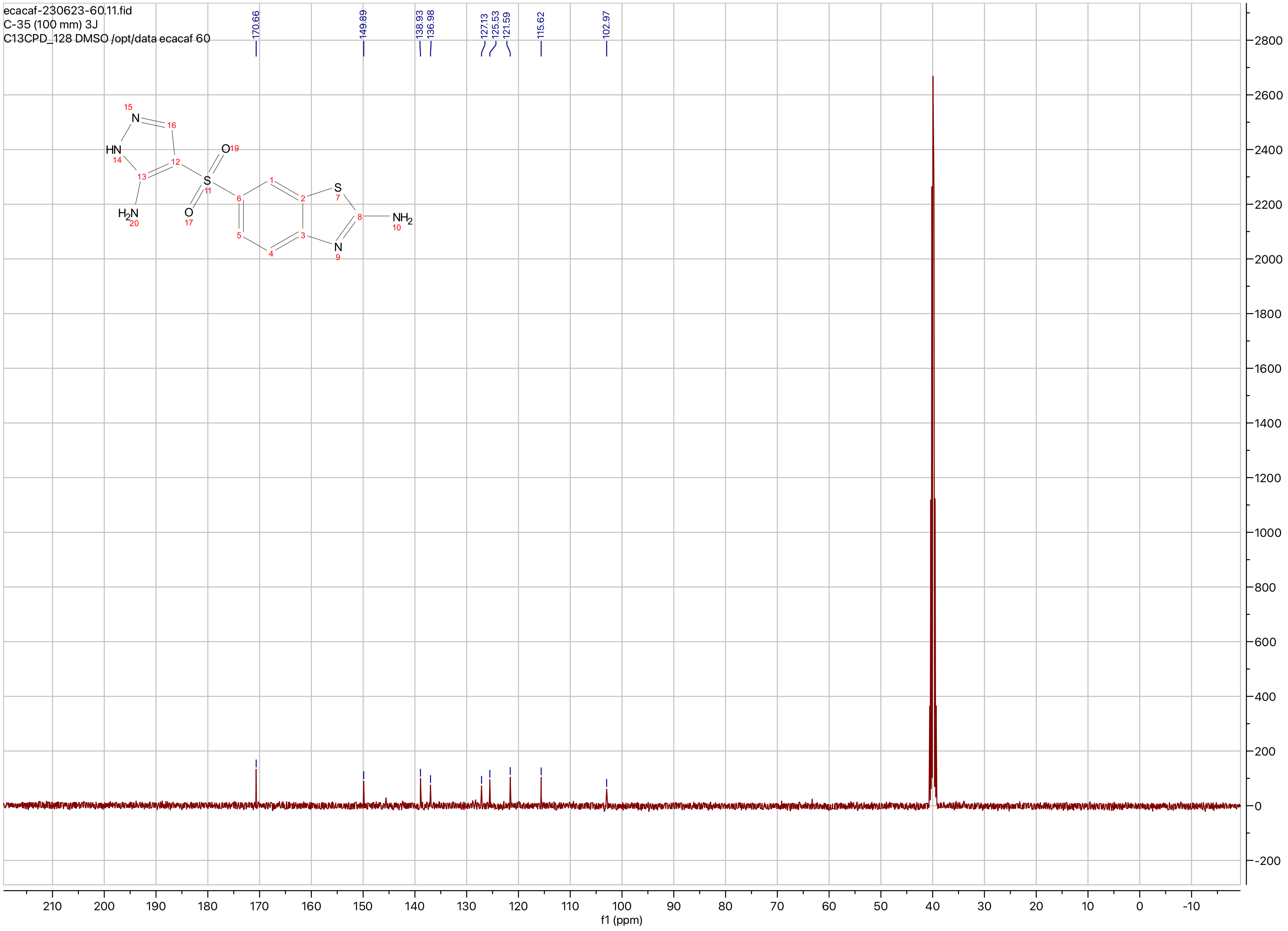
ecacaf-230623-60.11.fid
C-35 (100 mm) 3J
C13CPD_128 DMSO /opt/data ecacaf 60
170.66
149.89
138.93
136.98
127.13
125.53
121.59
115.62
102.97
N
HN
O
S
H2N
O
S
NH2
N
f1 (ppm)

**Compound ID: C-36**

ET27821-95-P1H1 DMSO Varian_Y_400MHz

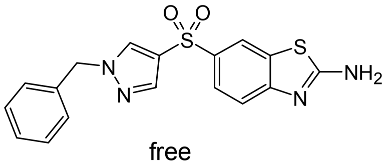


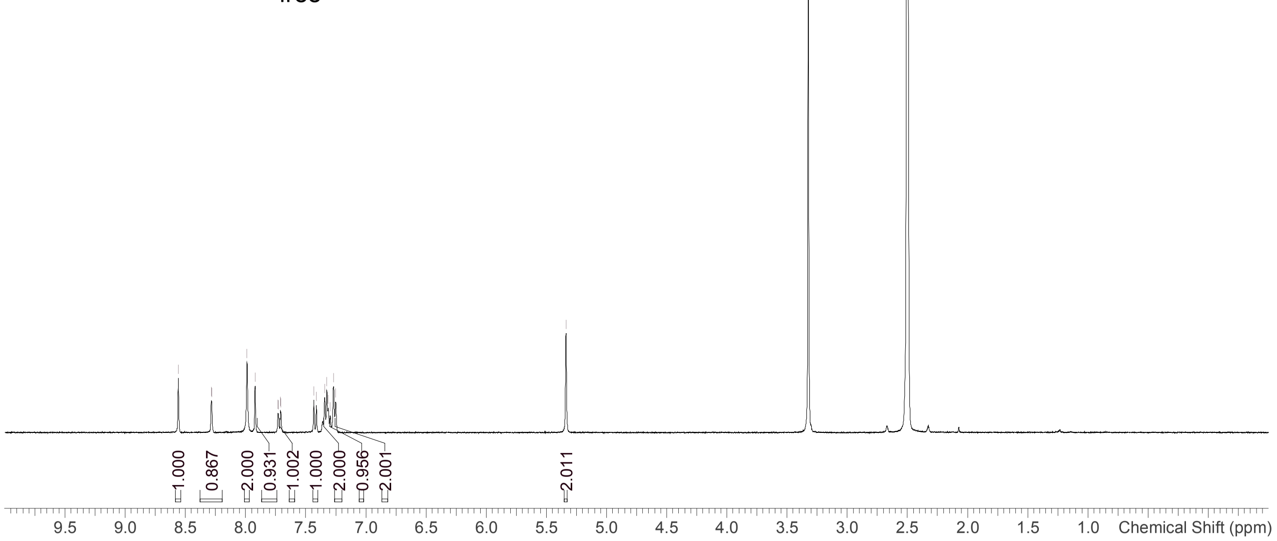


Supervisor: Tao Guo

| | |
|---|---|
| Acquisition Time (sec) | 2.0486 |
| Comment | ET27821-95-P1H1 DMSO Varian_Y_400MHz |
| Date | Jun 27 2019 |
| Frequency (MHz) | 399.6752 |
| Nucleus | 1H |
| Number of Transients | 8 |
| Original Points Count | 14802 |
| Points Count | 32768 |
| Pulse Sequence | s2pul |
| Receiver Gain | 42.00 |
| SW(cyclical) (Hz) | 7225.43 |
| Solvent | DMSO-d6 |
| Spectrum Offset (Hz) | 2807.8633 |
| Spectrum Type | standard |
| Sweep Width (Hz) | 7225.21 |
| Temperature (degree C) | 25.000 |

Operator:

Date:

Confidential, for research only not for regulatory filing

LCMS REPORT

->

Compound ID : C-36
Sample ID : ET27821-95-P1H
Injection Date : 27. Jun. 2019
Inj. Vol. : 3.00 ul
Location : P1-F-09
Acq Method : D:\DATA\190627-HD 5\5_95AB_6min-220.M
Data Filename : D:\DATA\190627-HD 5\ET27821-95-P1H.D
Instrument : H

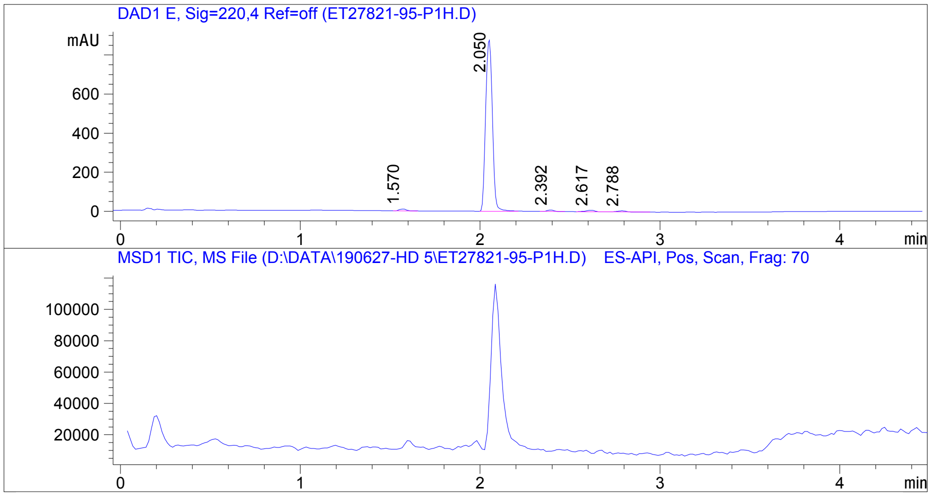


Integration Result

Signal 1 : DAD1 E, Sig=220,4 Ref=off

| Peak # | RT [min] | Area | Height | Height % | Width [min] | Area % |
|---|---|---|---|---|---|---|
| 1 | 1.570 | 25.920 | 9.490 | 1.040 | 0.044 | 1.083 |
| 2 | 2.050 | 2301.196 | 882.111 | 96.679 | 0.042 | 96.147 |
| 3 | 2.392 | 18.677 | 7.167 | 0.785 | 0.042 | 0.780 |
| 4 | 2.617 | 27.237 | 7.984 | 0.875 | 0.052 | 1.138 |
| 5 | 2.788 | 20.396 | 5.656 | 0.620 | 0.054 | 0.852 |

Operator:_____________

Date:_________________

Confidential. For research information only.

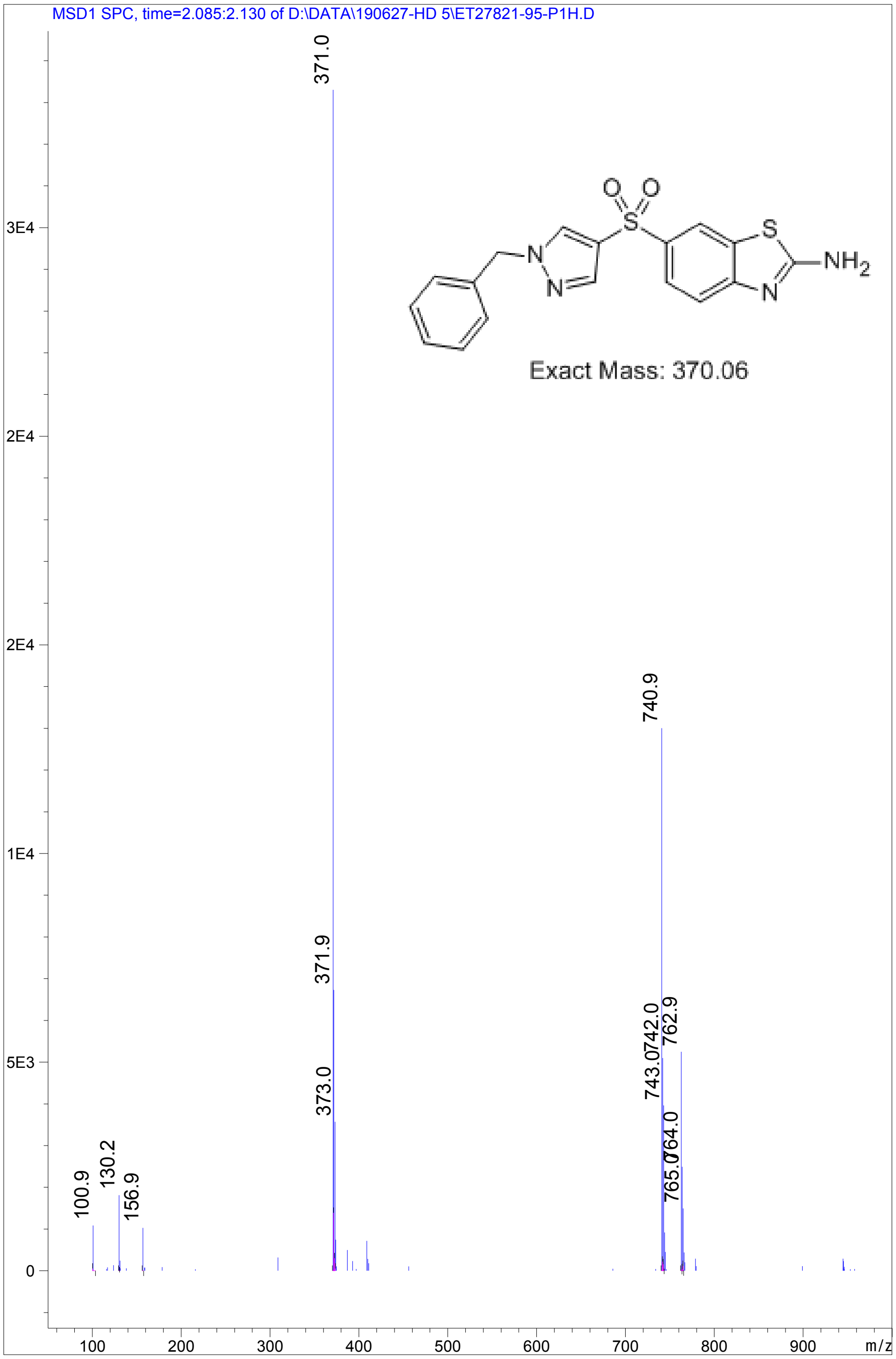


Confidential. For research information only.

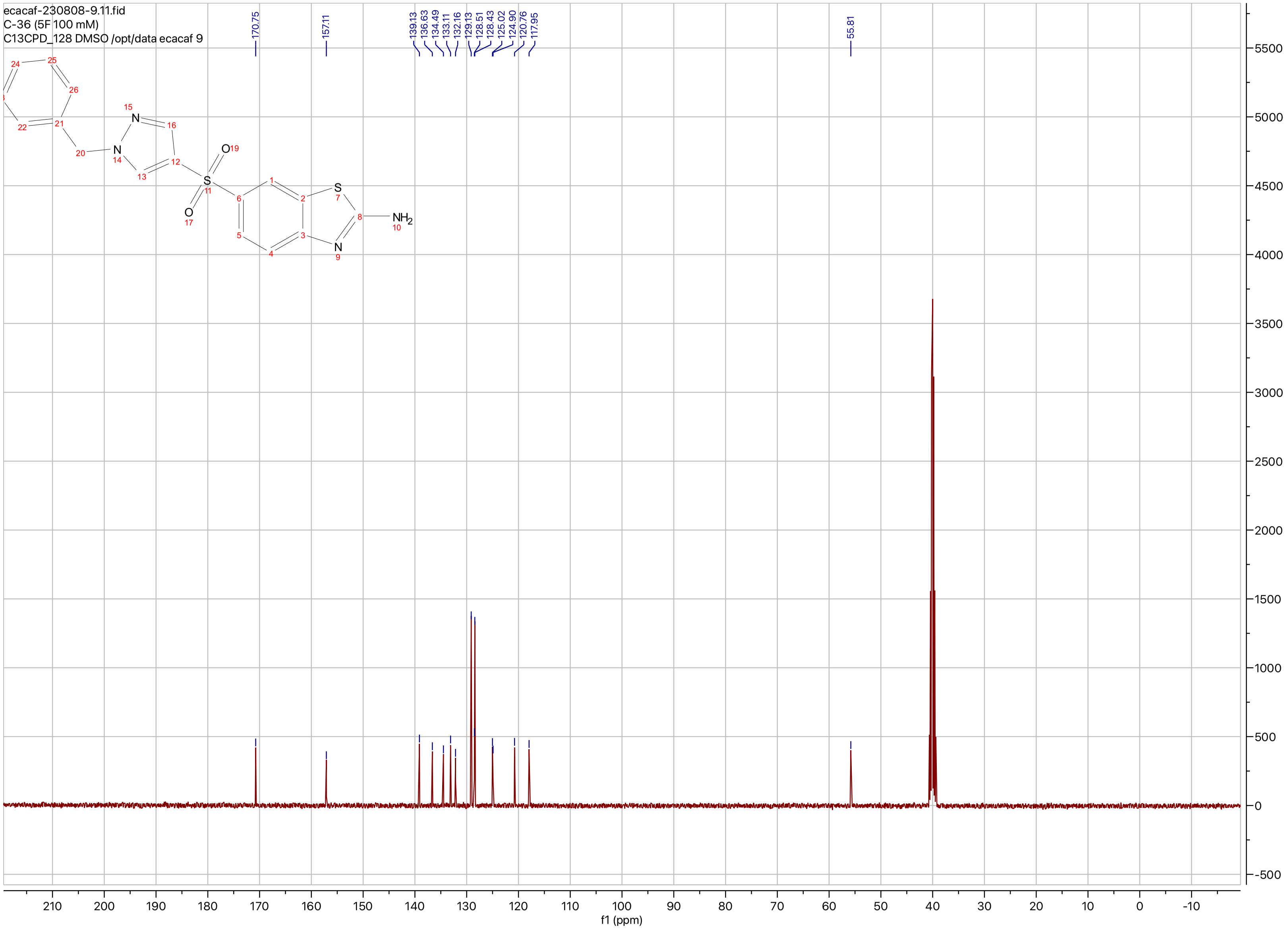

ecacaf-230808-9.11.fid
C-36 (5F 100 mM)
C13CPD_128 DMSO /opt/data ecacaf 9
170.75
157.11
139.13
136.63
134.49
133.11
132.16
129.13
128.51
128.43
125.02
124.90
120.76
117.95
55.81
NH2
f1 (ppm)

**Compound ID: C-37**

ET27821-155-P1H DMSO Bruker_K_400MHz

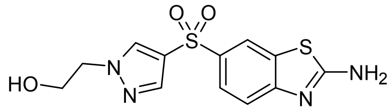


free

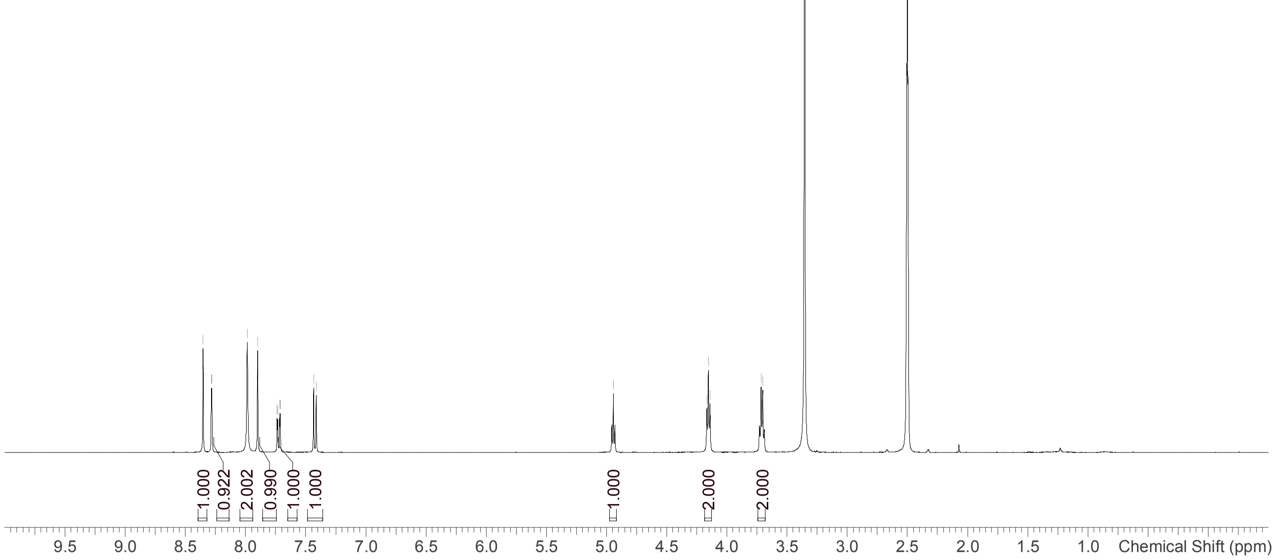


Supervisor: Tao Guo

| | |
|---|---|
| Acquisition Time (sec) | 1.9988 |
| Comment | ET27821-155-P1H DMSO Bruker_K_400MHz |
| Date | 24 Jul 2019 01:49:49 |
| Frequency (MHz) | 400.1300 |
| Nucleus | 1H |
| Number of Transients | 8 |
| Origin | Avance |
| Original Points Count | 16384 |
| Owner | nmr |
| Points Count | 65536 |
| Pulse Sequence | zg30 |
| Receiver Gain | 101.00 |
| SW(cyclical) (Hz) | 8196.72 |
| Solvent | DMSO-d6 |
| Spectrum Offset (Hz) | 2463.4790 |
| Spectrum Type | standard |
| Sweep Width (Hz) | 8196.60 |
| Temperature (degree C) | 23.349 |

Operator:

Date:

Confidential, for research only not for regulatory filing

LCMS REPORT

Compound ID : C-37 ->
Sample ID : ET27821-155-P1H
Injection Date : 24. Jul. 2019
Inj. Vol. : 0.70 ul
Location : P1-C-02
Acq Method : D:\DATA\190724-HD 7\5_95AB_6min-220.M
Data Filename : D:\DATA\190724-HD 7\ET27821-155-P1H.D
Instrument : H

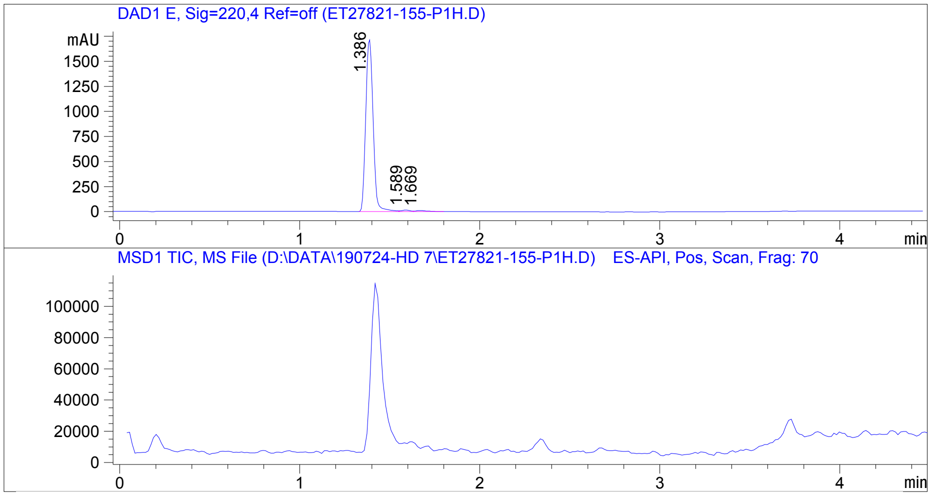


Integration Result

Signal 1 : DAD1 E, Sig=220,4 Ref=off

| Peak # | RT [min] | Area | Height | Height % | Width [min] | Area % |
|---|---|---|---|---|---|---|
| 1 | 1.386 | 4885.056 | 1721.853 | 98.607 | 0.045 | 98.135 |
| 2 | 1.589 | 47.374 | 14.166 | 0.811 | 0.051 | 0.952 |
| 3 | 1.669 | 45.457 | 10.154 | 0.582 | 0.064 | 0.913 |

Operator:_____________

Date:_________________

Confidential. For research information only.

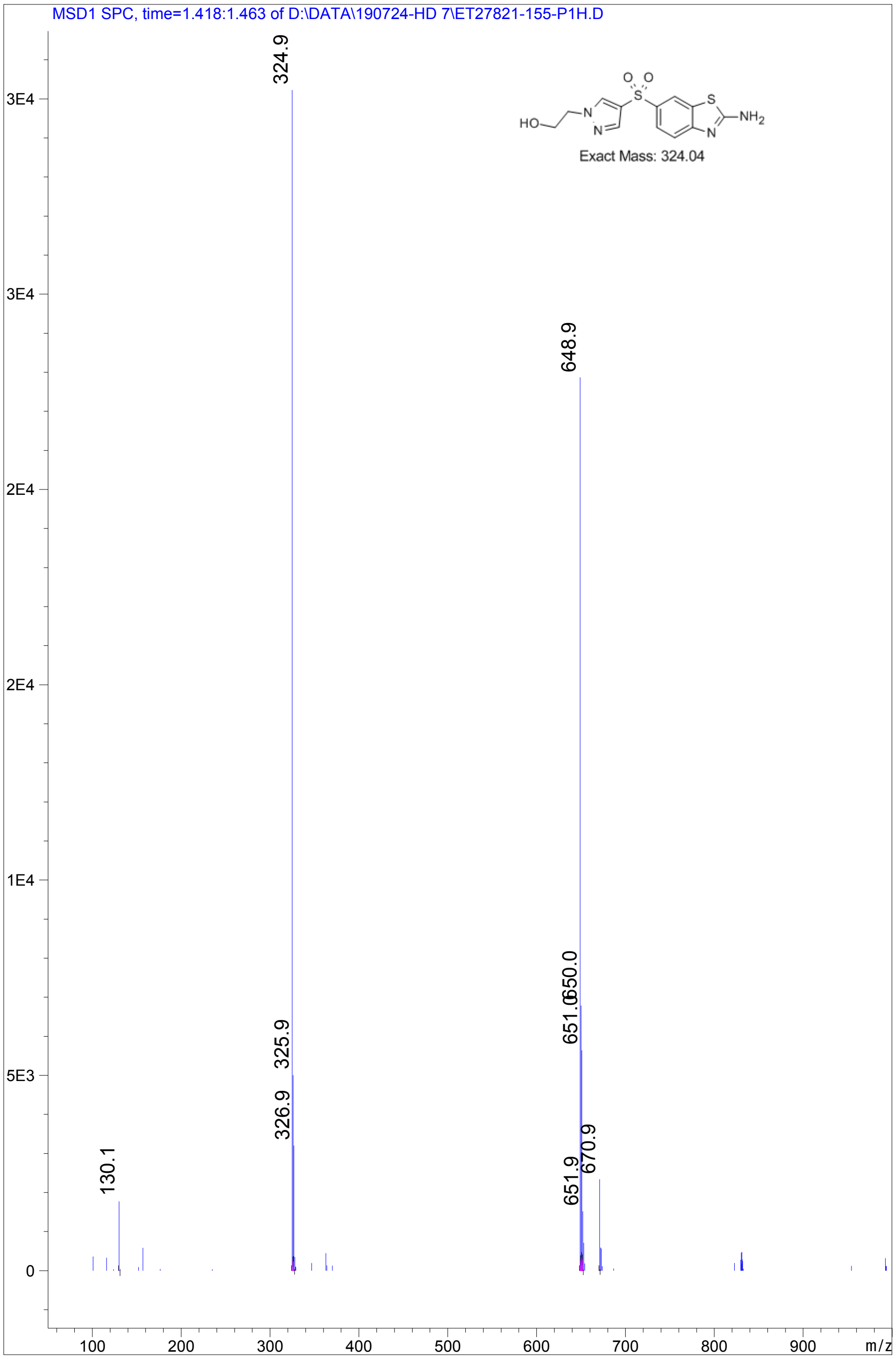


Confidential. For research information only.

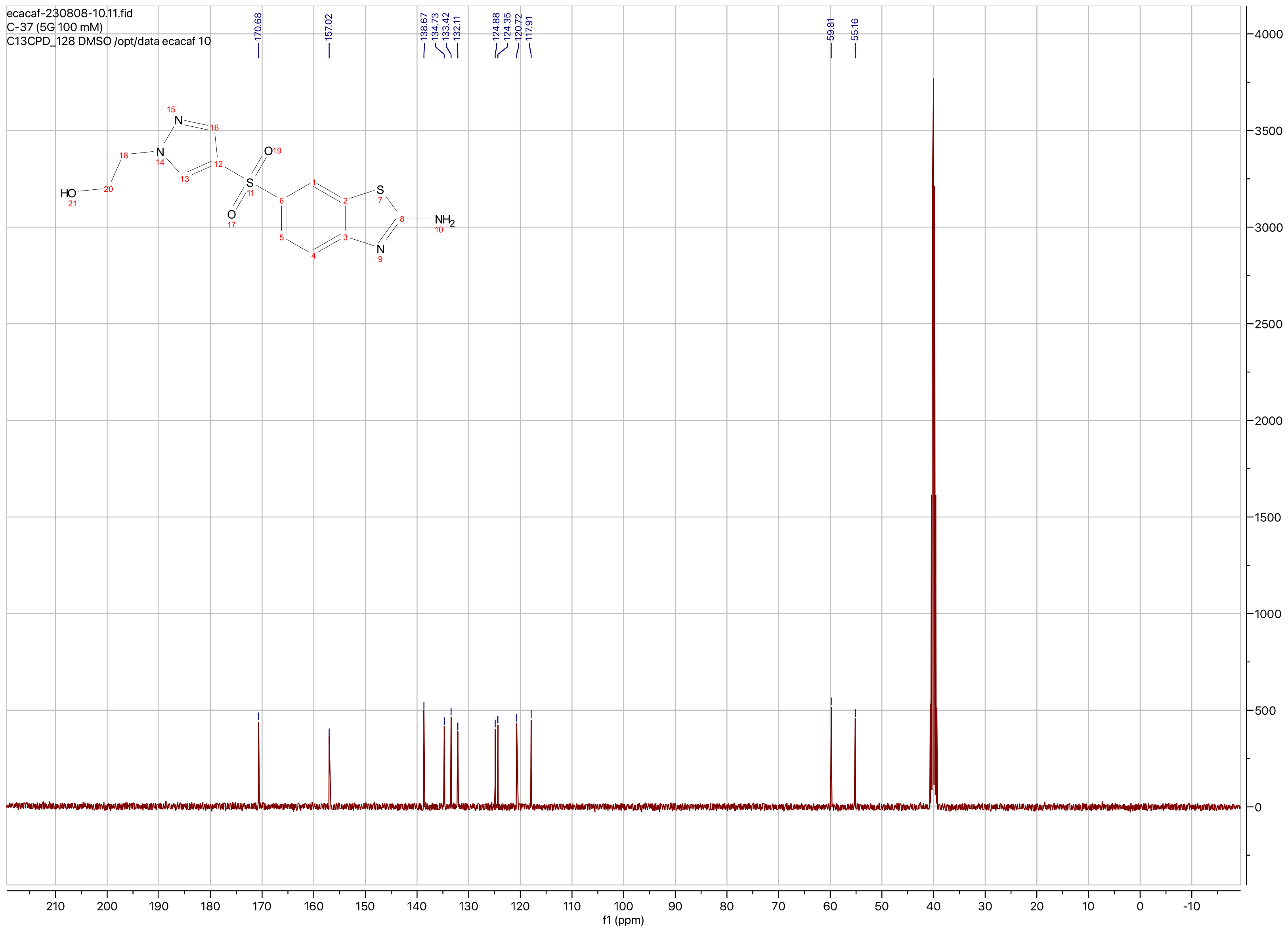
ecacaf-230808-10.11.fid
C-37 (5G 100 mM)
C13CPD_128 DMSO /opt/data ecacaf 10
170.68
157.02
138.67
134.73
133.42
132.11
124.88
124.35
120.72
117.91
59.81
55.16
HO
NH2
f1 (ppm)
210 200 190 180 170 160 150 140 130 120 110 100 90 80 70 60 50 40 30 20 10 0 -10
4000 3500 3000 2500 2000 1500 1000 500 0

**Compound ID: C-38**

ET27821-149-P1H DMSO Bruker_K_400MHz

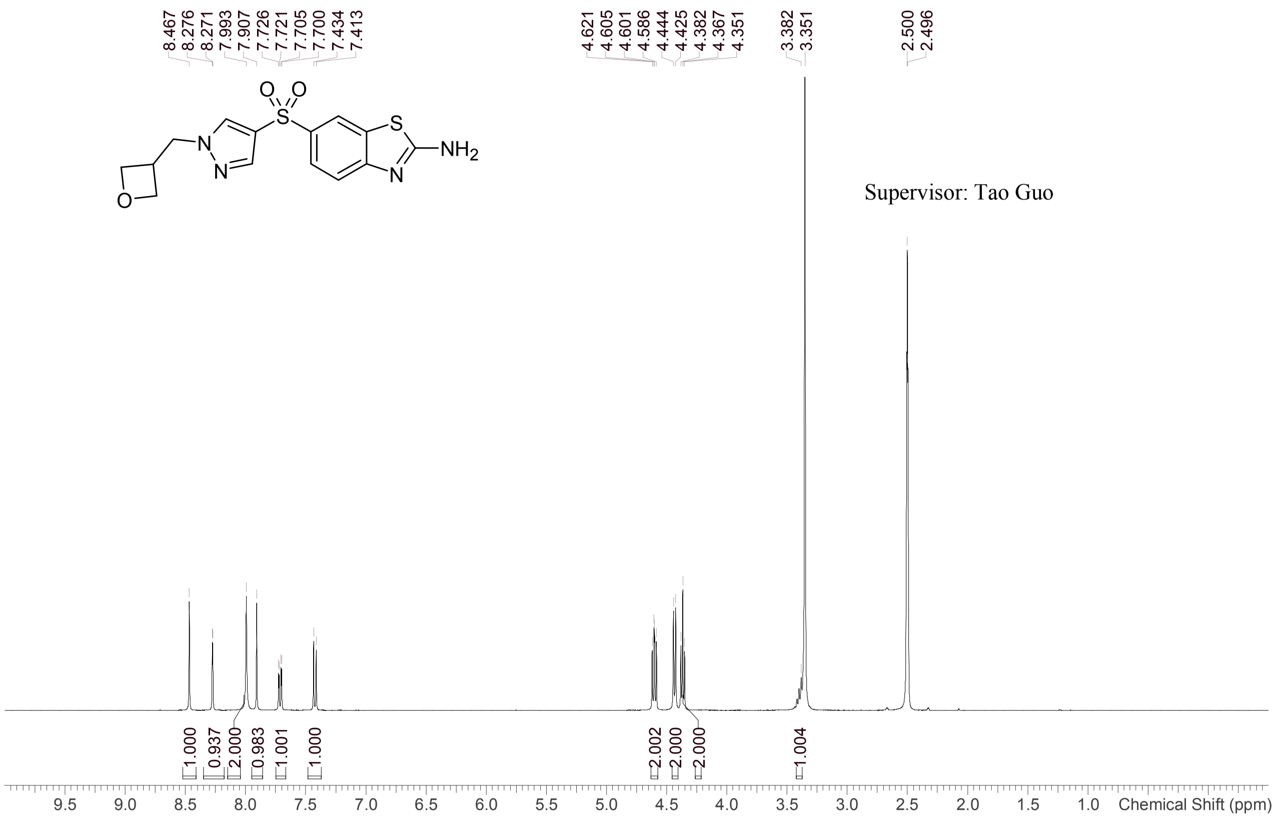


Supervisor: Tao Guo

| | |
|---|---|
| Acquisition Time (sec) | 1.9988 |
| Comment | ET27821-149-P1H DMSO Bruker_K_400MHz |
| Date | 24 Jul 2019 01:46:33 |
| Frequency (MHz) | 400.1300 |
| Nucleus | 1H |
| Number of Transients | 8 |
| Origin | Avance |
| Original Points Count | 16384 |
| Owner | nmr |
| Points Count | 65536 |
| Pulse Sequence | zg30 |
| Receiver Gain | 101.00 |
| SW(cyclical) (Hz) | 8196.72 |
| Solvent | DMSO-d6 |
| Spectrum Offset (Hz) | 2463.4790 |
| Spectrum Type | standard |
| Sweep Width (Hz) | 8196.60 |
| Temperature (degree C) | 23.396 |

Operator:

Date:

Confidential, for research only not for regulatory filing

LCMS REPORT

Compound ID : C-38 ->
Sample ID : ET27821-149-P1H
Injection Date : 23. Jul. 2019
Inj. Vol. : 2.00 ul
Location : P1-B-03
Acq Method : D:\DATA\190723-SD 7\5_95CD_6min-220.M
Data Filename : D:\DATA\190723-SD 7\ET27821-149-P1H.D
Instrument : S

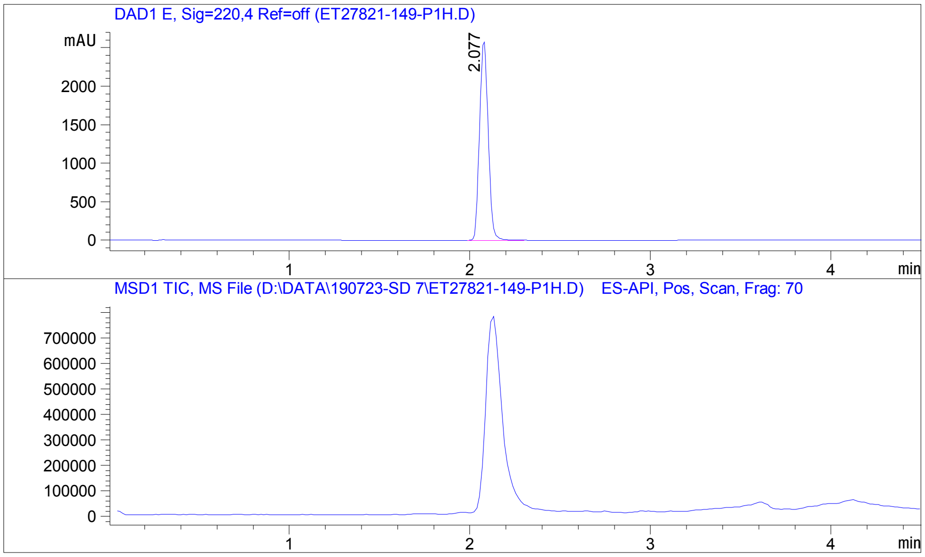


Integration Result

Signal 1 : DAD1 E, Sig=220,4 Ref=off

| Peak # | RT [min] | Area | Height | Height % | Width [min] | Area % |
|---|---|---|---|---|---|---|
| 1 | 2.077 | 8704.431 | 2590.153 | 100.000 | 0.053 | 100.000 |

Operator:_____________

Date:_________________

Confidential. For research information only
PDF created with pdfFactory Pro trial version www.pdffactory.com

MSD1 SPC, time=2.100:2.146 of D:\DATA\190723-SD 7\ET27821-149-P1H.D

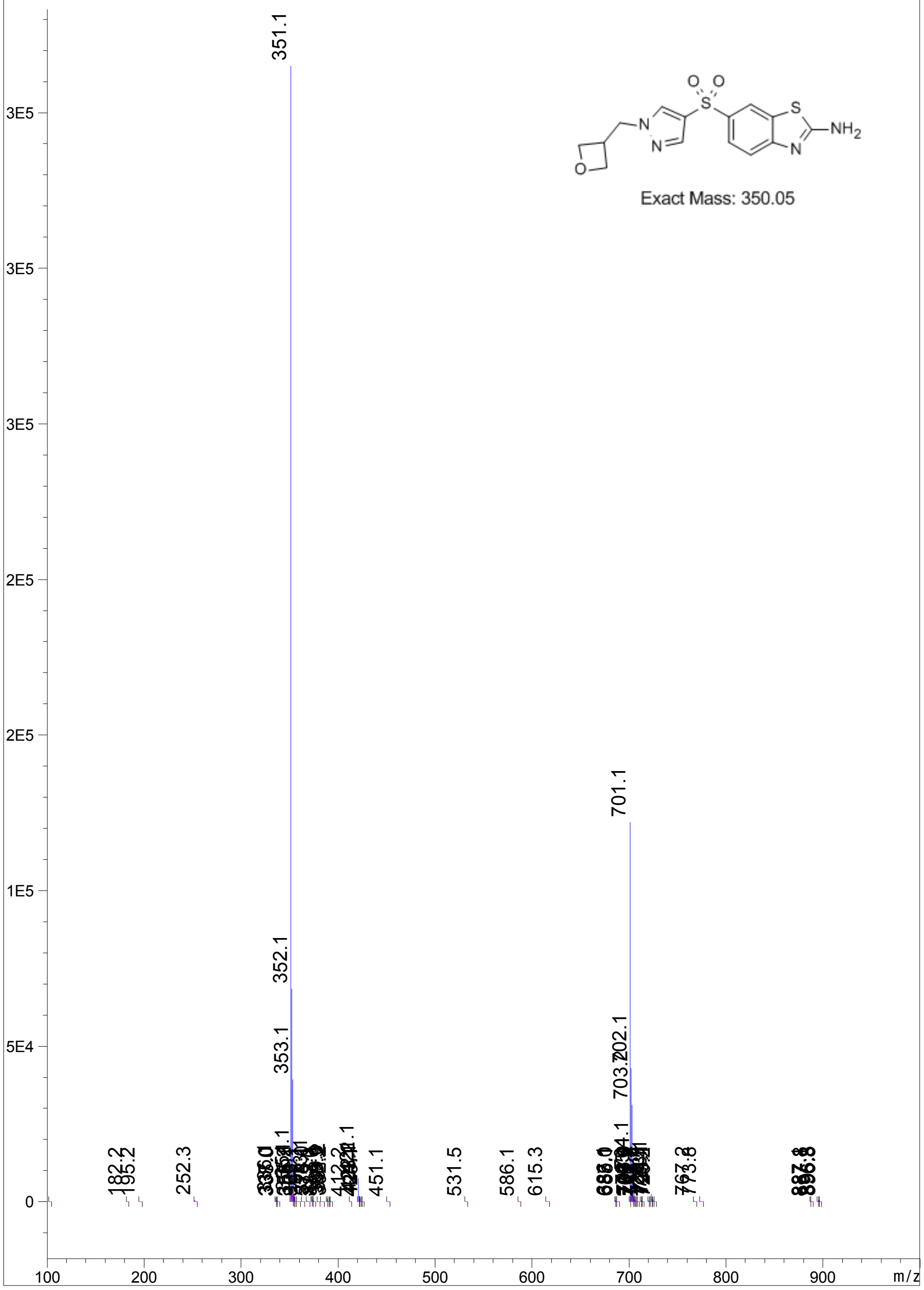


Confidential. For research information only
PDF created with pdfFactory Pro trial version www.pdffactory.com

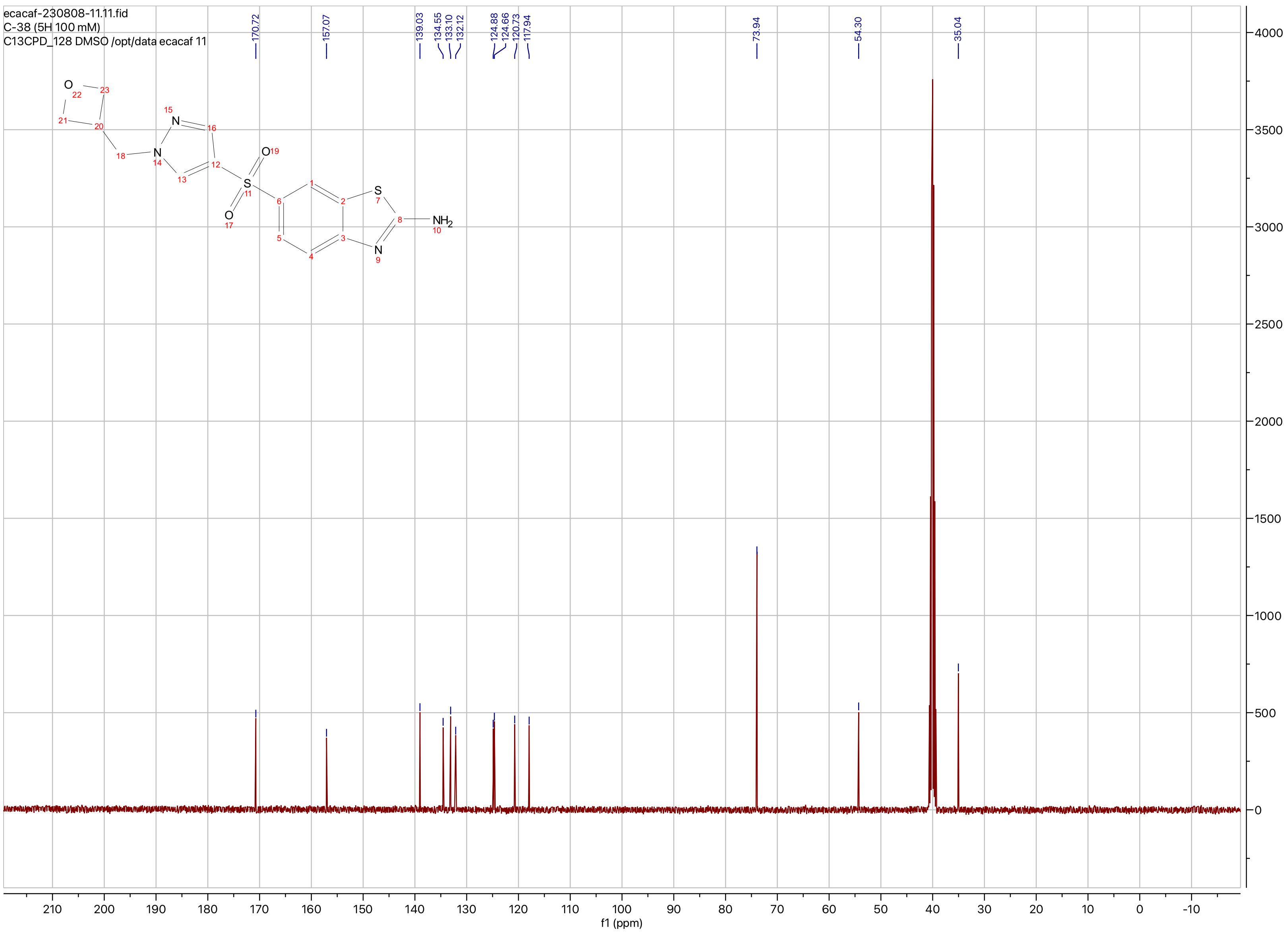
ecacaf-230808-11.11.fid
C-38 (5H 100 mM)
C13CPD_128 DMSO /opt/data ecacaf 11
170.72
157.07
139.03
134.55
133.10
132.12
124.88
124.66
120.73
117.94
73.94
54.30
35.04
NH2
f1 (ppm)
